# Fine-scale habitat partitioning of sympatric stingrays revealed by drone-based remote sensing and deep learning

**DOI:** 10.64898/2026.03.15.710512

**Authors:** Brian O. Nieuwenhuis, Charlotte Turlier, Ioana-Andreea Ciocănaru, Benja A. Blaschke, Malika Kheireddine, Guido Leurs, Jesse E.M. Cochran, Laura L. Govers, Burton H. Jones

**Affiliations:** Marine Science Program, Biological and Environmental Science and Engineering Division, King Abdullah University of Science and Technology (KAUST), 23955-6900 Thuwal, Saudi-Arabia; Conservation Ecology Group, Groningen Institute of Evolutionary Life Sciences (GELIFES), University of Groningen, 9747 AG Groningen, the Netherlands; Independent researcher; Center for Remote Sensing Applications (CRSA), Mohammed VI Polytechnic University (UM6P), 43150 Ben-Guerir, Morocco; Marine Animal Ecology, Wageningen University & Research, Wageningen, the Netherlands; Wageningen Marine Research, Wageningen University & Research, IJmuiden, the Netherlands; Department of Coastal Systems, Royal Netherlands Institute for Sea Research (NIOZ), Texel, the Netherlands

**Keywords:** Niche partitioning, UAV, artificial intelligence, *Taeniura lymma*, *Himantura uarnak*, YOLO-11, SDG-14, elasmobranch, habitat use, coastal ecosystems

## Abstract

Habitat partitioning supports the coexistence of sympatric species and shapes their ecological roles across coastal seascapes. Understanding how sympatric species move through and use coastal habitats therefore provides fundamental ecological insight. Aerial drones provide new opportunities to monitor fine-scale movement and habitat utilisation of elasmobranchs in shallow waters. Here, we use drones to investigate fine-scale habitat partitioning and foraging behaviour among stingrays in a coastal lagoon in the central Red Sea. We conducted 30 aerial transect surveys (~17 ha each) and tracked 40 rays and 1 shark (total tracking time > 23 h). Using a double-observer protocol (manual + AI-assisted), 1,468 rays (6 species) and 4 sharks (2 species) were recorded from the transect surveys. Transect detections were dominated by bluespotted ribbontail rays (*Taeniura lymma*; n = 1,221) and larger-bodied whiprays (predominantly *Himantura* uarnak; n = 187). AI-assisted image analysis outperformed human analysts detecting 97% of these observations, compared to 76% for human analysts. We found pronounced habitat partitioning at sub-kilometre scales: bluespotted rays occupied the shallowest (< 0.4 m deep) lagoonal areas, away from open water, with foraging-related digging concentrated along the mangrove edge, identifying this zone as a key feeding ground and bioturbation hotspot. Whiprays predominated on macroalgal reef flat habitats and appeared to forage non-disruptively on epifaunal prey. Both taxa aggregated with conspecifics. Together, our results demonstrate that contrasting micro-habitat preferences and foraging strategies structure the spatial ecology of sympatric stingrays and highlight how drone-based monitoring coupled with AI can scale ecological inference in nearshore ecosystems.

**Graphical abstract:** 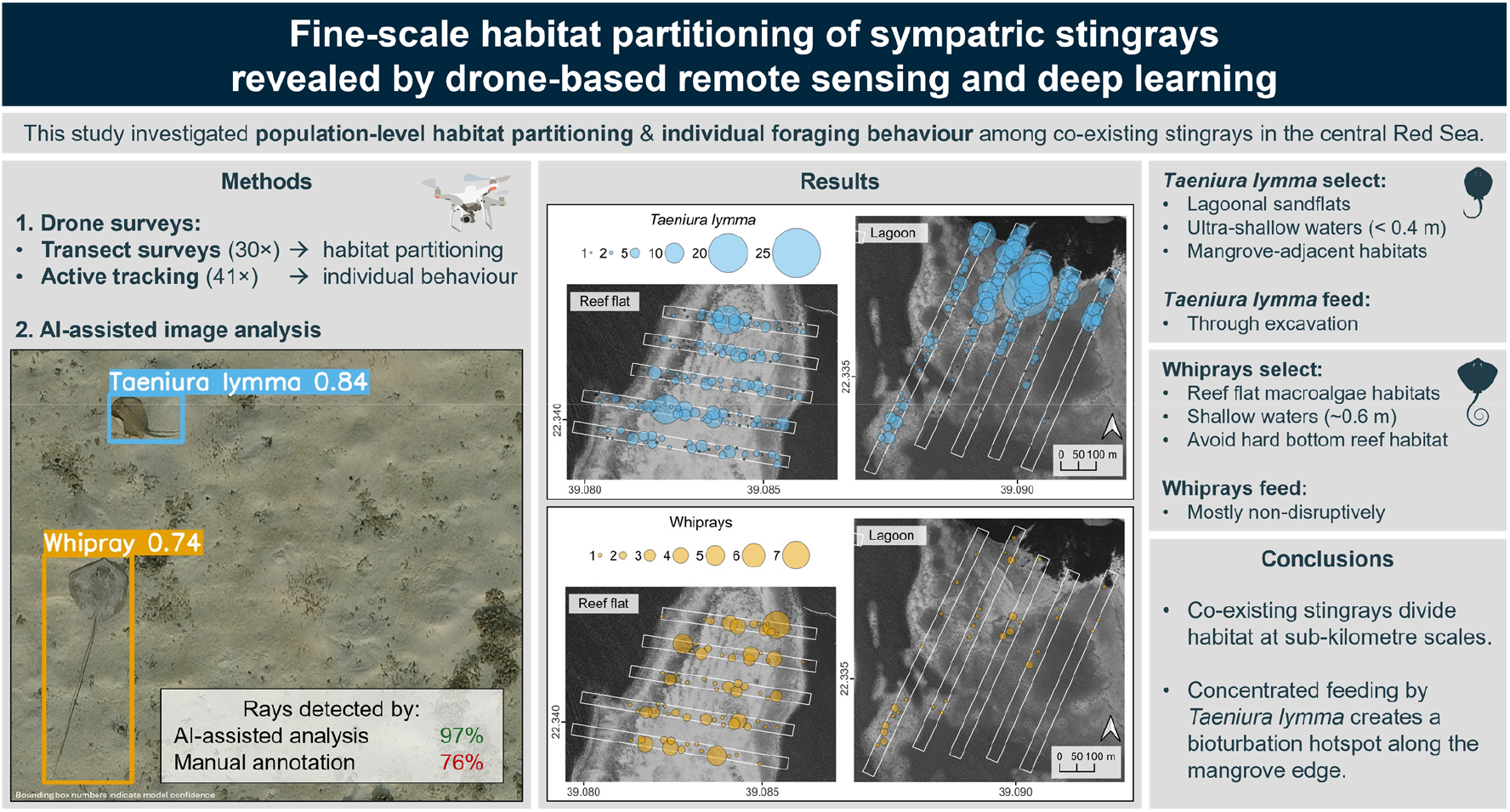

## 1 Introduction

Classical ecological niche theory predicts that ecologically similar species can only coexist when their resource use differs along at least one of three ecological axes, i.e., time, space, or diet (Hardin, 1960; Schoener, 1974). While more recent work has nuanced this view (Adler et al., 2007), habitat partitioning remains an important mechanism through which sympatric species reduce competitive overlap. Furthermore, by spatially concentrating ecological processes such as predation, bioturbation, or nutrient transfer, habitat partitioning modulates the functional roles of different species, thereby contributing to spatial heterogeneity at the ecosystem scale. The distribution of sympatric species across habitats in complex landscapes therefore remains of fundamental ecological interest and may inform both species-level conservation and broader ecosystem-based management (Kyne et al., 2023; Wang et al., 2021).

Elasmobranch populations have experienced widespread declines due to decades of overfishing, with many species now among the most threatened on the IUCN Red List (Dulvy et al., 2024; Sherman et al., 2023). To date, population declines have been most severe in top predatory, large-bodied sharks (Dulvy et al., 2024; Sherman et al., 2023). The loss of large sharks can trigger shifts in community structure through a mesopredator release. In some regions, smaller rays such as the bluespotted ribbontail ray (*Taeniura lymma*) indeed appear to be increasing despite their own sensitivity to overfishing (Dedman et al., 2024; Sherman et al., 2020, 2023; Simpfendorfer et al., 2023). As populations of these mesopredatory rays change, their consumptive and bioturbation-driven impacts may substantially alter the ecological functioning of nearshore ecosystems (Flowers et al., 2021; Grew et al., 2024; Leurs et al., 2023; Nauta et al., 2024). Even so, we still lack a clear understanding of how these rays move through coastal seascapes, connect habitats, and how habitats are partitioned within and across species (Flowers et al., 2021; Kanno et al., 2023; Leurs et al., 2023).

Conventional approaches to movement ecology, including acoustic and satellite telemetry, have provided extensive insights into broad-scale movements of elasmobranchs. However, these methods are often restricted to small sample sizes, have limited spatial resolutions, are difficult to apply in very shallow habitats, and rely on expensive technology (Bullock et al., 2024; Long et al., 2023; Matley et al., 2022; Renshaw et al., 2023). Over the past decade, Unoccupied Aerial Systems (UAS; hereafter drones), have emerged as an effective, complementary tool to study elasmobranch spatial ecology at much finer scales (Butcher et al., 2021; Oleksyn et al., 2021). Although direct drone-based monitoring of elasmobranchs is restricted to clear, shallow waters, it offers several key advantages, including minimal disturbance, high-resolution spatial data, and the ability to record undisturbed behaviour in natural settings (Bourke et al., 2023; Butcher et al., 2021; Crook et al., 2022; Oleksyn et al., 2020). In addition, drones have been used to indirectly quantify stingray foraging and bioturbation by mapping ray feeding pits on intertidal sandflats (Grew et al., 2024; Nauta et al., 2024). Thus, the adoption of drones has greatly expanded opportunities to monitor fine-scale habitat use and behaviour of elasmobranchs (Butcher et al., 2021; Oleksyn et al., 2021).

The volume of imagery produced by drone surveys, however, has outpaced our ability to process it, creating a major analytical bottleneck. Advances in artificial intelligence (AI) offer an avenue to alleviate this constraint (Axford et al., 2024; Katija et al., 2022). Computer vision models show considerable promise for automated detection of elasmobranchs from drone imagery, particularly in real-time shark detection (Chen and Liu, 2017; Desgarnier et al., 2022; Gorkin et al., 2020; Purcell et al., 2022; Saqib et al., 2018; Sharma et al., 2018). Yet the integration of AI in drone-based elasmobranch research remains limited, with most recent studies still relying on manual annotation (Ayres et al., 2021; Bullock et al., 2024; Colefax et al., 2020; Crook et al., 2022; McIvor et al., 2022; Myers, 2024; Oleksyn et al., 2020; Richardson et al., 2026; Sanchez et al., 2025). Key barriers to the wider adoption of AI include the scarcity of high-quality annotated datasets, limited transferability of existing models, the technical expertise required to successfully apply AI-based methods, the small size of individual animals in drone imagery, and marine-specific challenges like wave distortions, caustics, and sun glint (Axford et al., 2024; Belcher et al., 2023; Joyce et al., 2019; Katija et al., 2022; Oleksyn et al., 2021).

In this study, we investigate habitat partitioning among stingrays in the central Red Sea, where severe shark depletion has coincided with increased abundances of bluespotted rays (Simpfendorfer et al., 2023; Spaet et al., 2016; Spaet and Berumen, 2015). Despite their widespread distribution, the fine-scale habitat use of bluespotted rays and their interactions with sympatric species like the endangered coach whipray (*Himantura uarnak)* remain poorly understood (Sherman et al., 2021b, 2021a). We use the Red Sea’s setting to quantify fine-scale habitat partitioning between these species in a human-impacted environment and to evaluate the ecological and methodological benefits of integrating active tracking, transect surveys, and AI-assisted image analysis in drone-based elasmobranch research.

## 2 Materials and Methods

### 2.1 Study area

Aerial surveys were conducted over a small coastal lagoon and the adjacent reef flat (22.34° N, 39.09° E) in the central Red Sea (Fig. 1). The area lies within the King Abdullah University of Science and Technology campus, which prohibits commercial fishing but allows for recreational and research activities, including angling. Both the lagoon and reef flat feature extensive shallow (< 2 m deep) water that contains a mix of sand, reef, macroalgae, and seagrass habitats, with sand habitats being the most common (Fig. 1b,c; Nieuwenhuis et al., 2022). The lagoon is bordered by grey mangroves (*Avicennia marina* (Forssk.) Vierh.). Reef flat bathymetry is relatively uniform at about 0.6 m deep, while the lagoon’s bathymetry is more variable with deep sand habitats (> 1.5 m deep) interspersed with shallow reef framework (< 0.5 m deep), and an ultra-shallow sandflat (< 0.4 m deep) next to the mangroves (Nieuwenhuis et al., 2026). Tidal amplitudes in the area are generally less than 0.2 m (Pugh et al., 2019).

**Figure 1.**
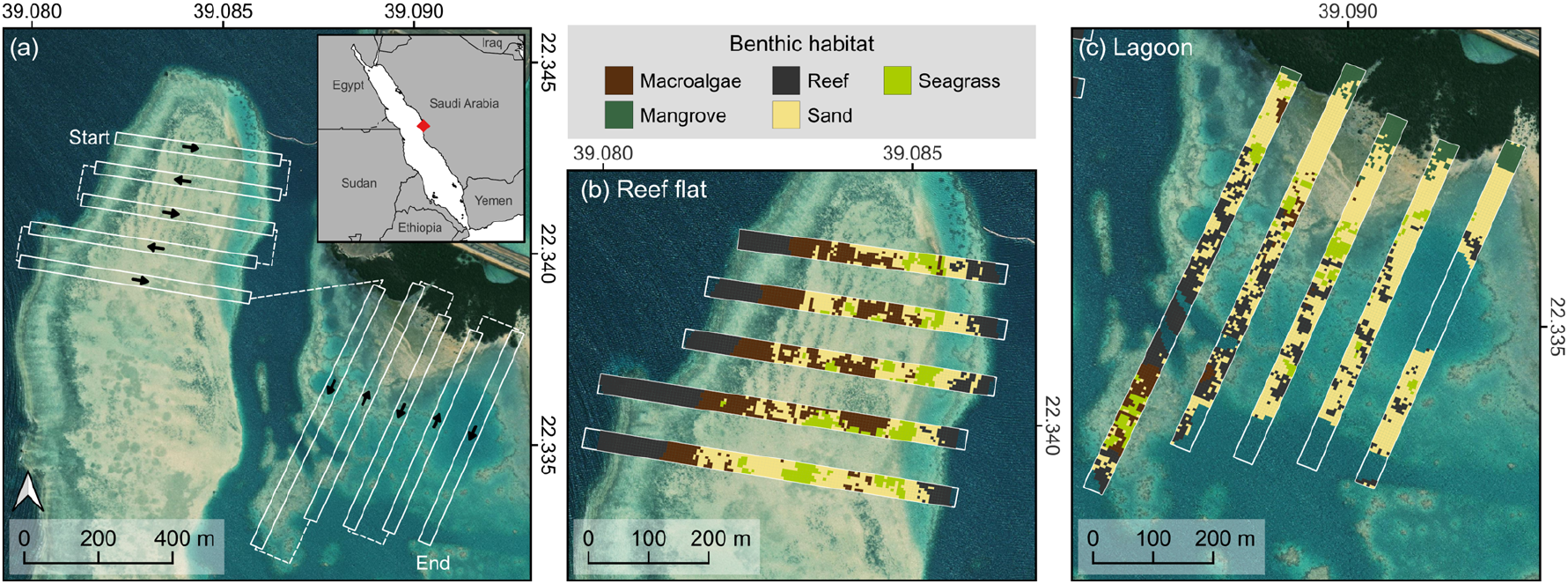
(a) Map of the study area with the transect surveys overlayed. White boxes indicate surveyed transects, arrows indicate flight direction, dashed lines indicate the transition routes between transects. Inset shows the general location of the study area on the Red Sea coast. (b, c) Benthic habitat maps of transect areas for the (b) reef flat and (c) lagoon, respectively. Map contains ESRI satellite imagery and Natural Earth data.

### 2.2 Data collection

Thirty aerial transect surveys were conducted between May and November 2024. Each survey consisted of ten transects evenly distributed across the lagoon and the reef flat (Fig. 1a). Surveys were flown at 35 m altitude using a DJI Matrice 300 RTK equipped with a 45-megapixel DJI Zenmuse P1 camera and a 35 mm lens, producing a 35 m image swath with a ground sampling distance (GSD) of 0.41 cm px^−1^ (SZ DJI Technology Co. Ltd., 2023a, 2021). Nadir images were captured every second with approximately 70 % frontal overlap and a shutter speed of 1/ 400 s. Transects were flown at 6 m s^−1^ to avoid motion blur. When transitioning between transects, which were spaced 100 m apart to reduce double counting, the drone accelerated to 15 m s^−1^, so that a full survey could be completed in 20 minutes. A DJI D-RTK2 base station supplied real-time kinematic corrections to ensure precise replication of transect paths (SZ DJI Technology Co. Ltd., 2023b). Each survey generated about 1074 images covering roughly 17 ha of shallow (< 2 m deep) habitat.

To complement the transect data, active tracking surveys were conducted between February and July 2024. The drone was flown manually until a ray or shark was located, at which point it was positioned directly above the ray or shark and set to capture a nadir image every second with a shutter speed of 1/ 400 s while the pilot kept the tracked animal centred in the live feed (Colefax et al., 2020; Oleksyn et al., 2020). Tracking altitude ranged from about 25 to 35 m depending on the prevailing weather conditions, resulting in a GSD of 0.3 to 0.4 cm px^−1^ (SZ DJI Technology Co. Ltd., 2023a, 2021). To extend the tracking duration beyond the drone’s battery-life (~ 20 min), a second drone temporarily assumed tracking while the battery of the primary drone was being replaced (Colefax et al., 2020). Because the secondary drone lacked RTK corrections and was equipped with a slower (maximum capture rate 2 s) and lower resolution (12 megapixels) DJI Zenmuse XT-2 camera, the drones were swapped back as soon as possible. This relay procedure was repeated until either visual contact with the ray was lost or all available batteries were depleted, enabling a maximum tracking duration of up to two hours. In total, 41 rays and 1 shark were tracked.

### 2.3 Image analysis

Transect imagery was annotated for the presence of six elasmobranch groups: bluespotted rays (*Taeniura lymma*), whiprays (encompassing *Himantura, Pateobatis*, and *Maculabatis spp*.), spotted eagle rays (*Aetobatus ocellatus*), mangrove whiprays (*Urogymnus granulatus*), cowtail rays (*Pastinachus sephen*), and sharks (Selachii). Whiprays were pooled due to wave distortions and sand cover frequently limiting species-level identification. All transect images were screened twice, once manually and once using an AI-assisted workflow based on a custom-trained YOLOv11x model (Jocher and Qiu 2024; Fig. 2). This double observer protocol allowed us to maximise detection accuracy and evaluate the potential of computer vision models to accelerate annotation of marine drone imagery.

**Figure 2.**
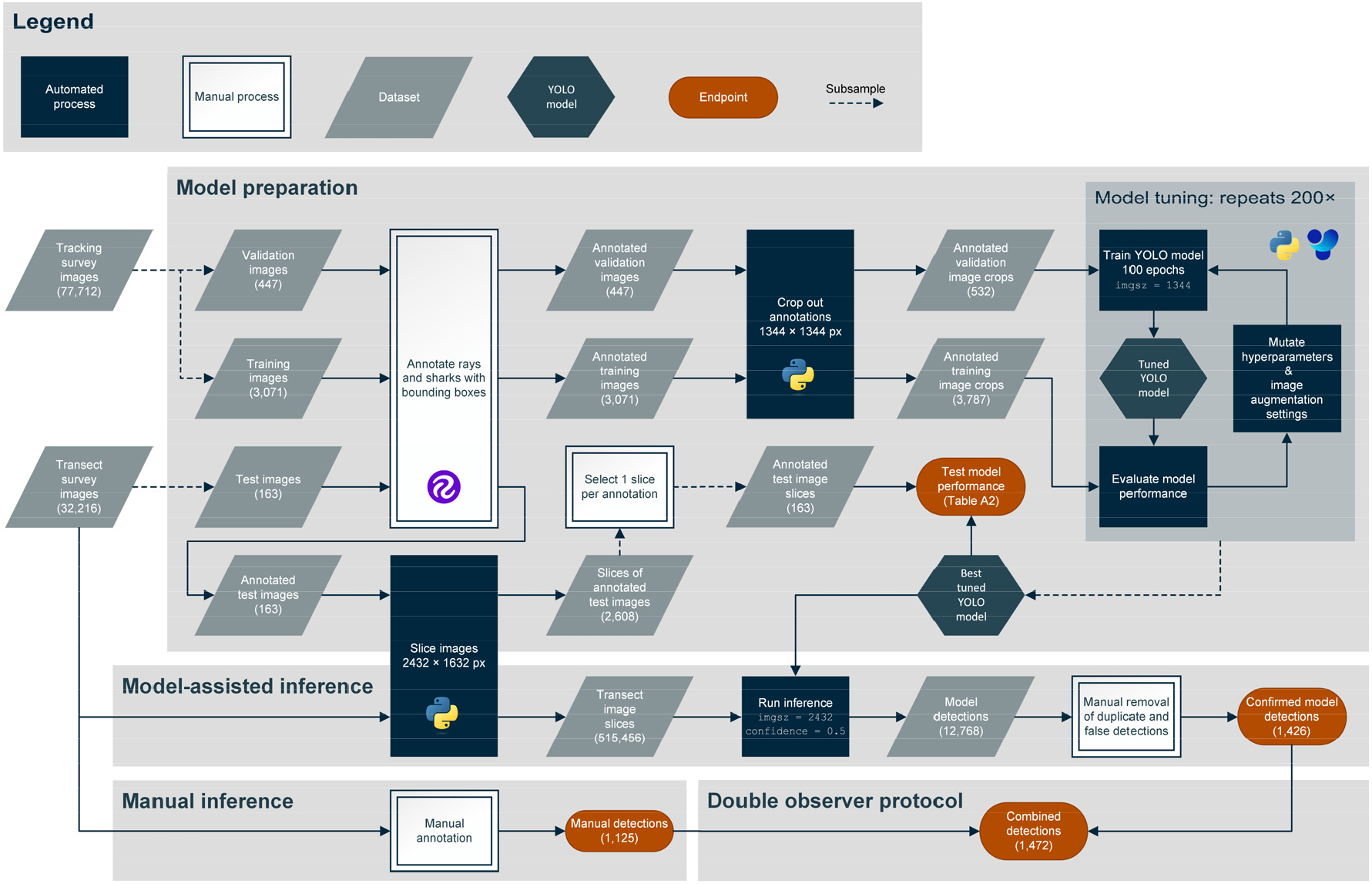
Overview of the image analysis conducted for the transect imagery. For visual clarity only the development of the final YOLO-based model is depicted. Note the terminology for different image sizes: ‘image’ = 8192 × 5460 px (original image), ‘image slice’ = 2432 × 1632 px, ‘image crop’ = 1344 × 1344 px. Logos indicate the reliance of different steps on Roboflow, Python, and Ultralytics’ YOLO software (Dwyer et al. 2024; Jocher and Qiu 2024). `imgsz`: image size parameter in the YOLOv11x model.

Three core strategies were implemented to optimise model accuracy. Firstly, the model was trained on imagery from the tracking surveys, which facilitated efficient annotation and provided high-quality, closely matching, but independent training data. Secondly, because bluespotted rays are very small (≤ 0.35 m disc width) relative to the 35 m swath of our drone images, standard YOLO downscaling procedures would have left insufficient visual detail to learn from (Akyon et al., 2022; Axford et al., 2024; Jocher and Qiu, 2024). Thus, to train and apply the model at the image’s native resolution, we implemented a custom “Slicing Aided Hyper Inference” workflow, whereby training images were automatically cropped to 1344 × 1344 pixels and transect images were divided into 16 overlapping tiles of 2432 × 1632 pixels before inference (Akyon et al., 2022). Thirdly, hyperparameters and image augmentation settings were iteratively tuned for 200 iterations of 100 epochs using YOLO’s automated tuning routine (Jocher and Qiu, 2024). The resulting model achieved a mean Average Precision (mAP-0.5) of 0.85 on a transect-derived test set, indicating good performance within the intended application (Supplementary table A.2). For optimal accuracy, inference was run at a confidence threshold of 0.5, after which all model detections were manually reviewed to remove false positives and duplicates resulting from image and tile overlap. Full workflow details are provided in Appendix A.

Detected individuals were geolocated by matching their location within the image to drone-derived orthomosaics of the survey area (Nieuwenhuis et al., 2026, 2025). We also recorded whether the individual was digging and, if visible, disc width, disc length, and total length were measured to the nearest pixel in FIJI v.1.54 (Schindelin et al., 2012). Pixel values were converted to standard linear units by multiplication with the GSD, which was calculated based on Equation 1 and calibrated with a submerged reference target.

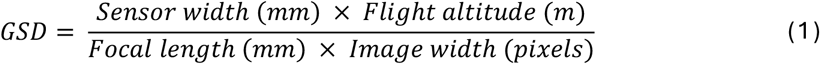

Tracking imagery was manually annotated for swimming, stationary, and digging behaviour. Stationary behaviour indicated that the individual remained in place but did not necessarily indicate the ray was at rest and could include surface feeding behaviour (Crook et al., 2022). Digging was recorded when sediment plumes indicated excavation or jetting behaviour (Crook et al., 2022). The benthic habitat beneath each individual was annotated as sand, seagrass, reef, macroalgae, mangroves, or rubble. Because the drone was kept straight above the tracked animal, the drone’s location served as a proxy for the animal’s position (Oleksyn et al., 2020). Geolocation data was extracted from the images using the *exiftoolr v.0.2.8* package in R (Harvey, 2024; O’Brien, 2018). Swimming speeds were calculated from a 10 second rolling average of the drone positions, excluding speeds below 0.05 m s^−1^ (Oleksyn et al., 2020). Size measurements were taken as described for the transect images.

### 2.4 Statistical analysis

Formal analyses focused on the two most abundant taxa, the bluespotted rays and whiprays, and relied on transect data because tracking locations were strongly influenced by the initial search path. To relate ray detections to environmental gradients and to explicitly define absence observations, transects were subdivided into 5 × 5 m grid cells in QGIS v3.34.4 (QGIS Association, 2024). For each cell, we extracted benthic habitat, distance to open water, distance to mangroves, tide-adjusted water depth, and distance to nearest conspecific, with the latter two calculated separately for each survey. Environmental variables were derived from drone-based datasets as described in appendix B. In brief, bathymetry was estimated using Stumpf’s bathymetric band ratio with data from Nieuwenhuis et al. (2026). Benthic habitat was manually classified based on drone orthomosaics from June 2024 (Fig. 1b,c). Distances to deep open water and mangroves were calculated using the *terra v.1.7-71* library in R following manual delineation of open water and a threshold-based mangrove classification, respectively (Hijmans et al., 2020).

Since multiple detections within a single cell on the same day were rare (1% of positive detections), spatial distribution was modelled with binomial presence-absence generalised additive mixed models (GAMMs; Wood 2017; Pedersen et al. 2019). Presence of each taxon was modelled as a function of survey area (reef flat or lagoon), water depth, benthic habitat, distance to open water, and distance to nearest conspecific, which was truncated at 100 m and included to assess aggregation and account for local spatial autocorrelation. A tensor-product smooth of the coordinates (shared across taxa) accounted for broad-scale spatial autocorrelation. Analyses were restricted to waters less than 1.5 m deep, beyond which bluespotted ray detections were rare (Supplementary fig. C.1). To accommodate survey-specific variation in water clarity and wave distortions, survey ID was included as a random-effect smooth interacting with depth (Pedersen et al., 2019; Wood, 2017). The resulting model structure is summarised in Equation 2.

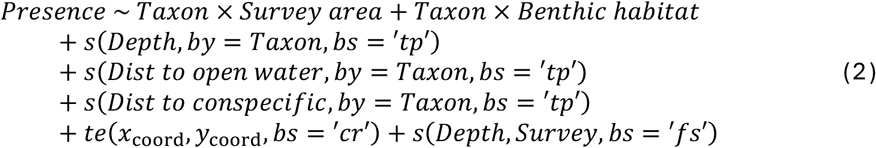

A second binomial GAMM modelled the probability that the observed bluespotted rays were digging as a function of water depth, distance to mangroves, and survey area. As above a tensor-product smooth was included to account for spatial autocorrelation. Survey ID was included as a random effect (Equation 3). Both models were fitted using restricted maximum likelihood with smooth selection enabled (`select = TRUE`) to allow unsupported smooth terms to be penalised to zero. Significance of model terms was calculated using Wald tests (Wood, 2017), while the significance of post-hoc pairwise comparisons was computed through delta method Z-tests with false discover rate adjusted p-values (Arel-Bundock et al., 2024; Benjamini and Hochberg, 1995). Analyses were conducted in R v4.2.2 (R Core Team, 2022), utilising *Tidyverse v2.0.0* for data preparation and visualization (Wickham et al., 2019), *mgcv v.1.9-1* to fit GAMMs (Wood, 2017), *DHARMa v0.4.6* for model diagnostics (Hartig, 2016), *marginaleffects v0.19.0* for post-hoc comparisons (Arel-Bundock et al., 2024), and *gratia v0.9.0* to plot model outputs (Simpson, 2024).

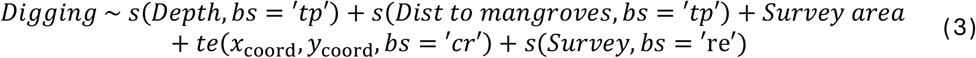

## 3 Results

### 3.1 Transect surveys

Across thirty transect surveys, we recorded 1,468 ray detections, dominated by 1,221 bluespotted rays and 187 whiprays. We also observed 24 spotted eagle rays, 23 mangrove whiprays, 9 cowtail rays, 3 unidentified rays, 3 blacktip reef sharks (*Carcharhinus melanopterus*) and 1 sicklefin lemon shark (*Negaprion acutidens*). Among whiprays, 59 individuals showed spotting consistent with *H. uarnak*, 16 were spotless resembling *P. fai*, while the remainder could not be identified further (Appendix D). The AI-assisted workflow detected 97% of all individuals (1,426 detections), while human analysts detected 76% (1,125 detections; Fig. 3a). Manual annotation of the drone imagery required up to fifteen hours per survey, while manual review of the model outputs in the AI-assisted workflow could be completed in under one hour. Although not trained for turtles or guitarfish, the model occasionally also flagged these taxa. Manual review confirmed the presence of 22 turtles and 3 guitarfish, including two critically endangered *Glaucostegus halavi* (Supplementary fig. C.2; Kyne and Jabado 2019).

**Figure 3.**
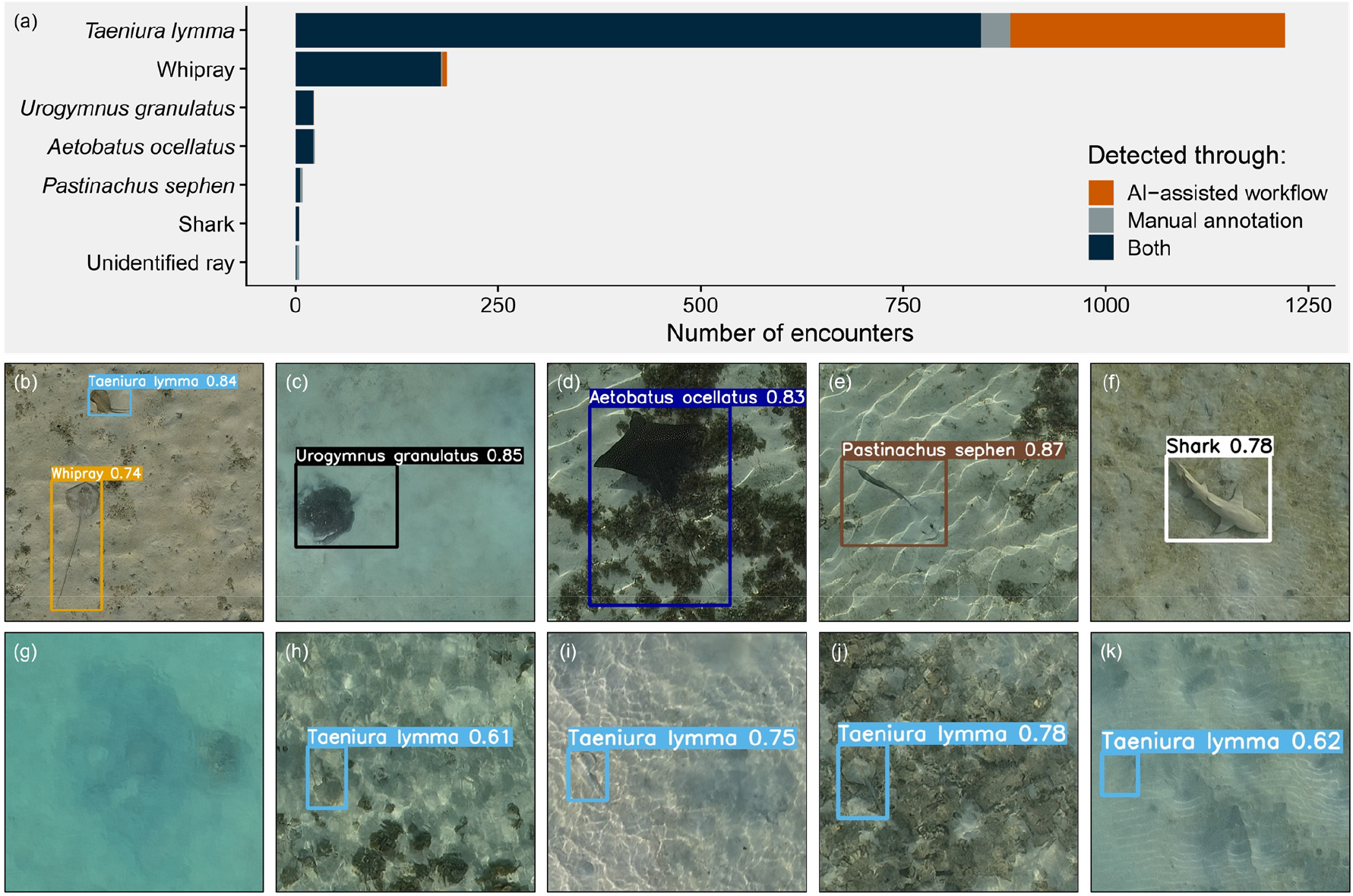
(a) The number of rays and sharks detected in the transect surveys through manual and/or AI-assisted annotation. Note that if the AI model correctly detects a ray or shark but assigns the wrong class this was corrected during the manual filtering step (Fig. 2); thus, such detections are counted as correct in this figure. (b-f) Cropped examples of AI-driven detections for each class. (g) Example of a ray that could not be identified due to the dust cloud it produced. (h-k) Examples of hard to spot bluespotted rays that were only detected in the AI-assisted workflow. (b-f, h-k) Numbers indicate model confidence.

Bluespotted rays averaged 40.7 (range = 12 – 67) individuals per survey (2.6 individuals ha^−1^), making them by far the most abundant taxon, while whiprays averaged 6.2 (range = 1 – 15) individuals per survey (0.4 individuals ha^−1^). Disc size revealed that most bluespotted rays were adults, whereas whiprays were mostly juveniles (Supplementary fig. C.3). The two taxa showed pronounced spatial segregation. Bluespotted rays were concentrated within the lagoon (*p* = 0.040, z = −2.1), especially on the ultra-shallow sandflat adjacent to the mangroves (Fig. 4a, 5d). Juveniles were almost exclusively recorded in this area; however, their small size may have limited detectability in deeper water (Supplementary fig. C.4). Conversely, whiprays primarily occupied the adjacent reef flat (*p* = 0.015, *z* = 2.7; Fig. 4b, 5d). Other taxa that were observed more frequently on the reef flat versus the lagoon included spotted eagle rays (17:7), blacktip reef sharks (3:0), and guitarfish (3:0). Conversely, mangrove whiprays (7: 16), cowtail rays (2:7), turtles (5:17), and the sicklefin lemon shark (0:1) were predominantly detected within the lagoon (Fig. 4c).

**Figure 4.**
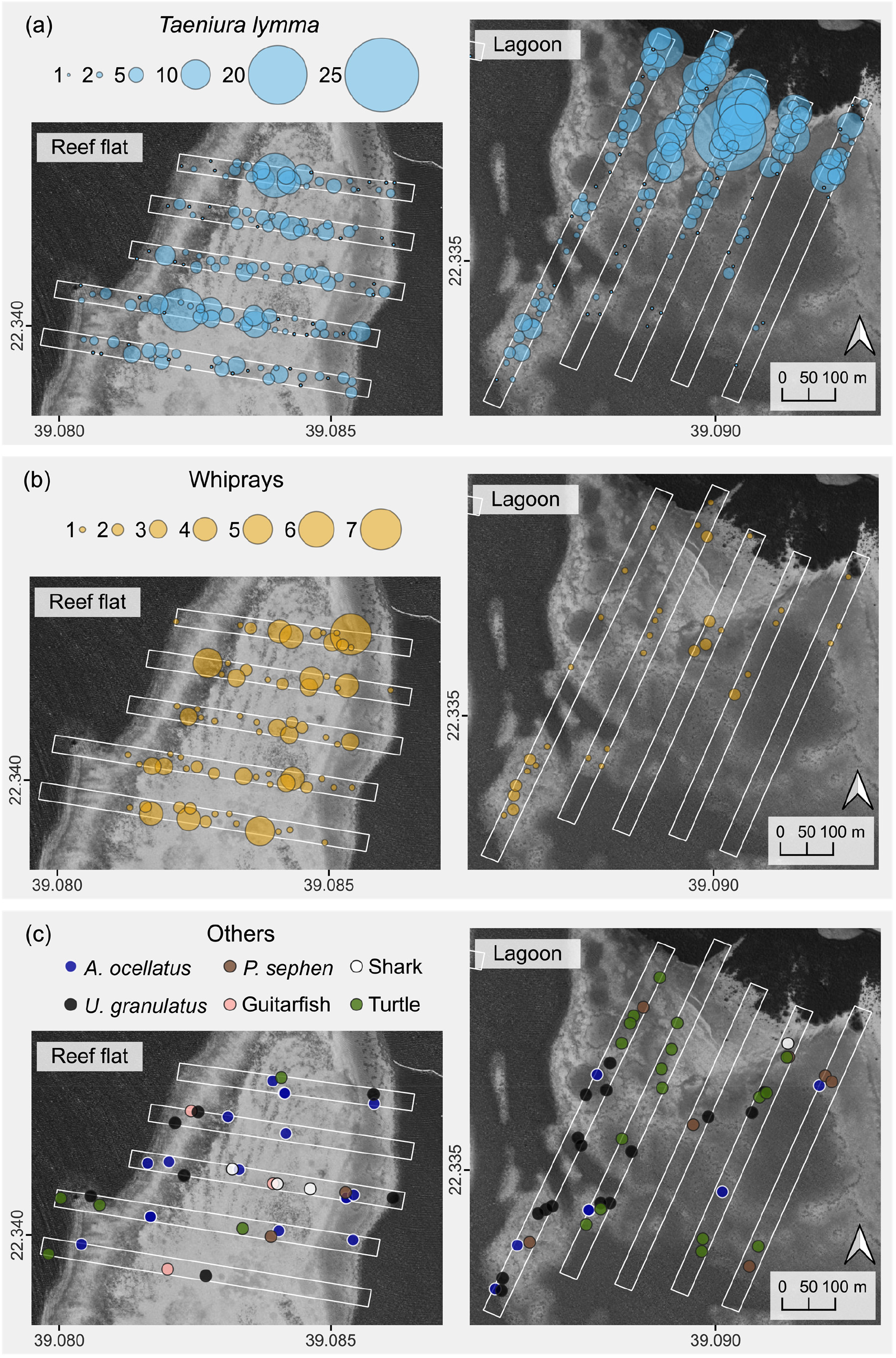
Bubble plots showing the relative observation densities across the transect surveys for (a) bluespotted rays, (b) whiprays, and (c) any other classes. Map contains ESRI satellite imagery.

Environmental models revealed clear taxon-specific responses to environmental gradients (Fig. 5). Bluespotted rays selected very shallow water of ~0.2 m deep, whereas the occurrence probability of whiprays peaked at ~0.6 m (Fig. 5a, Supplementary fig. C.1). Both taxa were positively associated with macroalgal habitat relative to sand (*p* < 0.009, *z* ≤ −2.9). Whiprays also showed a strong negative response to reef habitat (*p* < 0.001, *z* = 3.7; Fig. 5 b, Supplementary fig. C.5). Bluespotted ray presence increased with distance from open water, while this had no effect on the whiprays (Fig. 5c). Both taxa displayed evidence of aggregation, with higher occurrence probabilities at smaller distances to conspecifics (Fig. 5e). In addition, bluespotted rays were observed to be digging approximately twice as often as the whiprays (19.7% vs. 11.7%). Digging by bluespotted rays clustered on the mangrove-adjacent sandflat in the lagoon (Fig. 6). Digging probabilities were larger in the lagoon (*p* < 0.001, *z* = 3.4), close to the mangroves (*edf* = 0.8, *χ*^2^ = 4.1, *p* = 0.005) and in shallow waters (*edf* = 0.7, *χ*^2^ *=* 3.0, *p* = 0.050; Supplementary fig. C.6).

**Figure 5.**
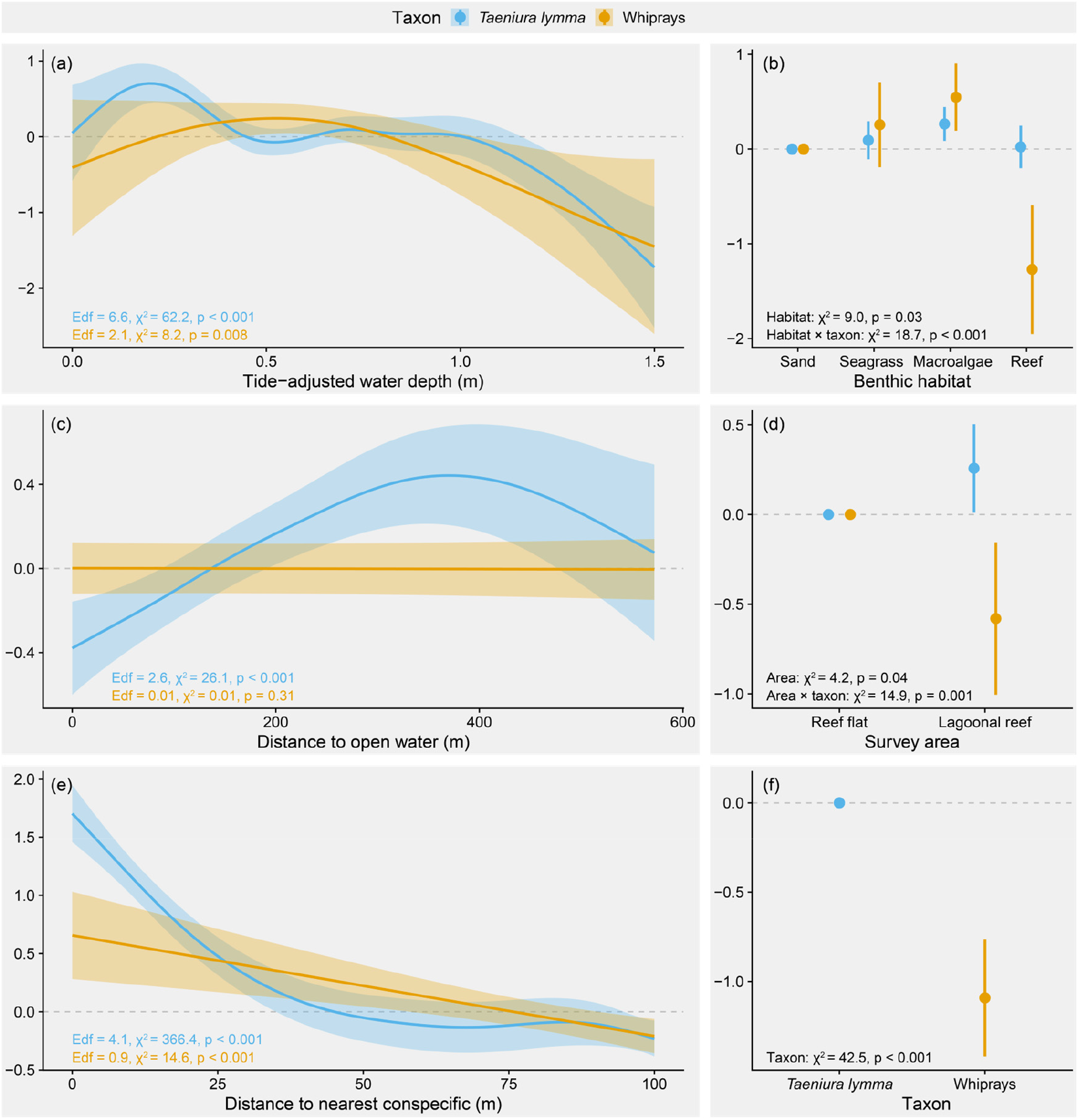
Partial effects of (a) bathymetry, (b) benthic habitat, (c) distance to open water, (d) survey area, (e) distance to nearest observed conspecific, and (f) ray taxon plots on ray encounter probability. Plots are on the link scale. Edf: effective degrees of freedom. Total deviance explained is 10.8%.

**Figure 6.**
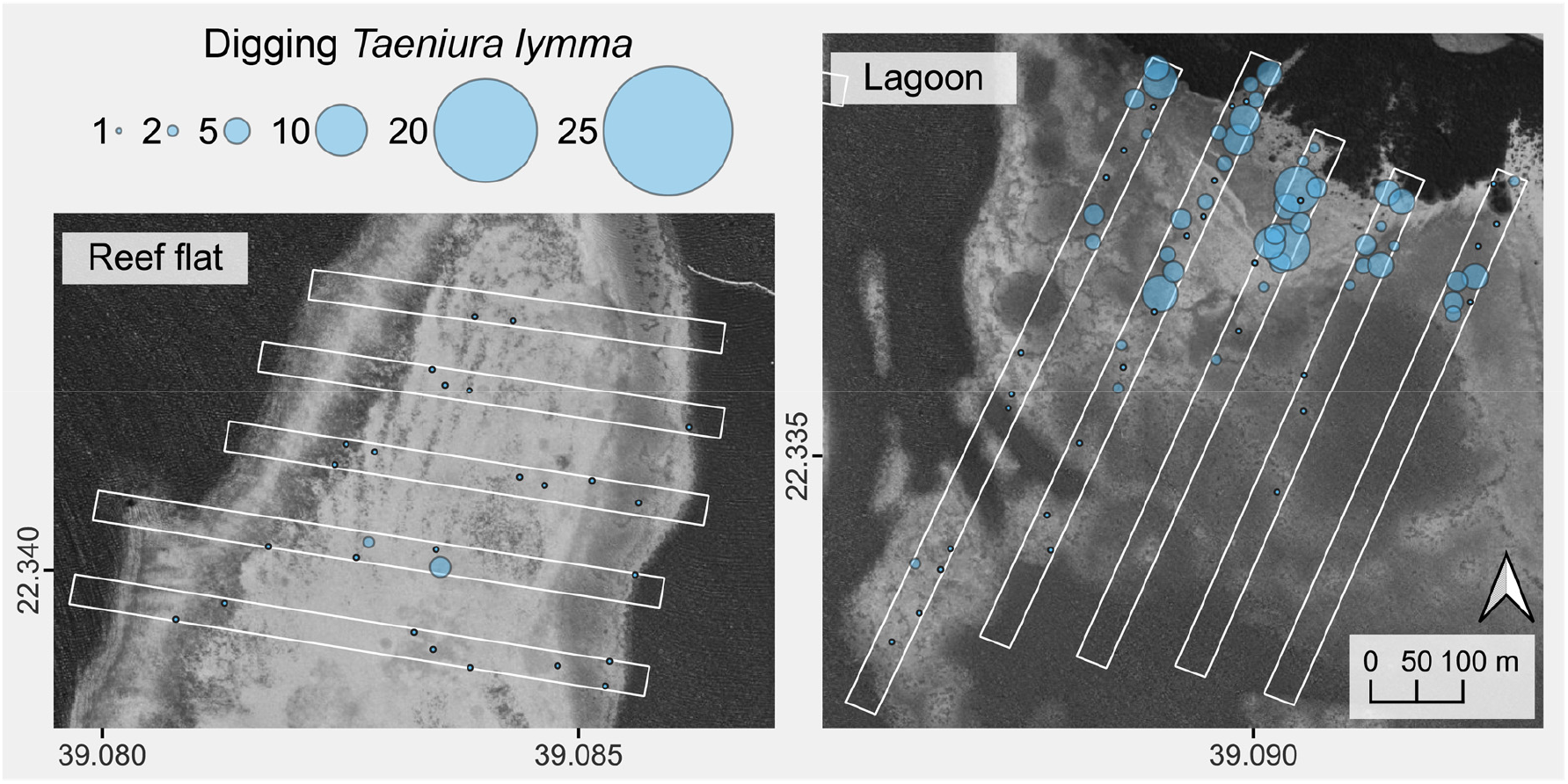
Bubble plot showing the relative observation densities across the transect surveys of digging bluespotted rays. Map contains ESRI satellite imagery.

### 3.2 Tracking surveys

We tracked 41 individuals for a total tracking time of 23 hours and 16 minutes (mean = 34 min, range = 0.6 – 114 min). Tracks included 20 bluespotted rays, 14 whiprays (10 of which were identified as *H. uarnak*), 3 mangrove whiprays, 2 spotted eagle rays, 1 cowtail ray, and 1 sicklefin lemon shark. Bluespotted rays were most often tracked in the lagoon (15 of 20), while most whiprays were tracked on the reef flat (12 of 14; Fig. 7). Swimming speeds of benthic rays were comparable and ranged from 0.25 to 0.33 m s^−1^. The shark and eagle rays swam notably faster, averaging 0.54 and 1.0 m s^−1^, respectively (Supplementary fig. C.7).

**Figure 7.**
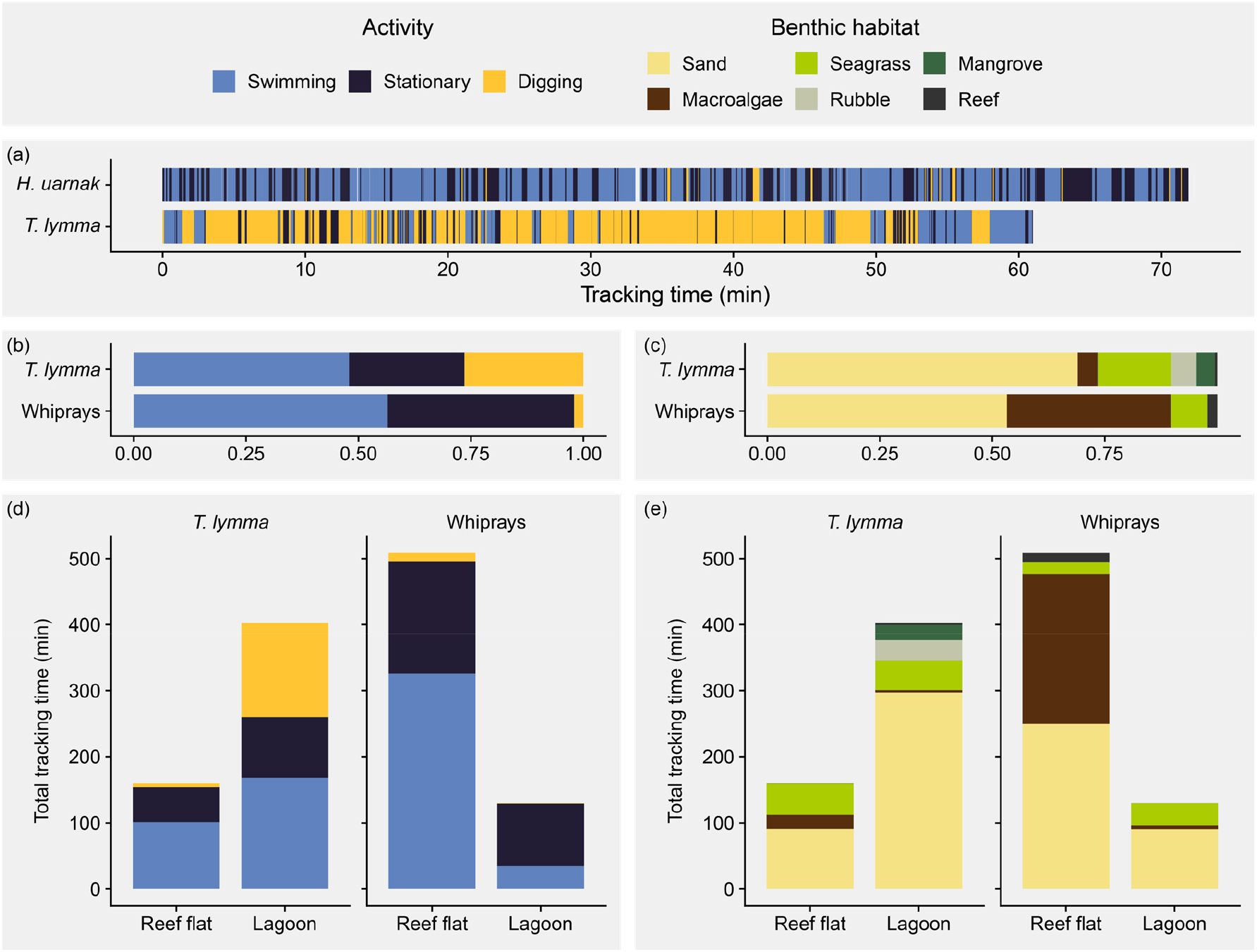
(a) Example activity patterns of a whipray (*Himantura uarnak*) tracked on the reef flat on May 9^th^ and a bluespotted ray tracked in the lagoon on May 10^th^, 2024. (b,c) Relative amount of time spent by both taxa (b) per activity and (c) in each benthic habitat across all tracking surveys. Total time spent by both taxa (d) per activity and (d) in each habitat separated by survey area (Fig. 1). *T.: Taeniura, H.: Himantura*.

Both bluespotted rays and whiprays frequently interrupted swimming with brief pauses indicative of foraging behaviour (Fig. 7a, Video E.1, E.2). In accordance with the transect surveys, excavation feeding was almost entirely restricted to bluespotted rays, which spent 26% of their tracked time digging compared to 2% for the whiprays (Fig. 7b). Almost all digging occurred on the shallow lagoonal sandflat adjacent to the mangroves, where digging accounted for 35% of all activity (Fig. 7d). Habitat use during tracking mirrored transect observations: bluespotted rays spent the most time over sand habitat, while whiprays spent the most time over sand and macroalgae (Fig. 7c). However, note that this was also a function of tracking location (Fig. 7e). Individual maps and activity patterns of the tracked individuals are provided in Appendix F.

### 3.3 Noteworthy observations

Drone footage provided further ecological insights through several incidental observations. For instance, digging bluespotted rays consistently attracted small fish feeding on resuspended material (Video E.1). Whiprays were repeatedly followed by needlefish and in one case by a pair of trevallies (Video E.2, E.3), suggesting opportunistic associations between predatory bony fish and foraging rays (de Carvalho-Souza et al., 2025; Saltzman et al., 2025). This observation was substantiated by 32 whiprays (17%) from the transect surveys being observed with a needlefish close-by (< 2 m). Although drone surveys primarily provided population-level snapshots, some individuals with distinct deformities or unique, stable spot patterns were re-identified across surveys. One bluespotted ray with a symmetrically deformed posterior pectoral fin matched a specimen documented two years earlier by Ciocănaru et al. (2024) on a sandflat four kilometres south of our site (Supplementary fig. C.8). A whipray with a uniquely coiled tail was re-identified five times from May to September 2024, with each observation occurring within a 200 m radius, thereby demonstrating high site-fidelity (Supplementary fig. C.9). Similar site-fidelity was also observed in two other whiprays identified by their unique spot patterns.

## 4 Discussion

Our study reveals fine-scale habitat partitioning between sympatric stingrays in a shallow Red Sea lagoon and shows that this segregation is closely tied to differences in foraging strategy. By integrating drone-based transect surveys with active tracking, we show both population-level habitat associations and high-resolution behavioural patterns. Bluespotted rays aggregated on a shallow, mangrove-adjacent sandflat where they fed through excavation, thereby concentrating their trophic impact and physical disturbance in a restricted area. Larger-bodied whiprays were associated with macroalgal habitat on the adjacent reef flat and mostly fed non-disruptively. Together, these contrasting patterns demonstrate how closely related mesopredators partition habitat at sub-kilometre scales, facilitating coexistence and structuring their ecological impact across the ecosystem. In total, we recorded 1,516 elasmobranch sightings, increasing documented records from the eastern Red Sea by over 50% (Frappi et al., 2024), thereby illustrating how drone-based remote sensing coupled with AI-assisted image analysis can scale ecological inference in nearshore ecosystems.

### 4.1 Ecological drivers & implications

Research on stingray niche partitioning has predominantly focused on dietary differences (e.g. Lim et al., 2019; Martins et al., 2022; Myers et al., 2025; O’Shea et al., 2013; Pardo et al., 2015; Queiroz et al., 2023). While spatiotemporal segregation has been inferred from both the presence (Martins et al., 2022; Myers et al., 2025; Queiroz et al., 2023) and absence (Lim et al., 2019; O’Shea et al., 2013) of dietary specialisation, empirical evidence of fine-scale habitat partitioning remains scarce. Recently, the integration of (drone-based) video monitoring has begun to bridge this gap, revealing nuanced patterns in stingrays’ microhabitat use. For example, in Pioneer Bay, Australia, *P. ater, U. granulatus*, and *P. fai* forage in overlapping areas but differ in their relative use of coral rubble, sand, and mangrove habitats (Kanno et al., 2019; Myers, 2024). Similarly, sympatric populations of *H. australis* and *P. ater* partitioned foraging habitats at sub-kilometre scales on a nearby intertidal sandflat (Crook et al., 2022). Although mixed-species aggregations of *P. ater* and *U. granulatus* at St. Joseph, the Seychelles, and at Shark Bay, Australia, seem to suggest that such spatial segregation is not universal, feeding behaviour was not quantified in these locations, potentially obscuring subtle differences in foraging habitat (Bullock et al., 2024; Semeniuk and Dill, 2006).

Our results extend this emerging evidence by demonstrating fine-scale habitat partitioning in the Red Sea, consistent with ecological trade-offs between prey availability, body-size constraints, and predation risks. Although we cannot directly quantify the contributions of these latent drivers, prey availability likely was a primary determinant of the observed spatial distributions. First and foremost, the mangrove-adjacent sandflat where bluespotted rays were most abundant, was also their primary foraging area (Fig. 6, Supplementary fig. C.6). While prey distributions were not characterised, mangrove proximity may alter infaunal sandflat communities (e.g. Meijer et al., 2021), potentially elevating resource availability for excavation-feeding rays. Furthermore, spatial segregation was linked to interspecific differences in foraging strategies (Fig. 7). The behavioural differences observed here are consistent with previous descriptions of epifaunal feeding in *P. fai* and *H. australis* (Crook et al., 2022; Myers, 2024) and known dietary contrasts between bluespotted rays and the *H. australis*, where bluespotted rays primarily target infaunal annelids while *H. australis* consumes more epifaunal decapods and penaeid shrimp (Myers et al., 2025; O’Shea et al., 2013). Thus, trophic specialization likely reinforced the observed spatial segregation.

Body-size effects further shaped habitat selection. Bluespotted rays (≤ 35 cm disc width) are more vulnerable to predation but less susceptible to stranding than whiprays (≤ 160 cm disc width; Last et al. 2016), which favours use of very shallow predator-refuge habitats. Their strong association with habitats < 0.4 m deep (Fig. 5a) and avoidance of deep open water (Fig. 5c) align with predator-avoidance strategies documented in small batoids (Elston et al., 2021; Flowers et al., 2021; Knip et al., 2010; Leurs et al., 2023). Both juvenile and adult bluespotted rays used this ultra-shallow, mangrove-adjacent habitat during an extremely hot summer (in situ water temperatures > 36°C), demonstrating very high thermal tolerance consistent with prior experimental work in juveniles of the species (Dabruzzi et al., 2013). Whiprays on the contrary did not avoid deep open water (Fig. 5c), which may reflect lower susceptibility to medium-sized predators like the observed blacktip reef sharks. The overall shallow nature of the surveyed area (< 2 m) likely limited access for large top predators. While the conspecific aggregations observed in both taxa may have anti-predatory benefits (Fig. 5e), they probably arose from shared resource use and common responses to environmental drivers rather than social behaviours per se (Jacoby et al., 2012; Semeniuk and Dill, 2006).

Concentrated excavation feeding by bluespotted rays on the mangrove-adjacent sandflat generated a persistent hotspot of predation pressure and bioturbation. Ray-induced sediment disturbance has previously been shown to shape landscape-scale geomorphology (Grew et al., 2024; Nauta et al., 2024; Takeuchi and Tamaki, 2014), restructure infaunal communities (Nauta et al., 2024; VanBlaricom, 1982), and alter nutrient fluxes at the sediment-water interface (D’Andrea et al., 2002; Thrush et al., 1991). Thus, the spatially focused disturbance created by a dense bluespotted ray aggregation may be an important structuring force within the lagoon (Flowers et al., 2021; Grew et al., 2024; Nauta et al., 2024). By comparison, whiprays’ impacts appeared more spatially diffuse and primarily mediated through trophic interactions. Nevertheless, occasional excavation feeding by whiprays resulted in substantial sediment disturbance and much larger feeding pits. Collectively, these observations illustrate how species-specific foraging and habitat selection interact to shape the ecological impact of stingrays across tropical seascapes.

### 4.2 Methodological innovation

This study also sought to address the analytical bottleneck associated with processing large volumes of drone imagery in optically complex marine environments (Axford et al., 2024; Belcher et al., 2023; Katija et al., 2022). The AI-assisted workflow we developed improved both processing efficiency and detection accuracy, identifying 347 rays that had been overlooked by human analysts (Fig. 3a). Most of these were bluespotted rays, which are easily missed due to their small size (≤ 0.35 m disc width) relative to the image footprint (20 × 35 m). Together with heterogeneous backgrounds, wave distortions, sun glint, caustics, and low object-background contrast, this made manual annotation cognitively demanding and prone to oversight (Fig. 3h-k). Our AI-assisted workflow specifically addressed the small-object detection problem, using automated cropping and slicing to enable model training and inference at native resolutions (Fig. 2; Akyon et al., 2022). Moreover, by changing the human role from detection to validation, we minimised the probability of oversight.

Another key factor supporting strong model performance was the use of tracking imagery for training. Tracking imagery closely matched the visual properties of the transect images, minimizing the distribution shift between training data and the intended application. This approach optimised model performance in our specific use-case at the cost of model generalisability. Although fine-tuning can overcome this issue, future research may focus on using larger, more diverse datasets to support transferability across broader geographic and environmental contexts (Axford et al., 2024). To support such efforts, all 109,928 images (of which 3,681 are annotated), have been made publicly available (see Data Availability Statement). In the interim, our combined tracking + transect survey approach provides a practical and reproducible template for rapid deployment of AI-assisted image analysis in targeted environmental monitoring applications.

### 4.3 Limitations and future directions

Variable water clarity, wave-induced distortions, and partial sand coverage hampered reliable species-level identification of whiprays. Several morphologically similar species (*H. uarnak, P. fai, P. jenkinsii*, and *M. ambigua*) co-occur in the Red Sea. Aggregating these species into a single subfamily taxon may obscure ecologically meaningful differences. Complementary approaches, such as telemetry, baited remote underwater video, or e DNA, could help resolve species-level patterns, albeit at coarser spatial resolutions and with smaller sample sizes than drones. Furthermore, to minimize wave-induced distortions and sun glint (Joyce et al., 2019), all drone flights were conducted in the morning hours (6 to 10 am), limiting our ability to detect diel shifts in habitat use and behaviour. Both bluespotted rays and whiprays have been proposed to feed primarily at night (Last et al., 2016; Vaudo and Heithaus, 2012). While bluespotted rays clearly foraged during the day, the absence of excavation feeding by whiprays could reflect unobserved nocturnal activity, particularly since surface feeding could not be identified from the drone footage with absolute certainty. Except for the occasional re-identification of unique individuals, drone-based surveys only capture short observational windows, which limits inference on long-term residency. Nevertheless, the efficiency gains provided by AI-assisted image analysis now pave the way to investigate seasonal or interannual patterns in abundance or habitat use through regularly repeated drone surveys. Finally, the method’s reliance on clear water and calm weather conditions largely constrains its application to tropical regions, but this is also where elasmobranch biodiversity and conservation risks are the highest (Dulvy et al., 2026, 2021; Sherman et al., 2023).

## 5 Conclusion

By combining drone-based remote sensing with AI-assisted image analysis, our study reveals stingrays partition habitat at sub-kilometre scales, demonstrating that ecologically meaningful segregation can arise at fine spatial scales even for highly mobile species. Diverging habitat selection between whiprays and bluespotted rays was linked to contrasting foraging strategies, stranding, and predation risks, and shaped species-specific ecological impacts. Thus, habitat partitioning can facilitate coexistence among sympatric mesopredators and shape the functional landscape of tropical lagoons. Additionally, the methodological advances demonstrated here show how integrating drones with computer vision can transform the spatial and temporal resolution of elasmobranch monitoring. This provides opportunities for population-level assessments and ecosystem-based management as anthropogenic pressures continue to reshape coastal communities.

## Supporting information

Appendix A: details AI workflow

Appendix B: details environmental data

Appendix D: images transect whiprays

Appendix F: maps tracking surveys

## 6 Acknowledgements

We thank Mustapha Ouhssain for assistance during the fieldwork, Ido Pen for advice regarding the statistical analysis, and Afnan K. AlBatati for proofreading the manuscript. For the development of the artificial intelligence model, this research made use of the Ibex high-performance computing cluster managed by the Supercomputing Laboratory at KAUST.

## 7 Author Contributions

**Conceptualization:** Brian O. Nieuwenhuis, Malika Kheireddine, Burton Jones; **Data curation:** Brian Nieuwenhuis, Charlotte Turlier, Ioana-Andreea Ciocănaru, Benja Blaschke; **Formal analysis:** Brian O. Nieuwenhuis; **Funding acquisition:** Burton H. Jones; **Investigation:** Charlotte Turlier, Brian Nieuwenhuis, Ioana-Andreea Ciocănaru; **Methodology:** Brian Nieuwenhuis, Charlotte Turlier; **Resources:** Burton H. Jones; **Supervision:** Malika Kheireddine, Burton H. Jones, Guido Leurs, Jesse Cochran, Laura Govers; **Validation:** Jesse Cochran, Guido Leurs; **Visualization:** Brian O. Nieuwenhuis; **Writing – original draft:** Brian O. Nieuwenhuis; **Writing – review and editing:** All authors.

## 8 Funding

This work was funded by KAUST through the baseline research funds awarded to B. Jones [BAS/1 /1032-01-01].

## 9 Declaration of competing interest

The authors declare that they have no known competing financial interests or personal relationships that could have appeared to influence the work reported in this paper.

## 10 Declaration of generative AI use

During the preparation of this work, the authors used ChatGPT-4 to and Gemini-2.5 to draft Python codes and improve sentence structure. After using these tools, the authors reviewed and edited the output as needed and take full responsibility for the content of the published article.

## 11 Data Availability Statement

The data that support the findings of this study are openly available in the KAUST repository at: http://hdl.handle.net/10754/708833, DOI: 10.25781/ KAUST-33KQ6. This dataset includes the videos of Appendix E. A copy of the annotated AI-dataset is also available on the Roboflow platform at: https://universe.roboflow.com/charliebrian/redsea_stingray_detection.

## Notes

https://universe.roboflow.com/charliebrian/redsea_stingray_detection

http://hdl.handle.net/10754/708833

### Competing Interest Statement

The authors have declared no competing interest.

