## Appendix A: details AI workflow for "Fine-scale habitat partitioning of sympatric stingrays revealed by drone-based remote sensing and deep learning"

### Appendix A:
### Automating drone-based ray detection

To enable efficient and accurate analysis of the 32,216 transect images, we developed an Artificial Intelligence (AI)-assisted workflow based around Ultralytics' YOLOv11x object detection model (Jocher and Qiu, 2024). An overview of the full workflow is given in Figure 2 of the main manuscript, which has been reprinted below for convenience. The following sections explains the individual steps in detail.

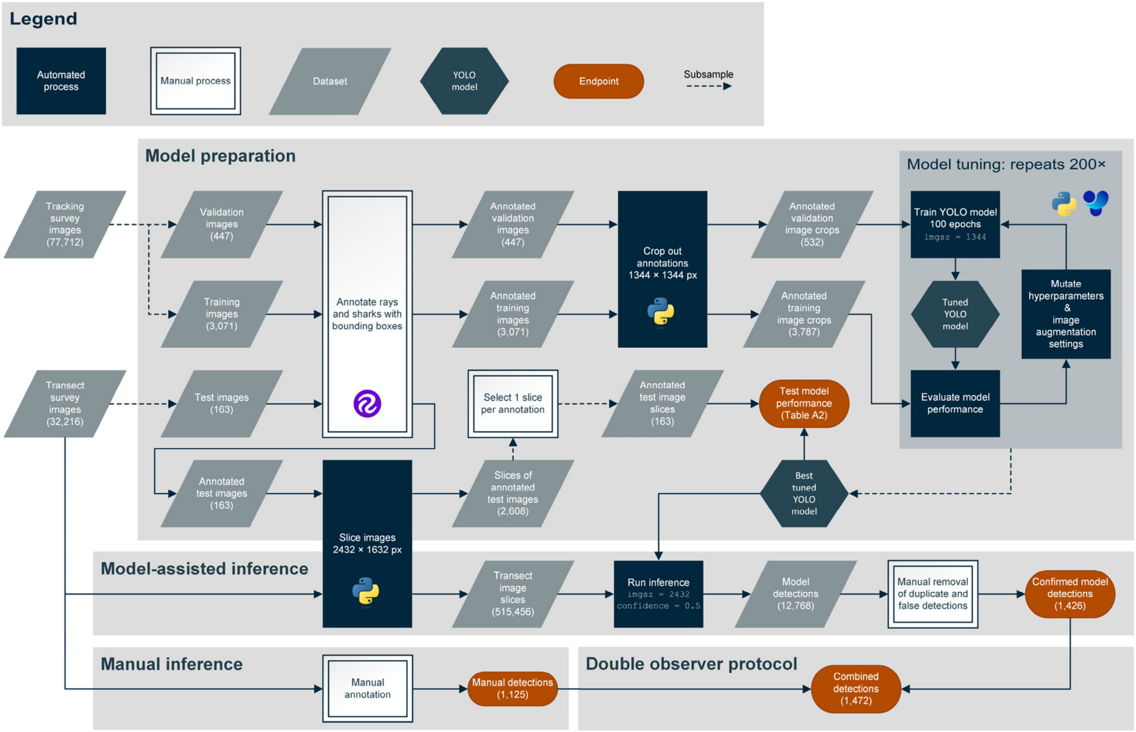


*Figure 2*. Overview of the image analysis conducted for the transect imagery. For visual clarity only the development of the final YOLO-based model is depicted. Note the terminology for different image sizes: "image" = 8192 × 5460 px (full size drone image), "image slice" = 2432 × 1632 px, "image crop" = 1344 × 1344 px. Logos indicate the reliance of different steps on Roboflow, Python, and Ultralytics' YOLO software (Dwyer et al. 2024; Jocher and Qiu 2024). '`imgsz`': image size parameter in YOLOv11x model.

#### Dataset creation

An AI-ready dataset consists of three components, the training, validation, and test partition. Since our ultimate aim was to automatically detect elasmobranchs in the transect imagery, the test dataset was constructed from five manually selected transect surveys that together encompassed all classes. To create an unbiased test set, only manually detected rays were included. The training and validation datasets were instead sourced from the tracking surveys. This allowed the model to be trained on imagery that was highly similar to, yet independent from the transects. To ensure consistency with the transect imagery, only images taken with the DJI Zenmuse P1 camera were included (SZ DJI Technology Co. Ltd., 2021). The use of tracking imagery also facilitated efficient annotation, because each image inherently contains a ray (or shark) in the centre of the frame. However, if any additional individuals were observed, these were also annotated. To reduce annotation effort and prevent overrepresentation of similar images, surveys tracking whiprays or bluespotted rays were subsampled at a frequency of 30 seconds, while tracks of data-limited classes like the spotted eagle rays, mangrove whiprays, cowtail rays, and sharks were subsampled at higher frequencies. To reduce overfitting, tracking imagery was partitioned between training and validation datasets at the survey level. Since cowtail rays and sharks were only tracked once, these surveys were randomly assigned to the training and validation datasets in a 90/10 ratio. Finally, to increase image diversity, the training dataset was supplemented with images known to contain stingrays from local Structure-from-Motion surveys (Nieuwenhuis, unpublished data) and (Ciocănaru et al., 2024). In total 3,683 images were annotated using the online Roboflow platform (Dwyer et al., 2026), yielding 4,043 training, 558 validation, and 209 test annotations, respectively (Table A.1).

*Table A.1*. The number of bounding box annotations and their source in the AI-ready dataset separated by class and dataset partition.

| Dataset partition | Image Source | Annotation count (bounding boxes) | | | | | | |
|---|---|---|---|---|---|---|---|---|
| | | Bluespotted rays | Whiprays | Eagle rays | Mangrove whiprays | Cowtail rays | Sharks | **Total** |
| Training | Tracking surveys[1] | 1683 | 1168 | 367 | 353 | 394 | 78 | **4043** |
| Validation | Tracking surveys | 274 | 182 | 19 | 31 | 44 | 8 | **558** |
| Test | Transect surveys | 154 | 40 | 7 | 5 | 4 | 3 | **213** |

[1] Supplemented with imagery known to contain stingrays from local Structure-from-Motion surveys (Nieuwenhuis, unpublished data) and (Ciocănaru et al., 2024).

### Image scaling

One of the primary challenges to the application of computer vision models to remotely sensed imagery such as our drone images is the “small object detection problem” (Akyon et al., 2022; Axford et al., 2024). Memory requirements increase exponentially with image size and thus computer vision models are generally trained on low-resolution or down-scaled images (Akyon et al., 2022; Axford et al., 2024; Jocher and Qiu, 2024). By default, YOLO-based models resize images to 640 × 640 px. With a maximum disc width of 0.35 m bluespotted rays only span ~1% of the full width of our drone images (image swath = 20 × 35 m). Thus, a bluespotted ray would be just 6 pixels wide in the downscaled image, leaving insufficient detail to effectively train a computer vision model (Axford et al., 2024).

To overcome this limitation, we developed a Python script that automatically cropped the annotated images based on the YOLO-formatted labels and outputs square crops (1344 × 1344 pixels) with updated label files (Fig. A.1). Crop size was determined to be approximately twice the total length of the largest ray and compatible with the 32-pixel stride of YOLO-based models. The script was set up to ensure that if multiple rays were present in a single image, each bounding box annotation was retained in at least one crop, without creating any crops that included partial or duplicated bounding boxes. Due to geometric constraints, the above logic failed once, and a single bluespotted ray annotation (0.06%) was duplicated. Cropped images were subsequently used to train/tune the YOLOv11x model with the image size parameter set to 1344, enabling model training at the images' original resolution. To also run inference at native resolution, transect images were tiled into 16 equal slices of 2432 × 1632 pixels with 20% overlap (Fig. A2), essentially implementing a 'Slicing Aided Hyper Inference' workflow (Akyon et al., 2022).

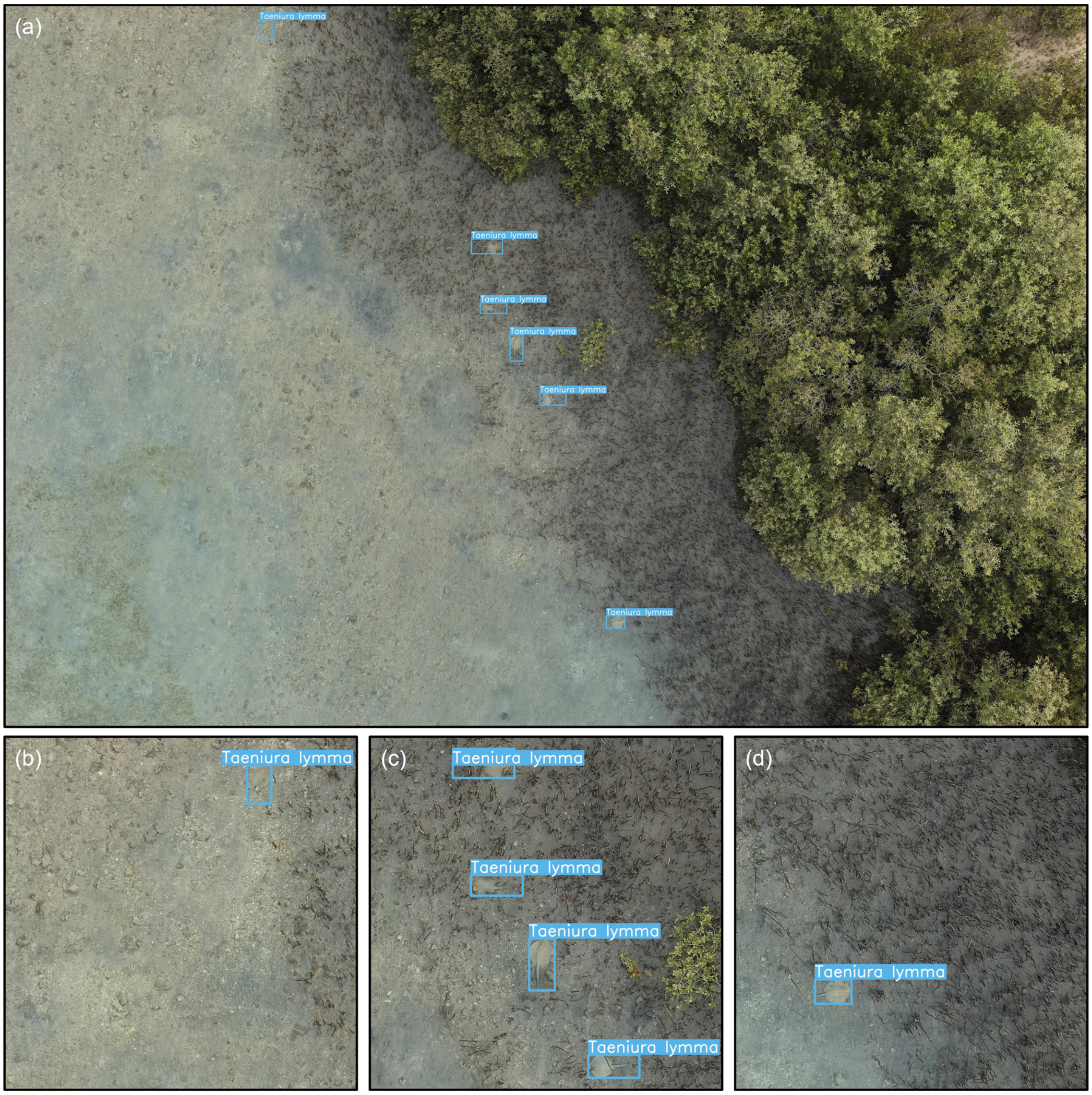


*Figure A.1.* Example of the cropping workflow. (a) The original 8192 × 5460 px drone image (DJI_20240624075641_0541.jpg) in which six bluespotted rays have been annotated with bounding boxes. (b-d) The 1344 × 1344 image crops used for model training.

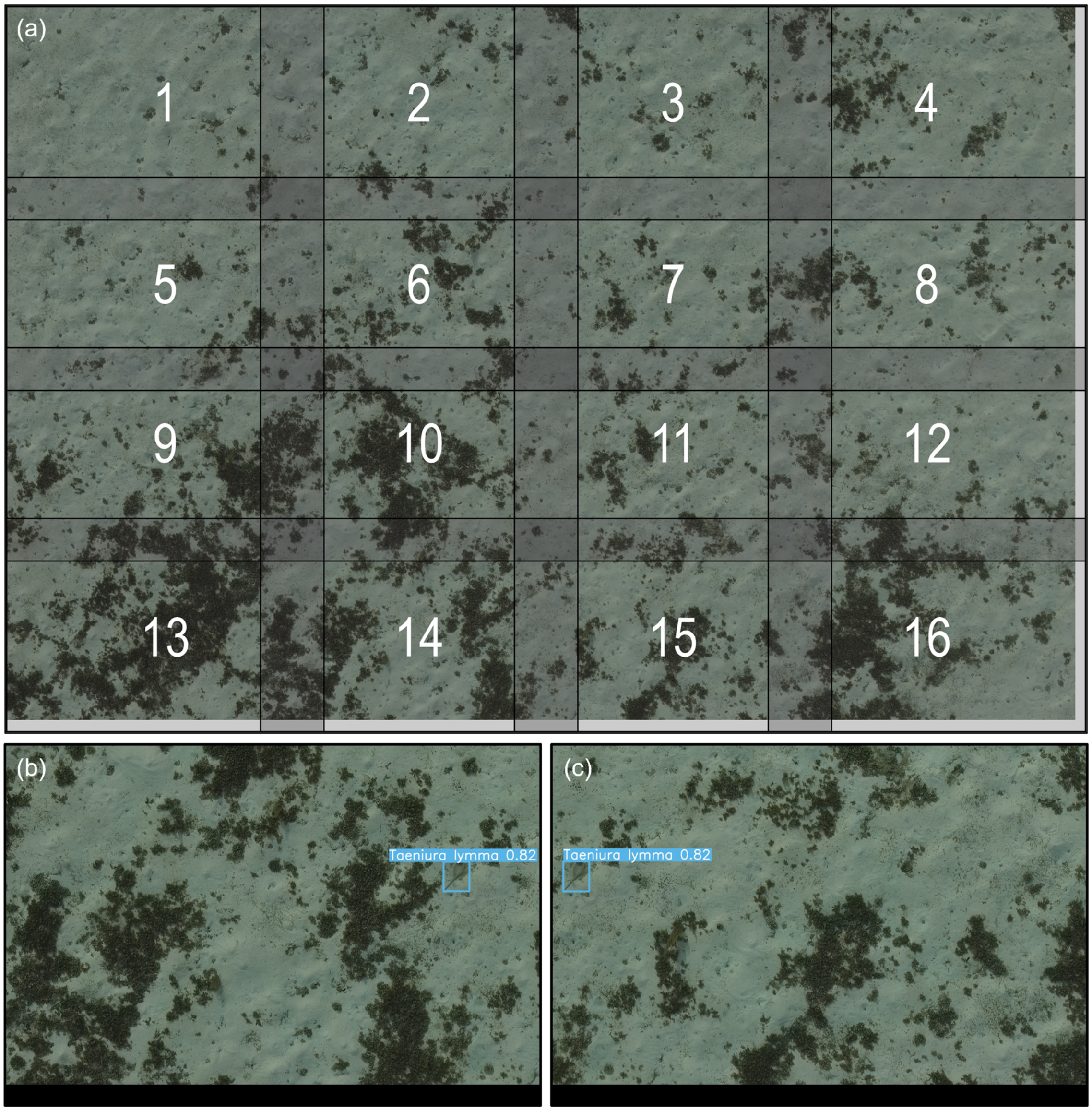


*Figure A.2.* Example of the slicing workflow. (a) The original 8192 × 5460 px drone image (DJI_20240510071325_0033.jpg) overlayed with the extend of the 16 slices. (b,c) The output of the YOLO-model after running inference on all 16 slices; (b) corresponds to slice 14, (c) corresponds to slice 15. Numbers indicate model confidence.

#### Model training and tuning

The YOLOv11x model was trained on KAUST's high-performance computing cluster, Ibex, utilizing NVIDIA A100 GPUs and a Python v3.13.3 environment with CUDA v11.8. To optimize the performance, the model's hyperparameters and image augmentation settings were iteratively tuned using YOLO's inbuilt tuning function (Fig. A.3; Jocher and Qiu, 2024). The model was tuned for 200 iterations of 100 epochs each, using an `AdamW` optimizer and a batch size of 16. We used the default hyperparameter/image augmentation search space except for the parameters '`degrees`', '`flipud`', and '`fliplr`', which were fixed to 180, 0.5, and 0.5, respectively. This ensured the image could be freely rotated during the tuning process to help the model recognise rays regardless of their orientation in the image.

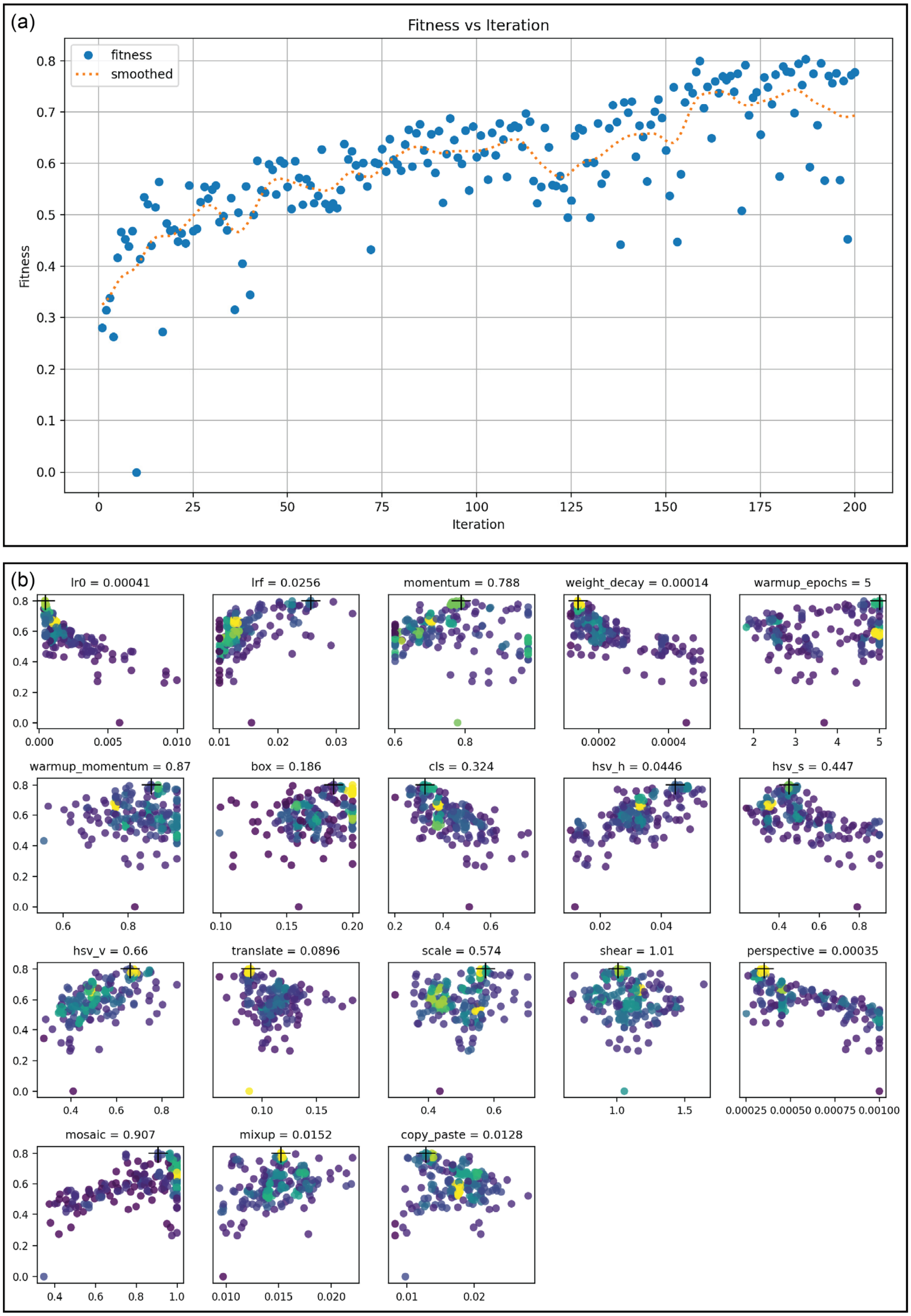


*Figure A.3.* Default plots output by the YOLO11-model to summarise the model tuning processes. (a) Development of model fitness (0.1 × mAP-0.5 + 0.9 × mAP-0.5:0.95) over the tuning iterations. (b) Relation between hyperparameter/image augmentation values and model fitness over the 200 tuning iterations, parameter values for best performing model are indicated in sub-plot titles. Fitness calculations are based on the validation dataset.

#### Model testing

To validate the benefit of our cropping and slicing approach and model tuning, we compared the performance of the tuned model to models trained on full-sized images and with default hyperparameters. If the model was trained on full-sized images, full sized images were also used in the test set. If the model was trained on cropped images, a 2432 × 1632 slice of the test image was used, emulating our sliced inference approach. Performance of the YOLO-based models increased substantially with image resolution (Table A.2). Increasing the image size setting ‘`imgsz`’, from the default 640 to 1344 pixels more than doubled mean Average Precision (mAP-0.5) from 0.22 to 0.51. The incorporation of the cropping and slicing workflow further increased mAP-0.5 to 0.65, finally hyperparameter tuning increased mAP-0.5 of the best model (iteration 187) to 0.85 (Table A.2).

*Table A2*. Evaluation of YOLOv11x models on the transect-based test set. All metrics are macro averages across all six classes. Precision and Recall are calculated at Intersection over Union = 0.5. ‘`imgsz`’: image size parameter in YOLOv11x model. The final model is indicated in bold.

| Training dataset | Training ‘`imgsz`’ | Test dataset | Test ‘`imgsz`’ | Hyper-parameters[1] | Precision | Recall | mAP-0.5 | mAP-0.5:0.95 |
|---|---|---|---|---|---|---|---|---|
| Full size[2] | 640 | Full size | 640 | Default | 0.500 | 0.227 | 0.217 | 0.088 |
| Full size | 1344 | Full size | 1344 | Default | 0.552 | 0.463 | 0.512 | 0.241 |
| Cropped[3] | 1344 | Sliced[4] | 2432 | Default | 0.758 | 0.600 | 0.648 | 0.294 |
| **Cropped** | **1344** | **Sliced** | **2432** | **Tuned 200 iterations** | **0.775** | **0.807** | **0.846** | **0.657** |

[1] Includes image augmentation settings. [2] Full size images: 8192 × 5460 px.[3] Cropped images: 1344 × 1344 px. [4] Sliced images: 2432 × 1632 px.
