## Appendix B: details environmental data for "Fine-scale habitat partitioning of sympatric stingrays revealed by drone-based remote sensing and deep learning"

### Appendix B: Remotely sensed environmental data

For the analysis of the transects, four environmental parameters were derived from drone-based data. Namely, bathymetry, benthic habitat, distance to open water, and distance to the mangroves. The following section describes how these environmental parameters were extracted.

- Bathymetry (water depth):

Bathymetric maps of the study area were generated following the methodology and using the data associated with Nieuwenhuis et al. (2026, 2025). For consistency we only use data that was generated with the 'Zenmuse P1' drone setup also used in this study. Specifically, we re-use the ground truth water depth measurements together with the green and red reflectance maps to create bathymetric maps of the reef flat and the lagoon through Stumpf's bathymetric band ratio (Eq. B.1) (Stumpf et al., 2003).

$$\text{Stumpf's bathymetric band ratio } = m_1 \times \frac{\ln\big(1000 \times R_{\mathrm{rs}}(Red)\big)}{\ln\big(1000 \times R_{\mathrm{rs}}(Green)\big)} - m_0 \qquad (B.1)$$

Where $m_1$ and $m_0$ are calibration coefficients determined by linear regression with ground truth data and $R_{\mathrm{rs}}$ is the remotely sensed reflectance in the green or red image band, respectively.

As the data from Nieuwenhuis et al. (2026, 2025) does not cover our full study area, we conducted two additional Structure-from-Motion (SfM) surveys that partially overlapped with the original surveys. SfM surveys were processed as described in Nieuwenhuis et al. (2026), but did not use Ground Control Points. Bathymetry in the extended areas was acquired by regressing the Stumpf's bathymetric band ratio of the newly created reflectance maps with the pre-existing bathymetric maps in the overlapping areas.

In areas shallower than 0.35 m SfM-derived bathymetry was used instead of the spectrally derived bathymetry described above. Bathymetry measurements were adjusted for temporal variations in water height by comparing the SfM-derived elevation at the water's edge in image 351 of each survey, which features a human-made, emerged structure. For analysis we extracted the median bathymetry value in each 5 × 5 m grid cell after removing any negative (above water) values, for example caused by mangrove trees (Fig. B.1a).

- Benthic habitat:

5 × 5 m grid cells were manually assigned one of four habitat classes: 'sand', 'seagrass', 'macroalgae', and 'reef' (Fig. B.1b). Habitat assignment was based on visual assessment of drone-derived orthomosaics of the study area from June 2024 (median survey date was June 17$^{\mathrm{th}}$). Since sand is the most common habitat, other habitats were considered as 'dominant' over sand if they covered at least 20% of the grid cell. Benthic habitat was only assigned to grid cells shallow enough to clearly see the bottom, if this was not the case grid cells were excluded from statistical analysis.

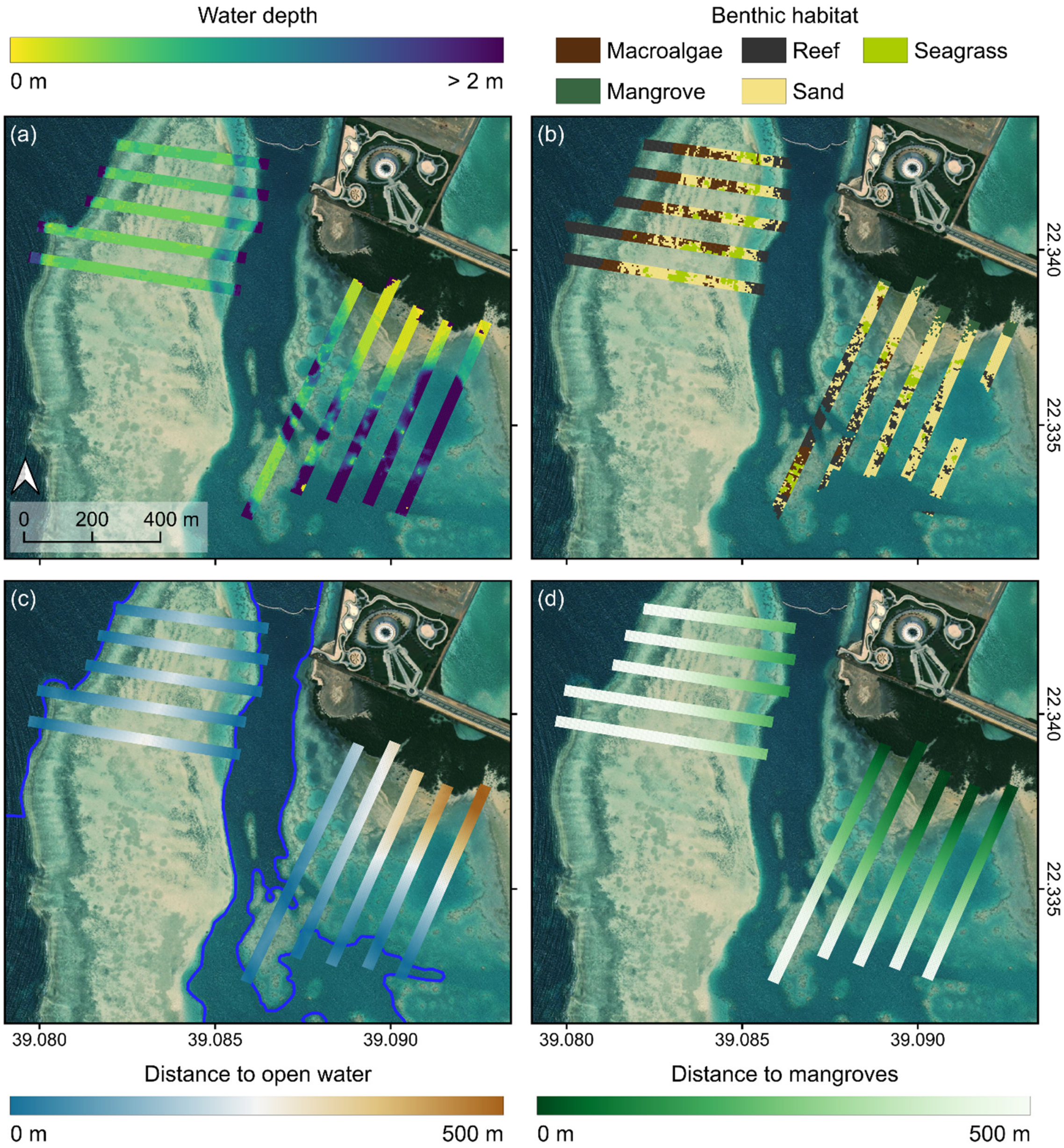


*Figure B.1*. Maps of the environmental parameters along the transect surveys (a) water depth, (b) benthic habitat, (c) distance to open water (delineated in dark blue), and (d) distance to mangroves. (b) Mangroves are indicated for visual clarity but were combined with the sand habitat in the analysis. Map contains ESRI satellite imagery.

- Distance to open water:

To calculate distance to open water, deep, open water was manually delineated in our study area in QGIS (Fig. B.1c). The distance from the centre of each grid cell to the edge of the polygon was subsequently calculated, utilizing the *terra v.1.7-71* library in R (Hijmans et al., 2020; R Core Team, 2022).

- Distance to mangroves:

To calculate distance to mangroves, a mangrove mask was created based on the UAV-derived orthomosaics and SfM-based elevation maps of the lagoon area. Mangrove area was delineated through a manually adjusted threshold classification based on SfM-derived elevation and the Excess Green vegetation Index (Eq. B.2).

$$\text{Excess Green Index (ExG)} = 2 \times R_{\text{rs}}(Green) - R_{\text{rs}}(Red) - R_{\text{rs}}(Blue) \qquad (B.2)$$

After conversion to a polygon, the distance from the centre of each grid cell to the edge of the polygon was calculated as described for the distance to open water (Fig. B.1d).
