## Appendix C: supplementary figures for "Fine-scale habitat partitioning of sympatric stingrays revealed by drone-based remote sensing and deep learning"

### Appendix C:
### Supplementary figures

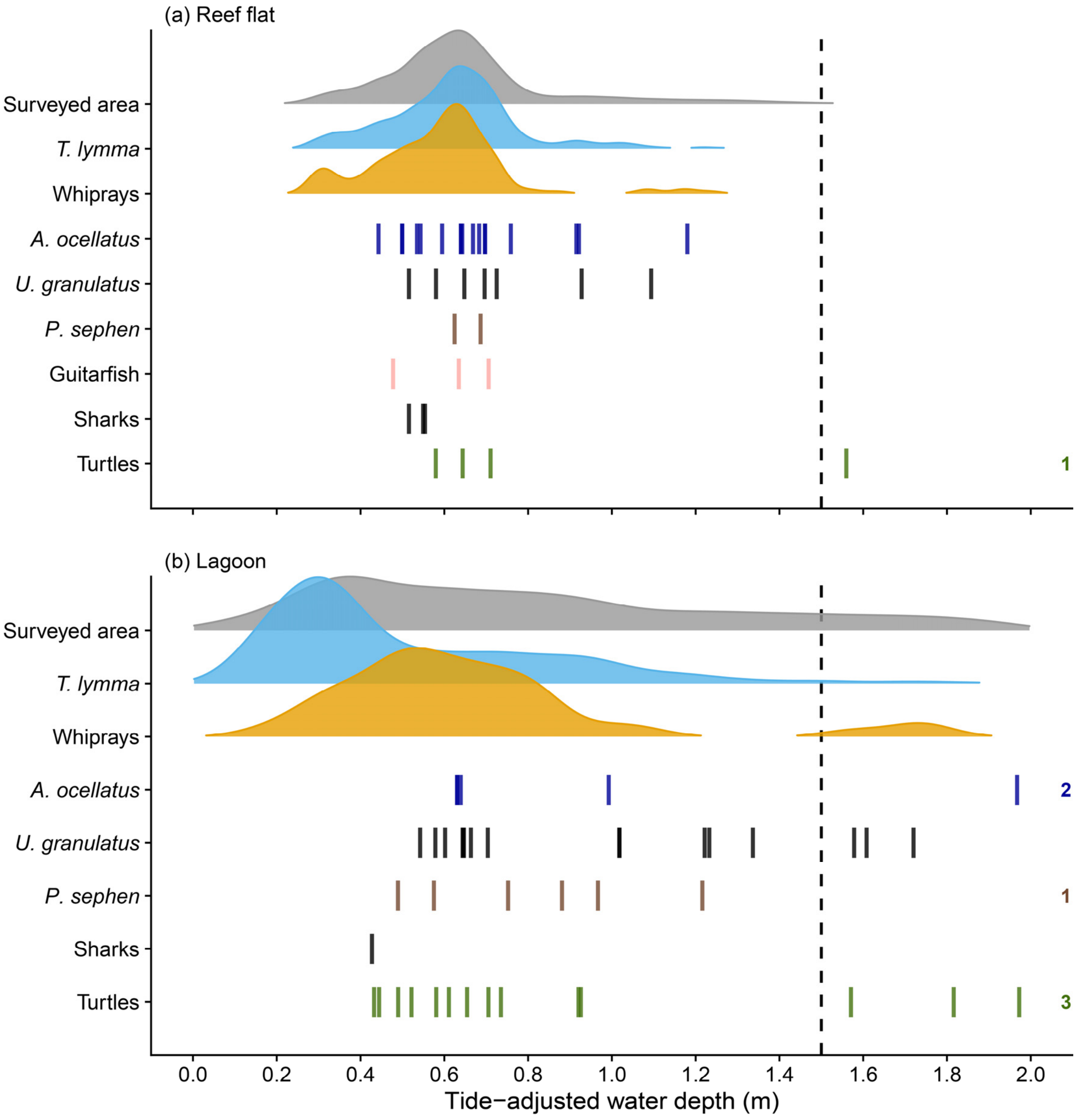


*Figure C1*. Density plots of the bottom depths of the animal detections in the transect surveys for the (a) reef flat, and (b) the lagoon. Depth distribution of the surveyed area is depicted in grey for reference. For the rarer taxa all observations are plotted separately. Dashed line at 1.5 m indicates depth cutoff for the statistical analysis. Numbers on the right indicate individuals observed in water deeper than 2 m.

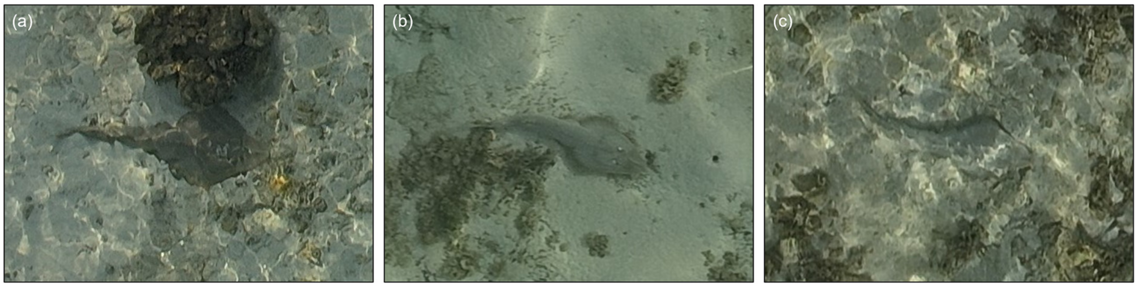


*Figure C2*. The three guitarfish observed on the reef flat during the transect surveys on (a) May 17th, (b) June 6th, and (c) July 3rd, 2024. The total length of the three individuals measured (a) 116 cm, (b) 90 cm, and (c) 88 cm, respectively. The individuals in (b, c) were identified as *Glaucostegus halavi*. The identity of the individual in (a) remained uncertain.

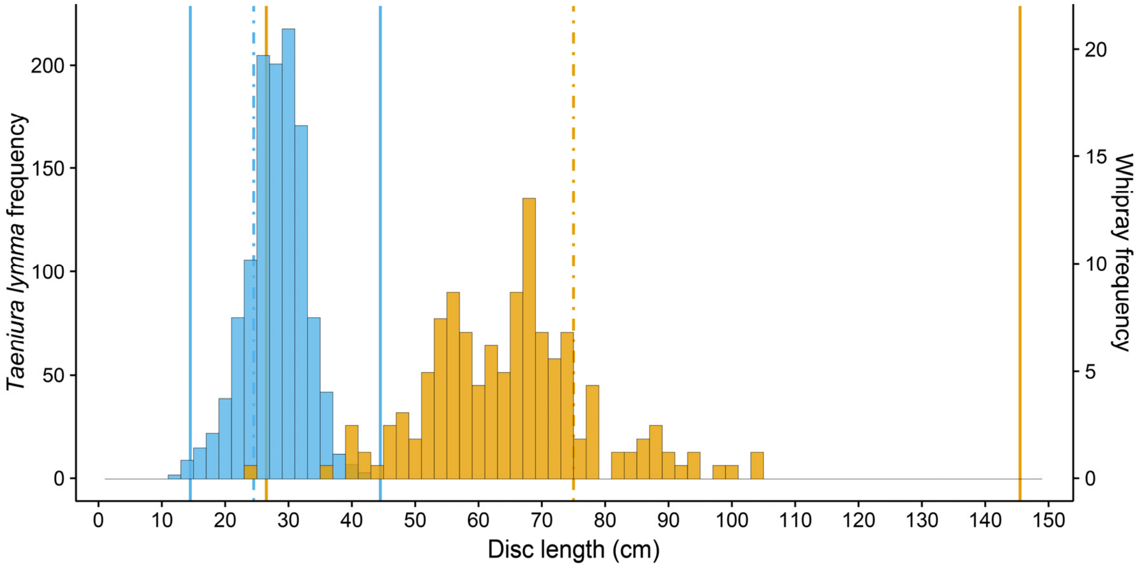


*Figure C3*. Histogram of the disc lengths measured for the bluespotted rays (*Taeniura lymma*) and whiprays observed in the transect surveys. Solid vertical lines depict minimum and maximum disc length,converted from disc widths in Last et al. (2016) using the co-published width:length ratios. Dashed vertical line indicates disc length at maturity. For the whiprays, the estimates for *Himantura uarnak* are displayed. Disc length is displayed instead of disc width because it could be measured more reliably from the drone's images.

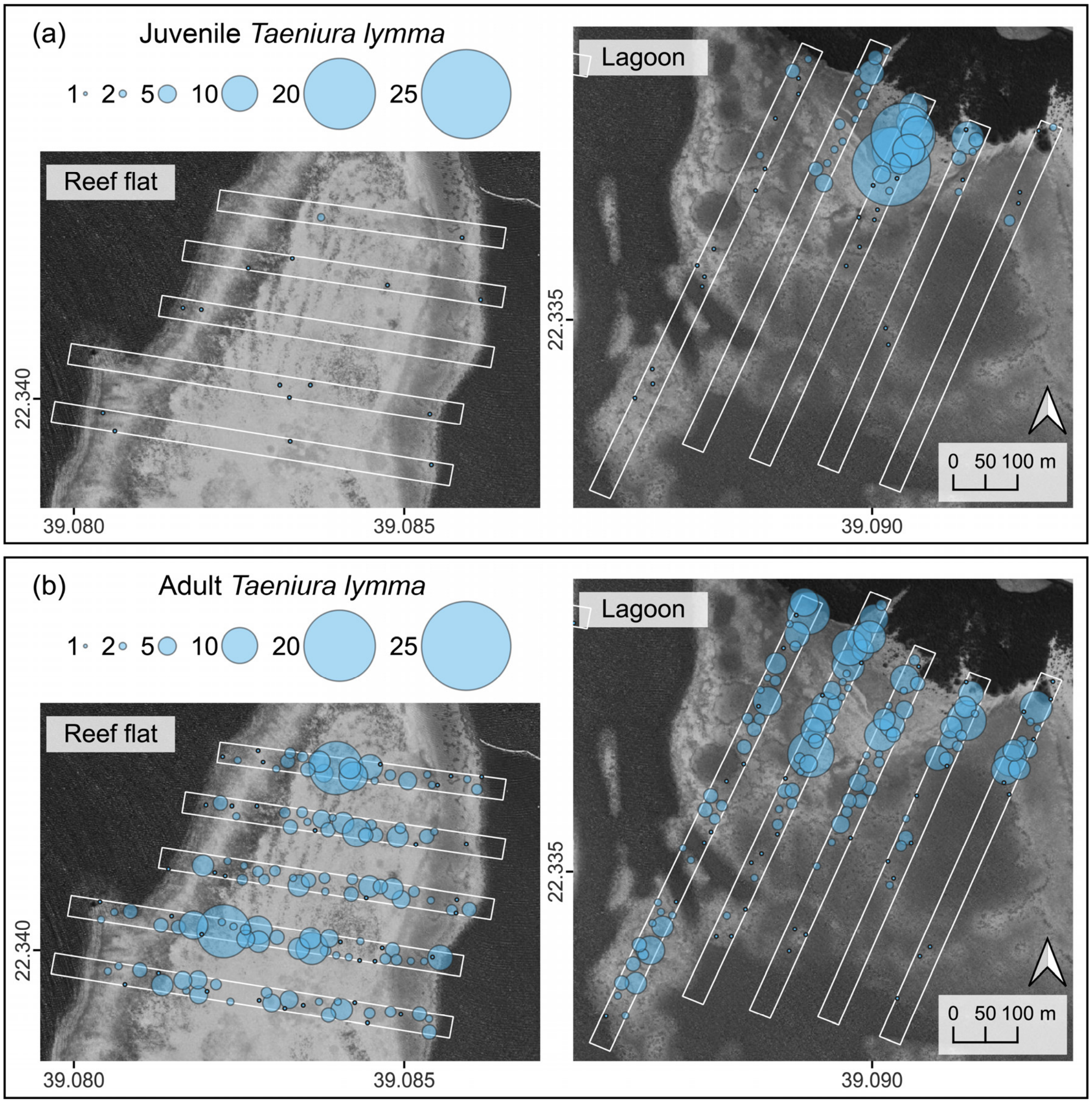


*Figure C4*. Bubble plots showing the relative observation densities across the transect surveys for (a) juvenile and (b) adult bluespotted rays (*Taeniura lymma*). Map contains ESRI satellite imagery.

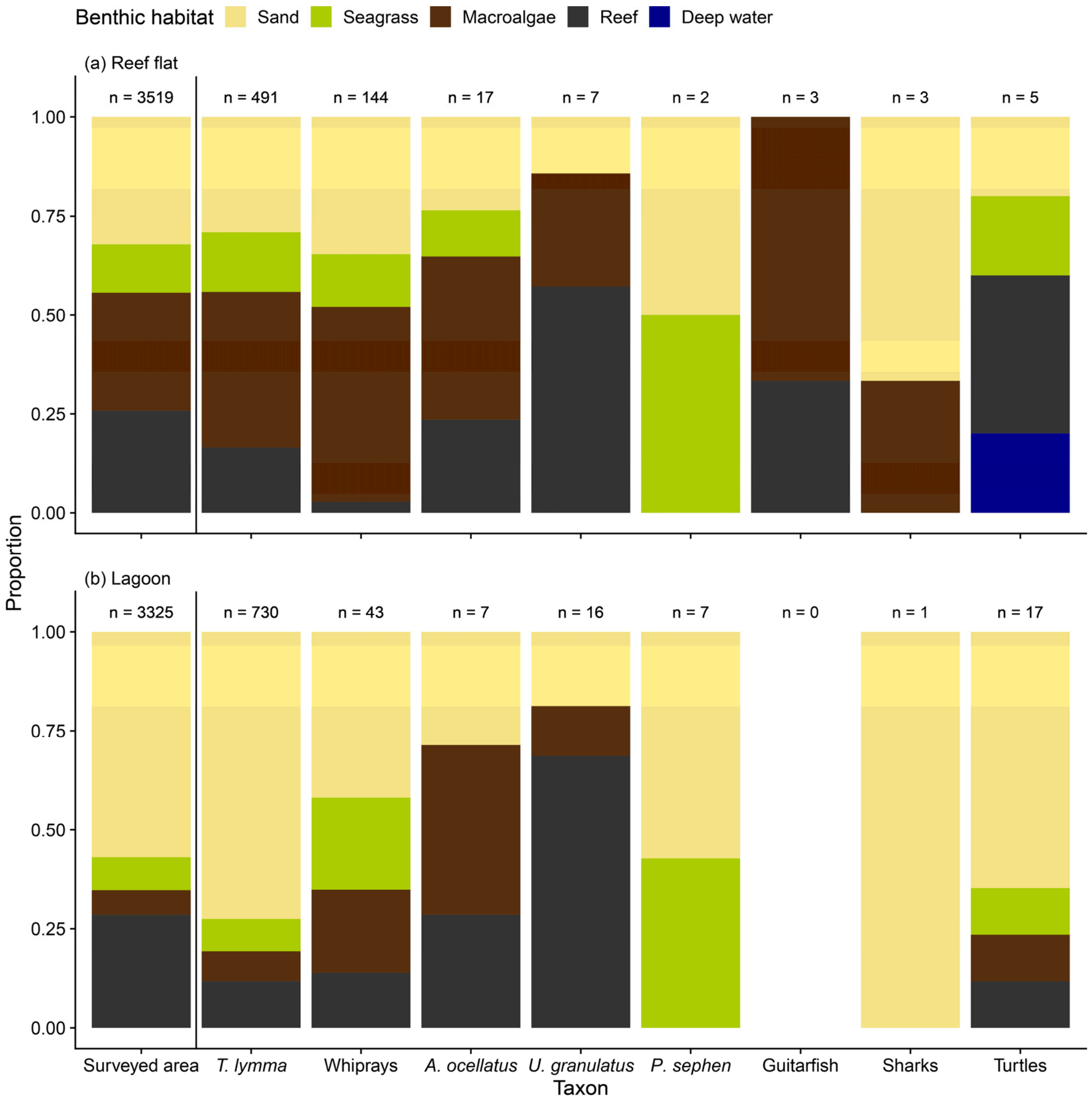


*Figure C5*. Benthic habitat use of different taxa observed along the transect surveys on (a) the reef flat and (b) in the lagoon. The available habitats in the surveyed area are indicated left of the vertical line. Deep water indicates animal was observed in water too deep for the bottom to be visible.

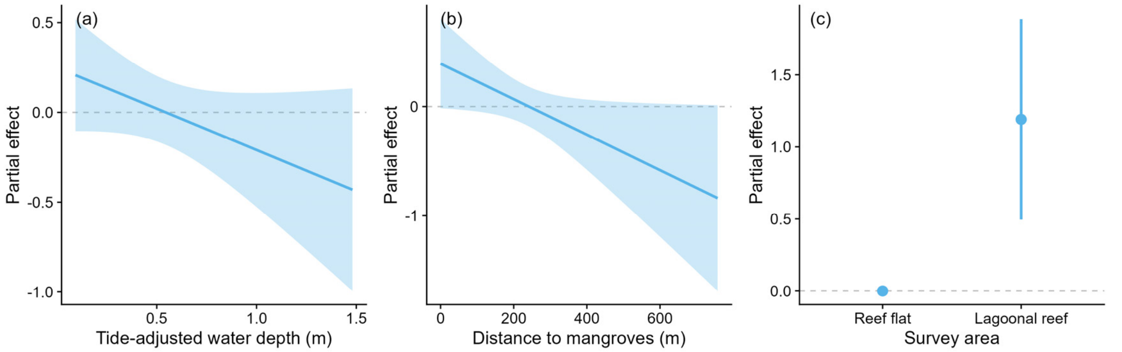


*Figure C6*. Selected partial effect plots showing the effect of (a) bathymetry, (b) distance to mangroves, and (c) survey area on the digging probability for bluespotted rays (*Taeniura lymma*) from the transect surveys. Partial effects are shown on the link scale.

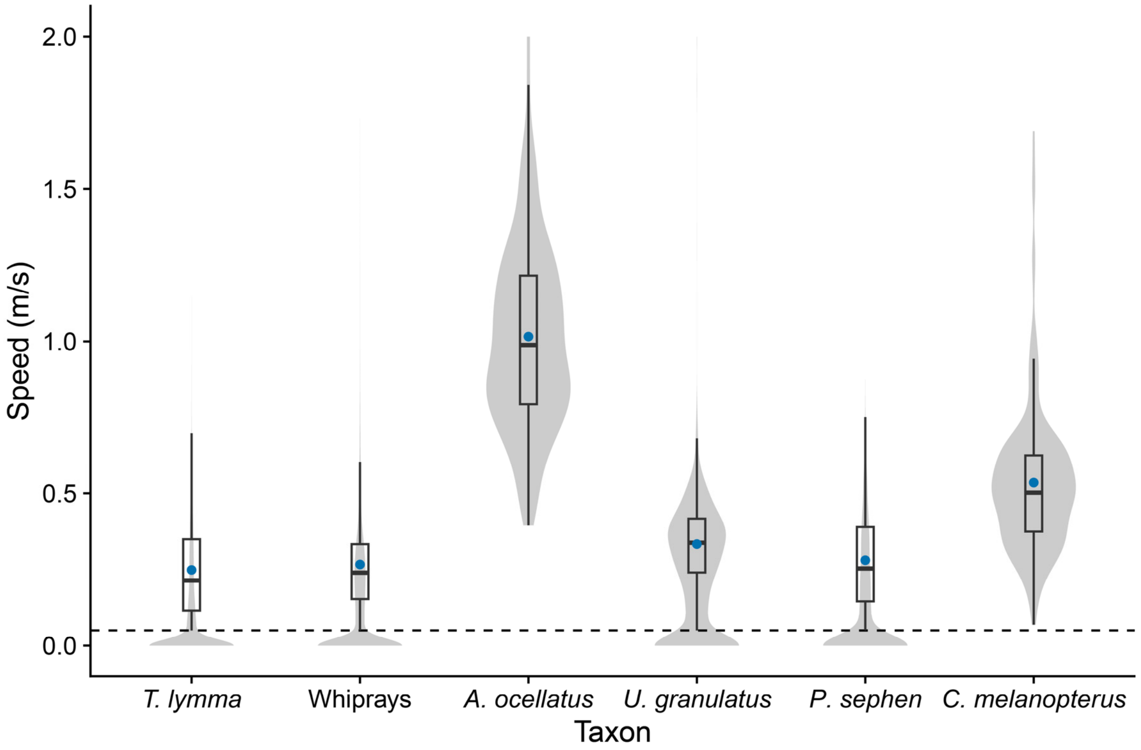


*Figure C7*. Approximate swimming speeds of the tracked rays extracted from the drone's geolocation data. Violin plots show the full distribution, boxplots and averages (blue dots) only consider speeds > 0.05 m $s^{-1}$ (dashed line) to exclude periods without movement.

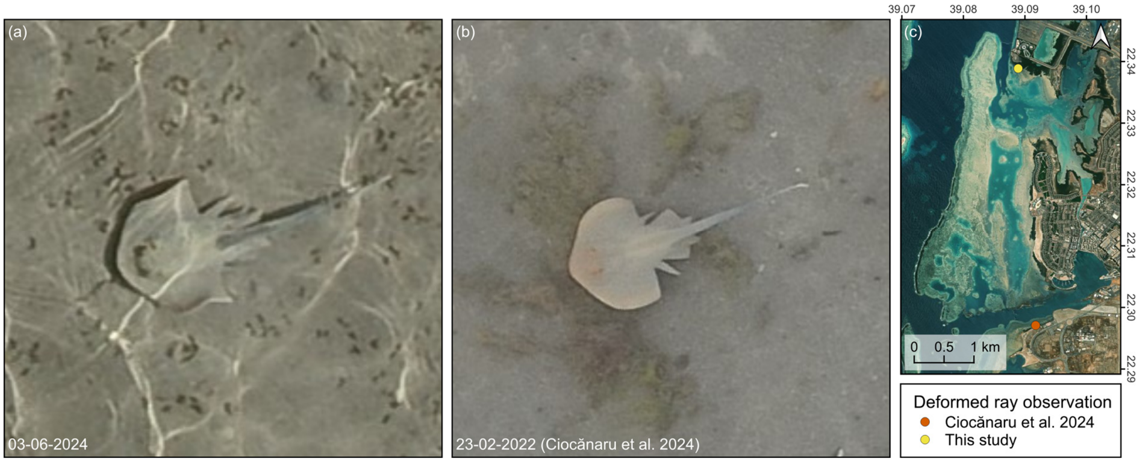


*Figure C8*. Photo comparison between (a) the deformed ray spotted during one of the tracking surveys conducted for this study and (b) the deformed ray described by Ciocănaru et al. (2024). (c) The respective locations of the observations.

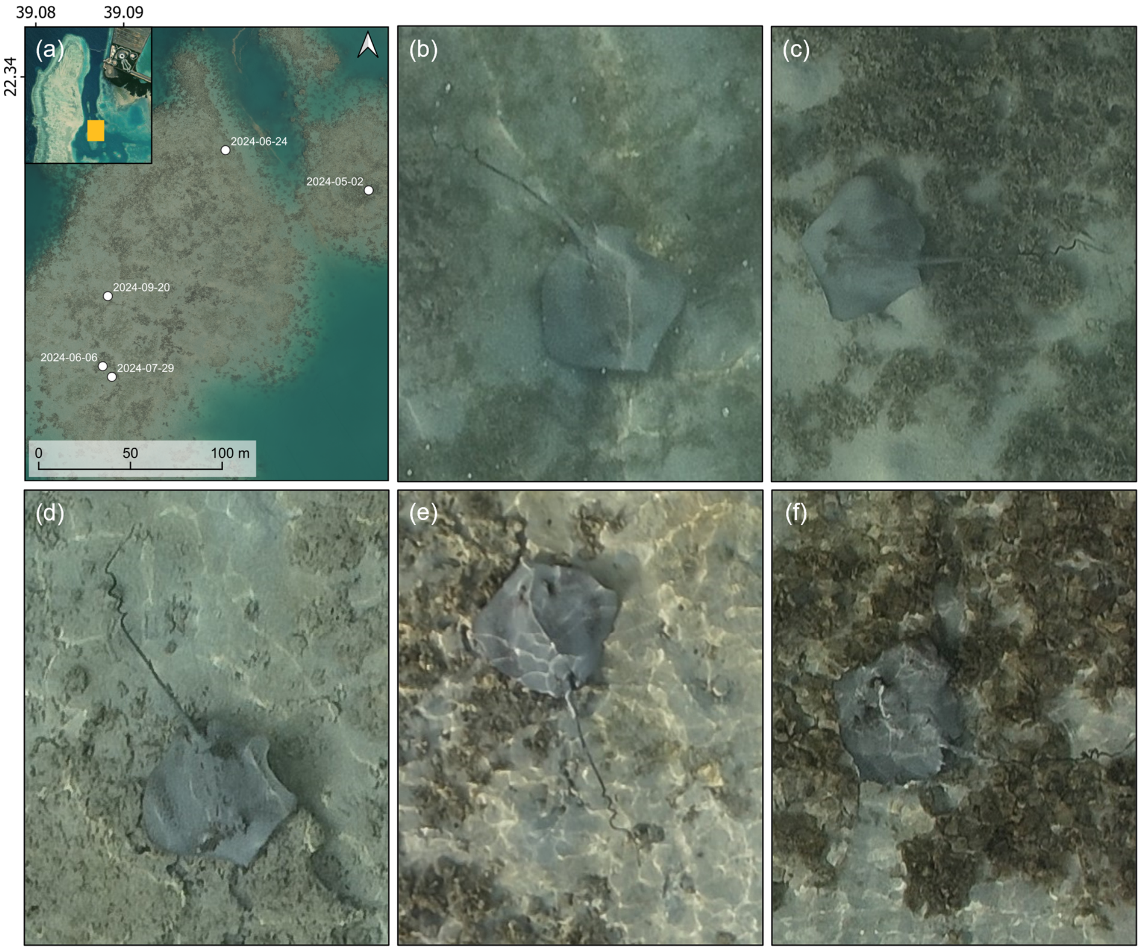


*Figure C9*. Repeated observations of a whipray with a coiled tail deformation. (a) The respective locations of the different observations with the inset showing the general location of the larger map within the study area (Fig. 1). (b-f) Pictures of the whipray on (b) May 2nd, (c) June 6th, (d) June 24th, (e) July 29th, and (f) September 20th, 2024.
