## Appendix D: images transect whiprays for "Fine-scale habitat partitioning of sympatric stingrays revealed by drone-based remote sensing and deep learning"

2024-05-02

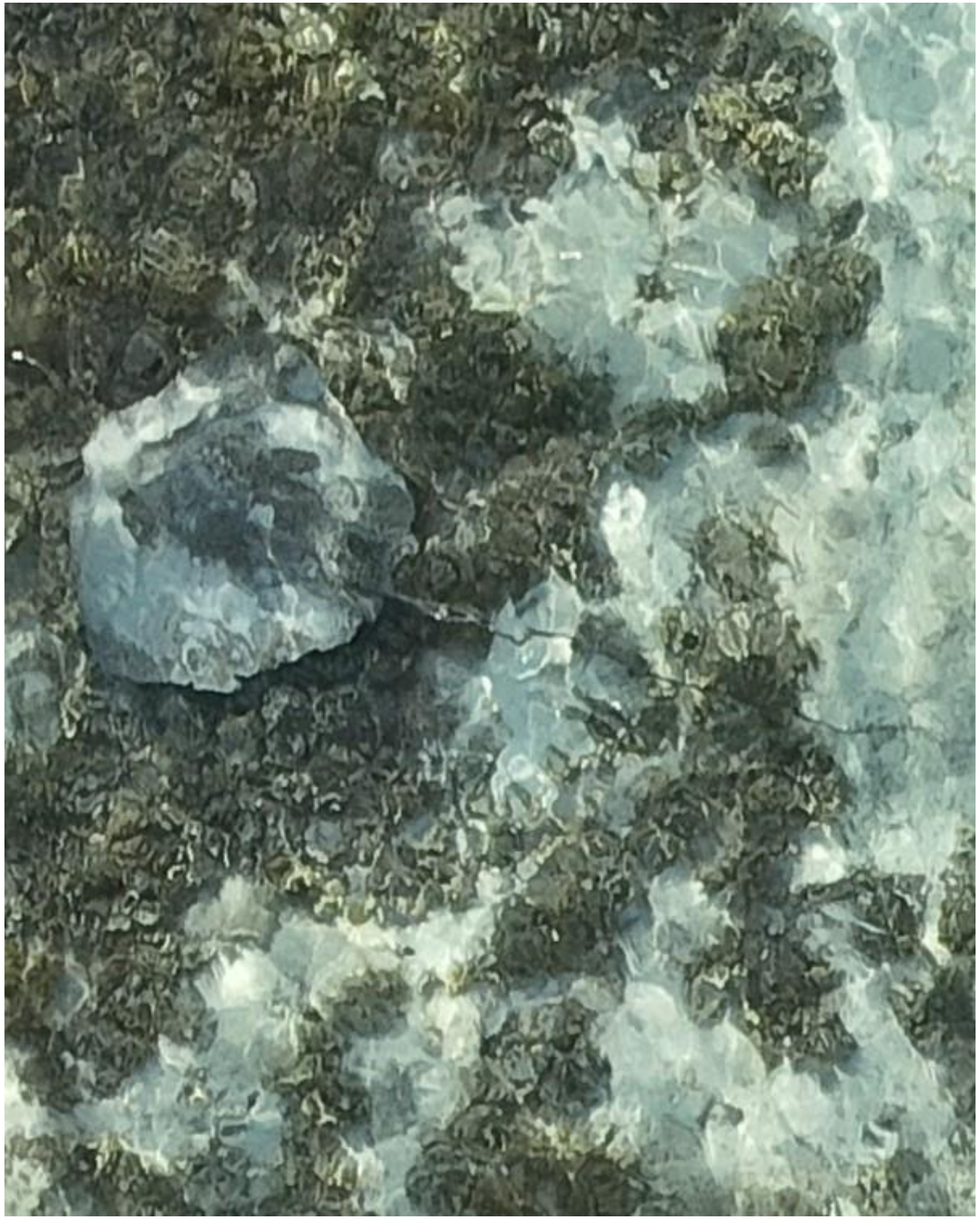

- Whipray

DJI_20240502071710_0178

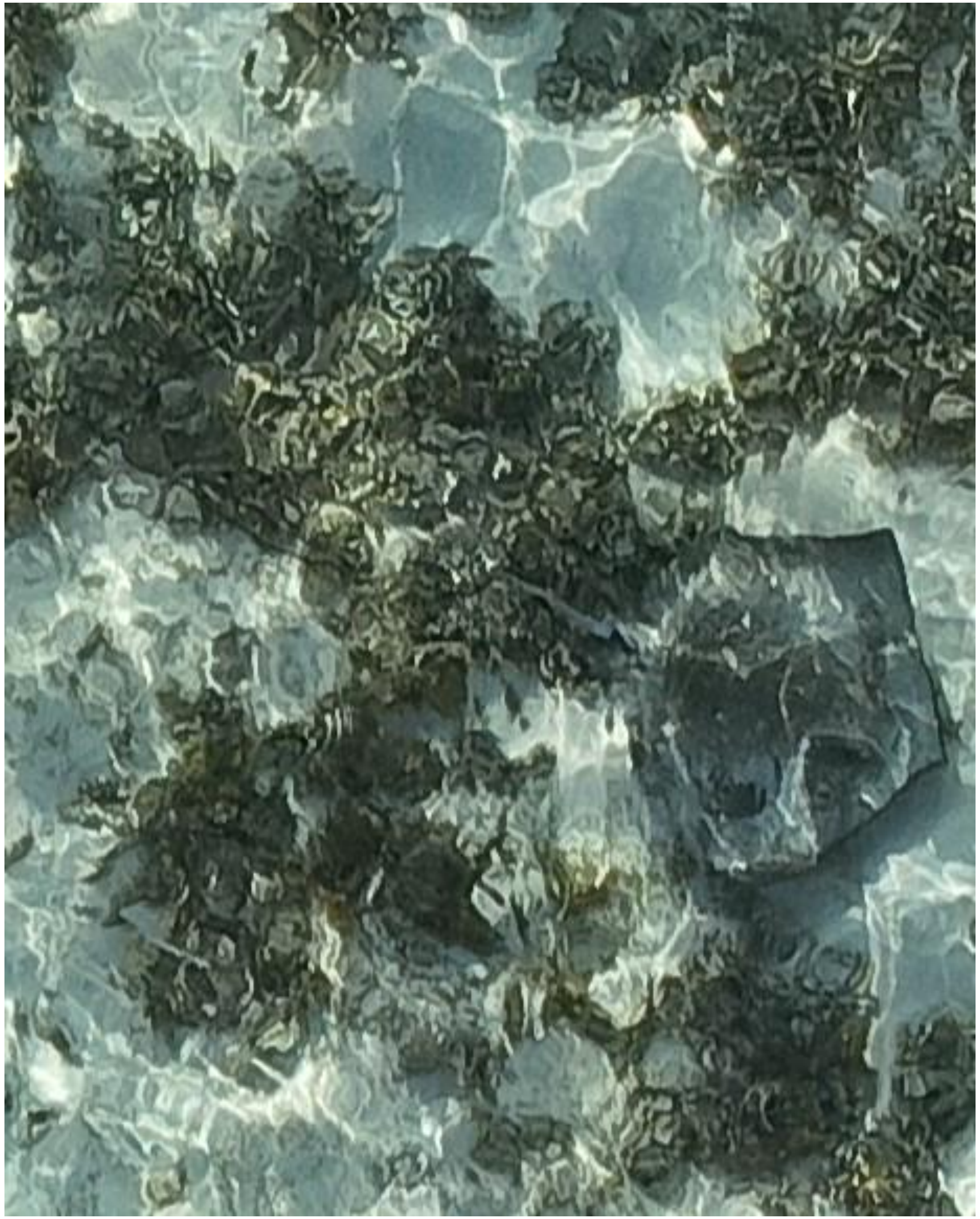

- Whipray

DJI_20240502072101_0386

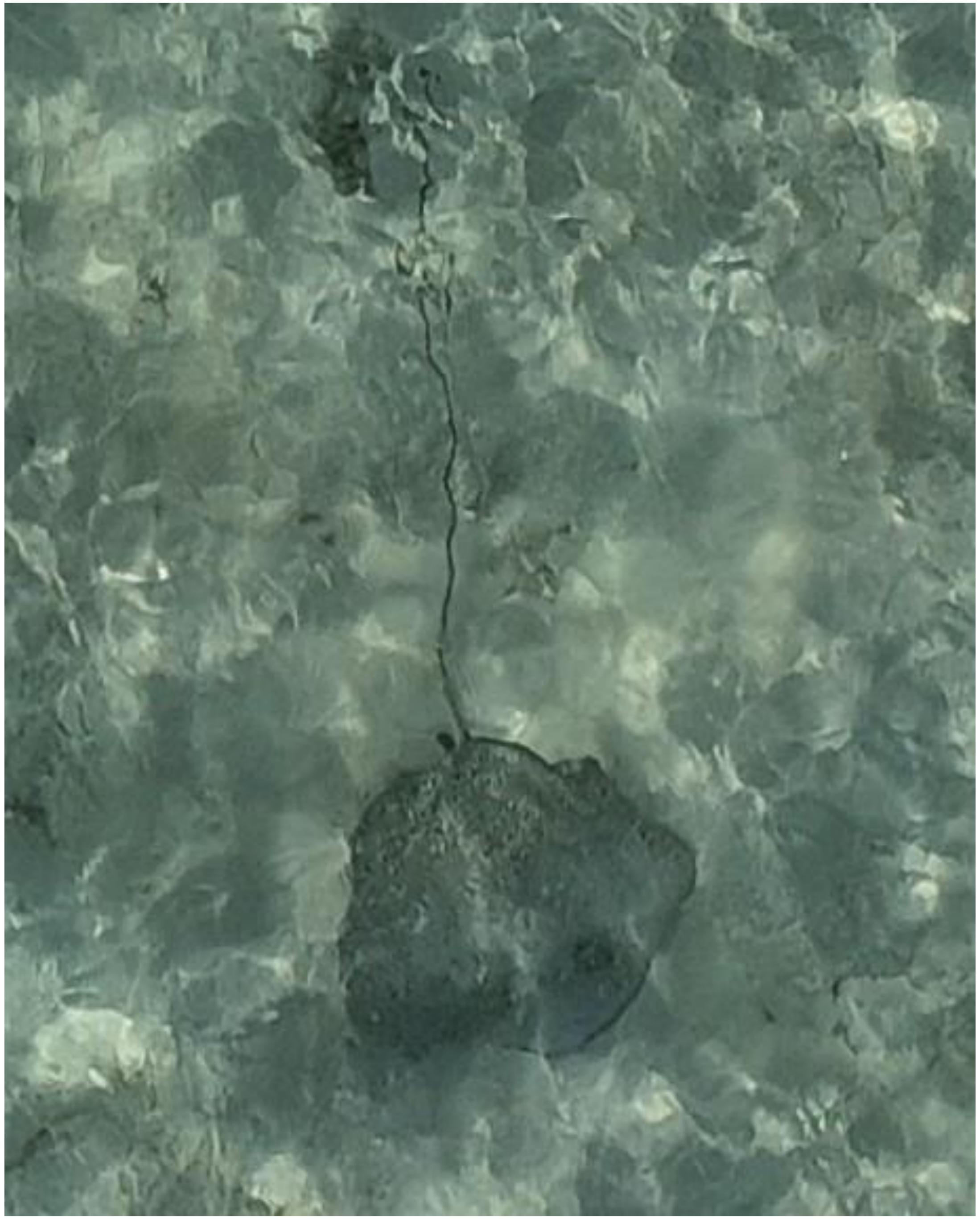

- Himantura uarnak

DJI_20240502072126_0411

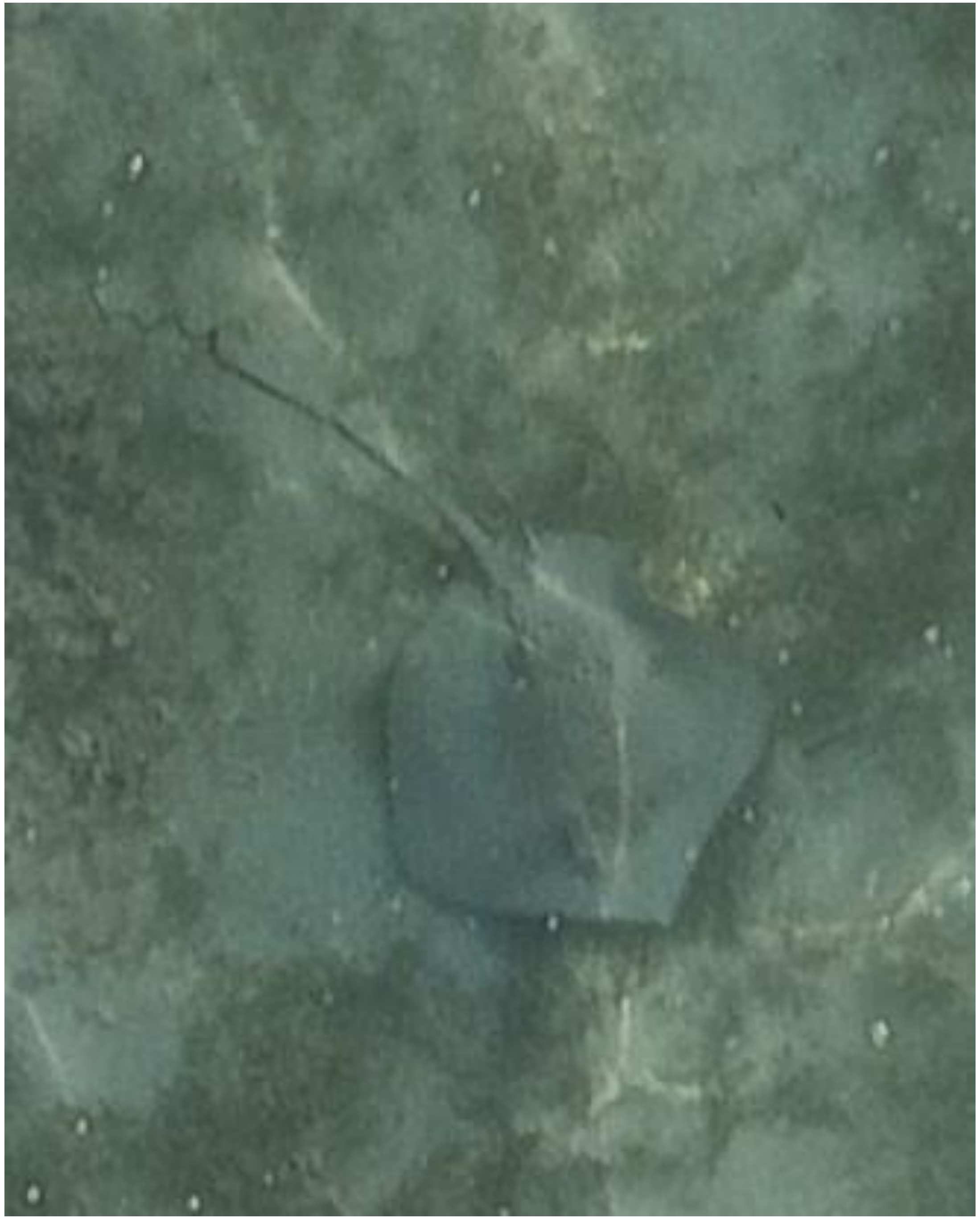

DJI_20240502072551_0630

- Whipray
- Also observed on 06-06, 06-24, 07-29, and 09-20 as identified by the coiled tail.
- Best guess: *H. uarnak*

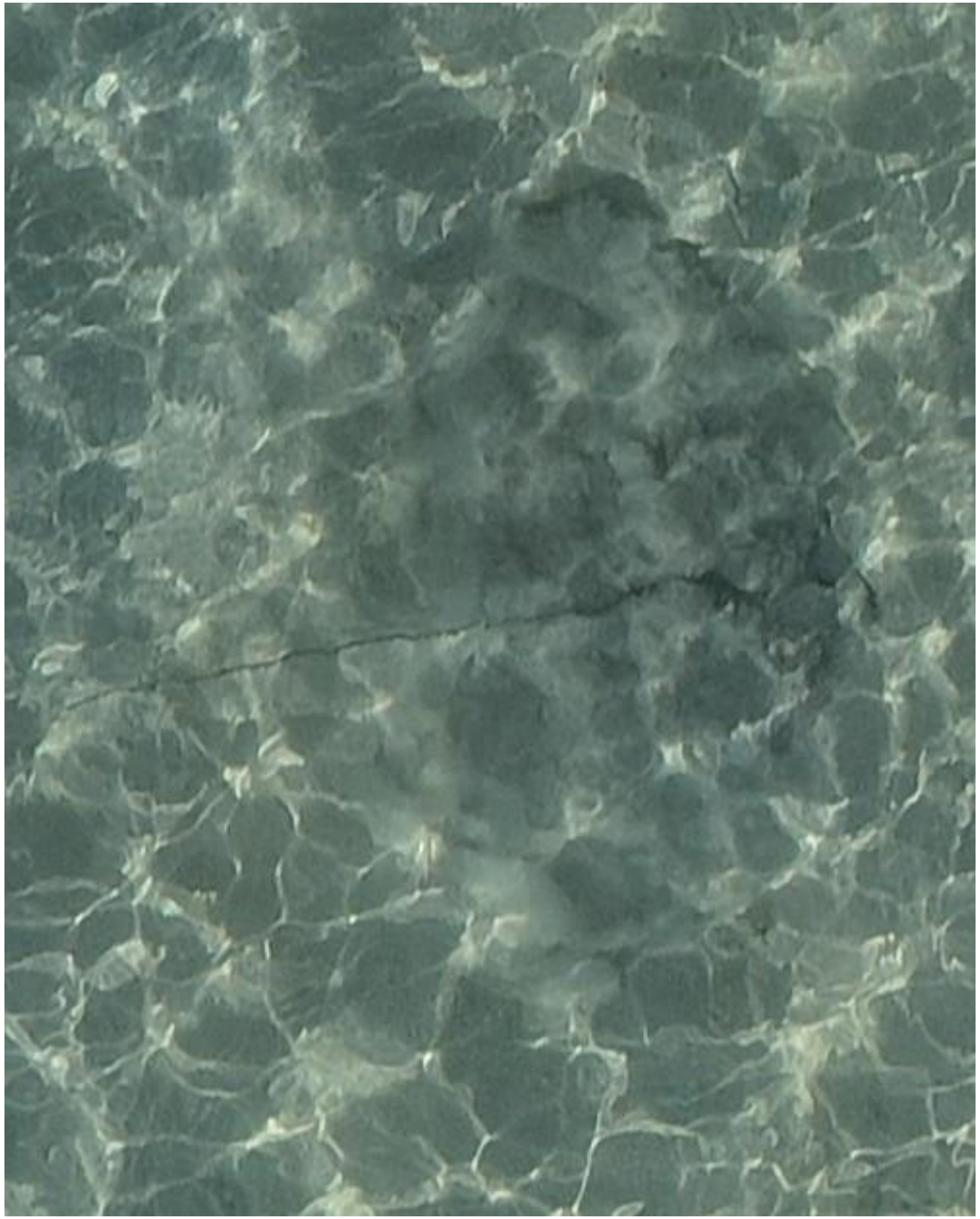
DJI_20240502072825_0772

- Whipray

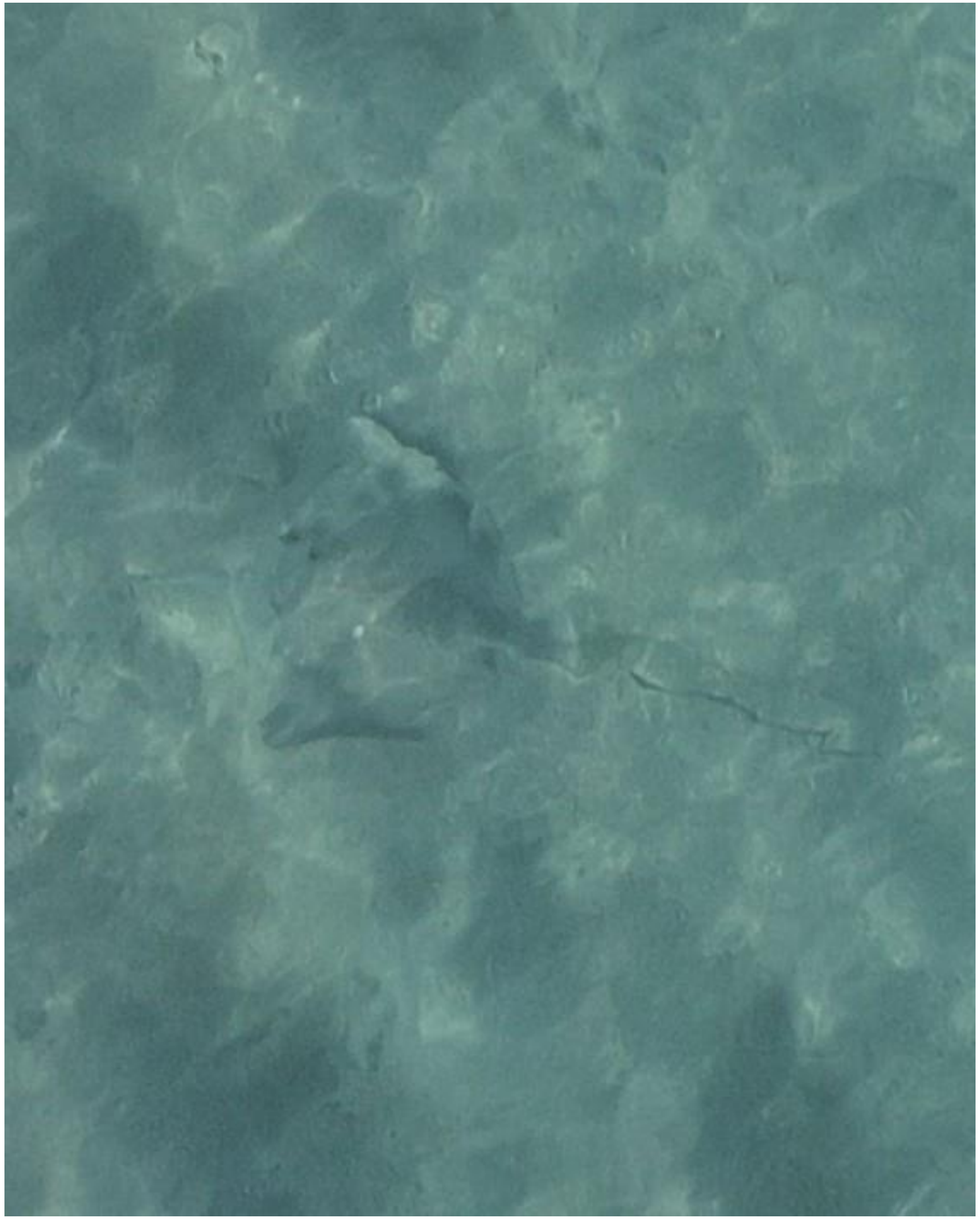

DJI_20240502073227_0990

- Whipray
- Accompanied by needlefish

2024-05-09

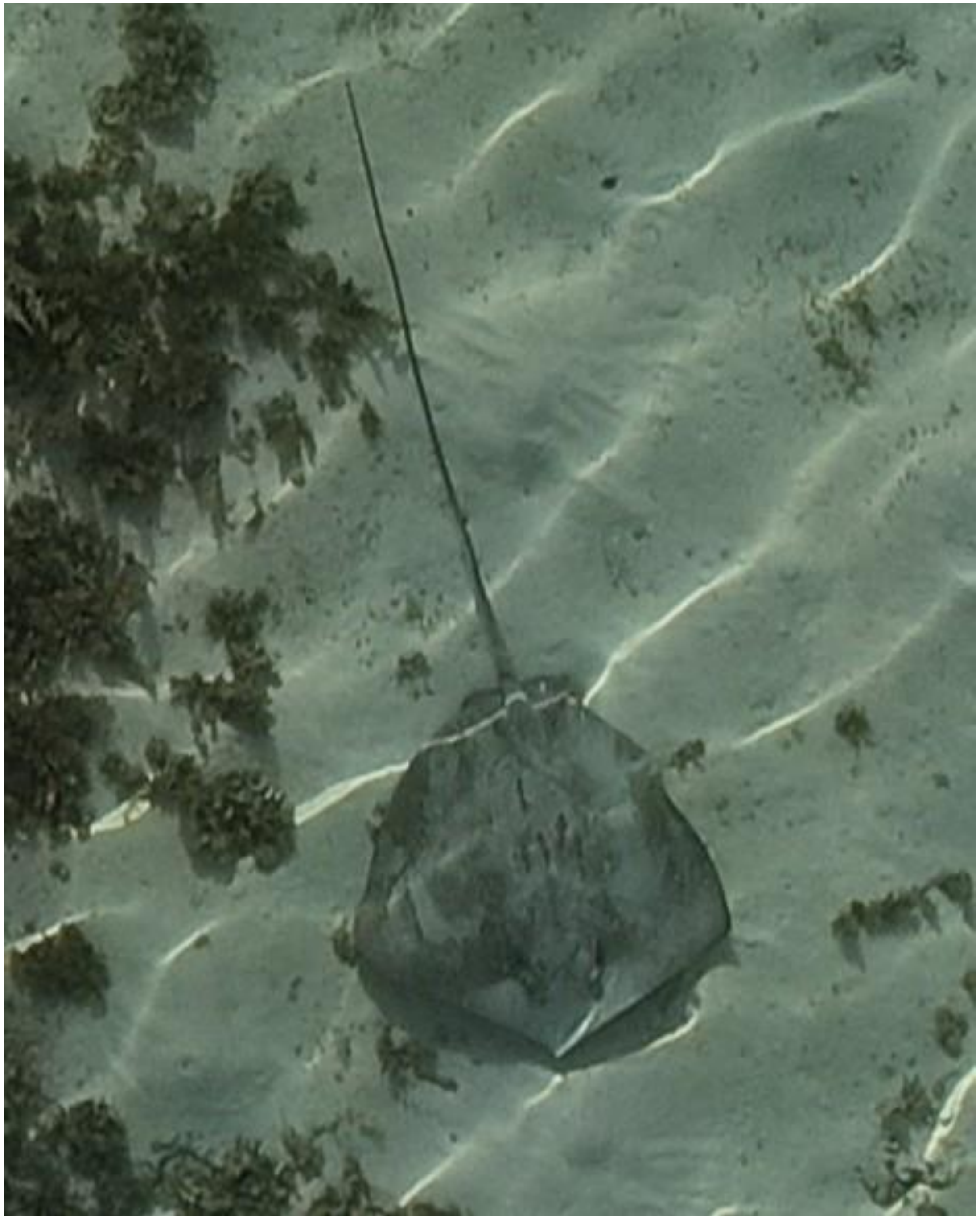

DJI_20240509073349_0036

- Whipray
- Best guess: cf. *P. fai*

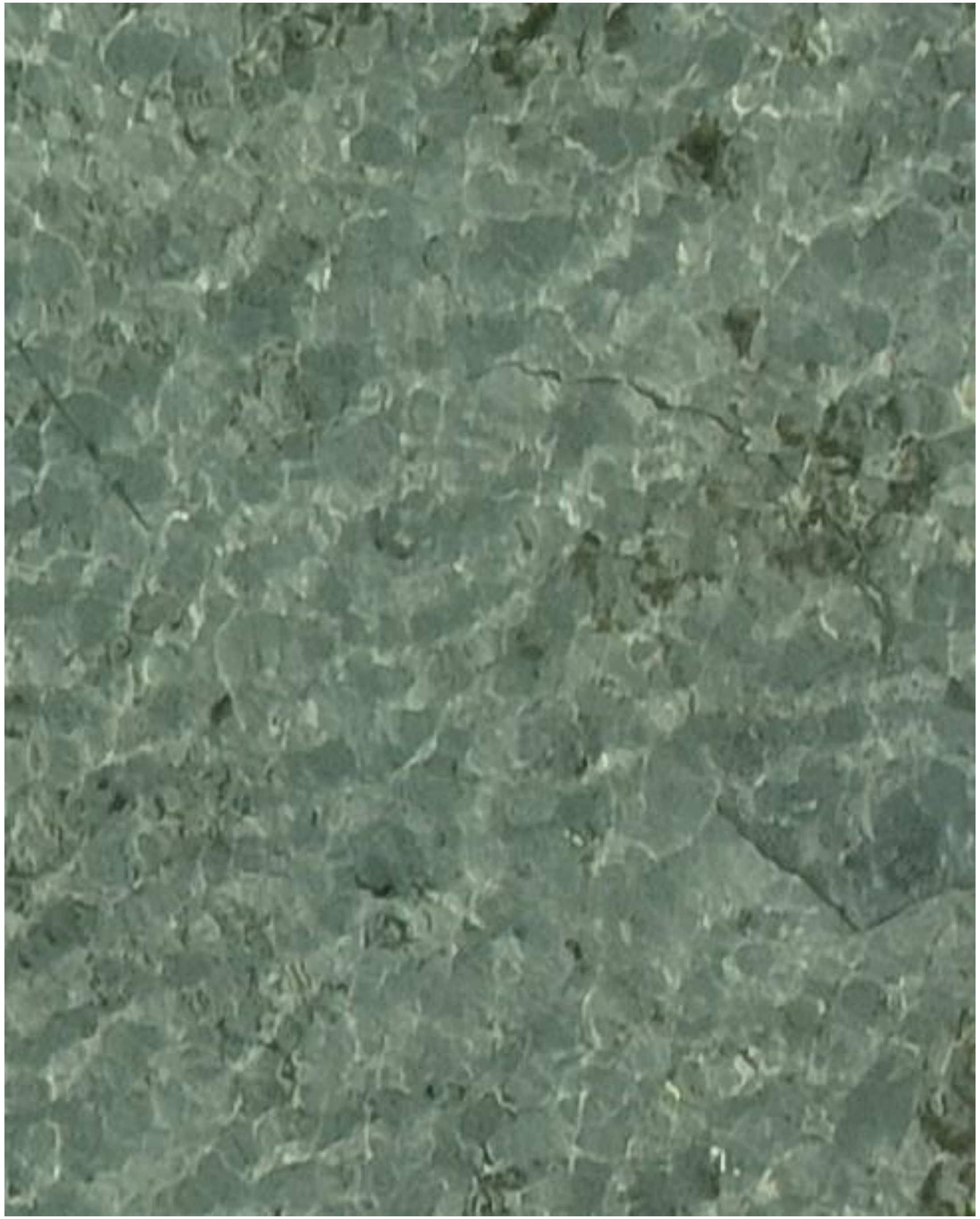

DJI_20240509073403_0050

- Whipray
- Best guess: *H. uarnak*
- Accompanied by needlefish

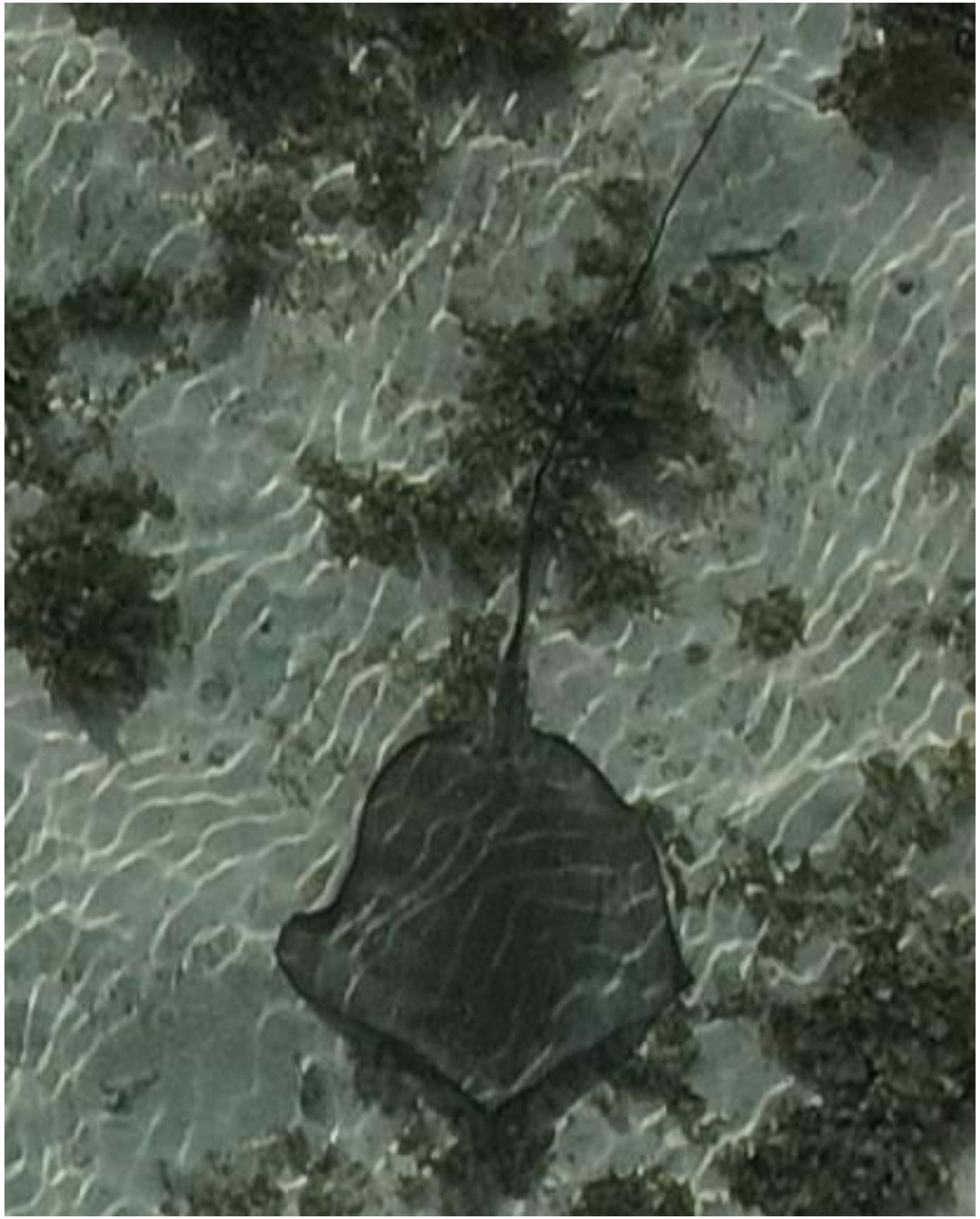

- Whipray
- Best guess: *H. uarnak*

DJI_20240509074034_0394

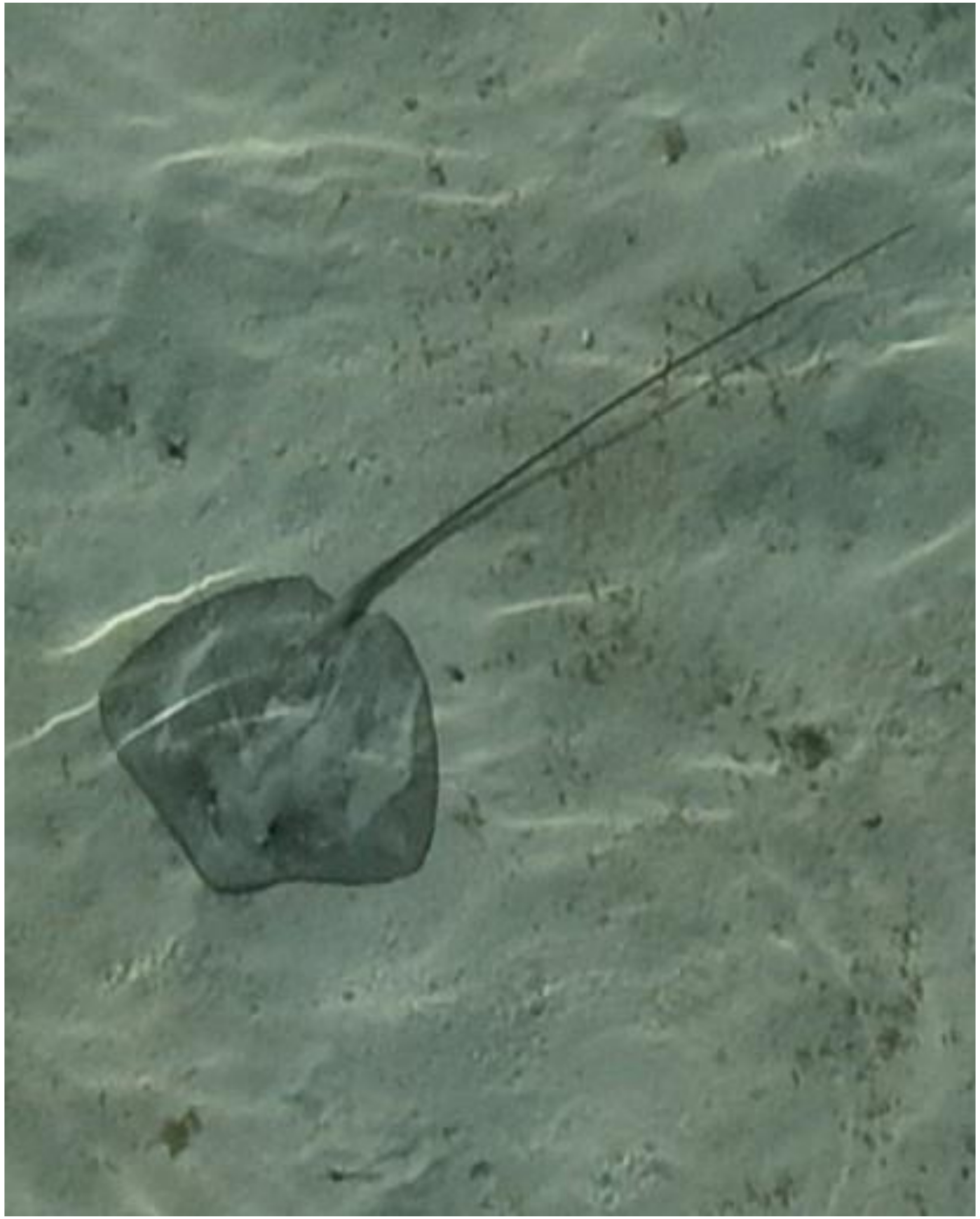

- Whipray
- Best guess: cf. *P. fai*

DJI_20240509074043_0403

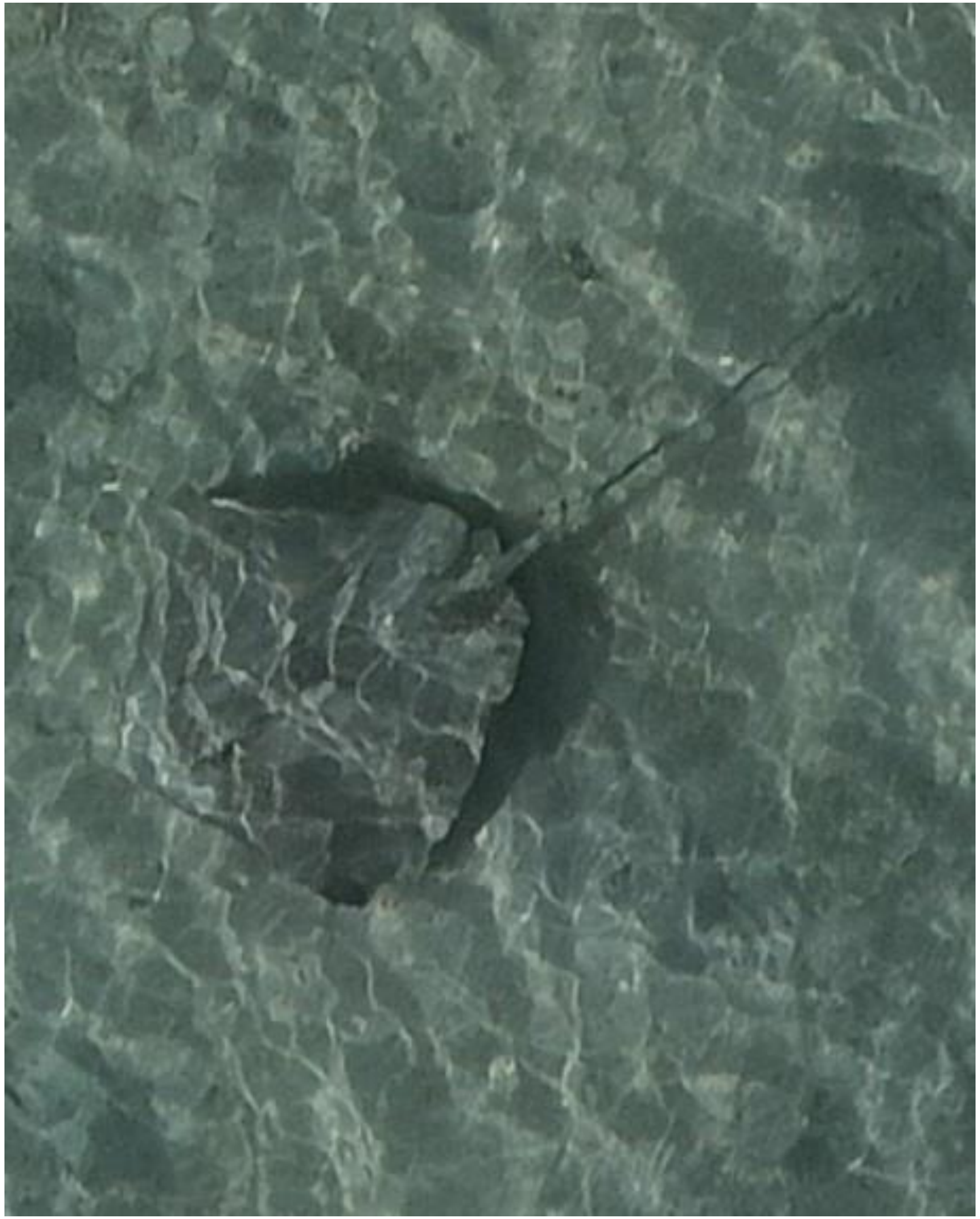

- Whipray

DJI_20240509074751_0773

2024-05-10

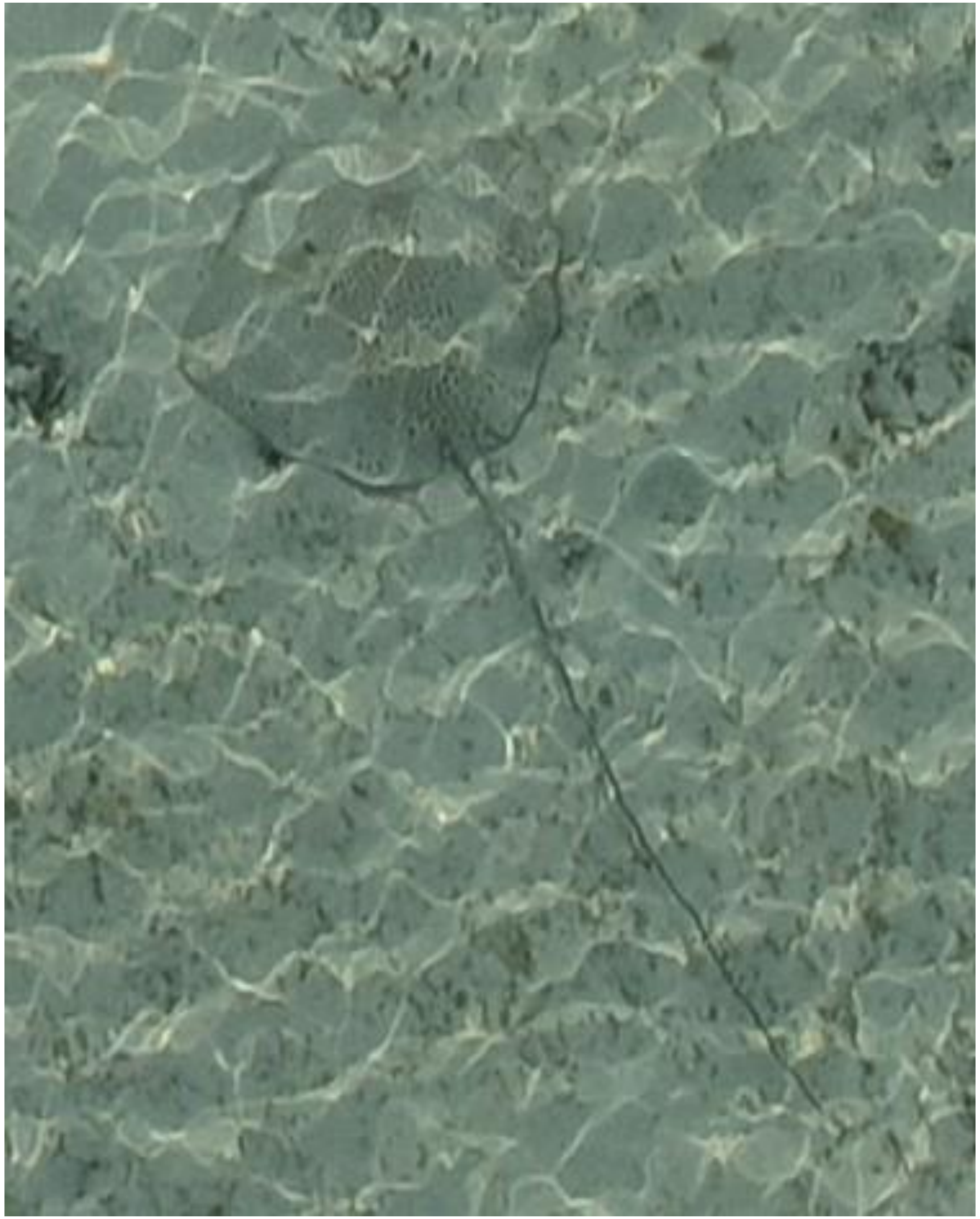
DJI_20240510071346_0054

- *Himantura uarnak*

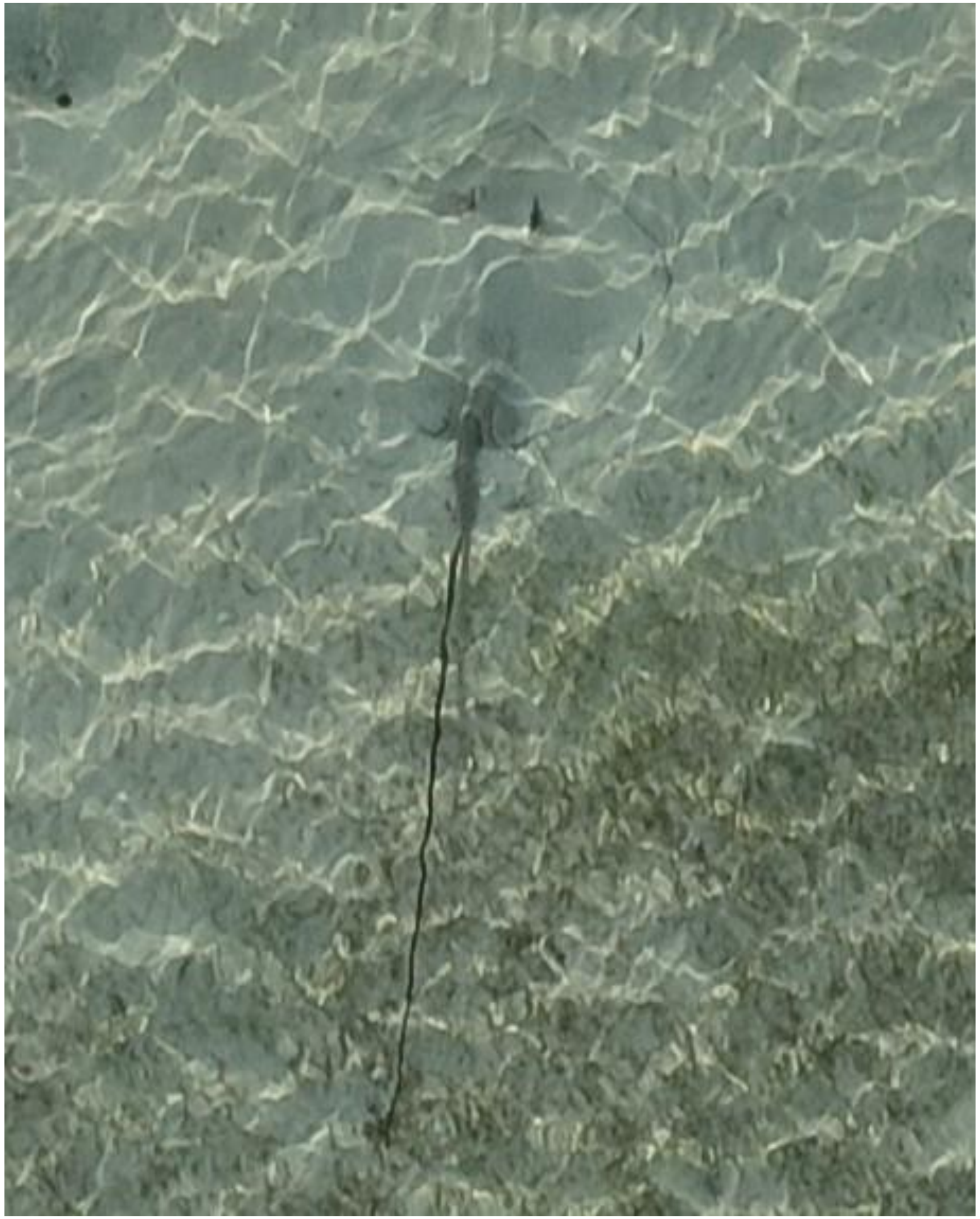

- Whipray

DJI_20240510071349_0057

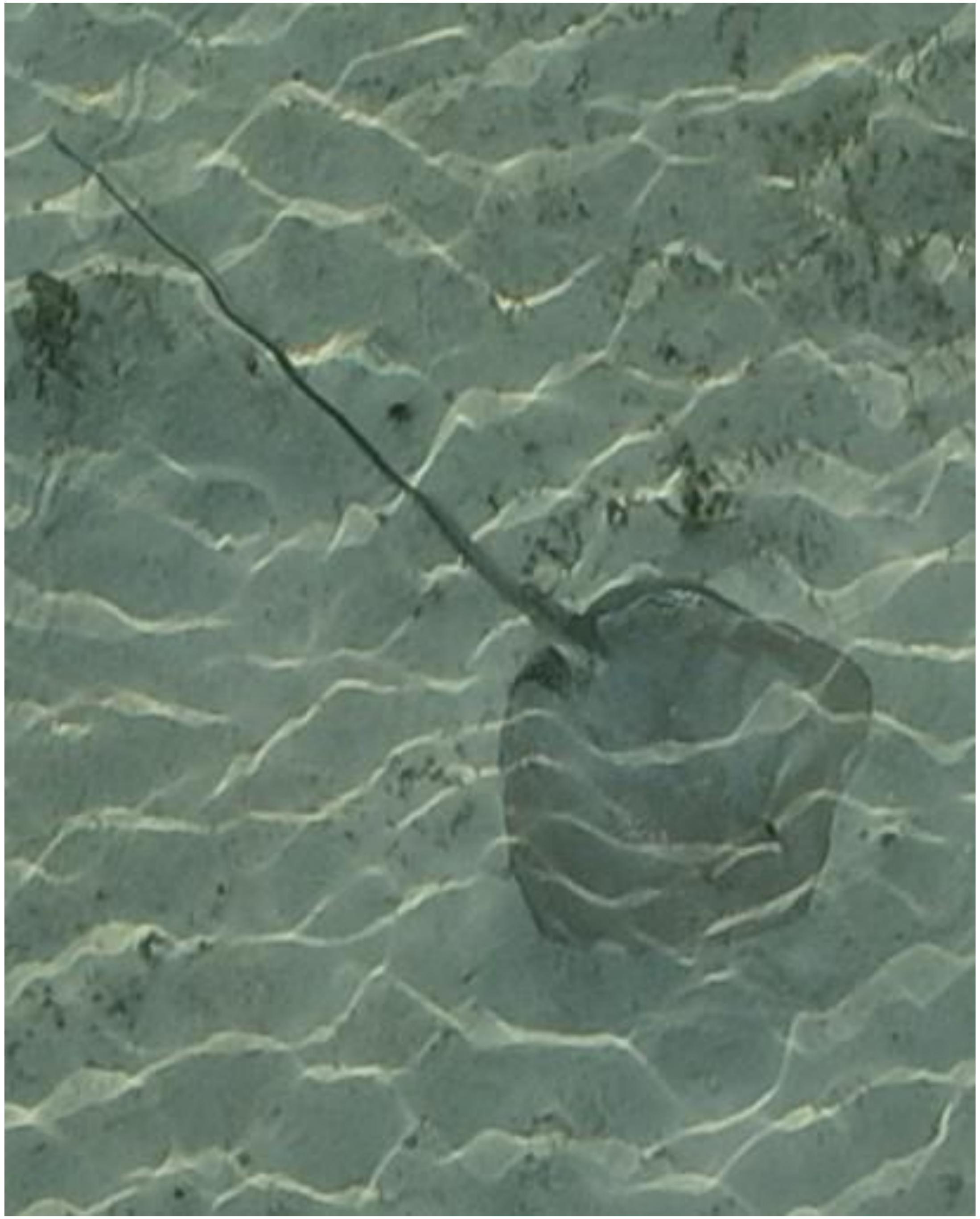

DJI_20240510071438_0095

- Whipray
- Best guess: *H. uarnak*

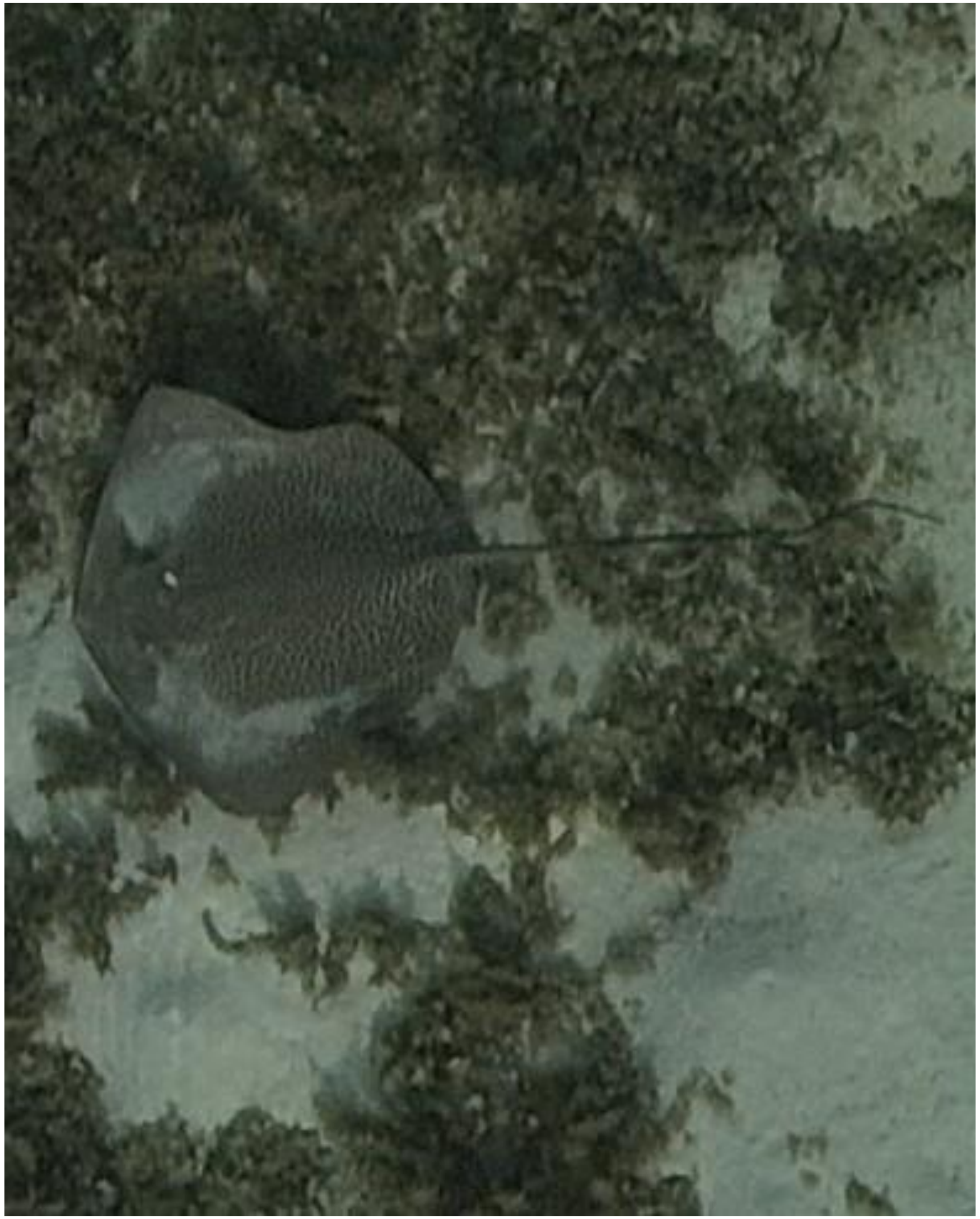
DJI_20240510071449_0106

- *Himantura uarnak*

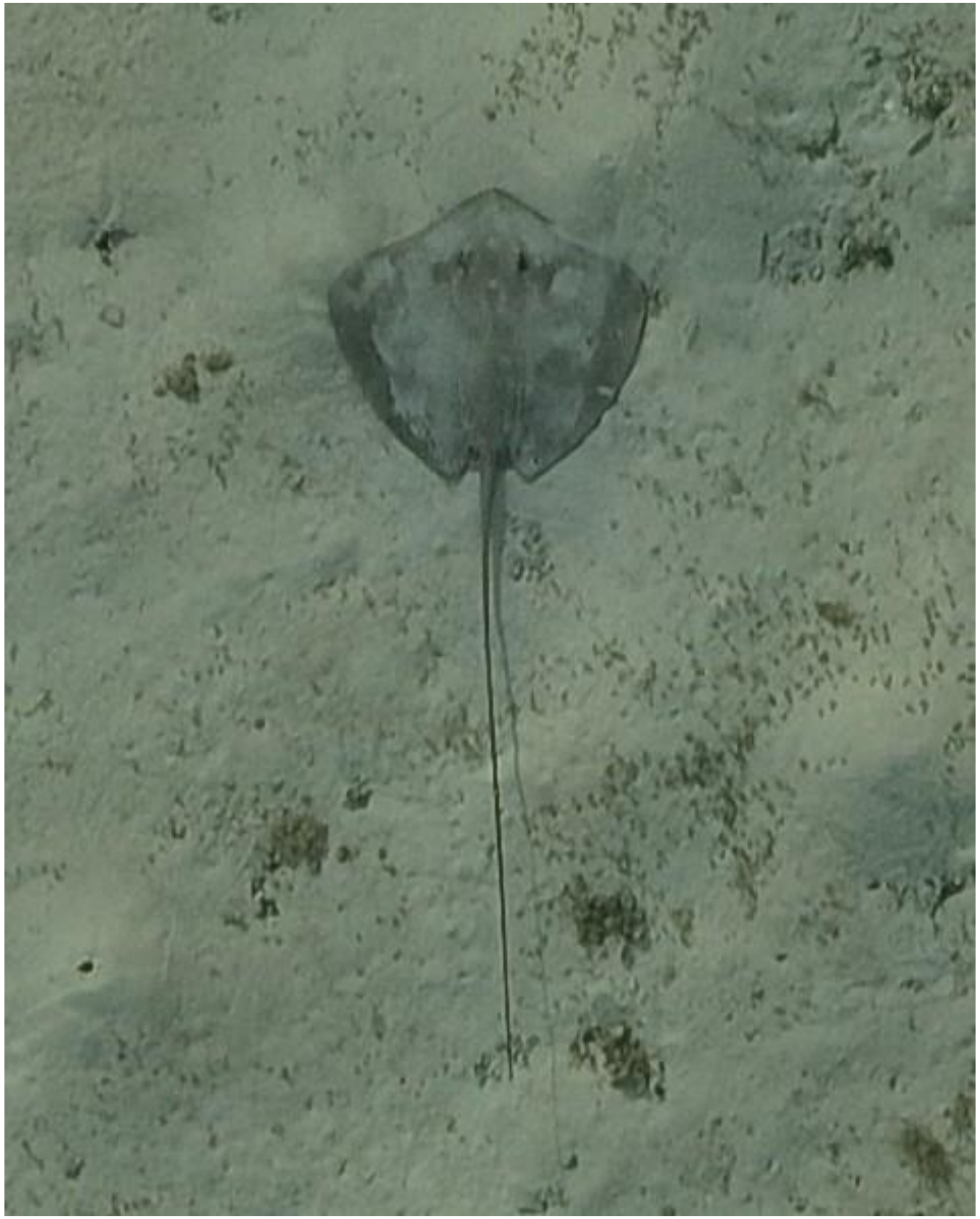

DJI_20240510071635_0200

- Whipray
- Best guess: *H. uarnak*

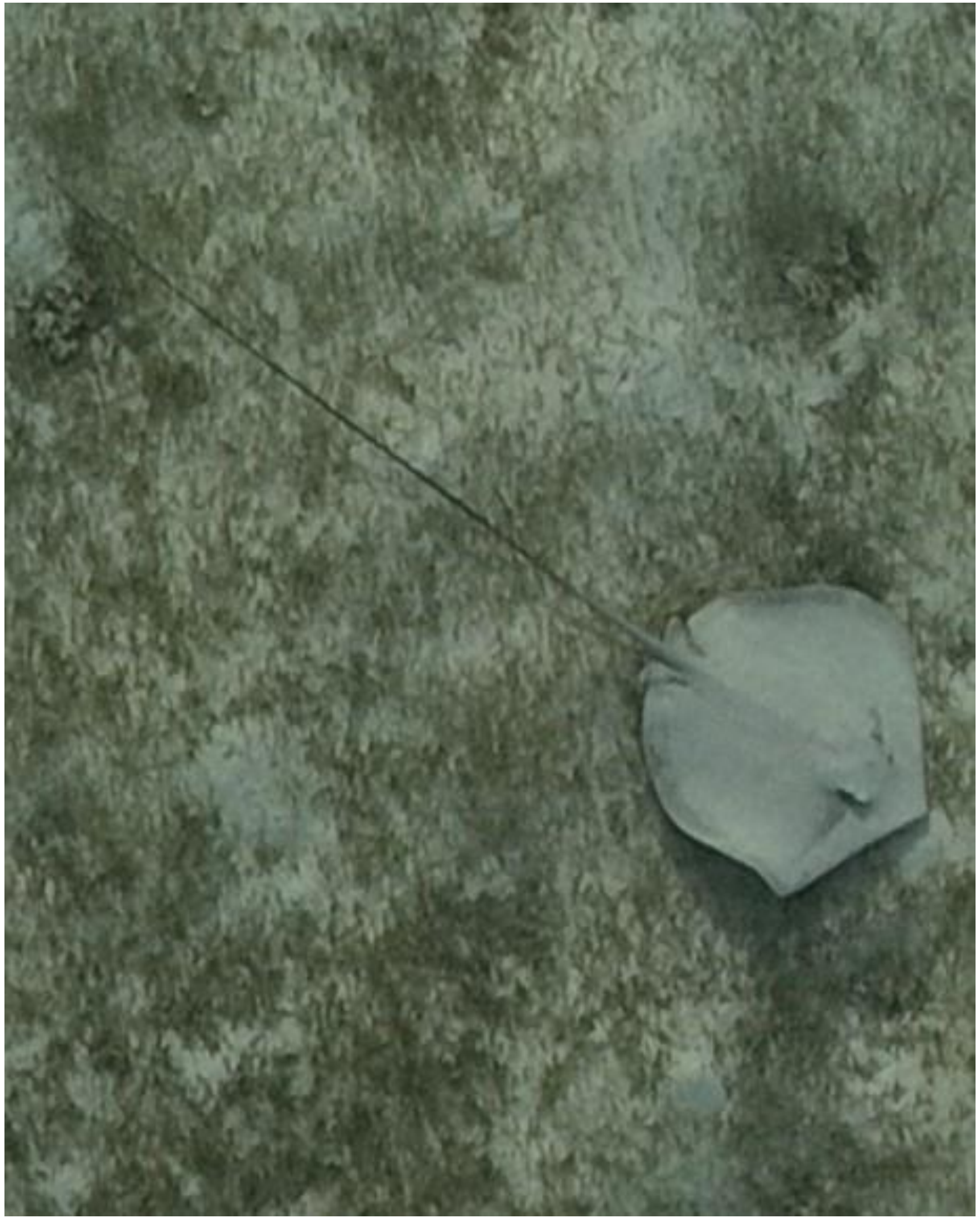

DJI_20240510071701_0226

- Whipray

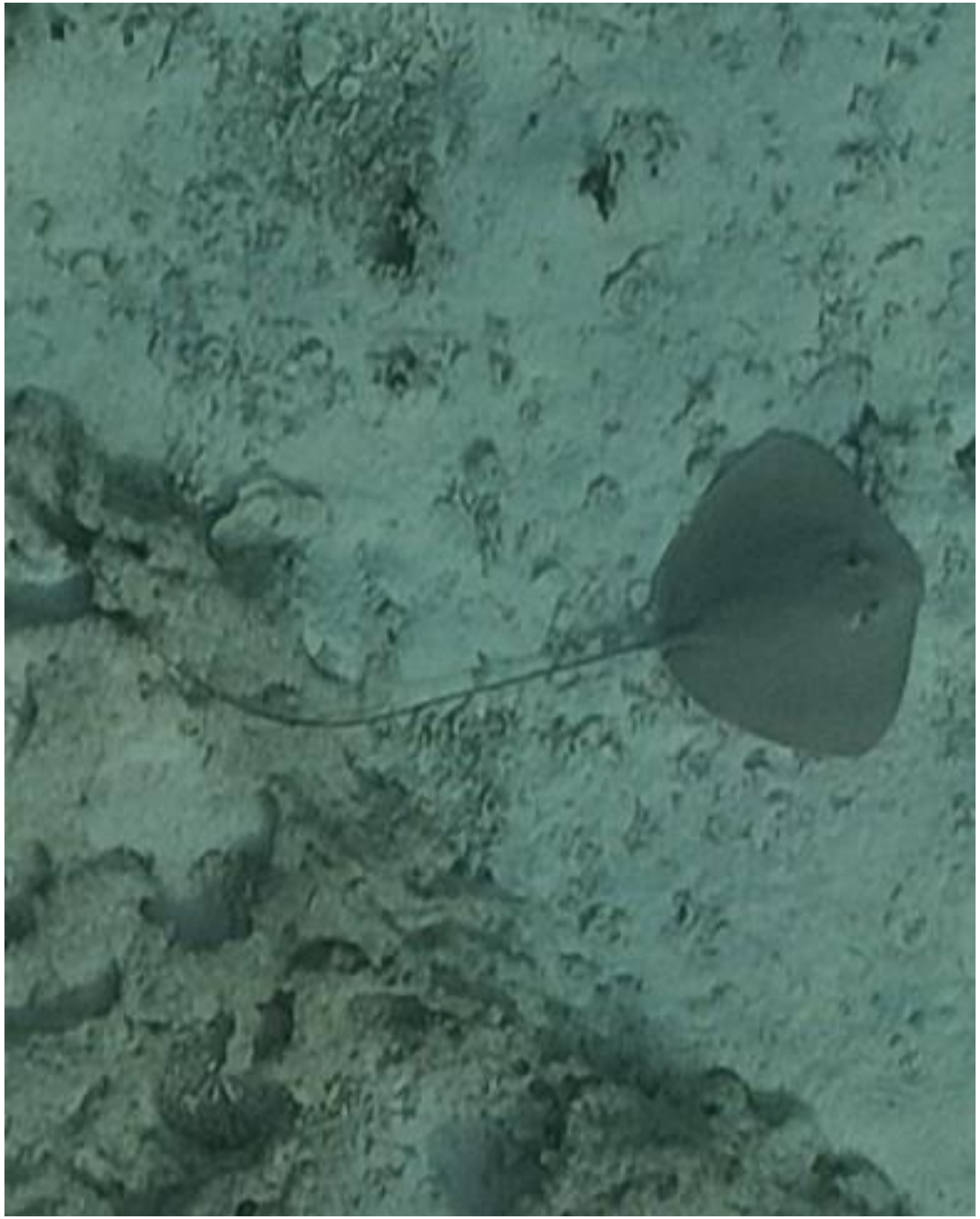

- cf. *Pateobatis fai*

DJI_20240510071749_0262

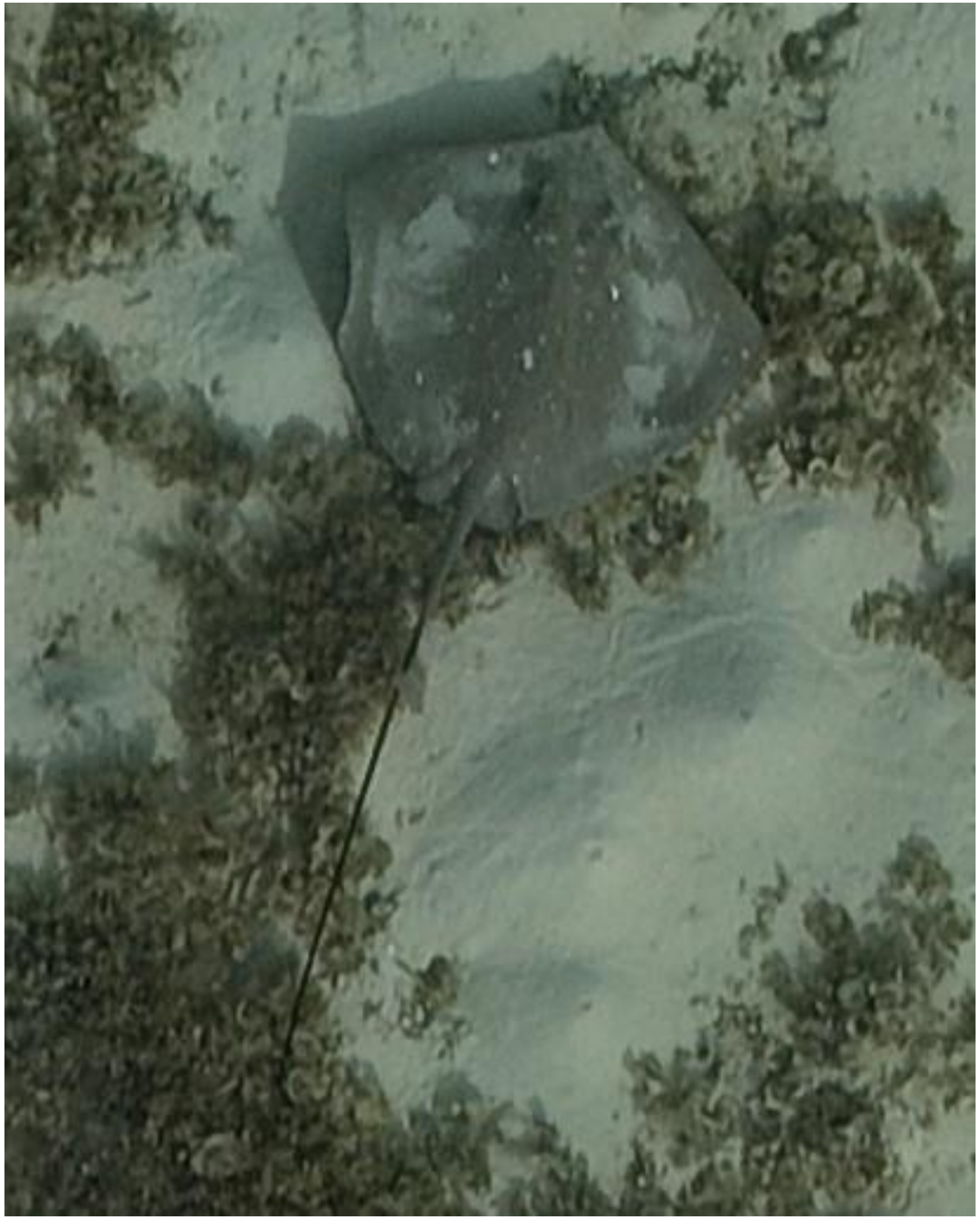

DJI_20240510071807_0280

- *Himantura uarnak*
- Identified as the individual that was also tracked on 05-22 and 06-11. Also observed on 05-17 and 06-03.

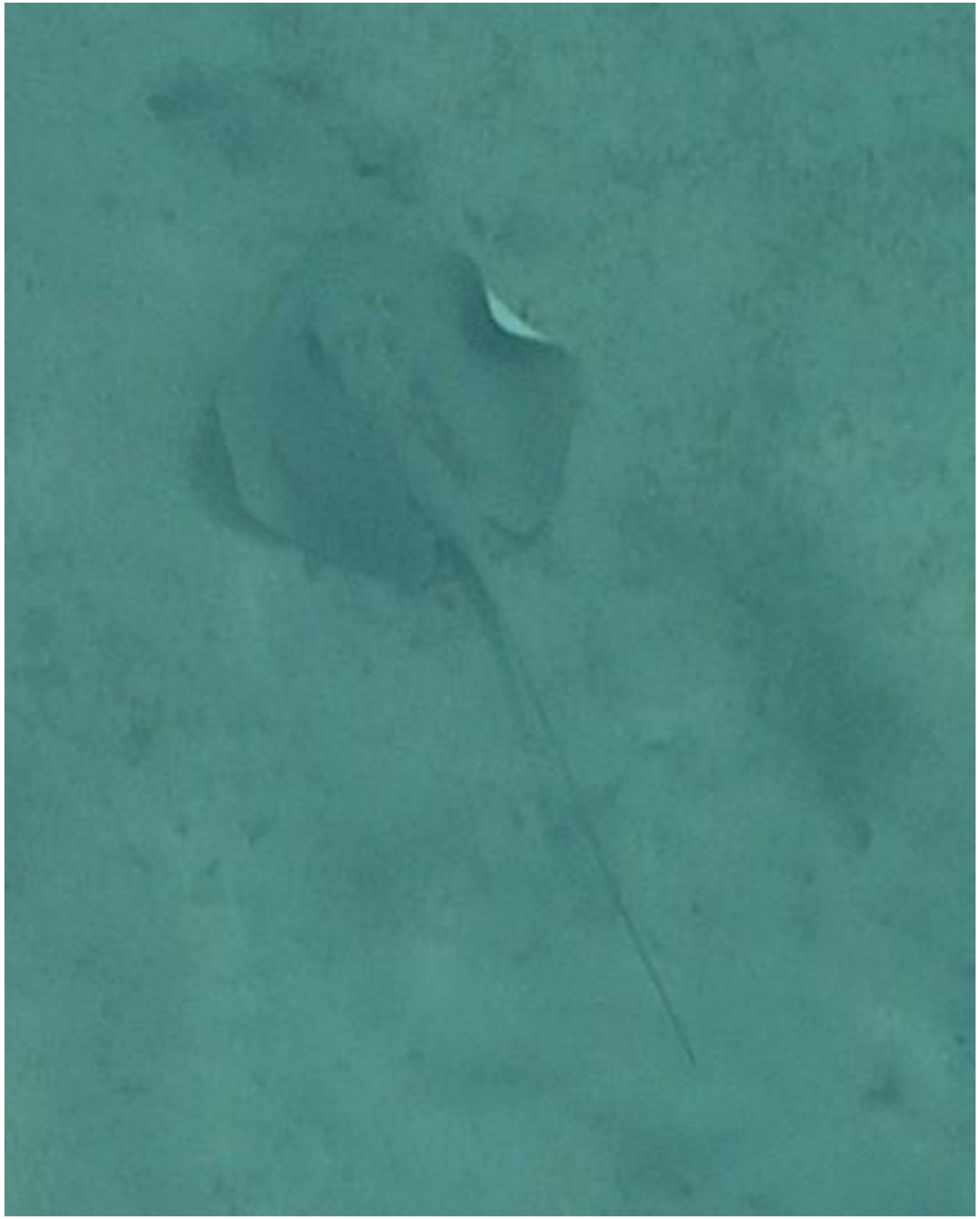
DJI_20240510072955_0905

- Whipray

2024-05-12

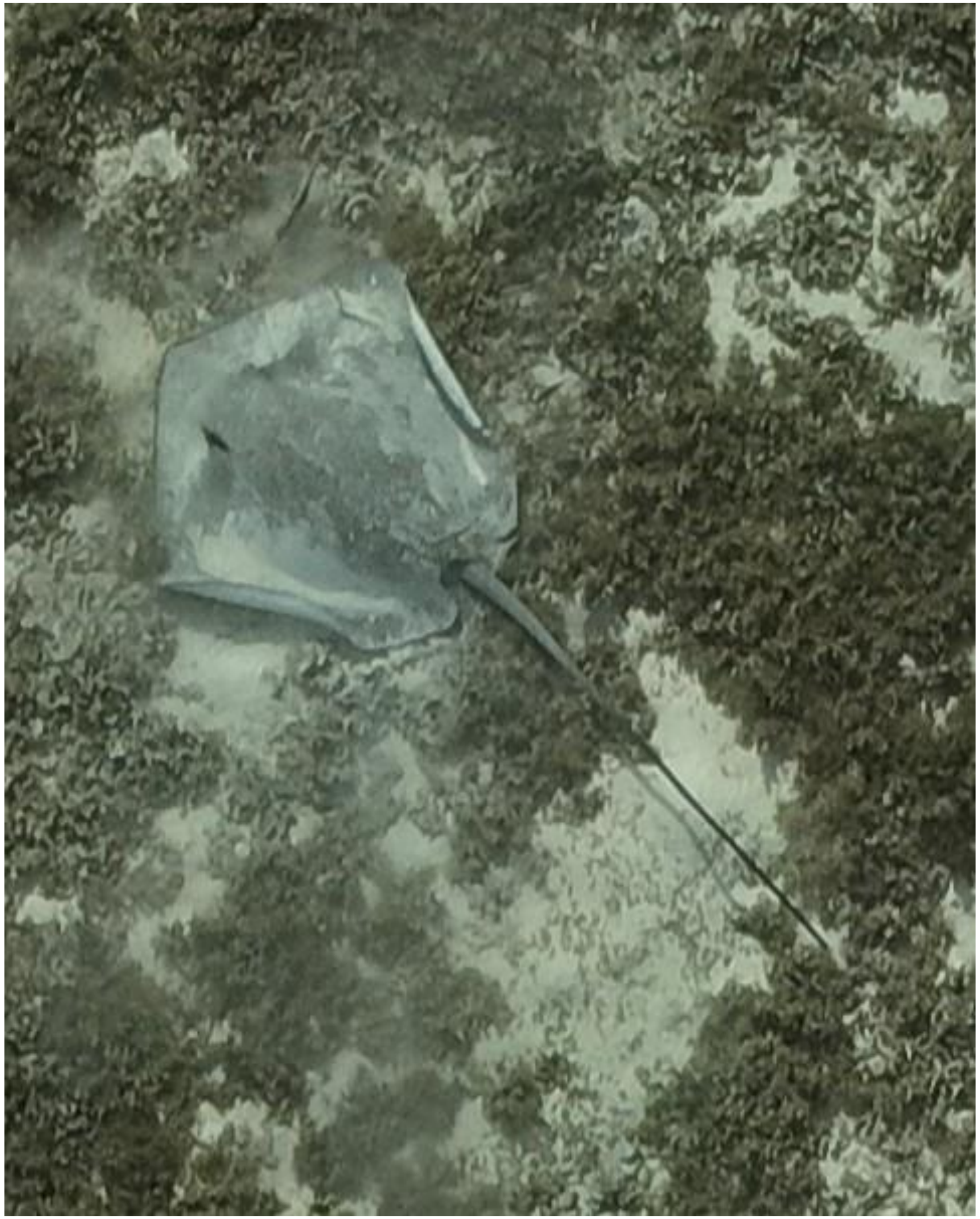

- Whipray

DJI_20240512090654_0021

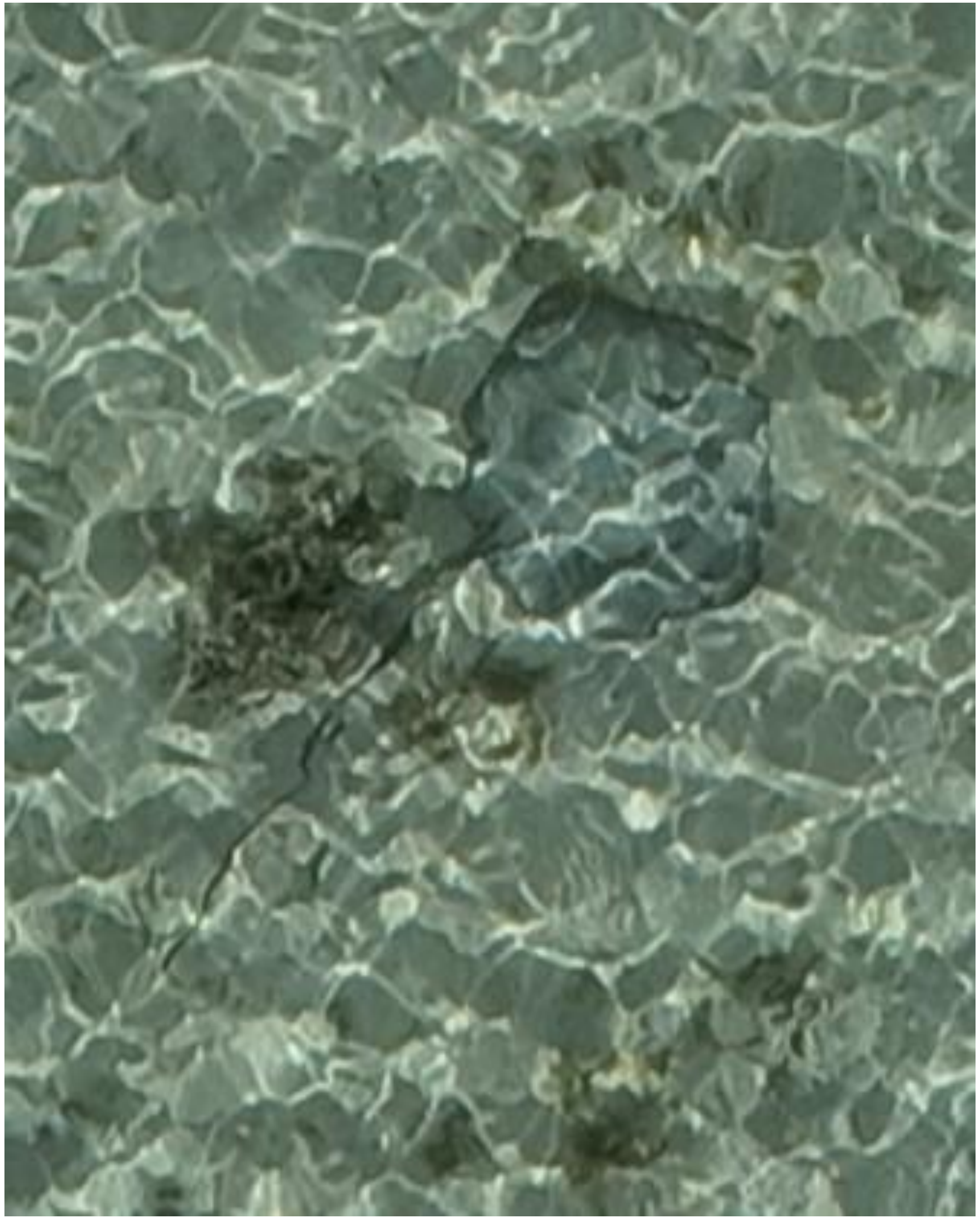

DJI_20240512090849_0125

- Whipray

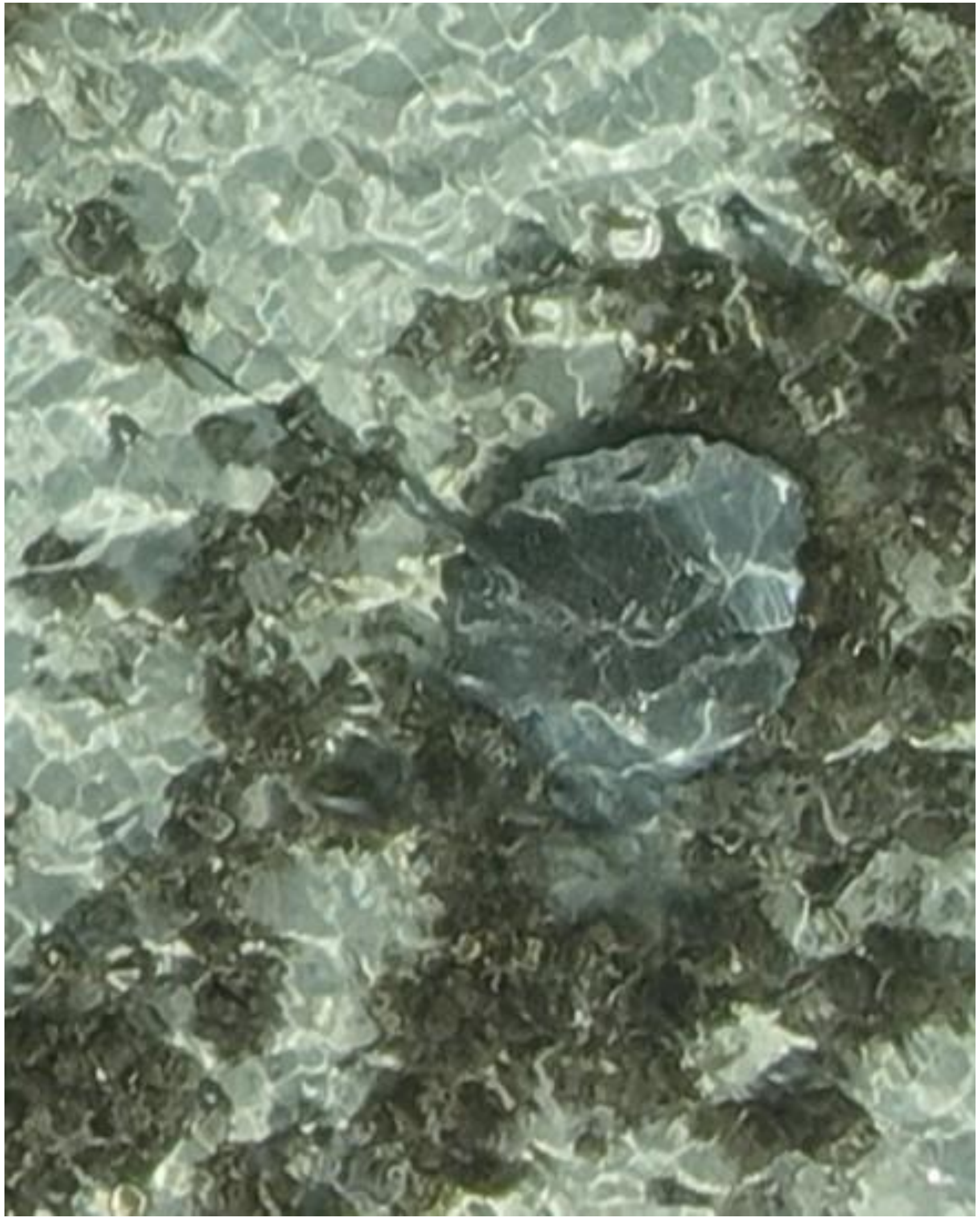

DJI_20240512090859_0135

- Whipray

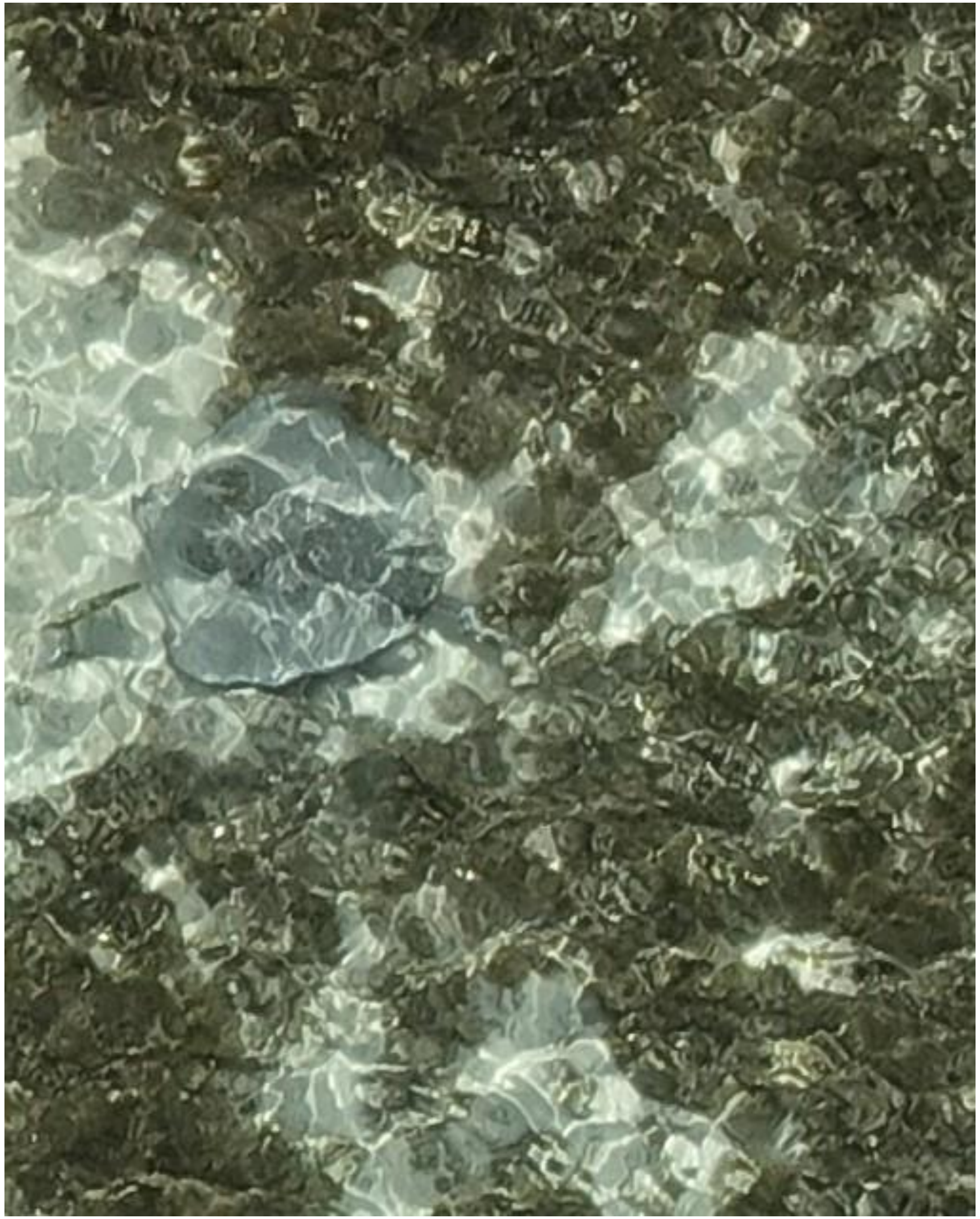

DJI_20240512090954_0178

- Whipray

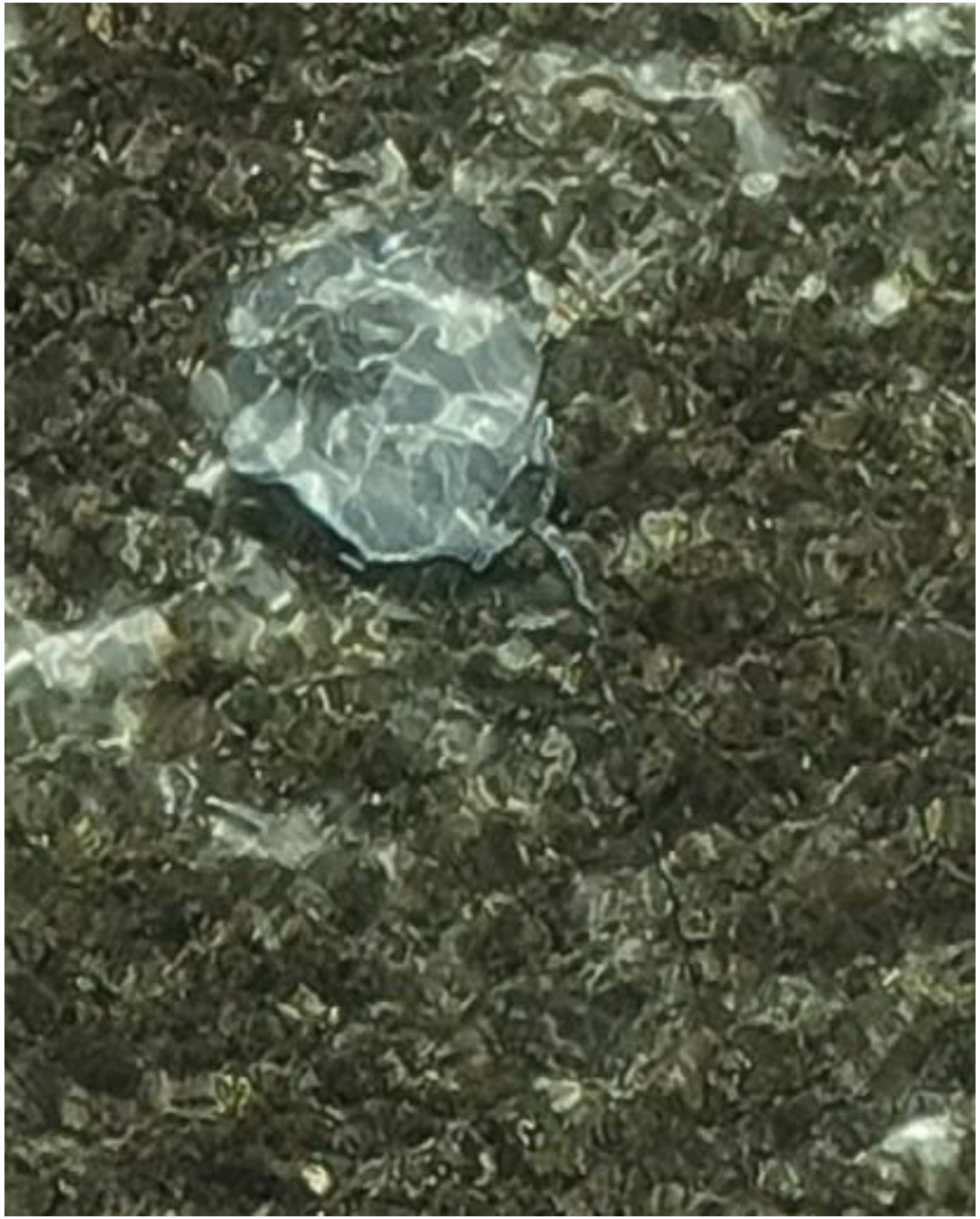

- Whipray

DJI_20240512090954_0178

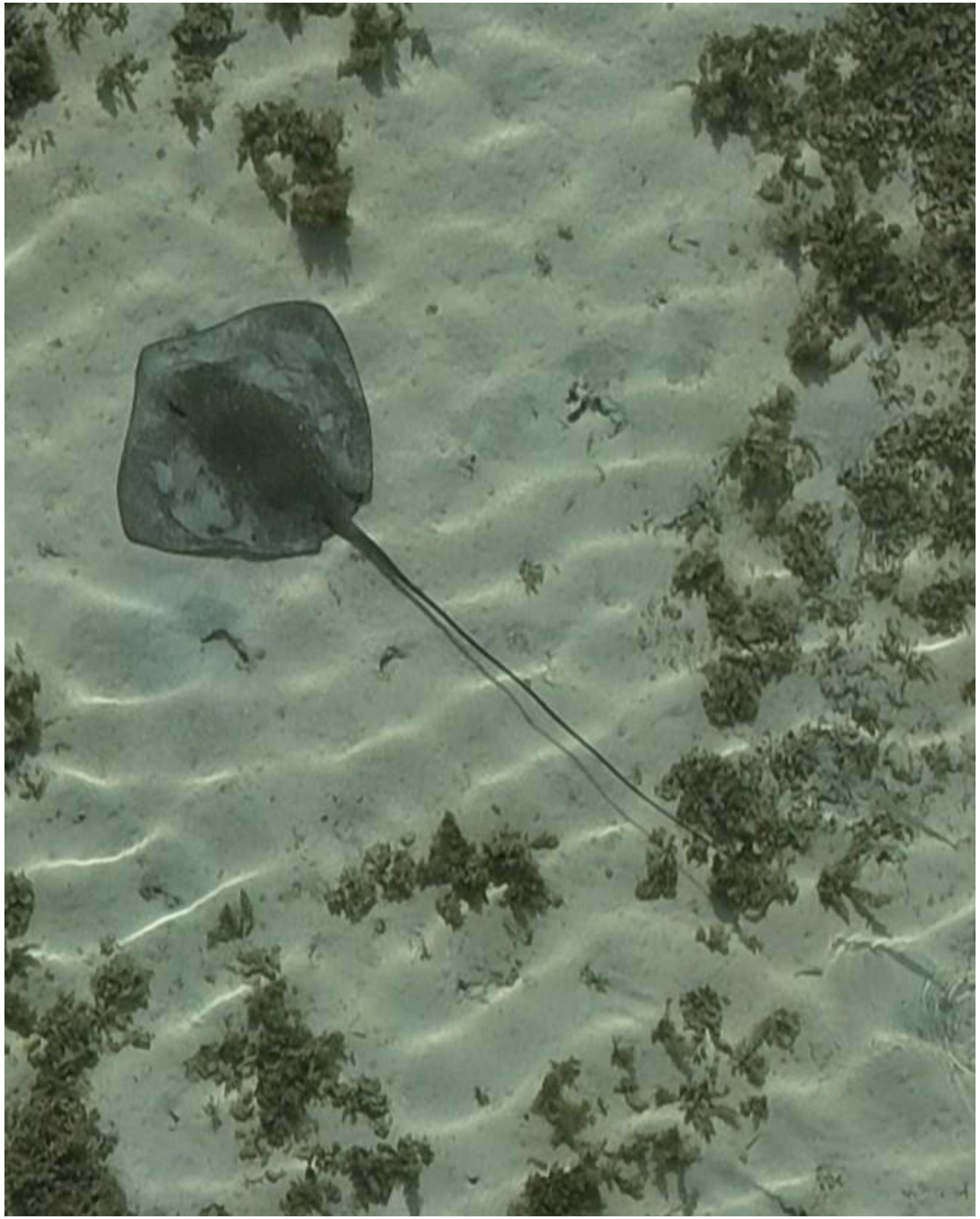

- cf. *Pateobatis fai*
- Accompanied by needlefish

DJI_20240512091023_0207

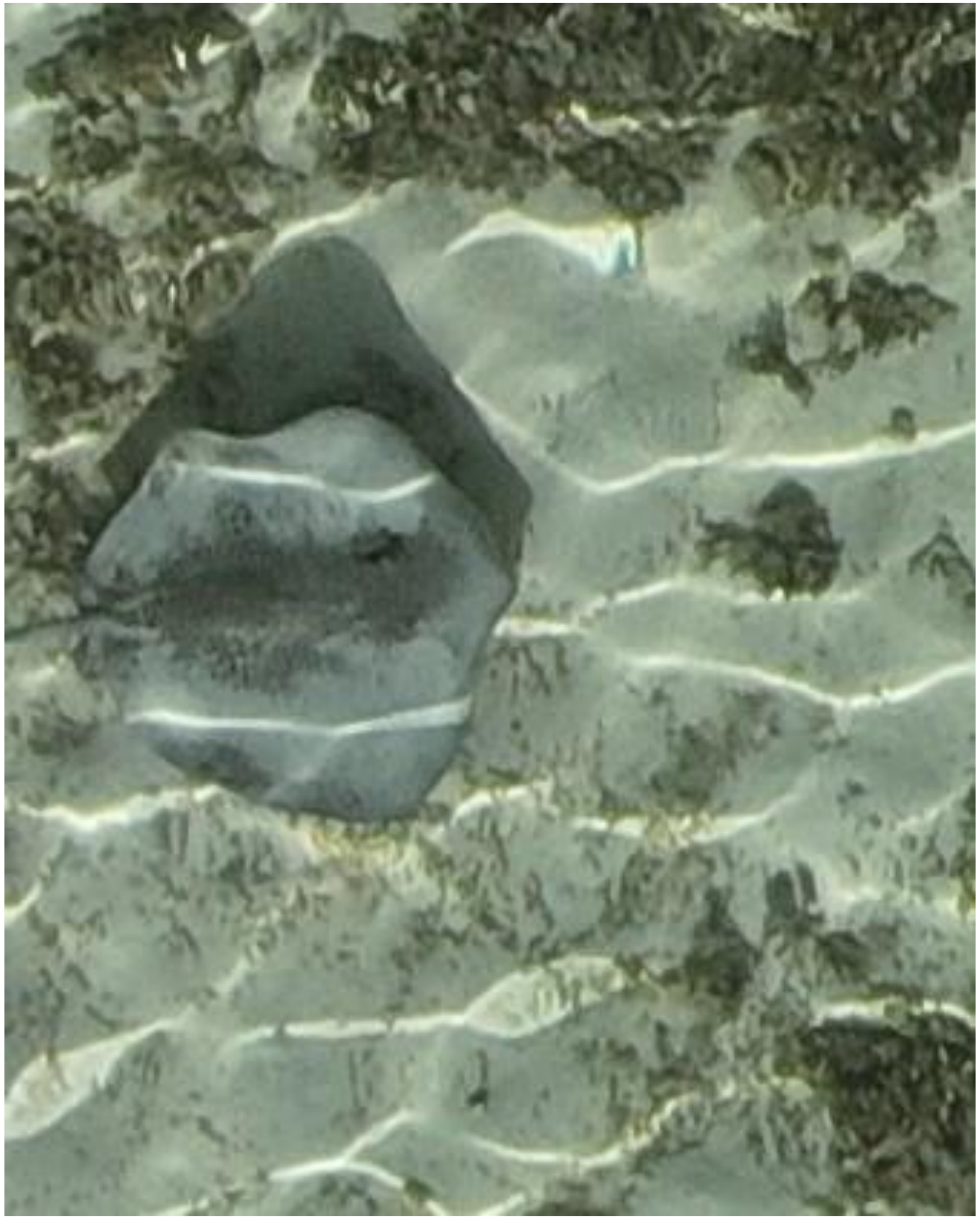

DJI_20240512091232_0324

- *Himantura uarnak*

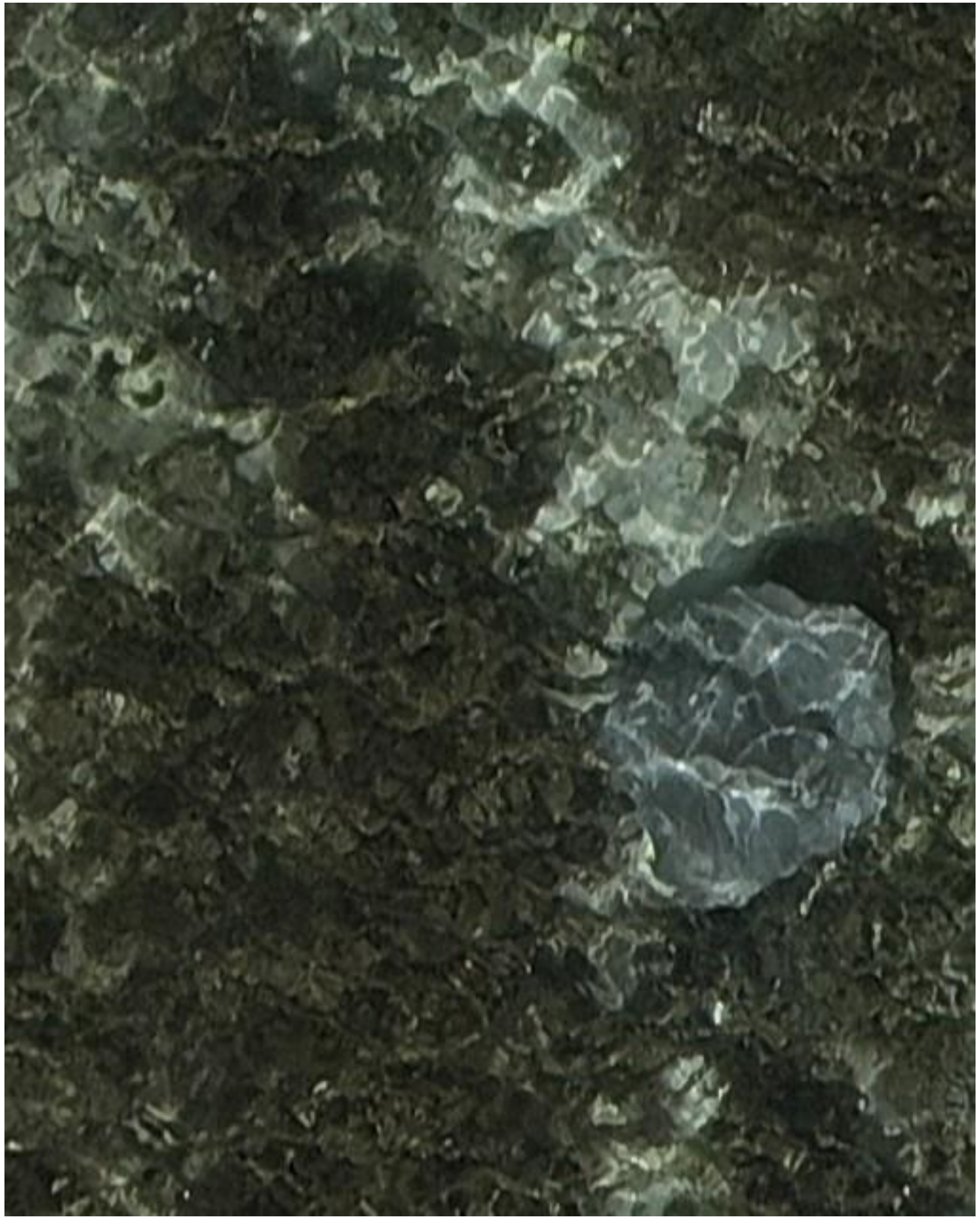

- Whipray

DJI_20240512091238_0330

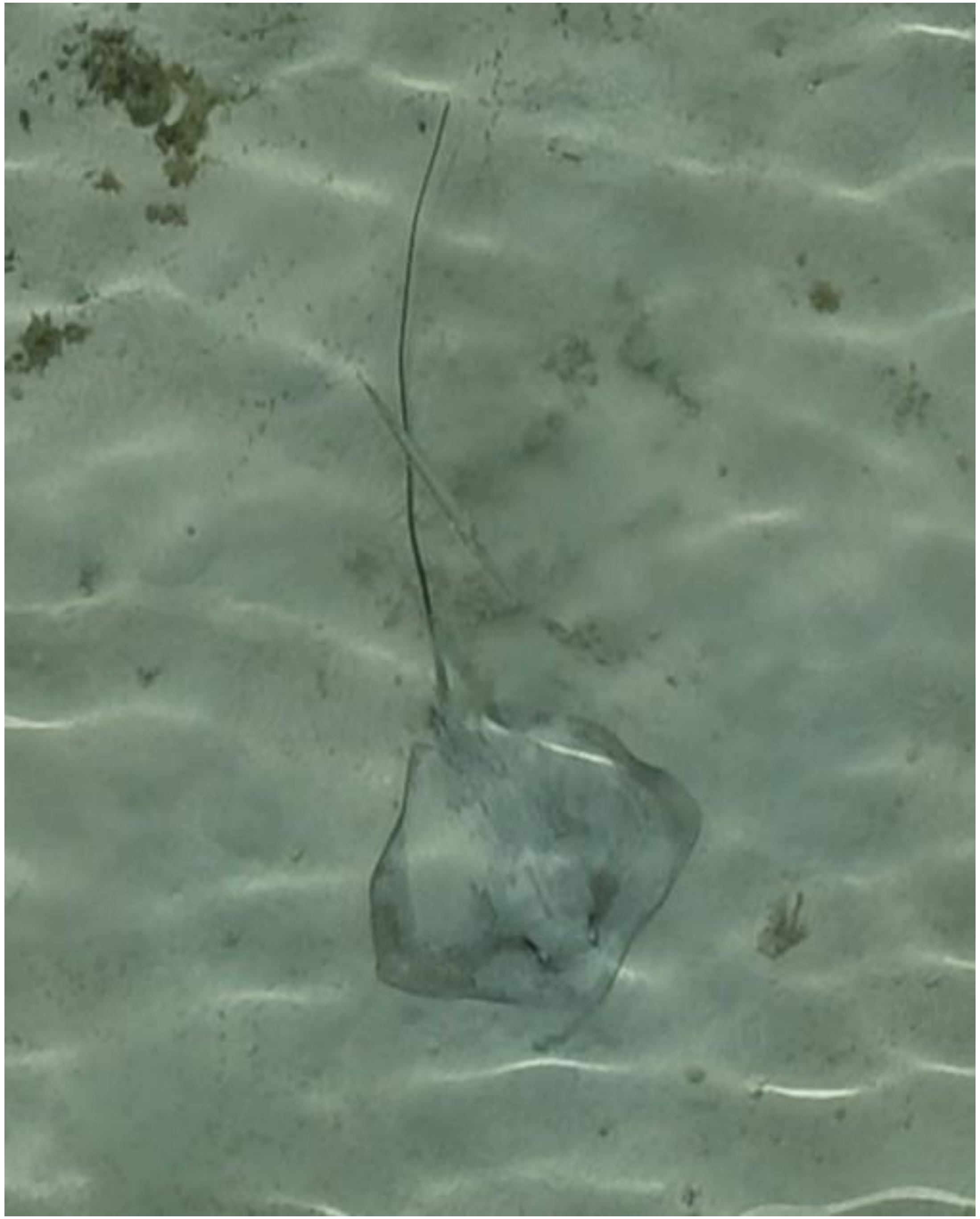

DJI_20240512091436_0436

- Whipray
- Accompanied by needlefish

2024-05-17

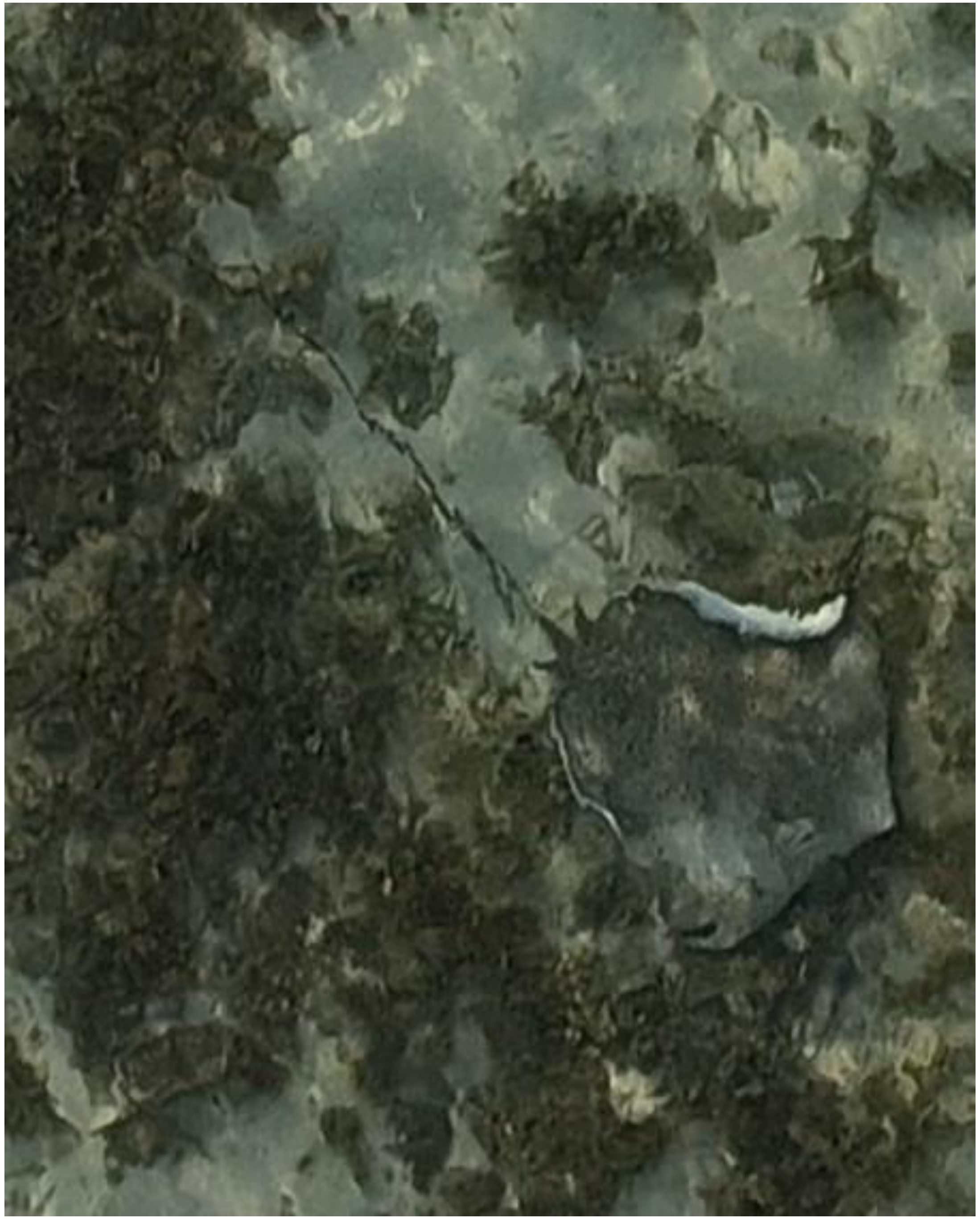

DJI_20240517071346_0046

- *Himantura uarnak*

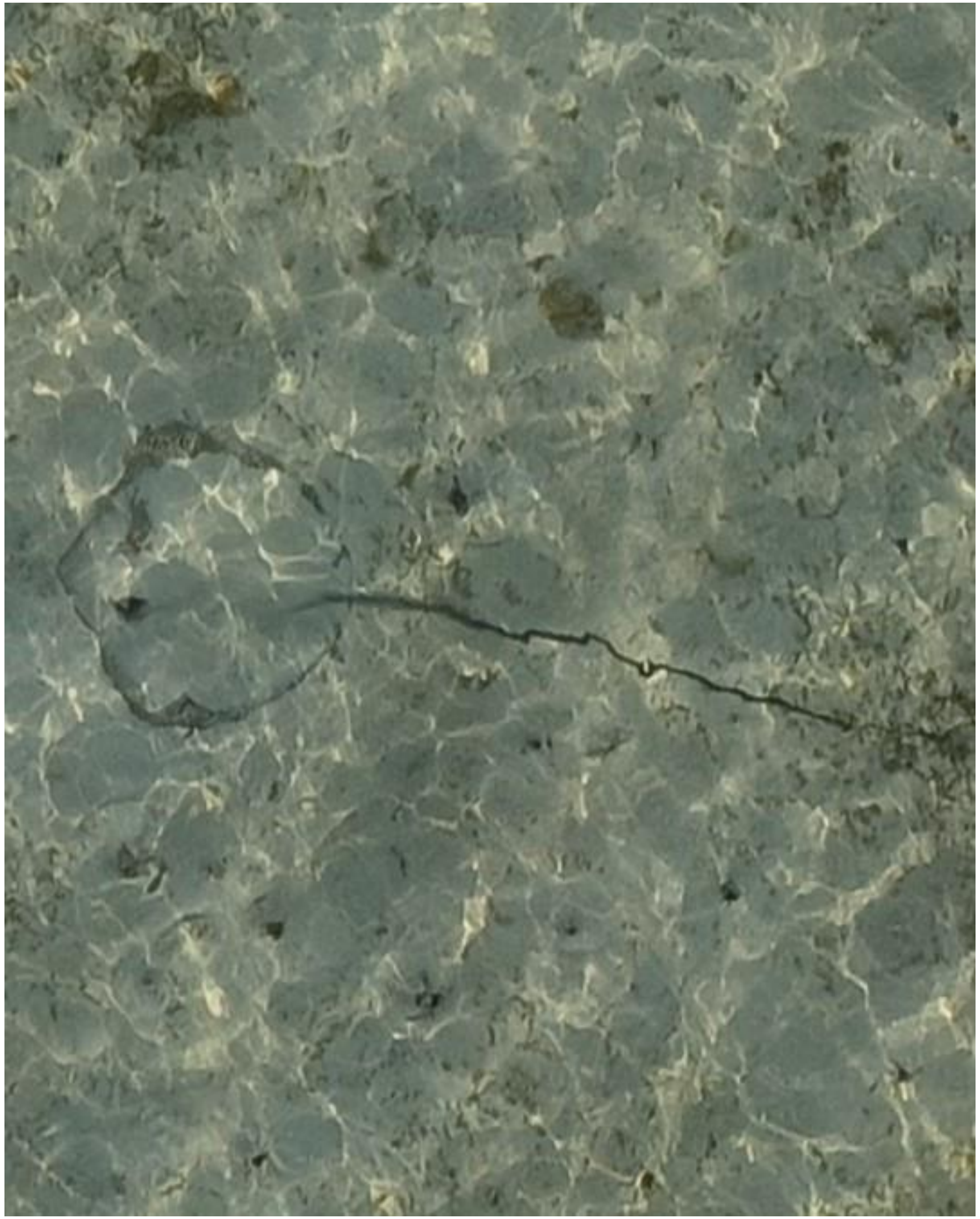

DJI_20240517071350_0050

- Whipray
- Best guess: *H. uarnak*

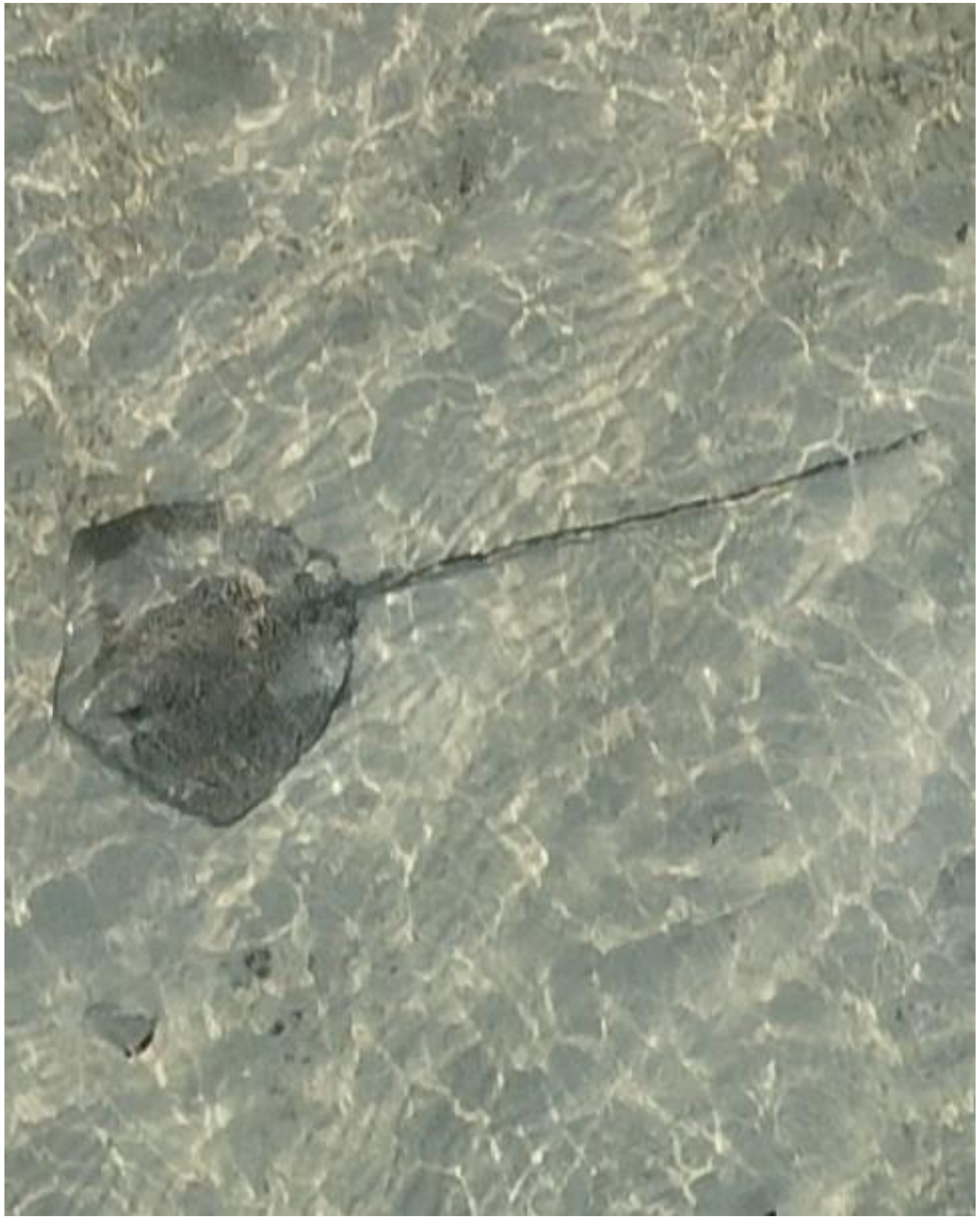

DJI_20240517071357_0057

- *Himantura uarnak*

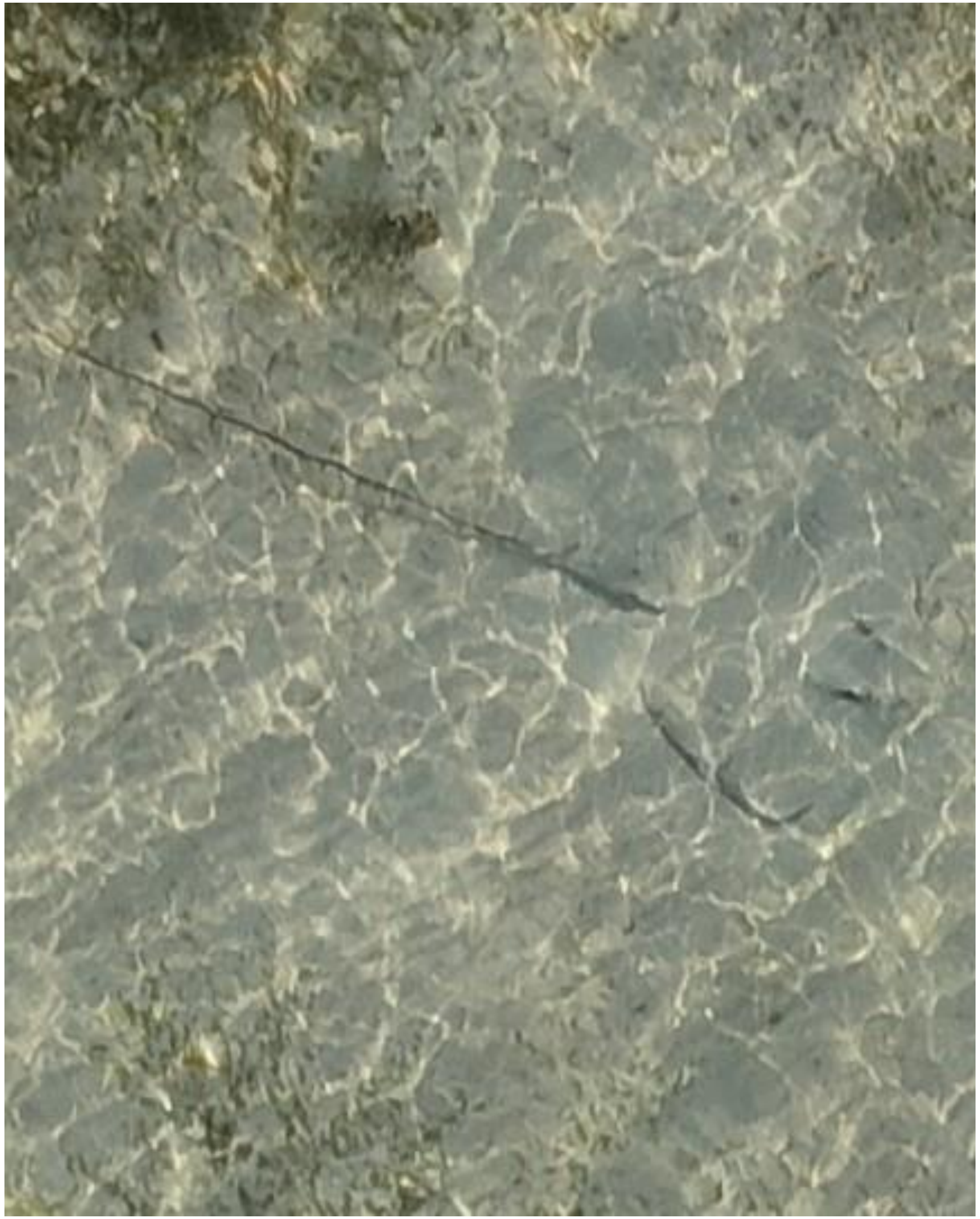

- Whipray

DJI_20240517071357_0057

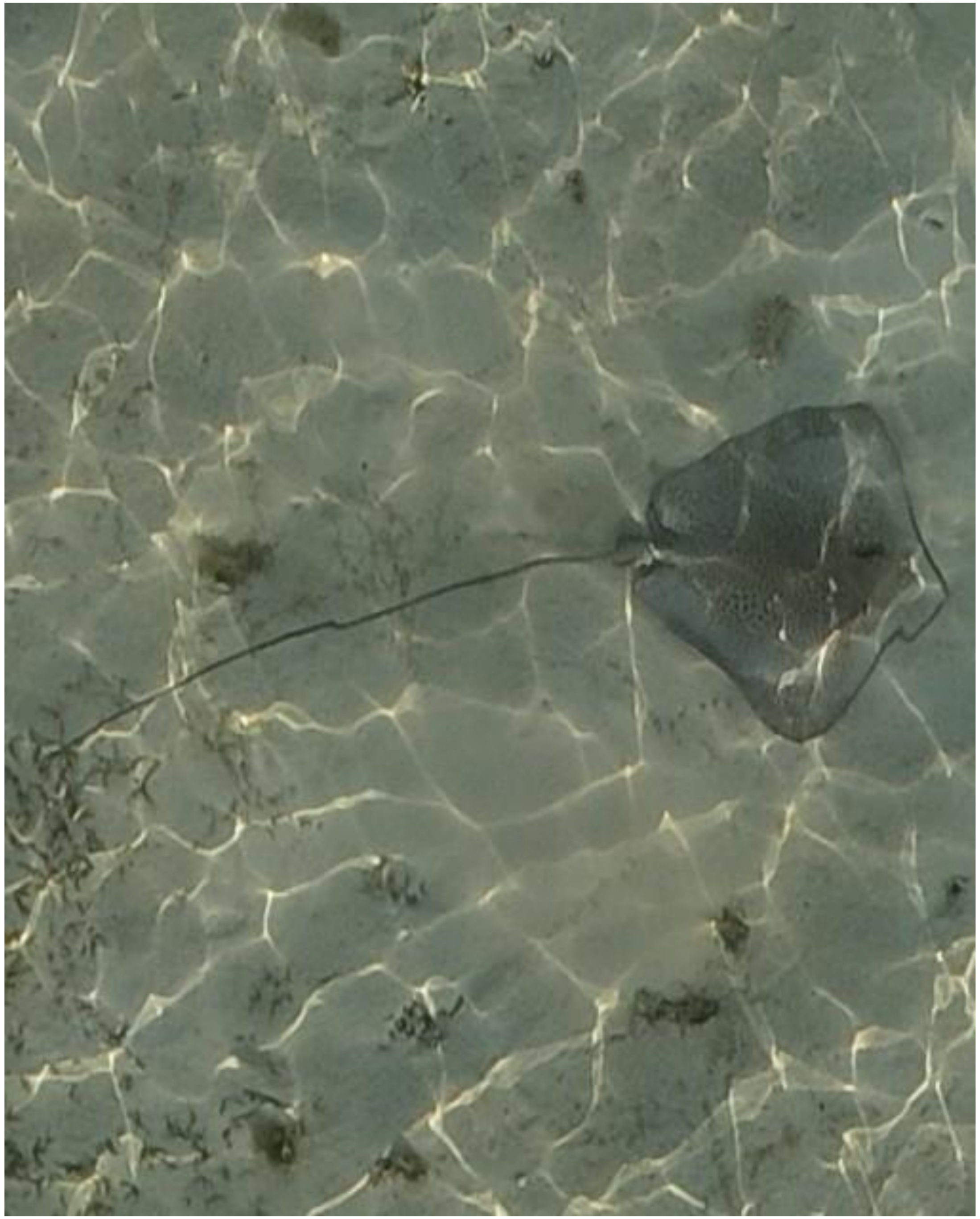

DJI_20240517071450_0098

- *Himantura uarnak*

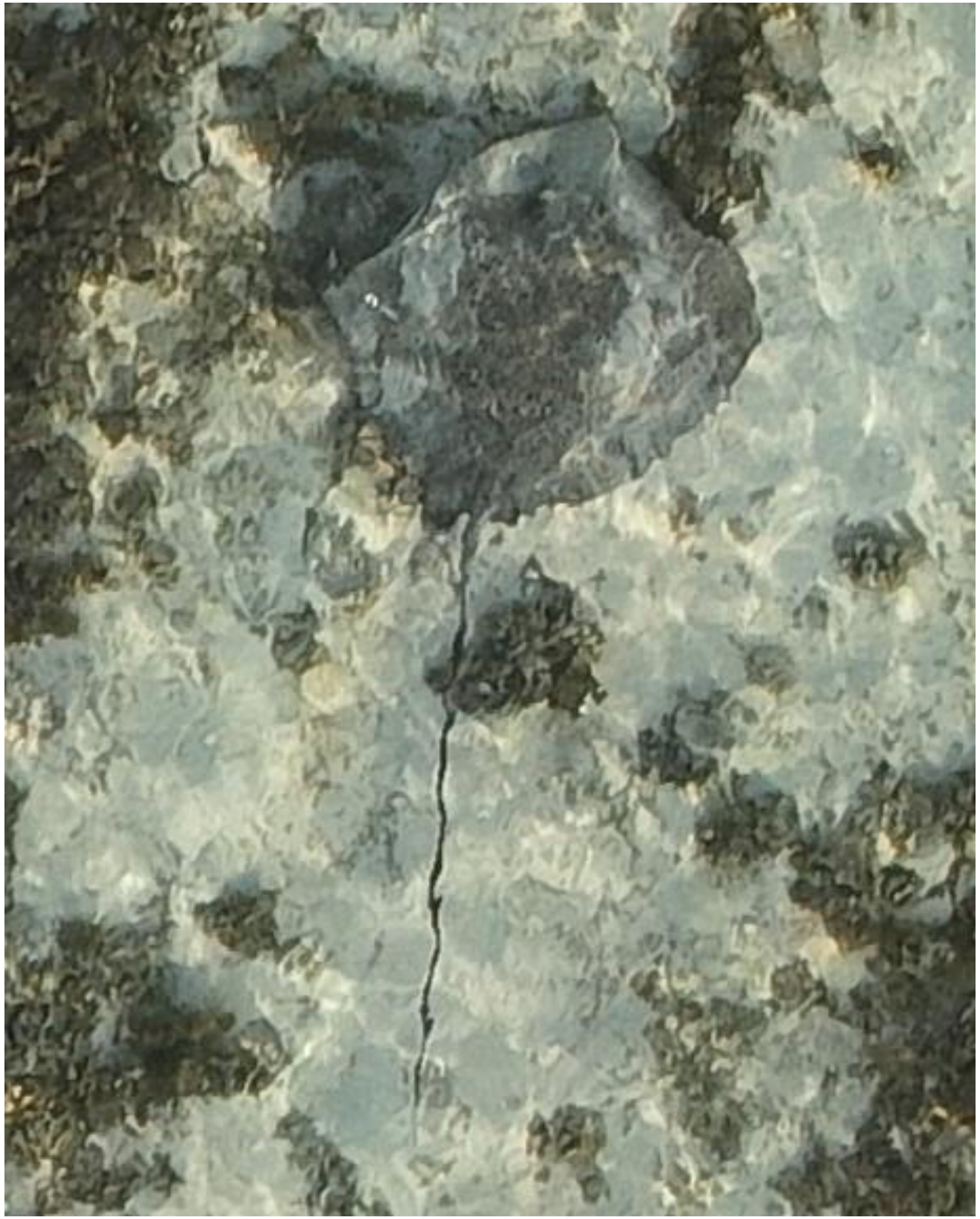
DJI_20240517071531_0139

- *Himantura uarnak*

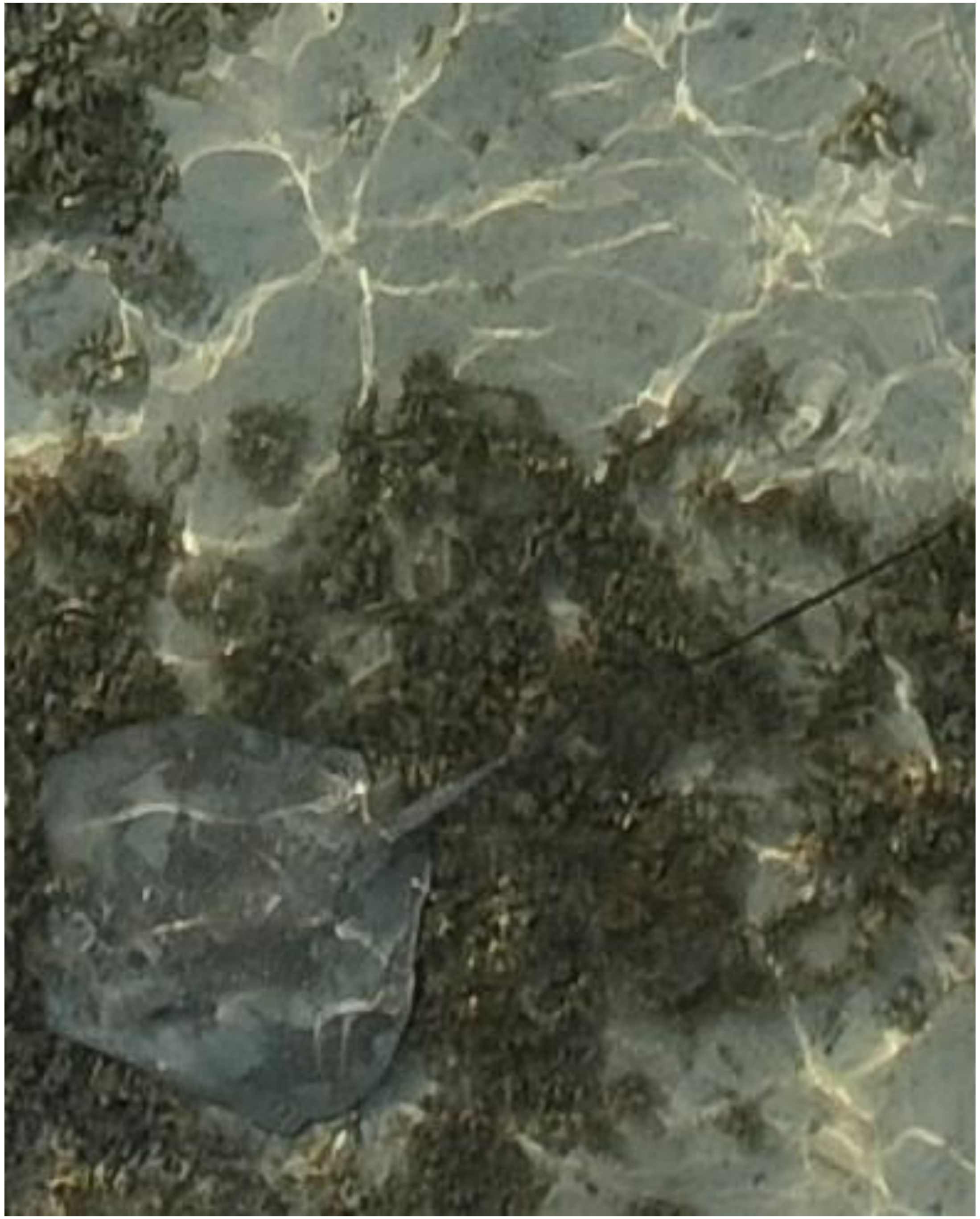

DJI_20240517071653_0209

- *Himantura uarnak*
- Identified as the individual that was also tracked on 05-22 and 06-11. Also observed on 05-10 and 06-03.

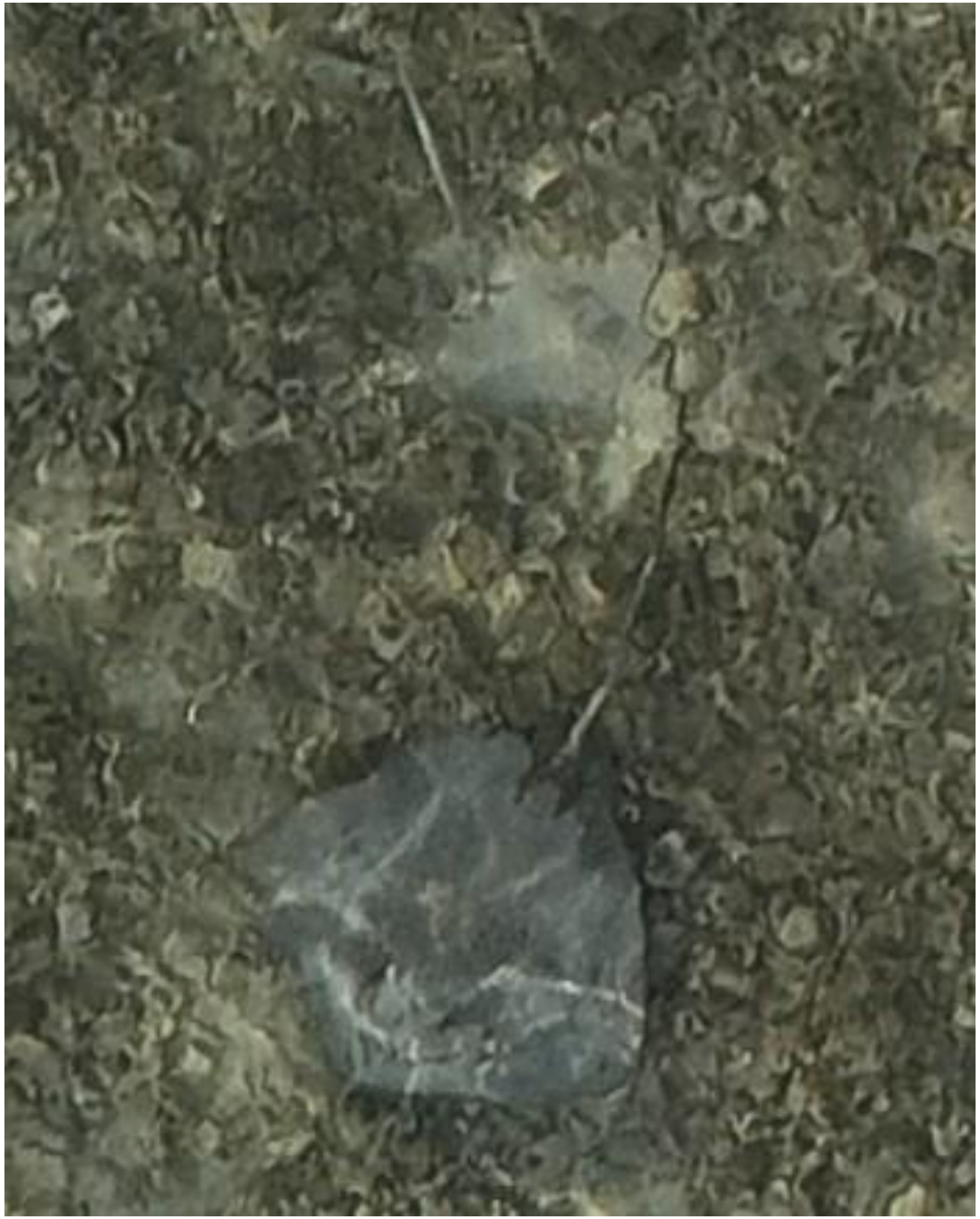
DJI_20240517072352_0575

- Whipray
- Best guess: *H. uarnak*
- Accompanied by needlefish

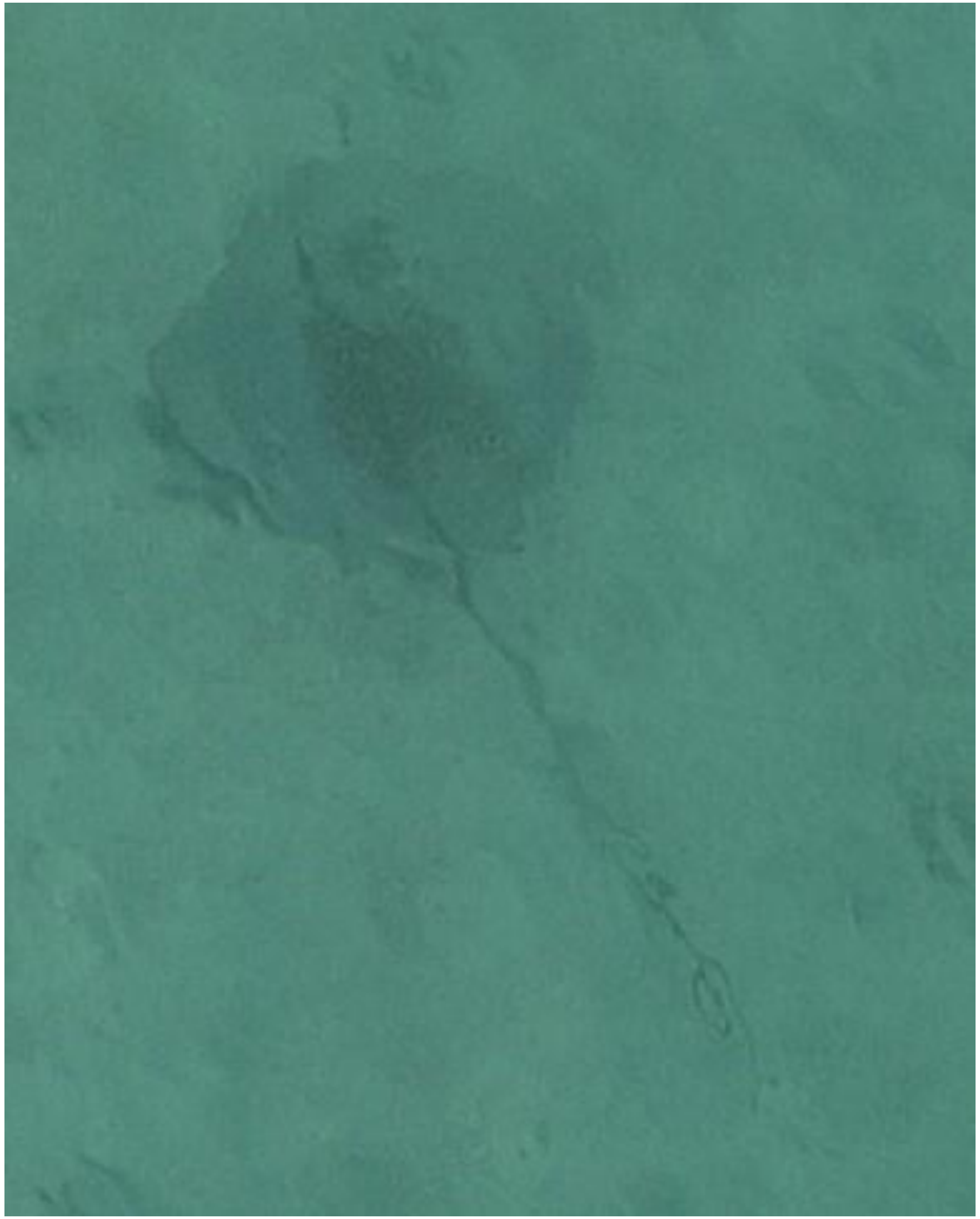

DJI_20240517073012_0915

- Whipray

2024-05-20

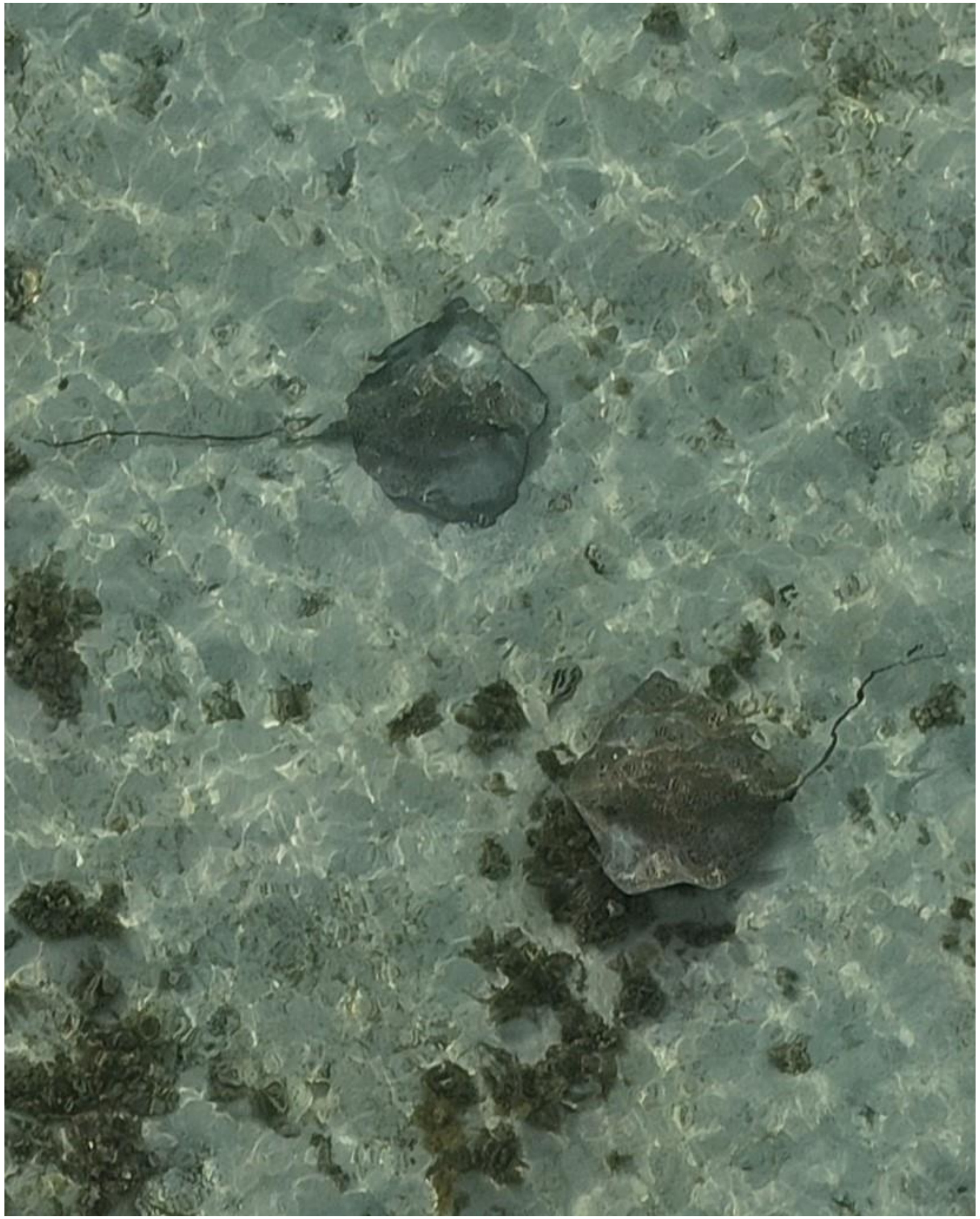
DJI_20240520071418_0033

- *Himantura uarnak* (2x)

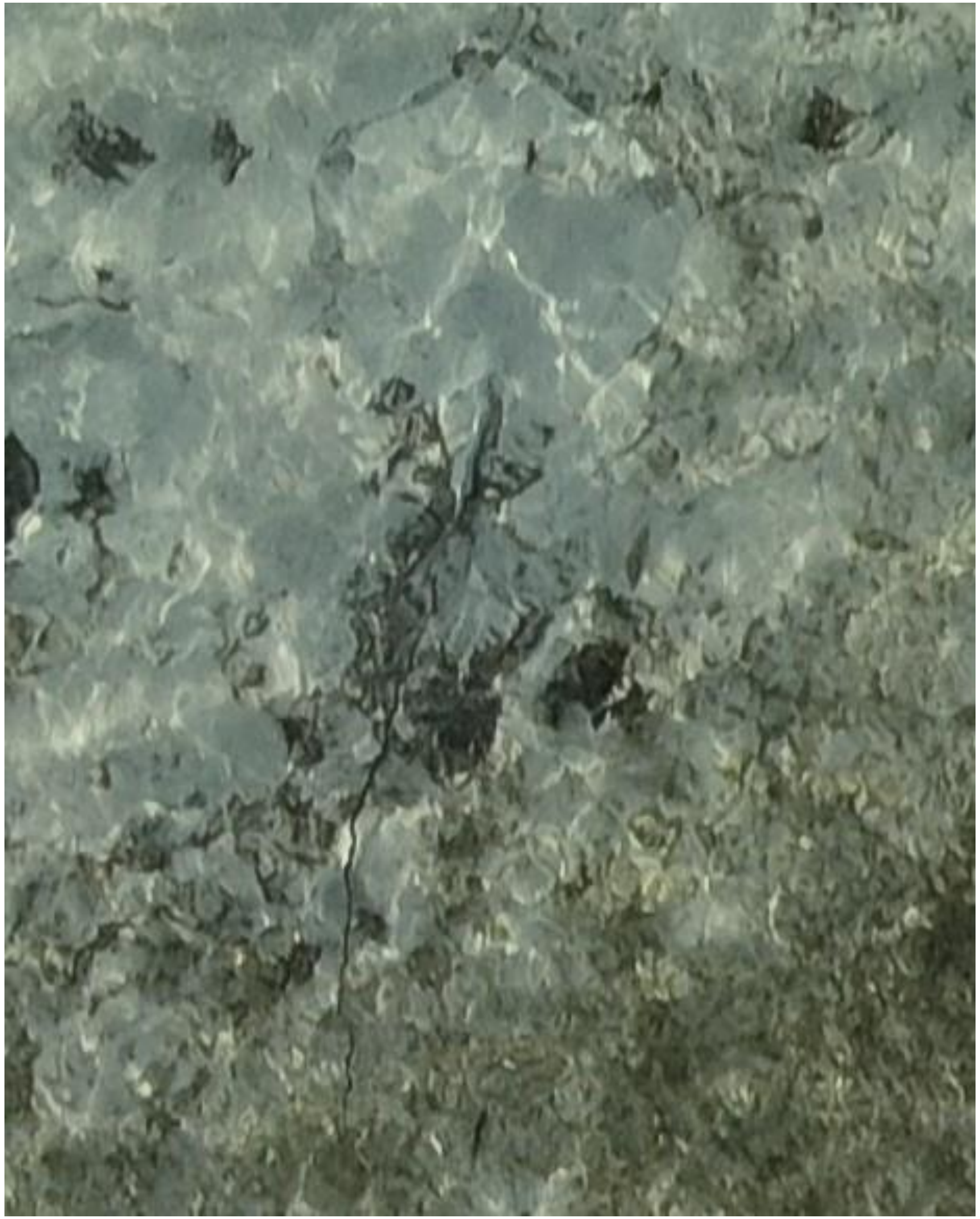
DJI_20240520071441_0056

- Whipray

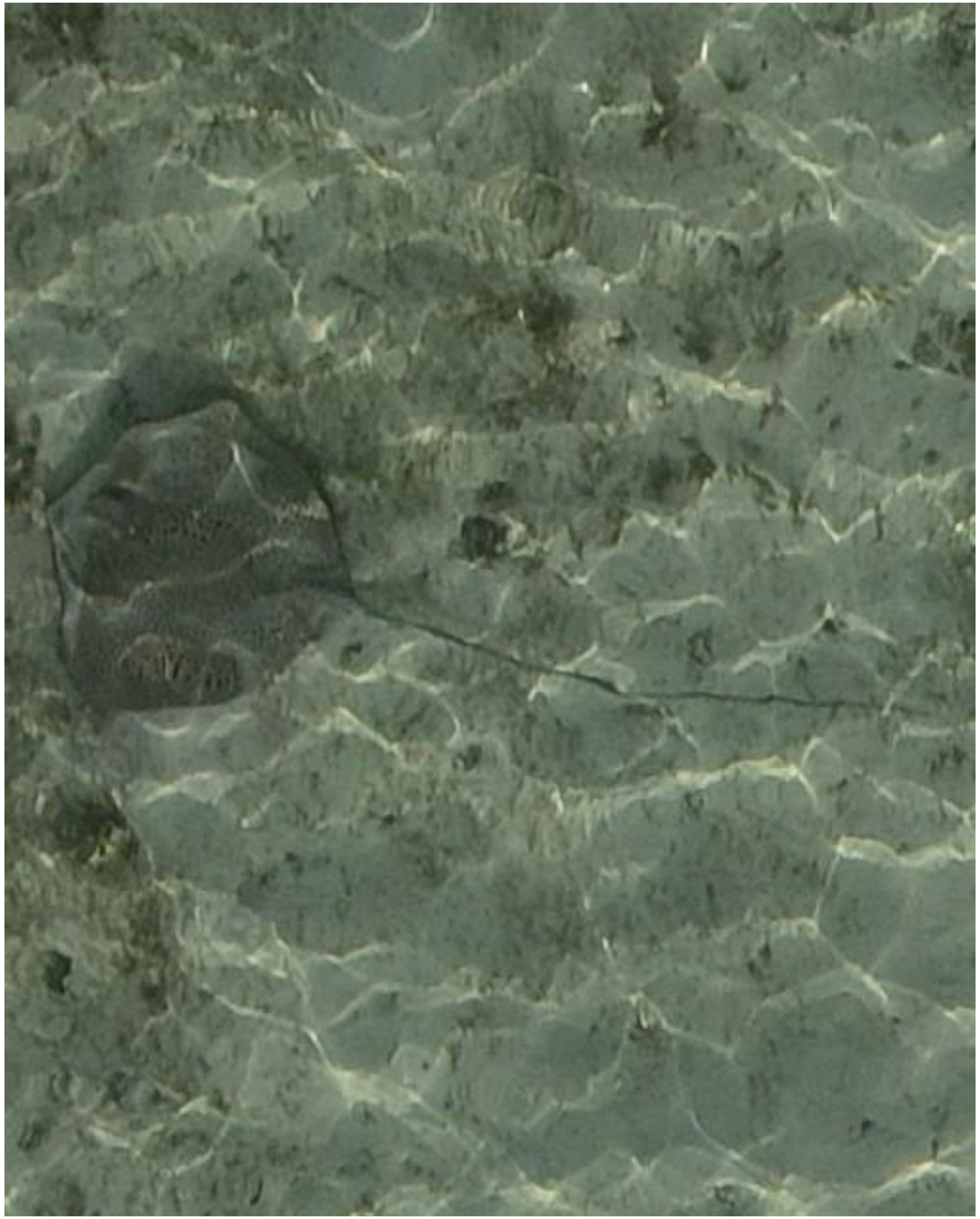

DJI_20240520071855_0275

- *Himantura uarnak*

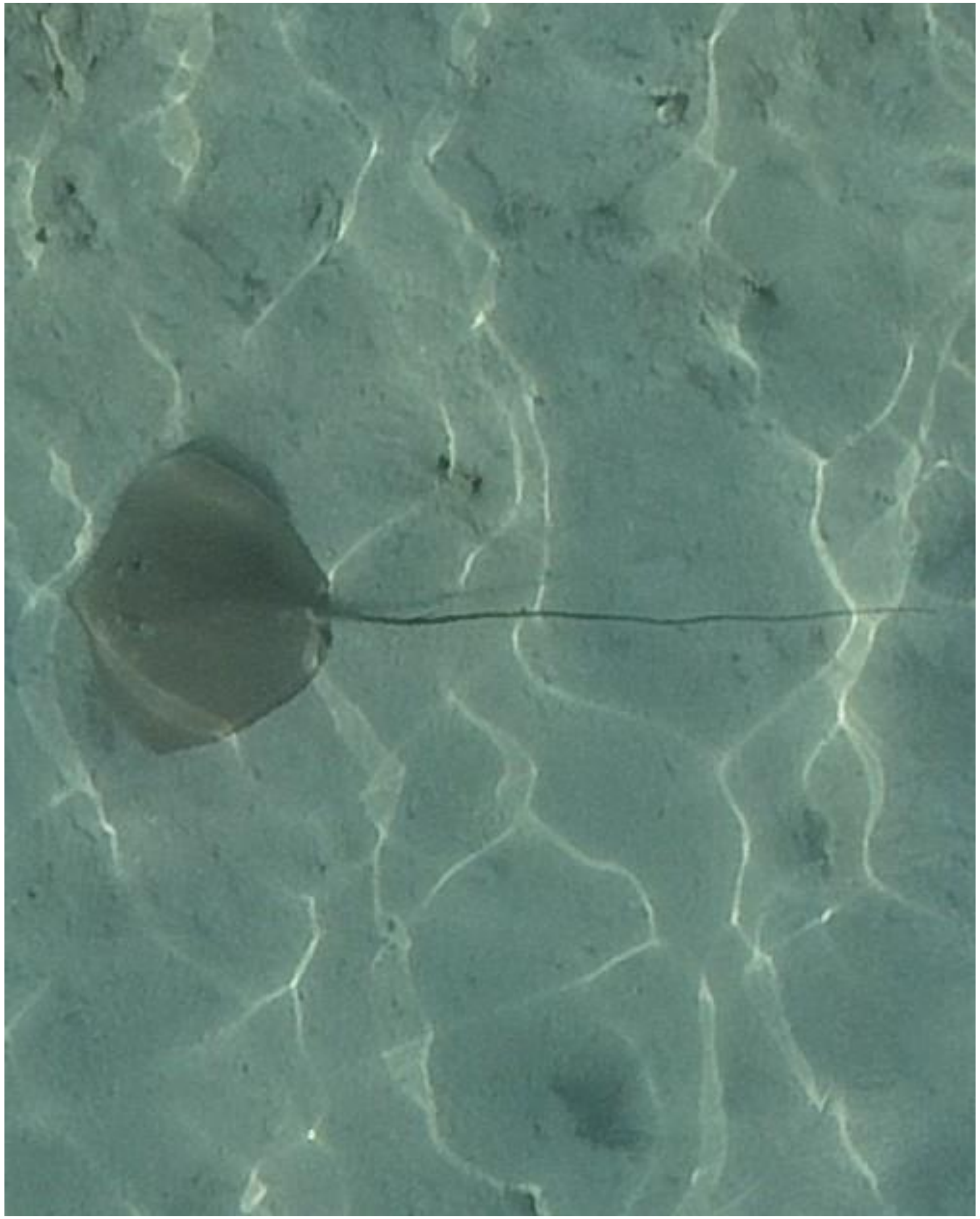

DJI_20240520072321_0500

- cf. *Pateobatis fai*

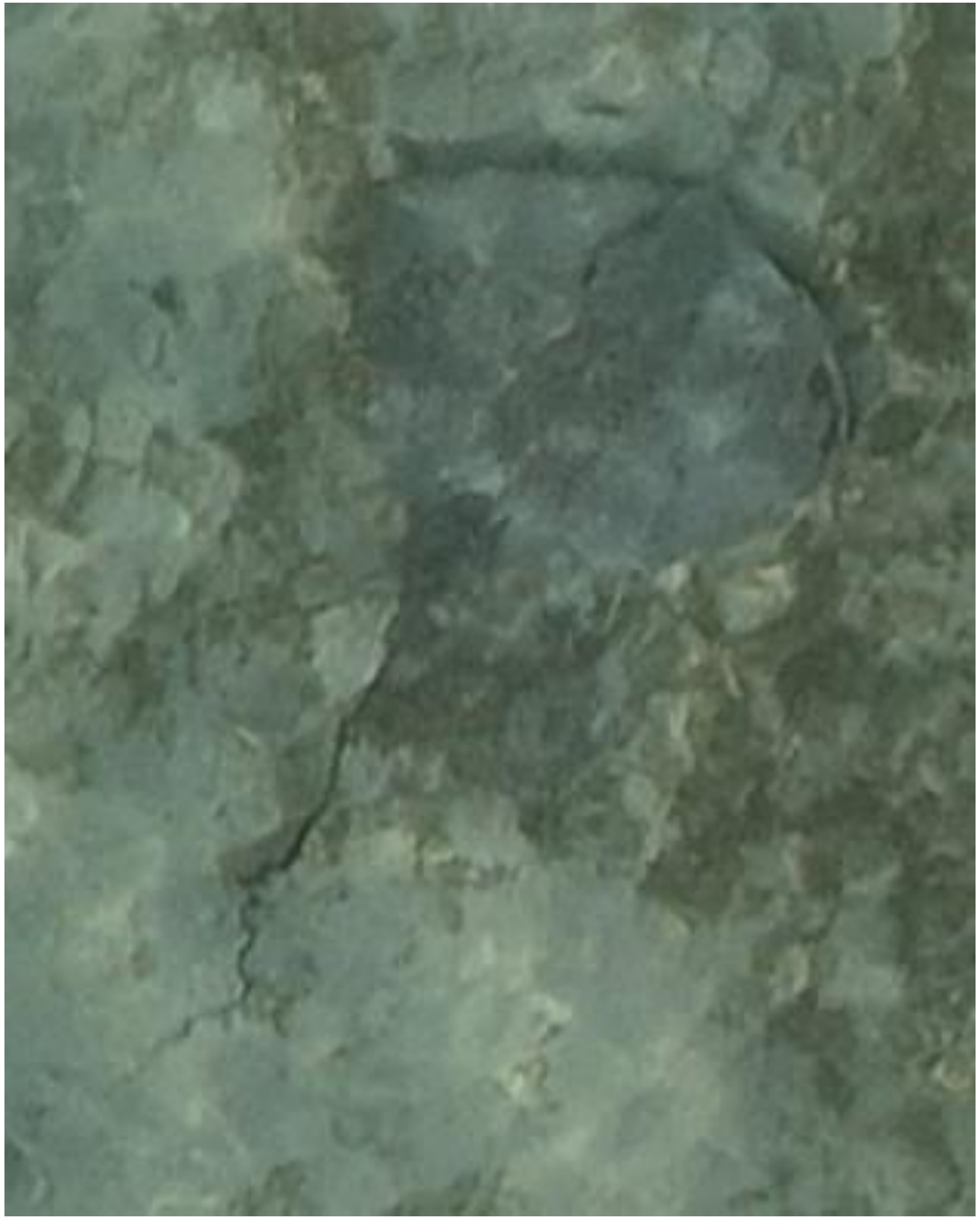

DJI_20240520072448_0587

- Whipray
- Best guess: *H. uarnak*

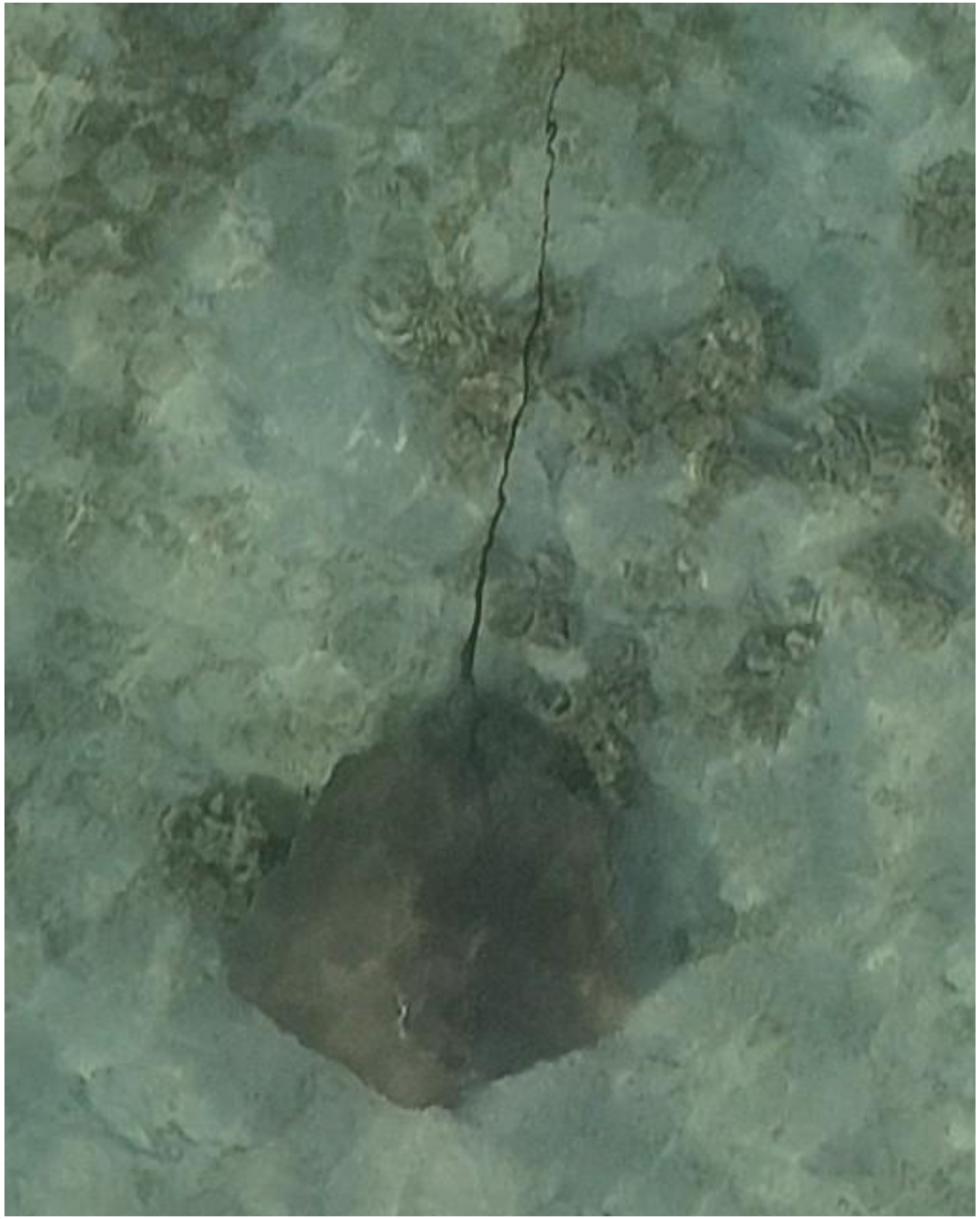

- cf. *Pateobatis fai*

DJI_20240520072453_0592

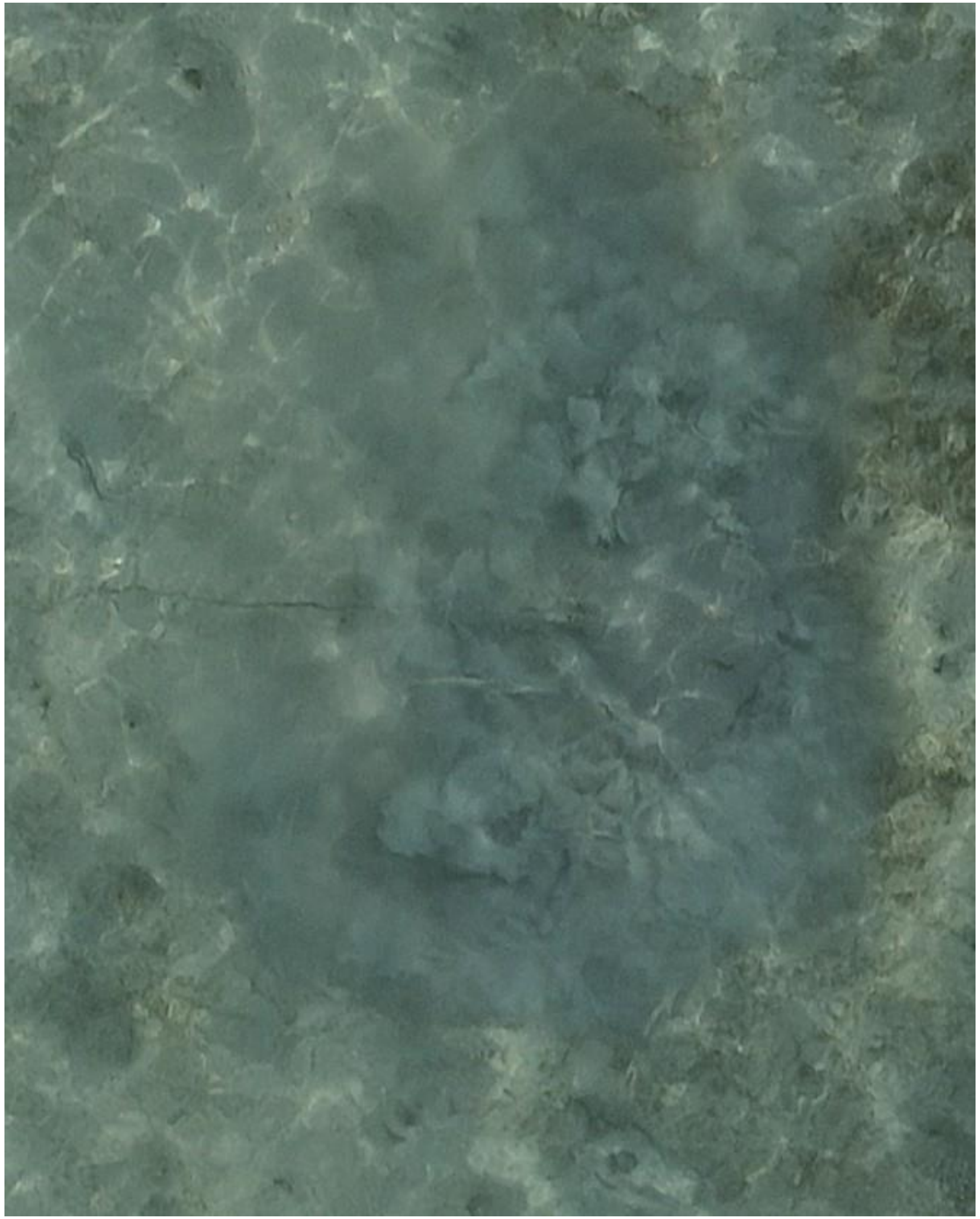

DJI_20240520072645_0687

- Whipray
- Accompanied by needlefish

2024-05-22

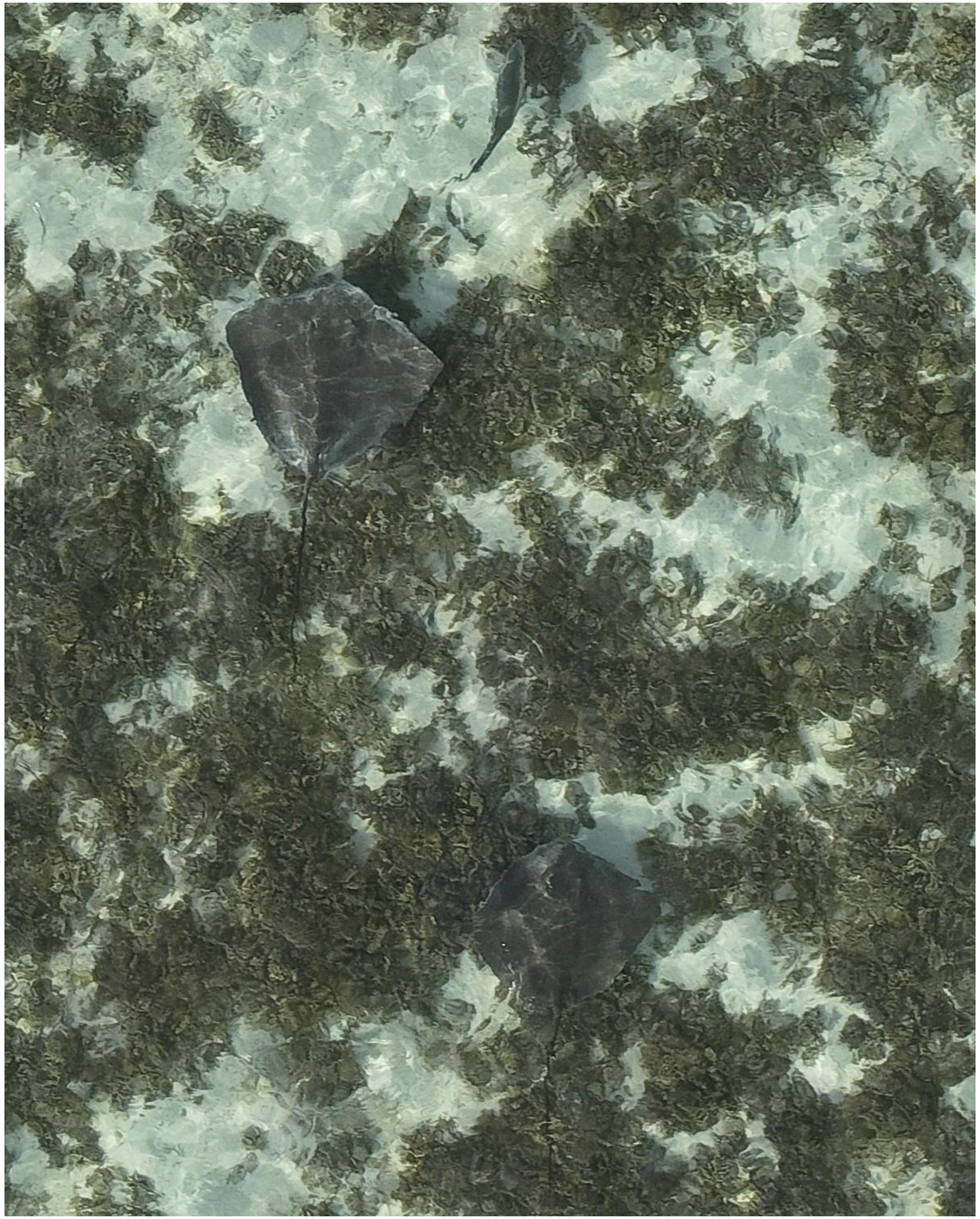

DJI_20240522071804_0140

- cf. *Pateobatis fai* (2x)
- Appears to be the same duo as was tracked on 05-10. Again accompanied by a trevally.

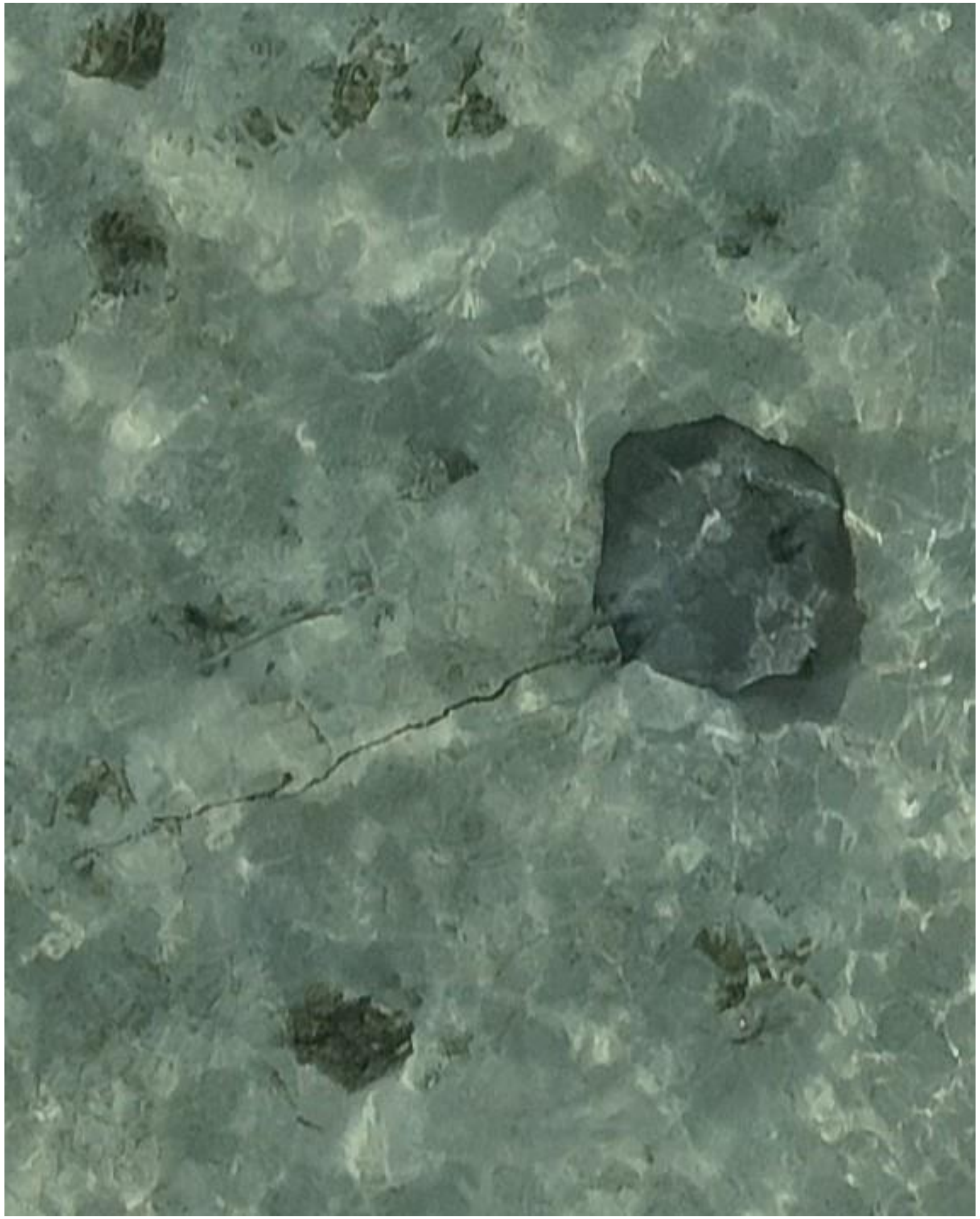

- Whipray
- Best guess: cf. *P. fai*
- Accompanied by needlefish

DJI_20240522072315_0415

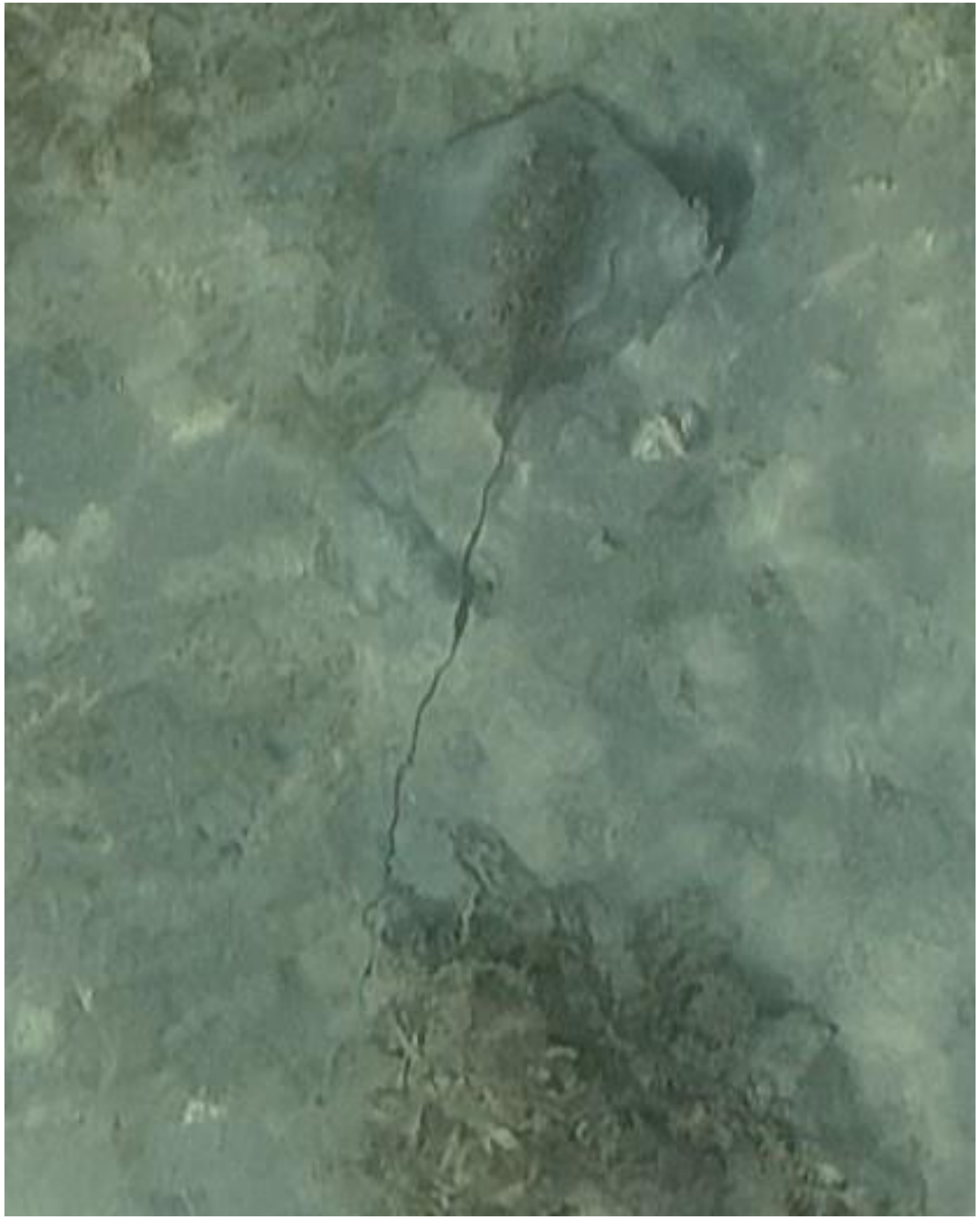

- *Himantura uarnak*

DJI_20240522072633_0583

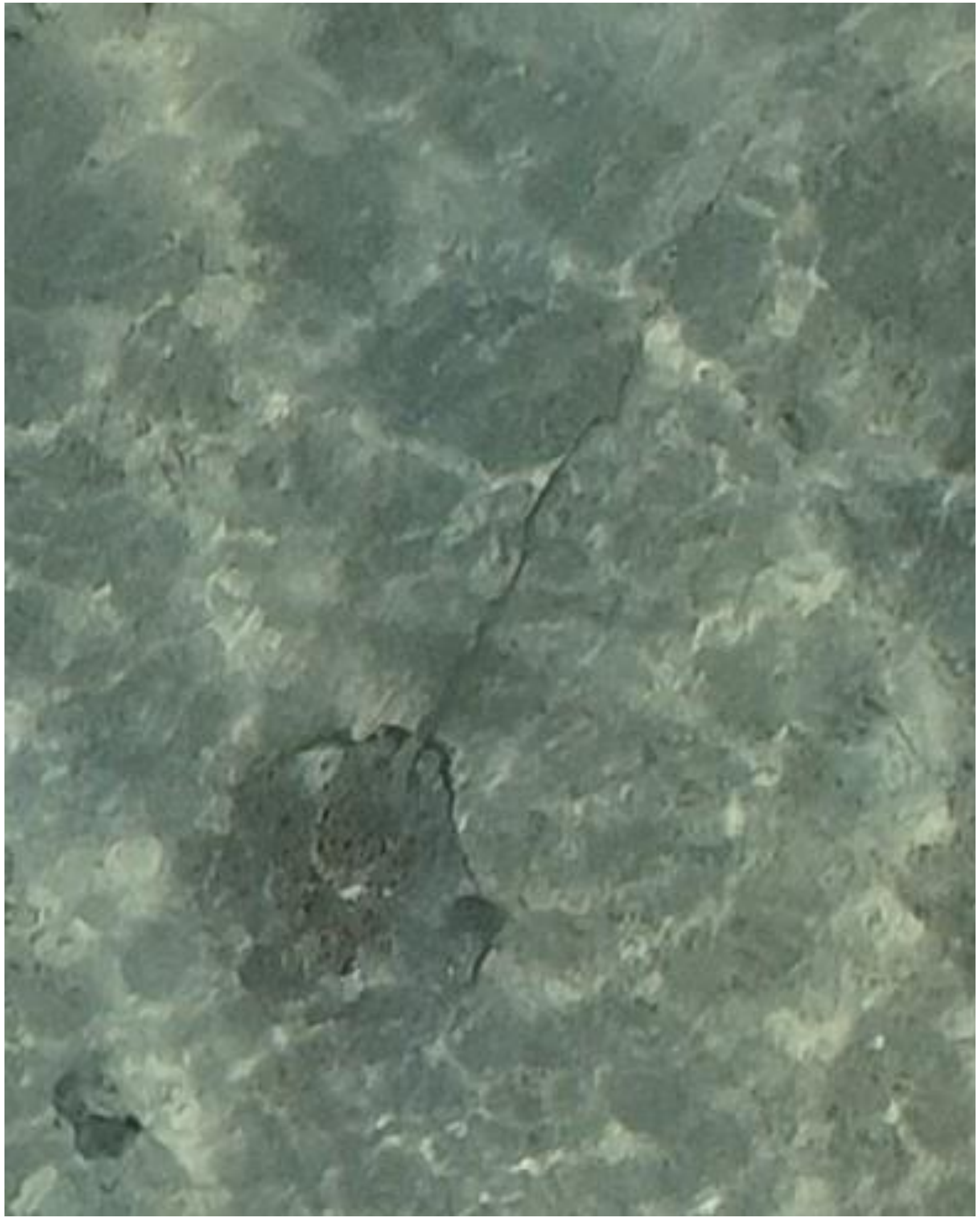

- *Himantura uarnak*

DJI_20240522073012_0772

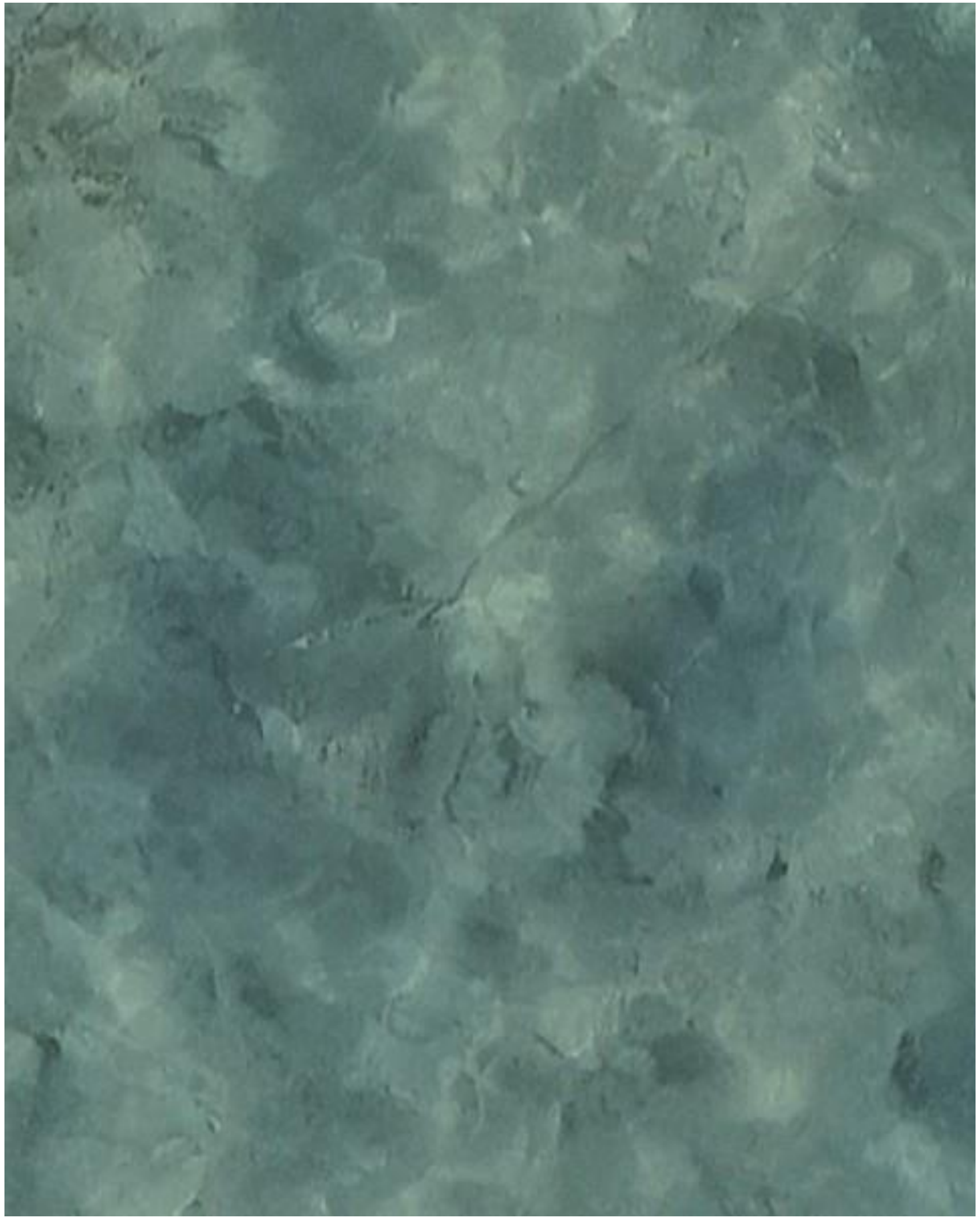

- Whipray

DJI_20240522073018_0778

2024-05-23

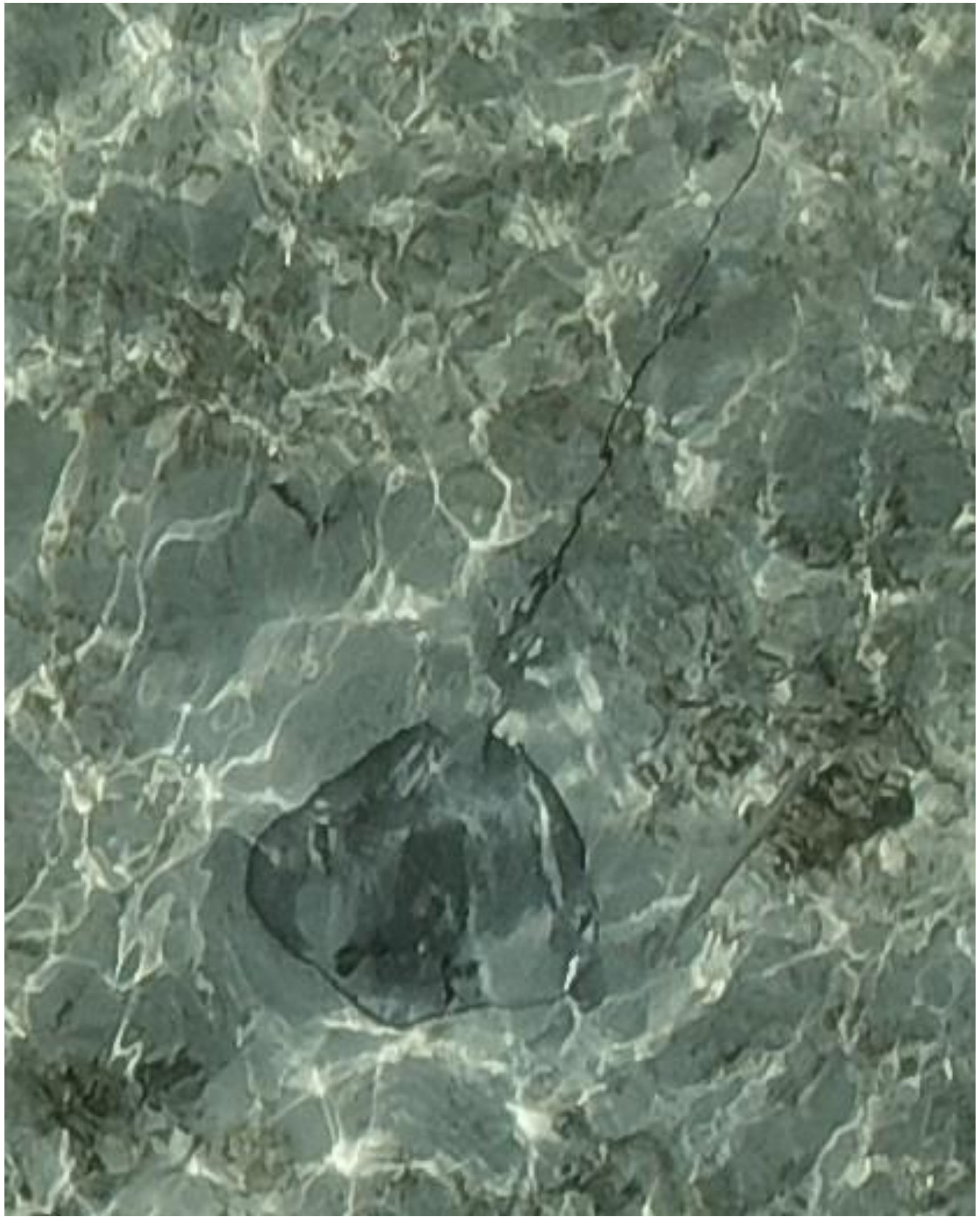

- Whipray
- Best guess: cf. *P. fai*
- Accompanied by needlefish

DJI_20240523075237_0397

2024-05-24

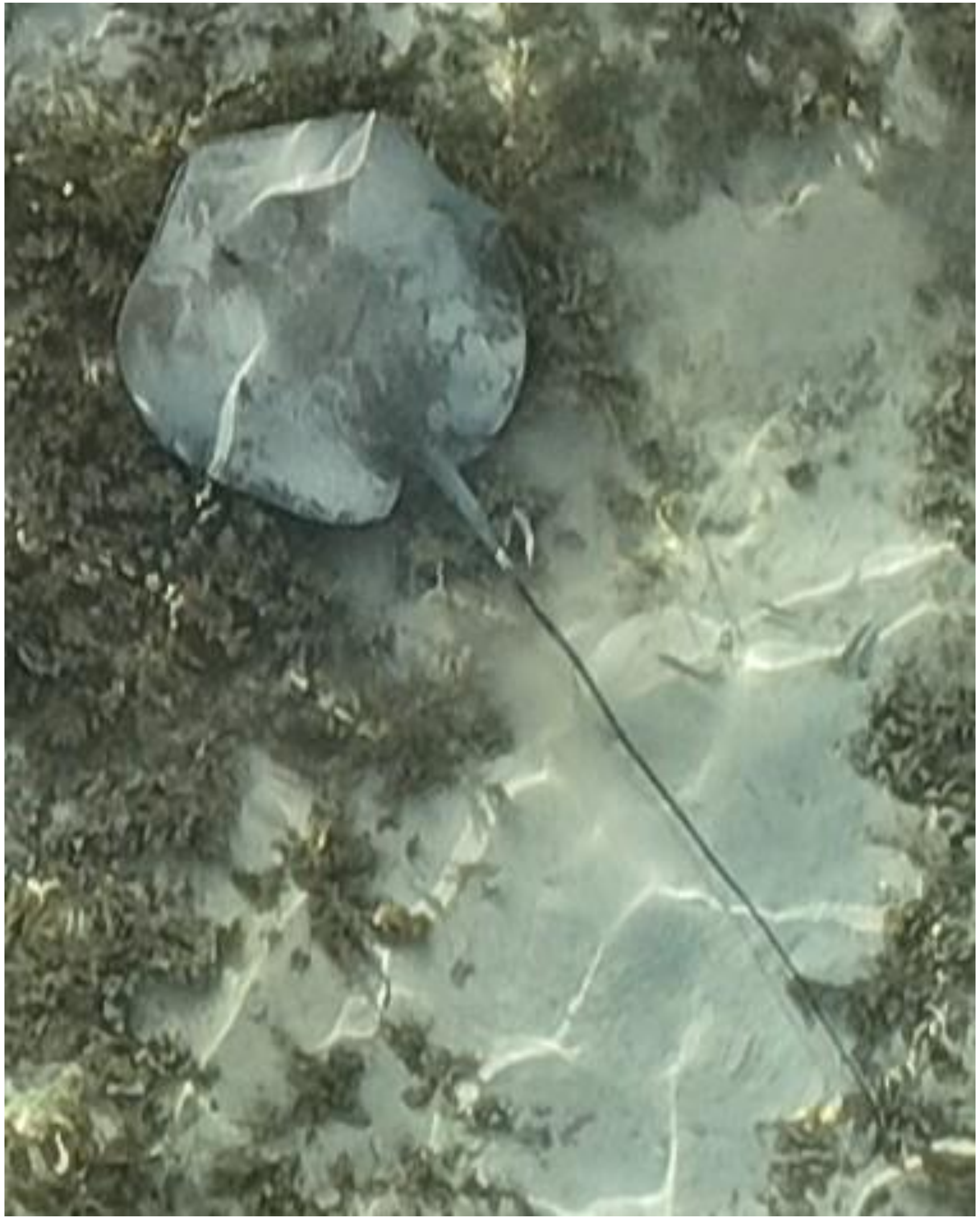

DJI_20240524072431_0179

- Whipray

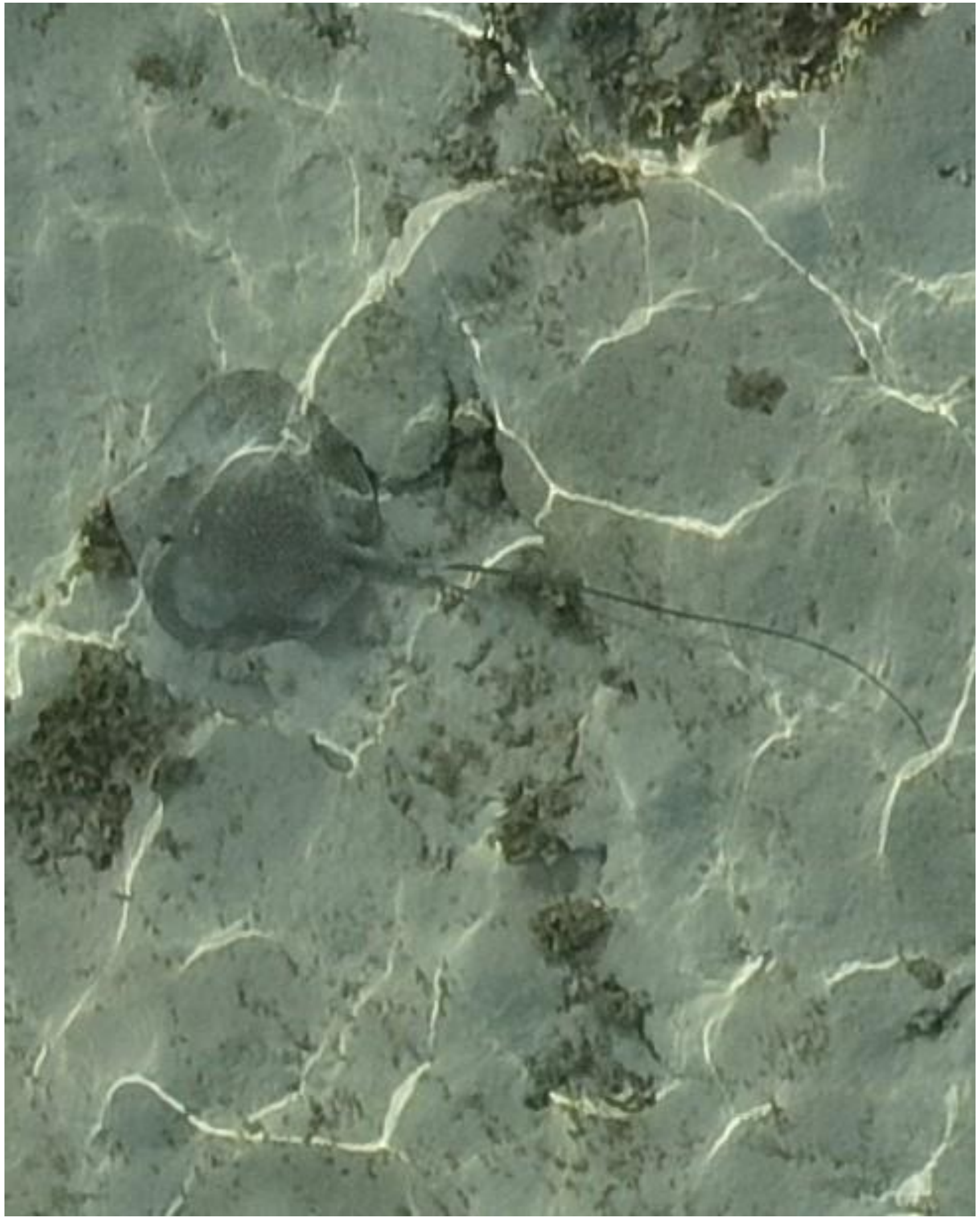

DJI_20240524072447_0195

- *Himantura uarnak*

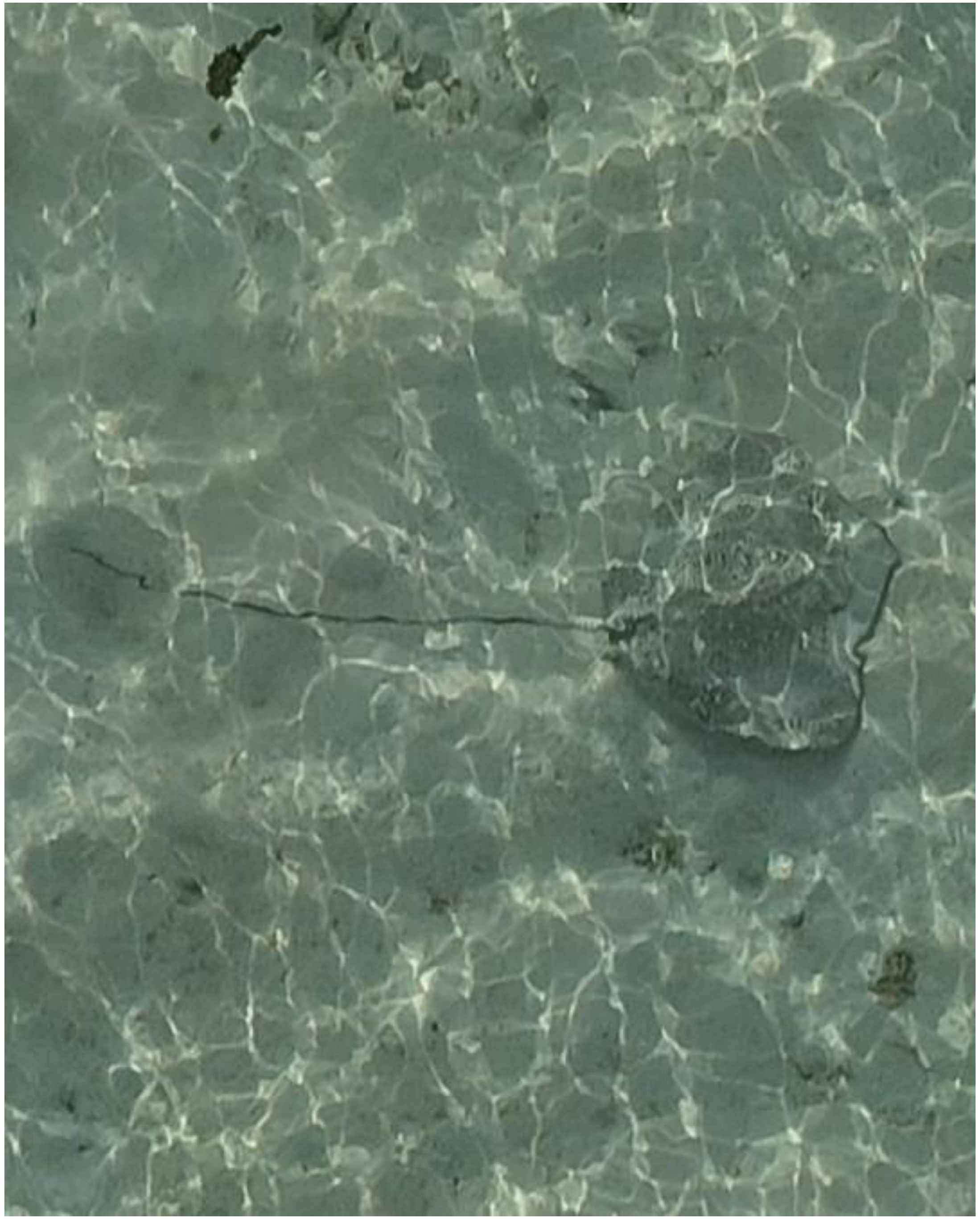

DJI_20240524072845_0409

- *Himantura uarnak*

2024-06-03

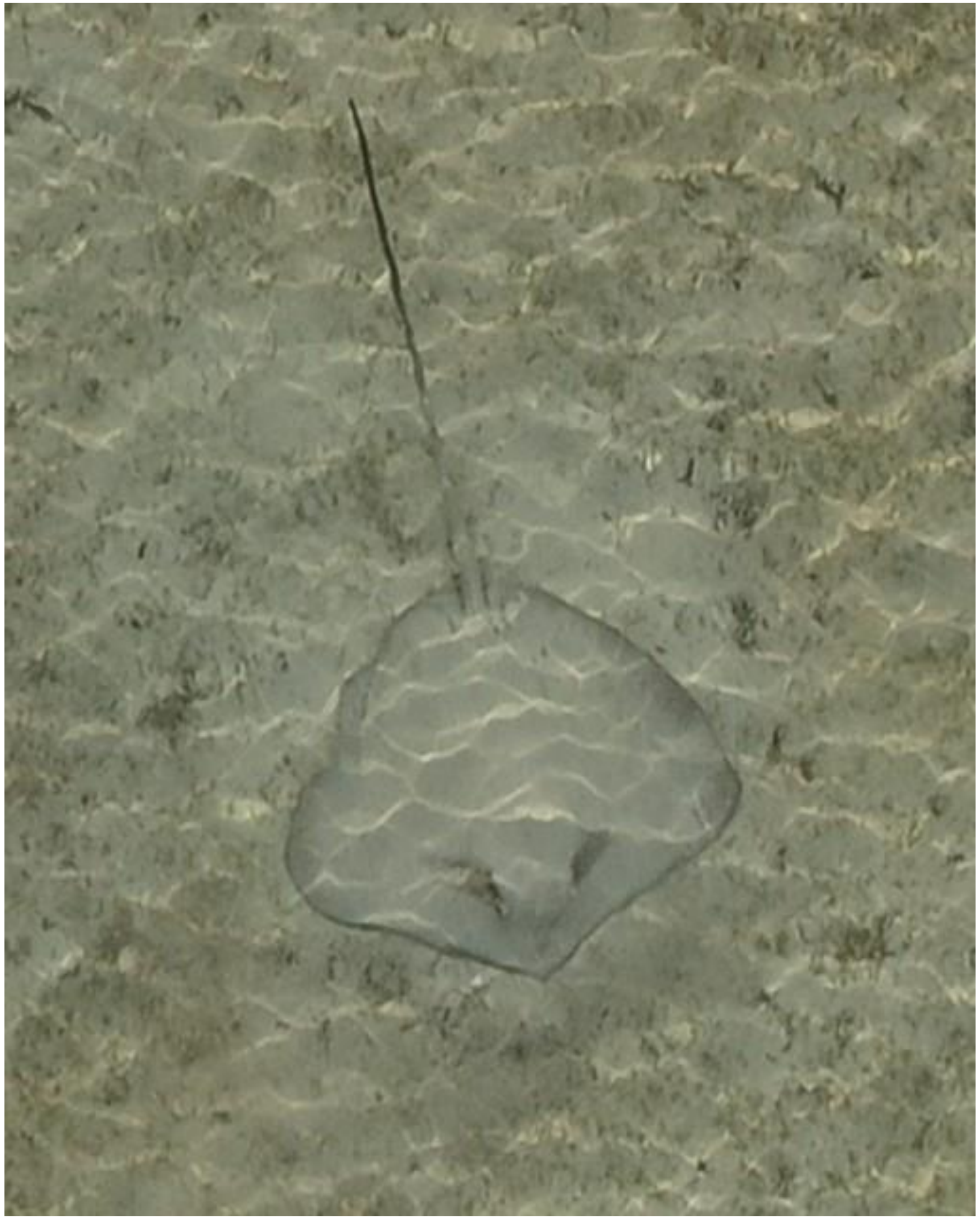

DJI_20240603072558_0056

- Whipray

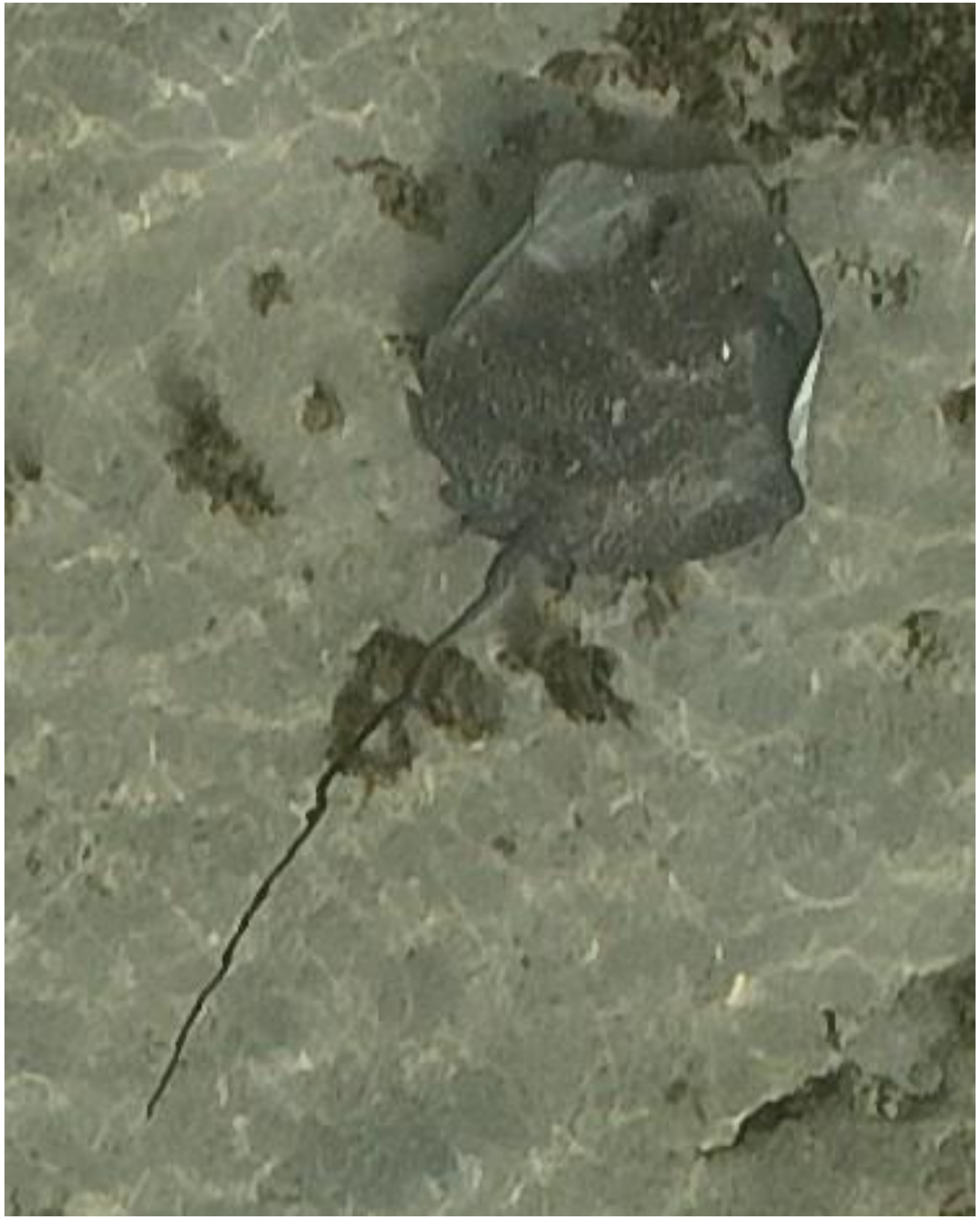

DJI_20240603072659_0106

- *Himantura uarnak*
- Identified as the individual that was also tracked on 05-22 and 06-11. Also observed on 05-10 and 05-17.

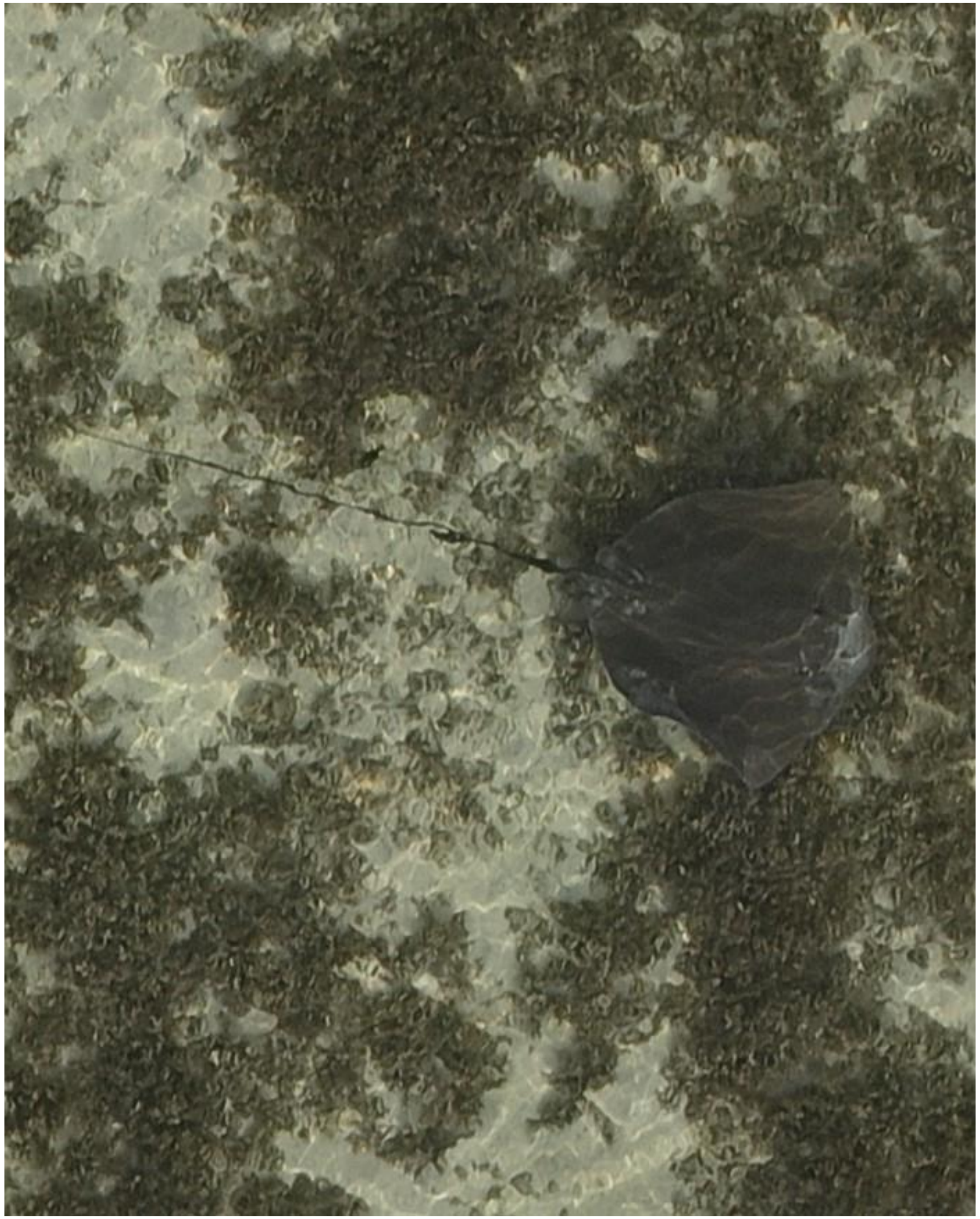

- cf. *Pateobatis fai*

DJI_20240603072731_0138

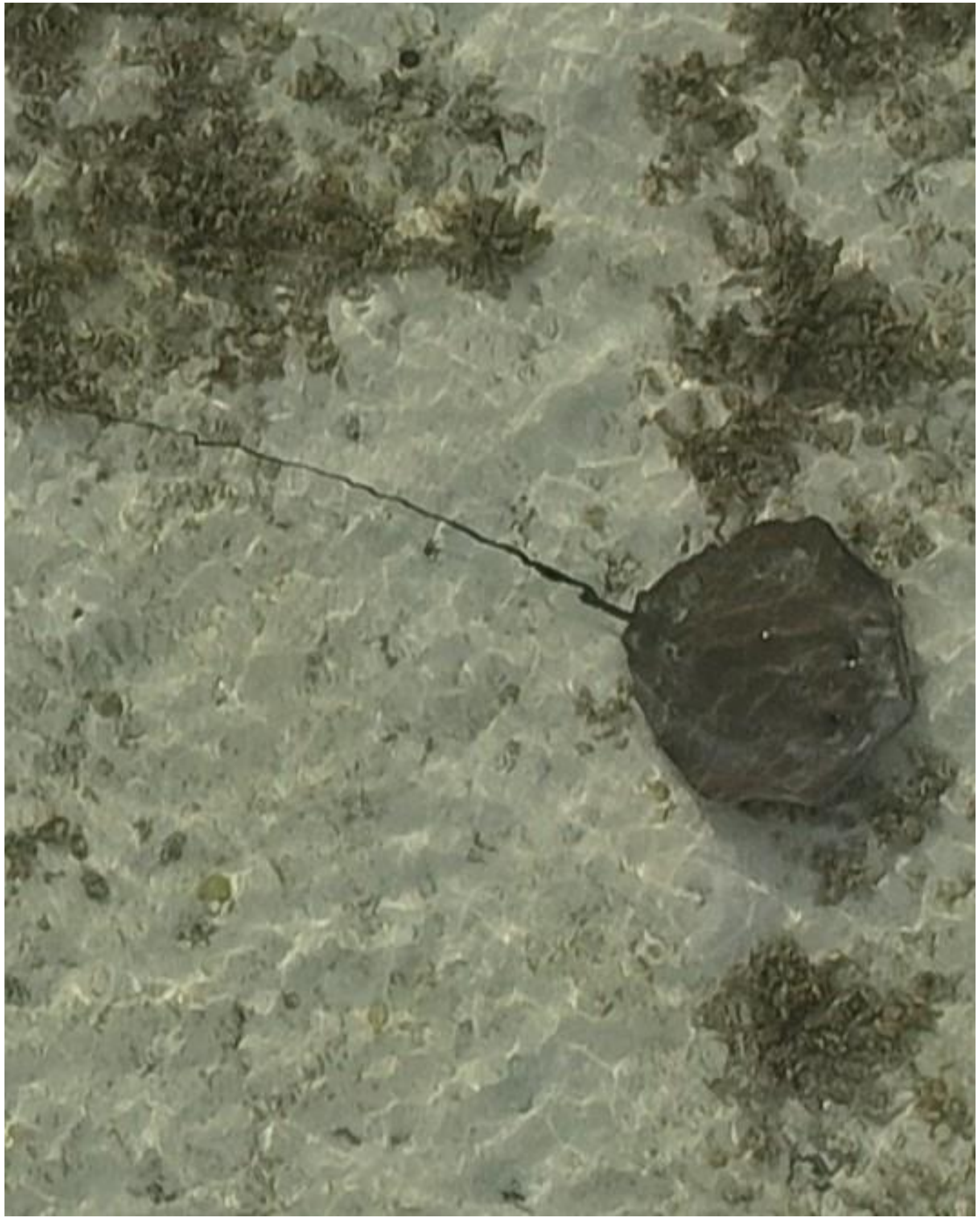

DJI_20240603072828_0183

- cf. *Pateobatis fai*

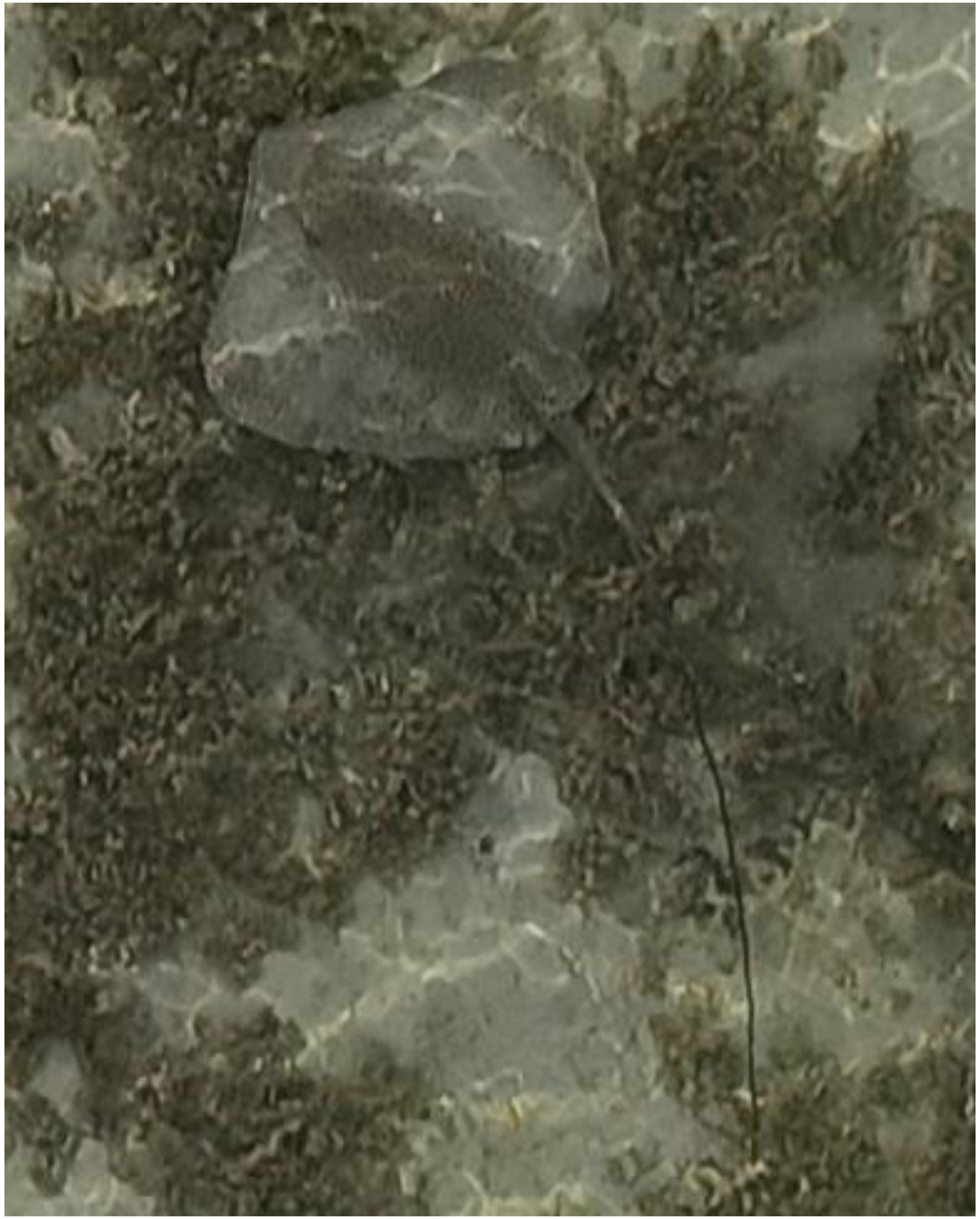

DJI_20240603072856_0211

- *Himantura uarnak*

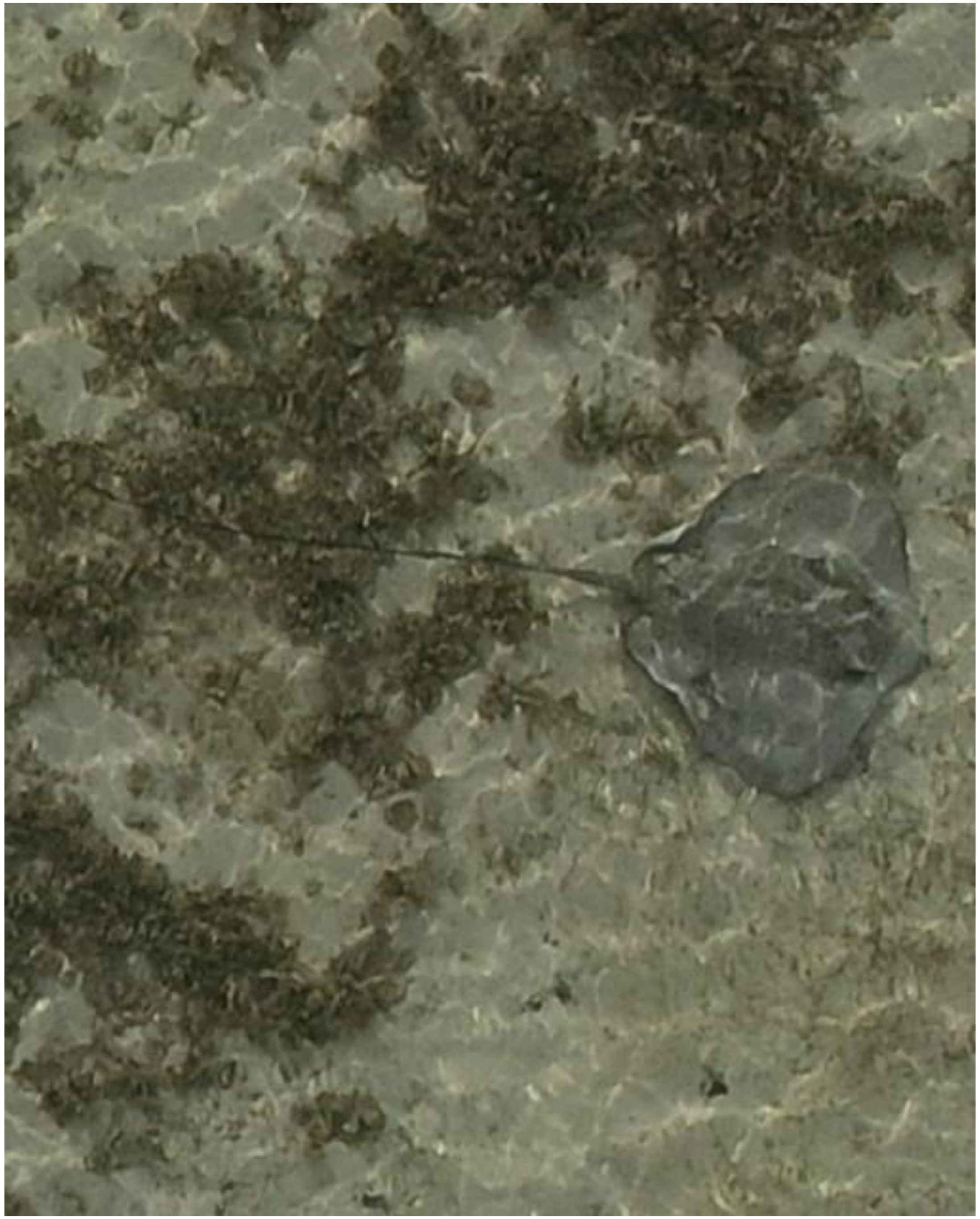

- Whipray

DJI_20240603072857_0212

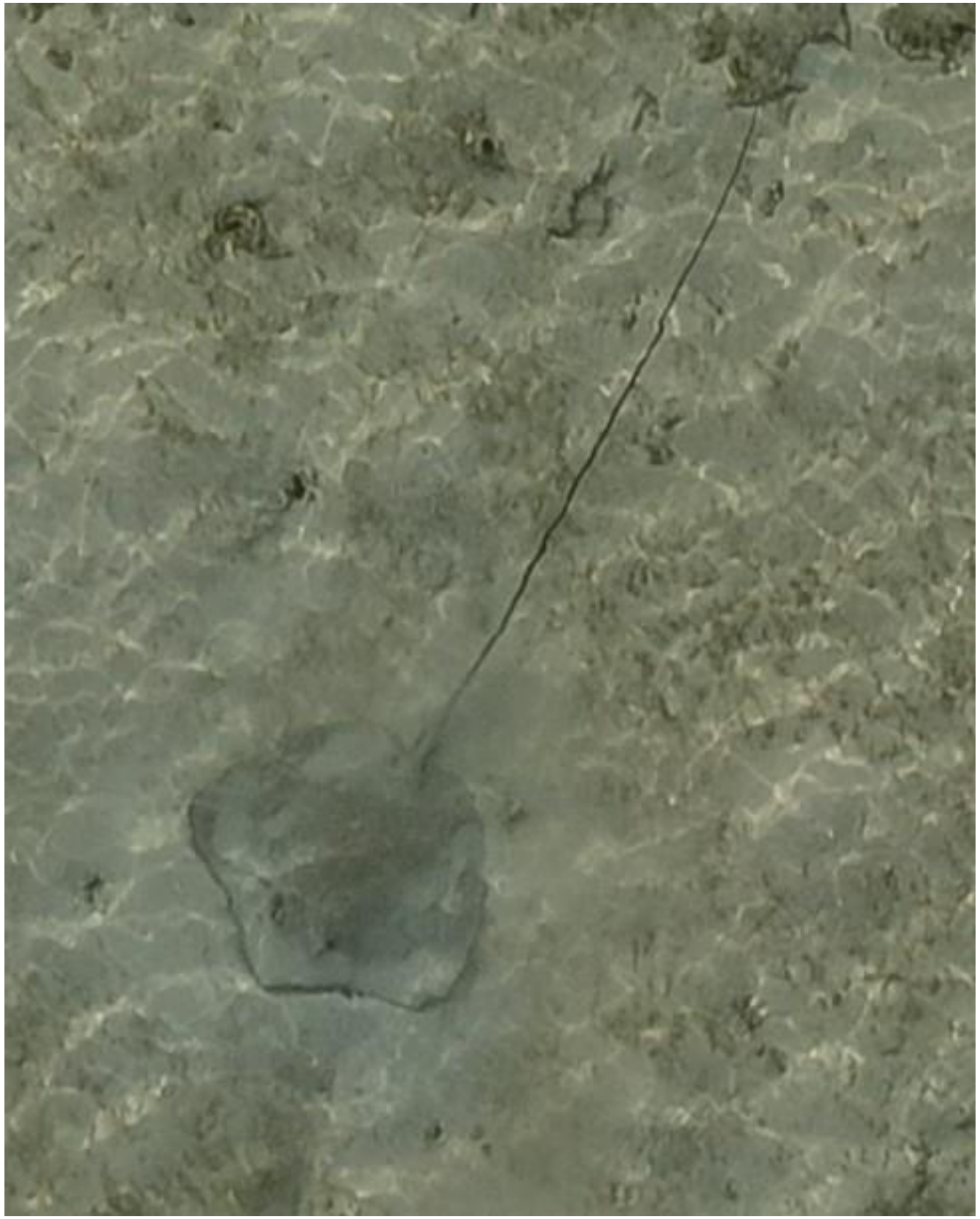

- Whipray

DJI_20240603072901_0216

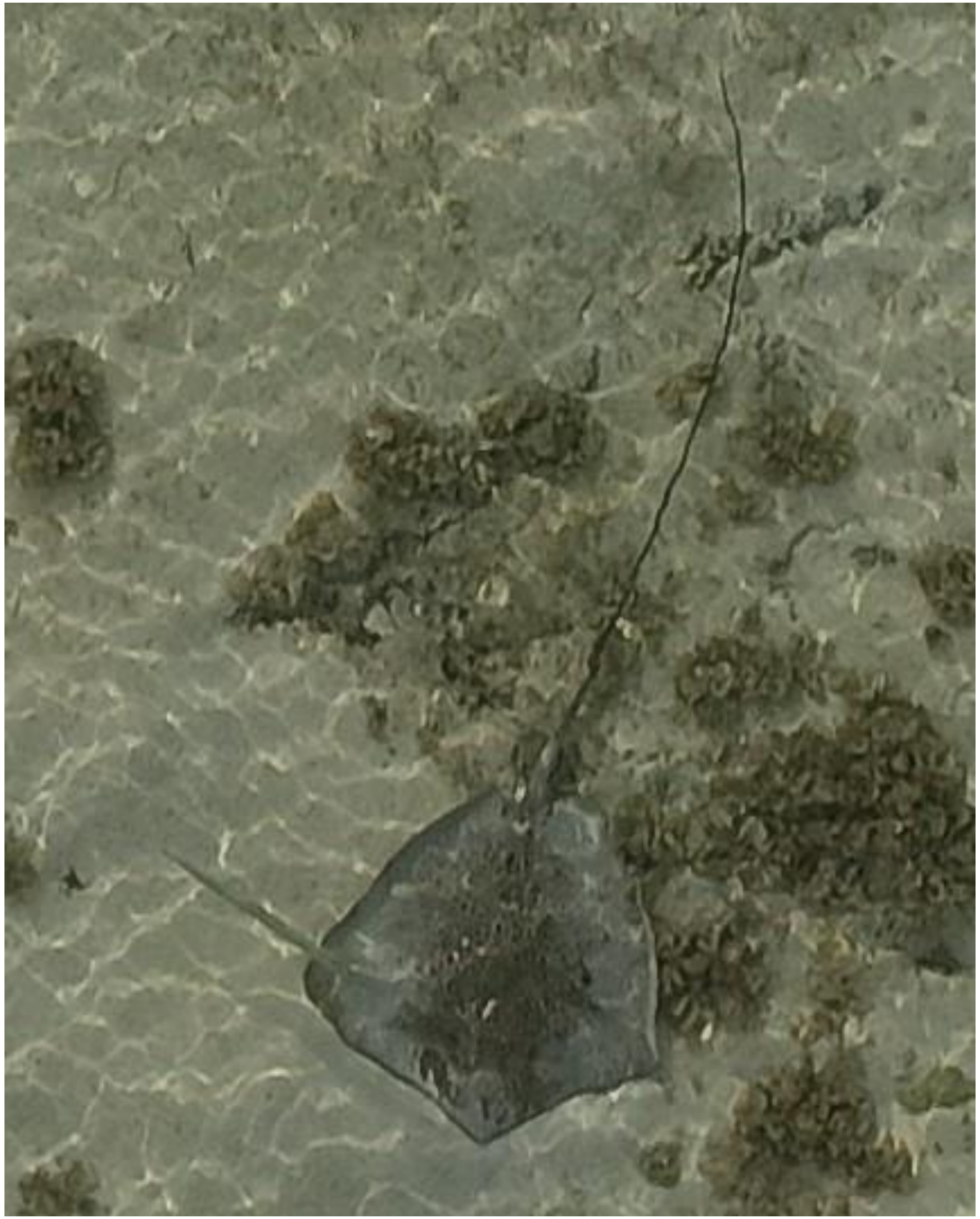

DJI_20240603072908_0223

- *Himantura uarnak*
- Accompanied by needlefish

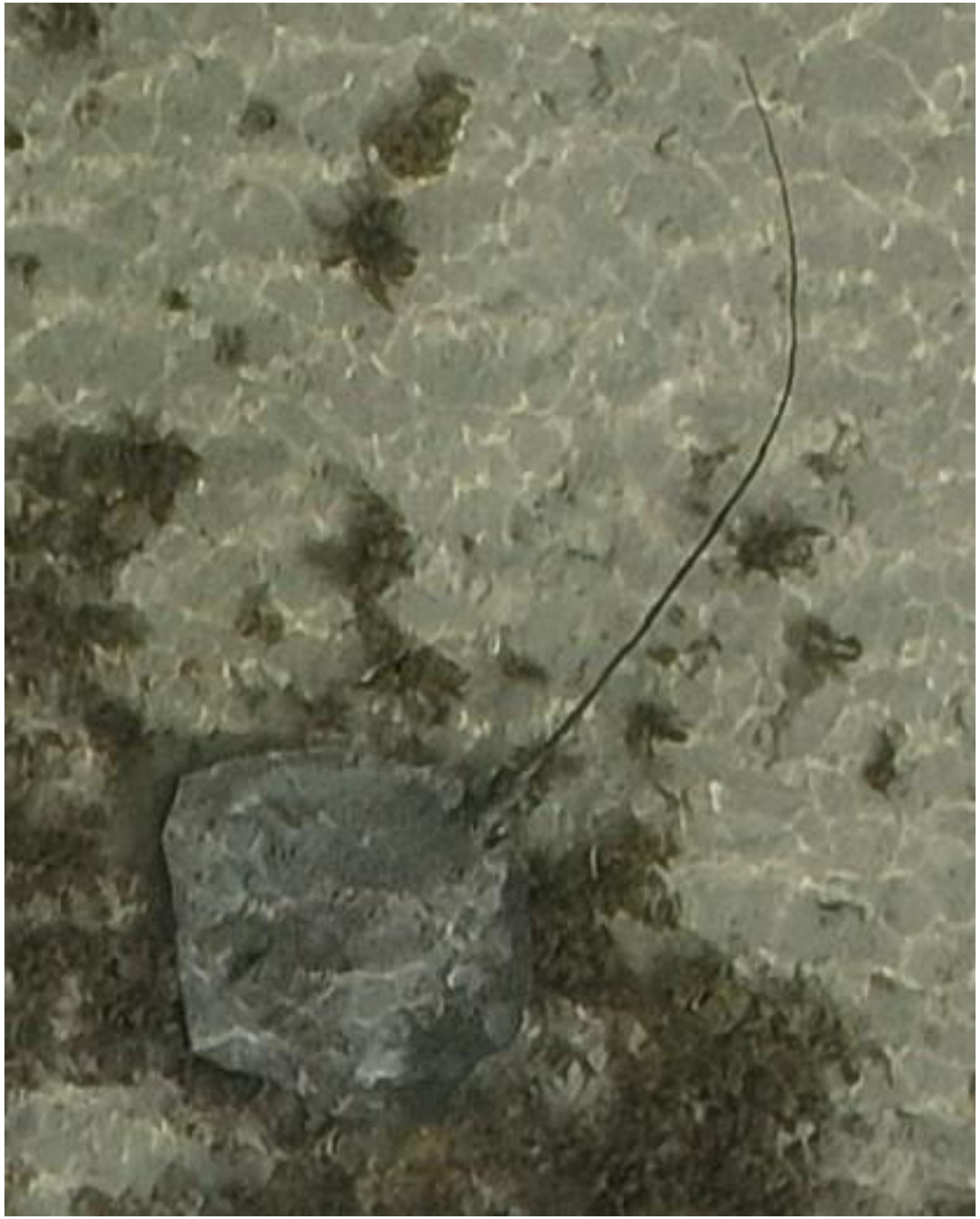

- *Himantura uarnak*

DJI_20240603073019_0282

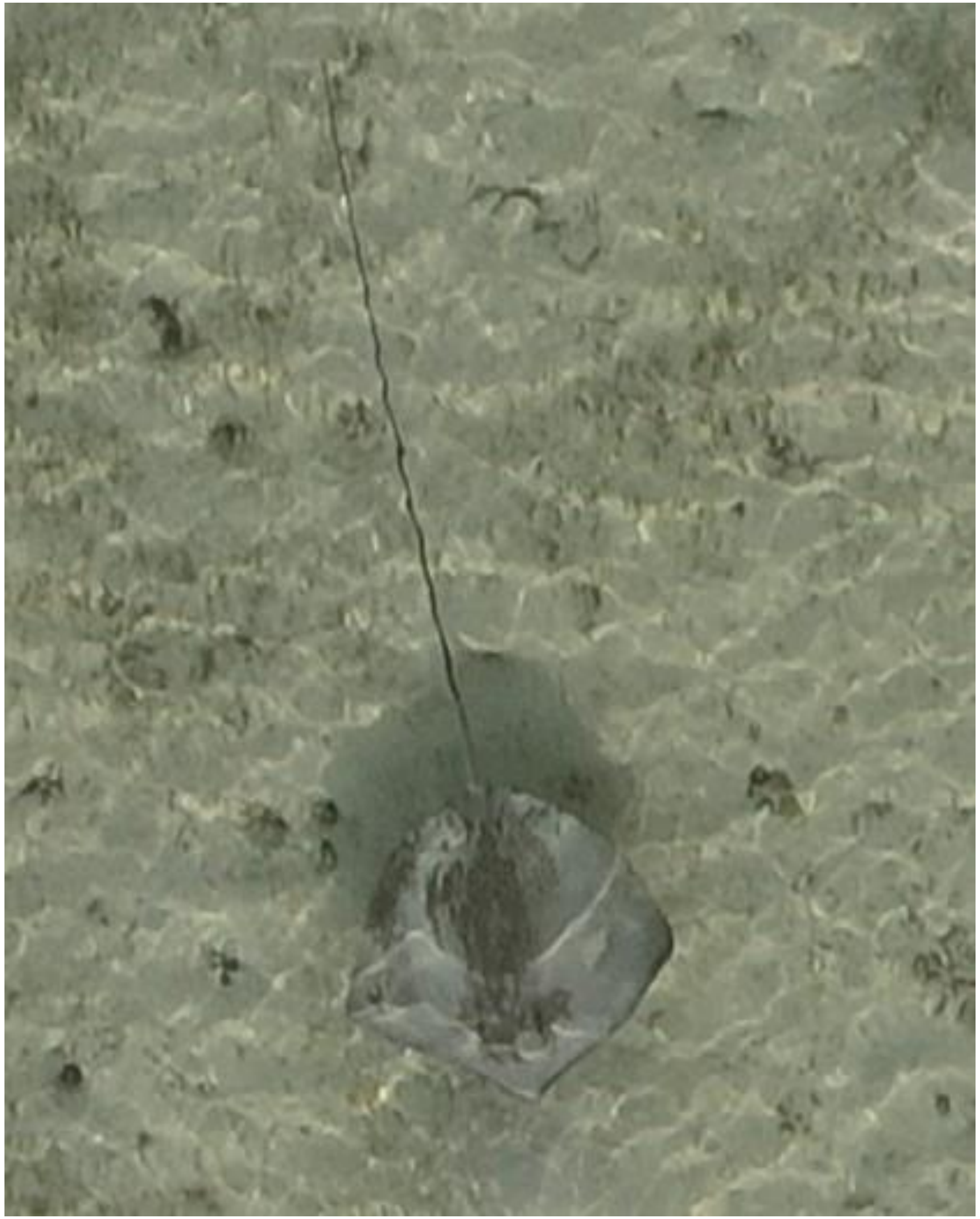

DJI_20240603073022_0285

- *Himantura uarnak*

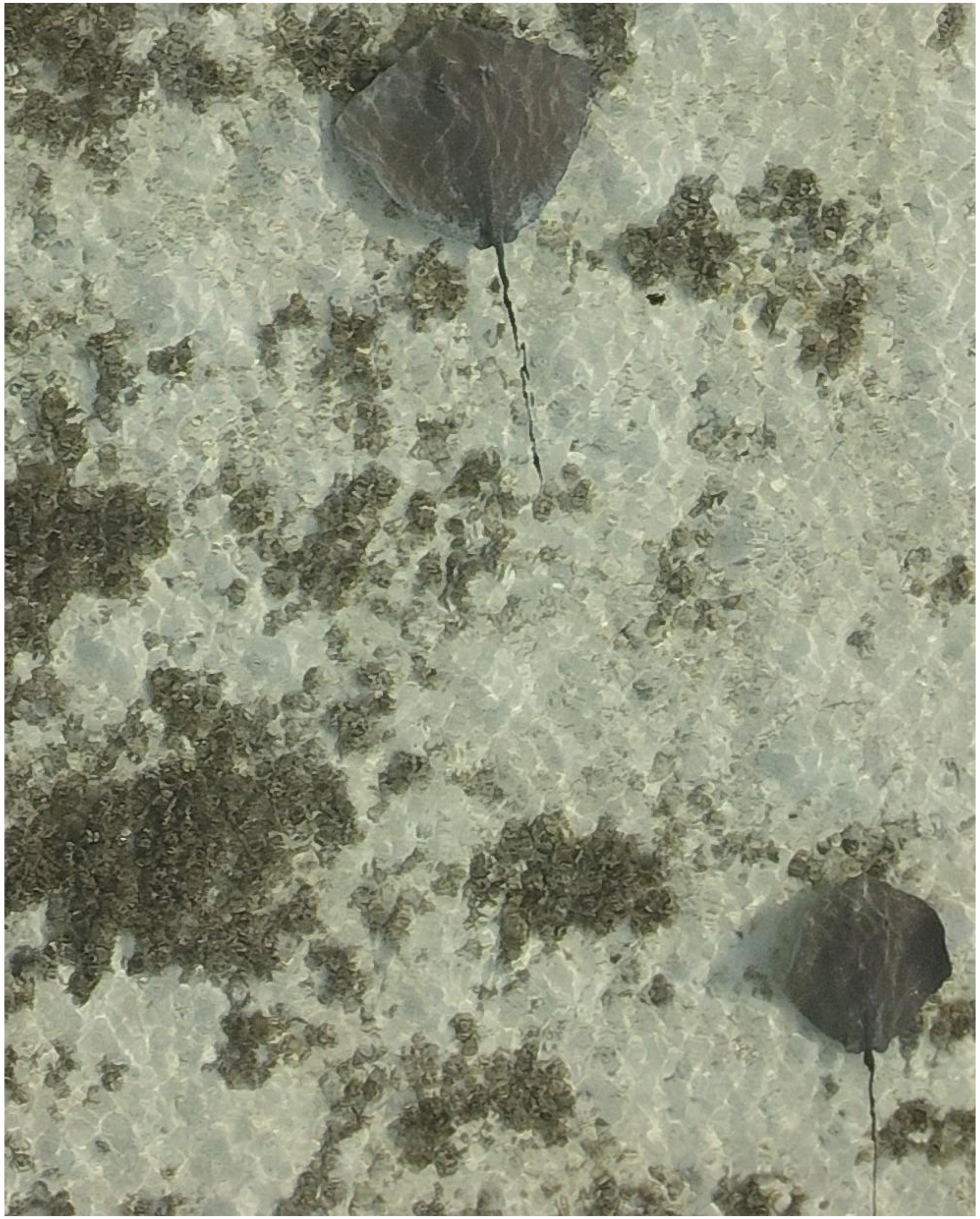

- cf. *Pateobatis fai* (2x)
- Appear to be same duo as was tracked on 05-10.

DJI_20240603073104_0327

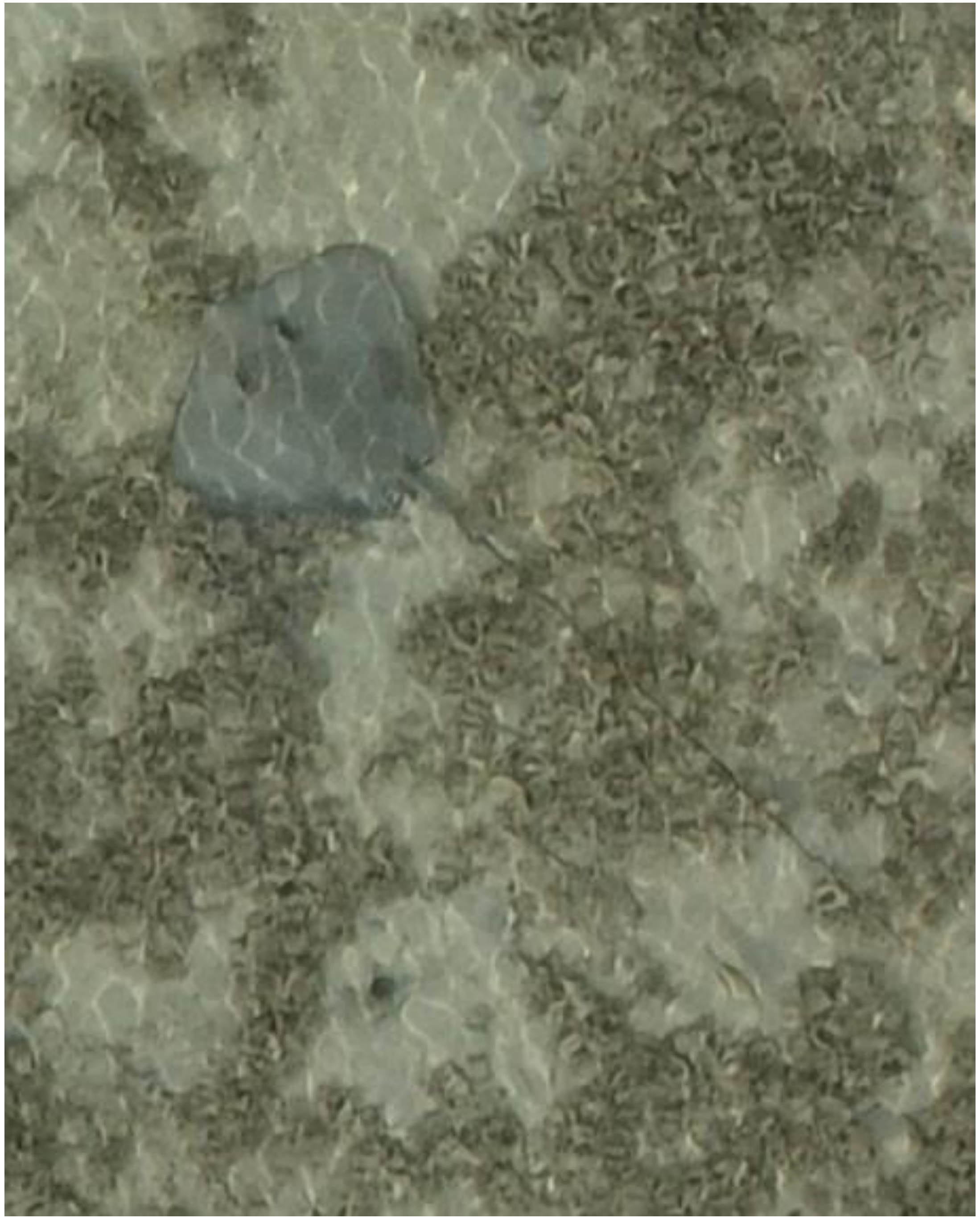

- Whipray

DJI_20240603073549_0572

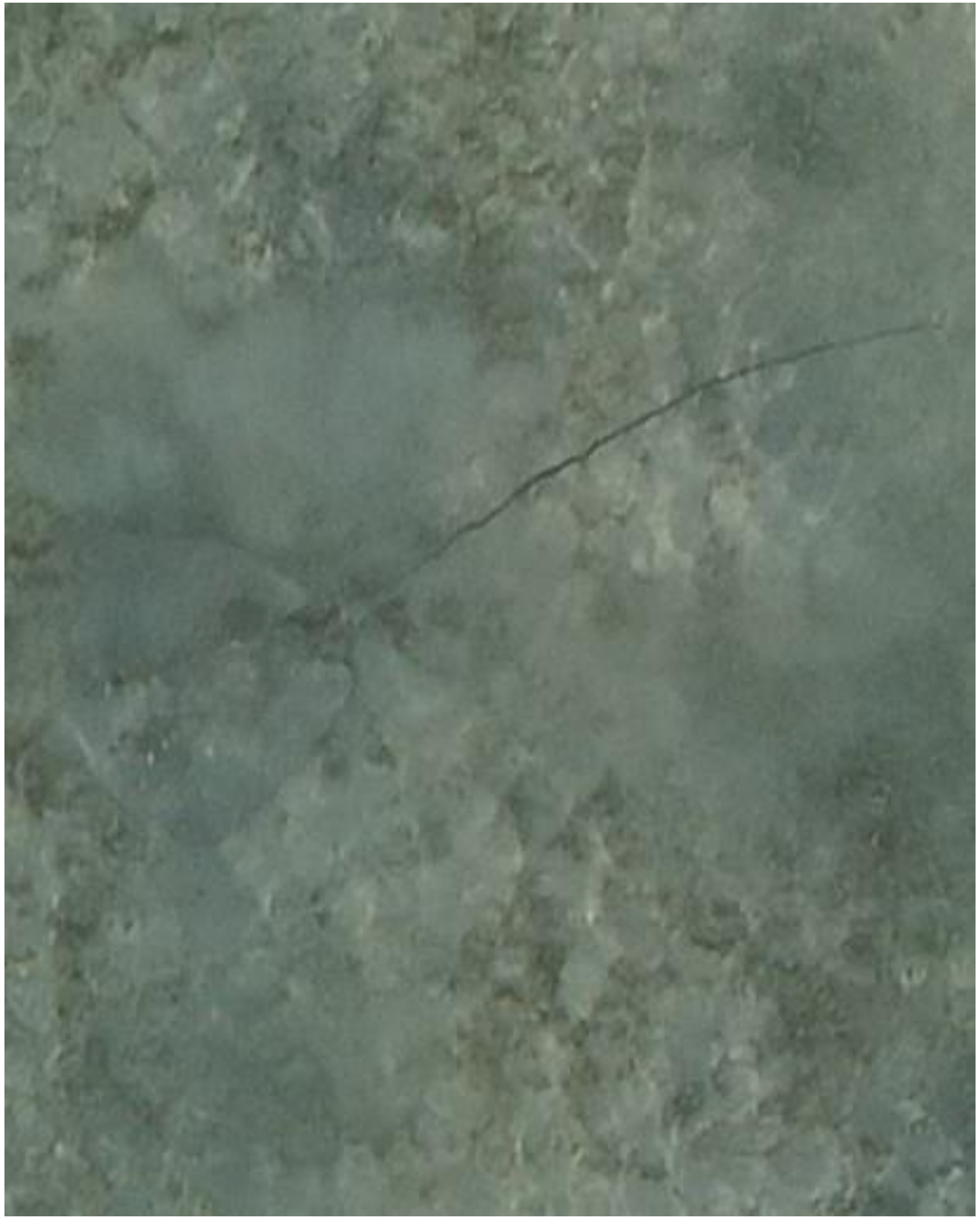

- Whipray

DJI_20240603073950_0784

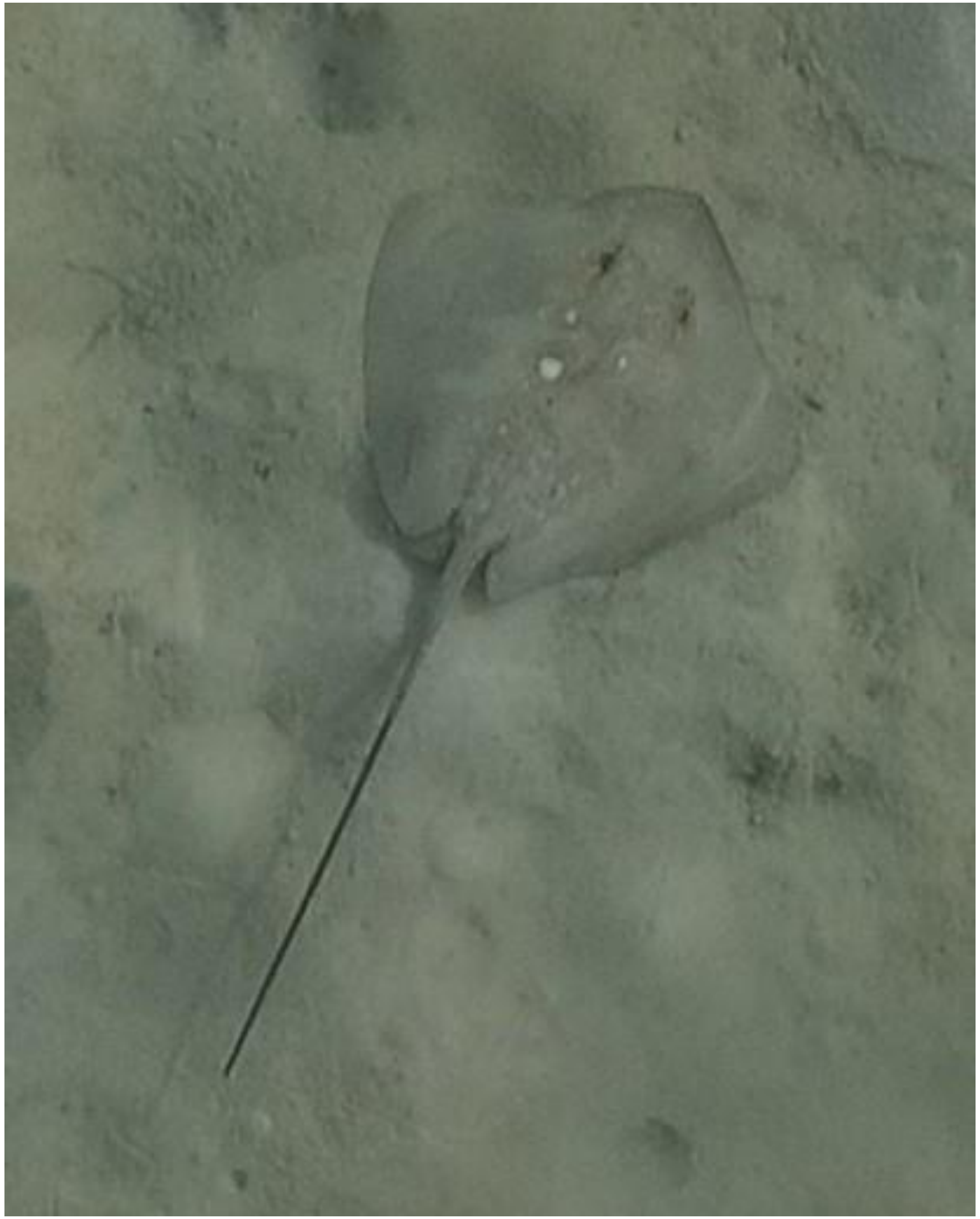

DJI_20240603074238_0940

- Whipray
- Somewhat resembles *Maculabatis randalli*. *T*he same individual was also observed on 09-20.

2024-06-06

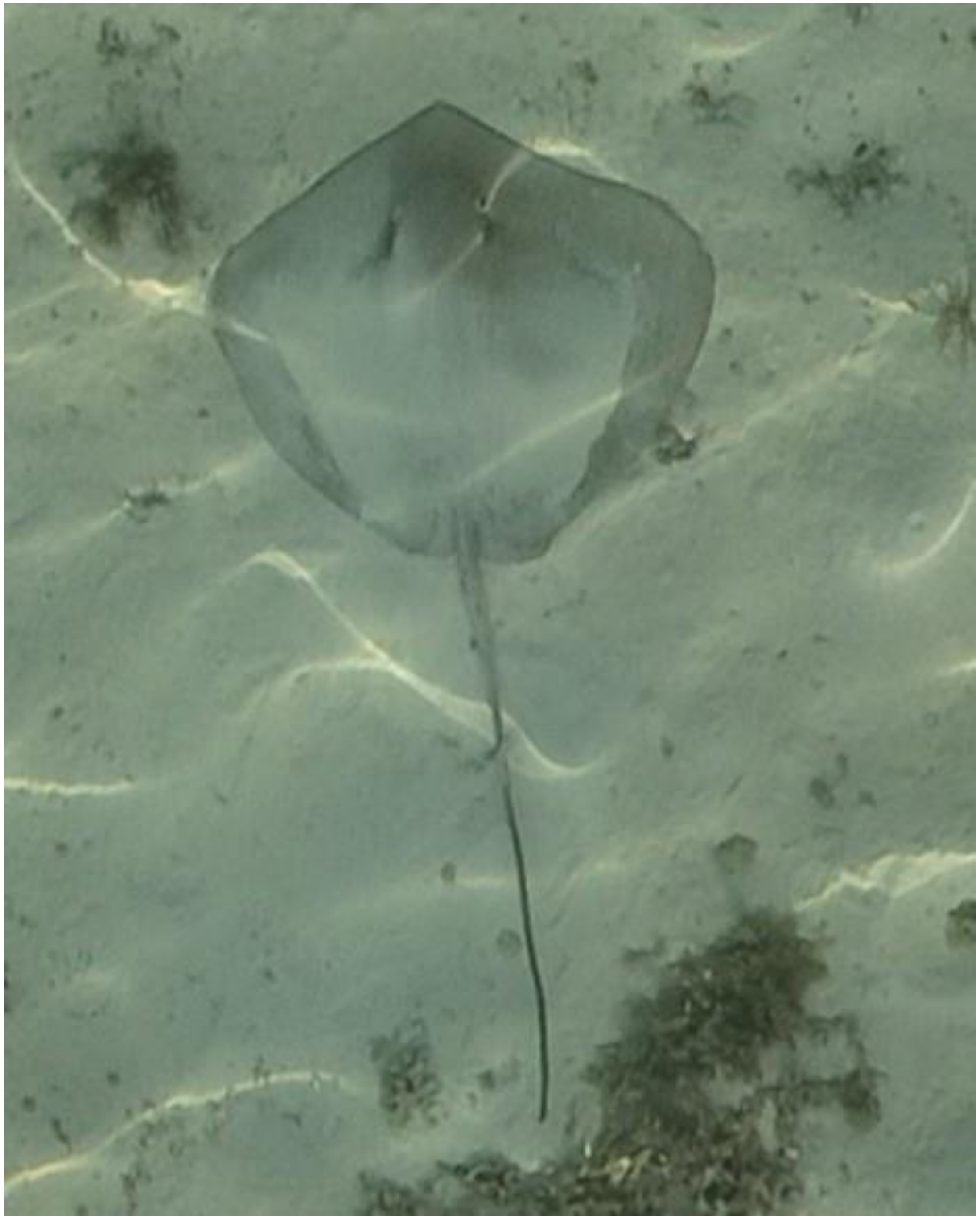

- cf. *Pateobatis fai*

DJI_20240606071952_0282

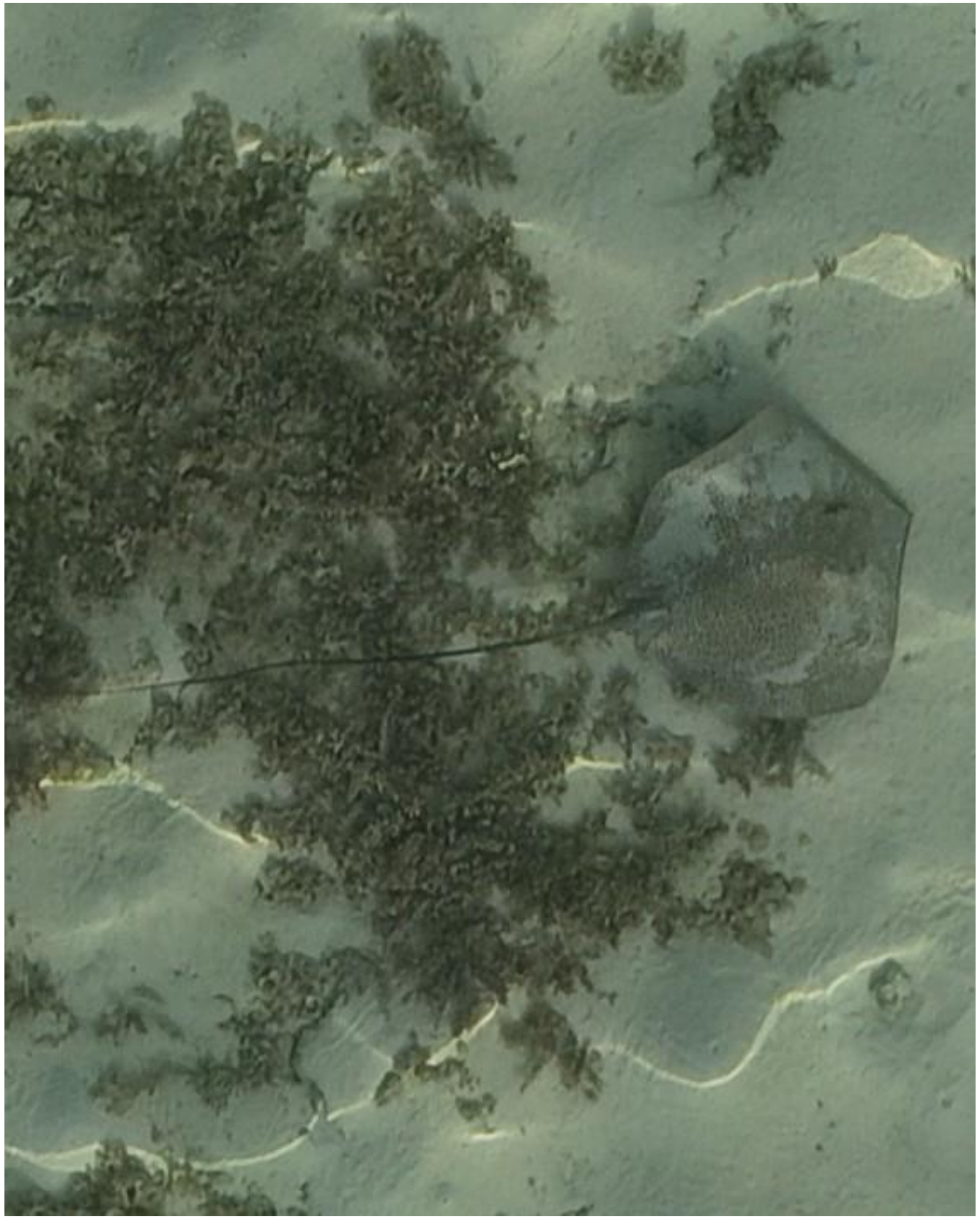
DJI_20240606071952_0282

- *Himantura uarnak*

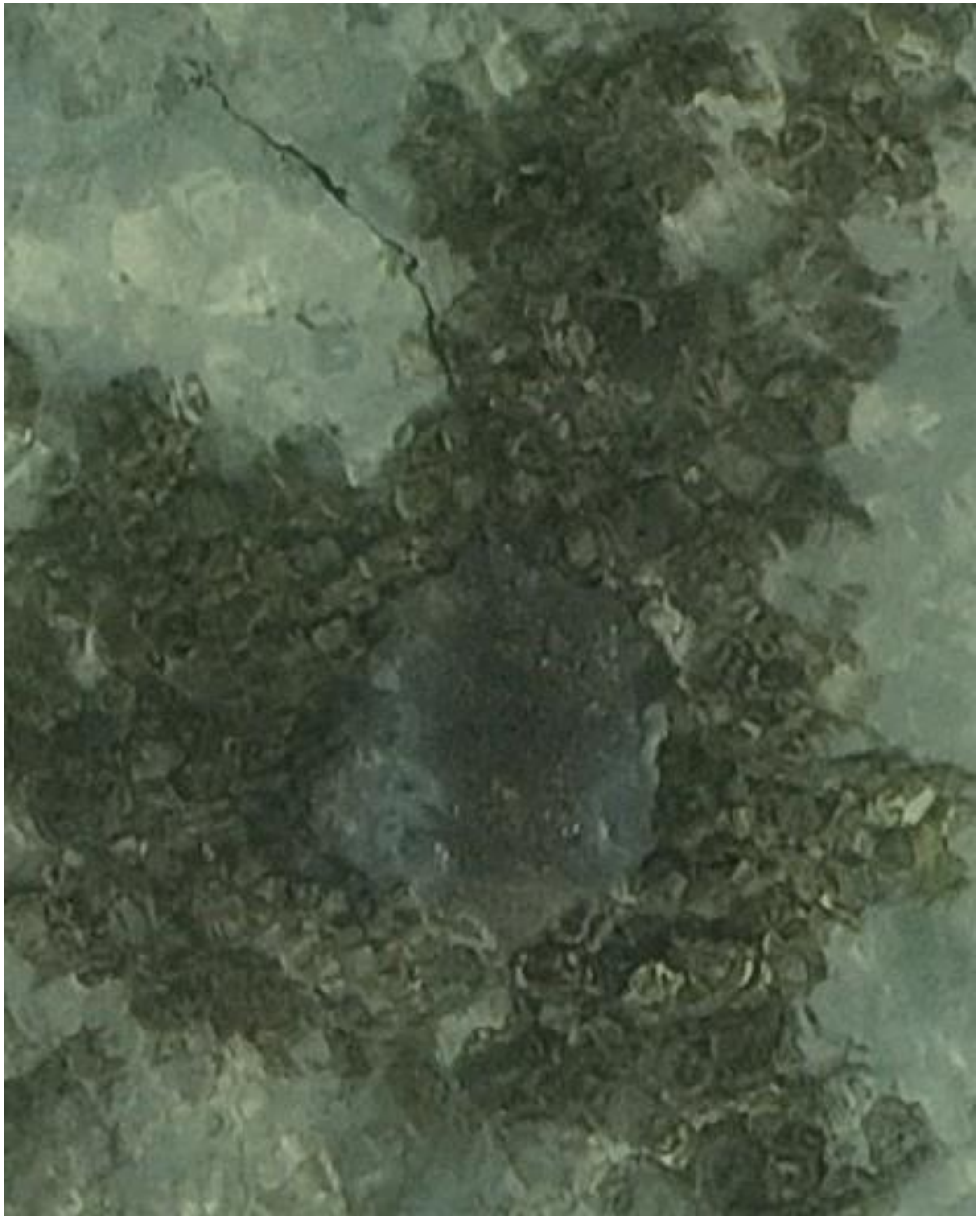

DJI_20240606071959_0289

- Whipray
- Best guess: *H. uarnak*

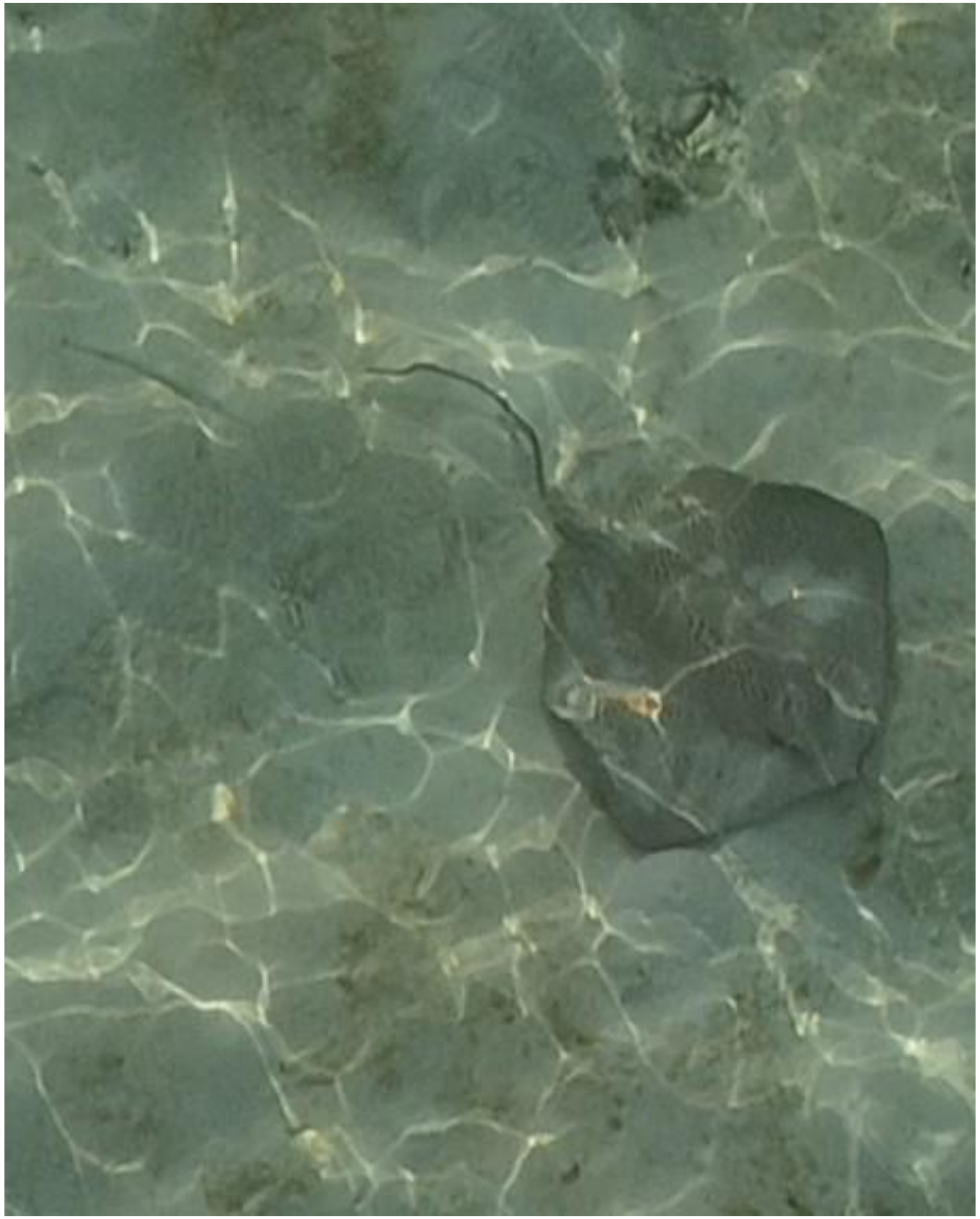

DJI_20240606072232_0430

- *Himantura uarnak*
- Accompanied by needlefish

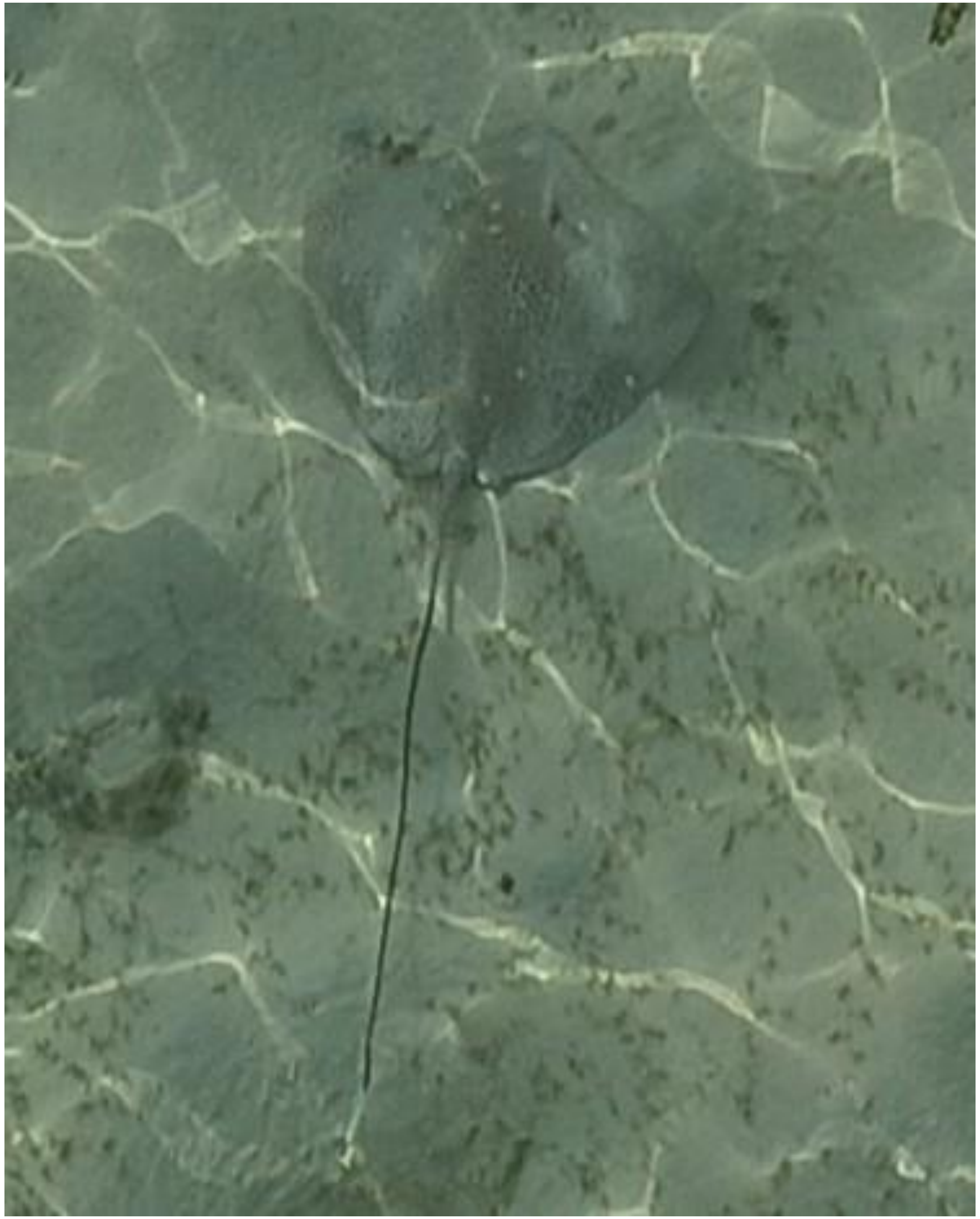
DJI_20240606072241_0439

- *Himantura uarnak*

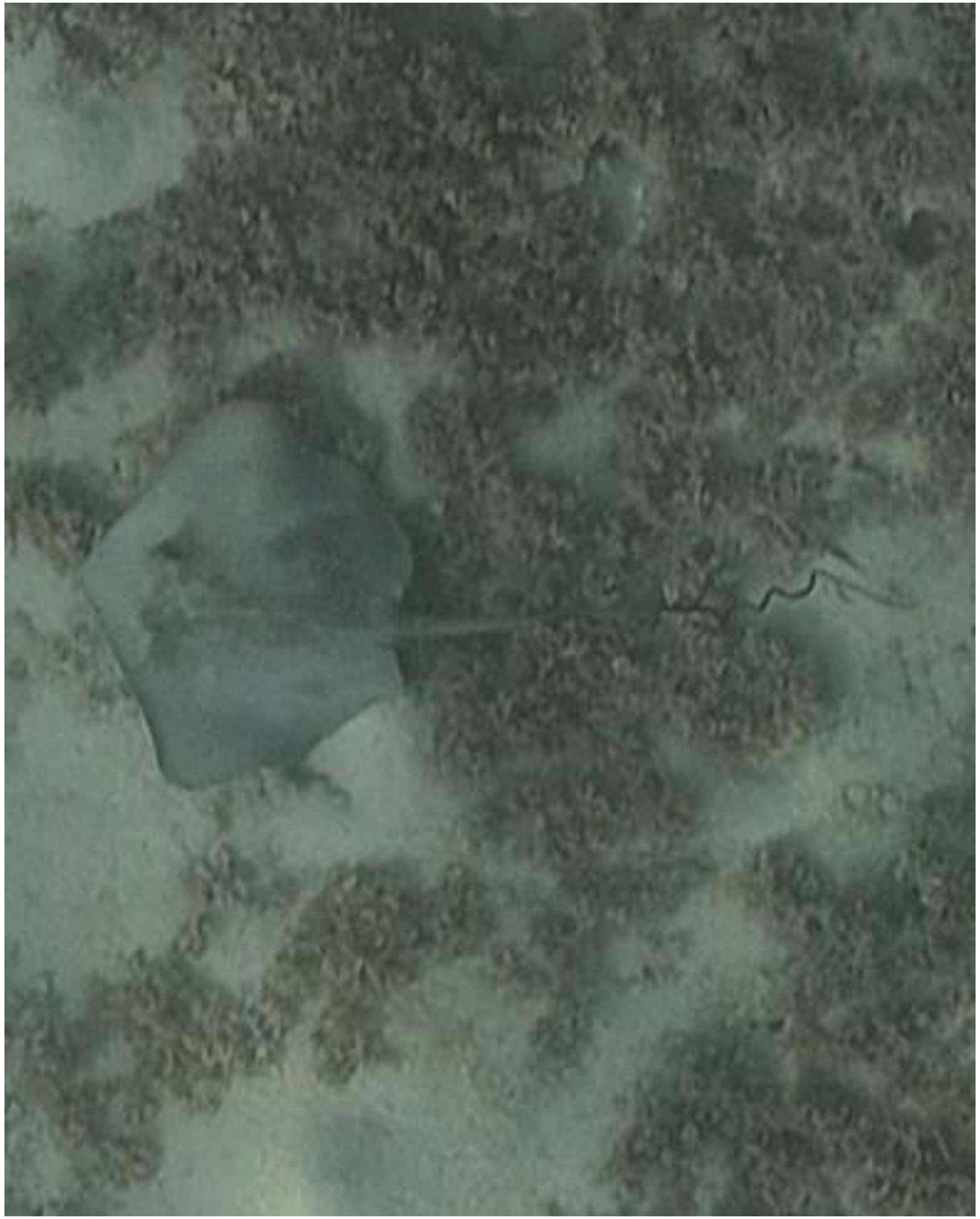

DJI_20240606072539_0588

- Whipray
- Also observed on 05-02, 06-24, 07-29, and 09-20 as identified by the coiled tail.
- Best guess: *H. uarnak*

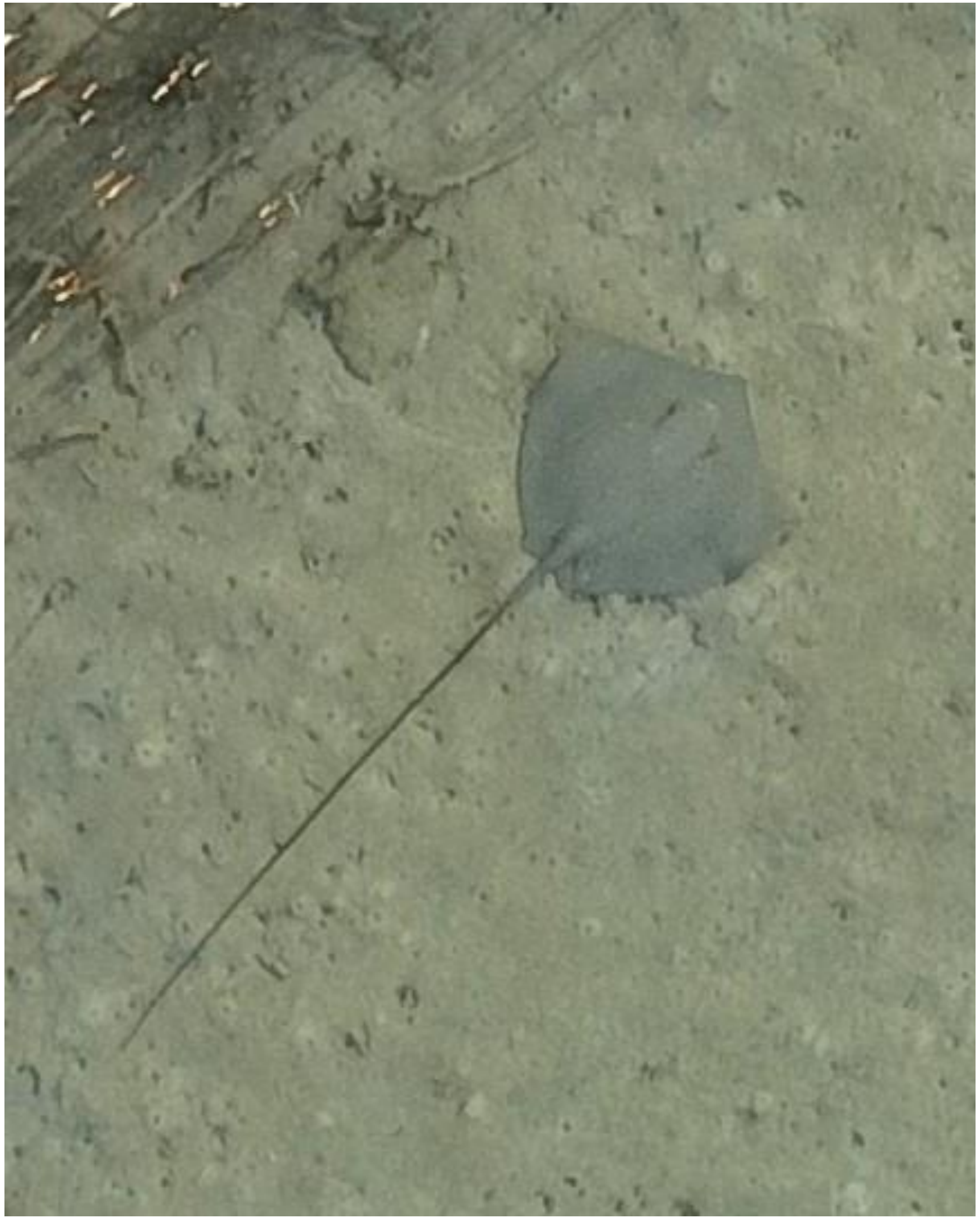
DJI_20240606072814_0727

- Whipray
- Best guess: *H. uarnak*

2024-06-10

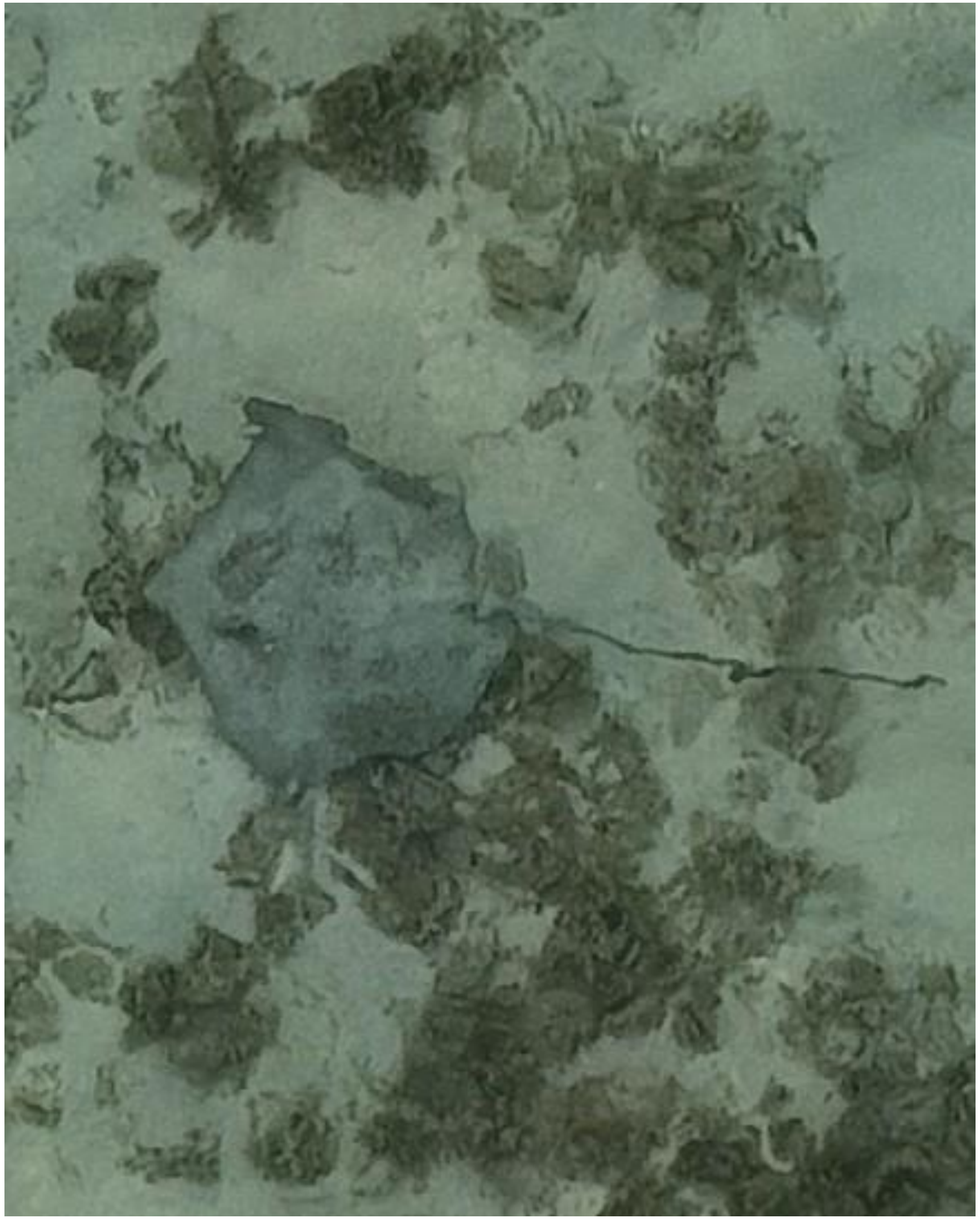

- Whipray

DJI_20240610064232_0039

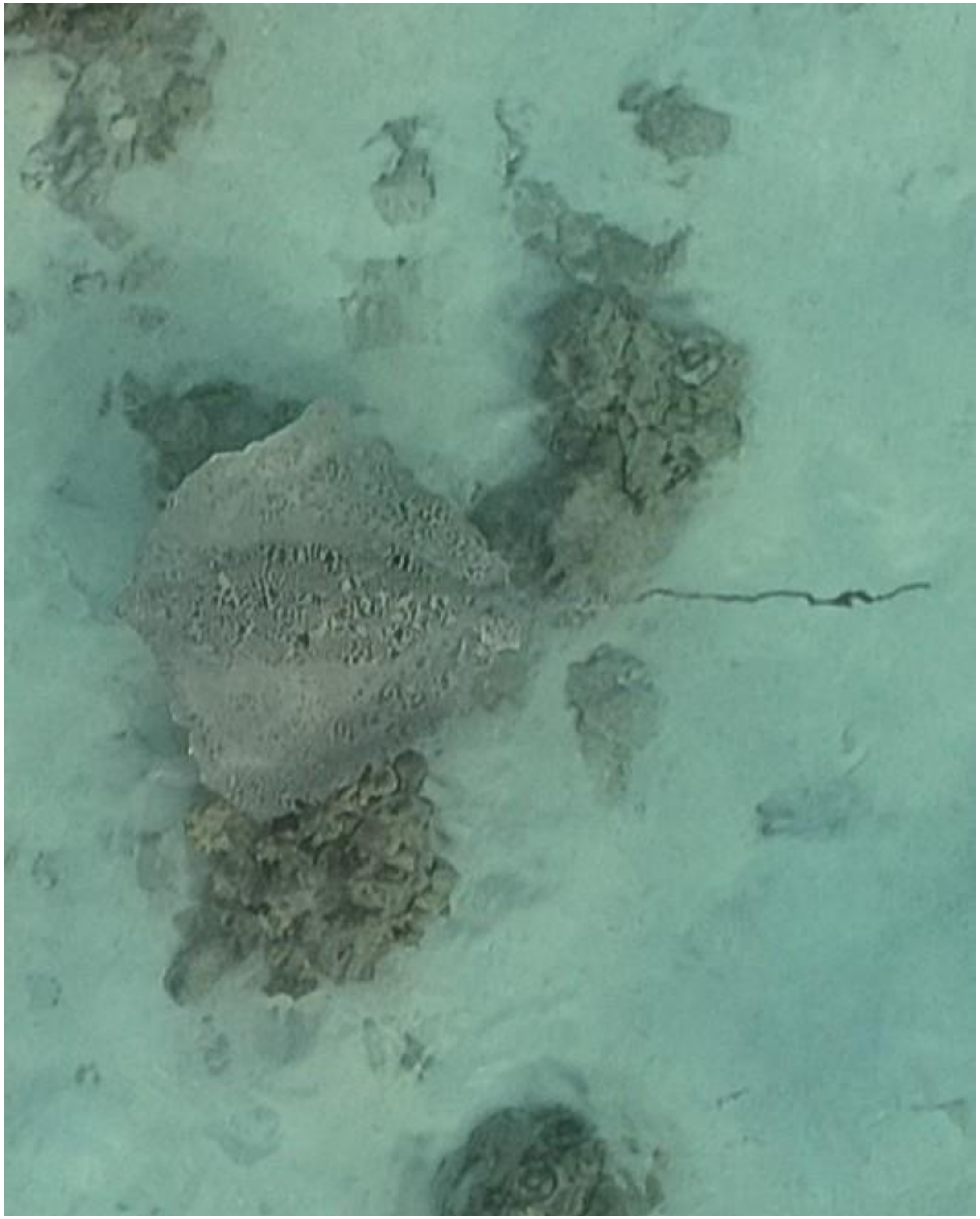

DJI_20240610064326_0081

- *Himantura uarnak*

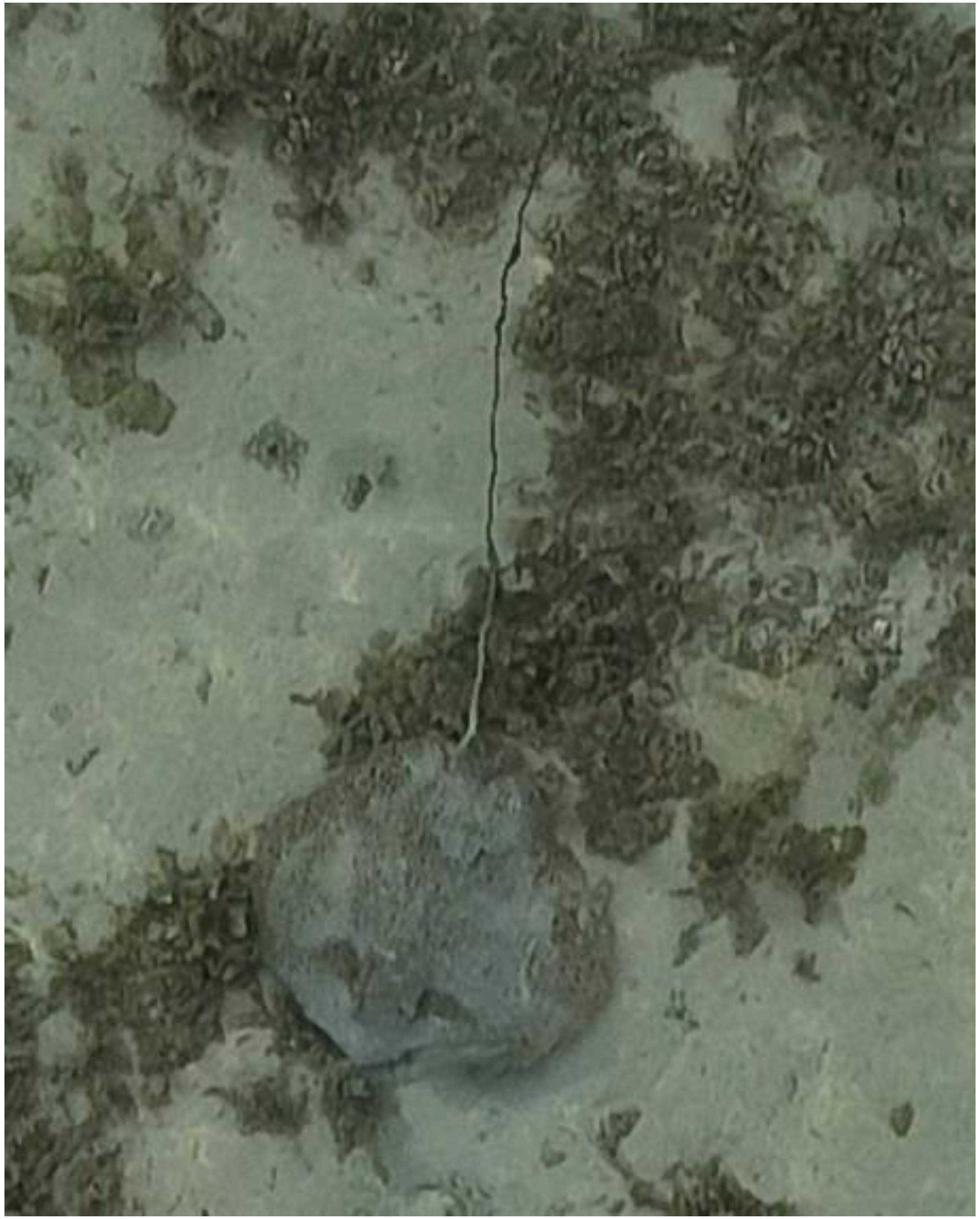

DJI_20240610064542_0206

- *Himantura uarnak*

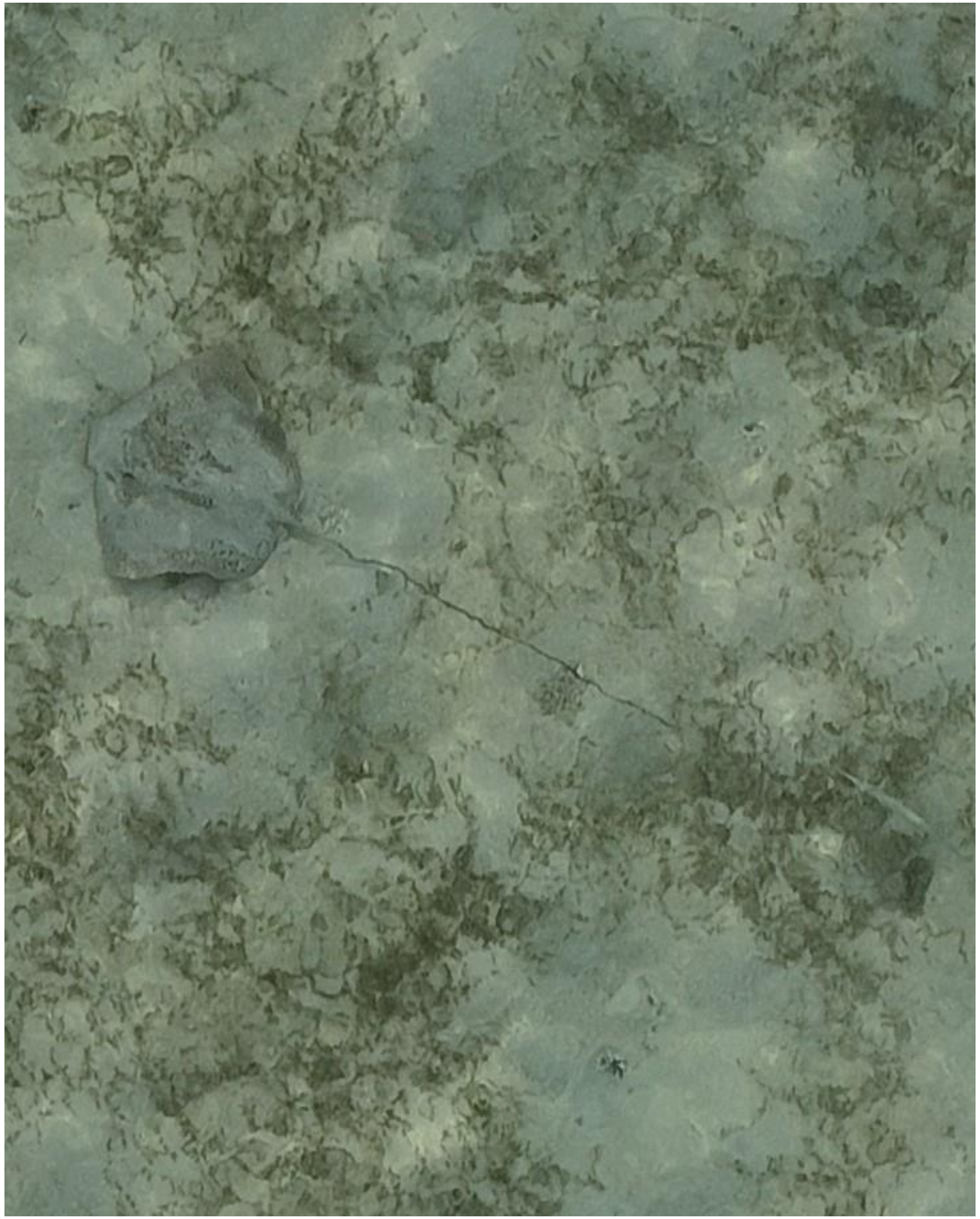

- *Himantura uarnak*
- Accompanied by needlefish

DJI_20240610064942_0422

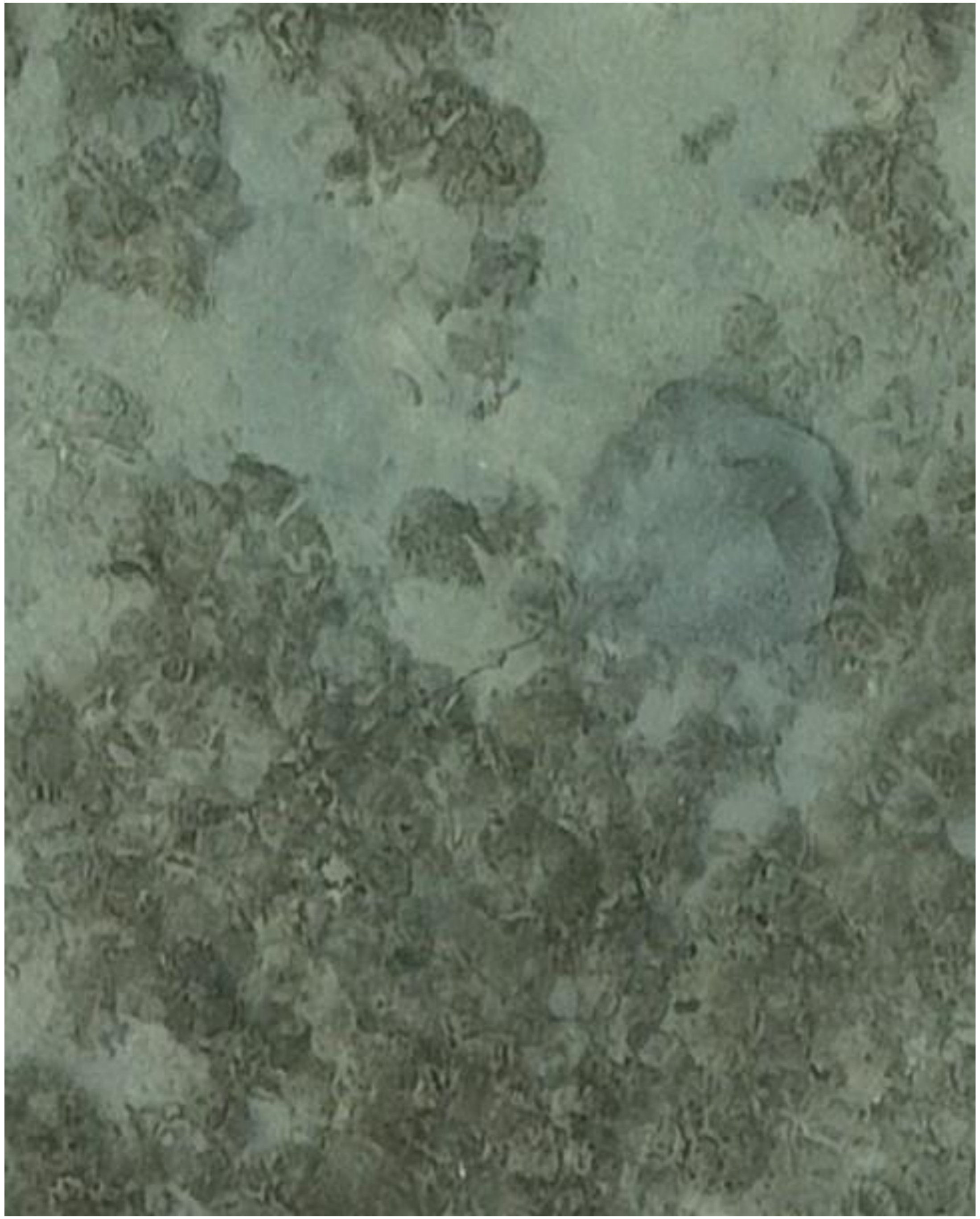

DJI_20240610065241_0572

- Whipray
- Best guess: *H. uarnak*

2024-05-11

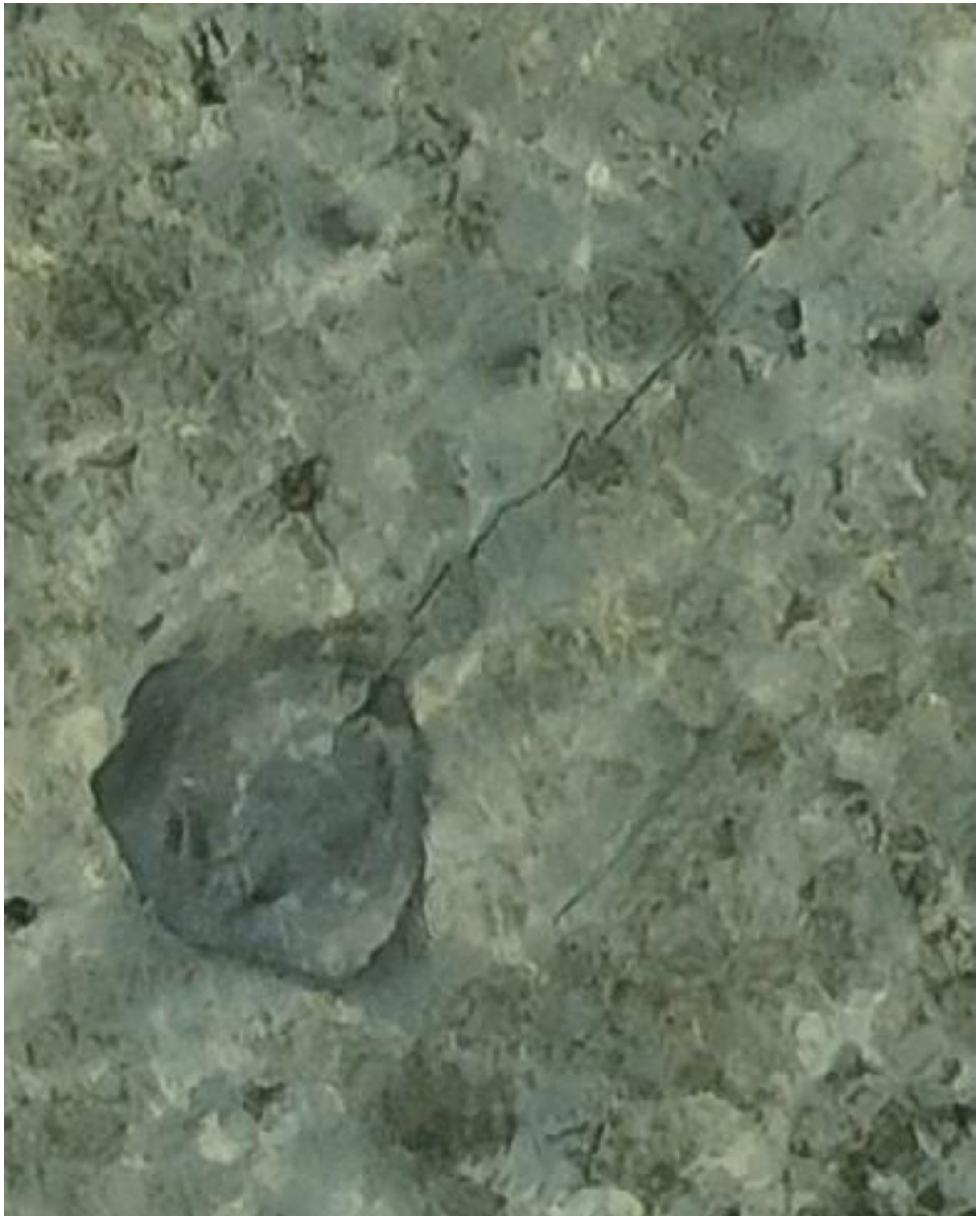

DJI_20240611070213_0054

- Whipray
- Accompanied by needlefish

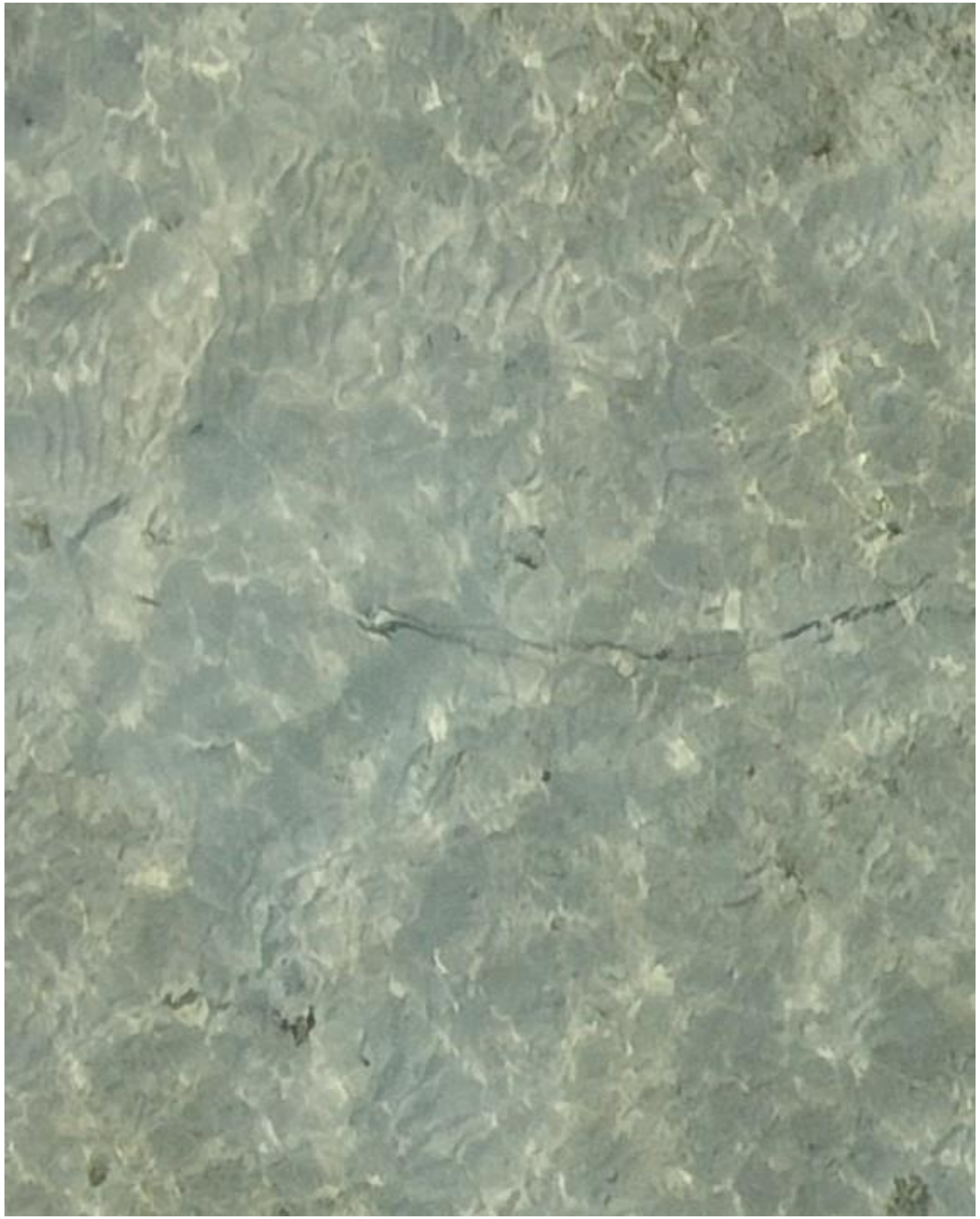

DJI_20240611070216_0057

- Whipray

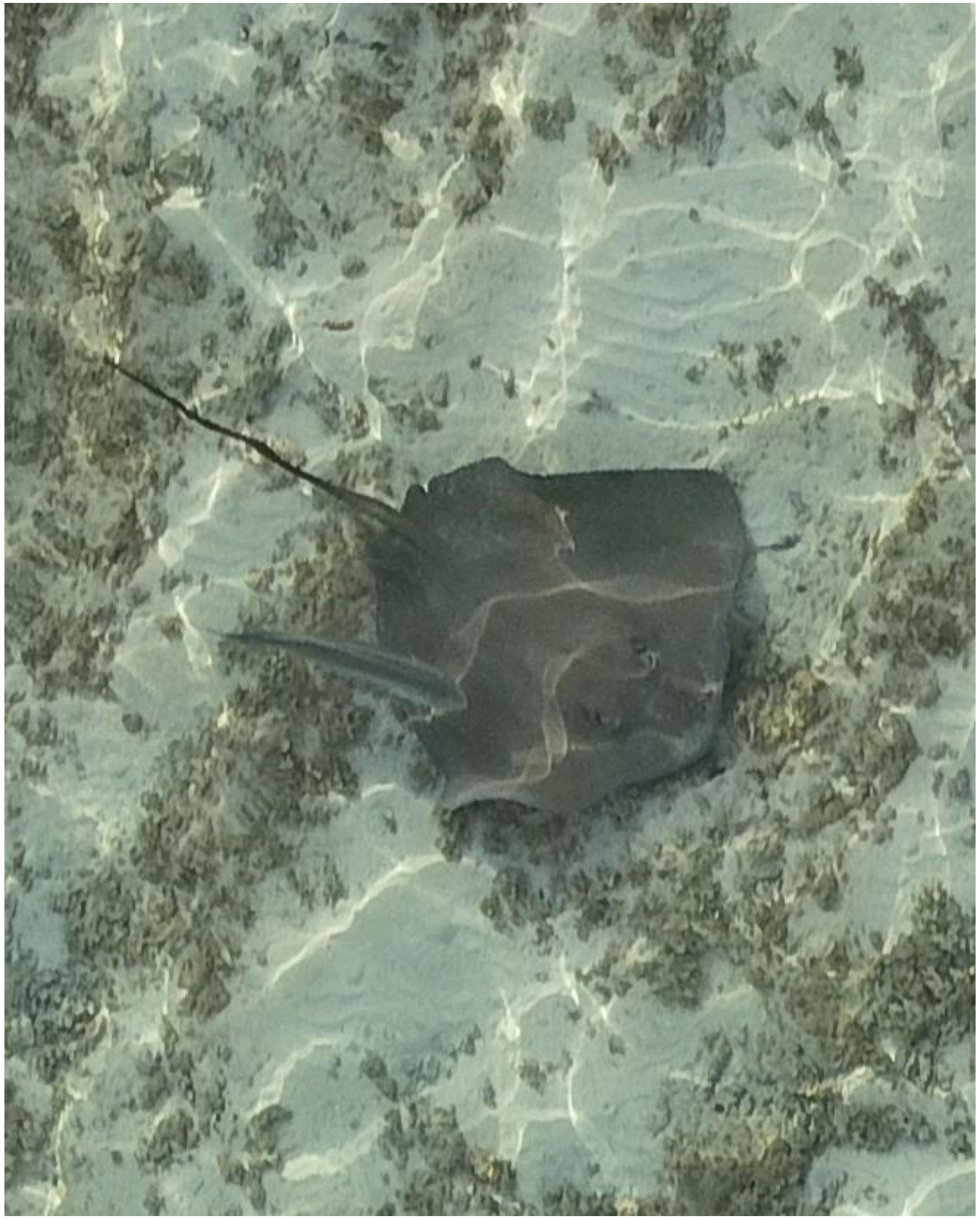

DJI_20240611070436_0174

- cf. *Pateobatis fai*
- Accompanied by trevally
- Resembles the ray tracked on 05-10.

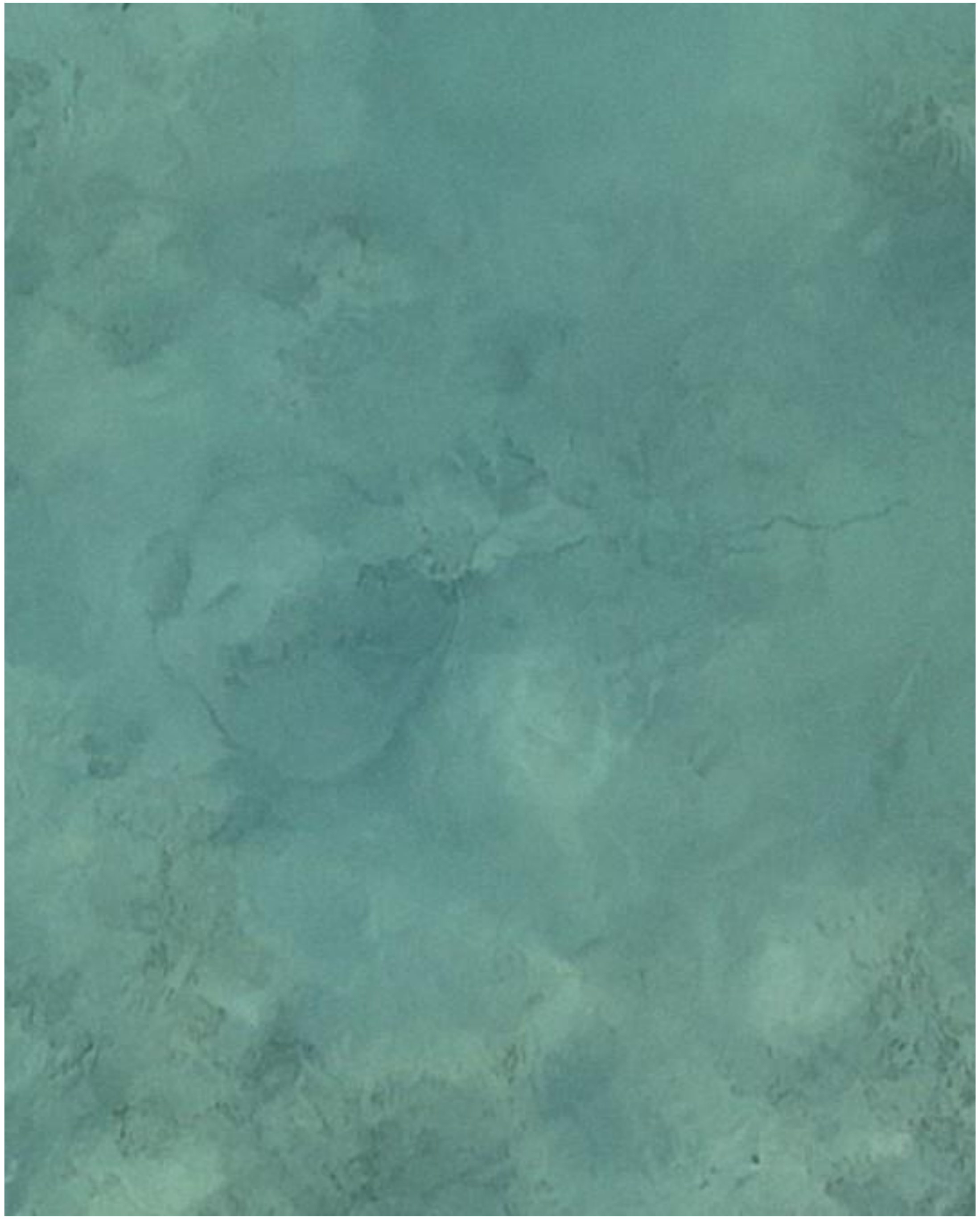

DJI_20240611070533_0231

- Whipray

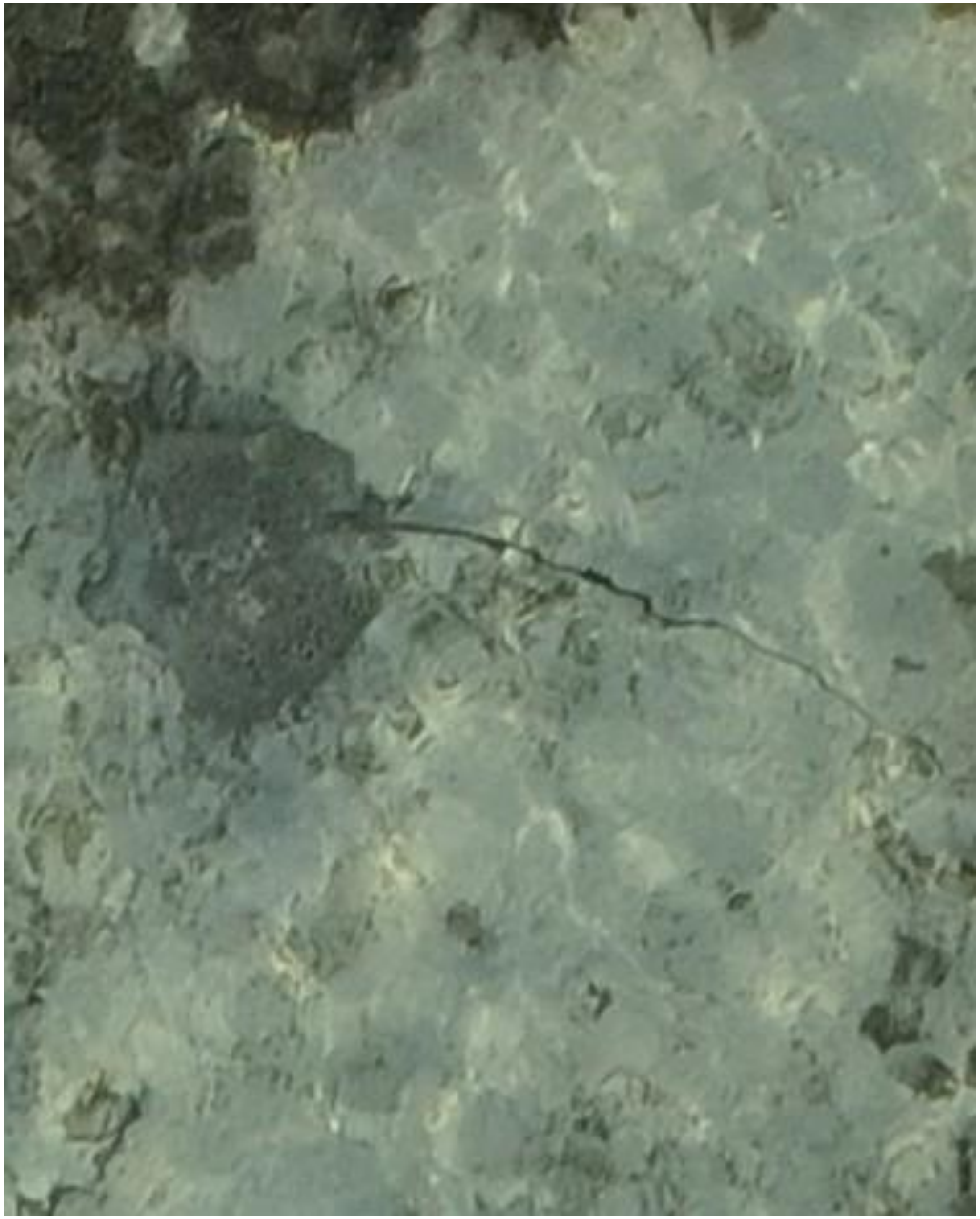

DJI_20240611070634_0280

- *Himantura uarnak*

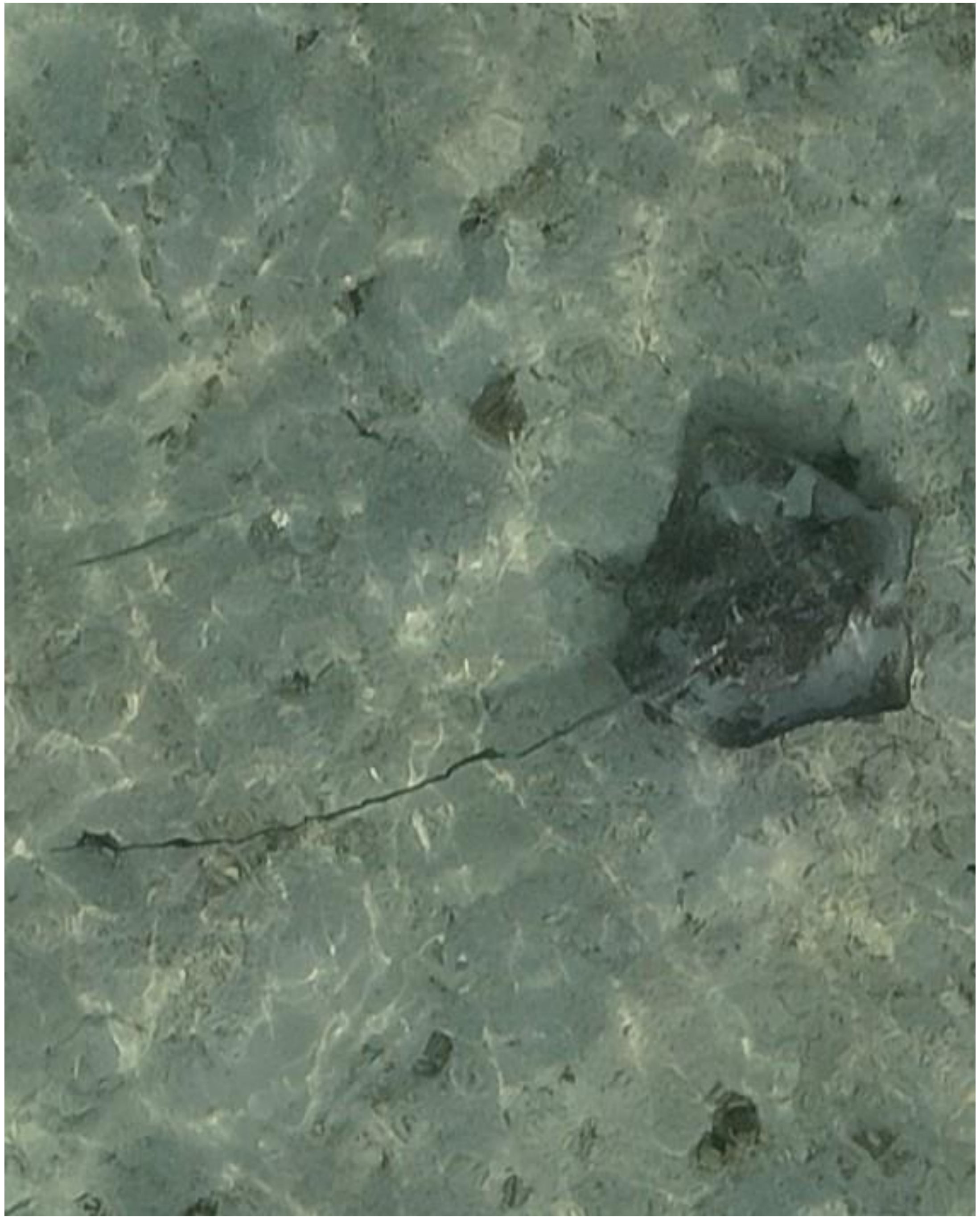

DJI_20240611070702_0308

- *Himantura uarnak*
- Accompanied by needlefish

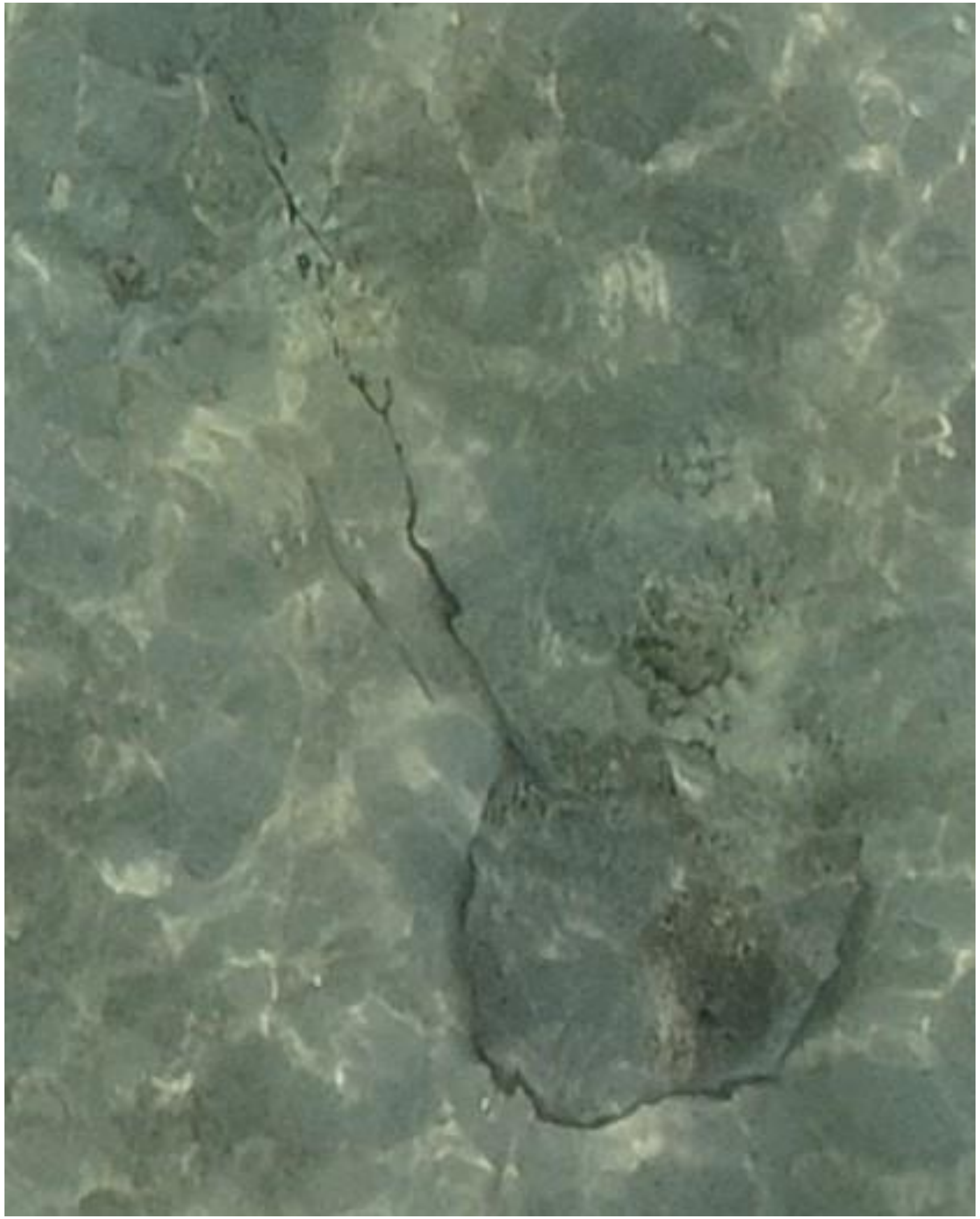

DJI_20240611070916_0430

- *Himantura uarnak*
- Accompanied by needlefish

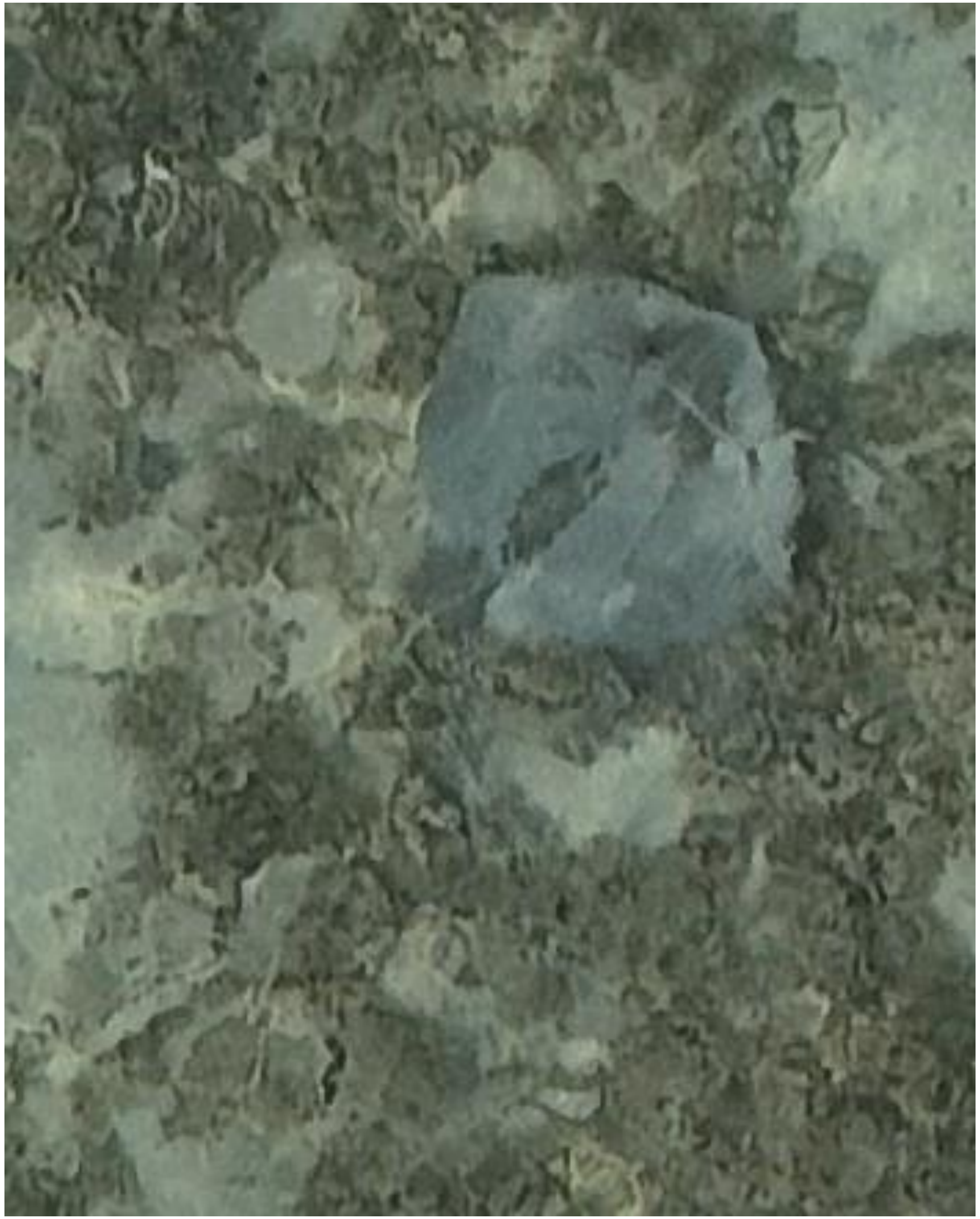

DJI_20240611071214_0578

- Whipray

2024-06-17

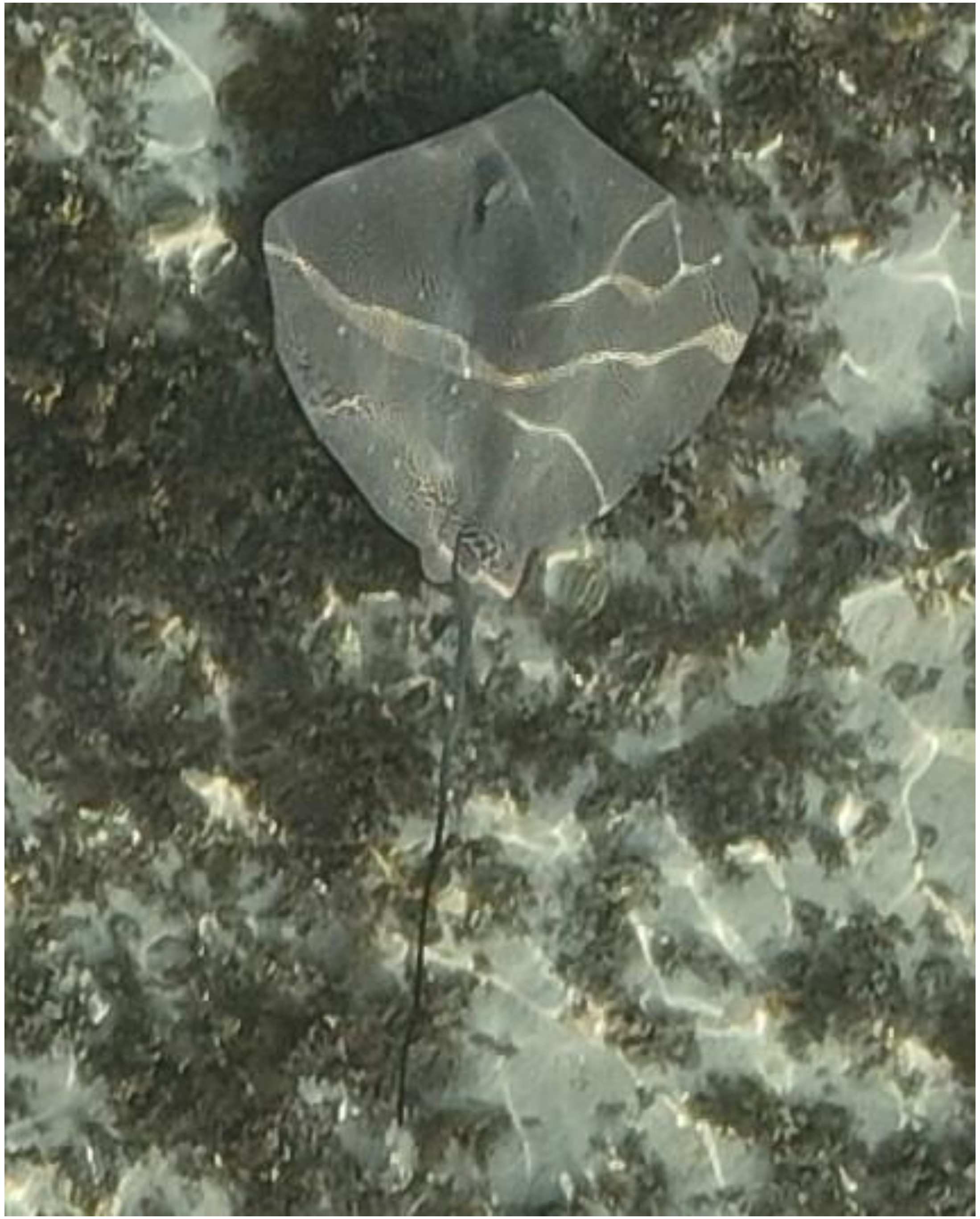

DJI_20240617071650_0141

- *Himantura uarnak*

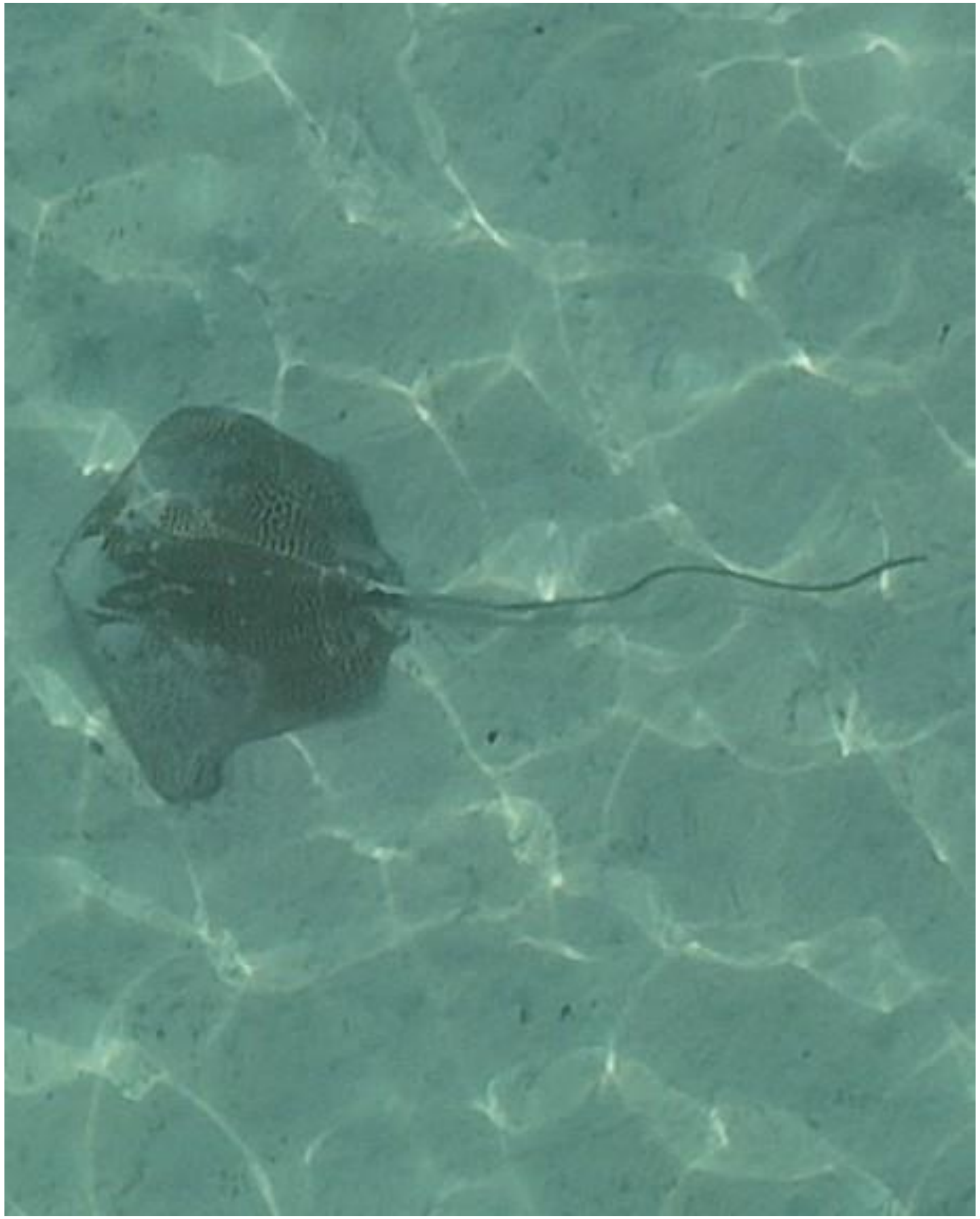

- *Himantura uarnak*

DJI_20240617071833_0232

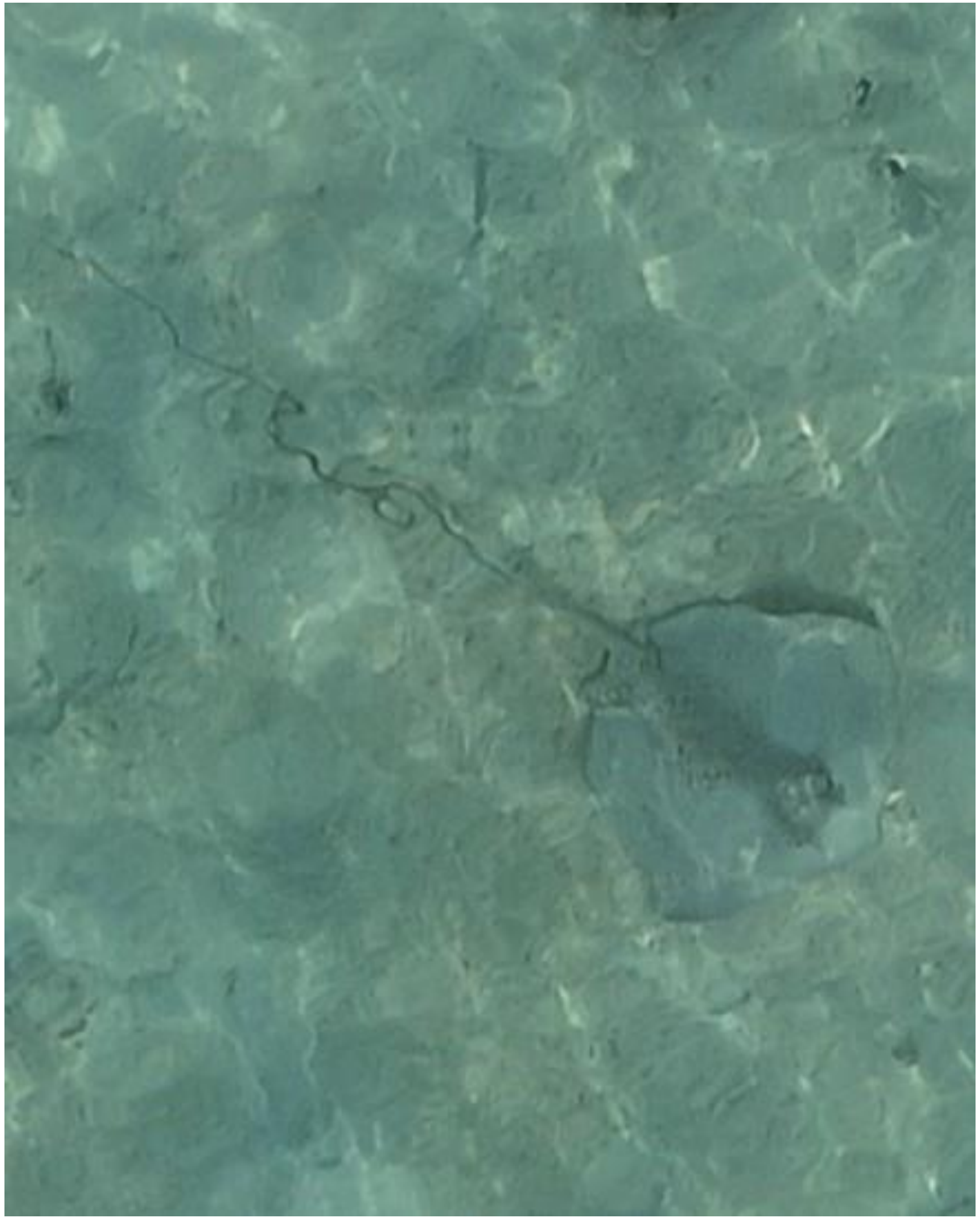

DJI_20240617071921_0268

- Whipray
- Best guess: *H. uarnak*

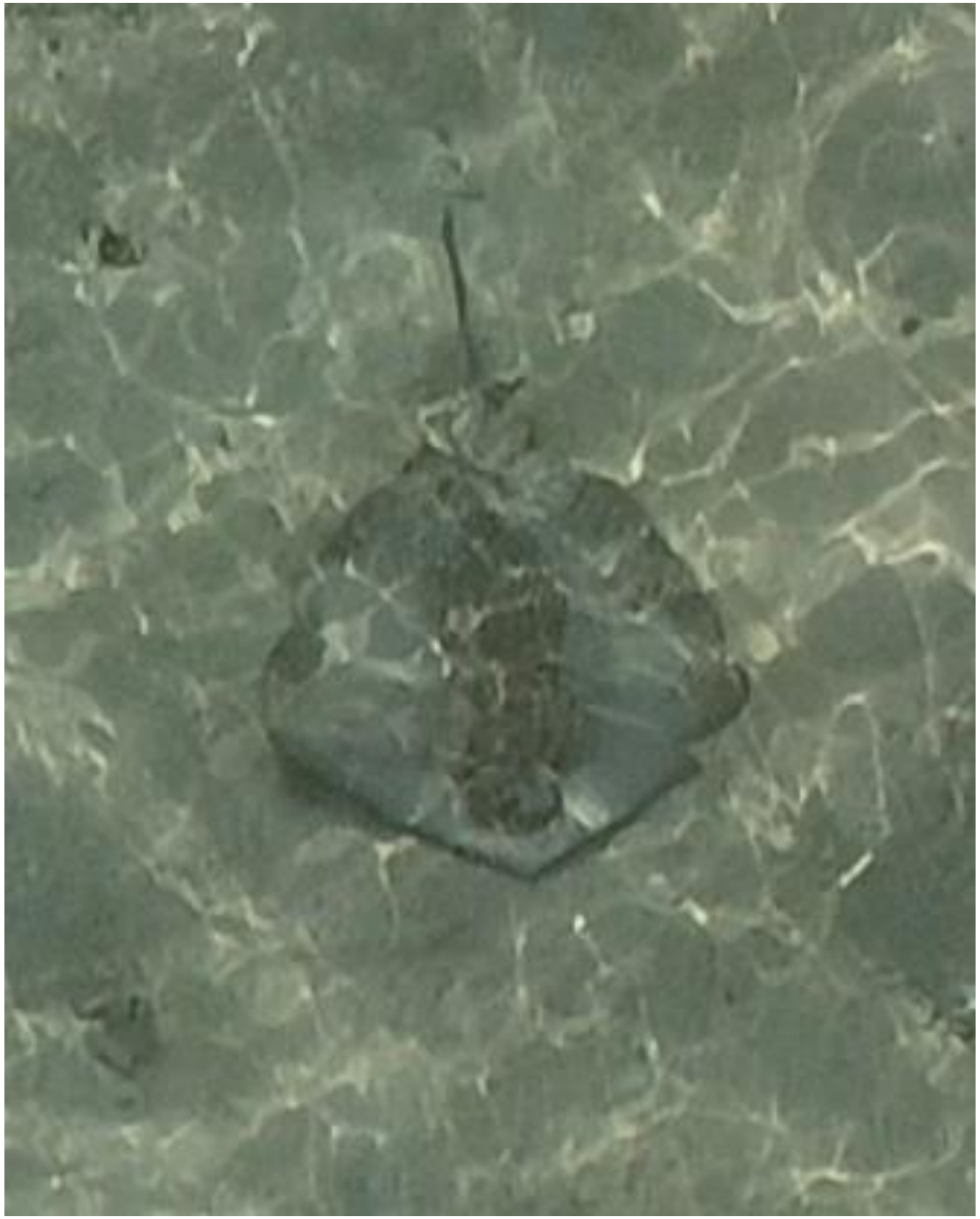

- Whipray

DJI_20240617072214_0429

2024-06-18

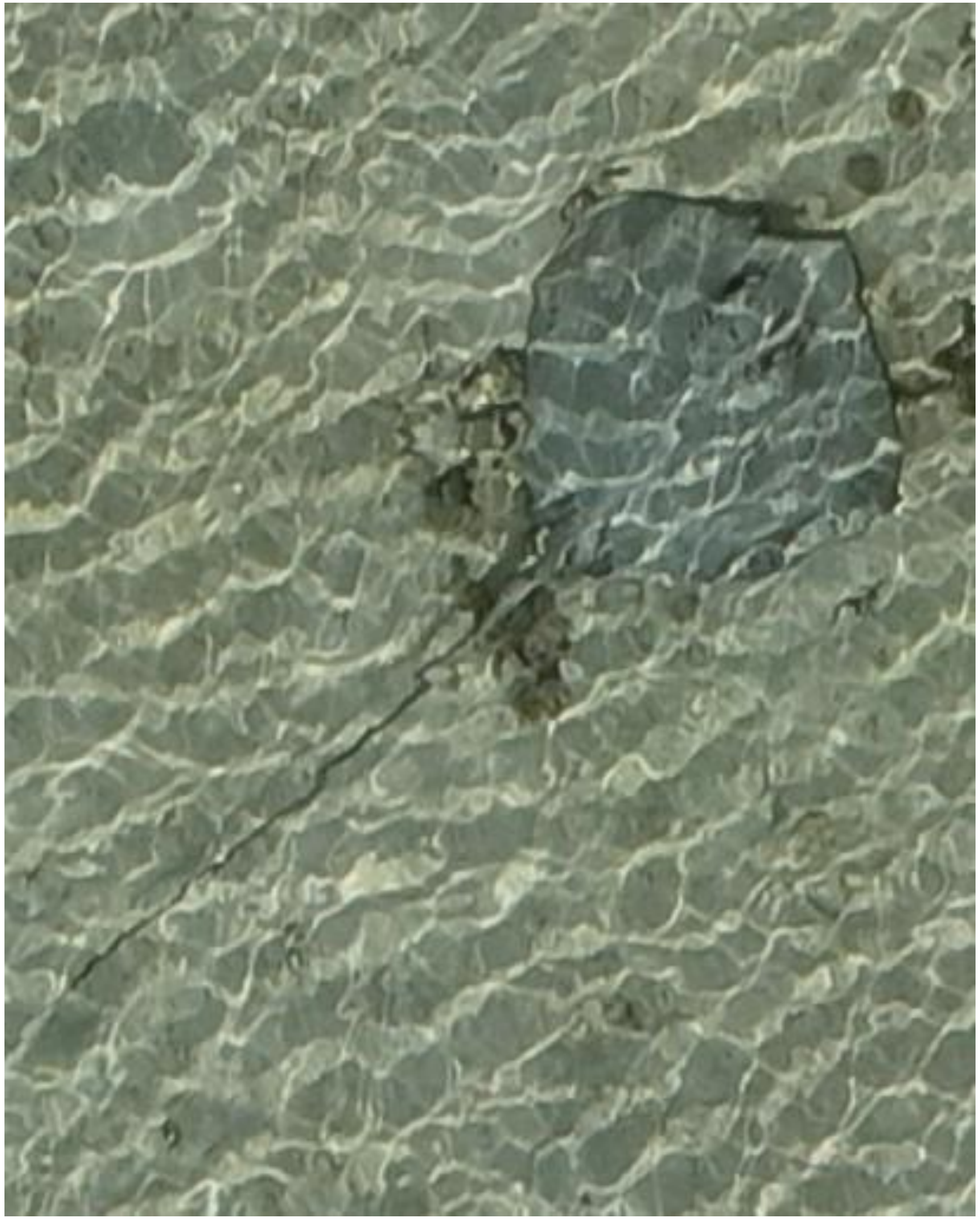

DJI_20240618084734_0131

- Whipray

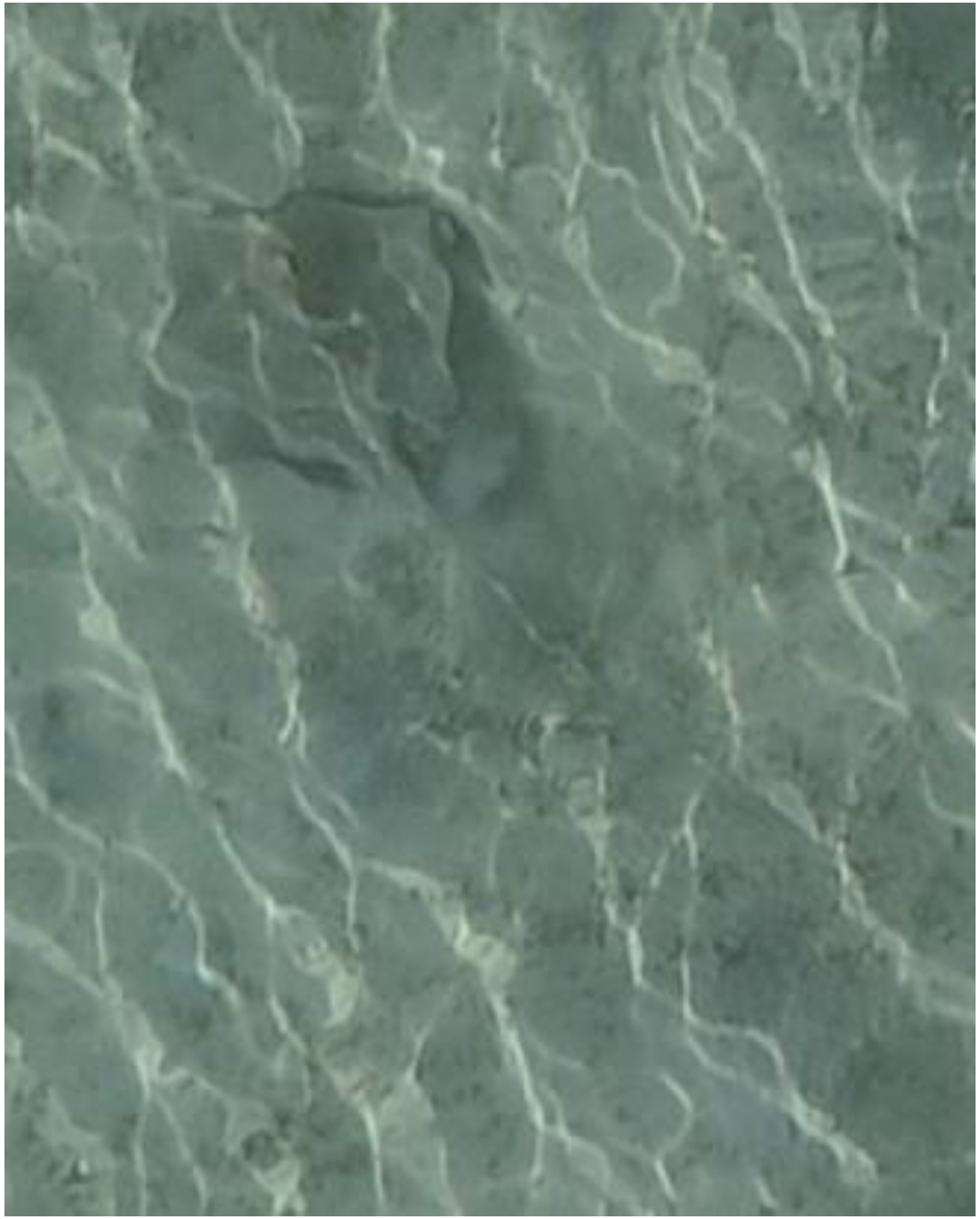

DJI_20240618085958_0781

- Whipray
- Best guess: cf. *P. fai*

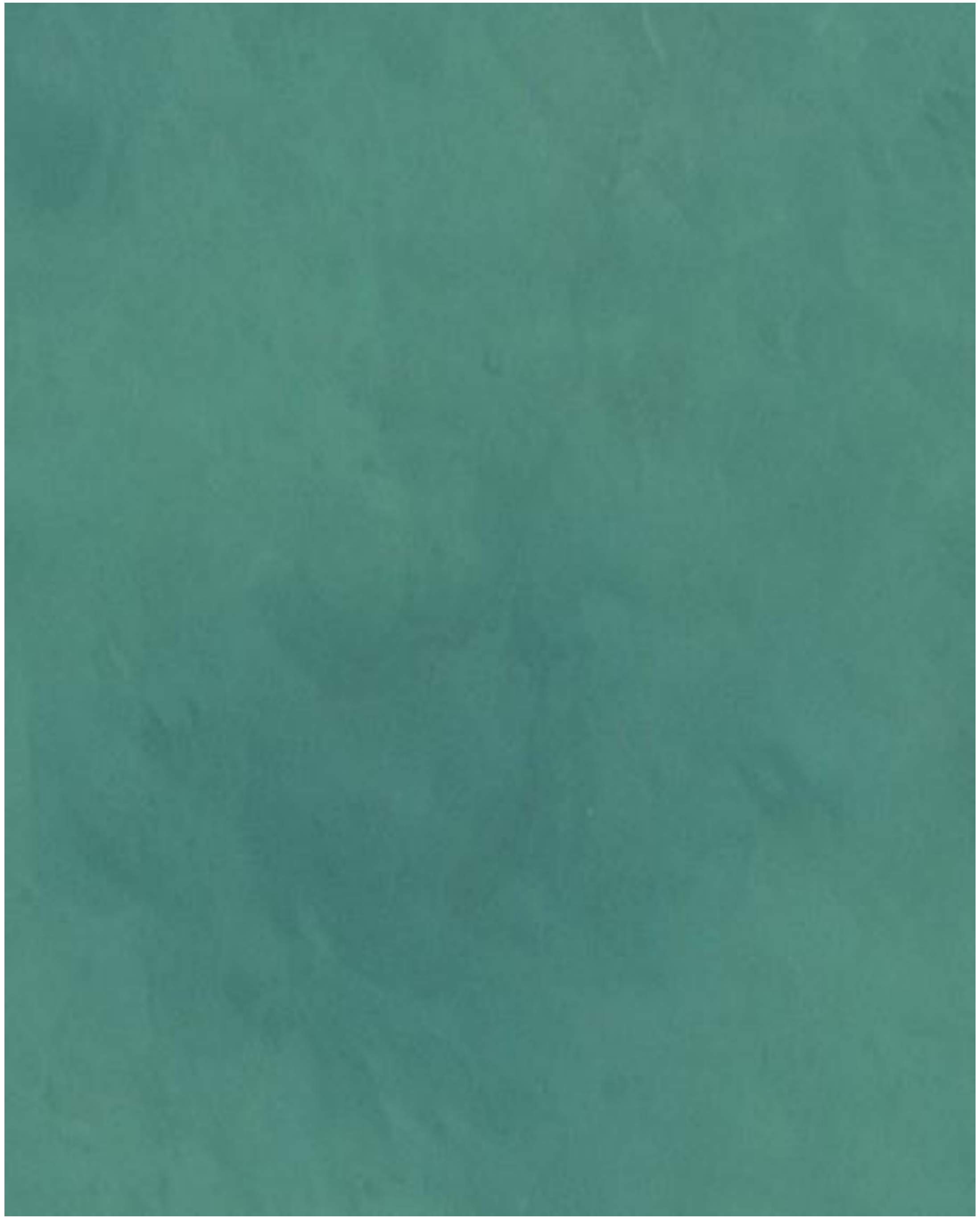

DJI_20240618090215_0906

- Whipray
- There really is one! The darker patch is the body, from which the tail points upwards and slightly to the right.

2024-06-19

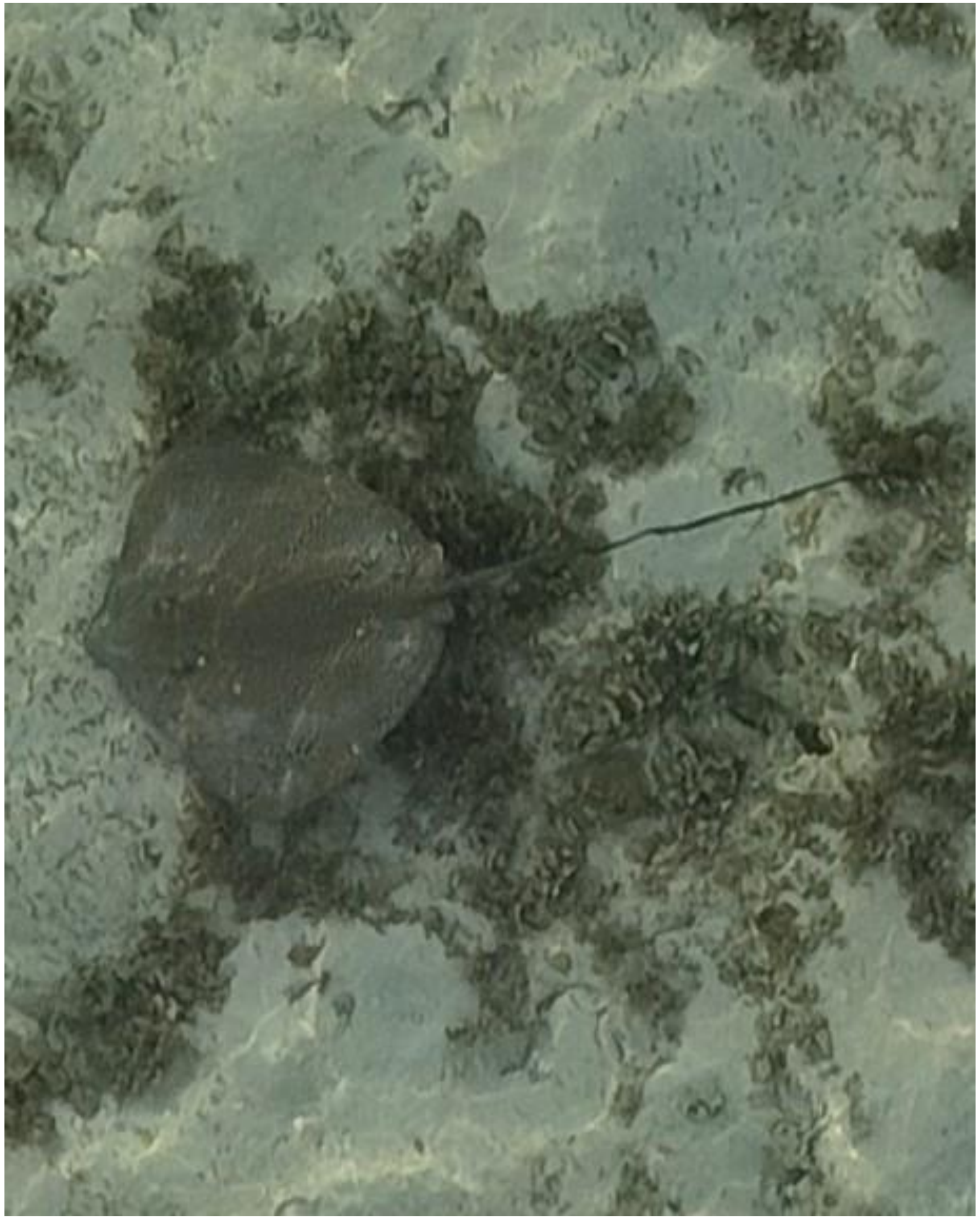

- *Himantura uarnak*

DJI_20240619065828_0026

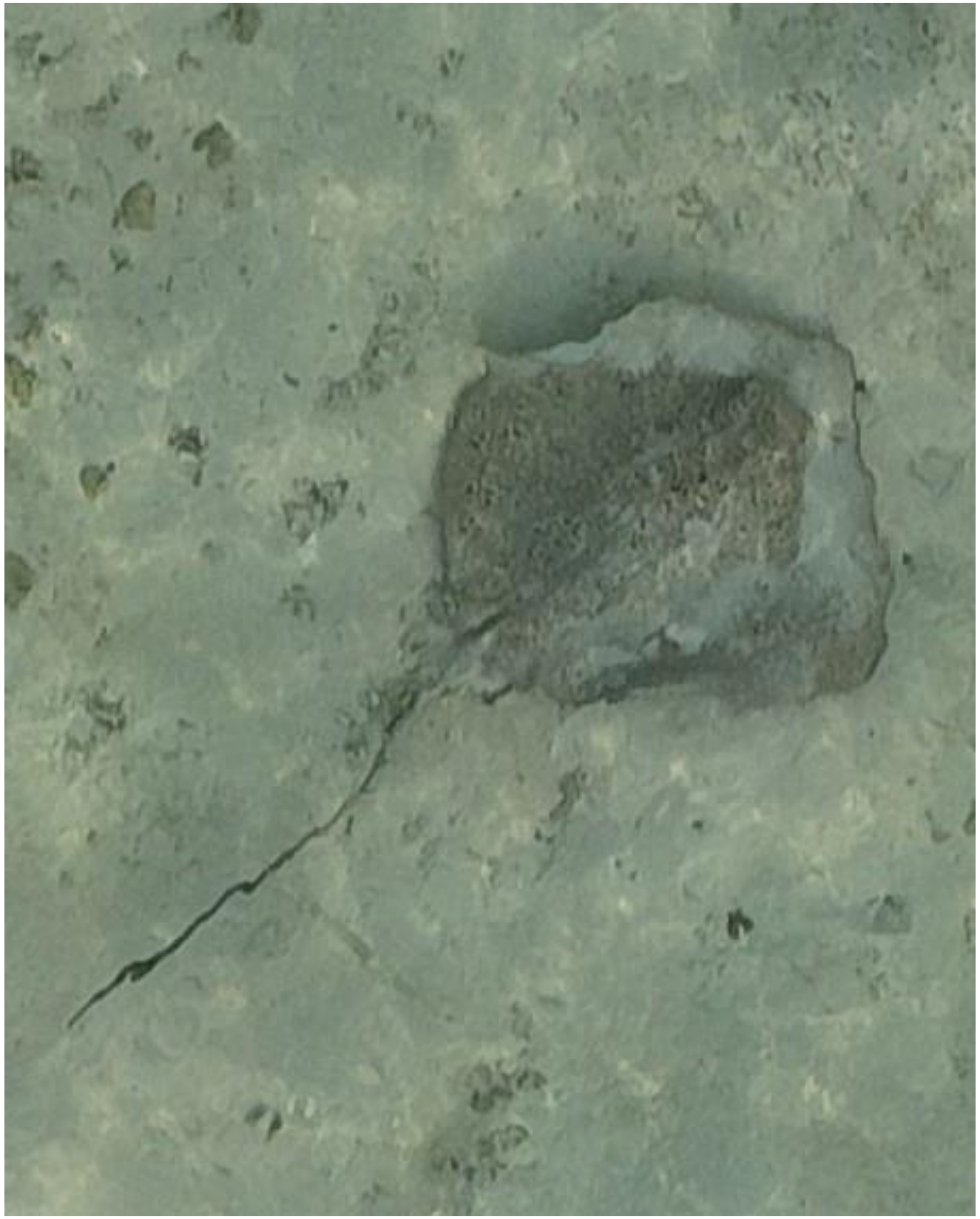

DJI_20240619070325_0288

- *Himantura uarnak*

2024-06-23

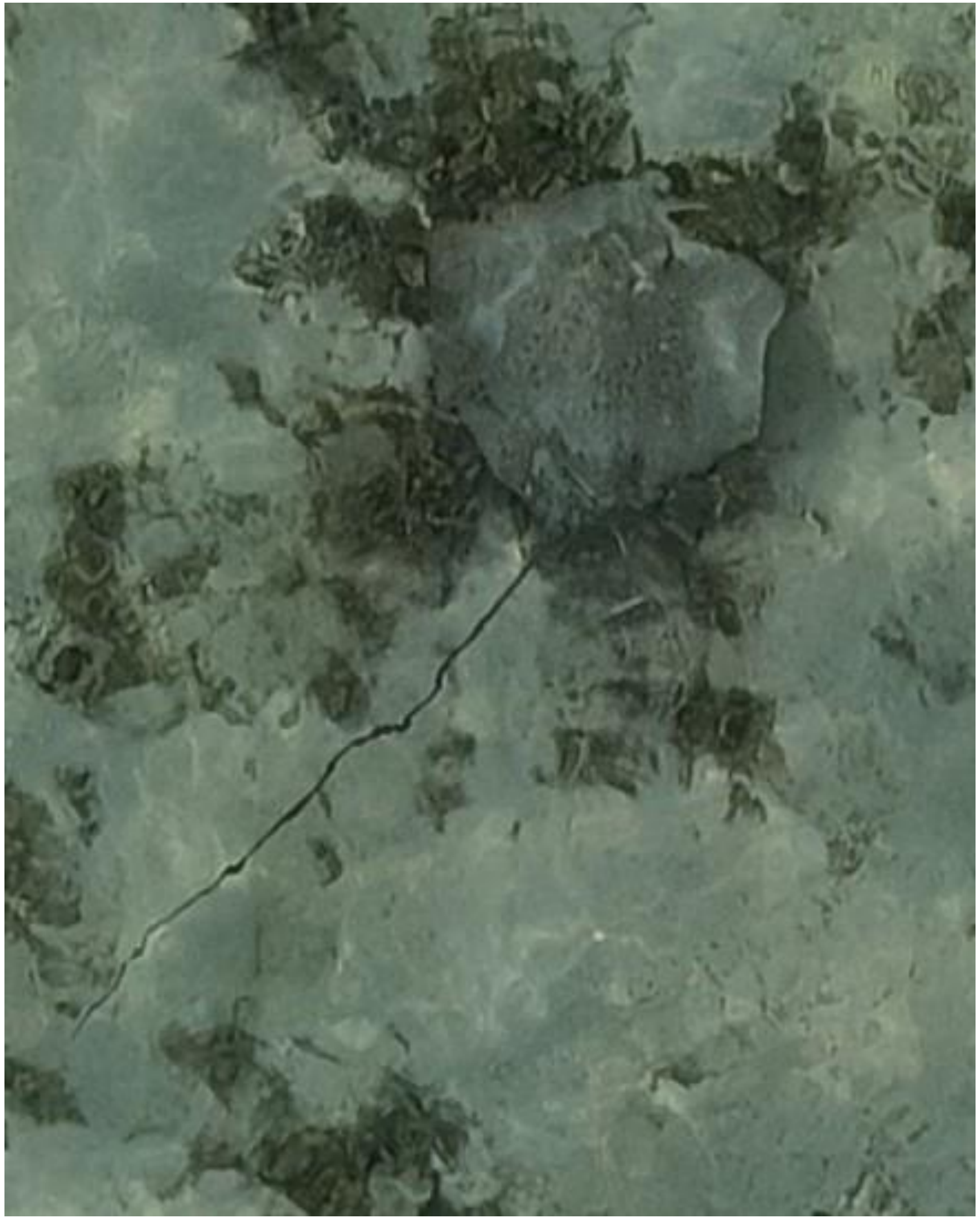

DJI_20240623071705_0047

- *Himantura uarnak*

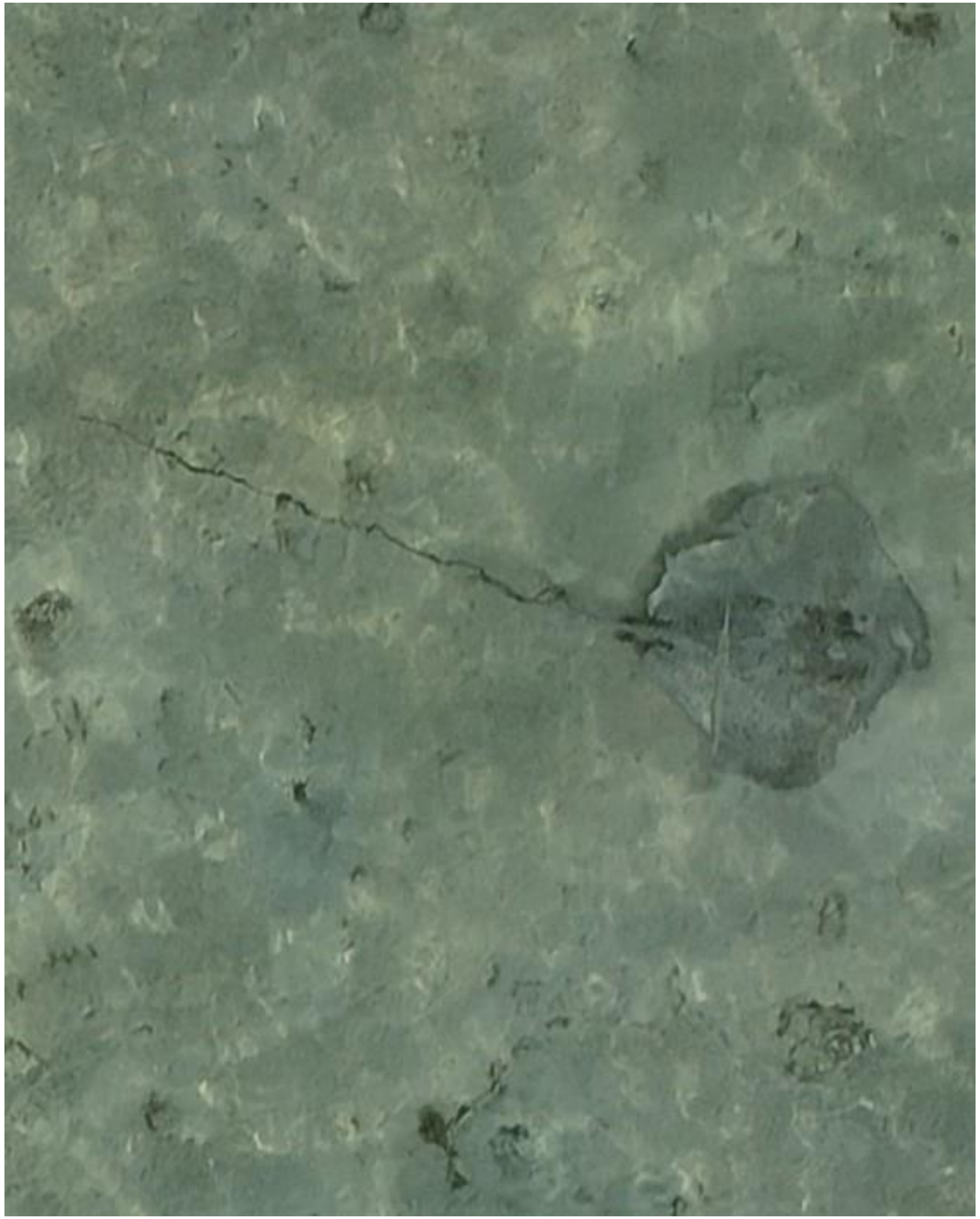

DJI_20240623071806_0096

- Whipray
- Best guess: *H. uarnak*
- Accompanied by needlefish

2024-06-24

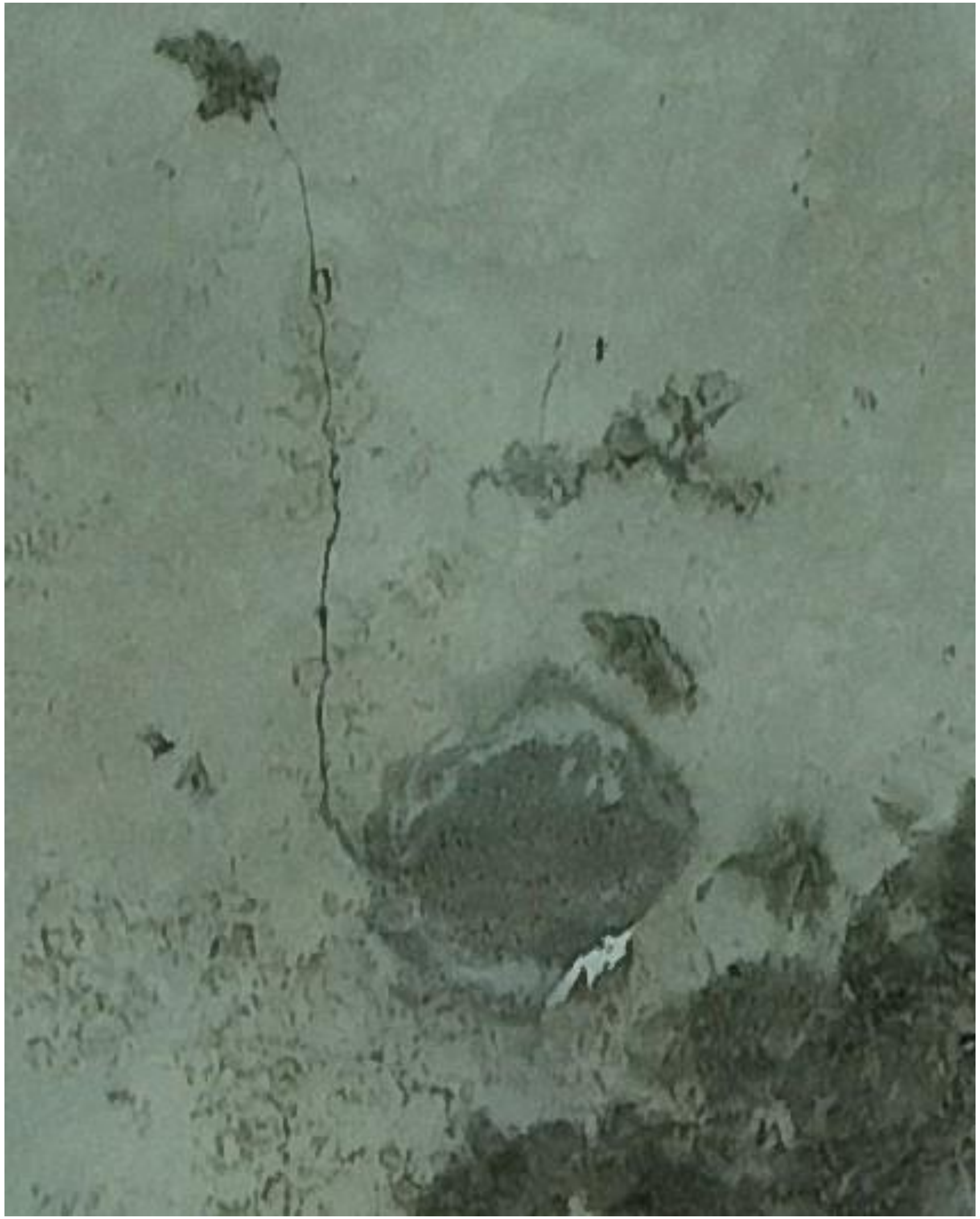

DJI_20240624072158_0106

- *Himantura uarnak*

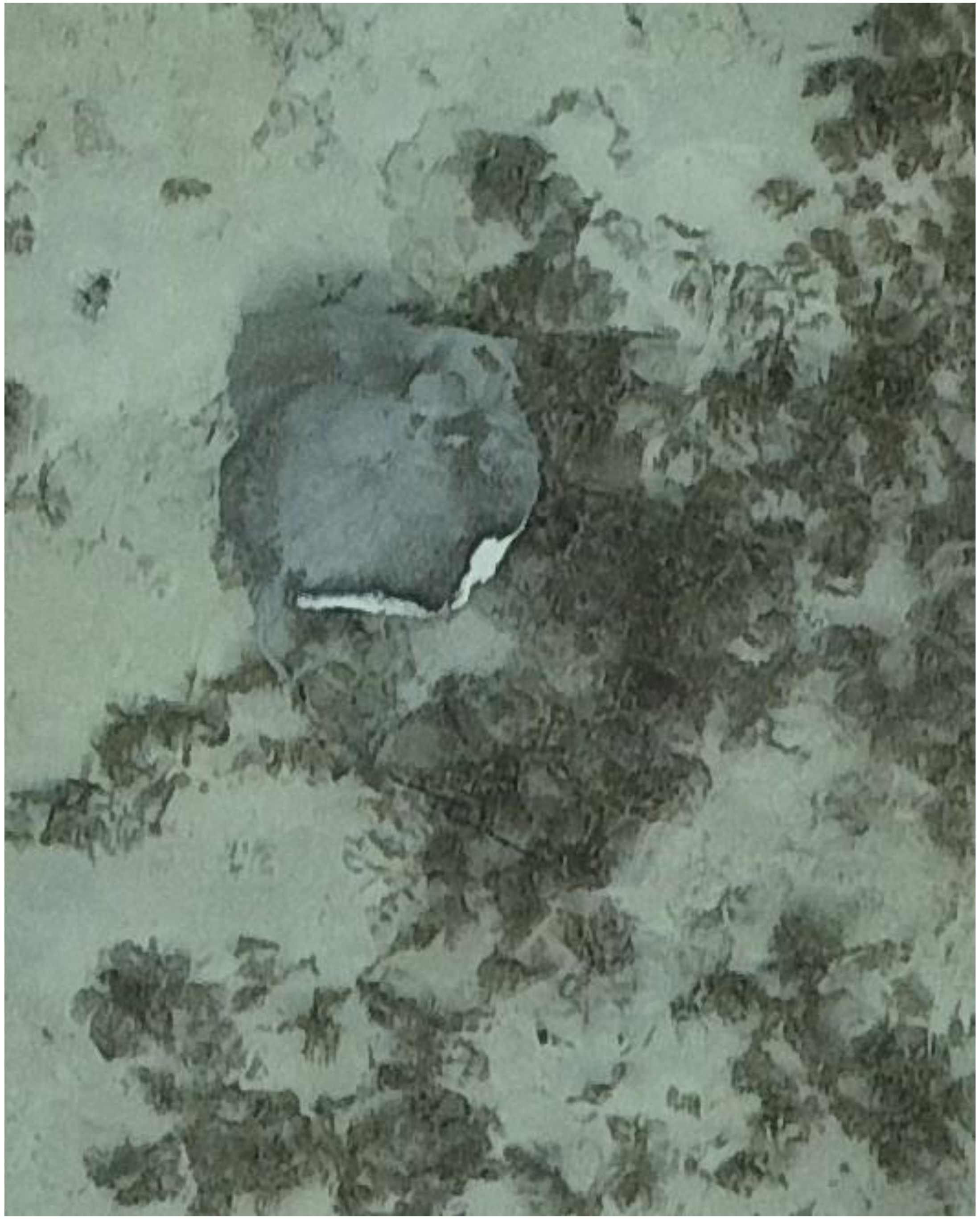

- Whipray

DJI_20240624072226_0134

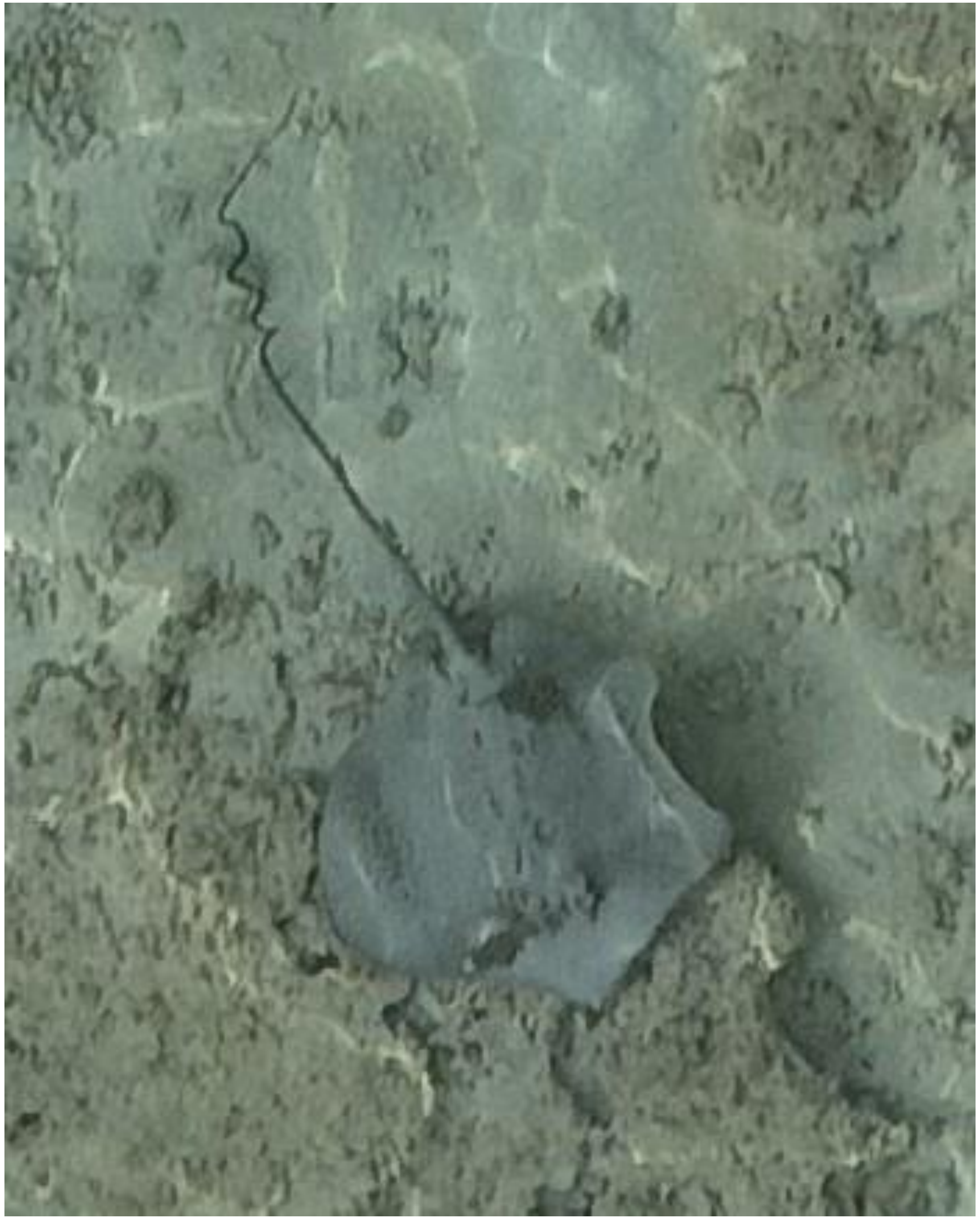

DJI_20240624073040_0563

- Whipray
- Also observed on 05-02, 06-06, 07-29, and 09-20 as identified by the coiled tail.
- Best guess: *H. uarnak*

2024-06-25

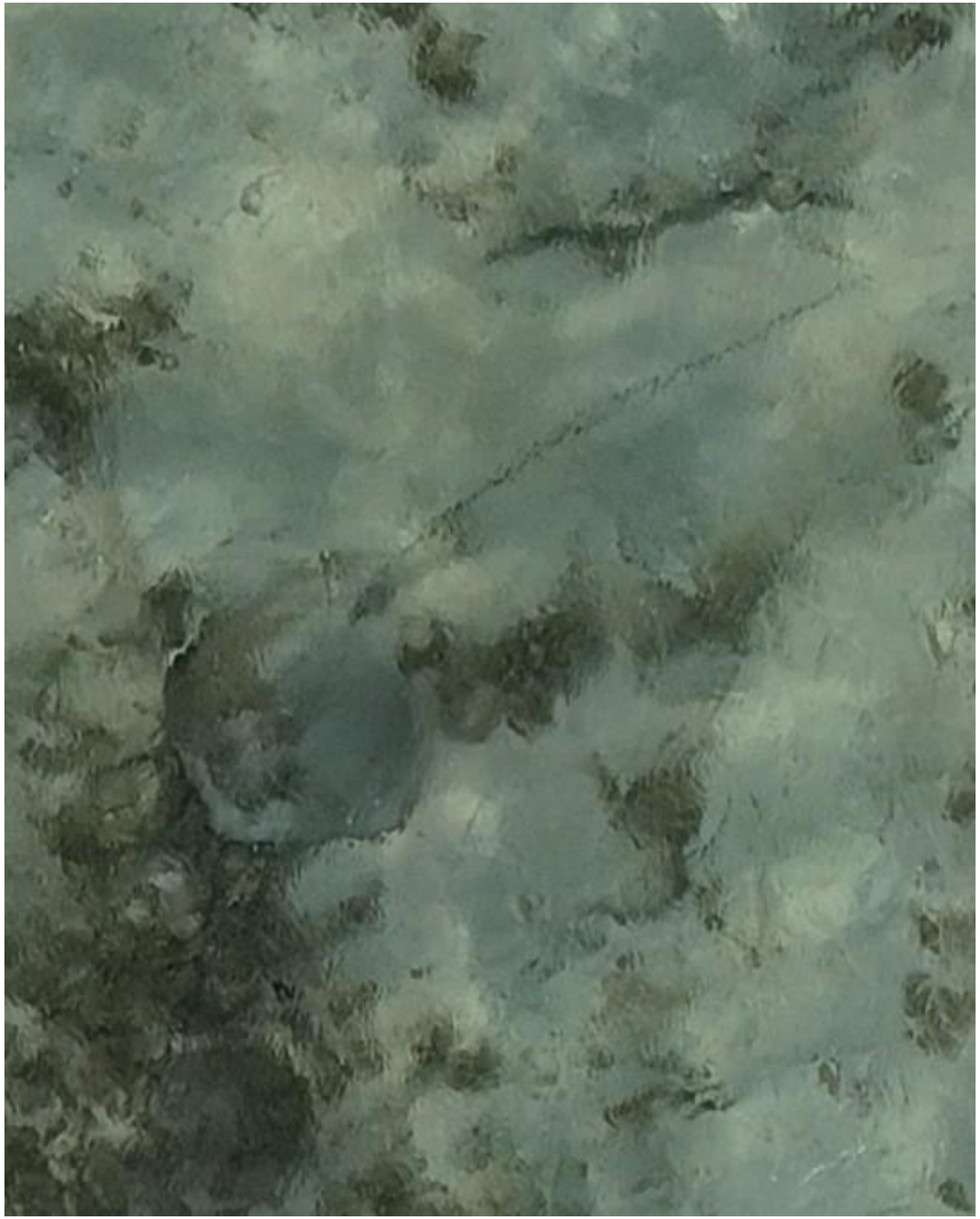

- Whipray

DJI_20240625071841_0039

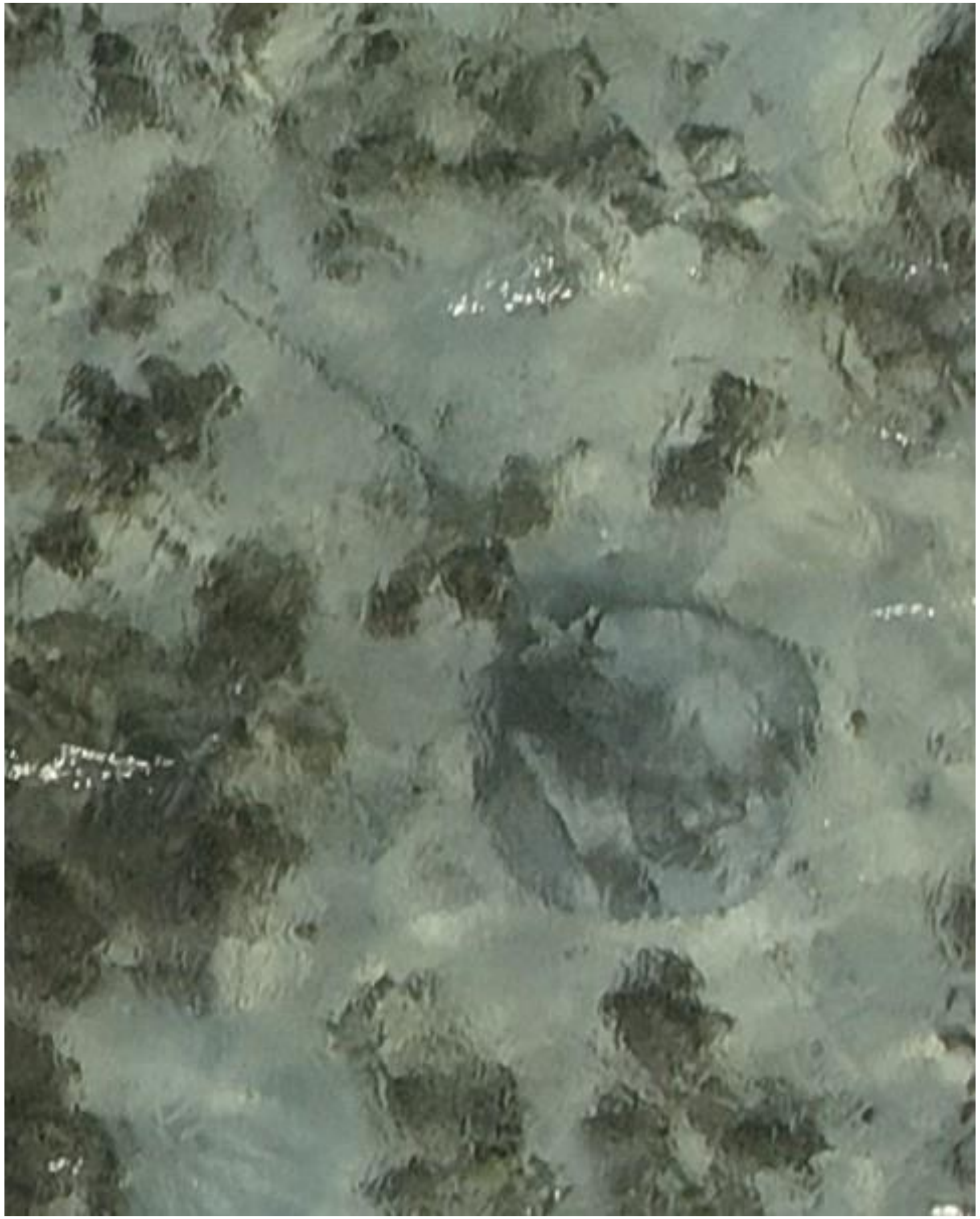

DJI_20240625072400_0322

- Whipray

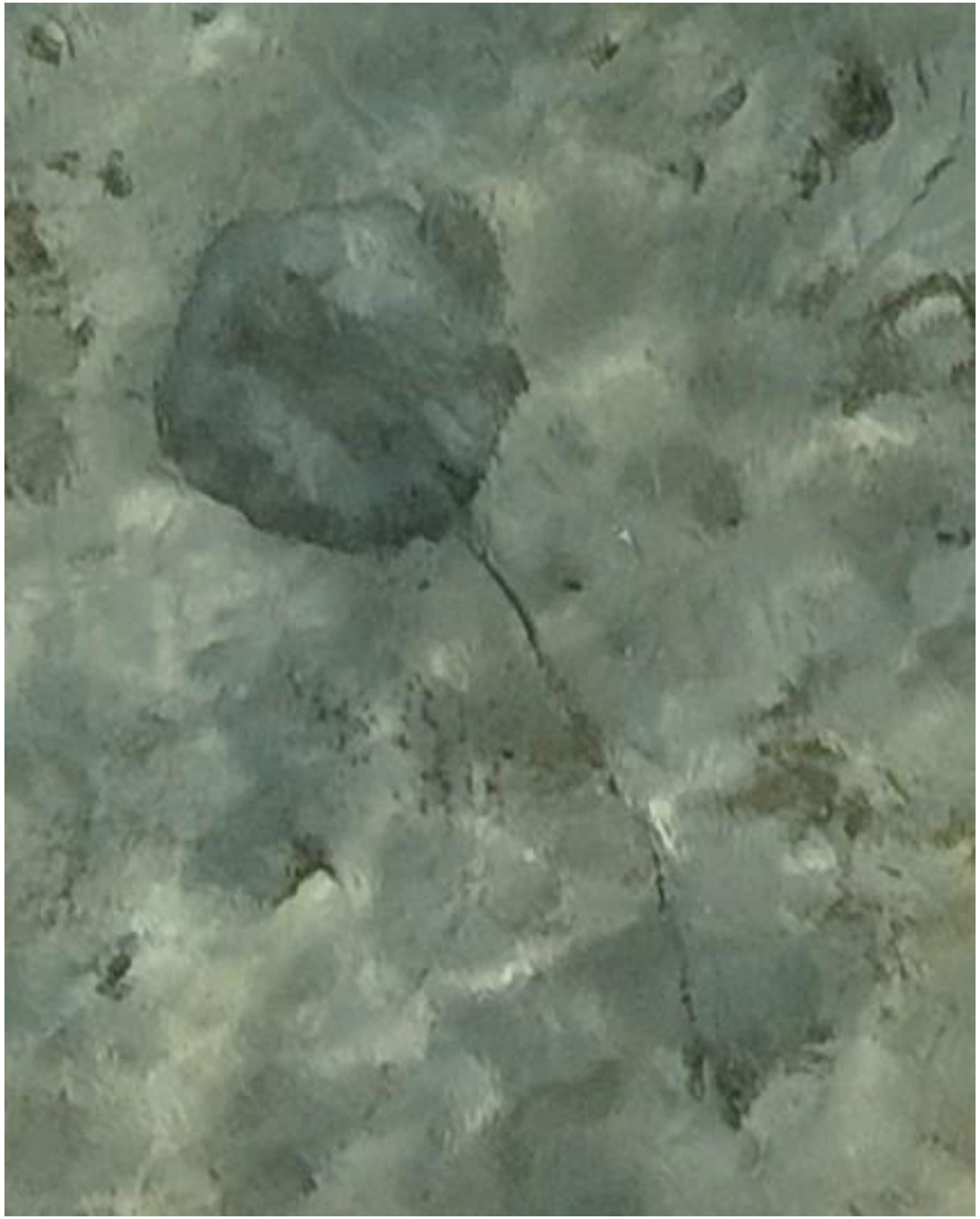

DJI_20240625072536_0406

- Whipray

2024-07-02

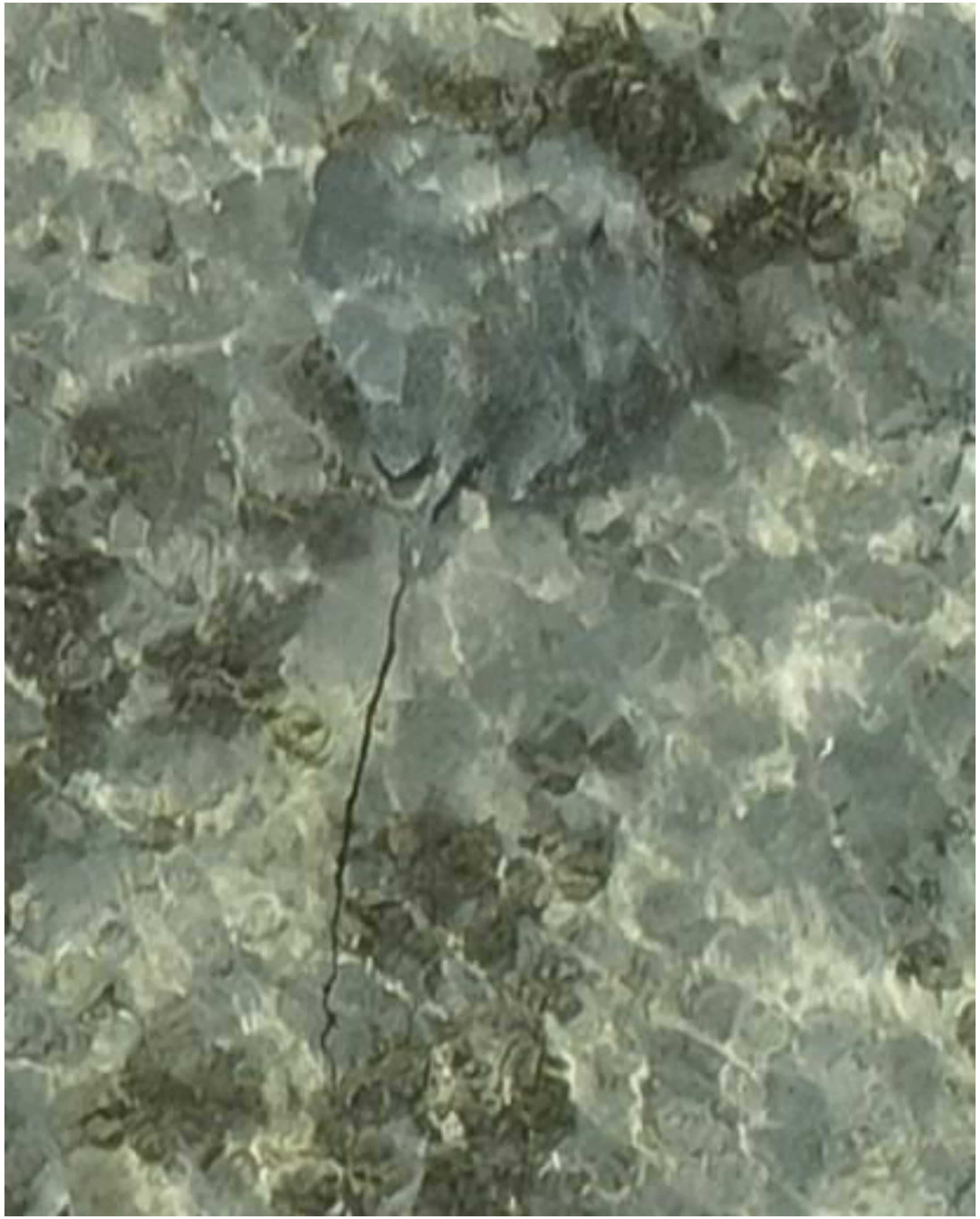

DJI_20240702072456_0025

- Whipray
- Best guess: *H. uarnak*

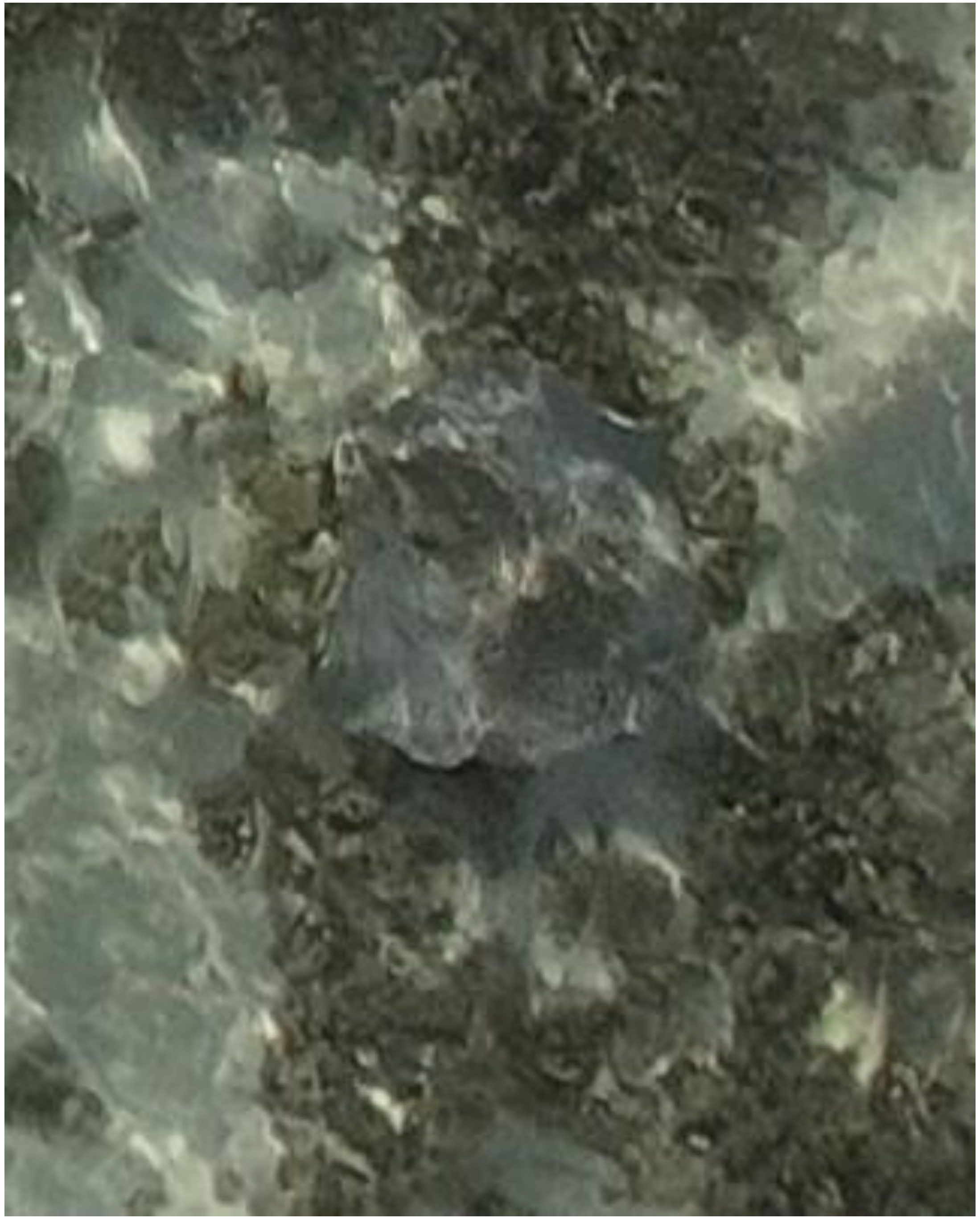

- Whipray

DJI_20240702072510_0039

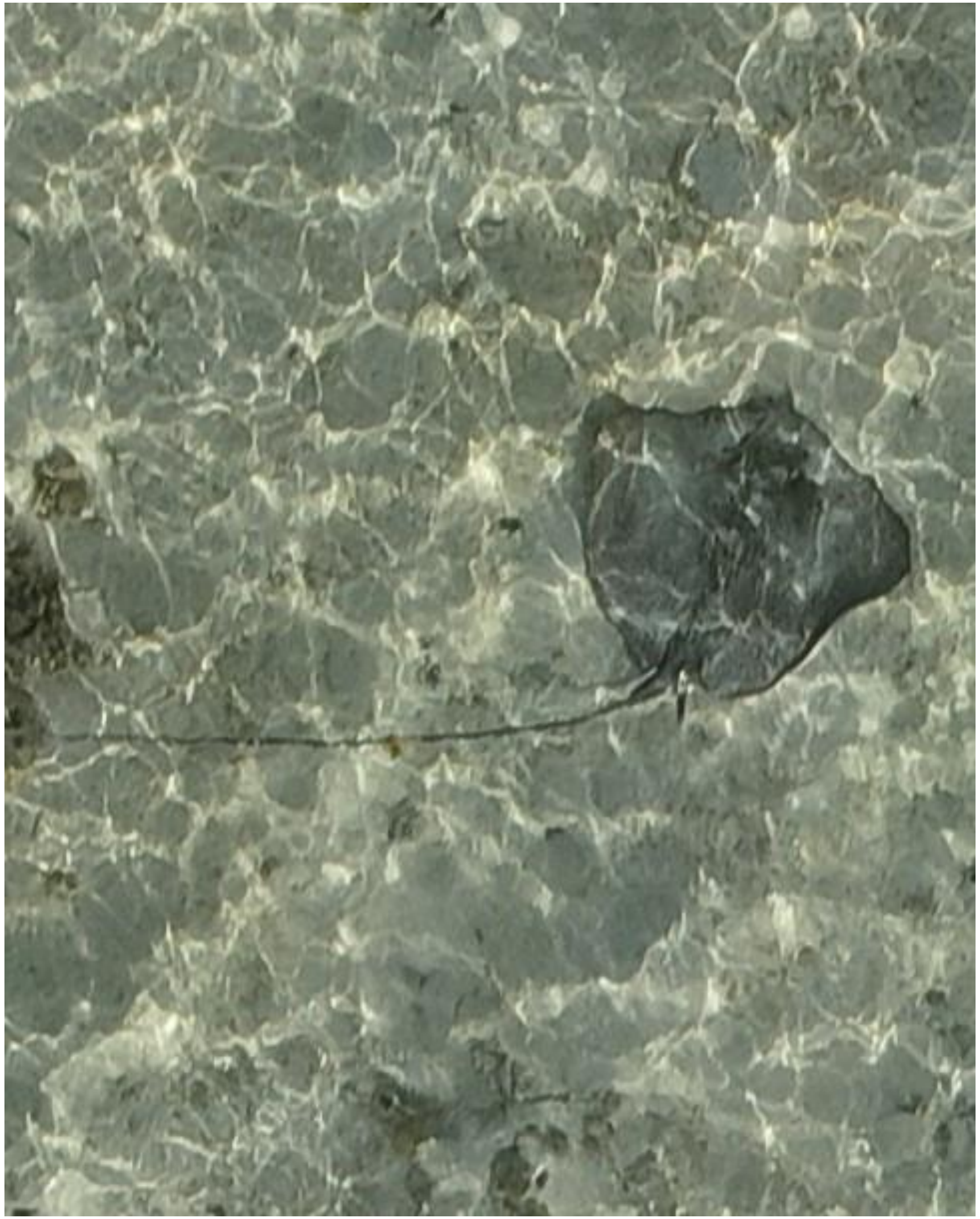

- Whipray
- Best guess: *H. uarnak*

DJI_20240702072643_0120

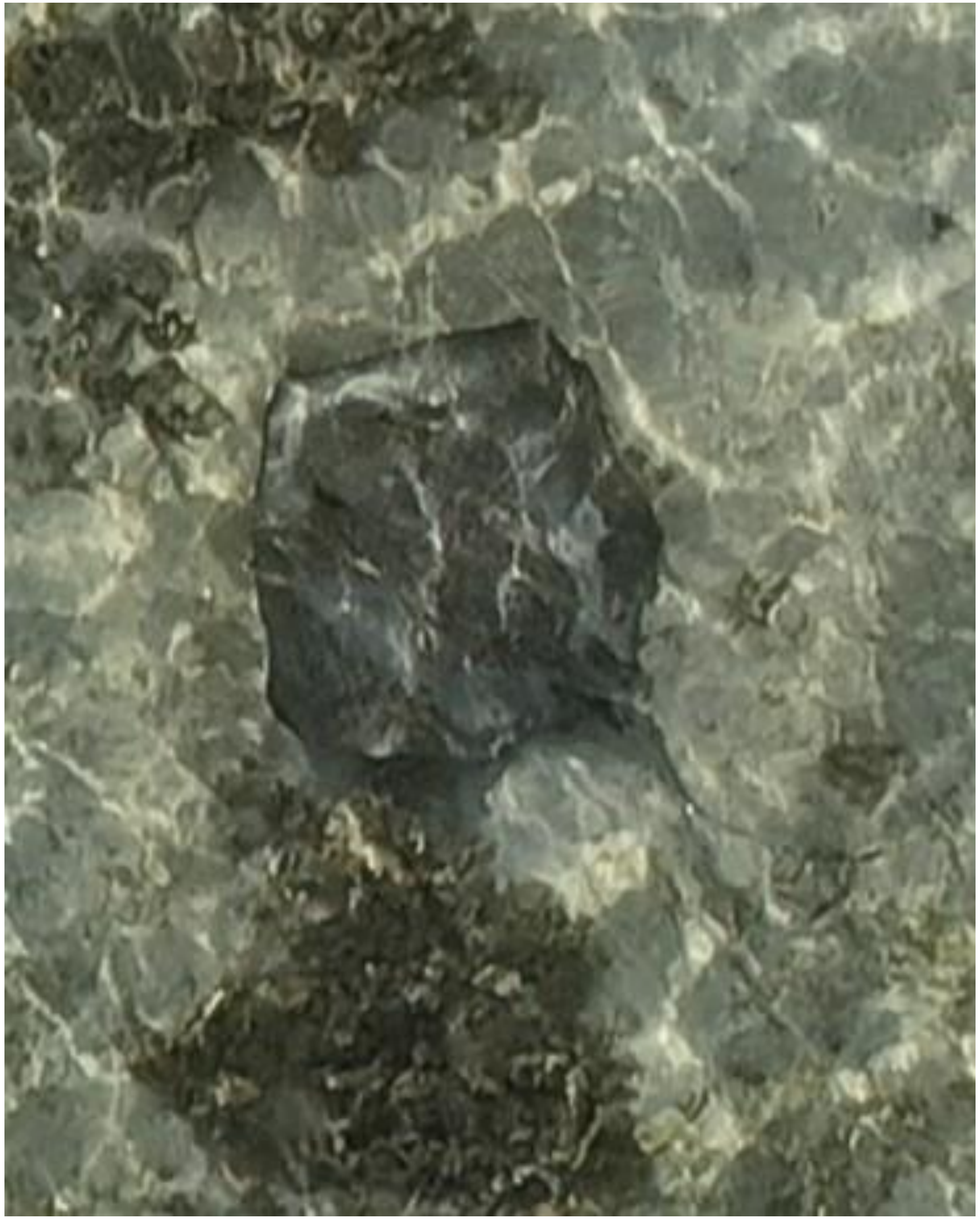

- Whipray
- Best guess: *H. uarnak*

DJI_20240702072802_0187

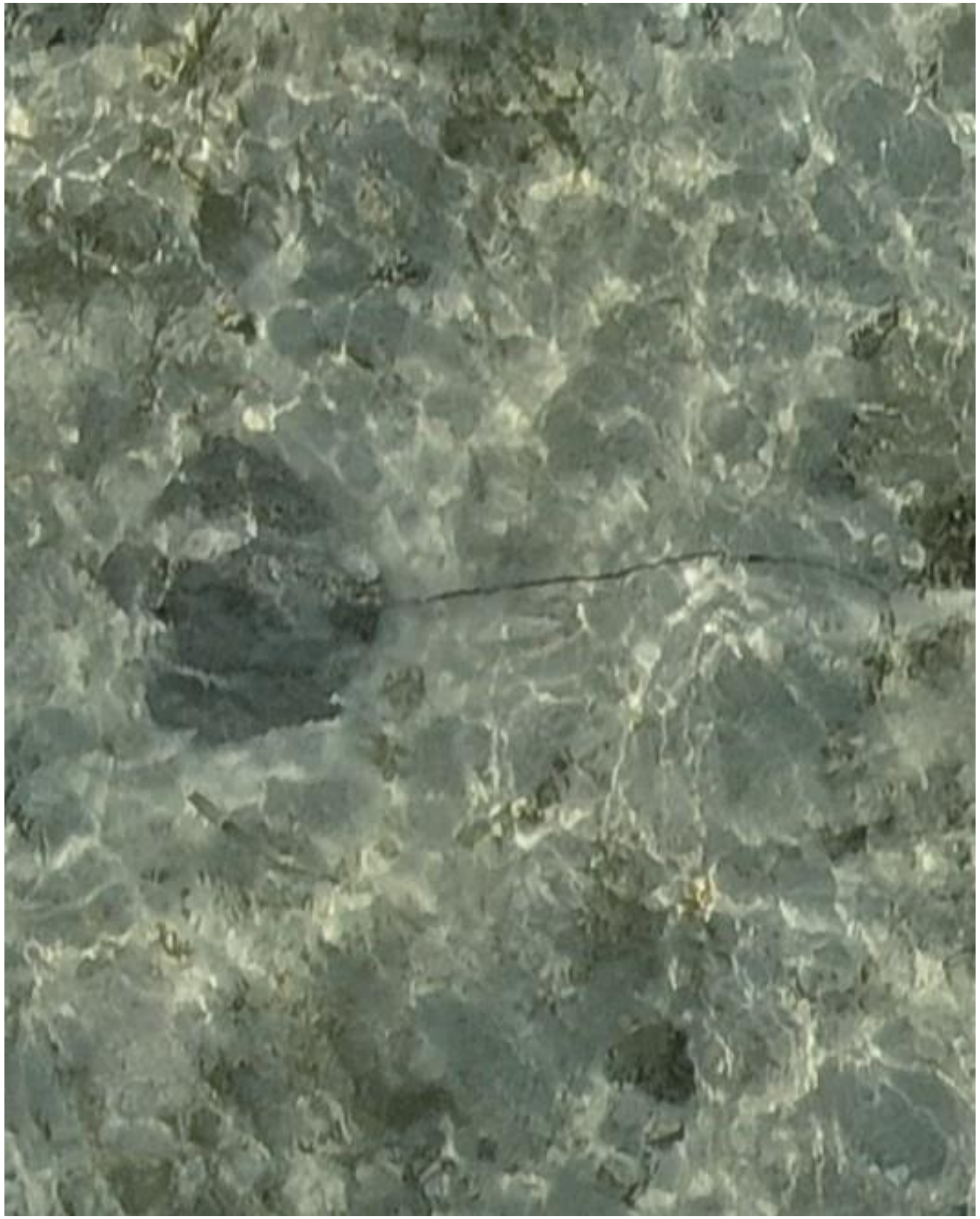

DJI_20240702072828_0213

- *Himantura uarnak*
- Accompanied by needlefish

2024-07-03

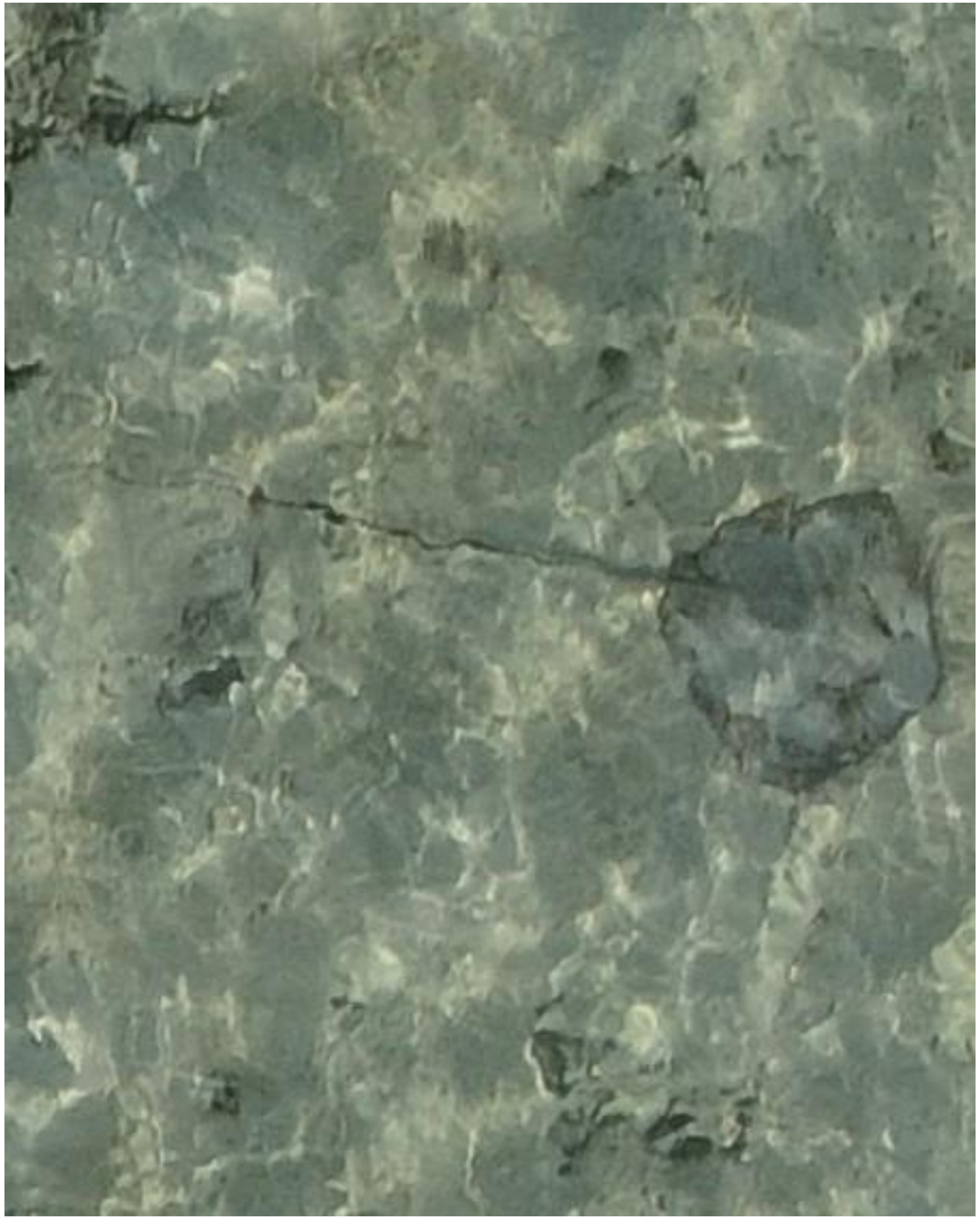

DJI_20240703070827_0102

- Whipray

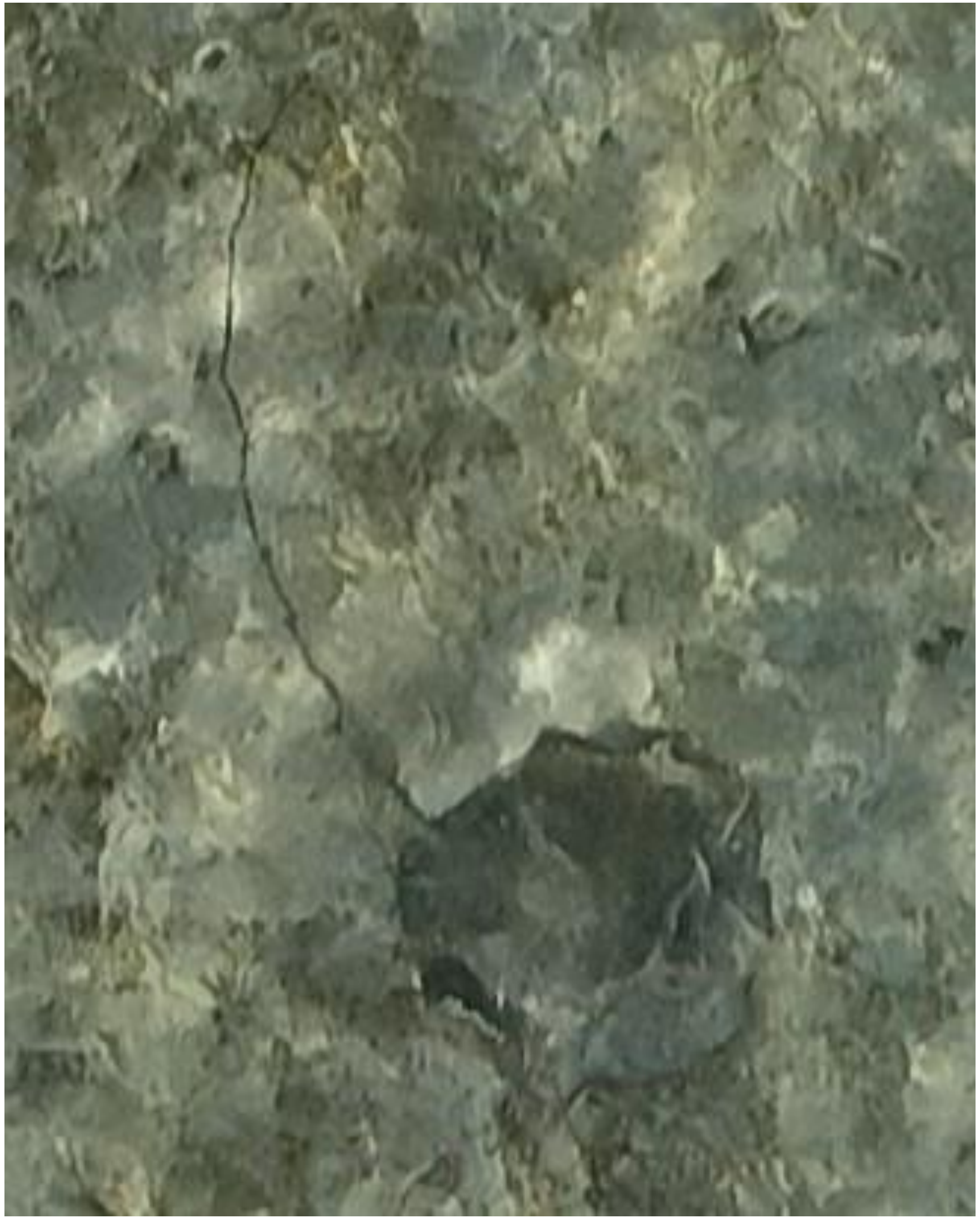

DJI_20240703071202_0293

- Whipray

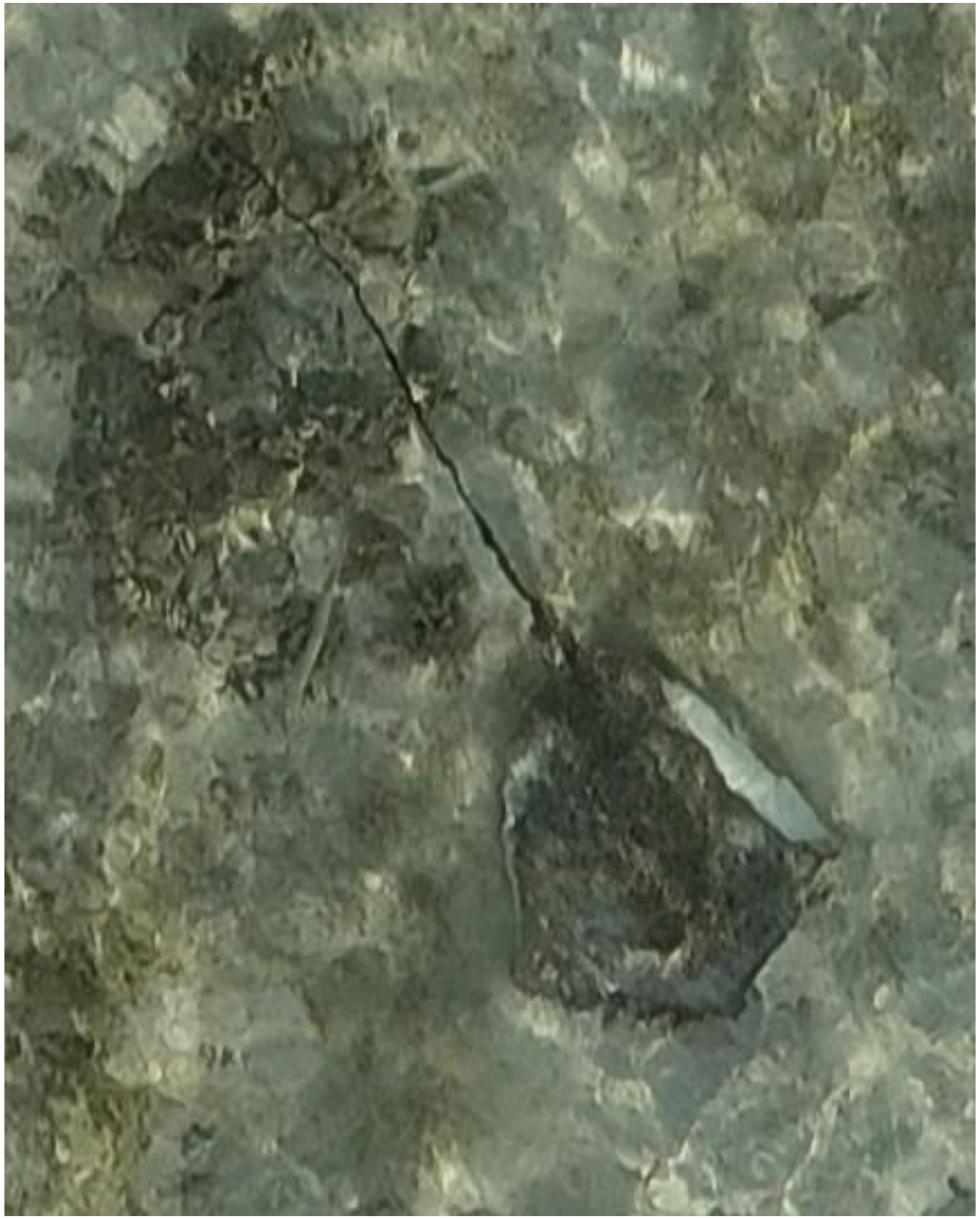

DJI_20240703071207_0298

- *Himantura uarnak*
- Accompanied by needlefish

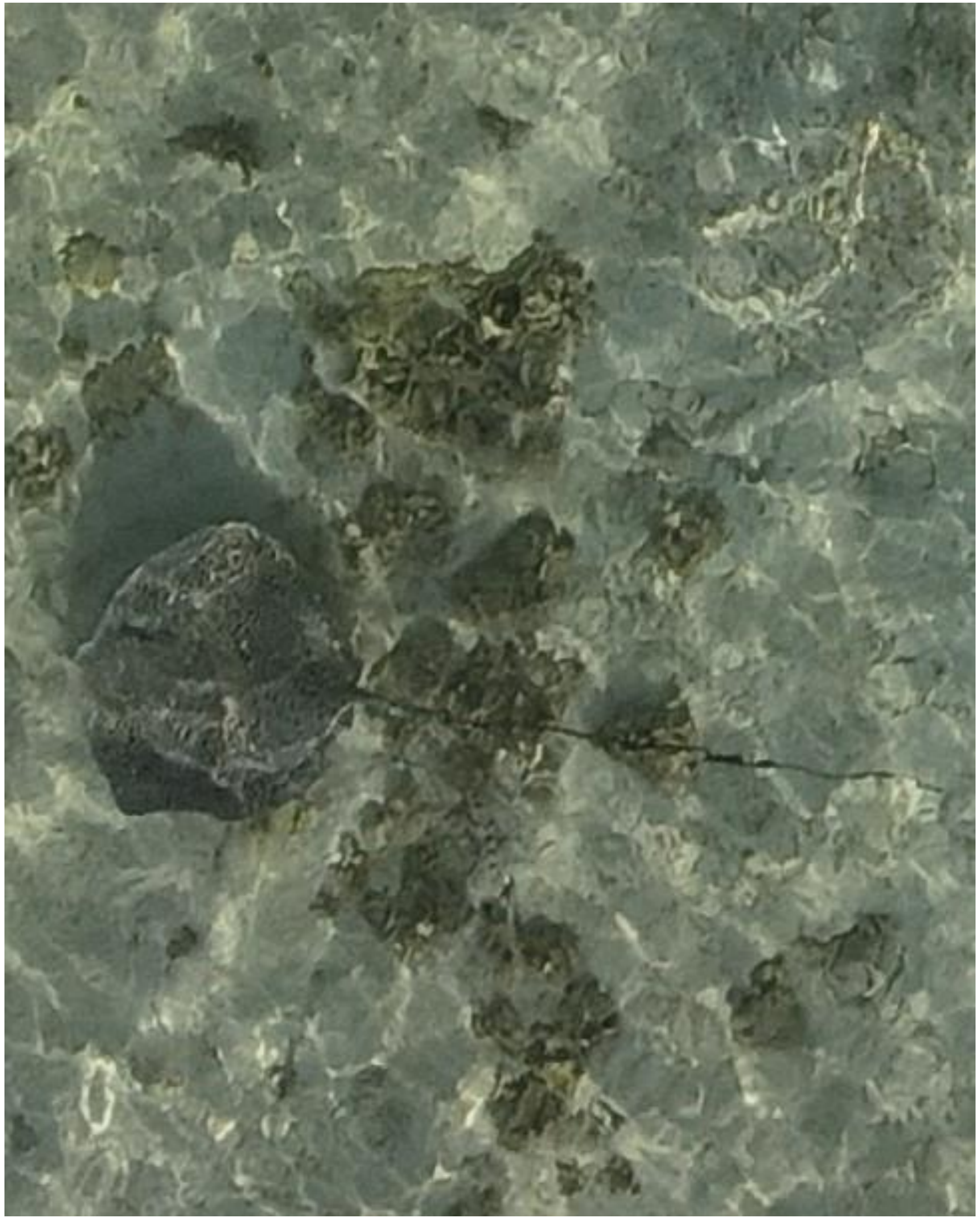

DJI_20240703071222_0313

- *Himantura uarnak*

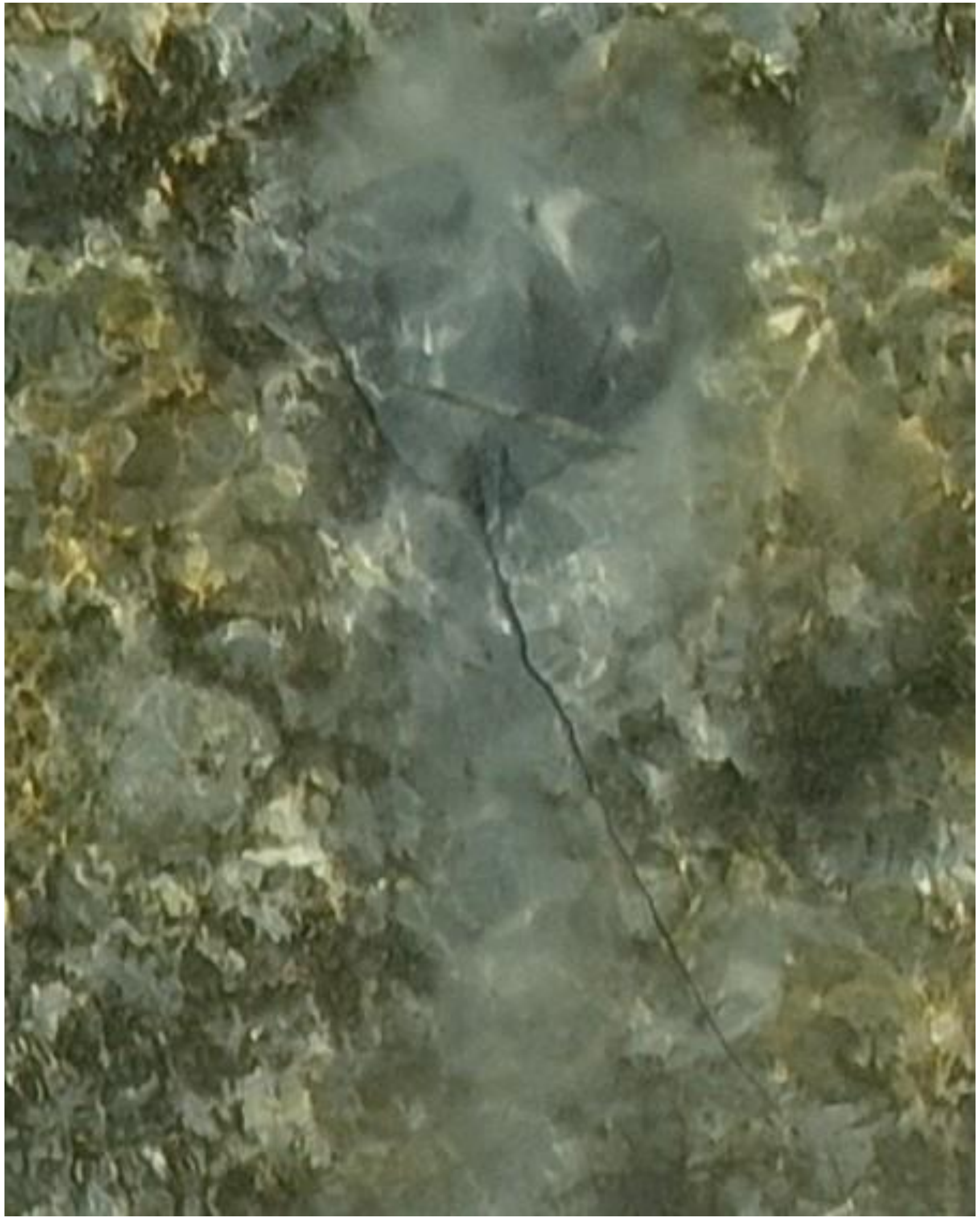

DJI_20240703071406_0405

- Whipray
- Accompanied by needlefish

2024-07-29

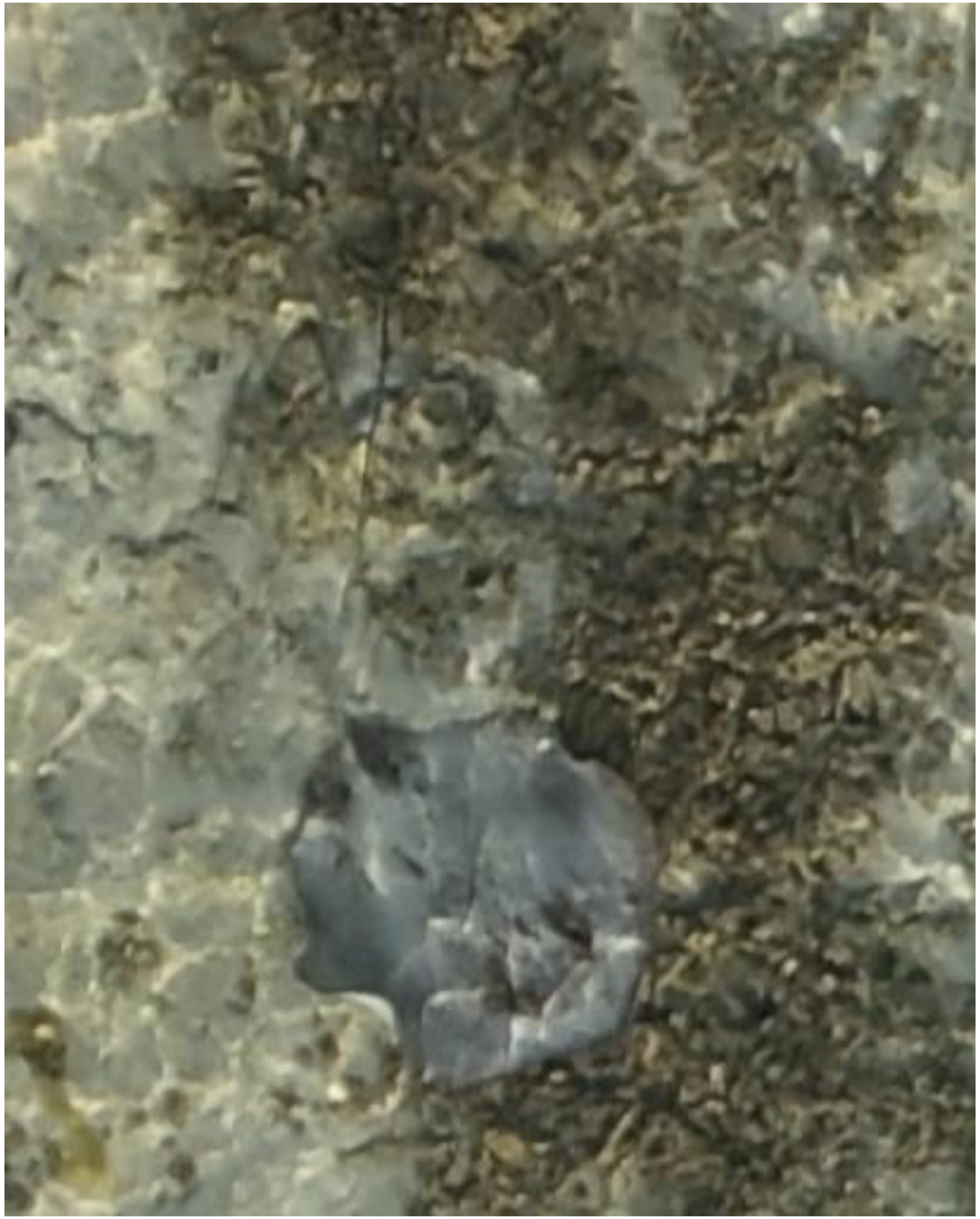

- Whipray

DJI_20240729071855_0115

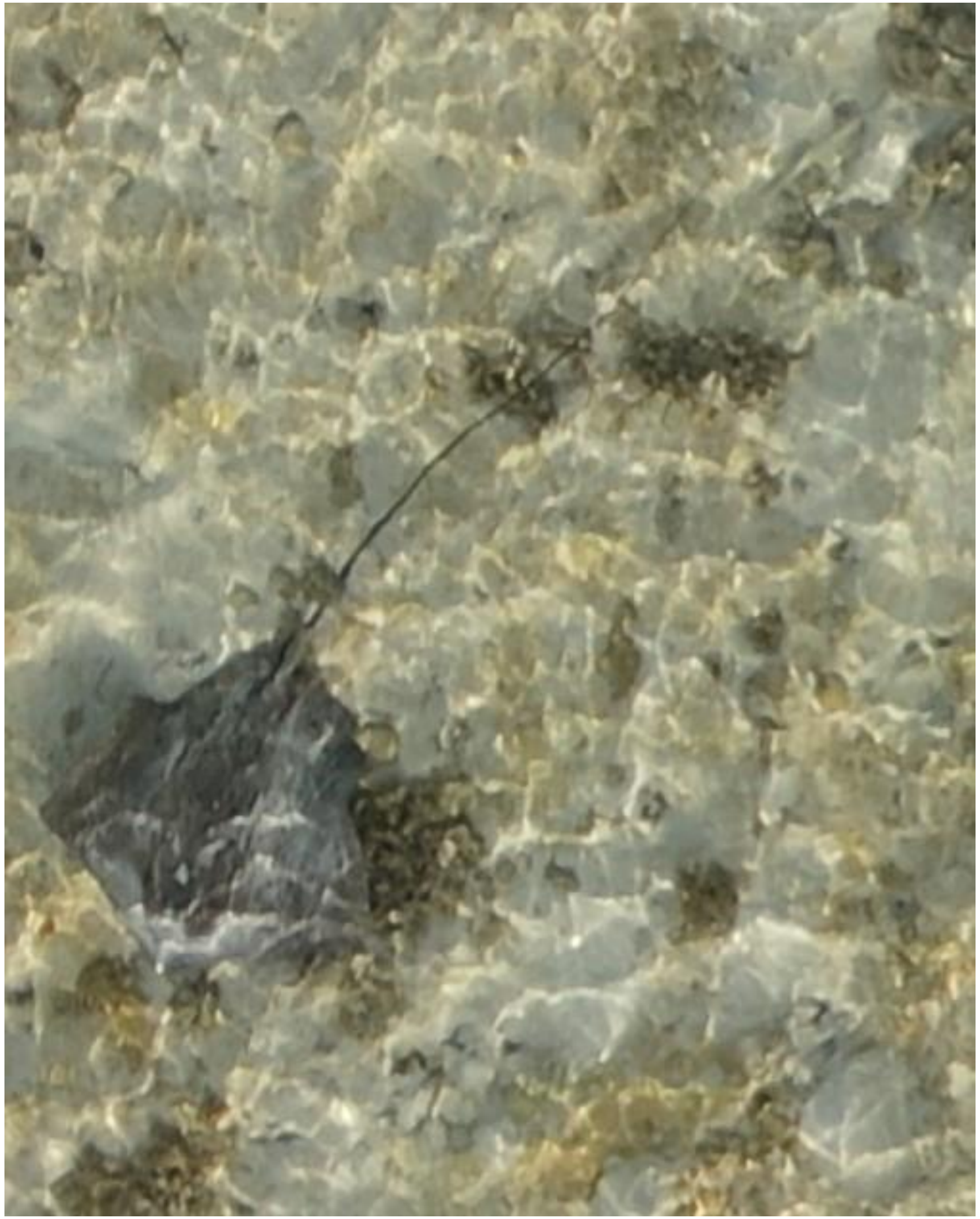

- Whipray
- Accompanied by needlefish

DJI_20240729071909_0129

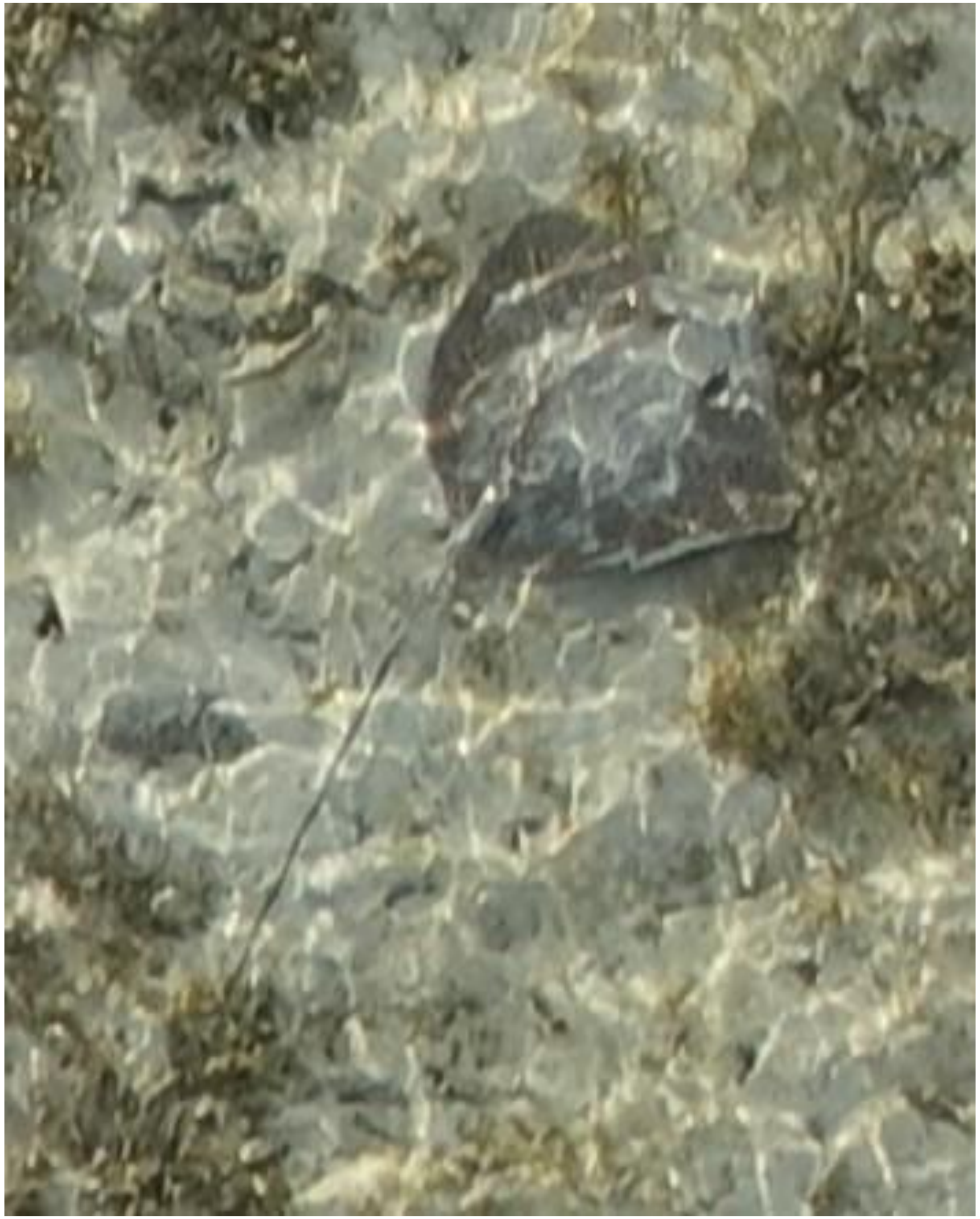

- Whipray
- Best guess: *H. uarnak*

DJI_20240729072042_0210

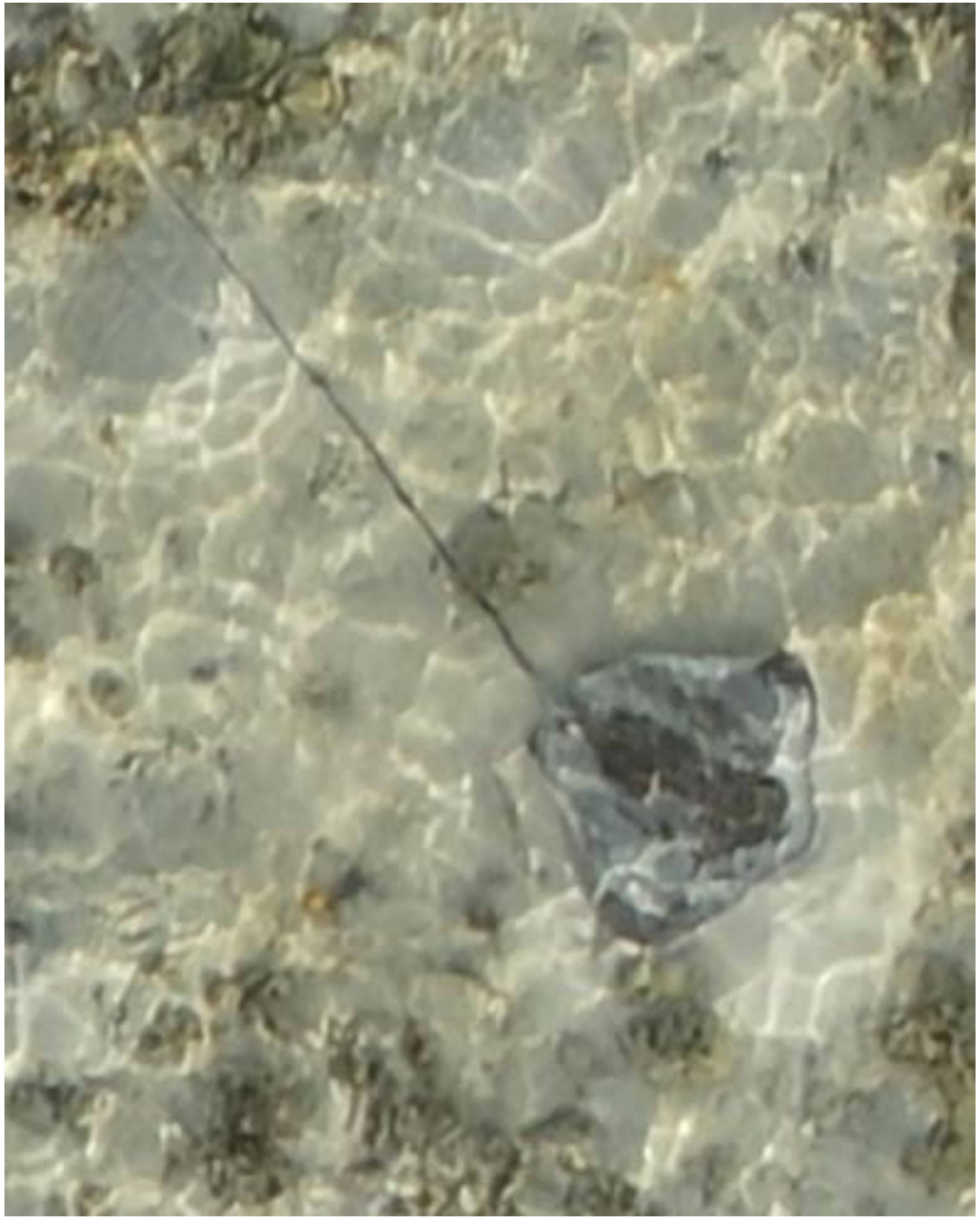
DJI_20240729072241_0317

- Whipray

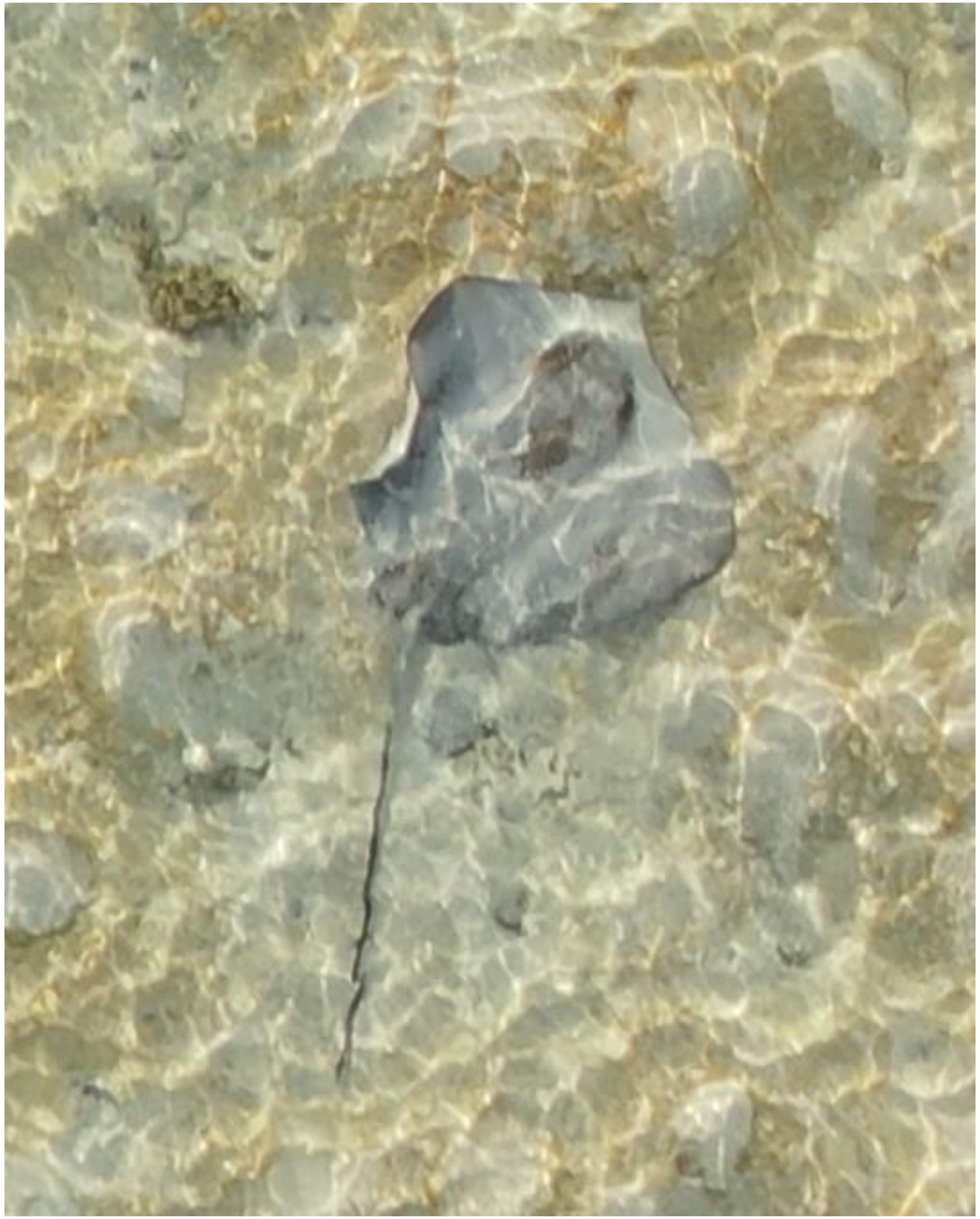

- Whipray

DJI_20240729072428_0412

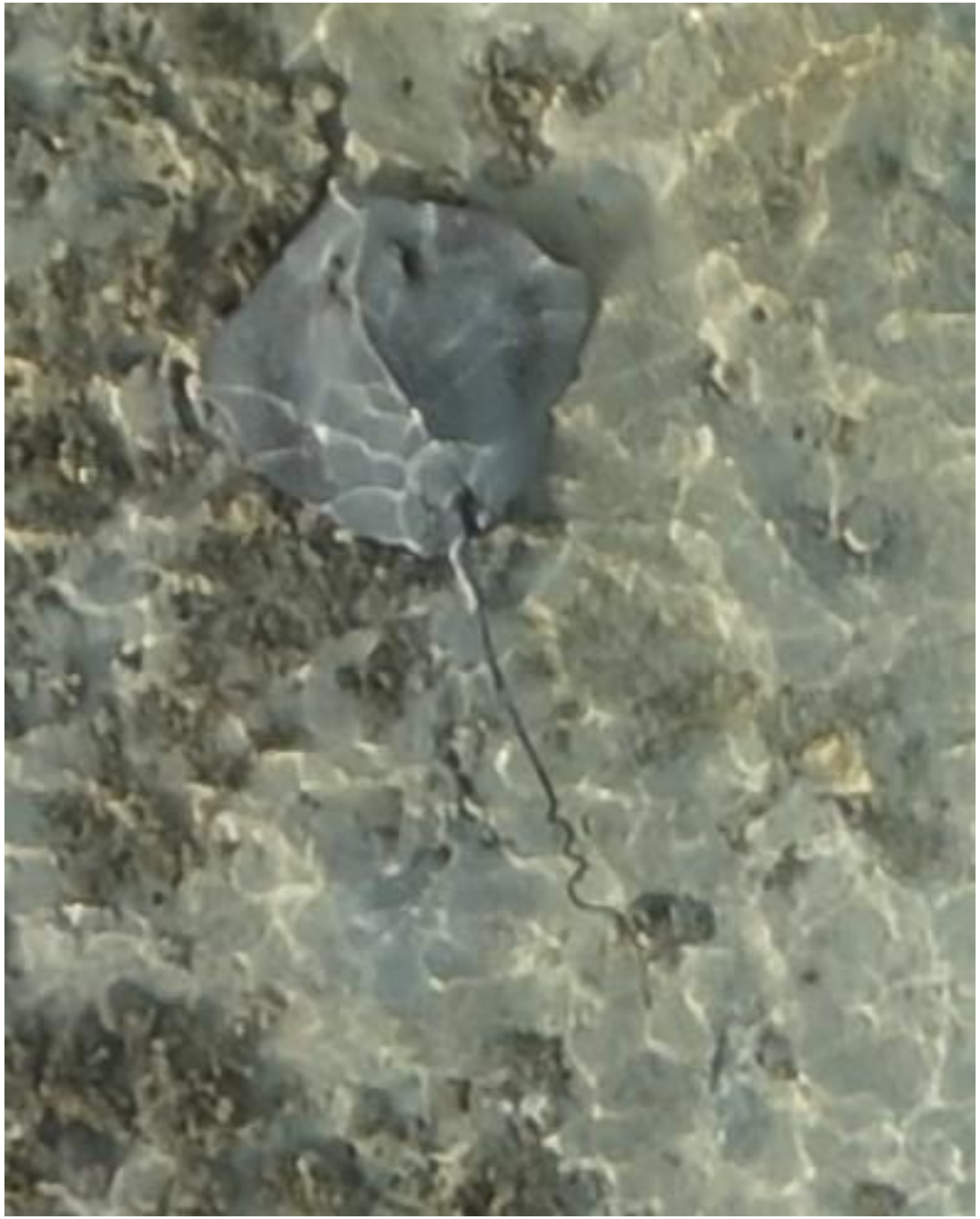

- Whipray
- Also observed on 05-02, 06-06, 06-24, and 09-20 as identified by the coiled tail.
- Best guess: *H. uarnak*

DJI_20240729072754_0588

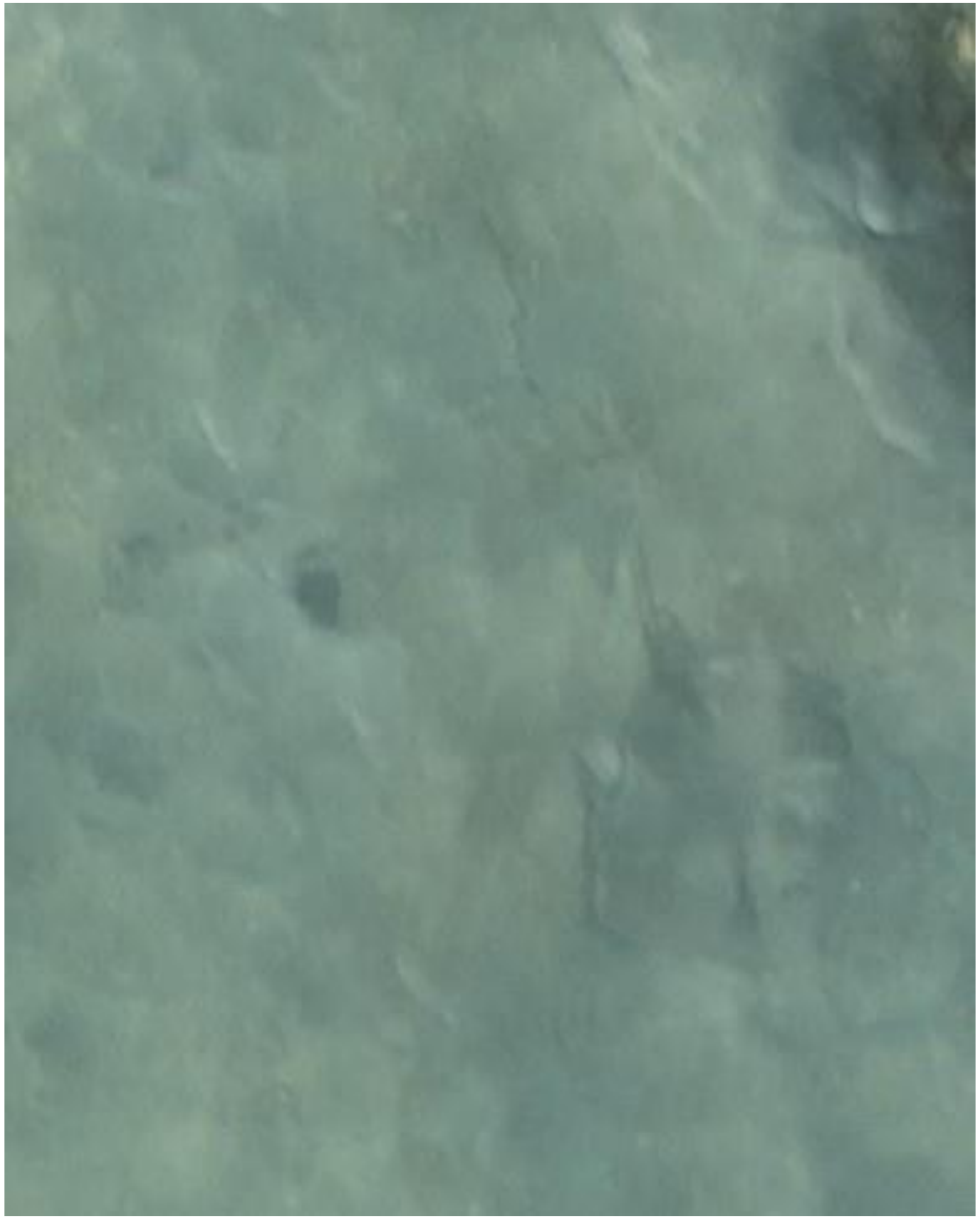

DJI_20240729072852_0628

- Whipray

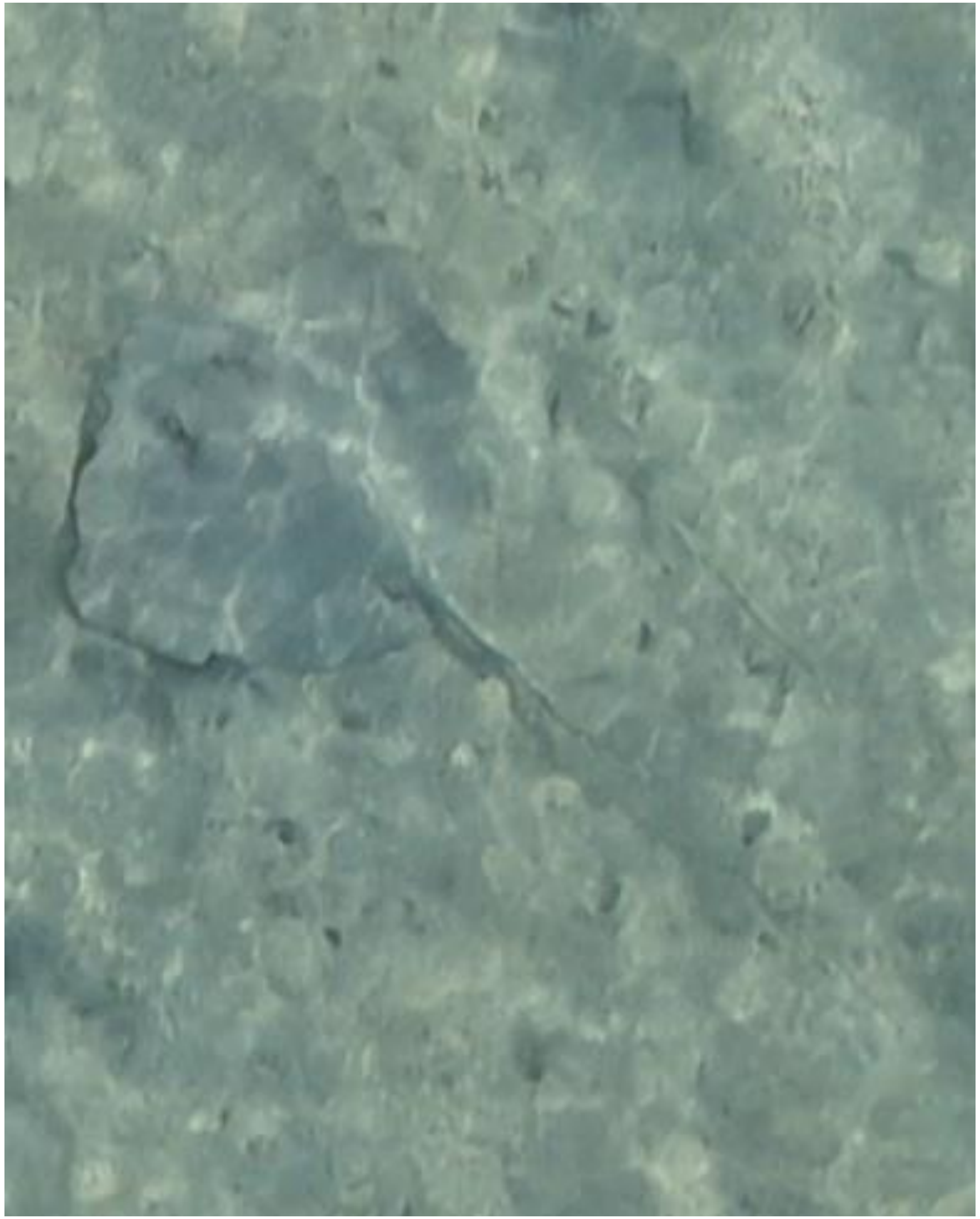

DJI_20240729073420_0932

- Whipray
- Accompanied by needlefish

2024-08-14

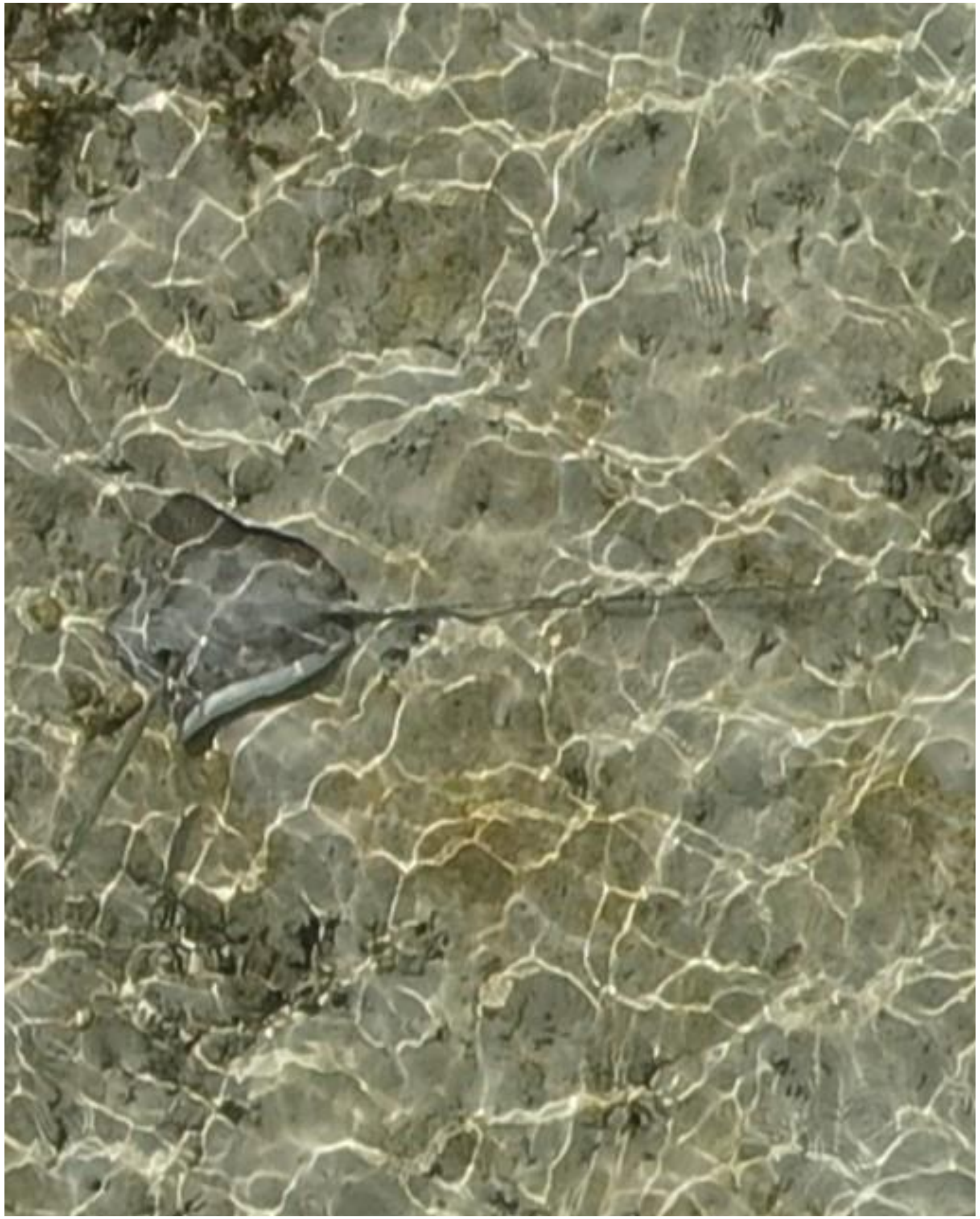

DJI_20240814085407_0210

- Whipray
- Accompanied by needlefish

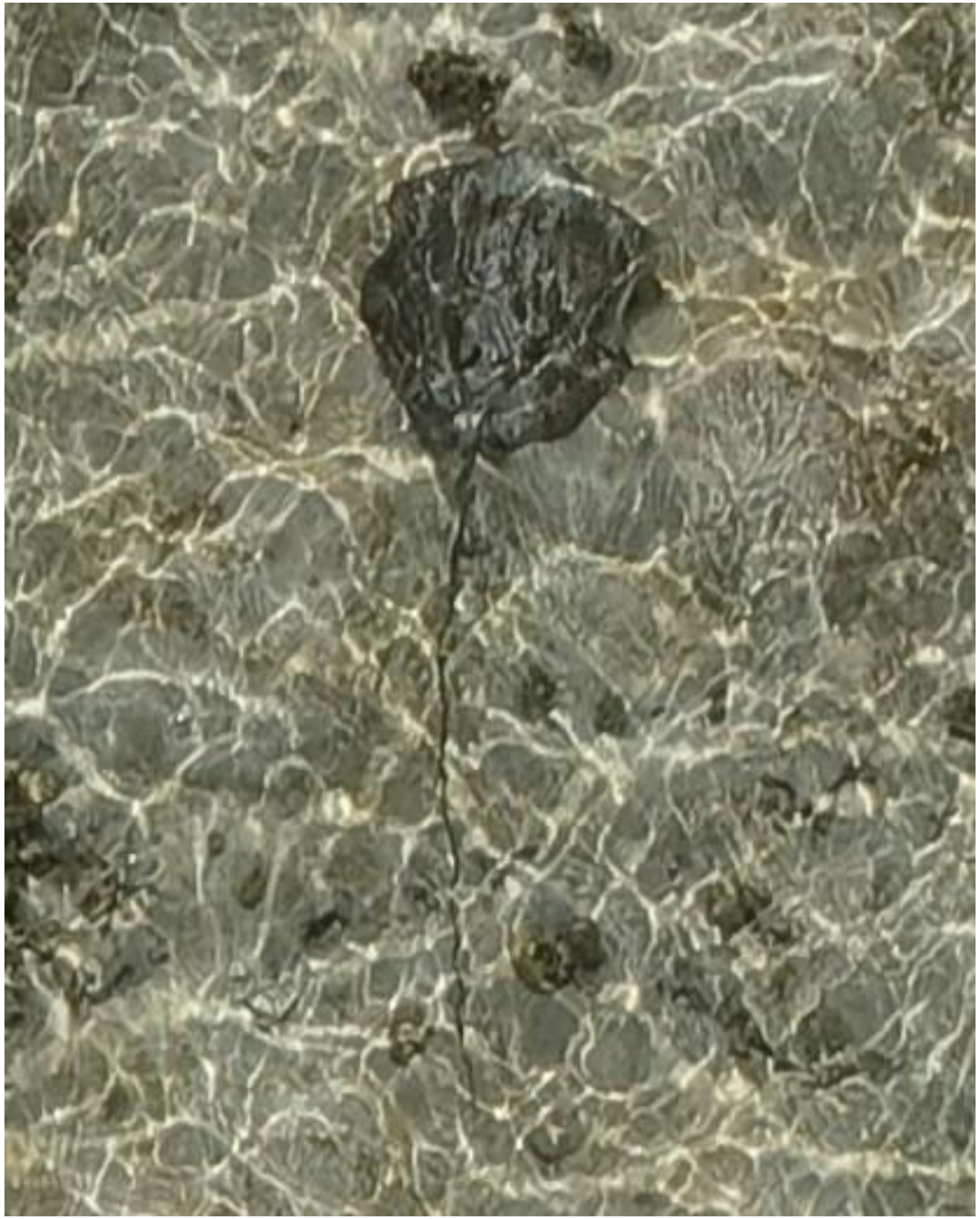

DJI_20240814085734_0393

- Whipray

2024-08-18

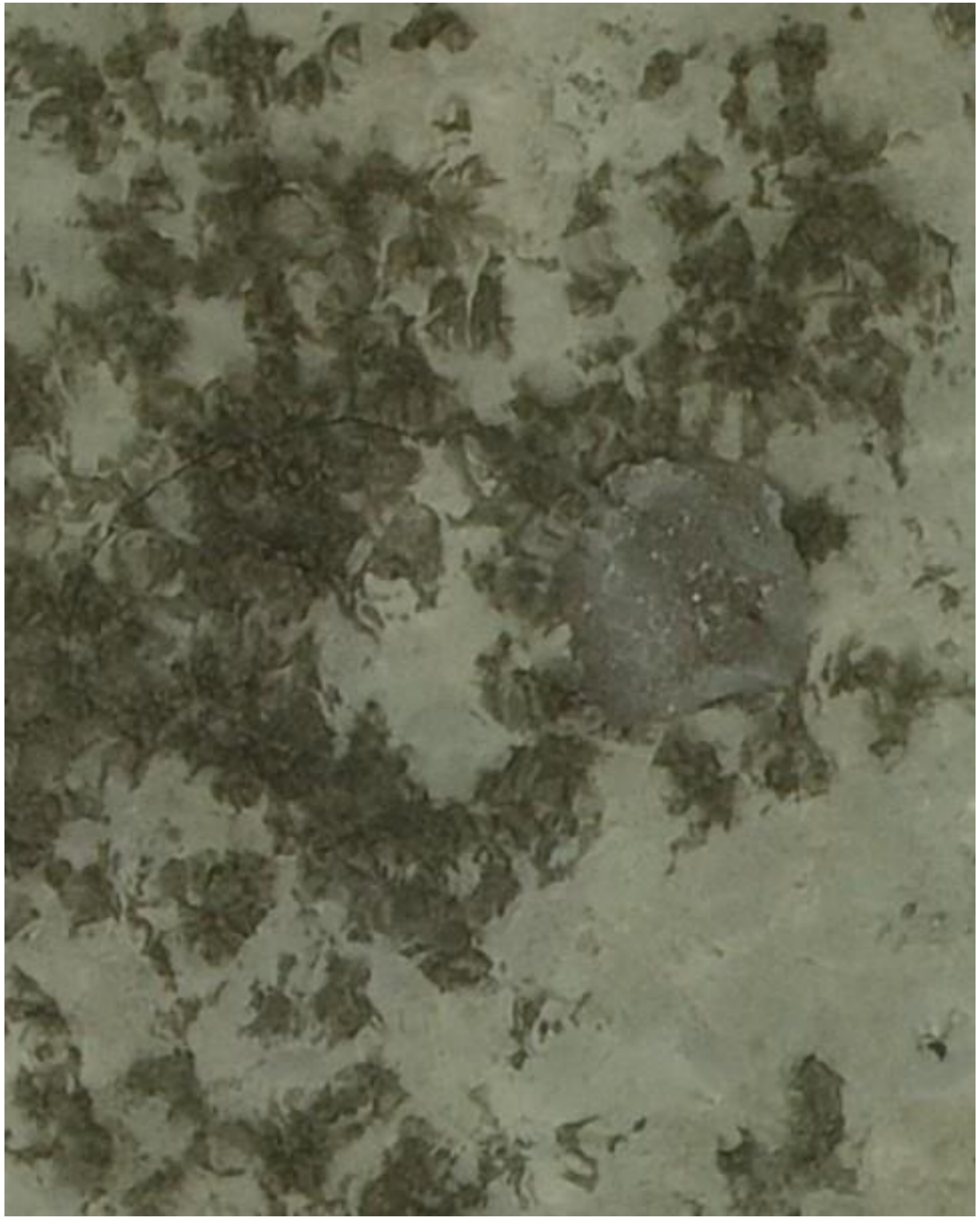

- Whipray

DJI_20240818080133_0116

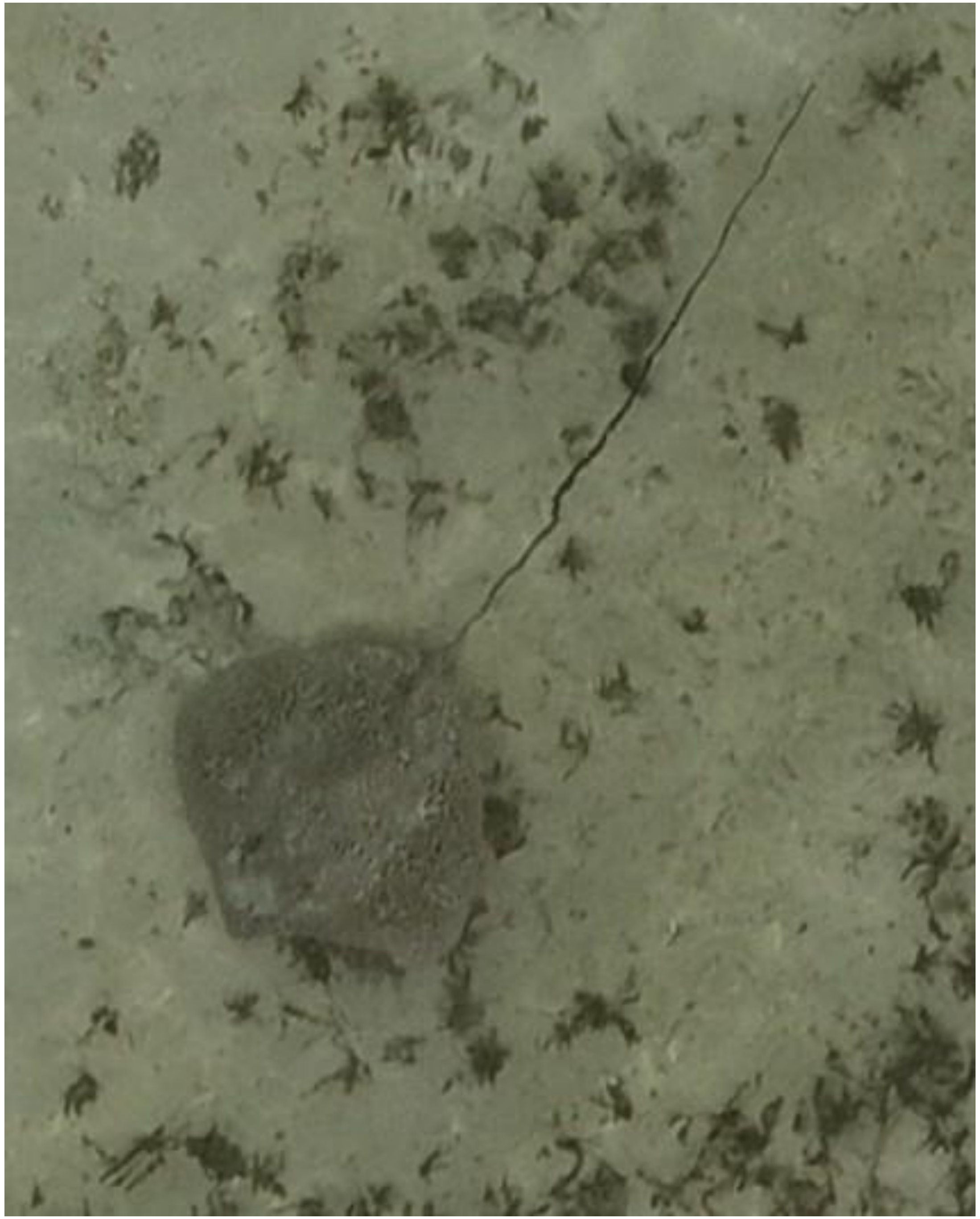

DJI_20240818080442_0281

- *Himantura uarnak*

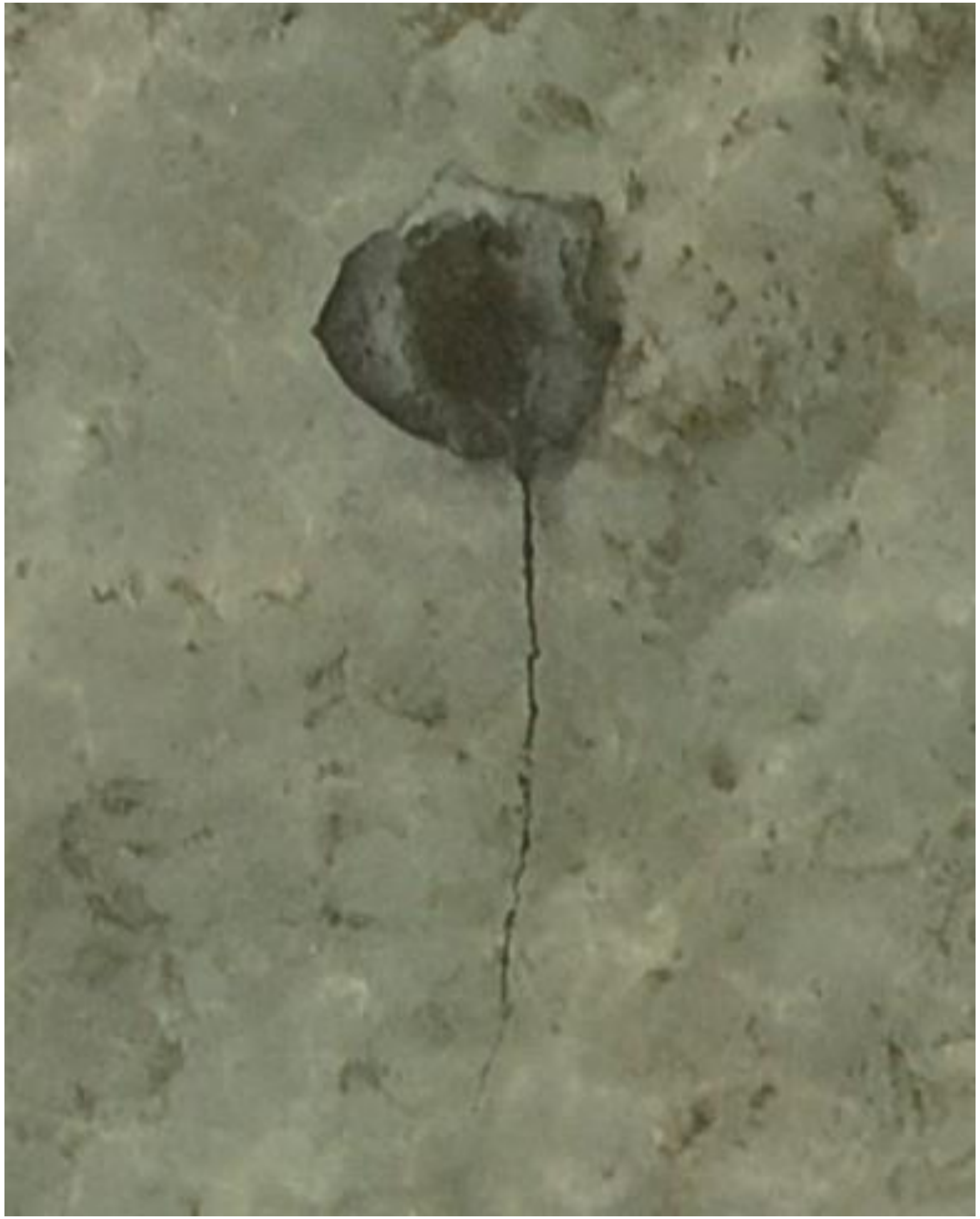

- Whipray

DJI_20240818080712_0419

2024-08-24

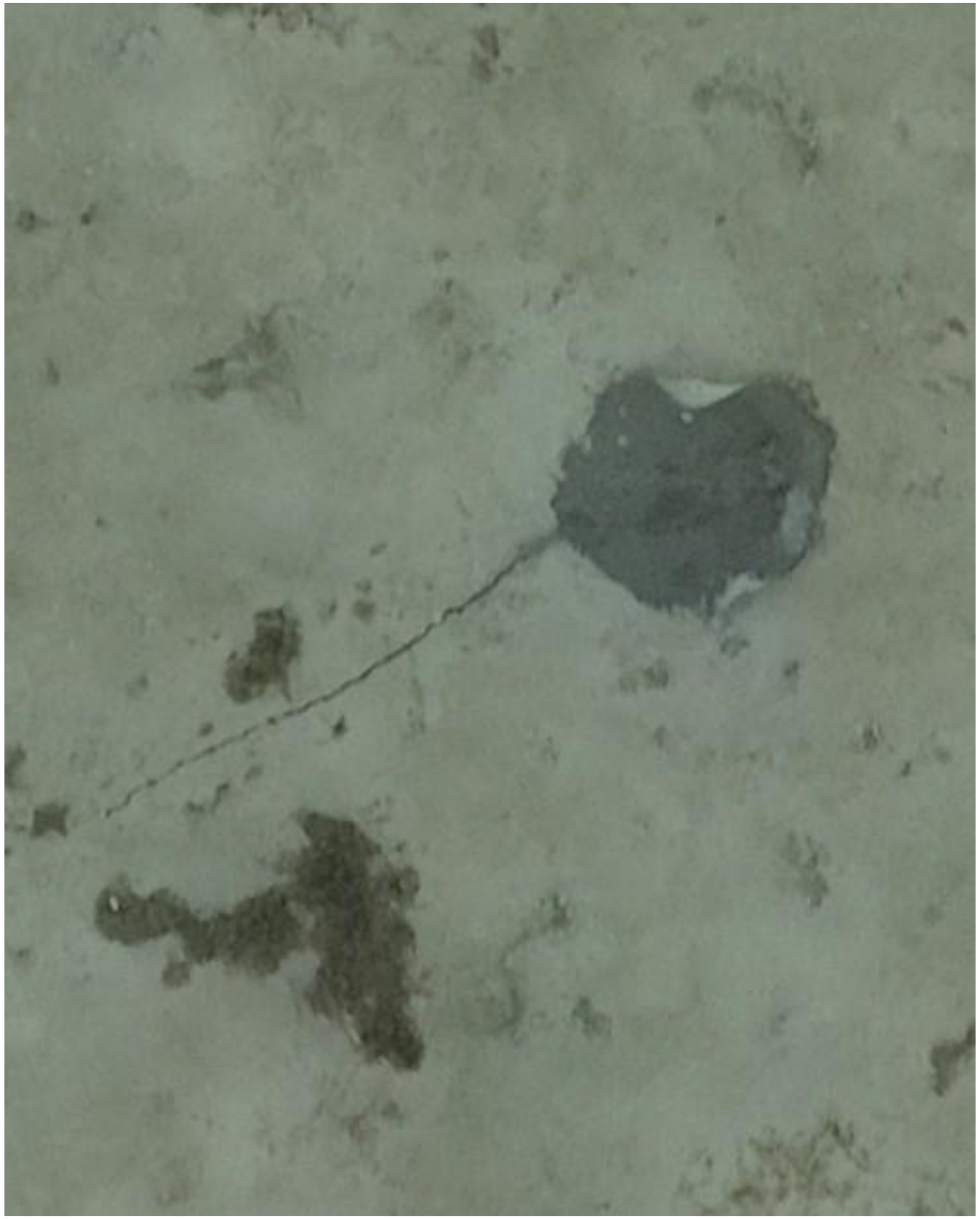

DJI_20240824073510_0035

- Whipray

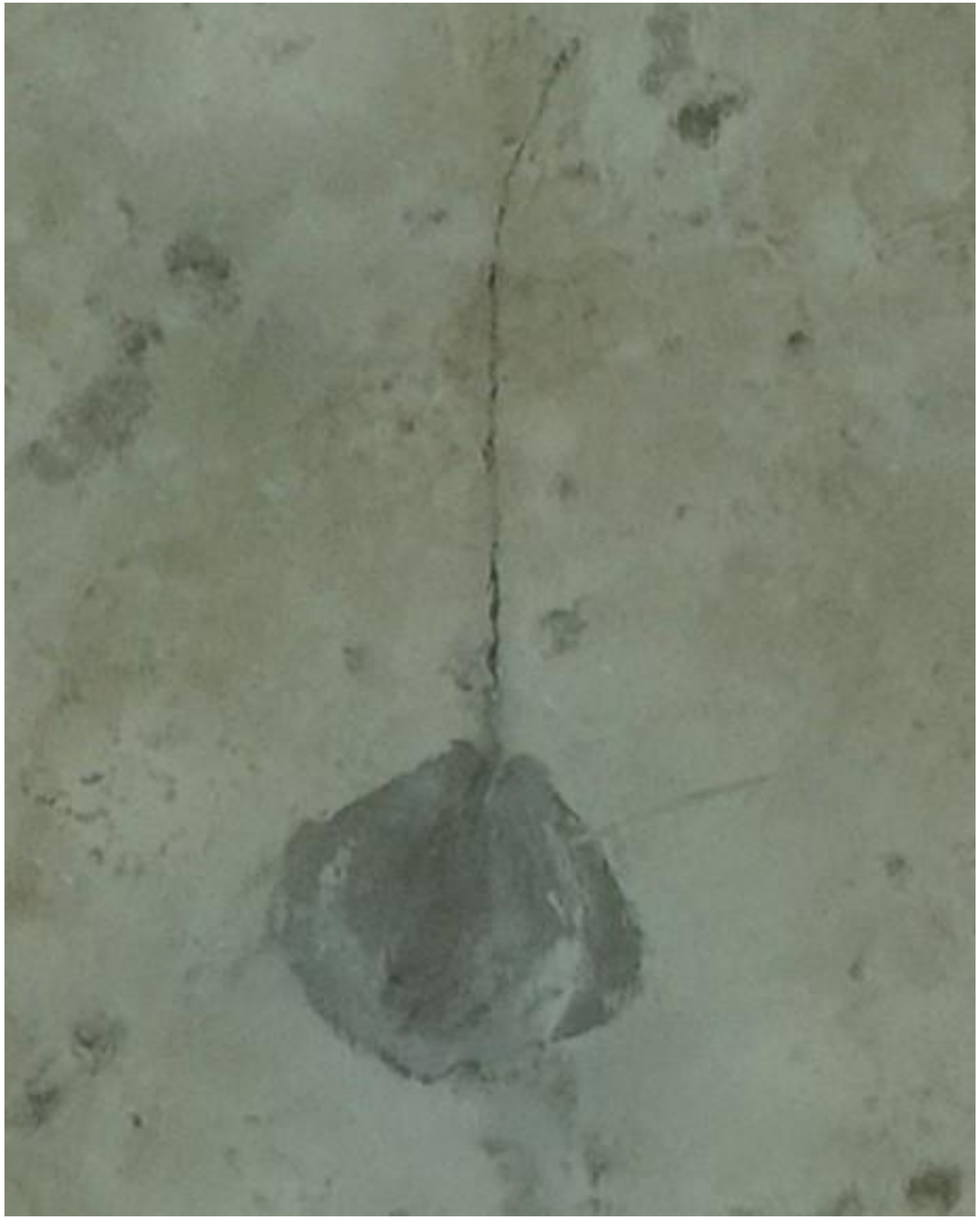

DJI_20240824073656_0128

- Whipray
- Accompanied by needlefish

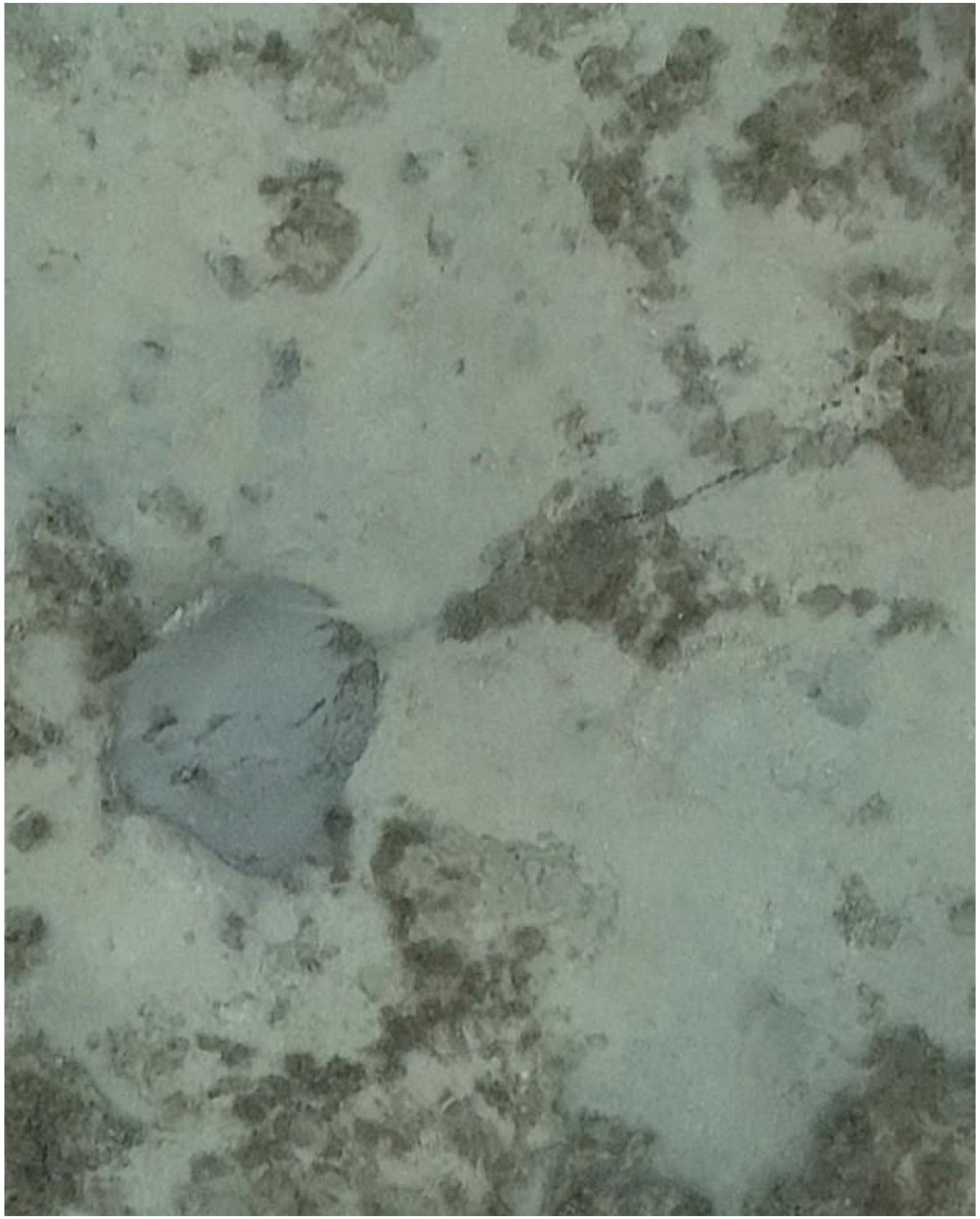

DJI_20240824074525_0569

- Whipray

2024-09-20

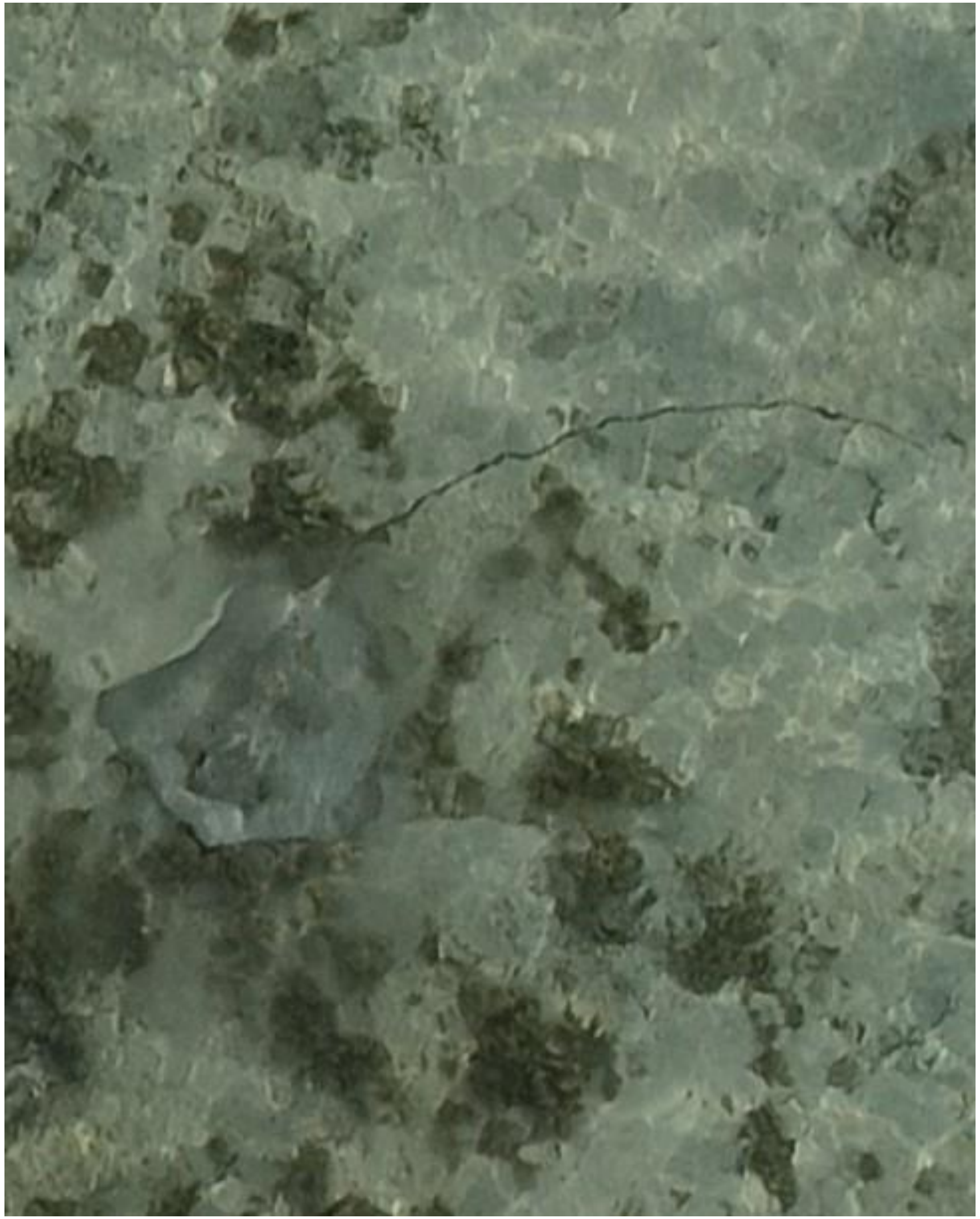

DJI_20240920075430_0039

- Whipray

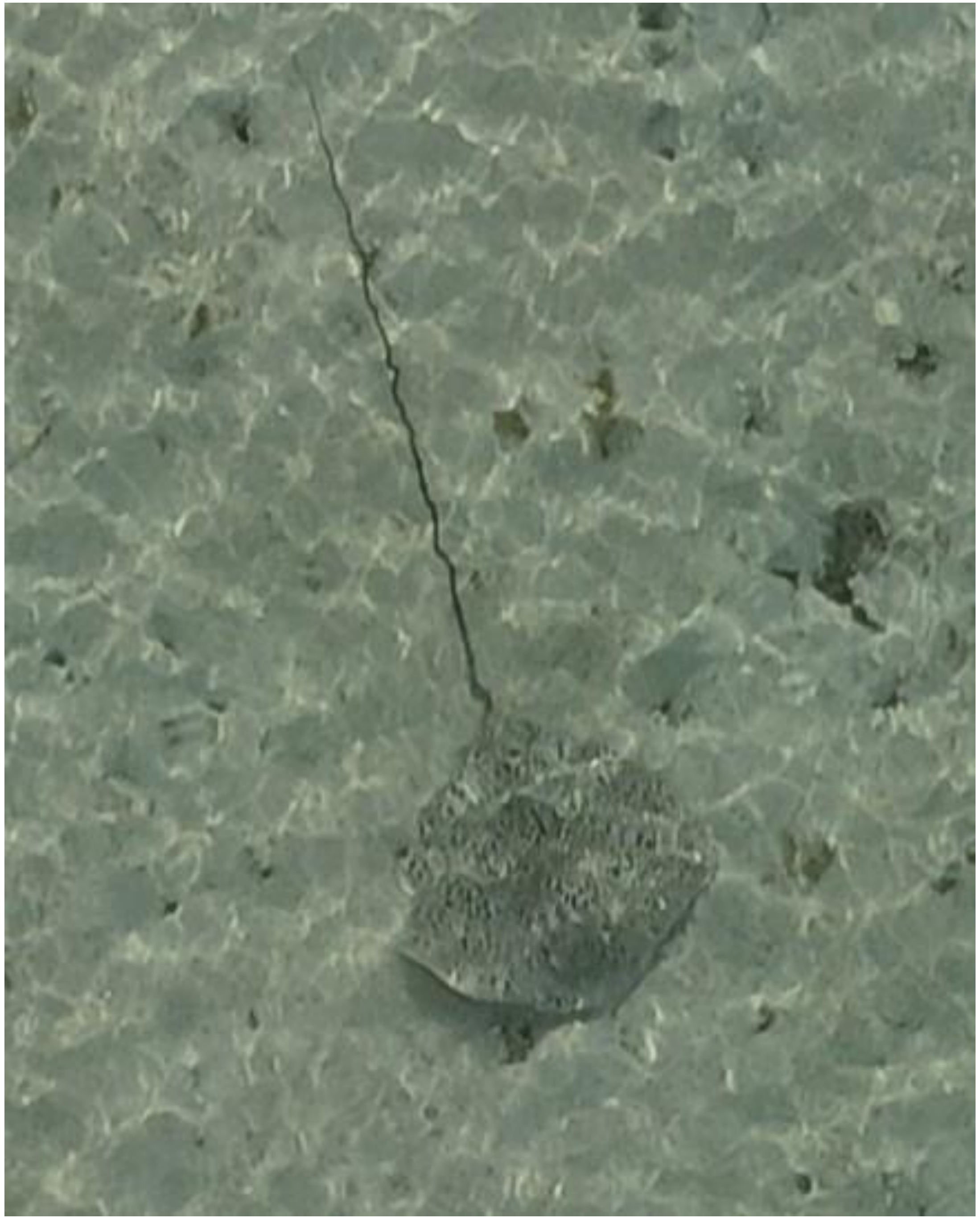

DJI_20240920075439_0048

- *Himantura uarnak*

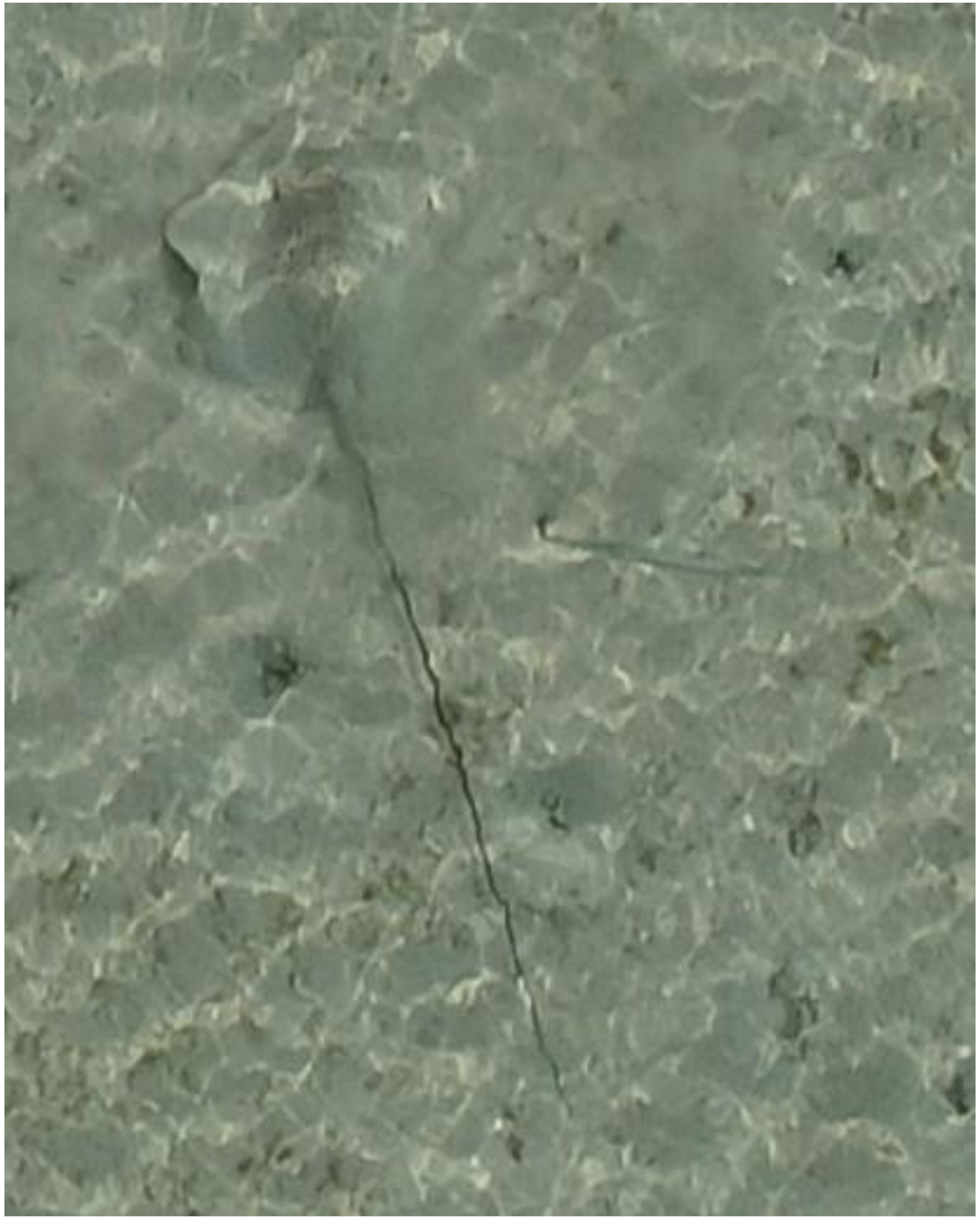

DJI_20240920075447_0055

- Whipray
- Best guess: *H. uarnak*
- Accompanied by needlefish

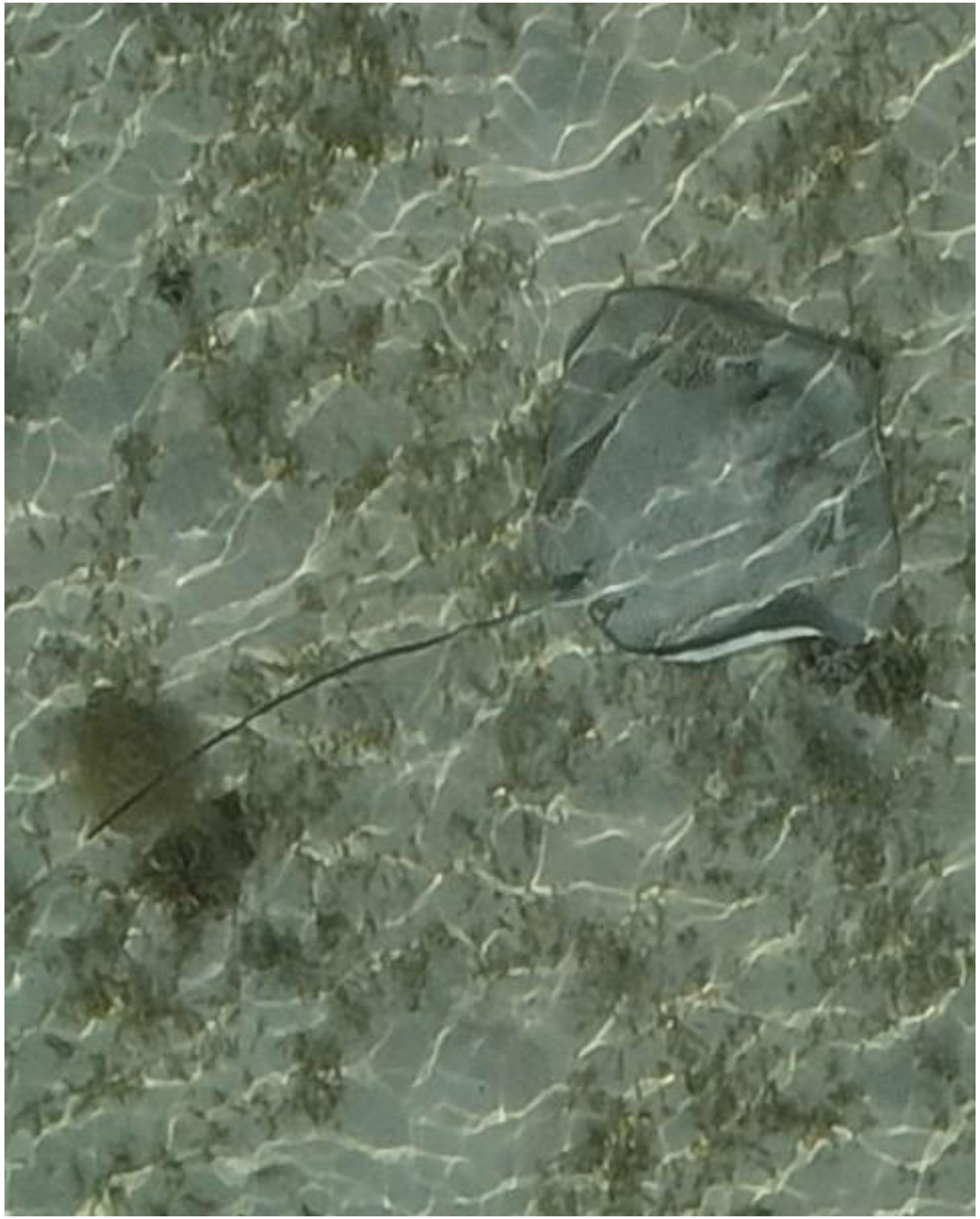

- *Himantura uarnak*

DJI_20240920075536_0093

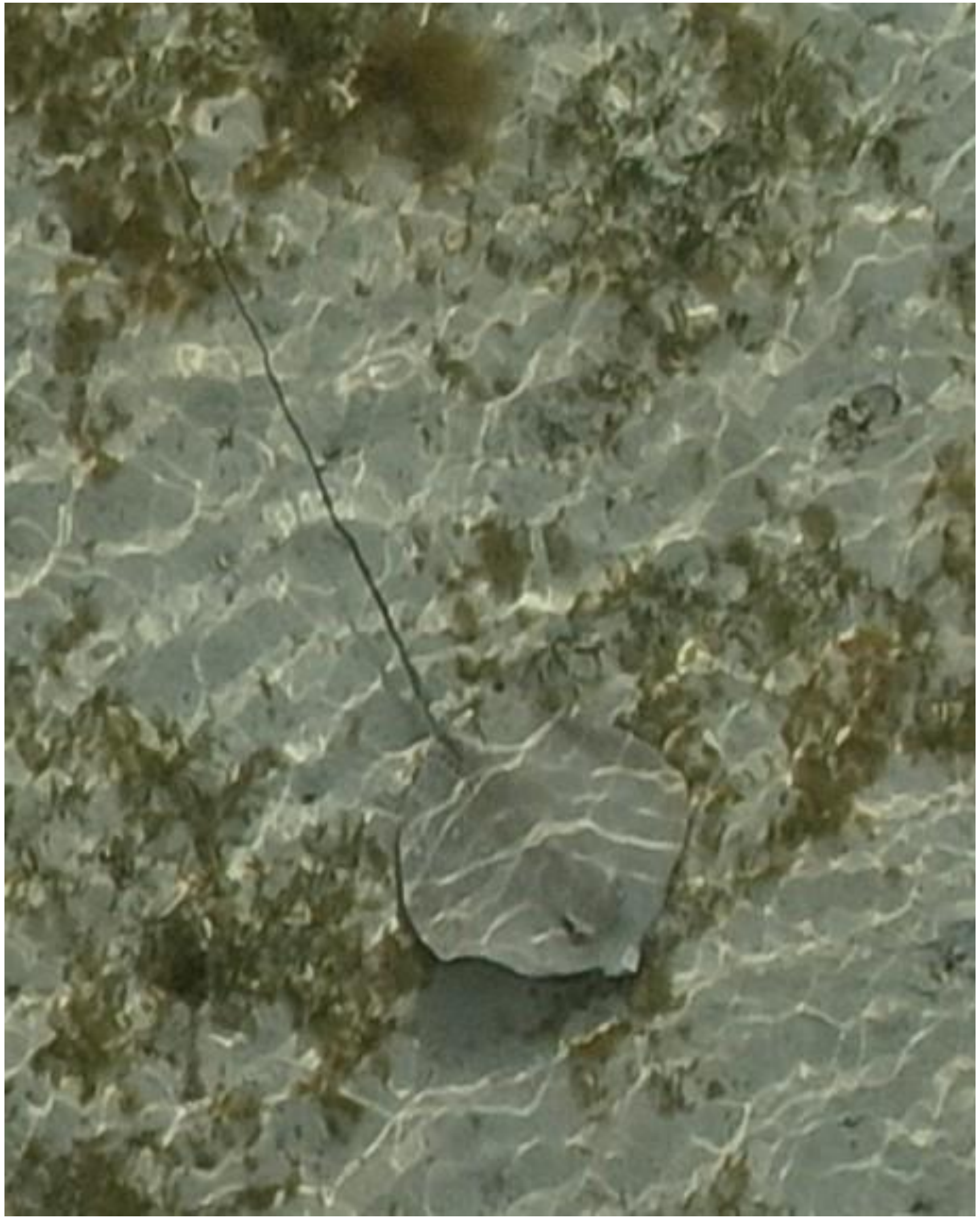

- Whipray

DJI_20240920075736_0201

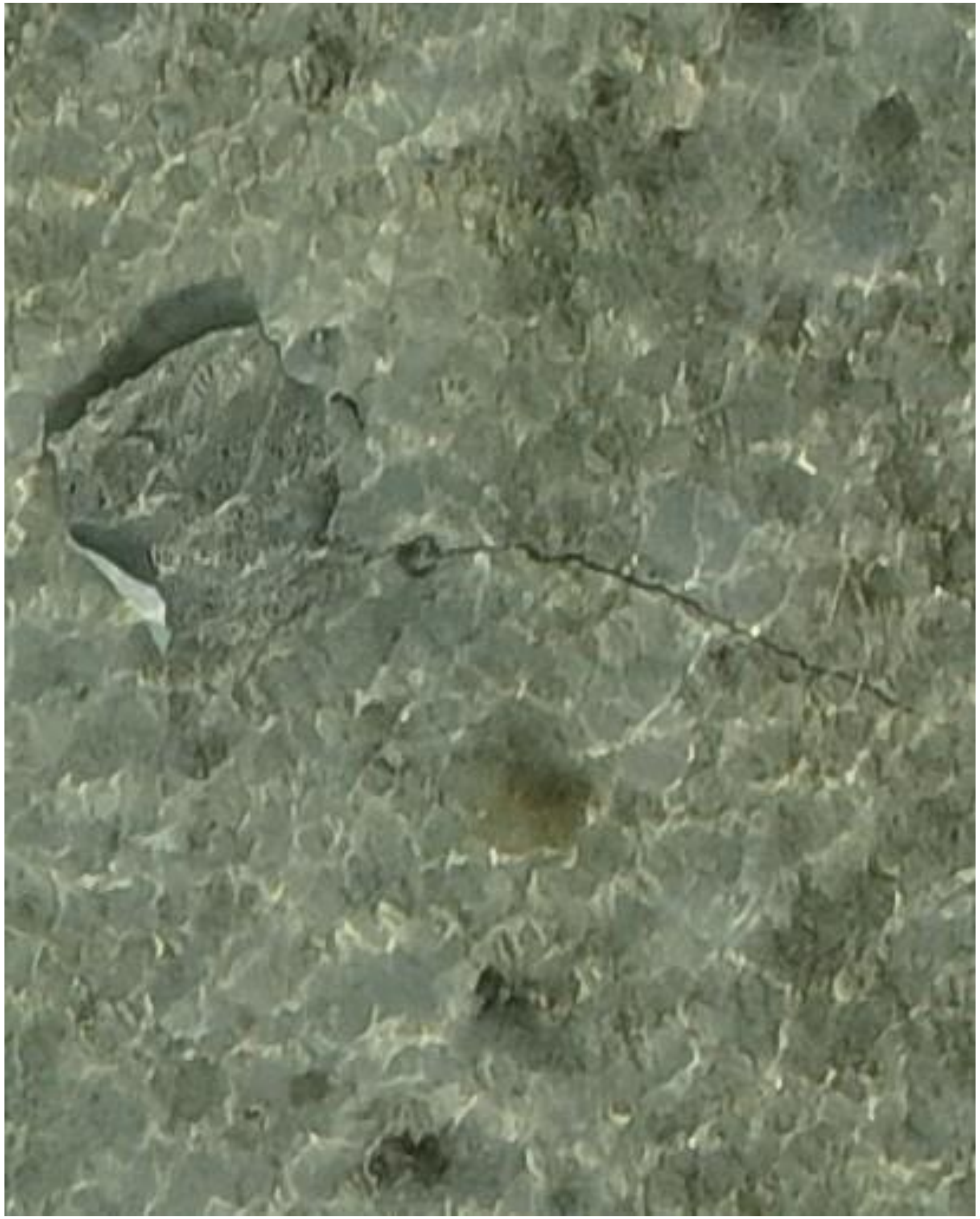

DJI_20240920075859_0272

- *Himantura uarnak*

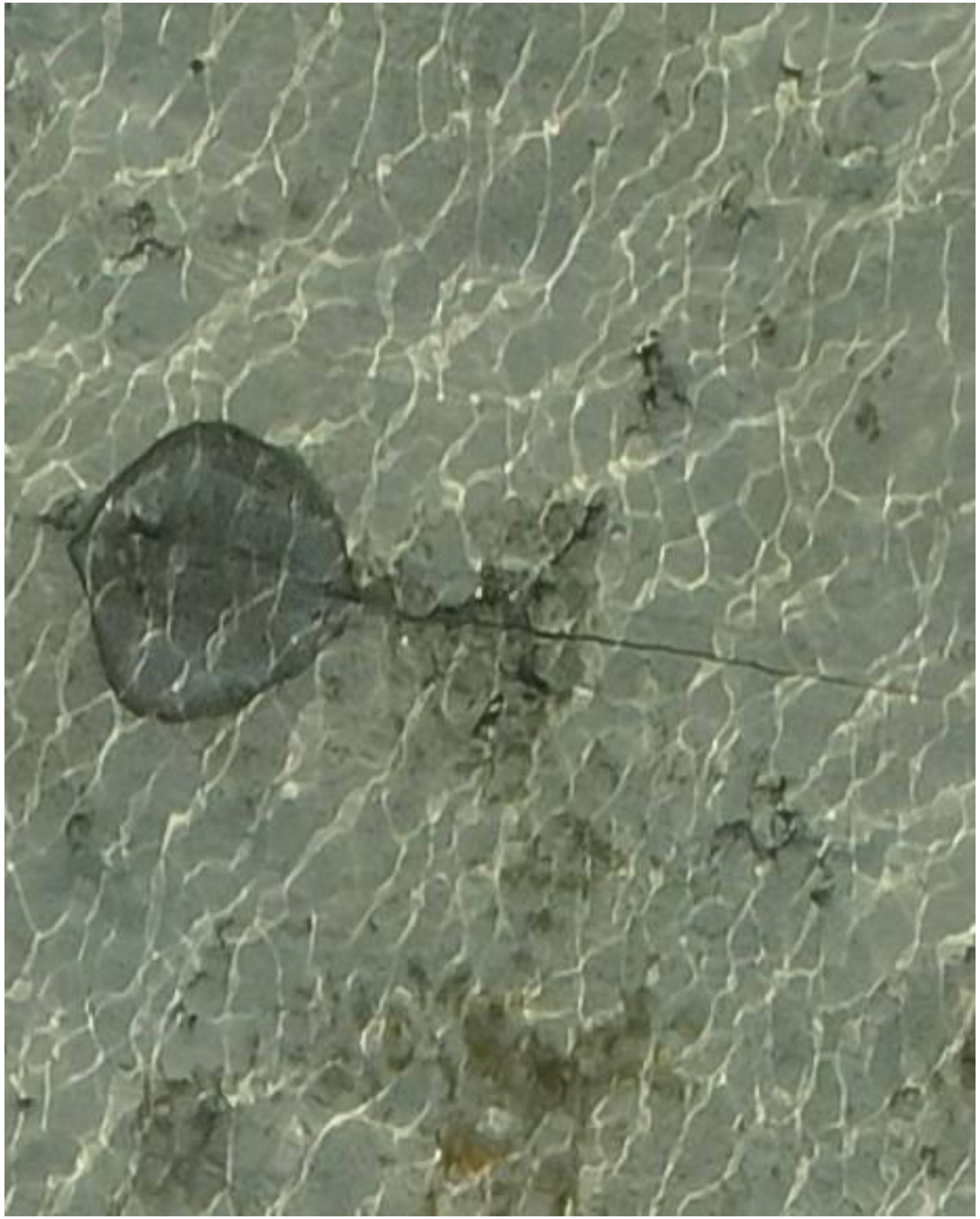

DJI_20240920075924_0297

- Whipray

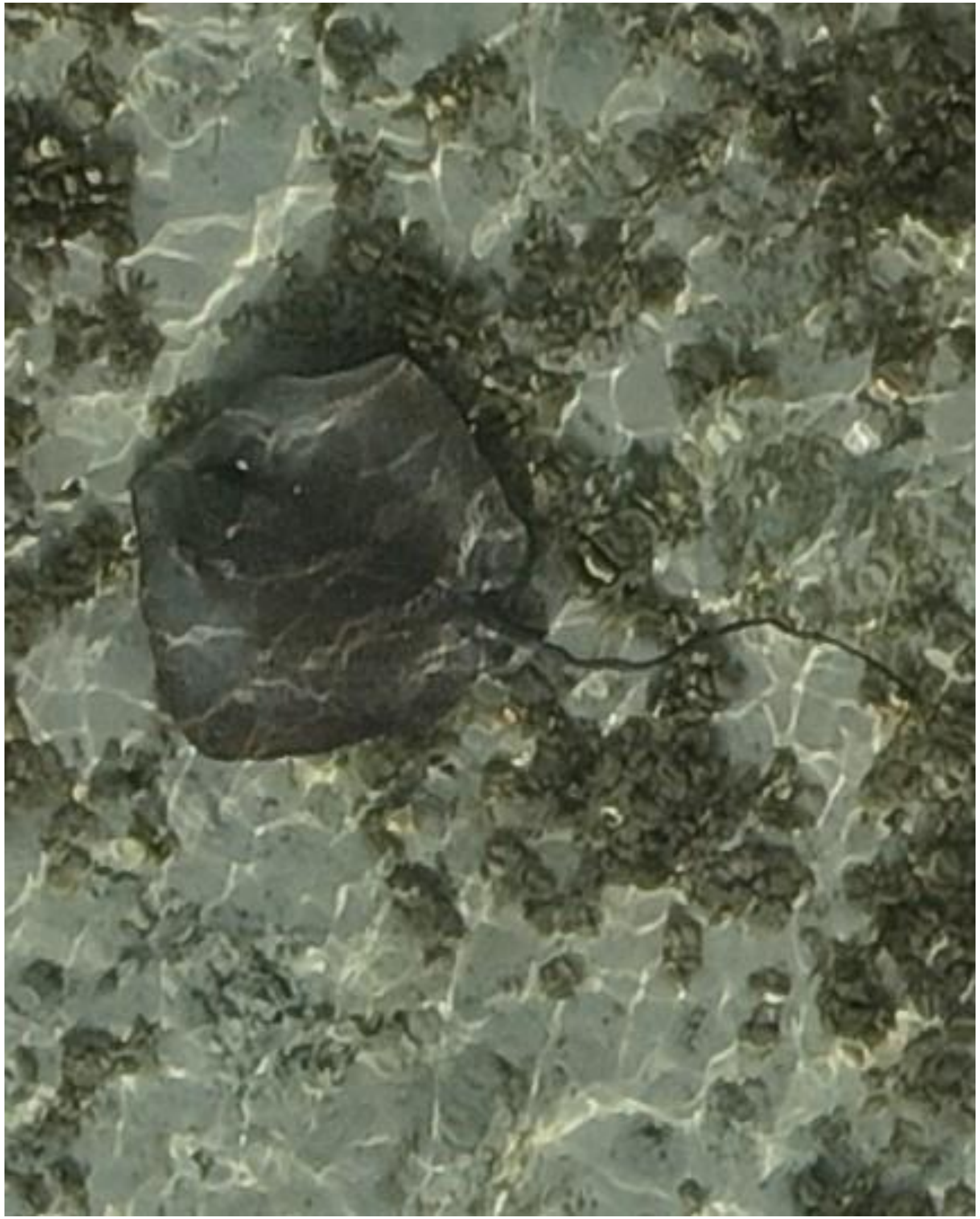

- Whipray
- Best guess: *H. uarnak*

DJI_20240920075949_0322

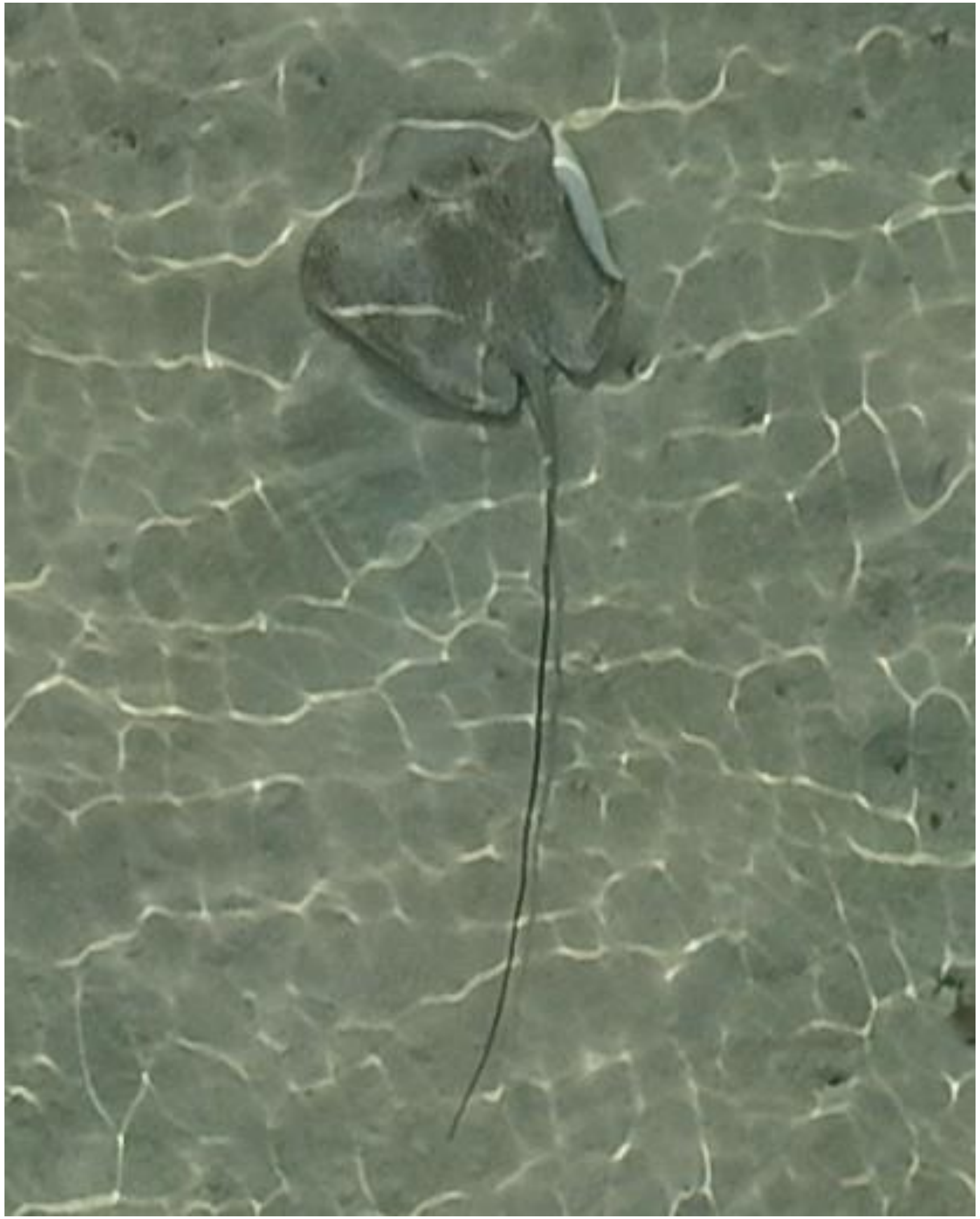

DJI_20240920080150_0431

- *Himantura uarnak*

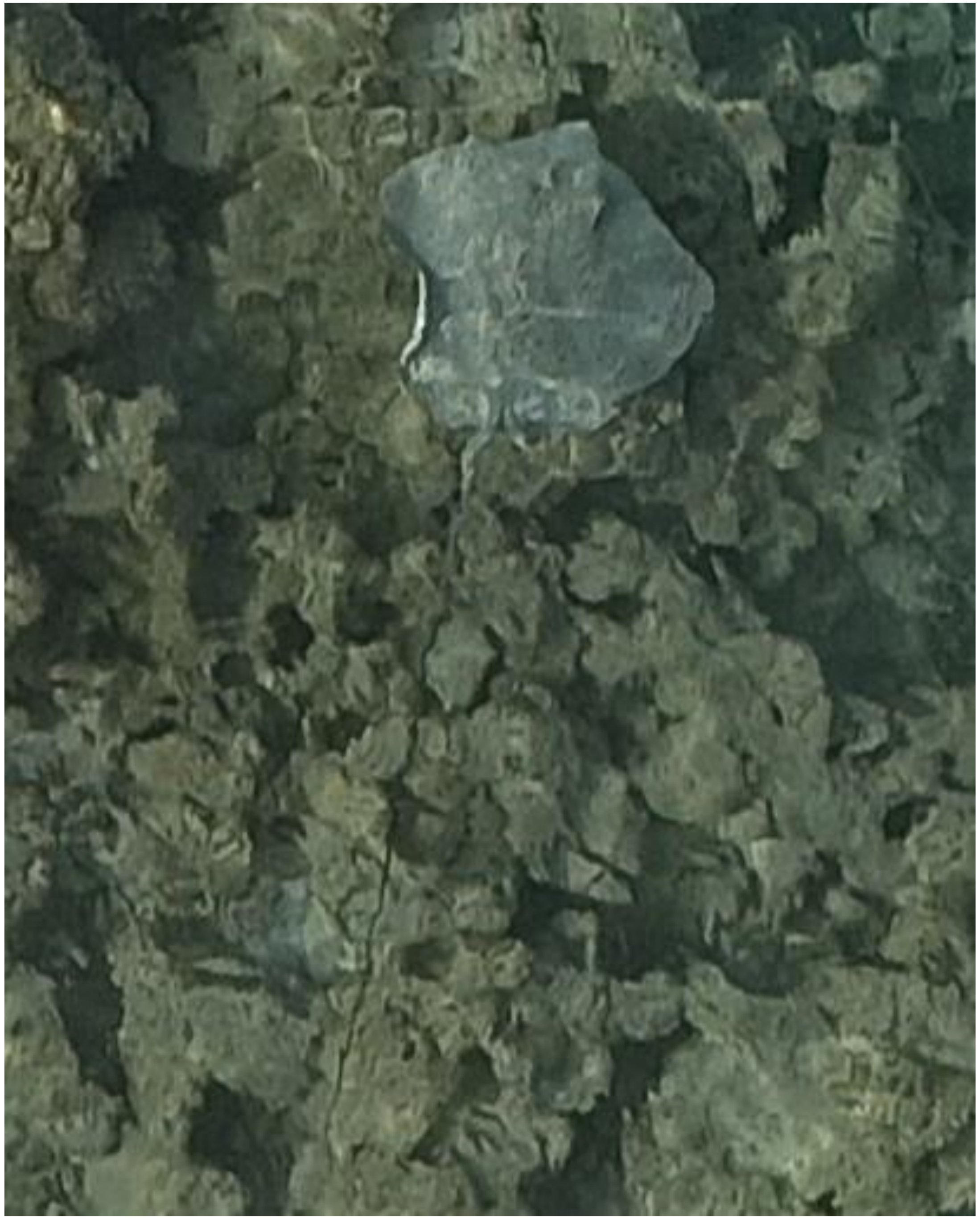

- *Himantura uarnak*

DJI_20240920080344_0517

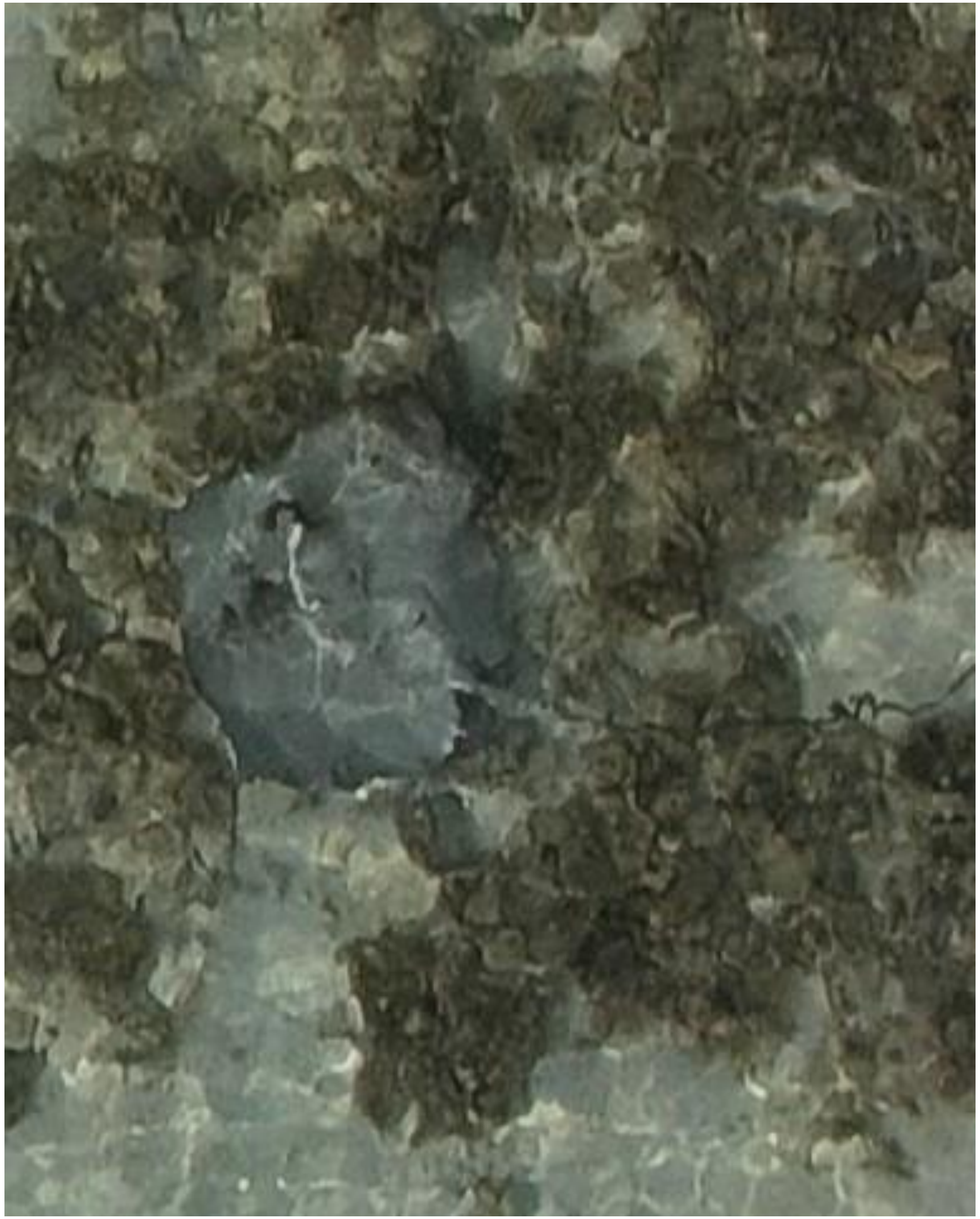

DJI_20240920080450_0583

- Whipray
- Also observed on 05-02, 06-06, 06-24, and 07-29 as identified by the coiled tail.
- Best guess: *H. uarnak*

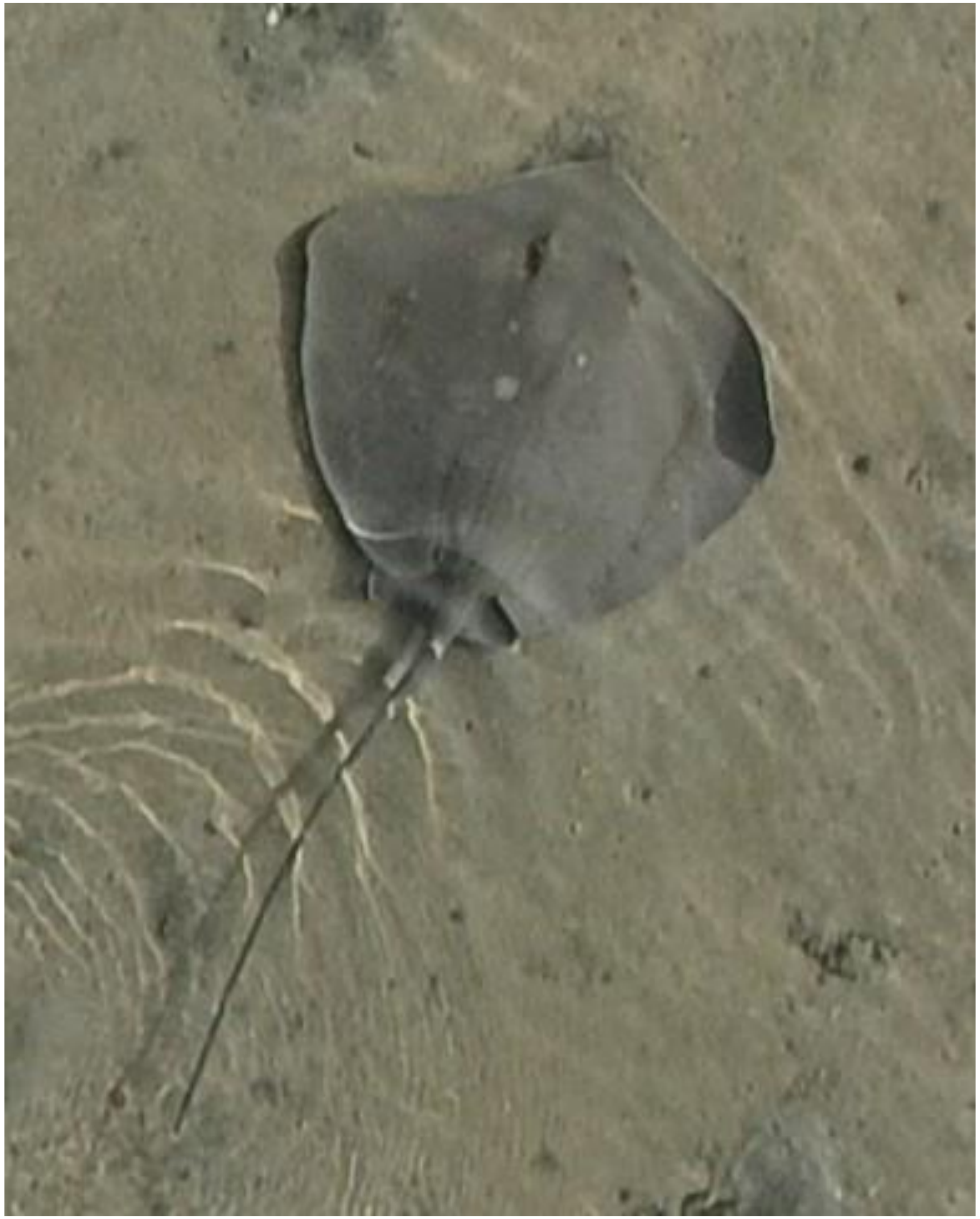

DJI_20240920080720_0716

- Whipray
- Somewhat resembles *Maculabatis randalli*. Individual was also observed on 06-03.

2024-09-27

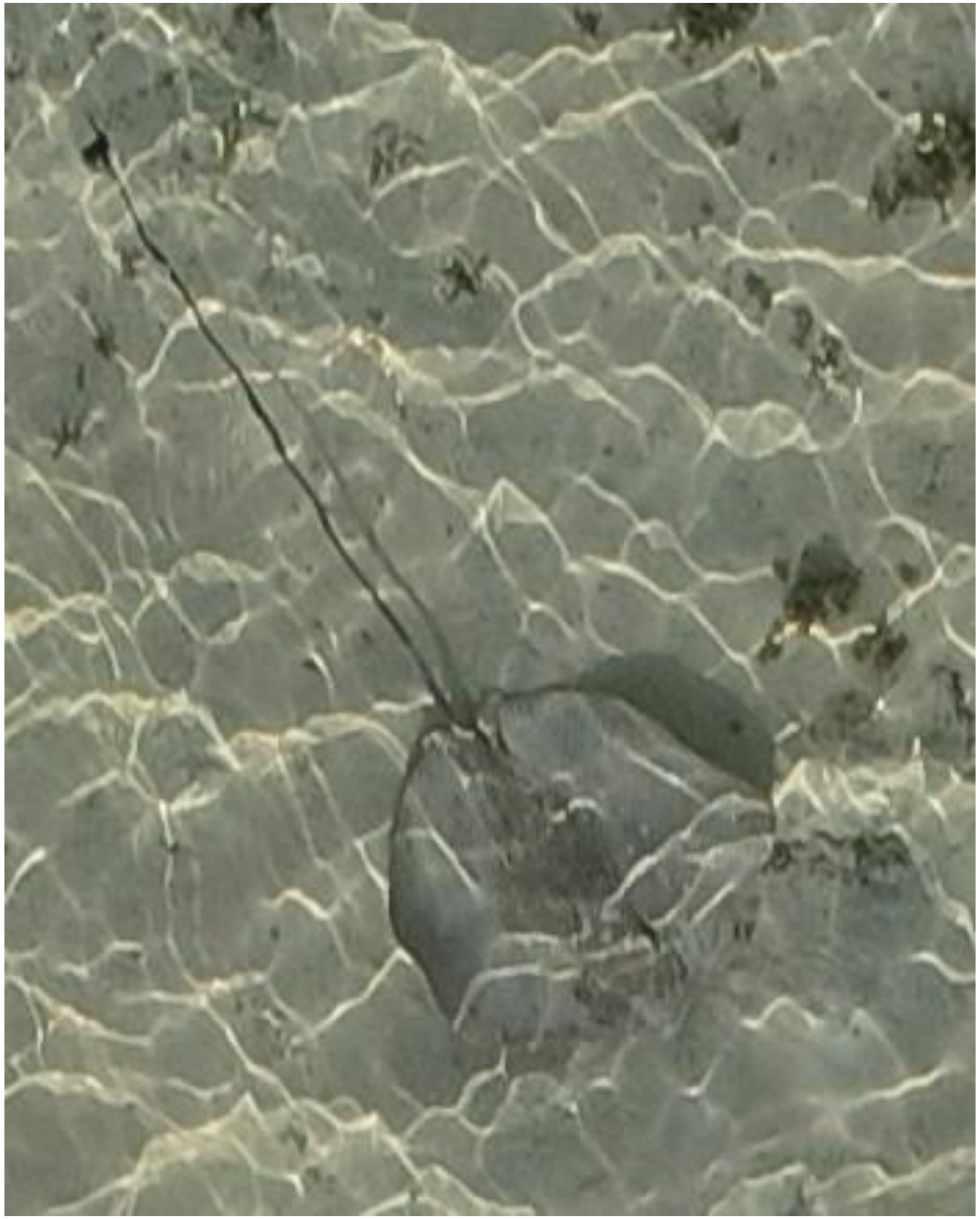
DJI_20240927075429_0277

- Whipray

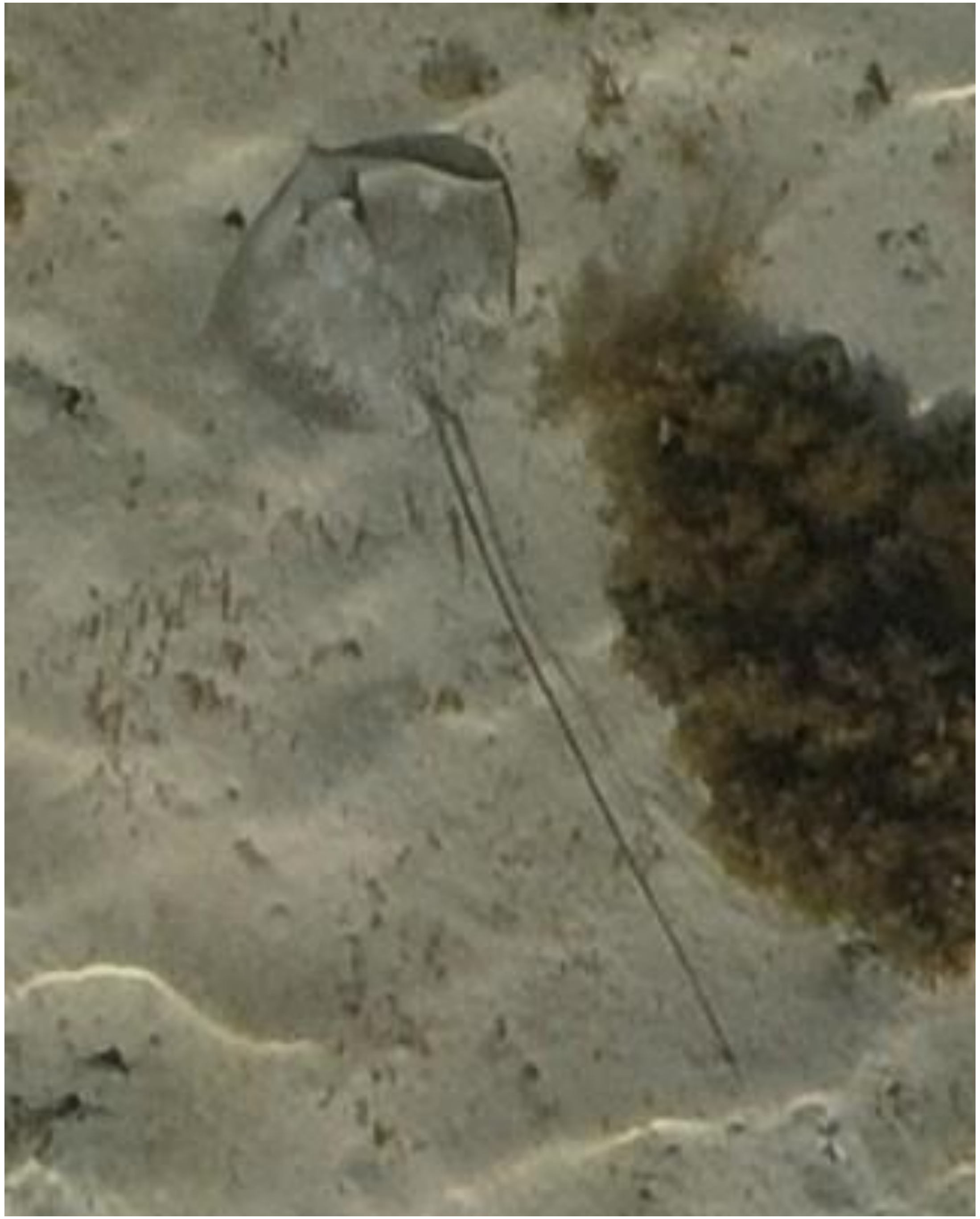

DJI_20240927075457_0305

- *Himantura uarnak*

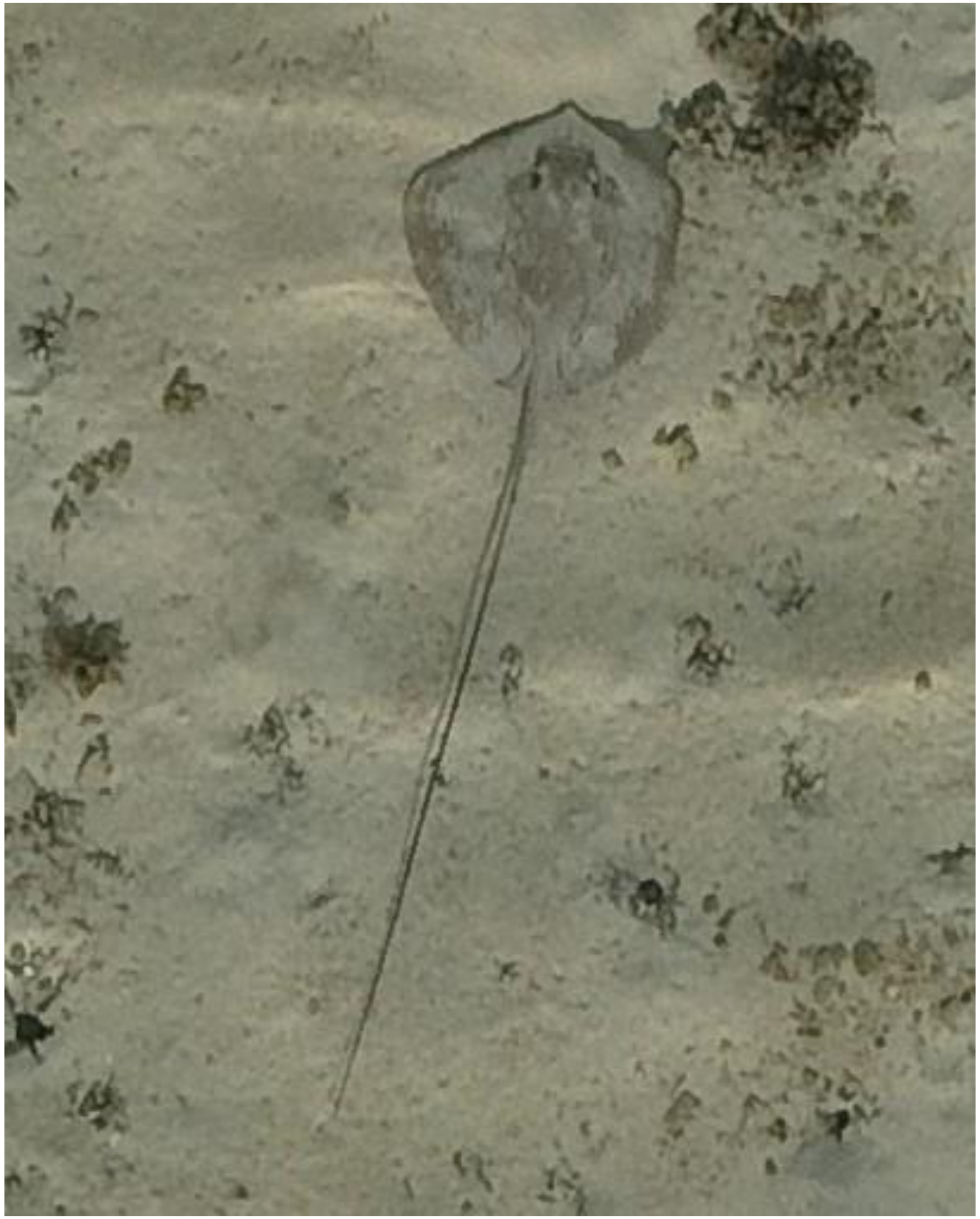
DJI_20240927075504_0312

- Whipray

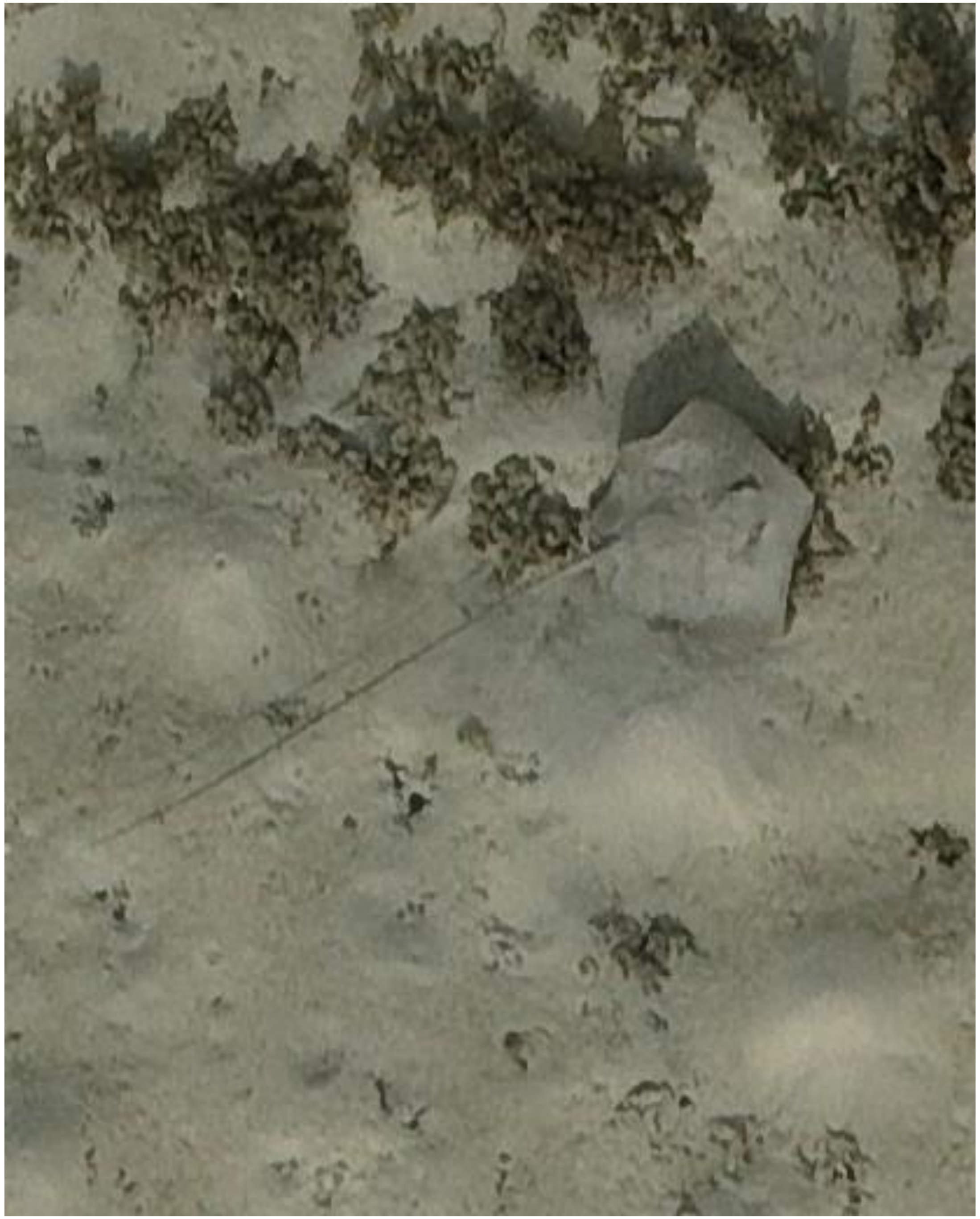

- Whipray

DJI_20240927075506_0314

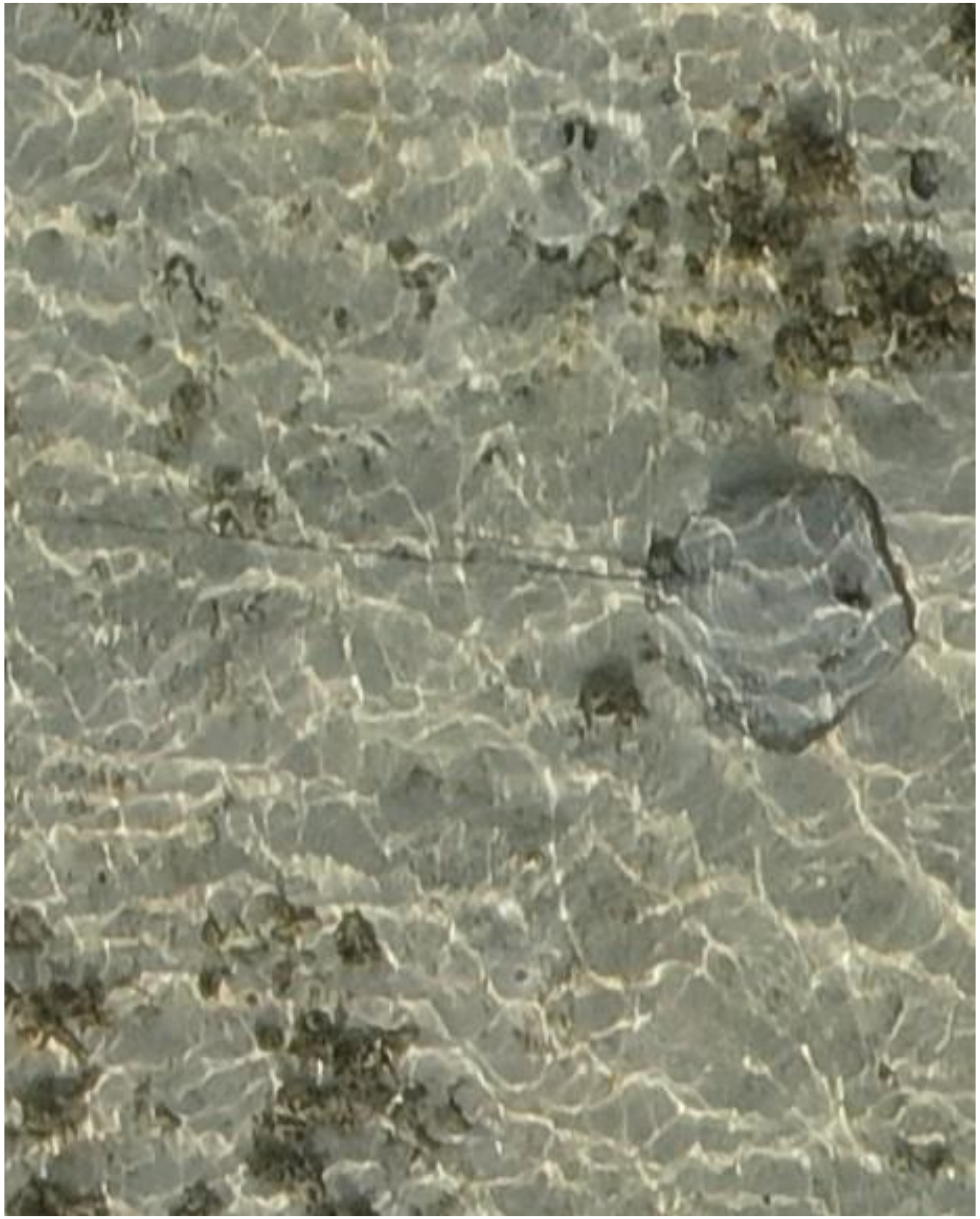

- Whipray

DJI_20240927075514_0322

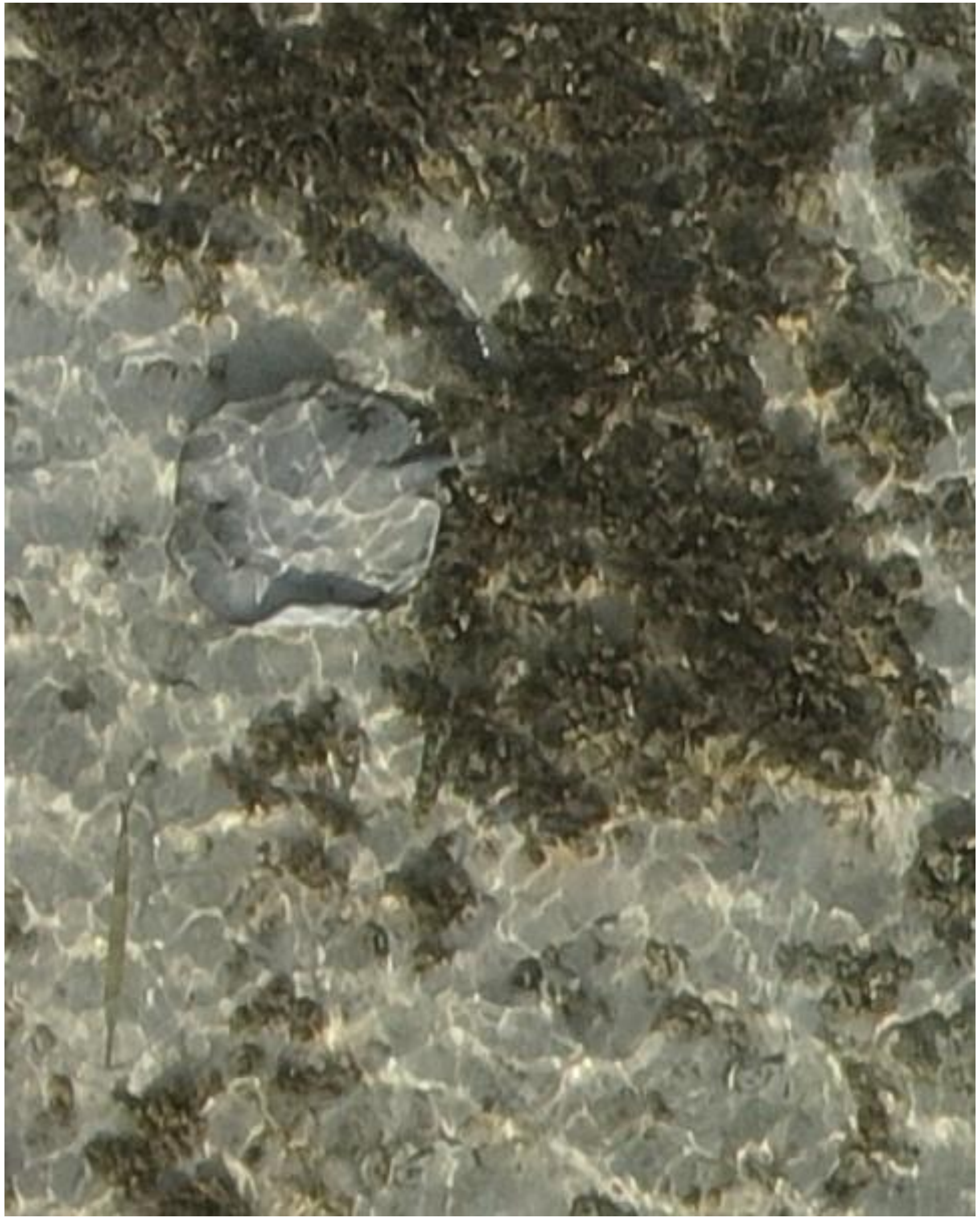

- Whipray
- Accompanied by needlefish

DJI_20240927075517_0325

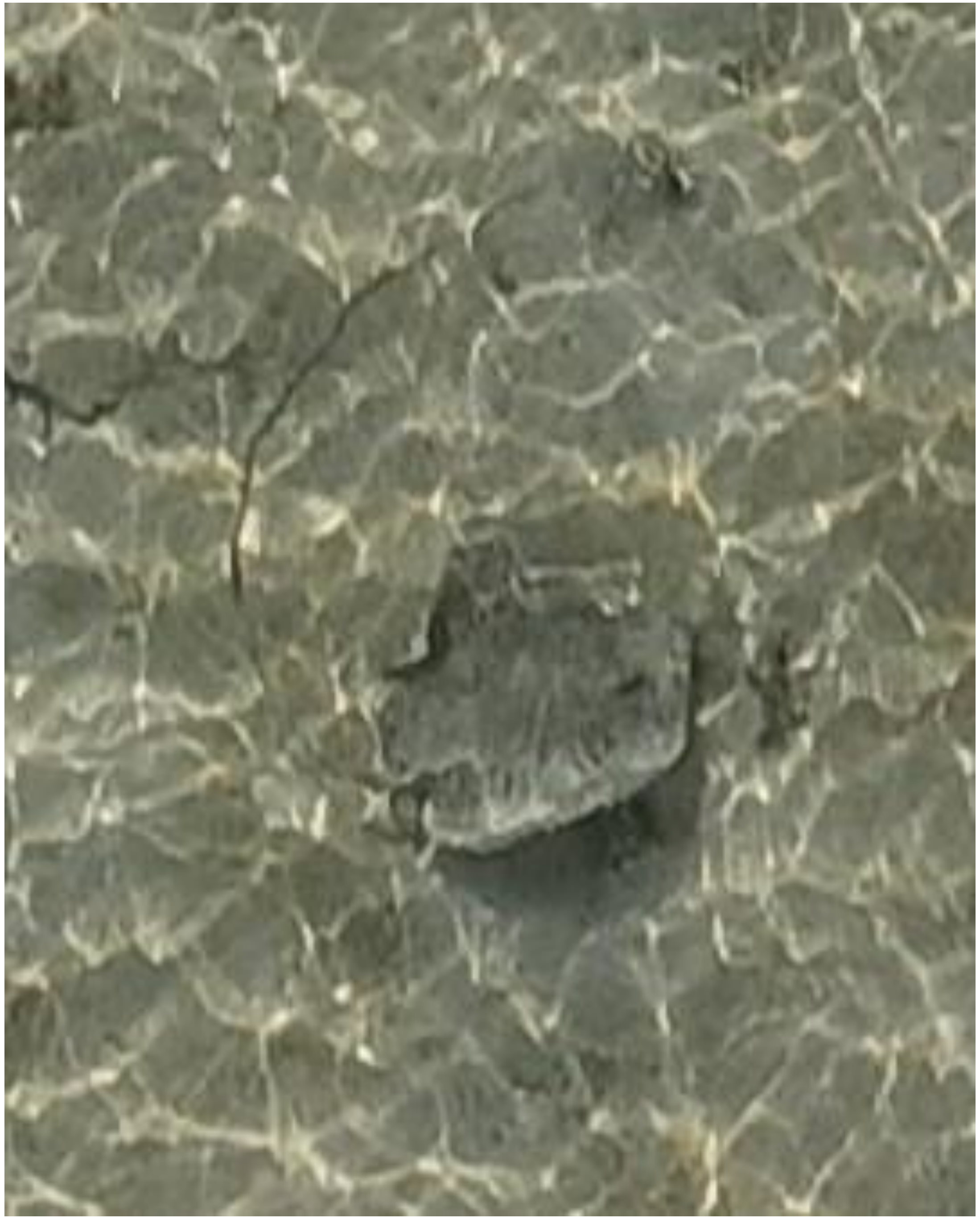

- Whipray
- Best guess: *H. uarnak*

DJI_20240927075638_0394

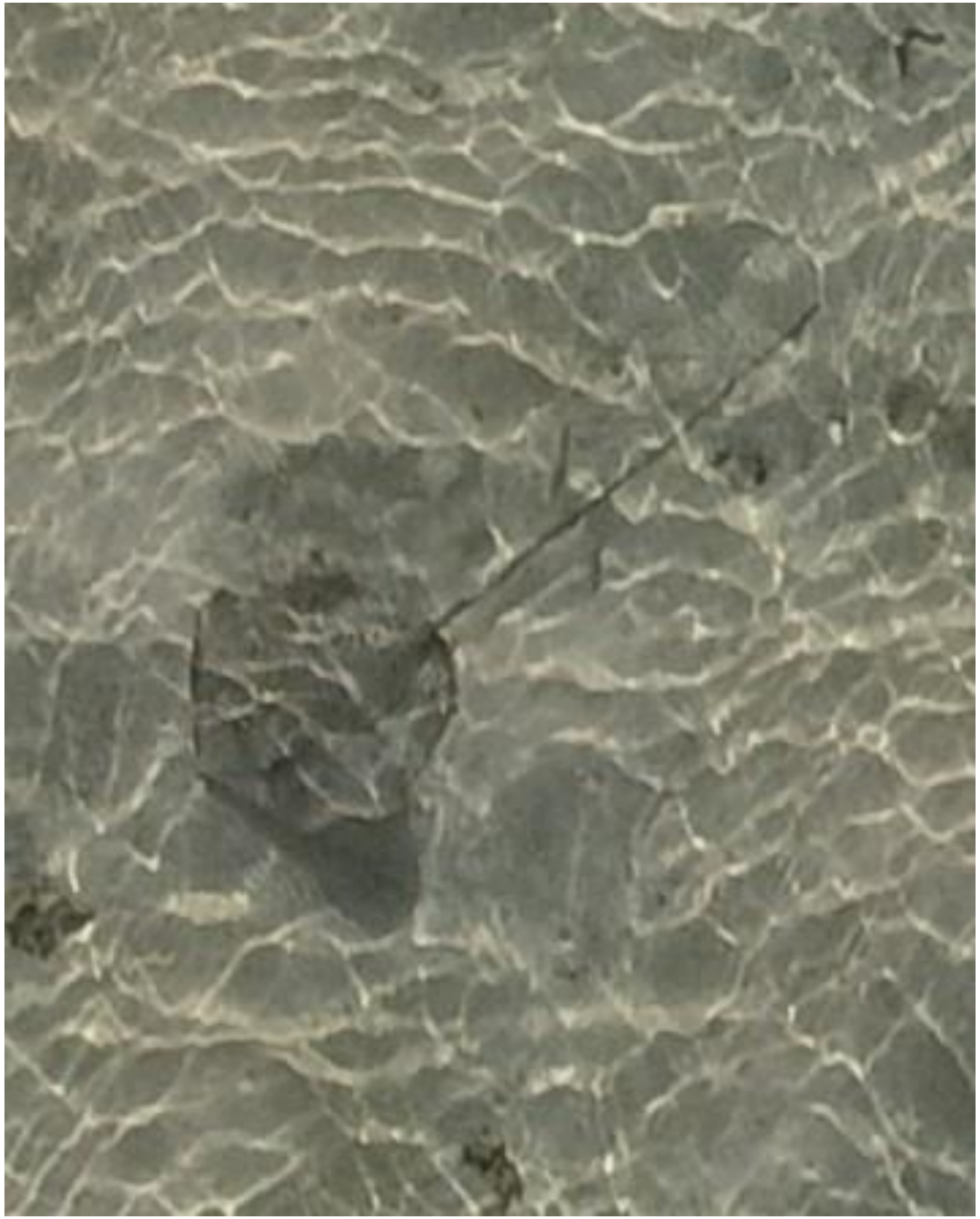
DJI_20240927075651_0407

- Whipray

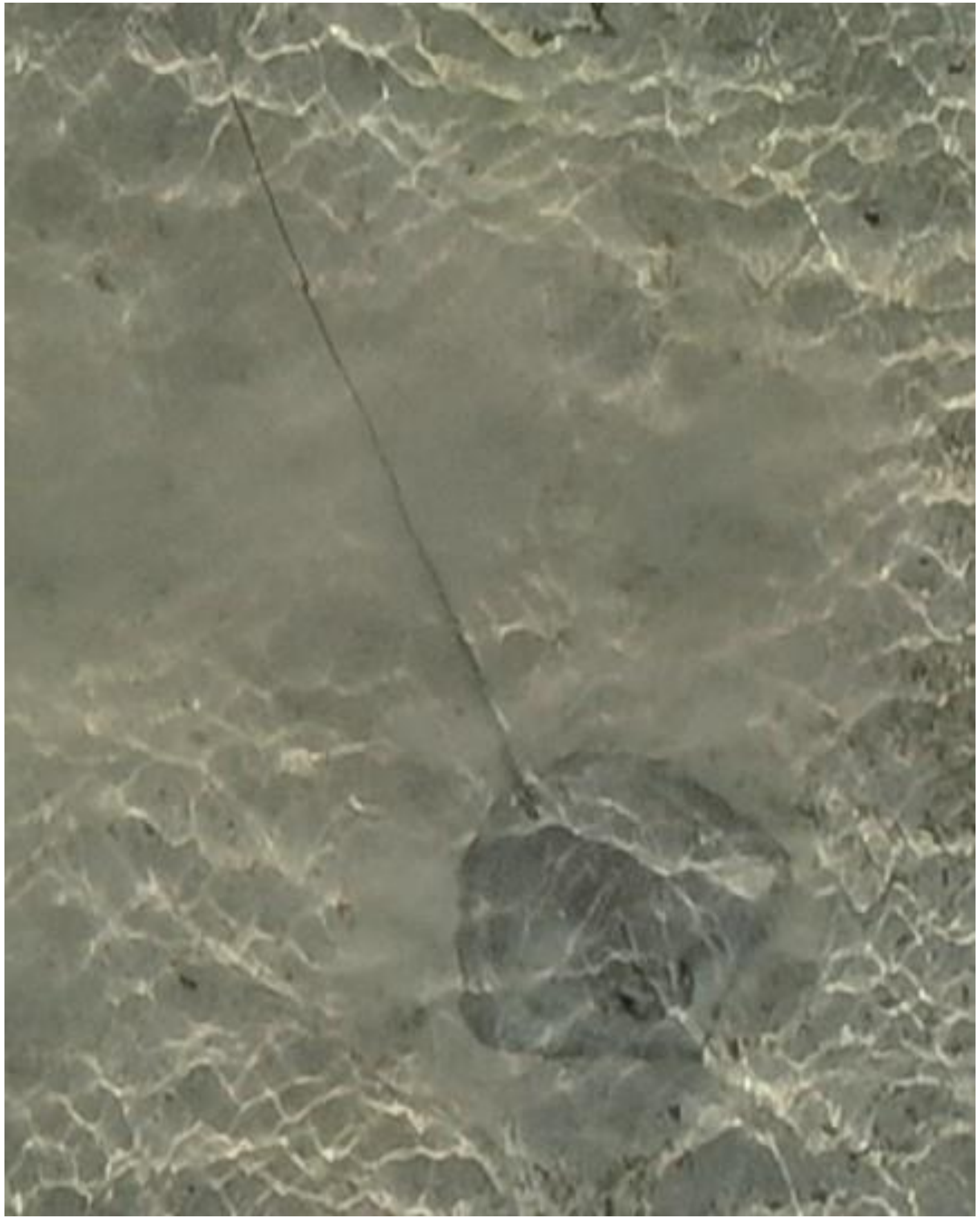

DJI_20240927075652_0408

- Whipray

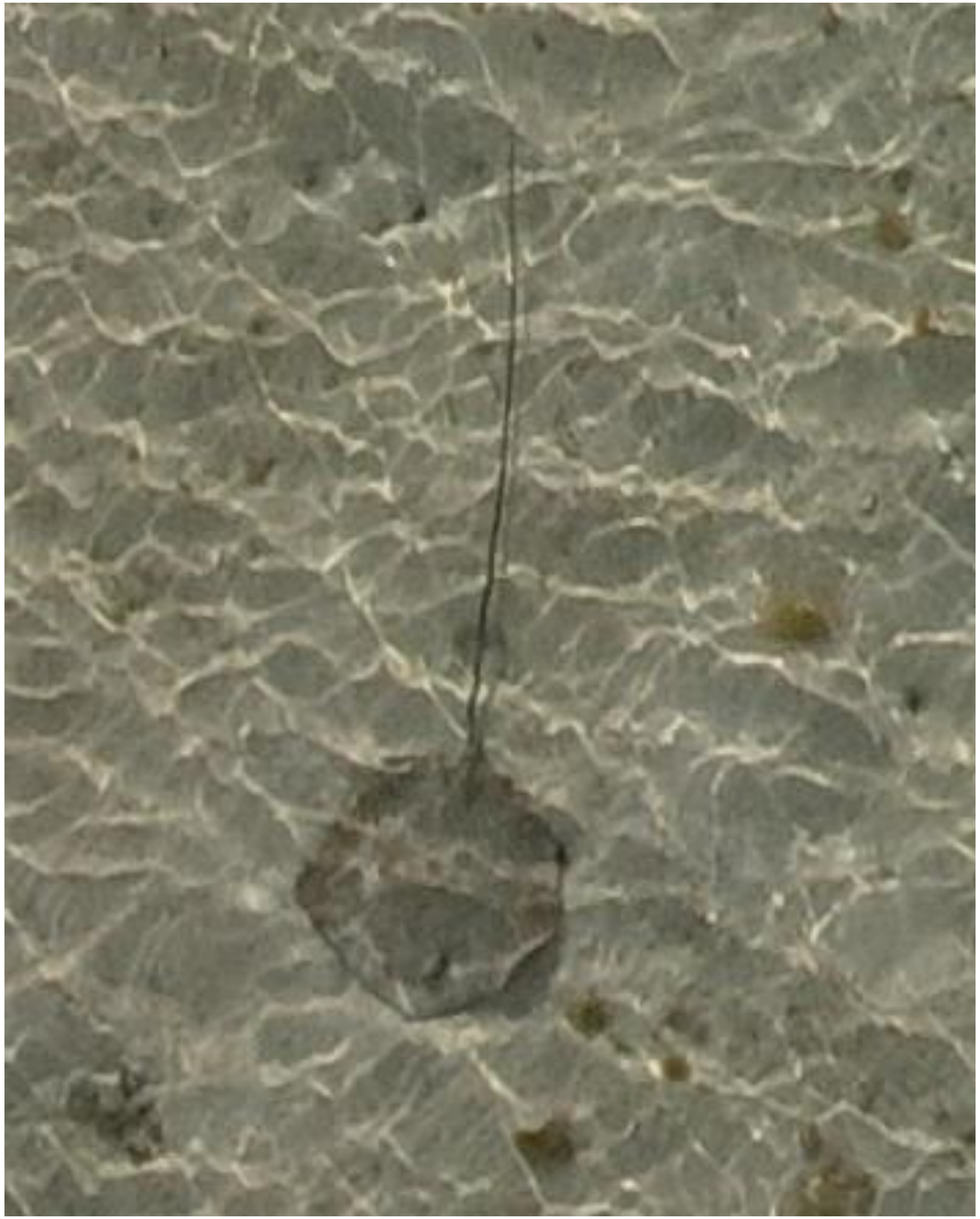

DJI_20240927075653_0409

- Whipray

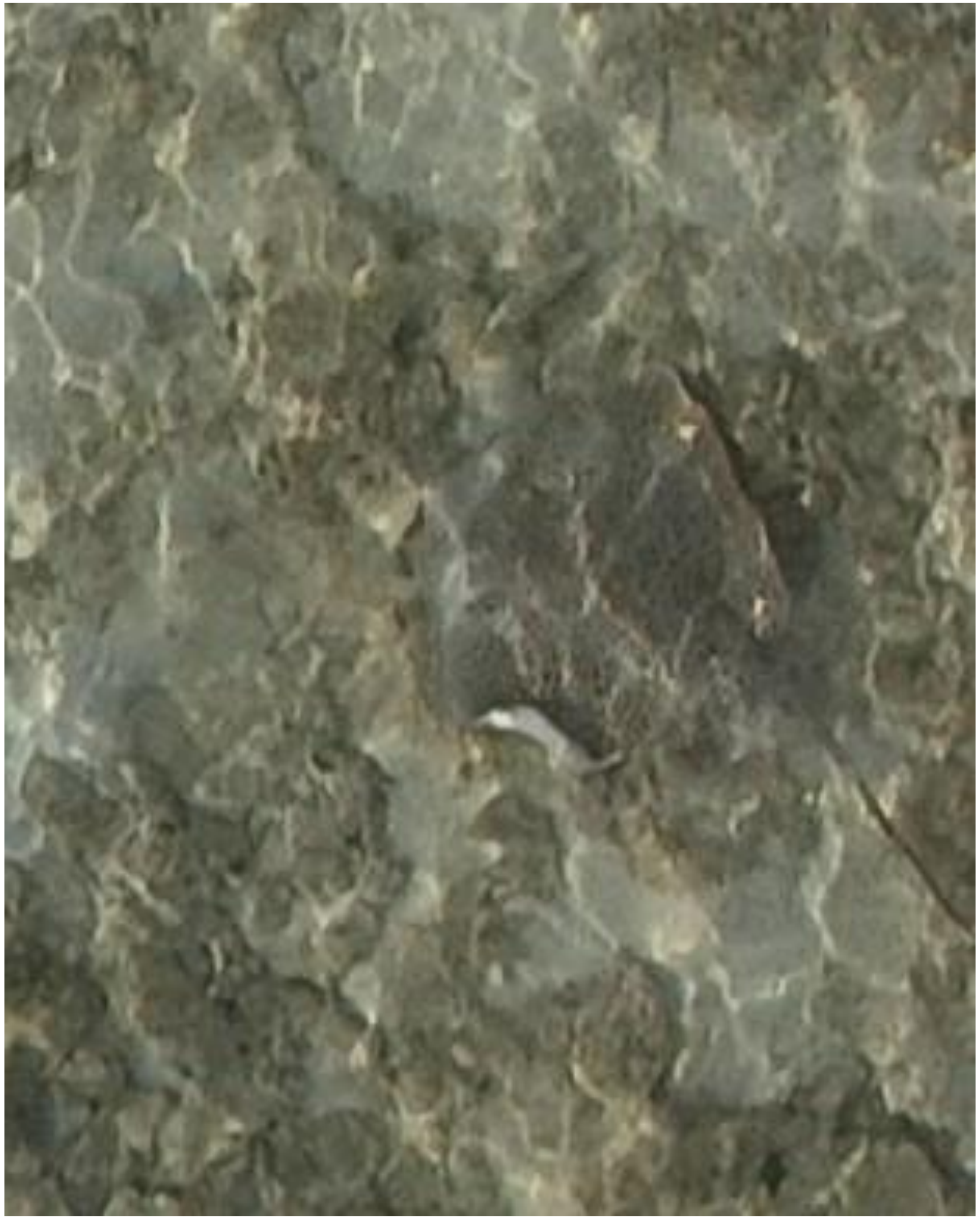

DJI_20240927075930_0536

- *Himantura uarnak*

2024-10-09

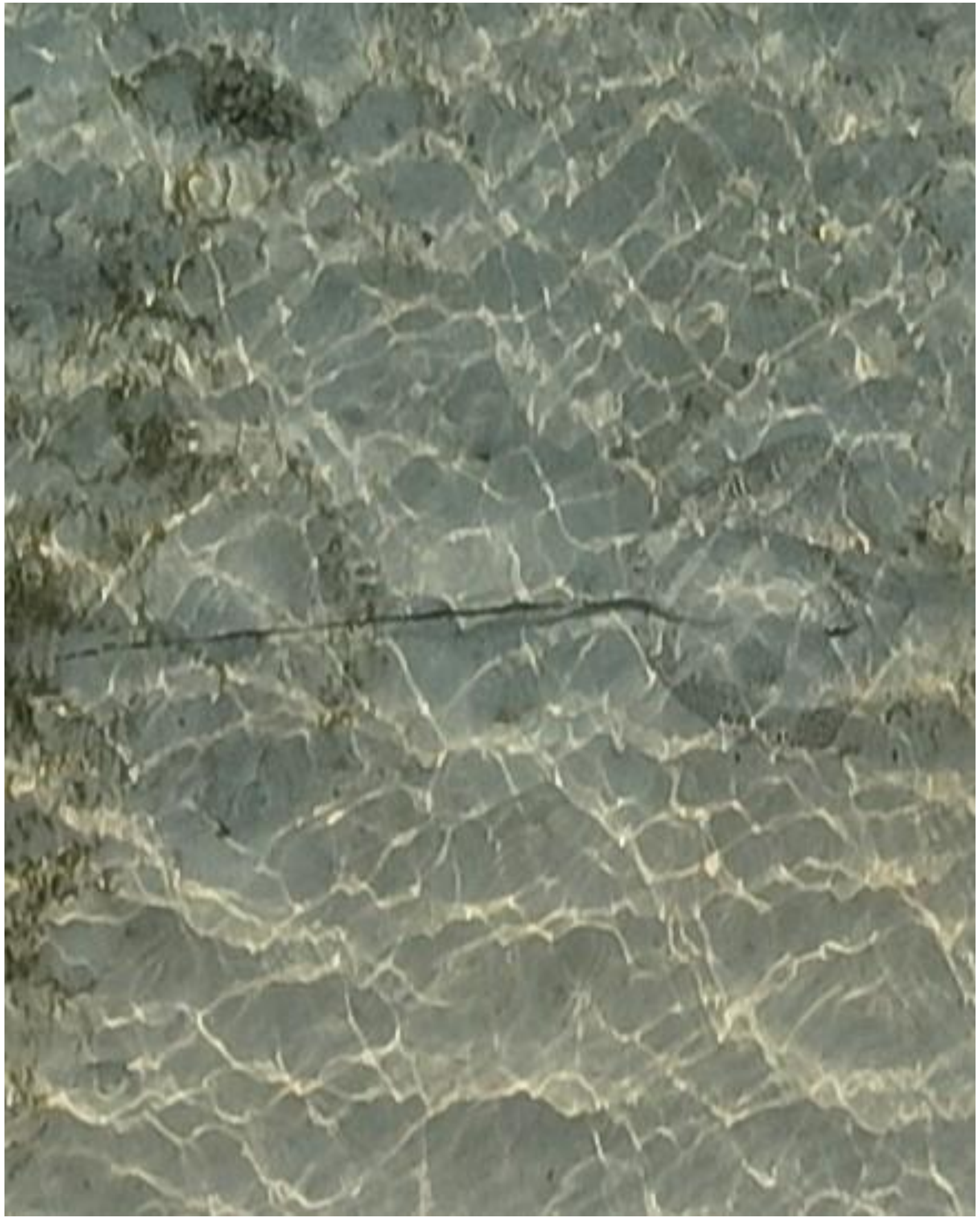

- *Himantura uarnak*

DJI_20241009080206_0058

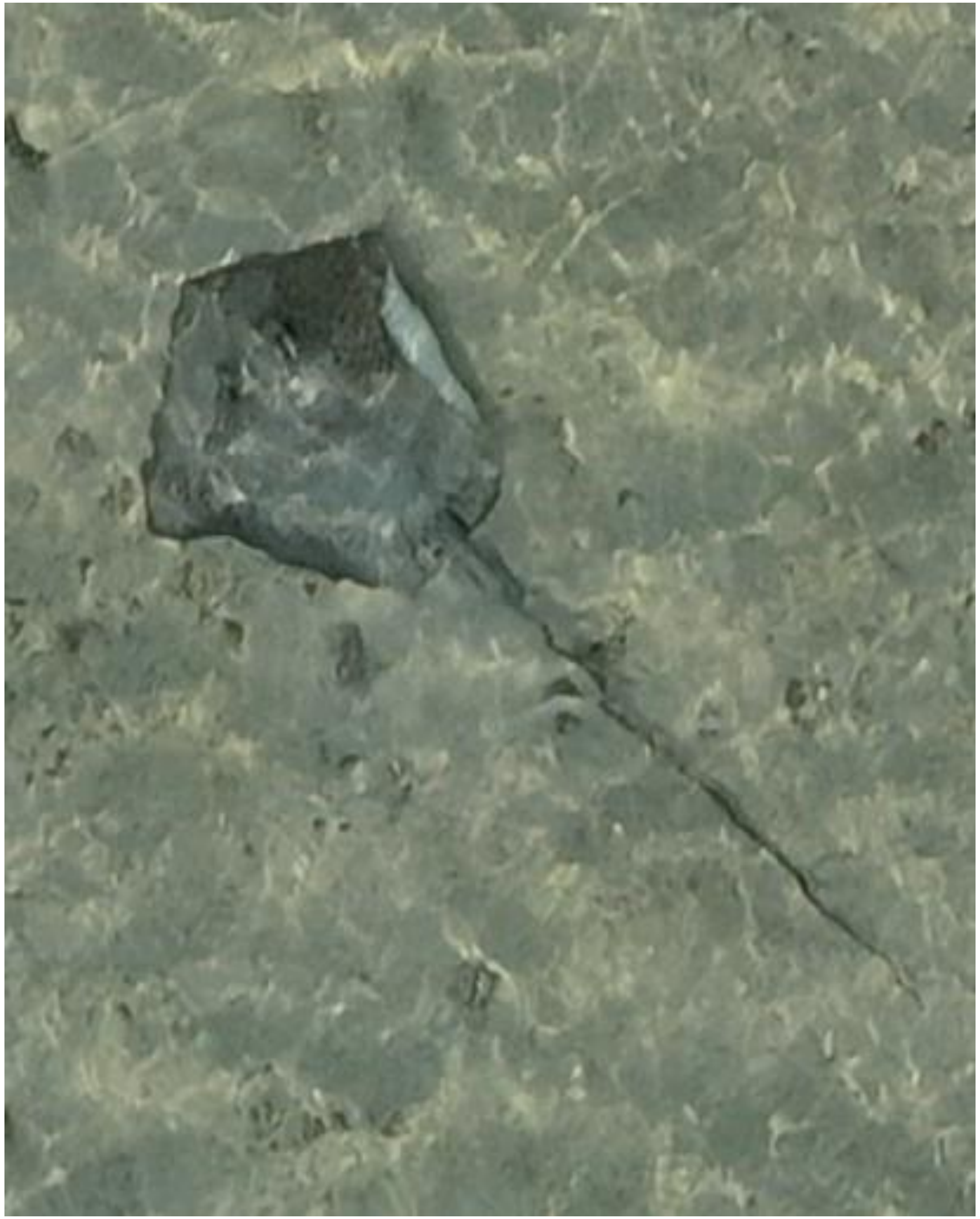

DJI_20241009080305_0106

- *Himantura uarnak*

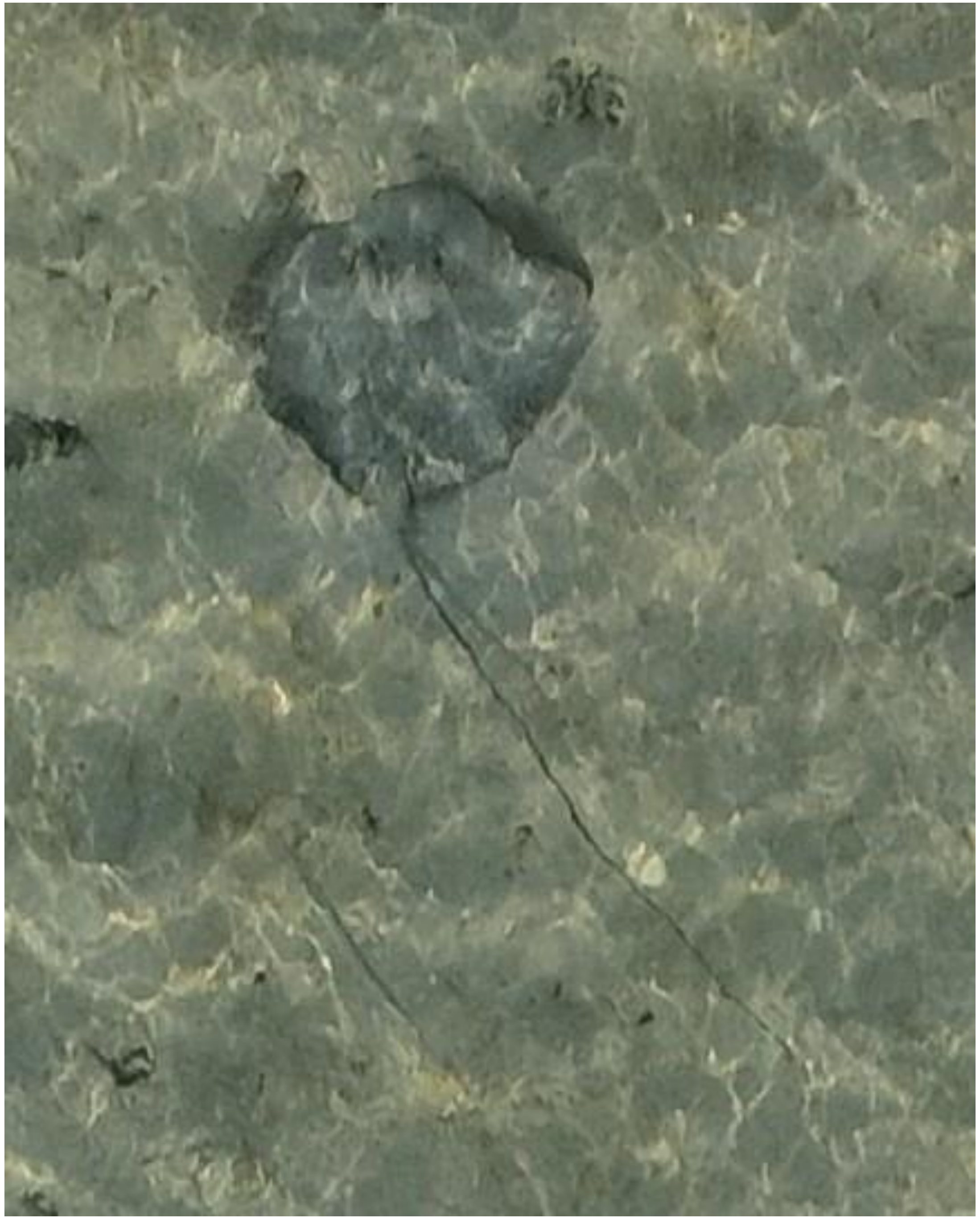

DJI_20241009080307_0108

- Whipray
- Accompanied by needlefish

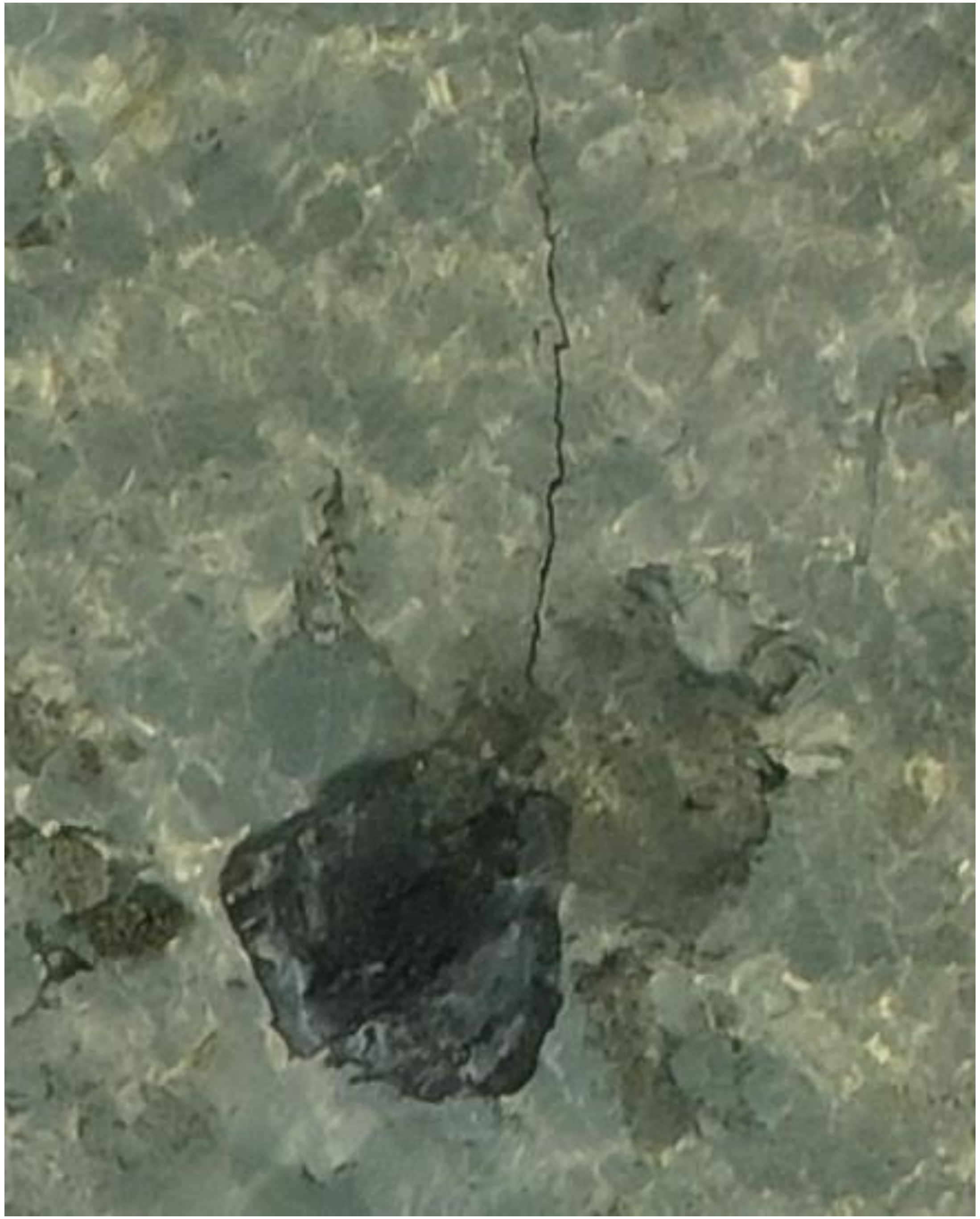
DJI_20241009080307_0108

- Whipray
- Accompanied by needlefish

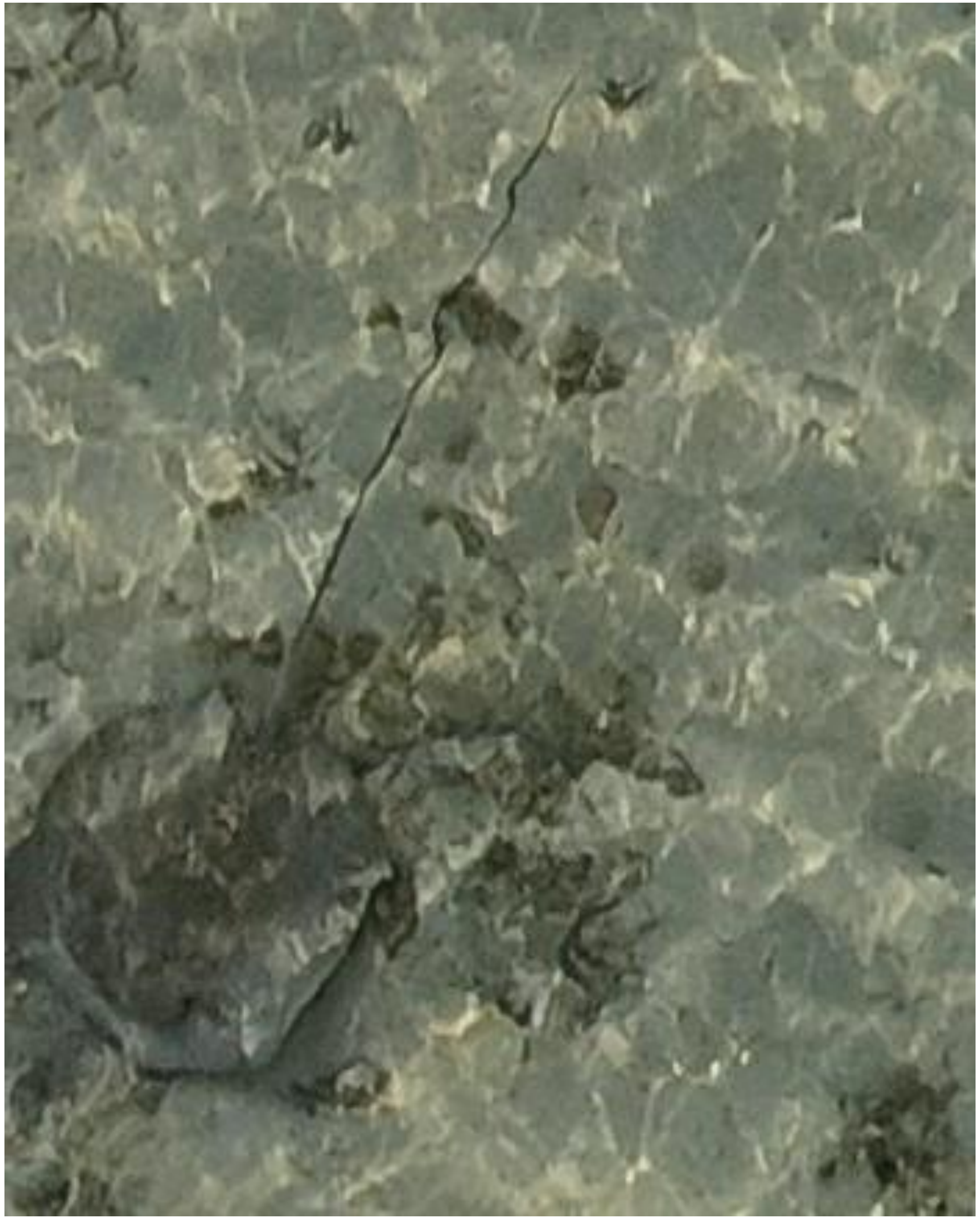

DJI_20241009080504_0213

- Whipray

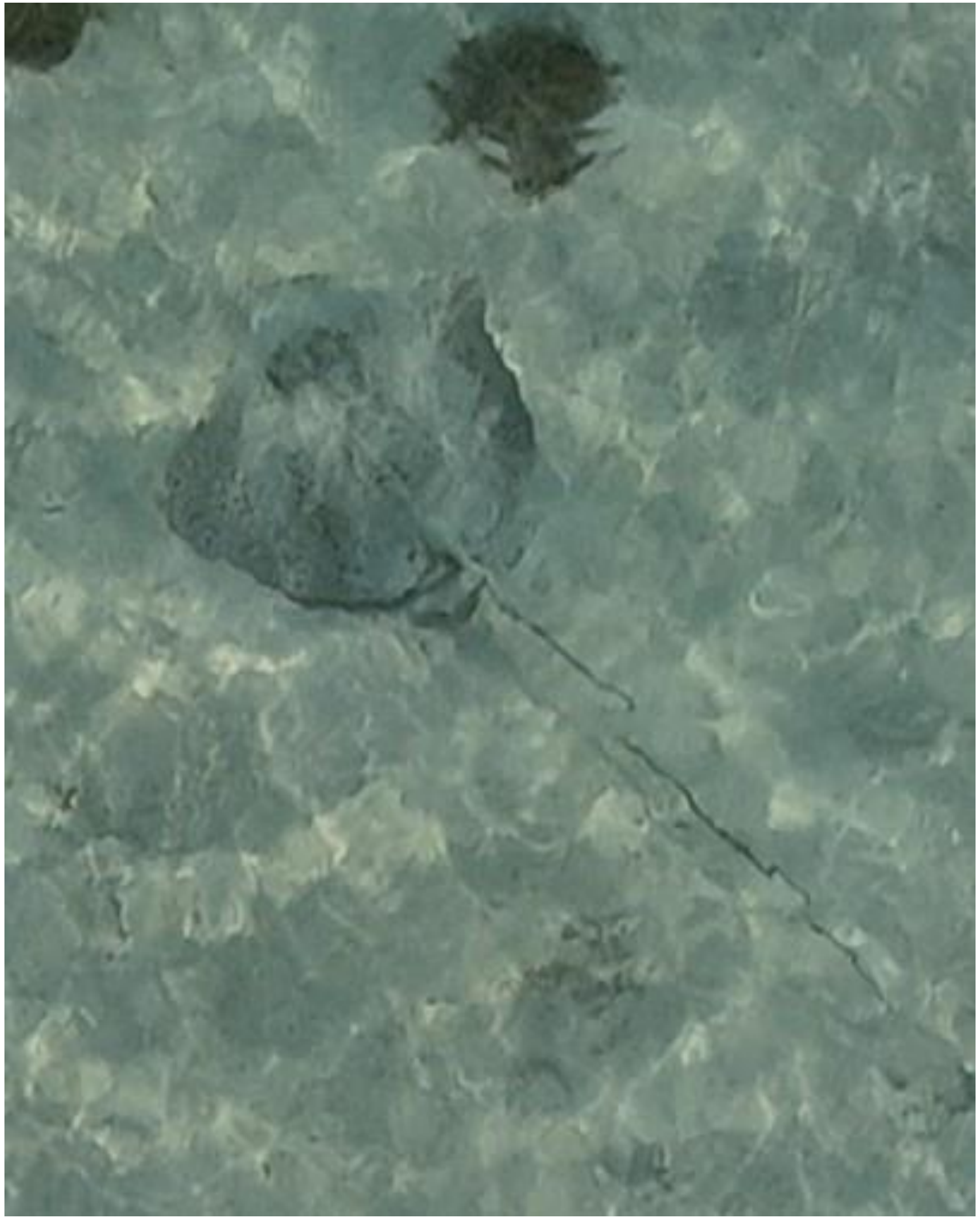
DJI_20241009080519_0228

- *Himantura uarnak*

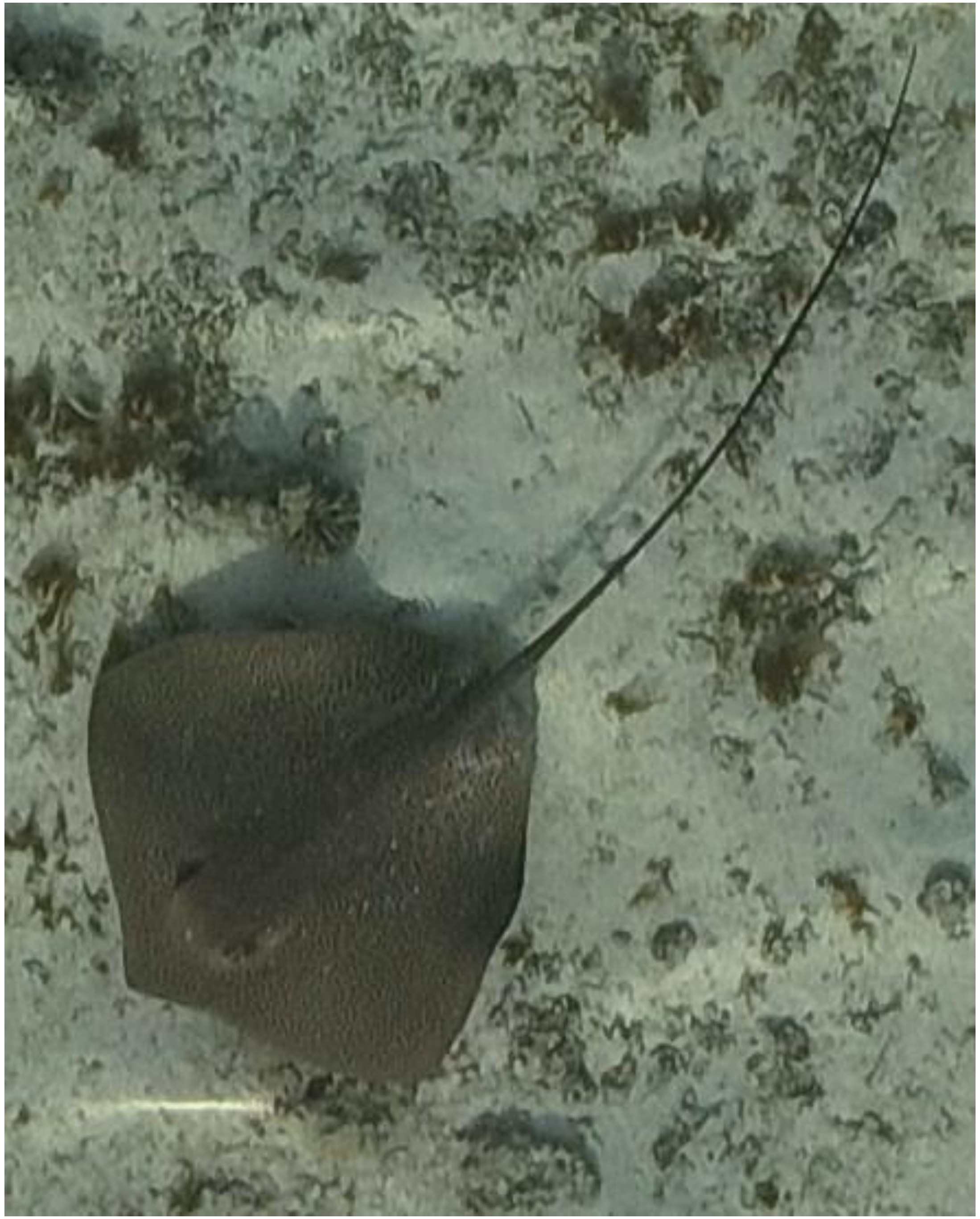

- *Himantura uarnak*

DJI_20241009080718_0335

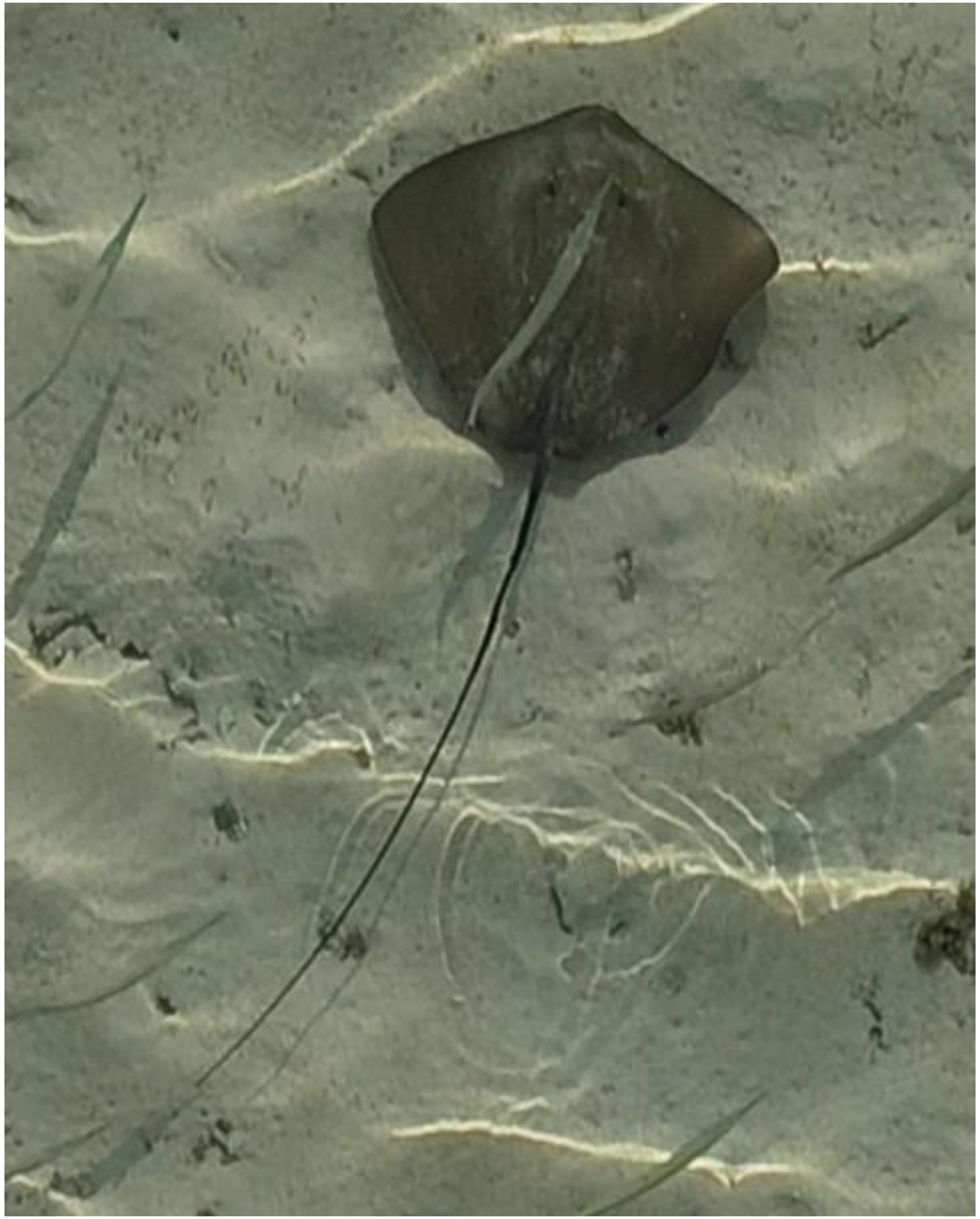

- cf. *Pateobatis fai*
- Accompanied by needlefish

DJI_20241009080827_0392

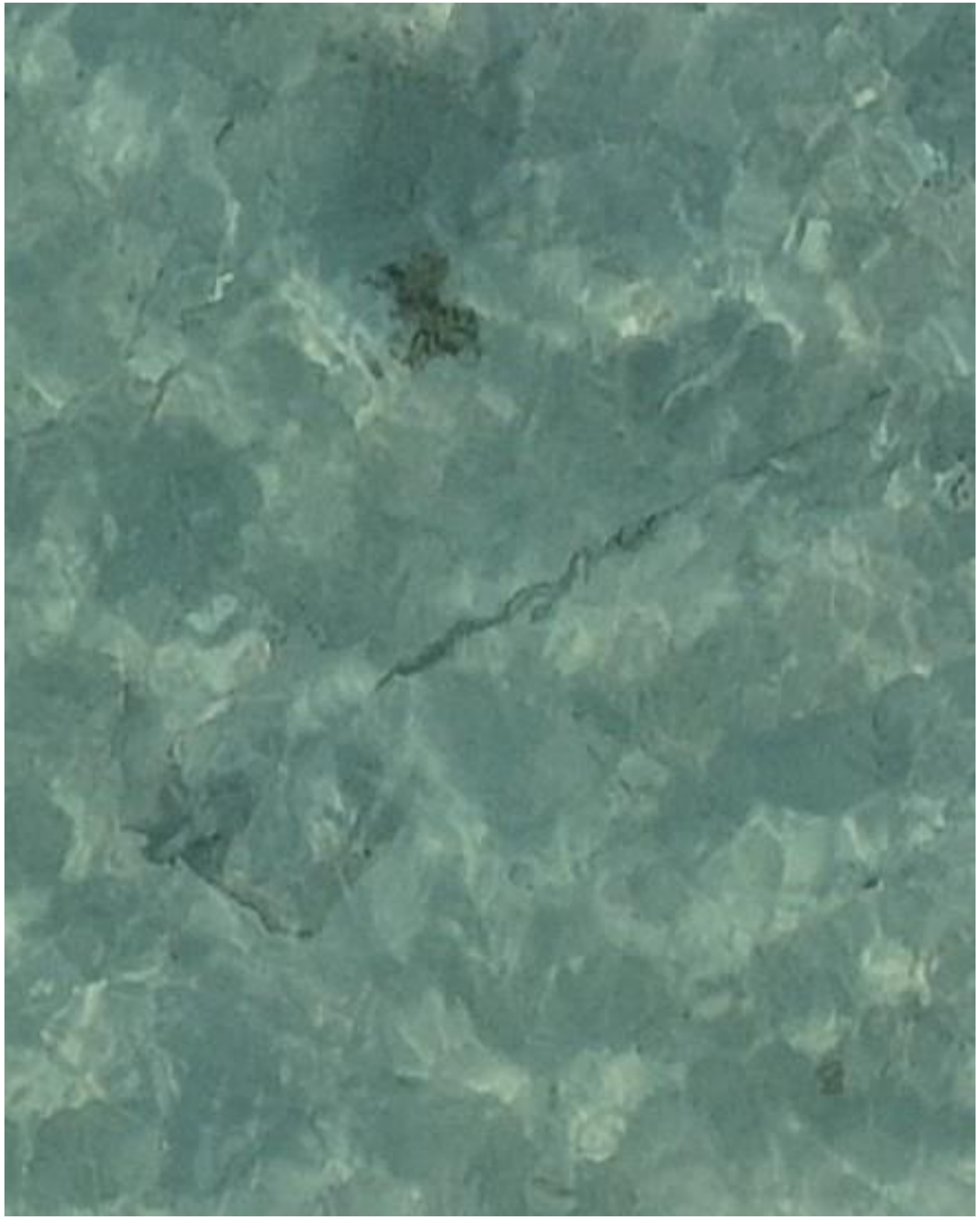

- Whipray

DJI_20241009080926_0451

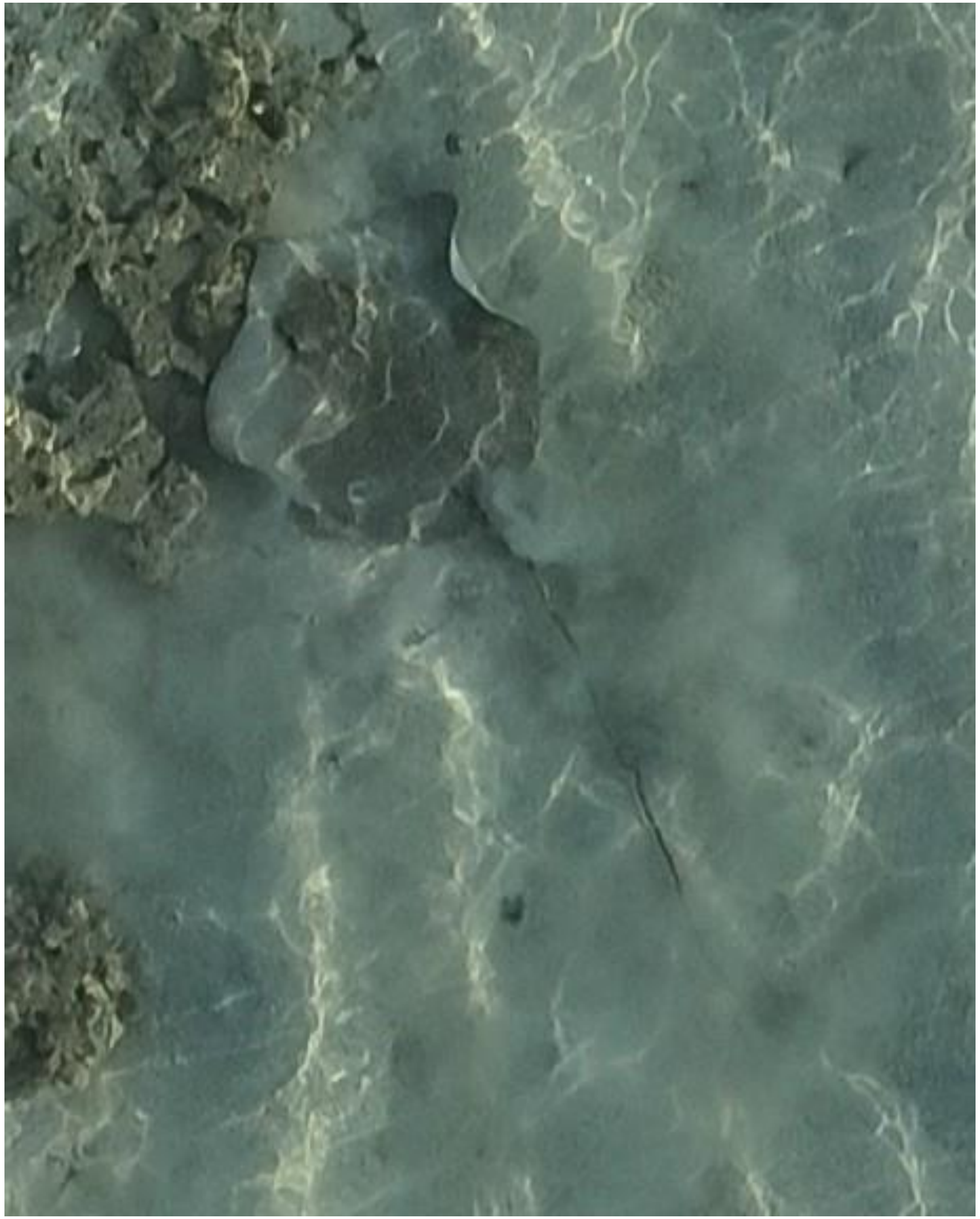

- Whipray

DJI_20241009081352_0670

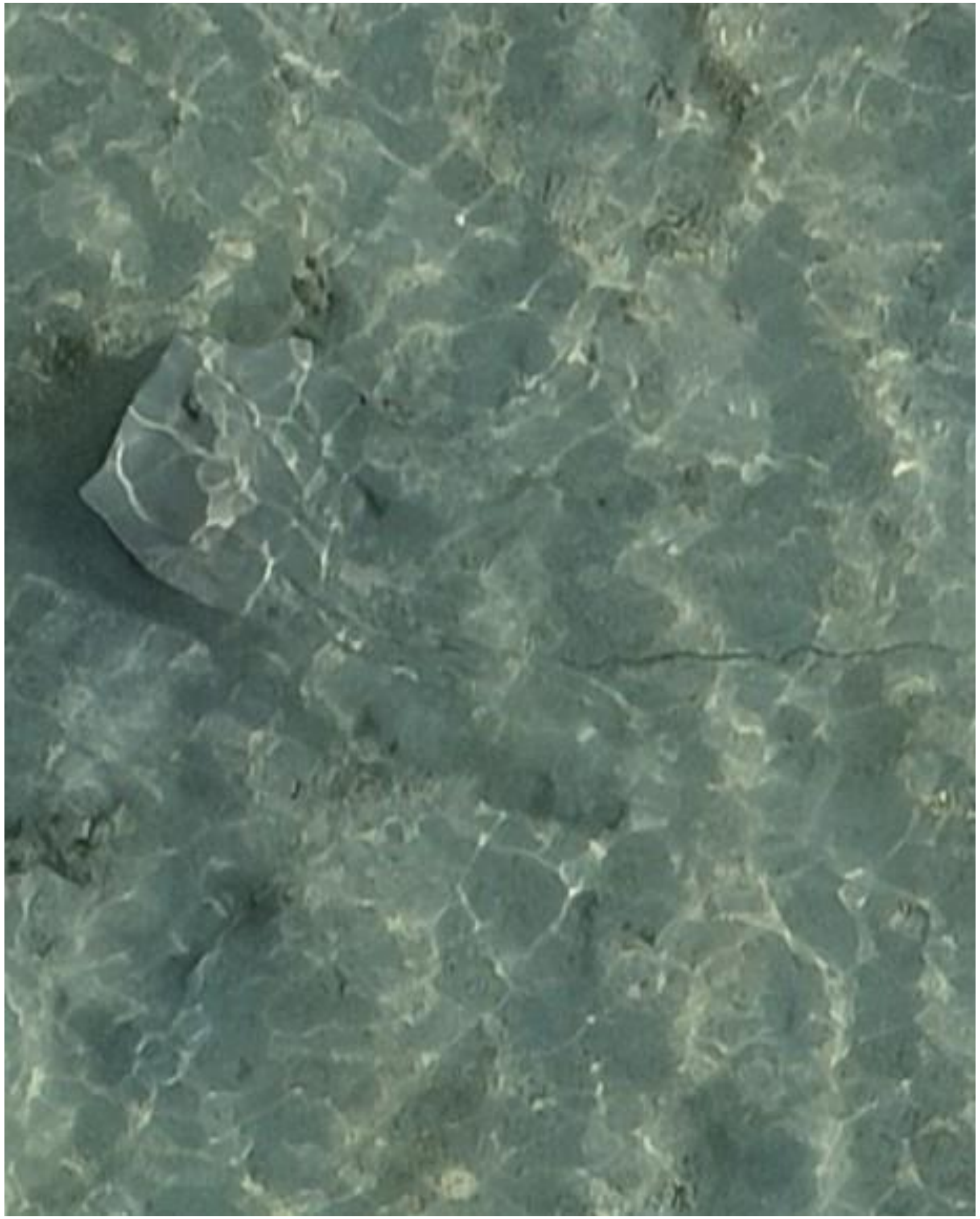

- Whipray

DJI_20241009081403_0681

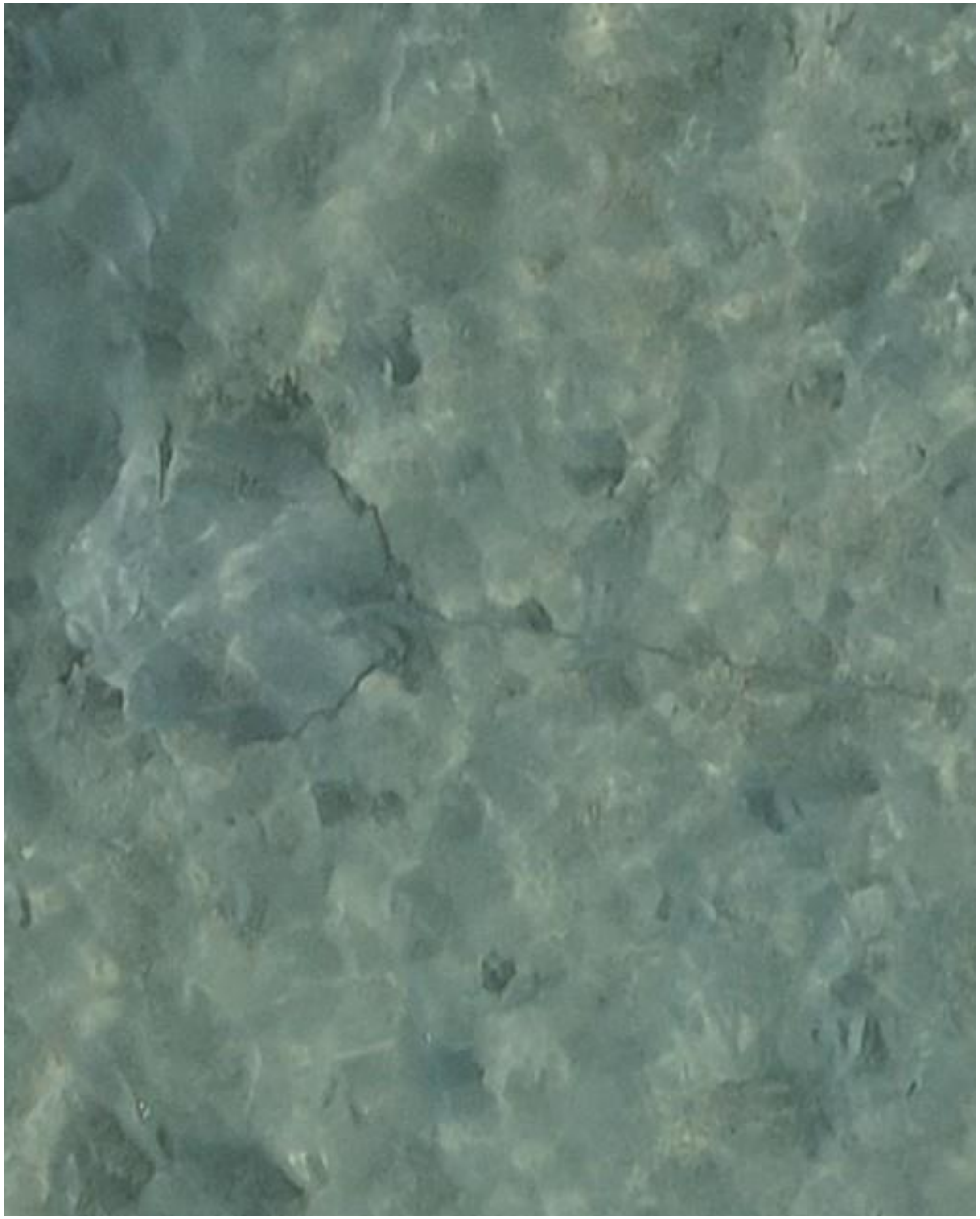

- Whipray

DJI_20241009081953_0994

2024-10-26

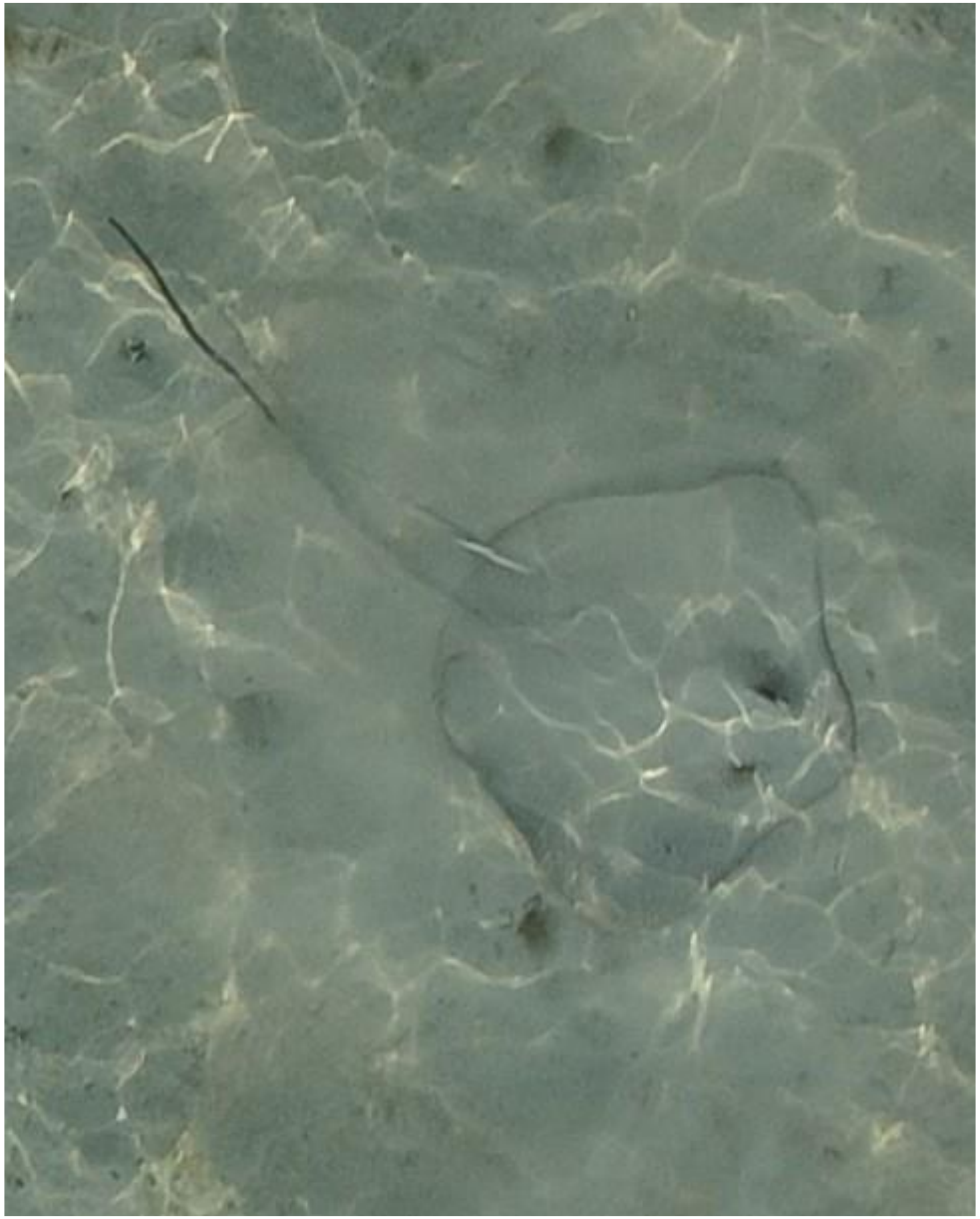

- Whipray
- Accompanied by needlefish

DJI_20241026080428_0096

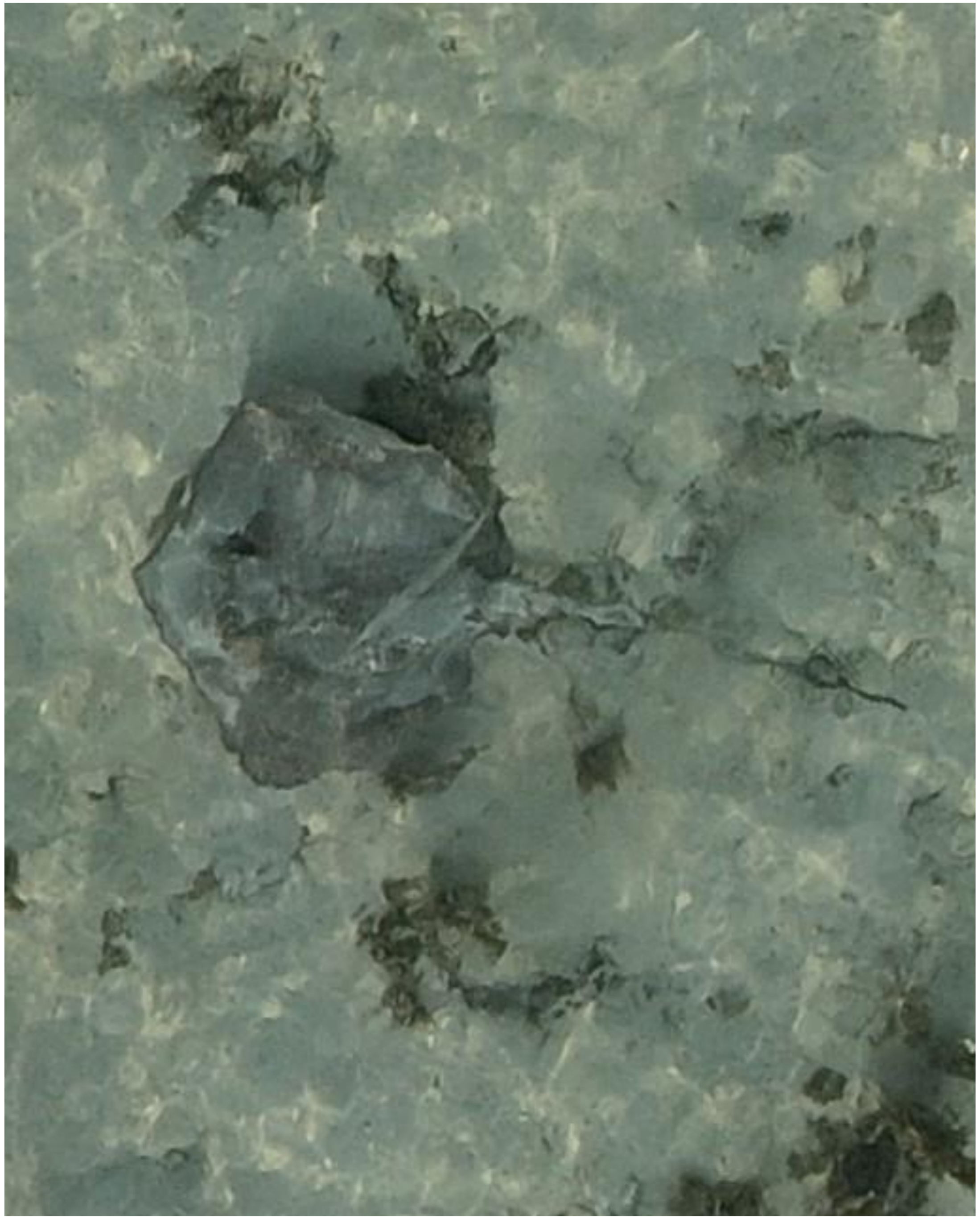

DJI_20241026080439_0107

- Whipray
- Accompanied by needlefish

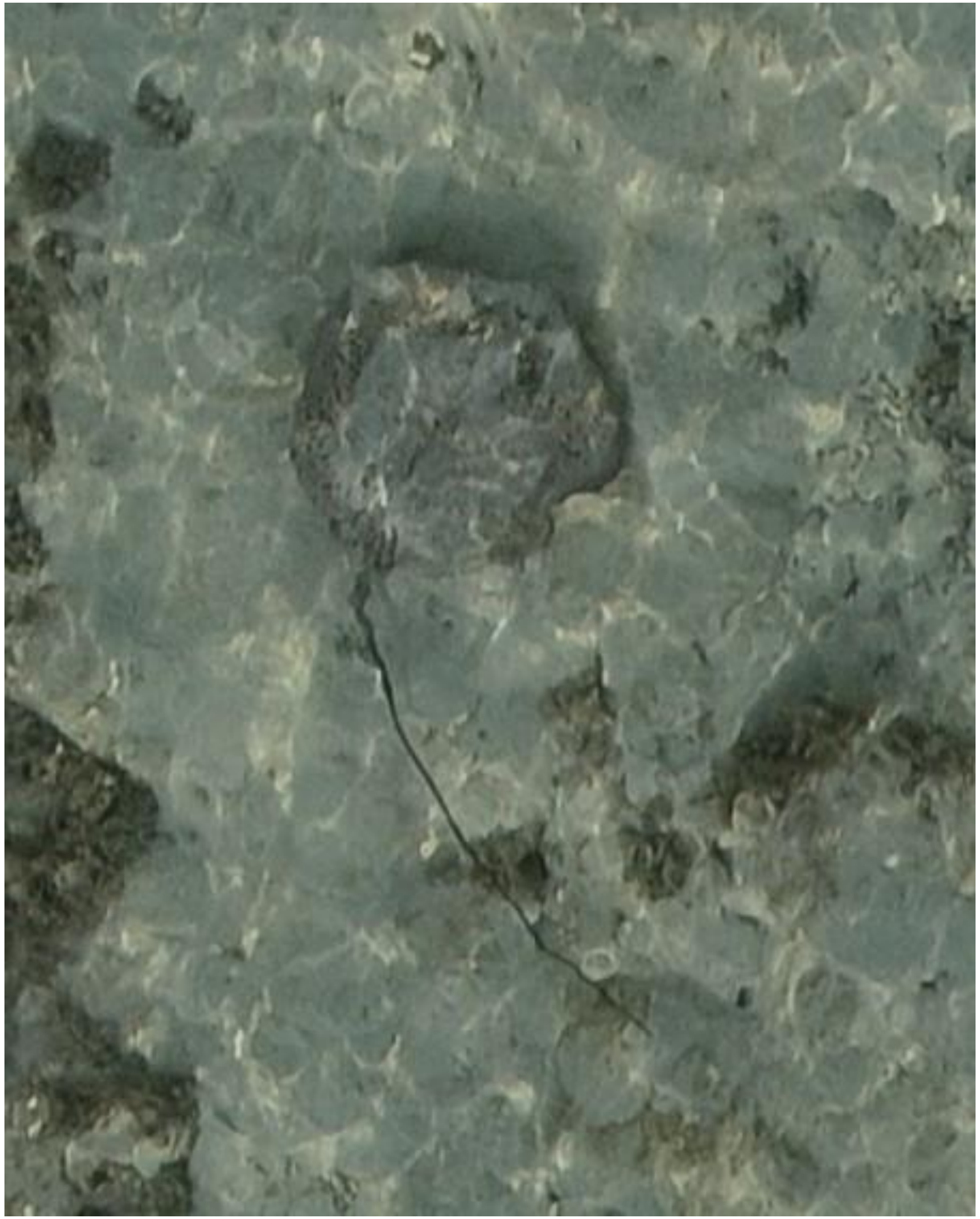

- *Himantura uarnak*

DJI_20241026080810_0294

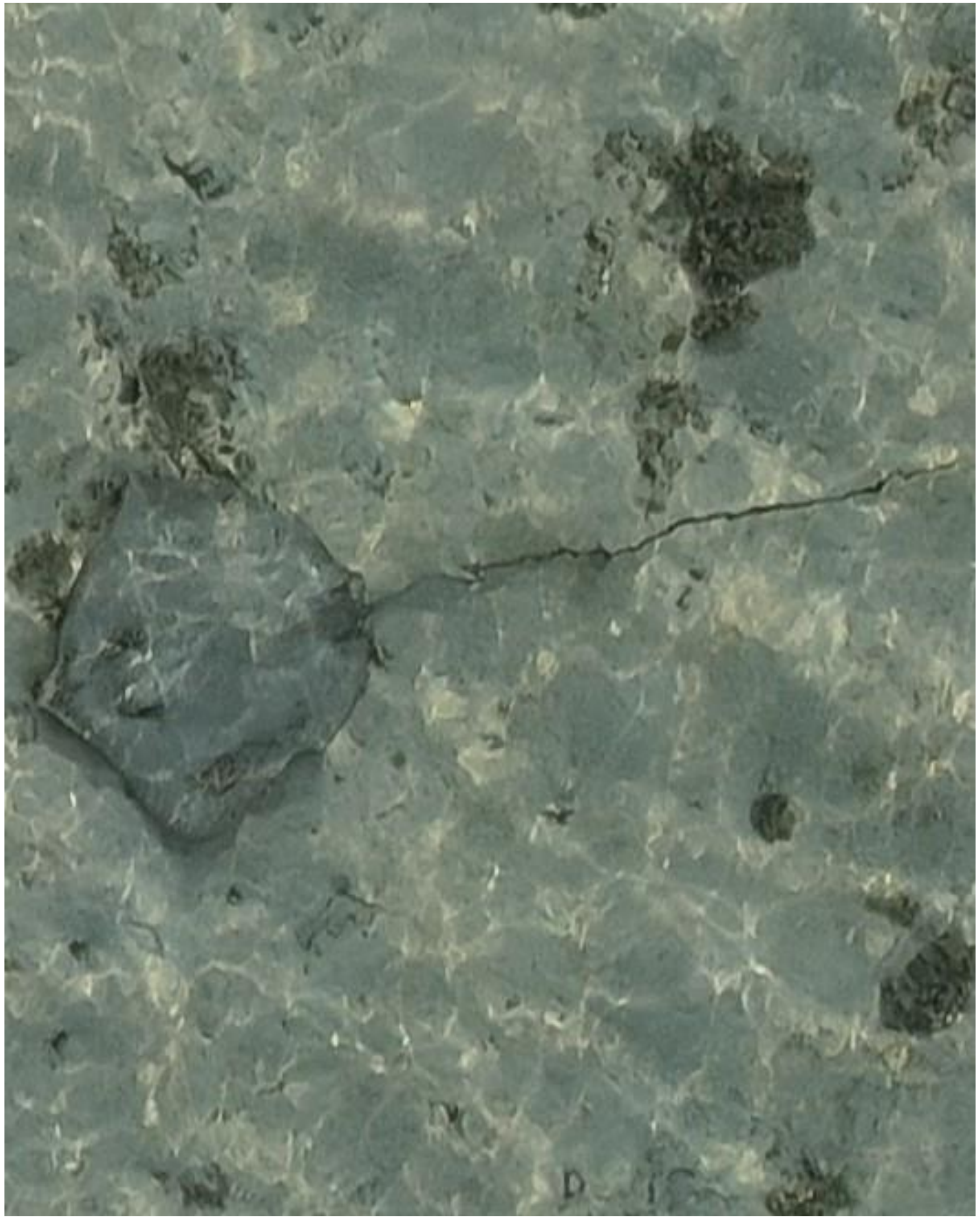

DJI_20241026081004_0395

- Whipray
- Best guess: *H. uarnak*

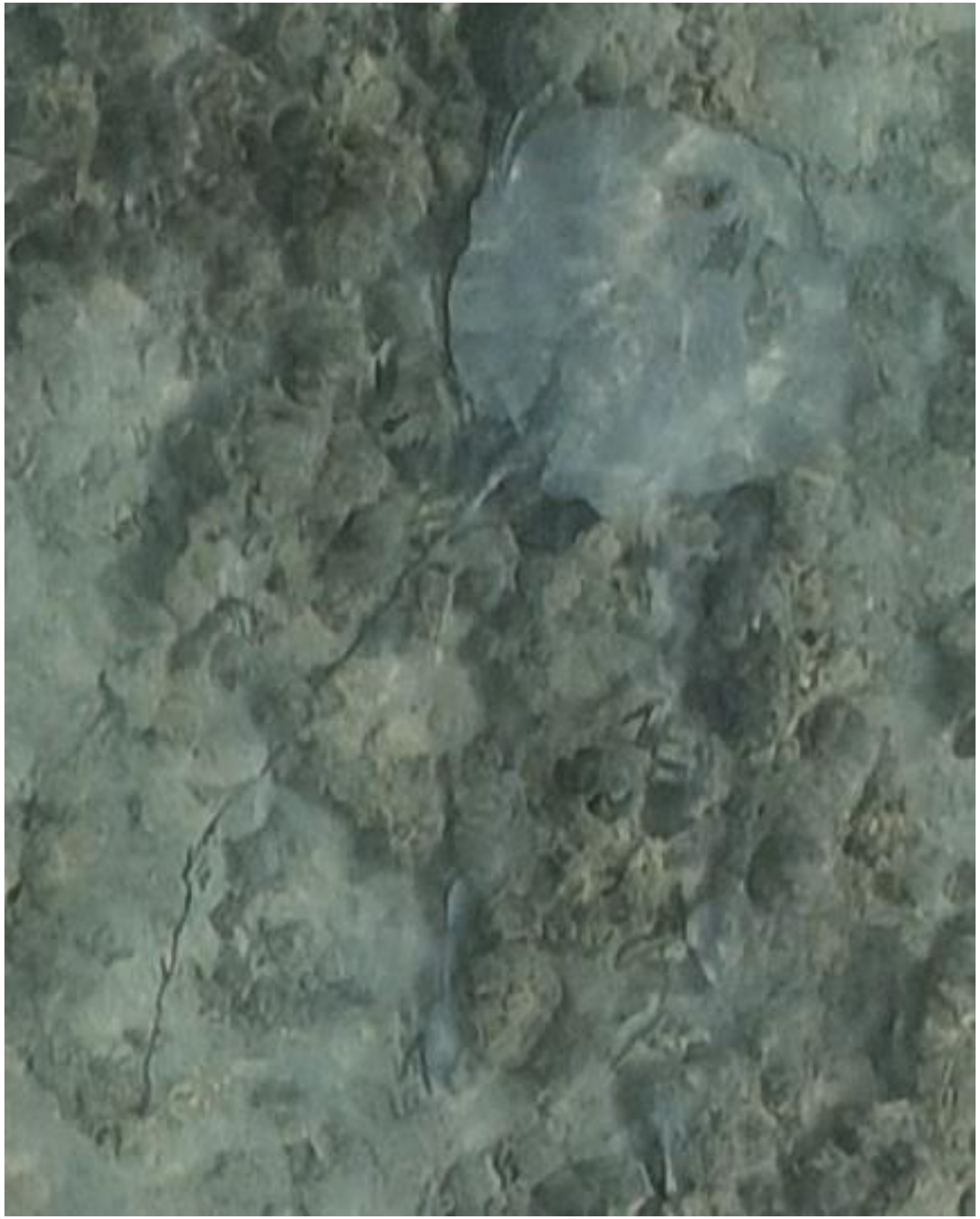

DJI_20241026081449_0635

- Whipray

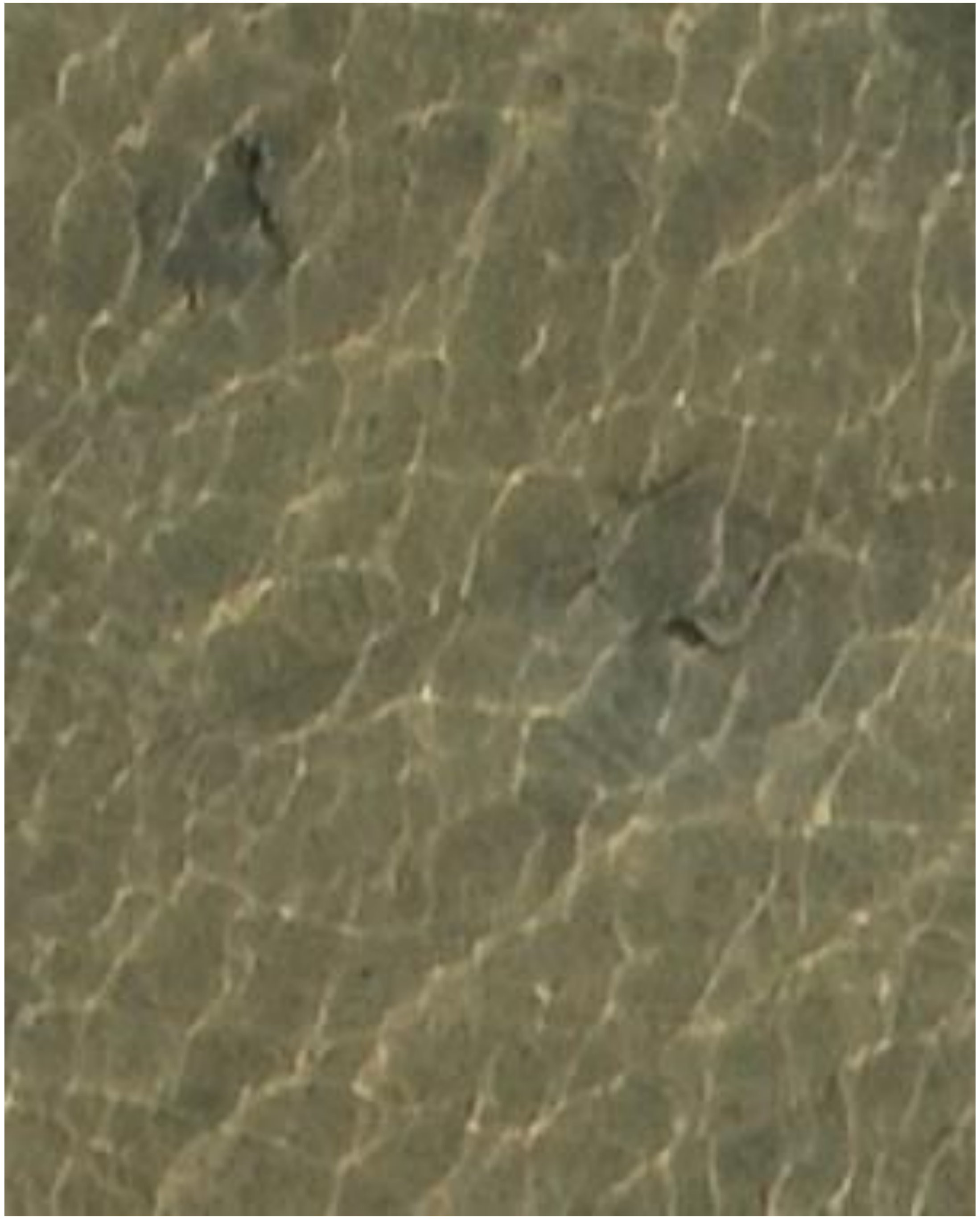

DJI_20241026081610_0716

- Whipray
- Very small, 24 cm disc width

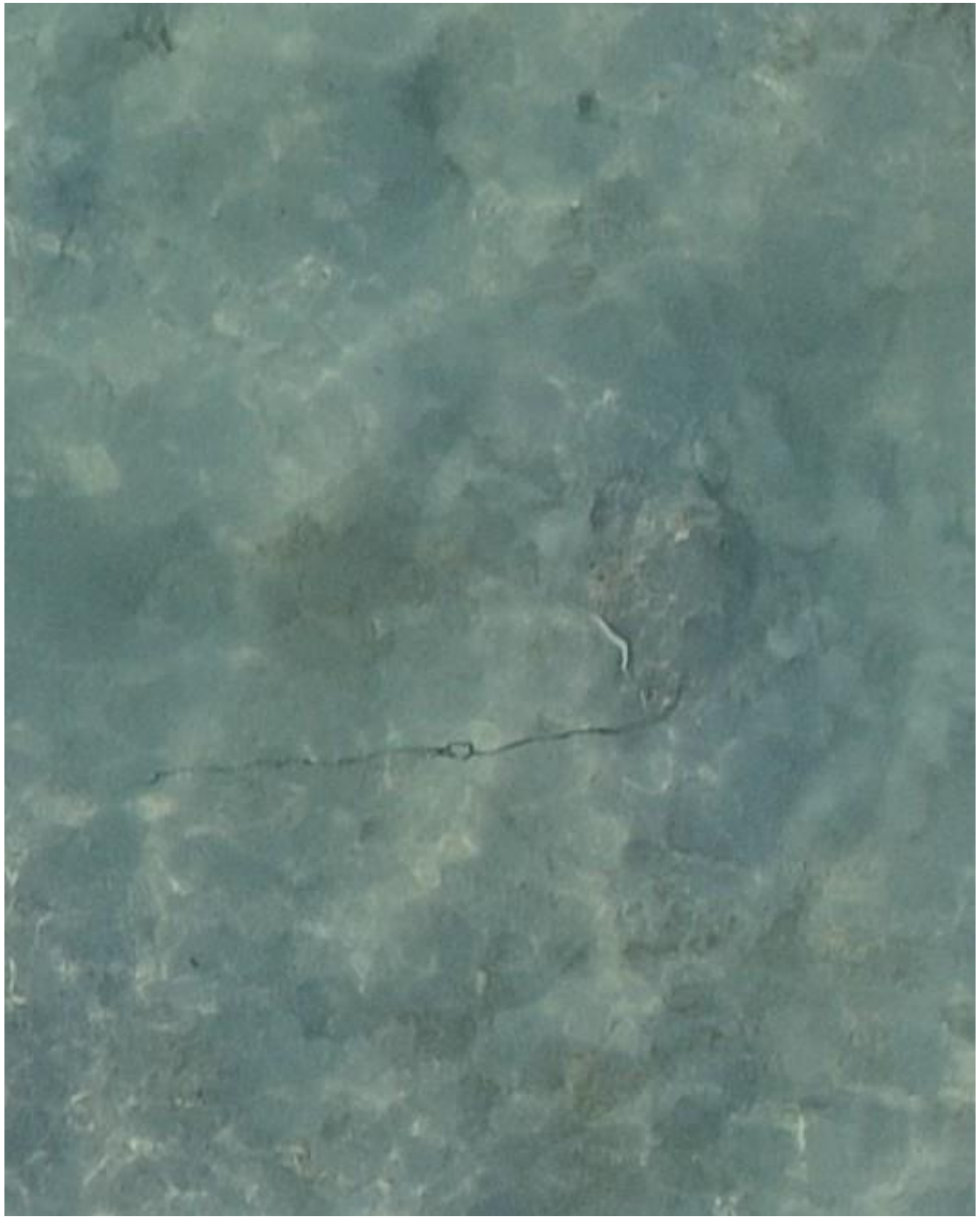

- Whipray
- Best guess: *H. uarnak*

DJI_20241026081730_0783

2024-11-18

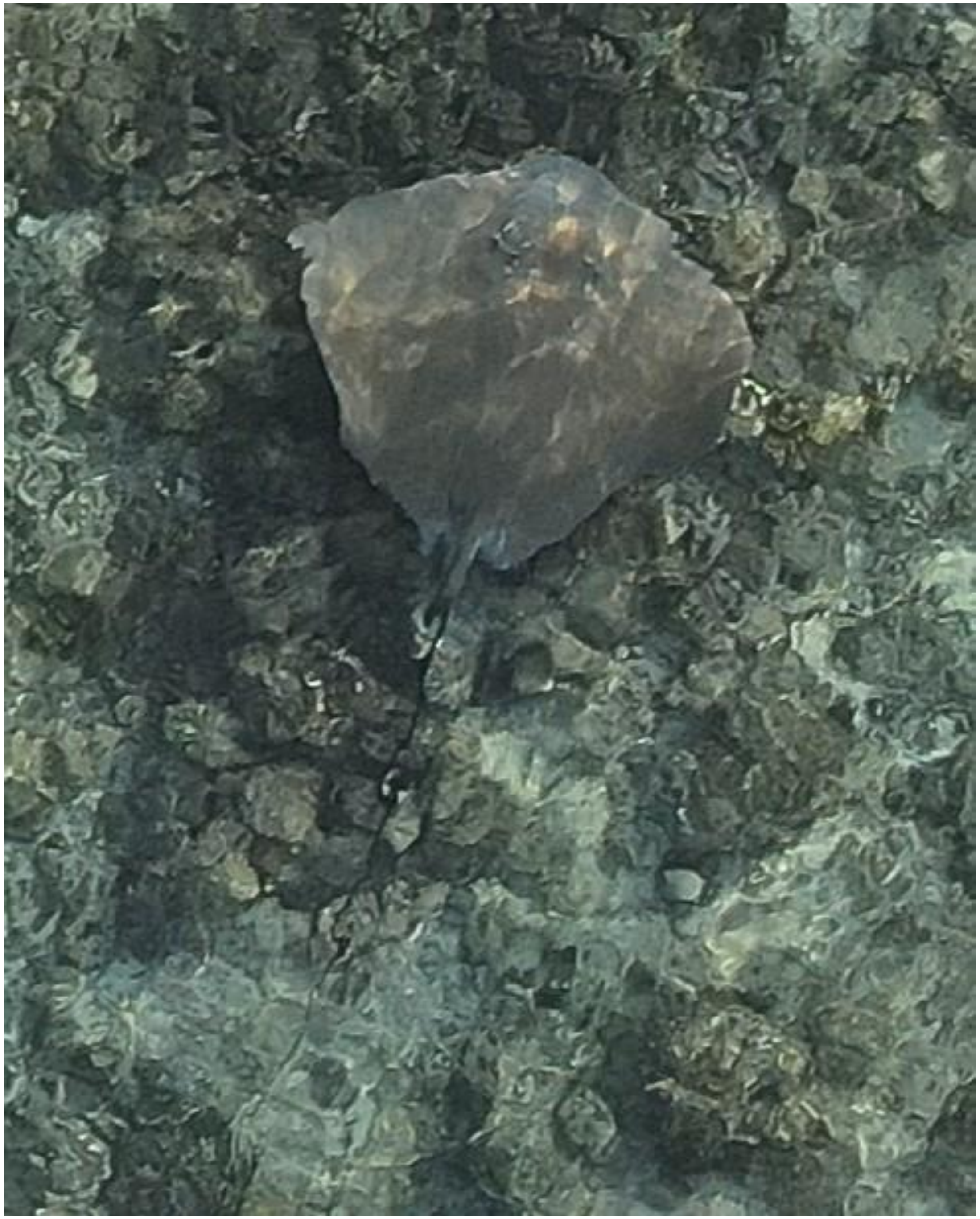

- cf. *Pateobatis fai*

DJI_20241118082014_0001

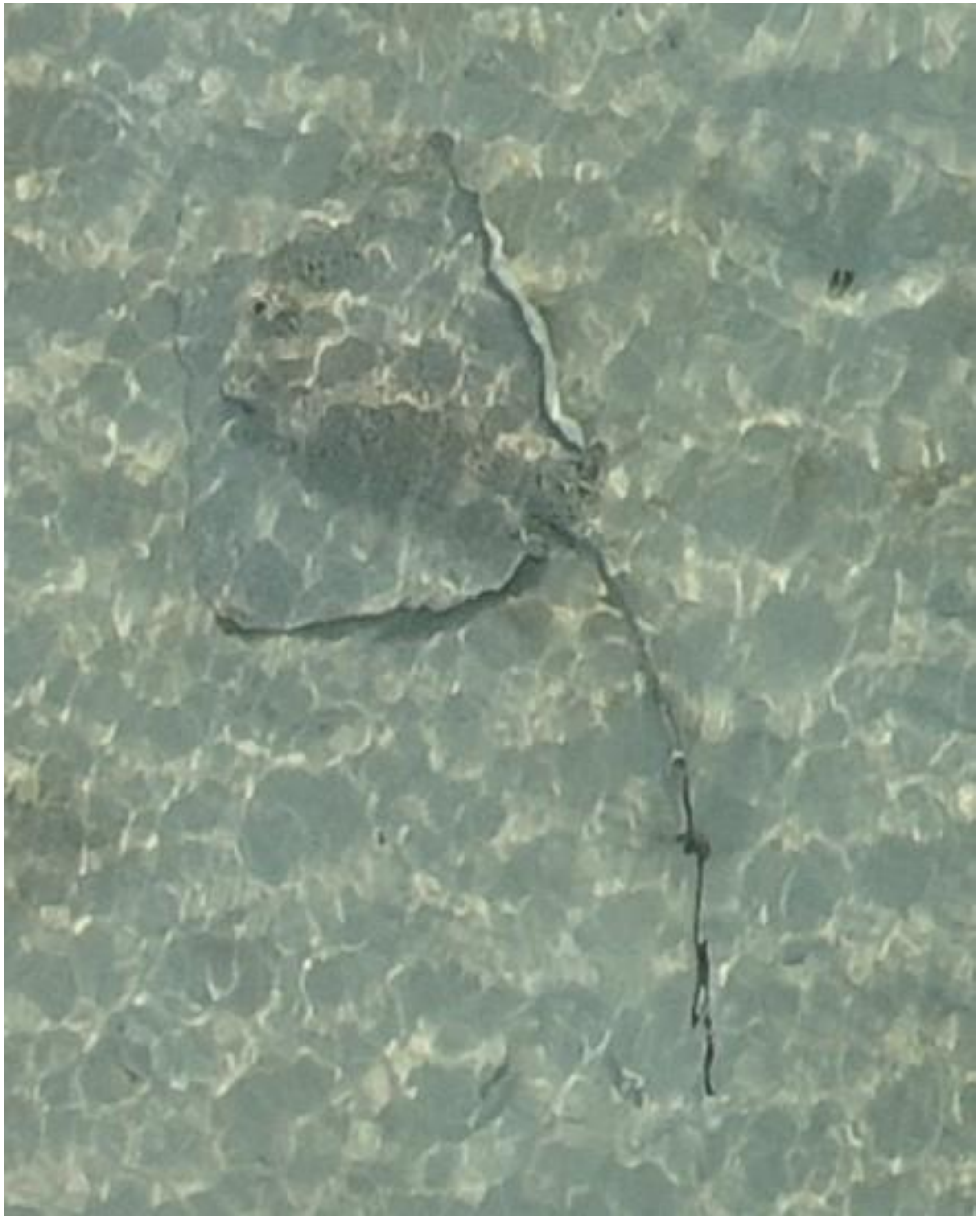

DJI_20241118082111_0058

- *Himantura uarnak*

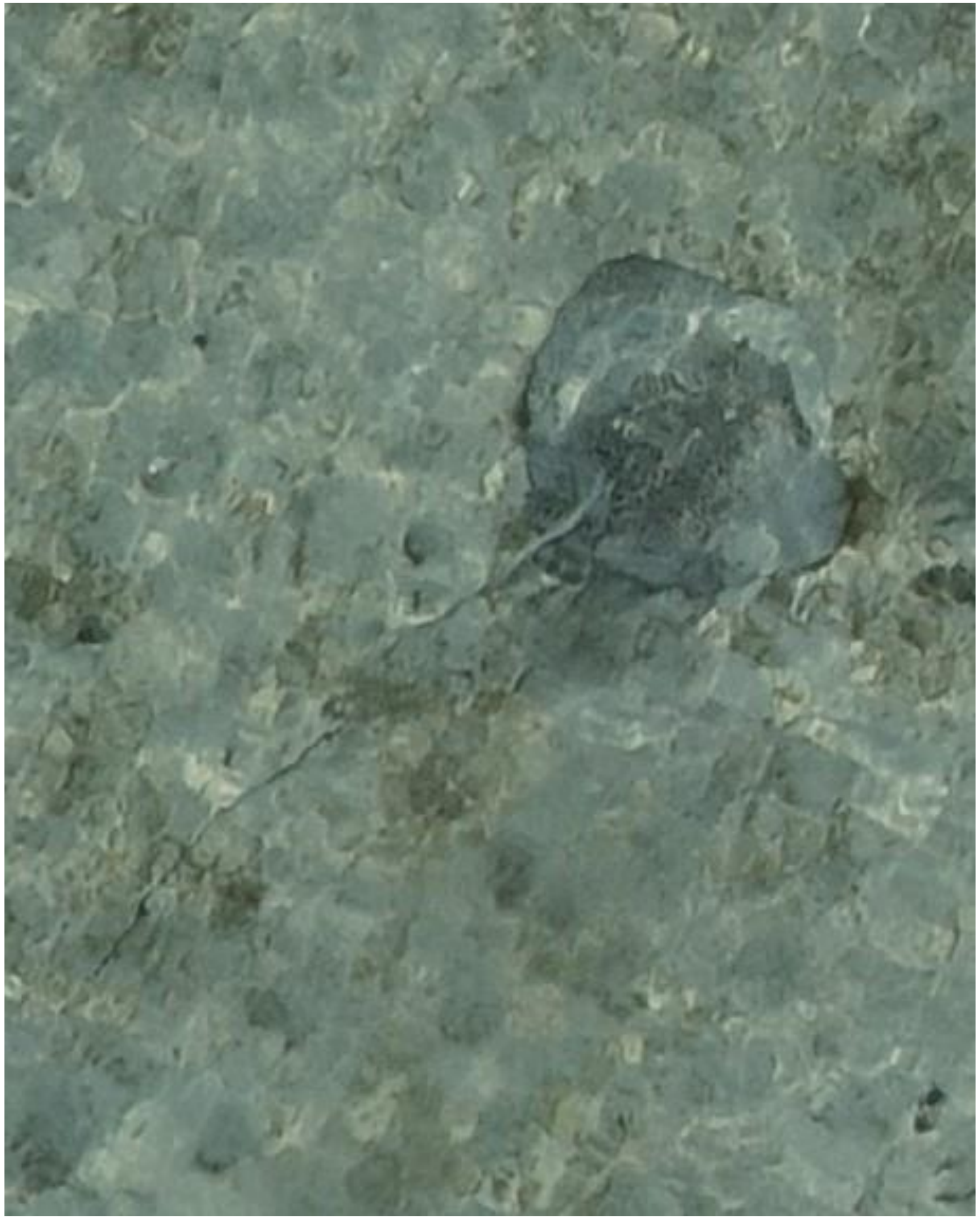

DJI_20241118082110_0057

- *Himantura uarnak*

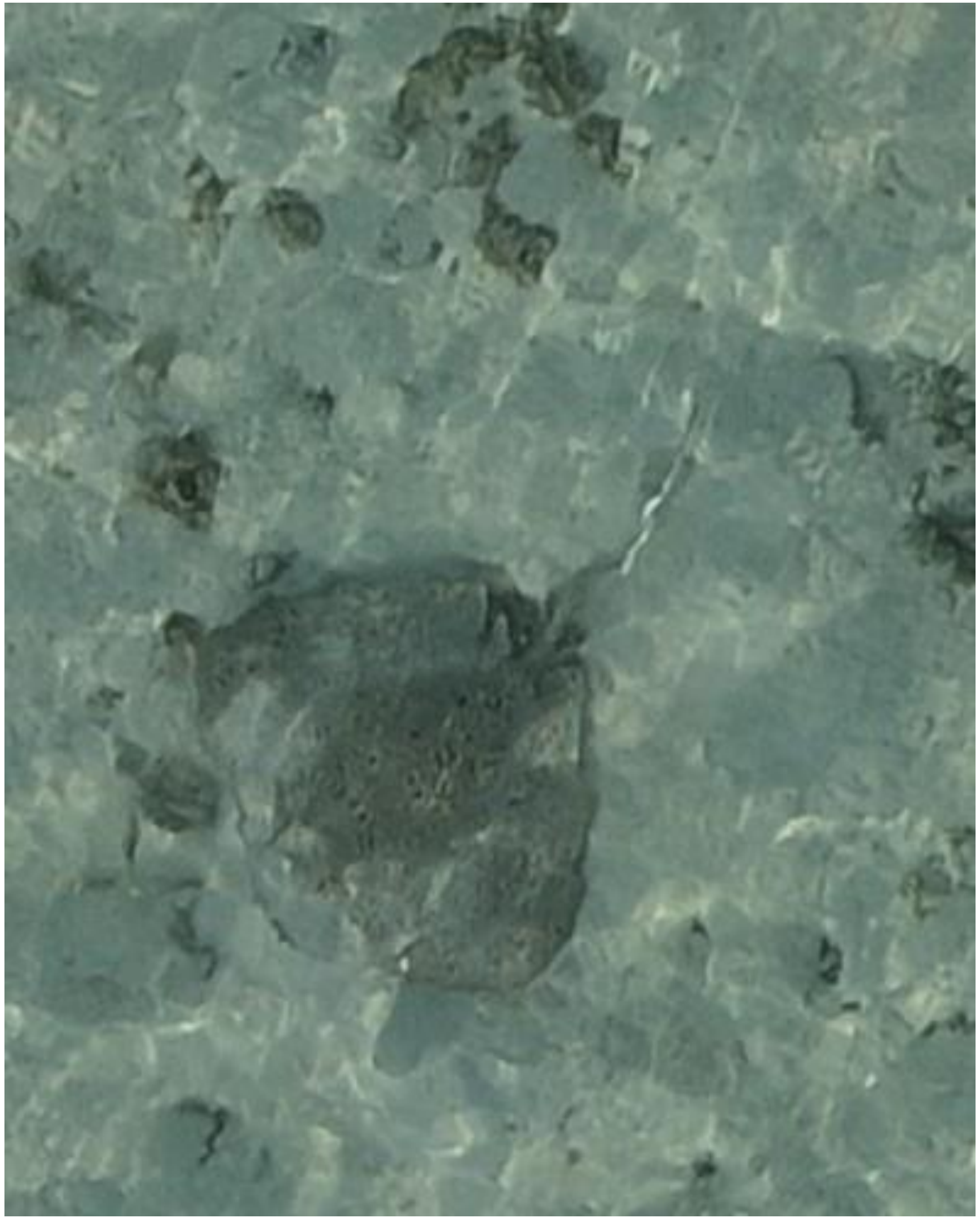

DJI_20241118082235_0130

- *Himantura uarnak*

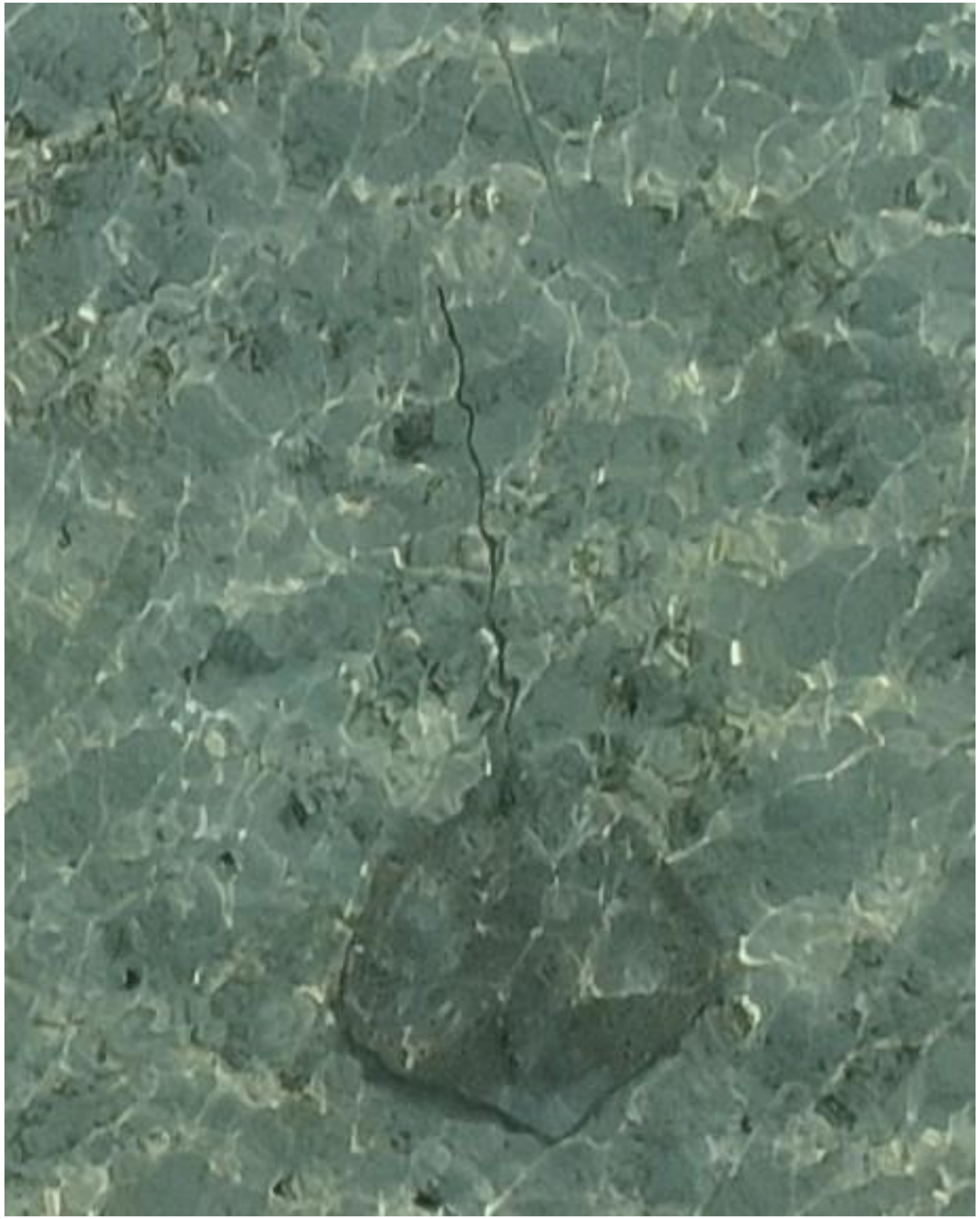

DJI_20241118082748_0406

- *Himantura uarnak*
- Accompanied by needlefish

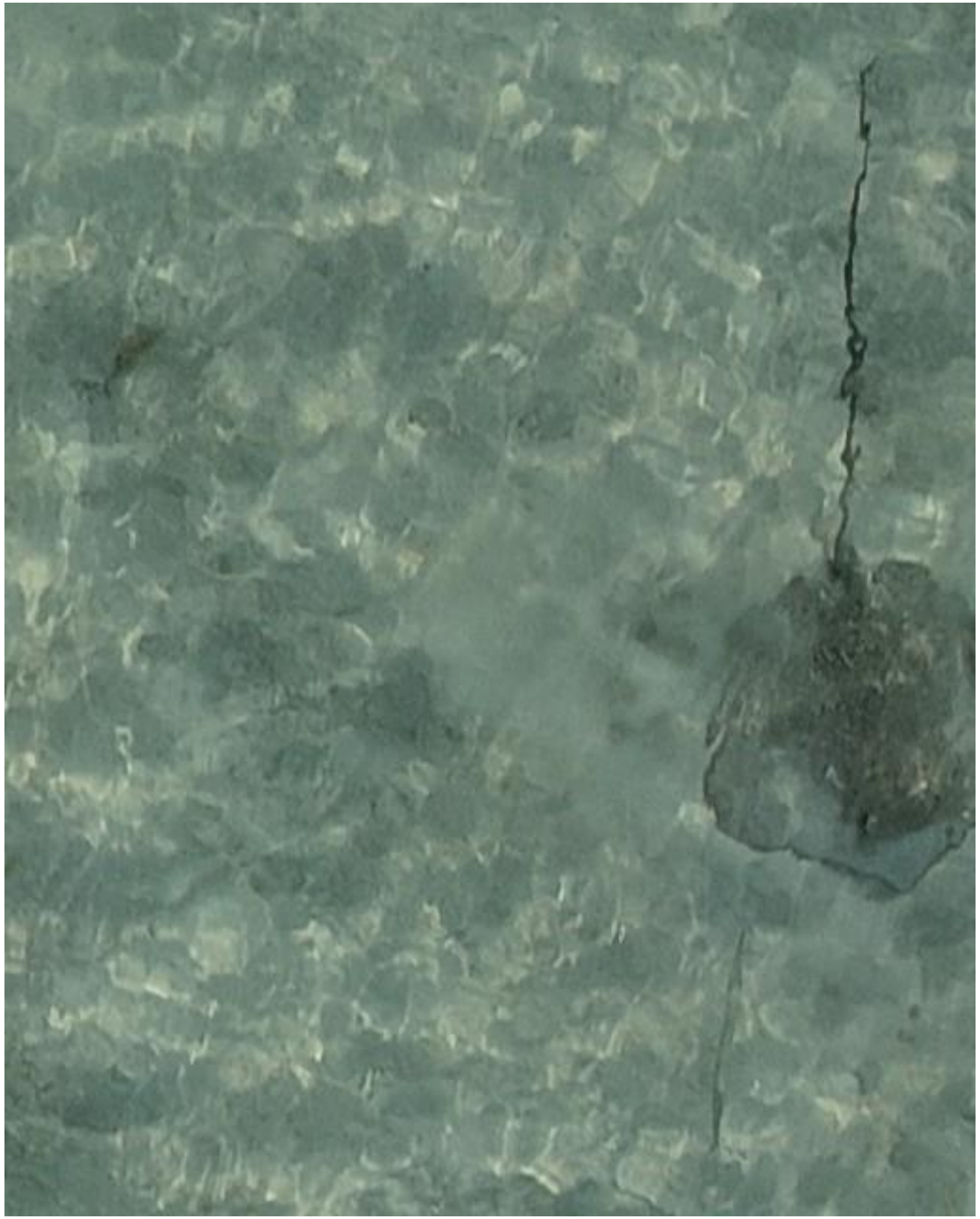

DJI_20241118082810_0428

- *Himantura uarnak*
- Accompanied by needlefish

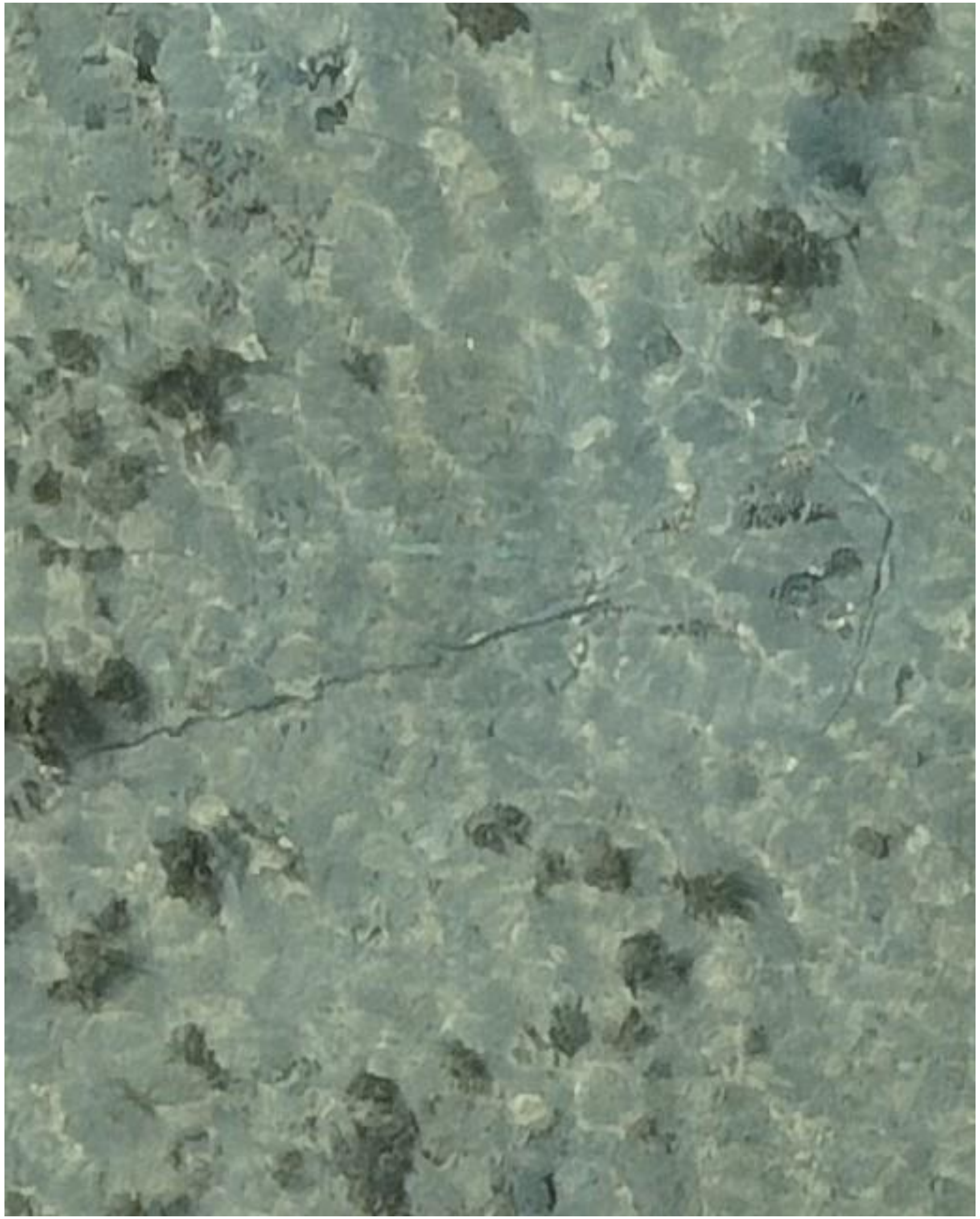

DJI_20241118082932_0482

- Whipray
- Best guess: *H. uarnak*

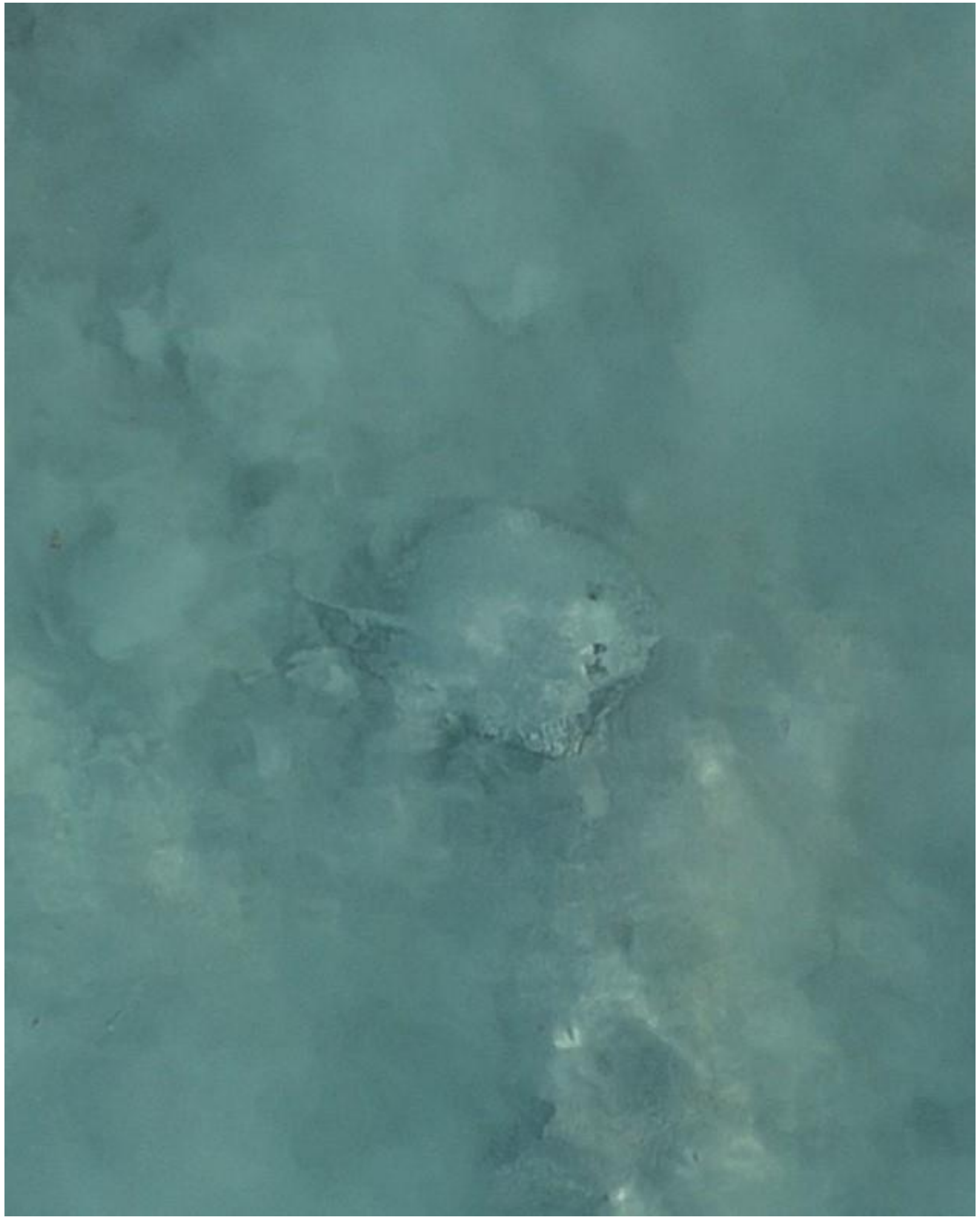

DJI_20241118083303_0676

- Whipray

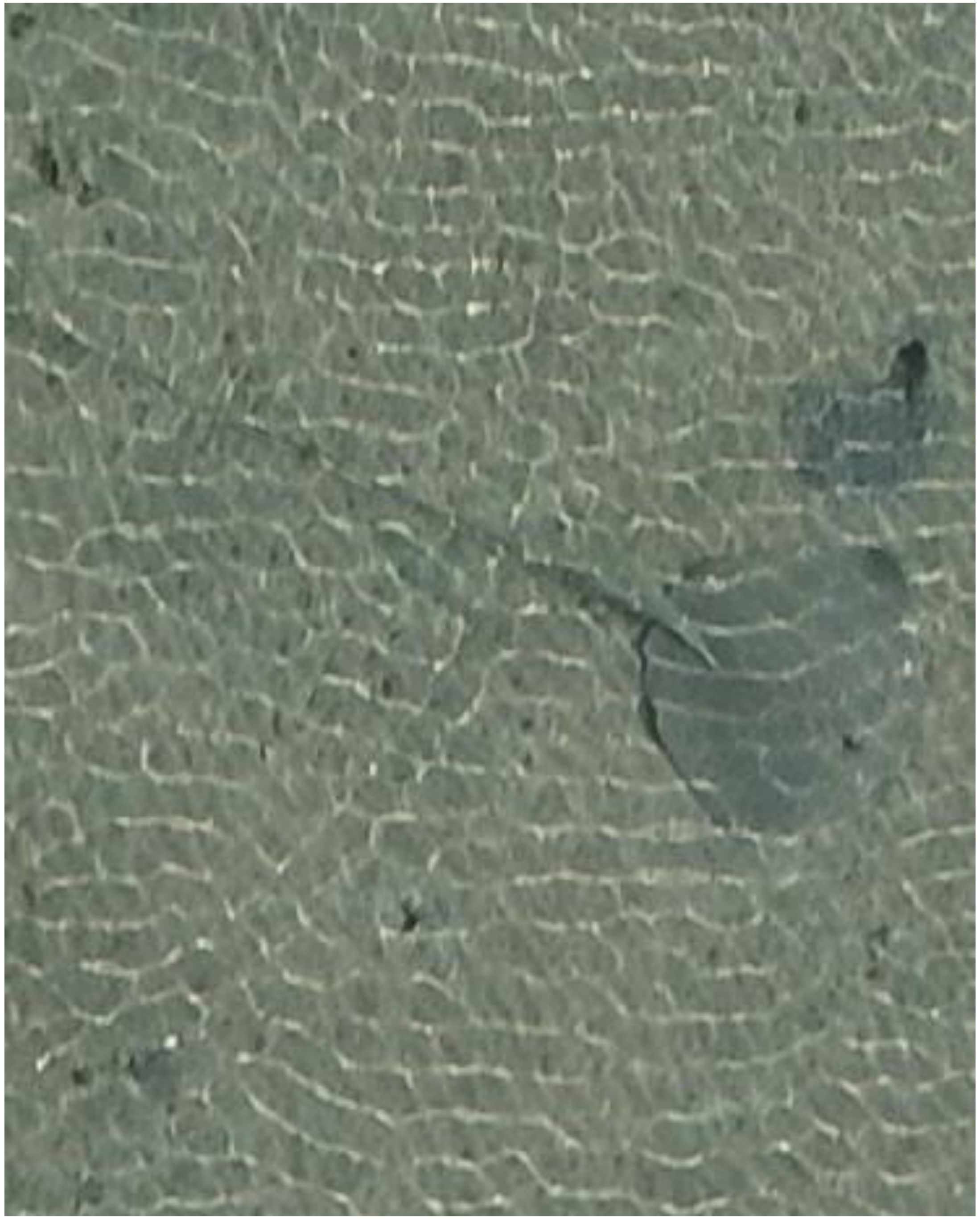
DJI_20241118083339_0712

- Whipray
- Accompanied by needlefish

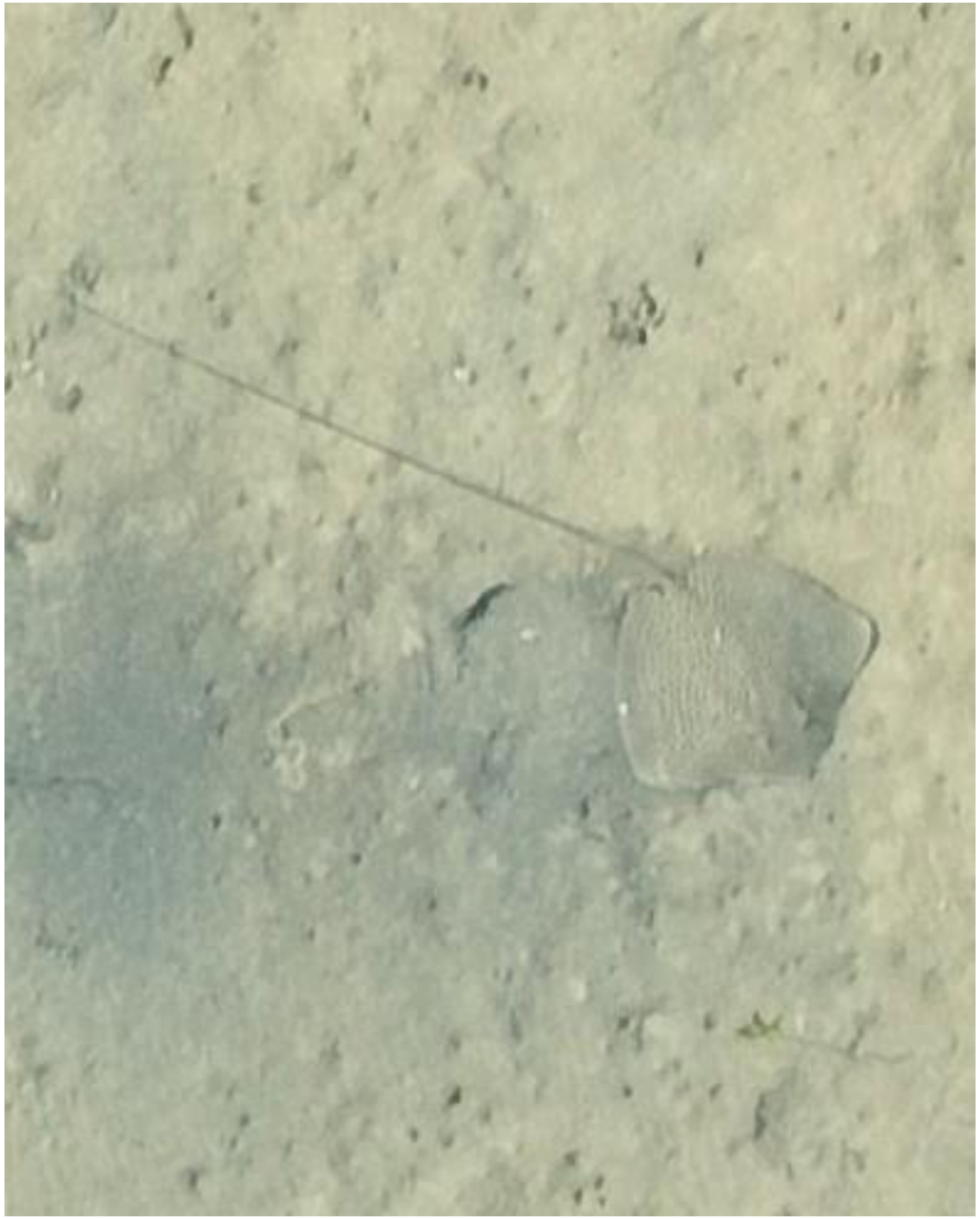

DJI_20241118083419_0740

- *Himantura uarnak*

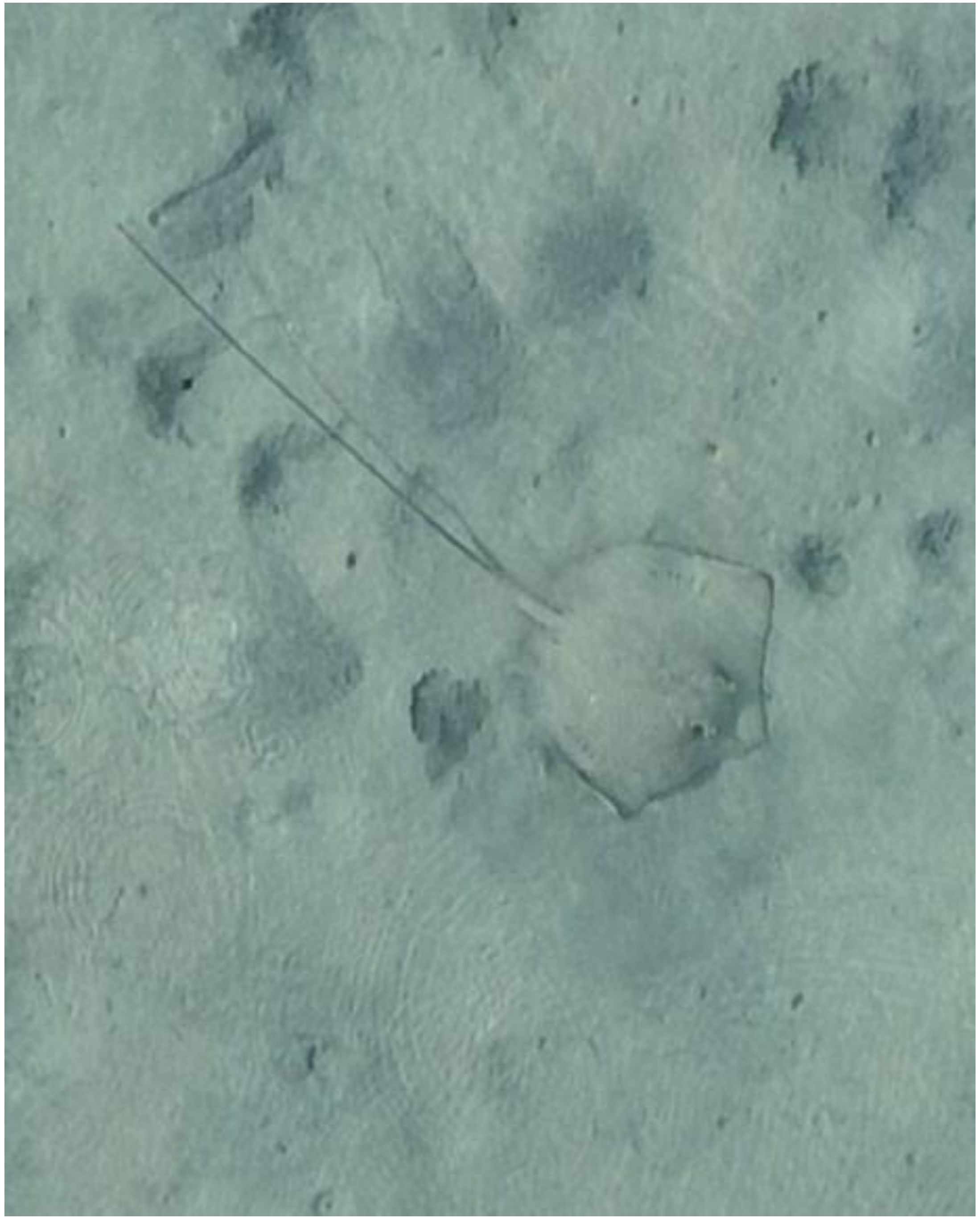

- Whipray

DJI_20241118083836_0972
