## Appendix F: maps tracking surveys for "Fine-scale habitat partitioning of sympatric stingrays revealed by drone-based remote sensing and deep learning"

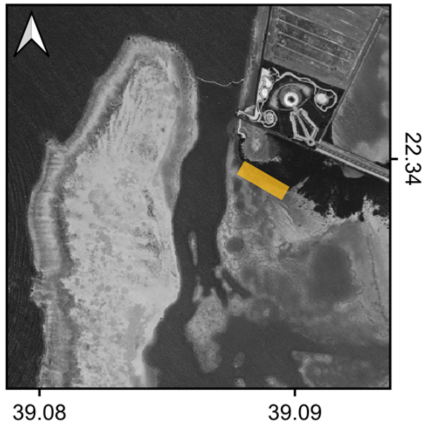


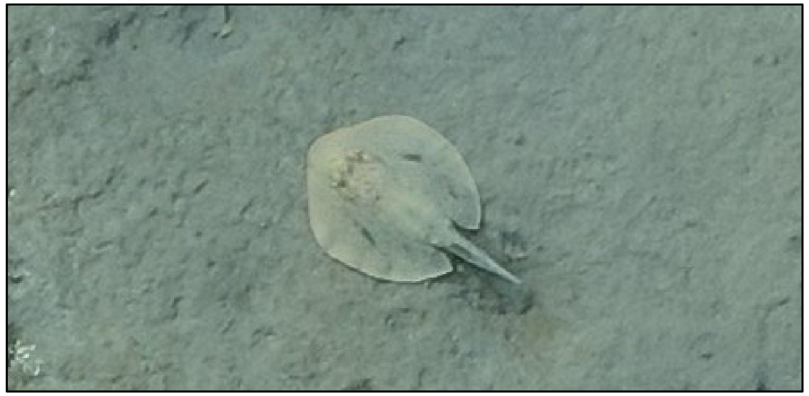

Species: *Taeniura lymma*

Date: 2024-04-28

Disc width: 26 cm

Disc length: 26 cm

Total length: 40 cm

Notes:

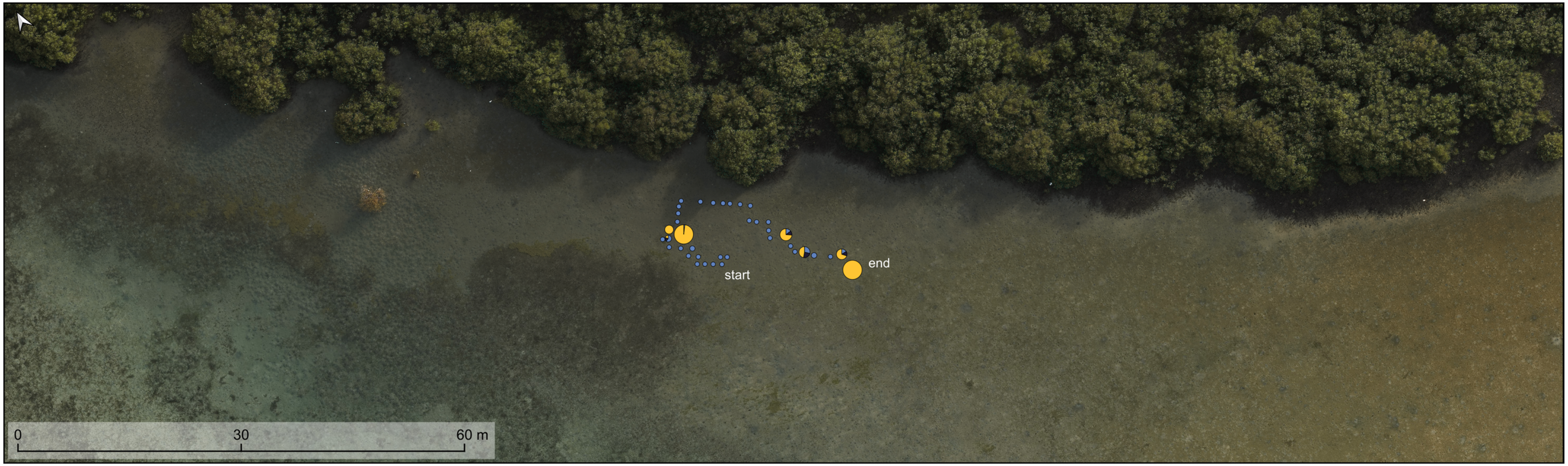


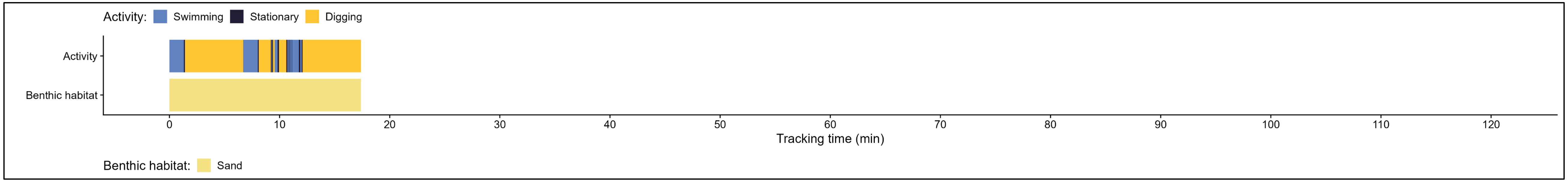

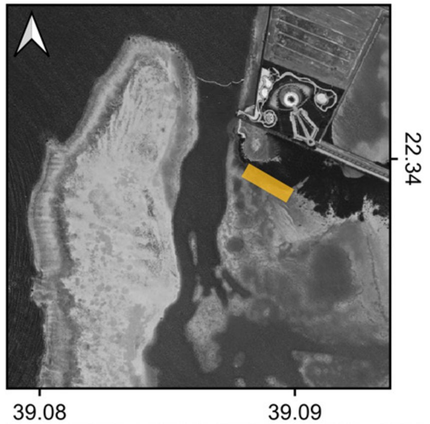


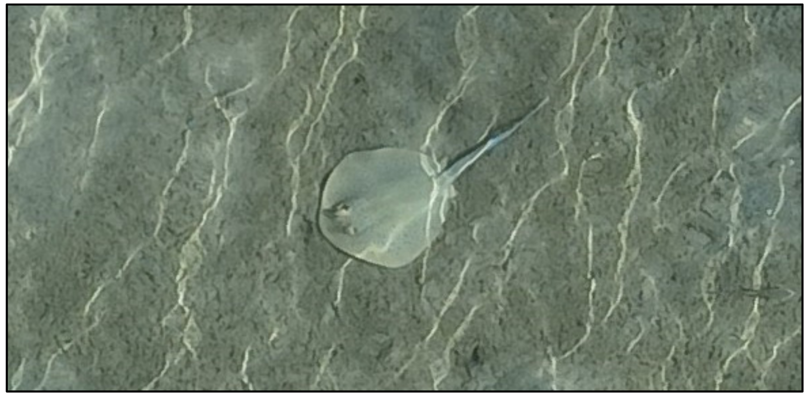

| | |
|---|---|
| Species: | *Taeniura lymma* |
| Date: | 2024-05-01 |
| Disc width: | 29 cm |
| Disc length: | 30 cm |
| Total length: | 61 cm |
| Notes: | |

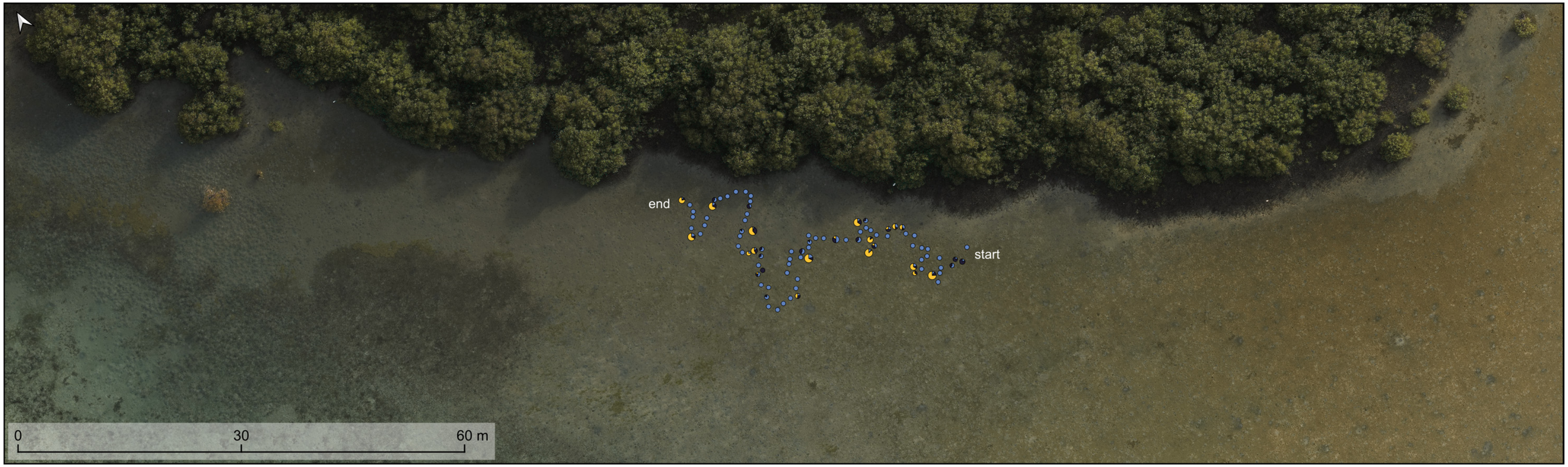


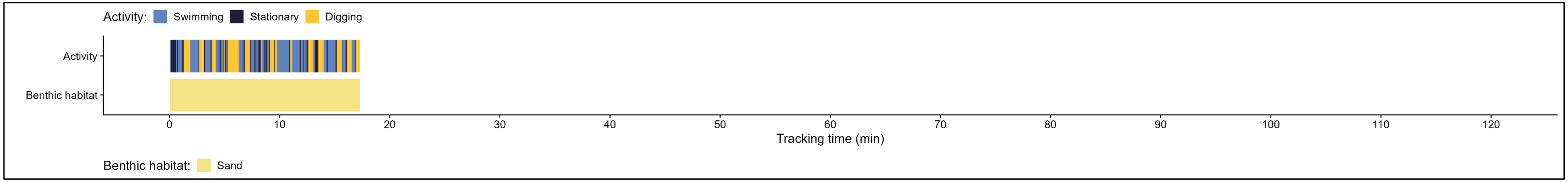

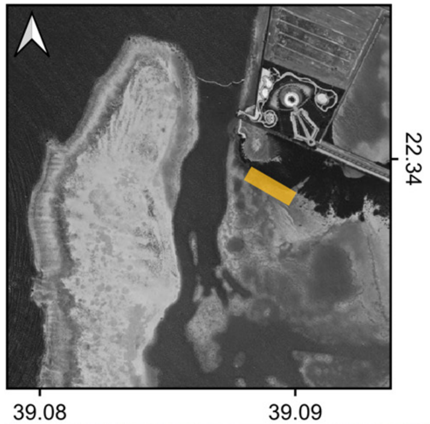


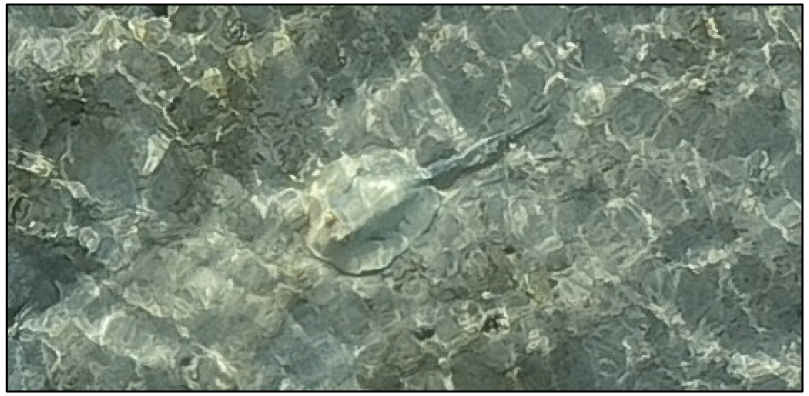

Species: *Taeniura lymma*

Date: 2024-05-02

Disc width: 25 cm

Disc length: 29 cm

Total length: 62 cm

Notes:

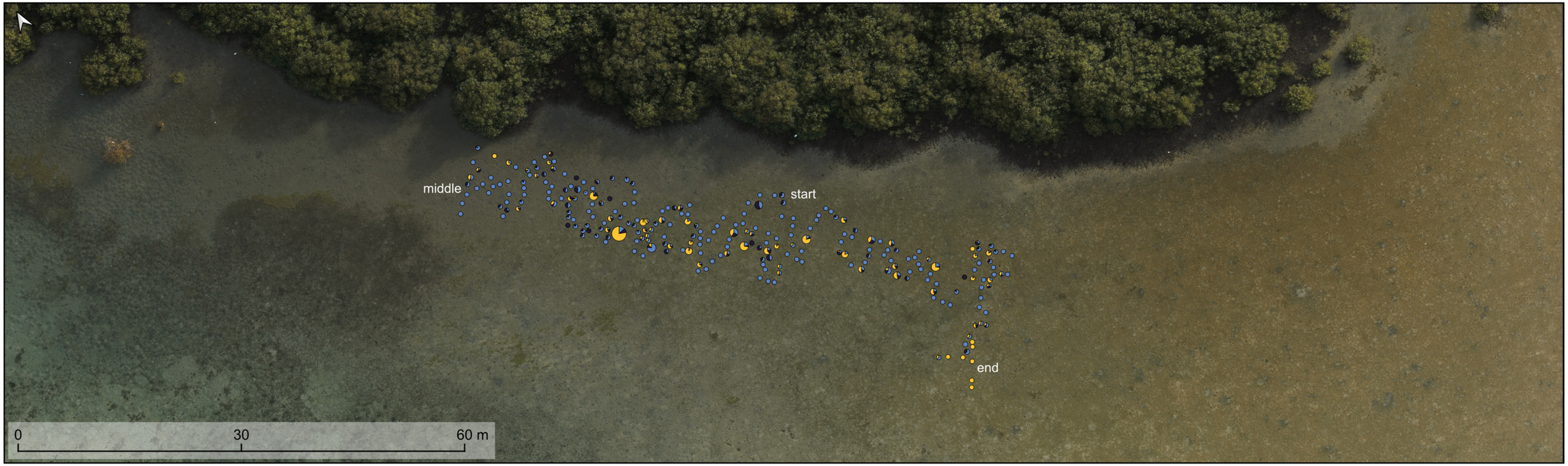


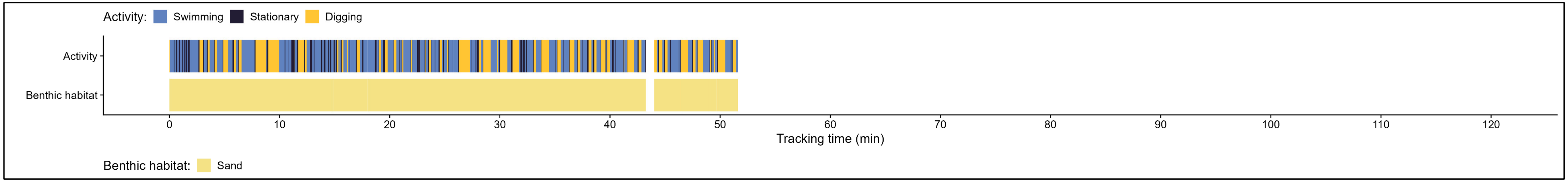

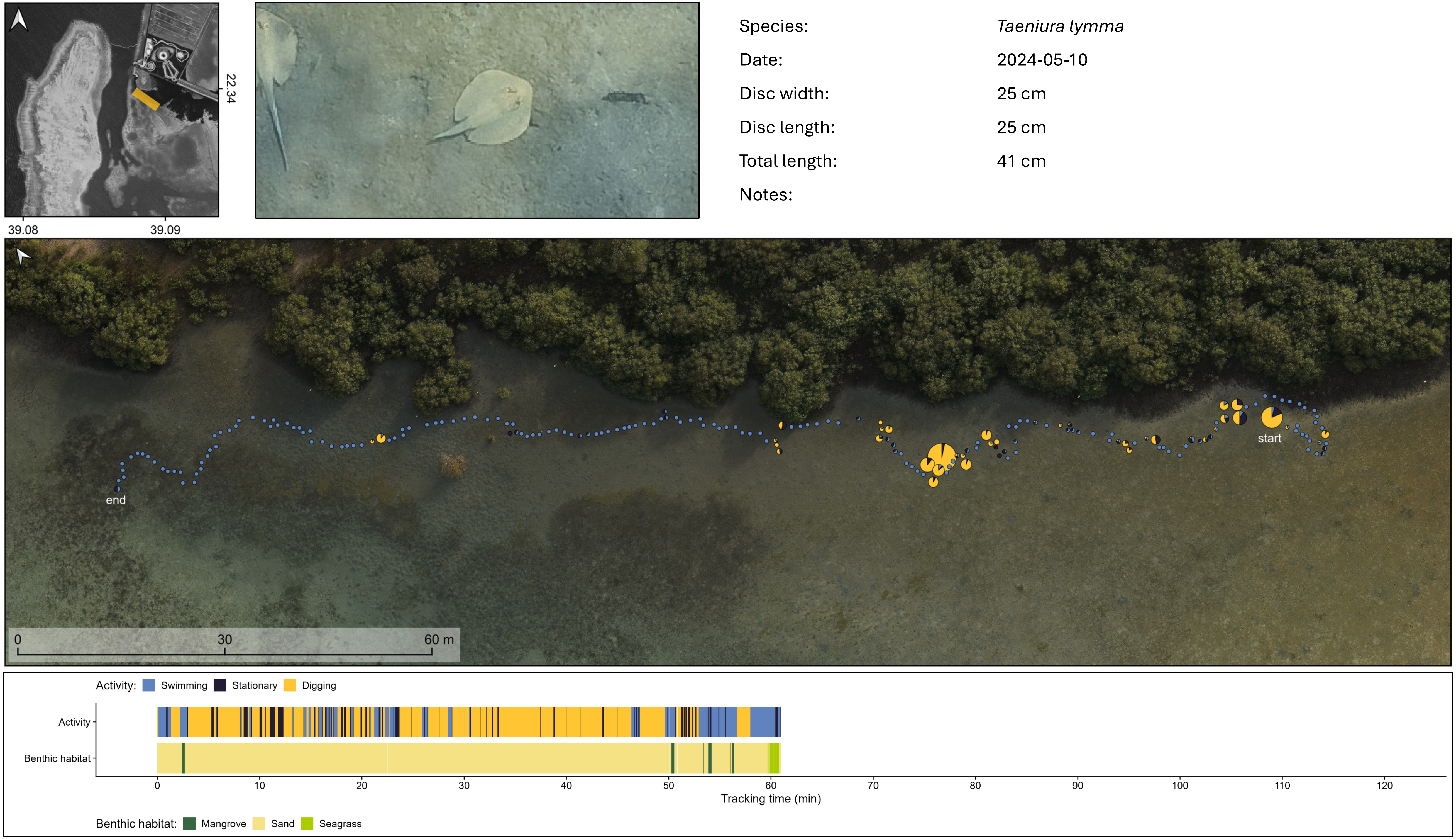

22.34
39.08
39.09
Species:
Taeniura lymma
Date:
2024-05-10
Disc width:
25 cm
Disc length:
25 cm
Total length:
41 cm
Notes:
start
end
0
30
60 m
Activity: Swimming Stationary Digging
Activity
Benthic habitat
0 10 20 30 40 50 60 70 80 90 100 110 120
Tracking time (min)
Benthic habitat: Mangrove Sand Seagrass

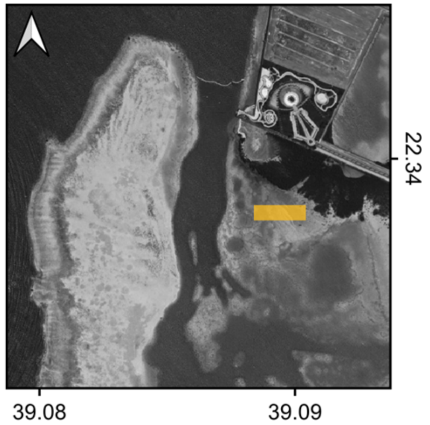


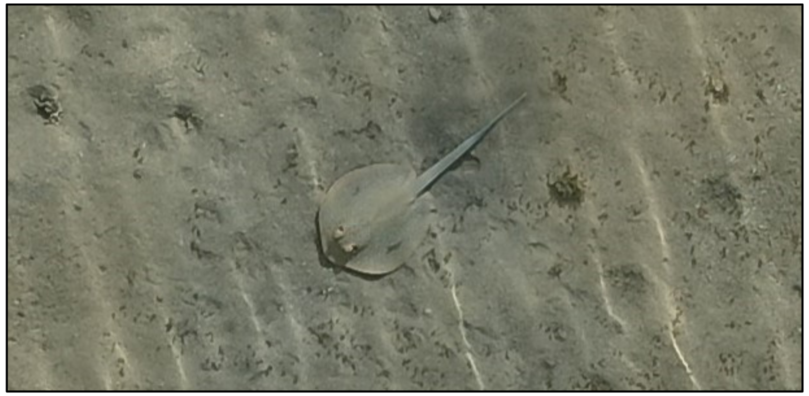

Species: *Taeniura lymma*

Date: 2024-05-17 (a)

Disc width: 23 cm

Disc length: 26 cm

Total length: 60 cm

Notes:

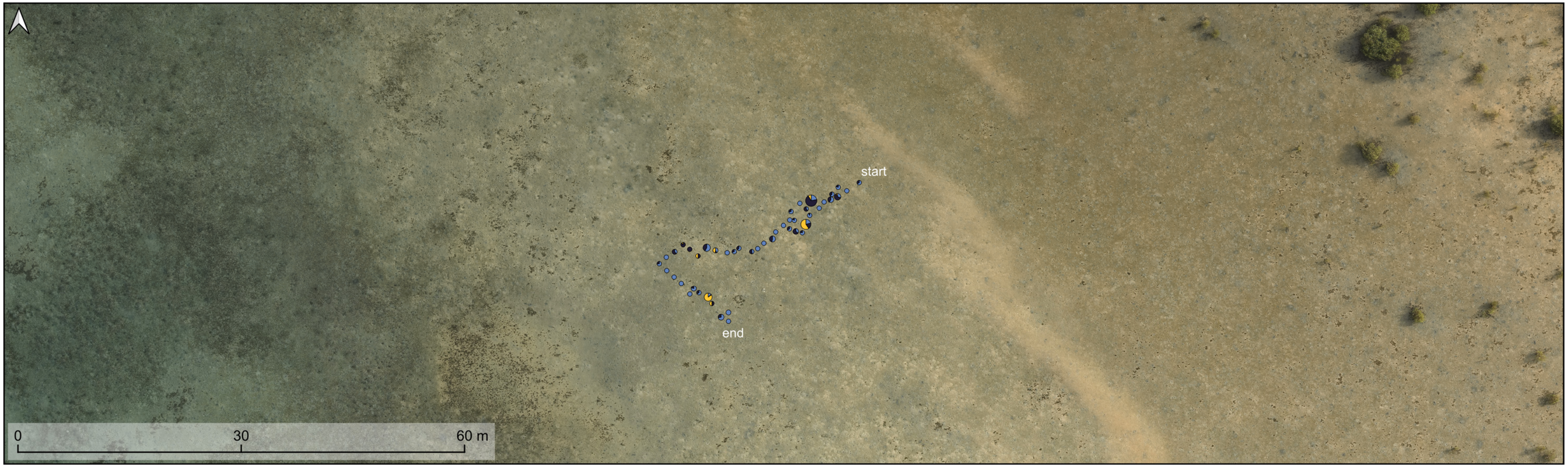


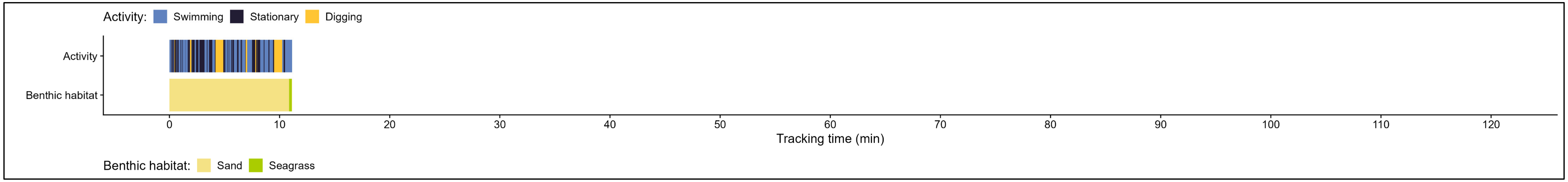

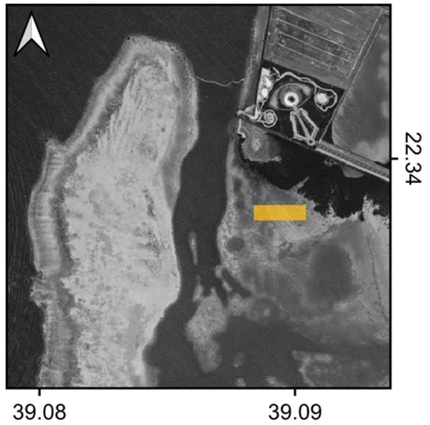


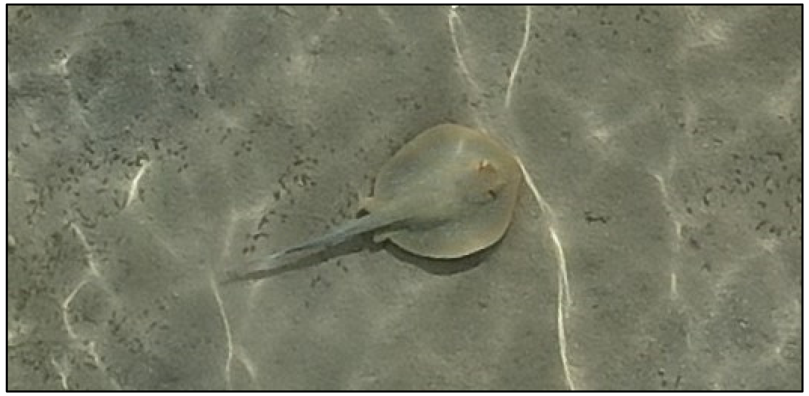

| | |
|---|---|
| Species: | *Taeniura lymma* |
| Date: | 2024-05-17 (b) |
| Disc width: | 25 cm |
| Disc length: | 27 cm |
| Total length: | 62 cm |
| Notes: | |

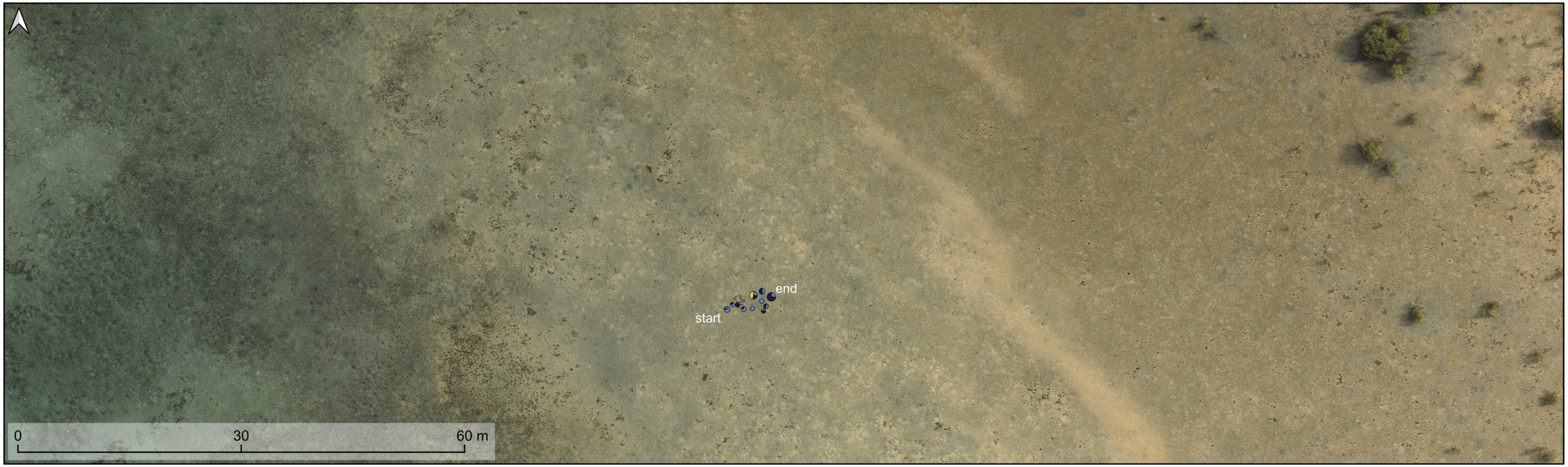


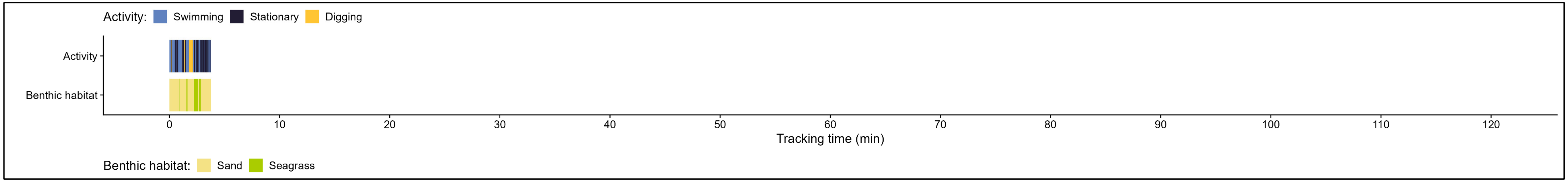

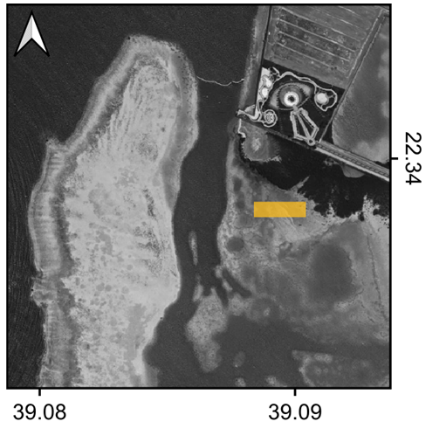


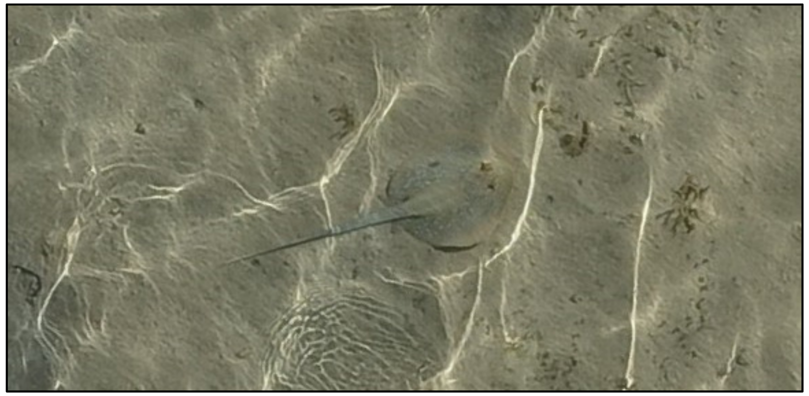

Species: *Taeniura lymma*

Date: 2024-05-17 (c)

Disc width: 25 cm

Disc length: 24 cm

Total length: 66 cm

Notes:

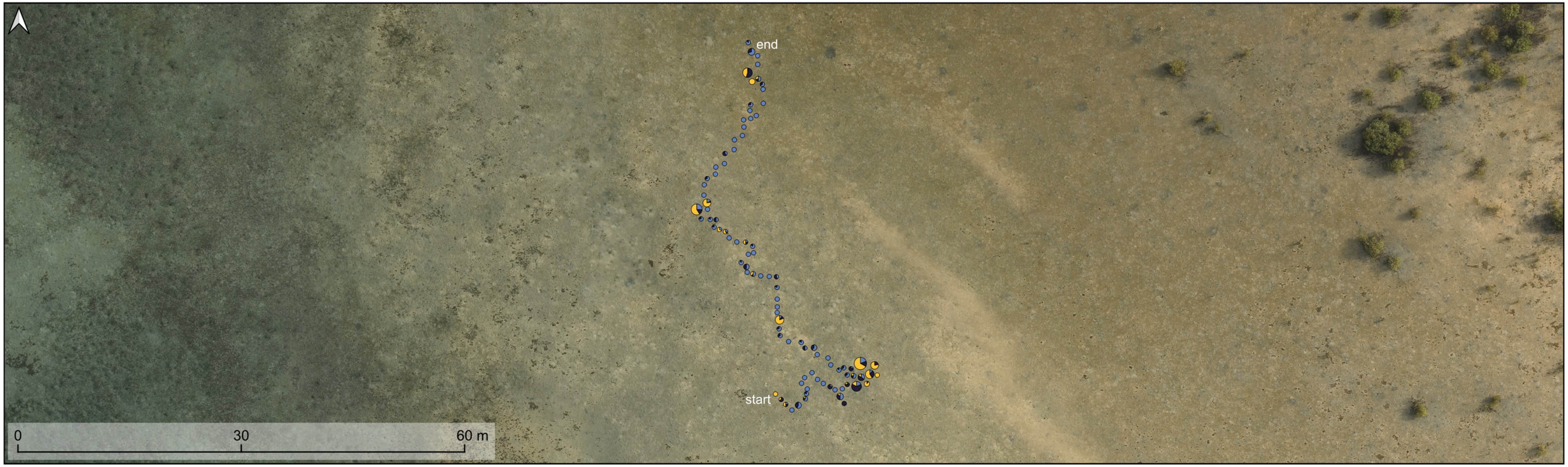


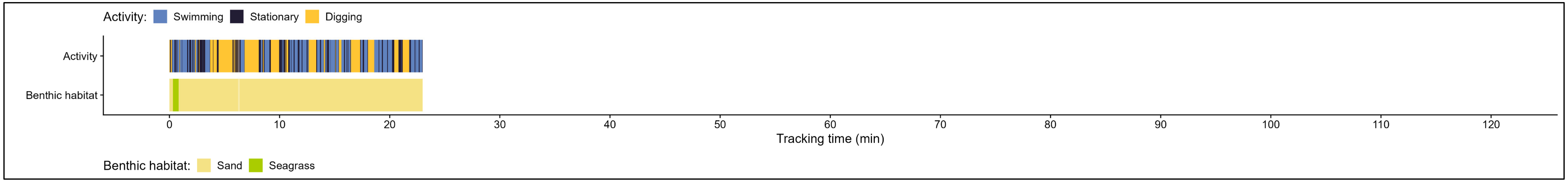

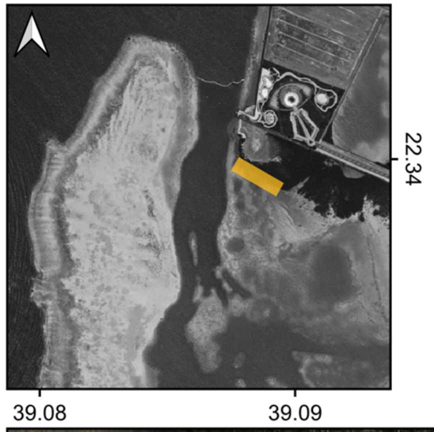


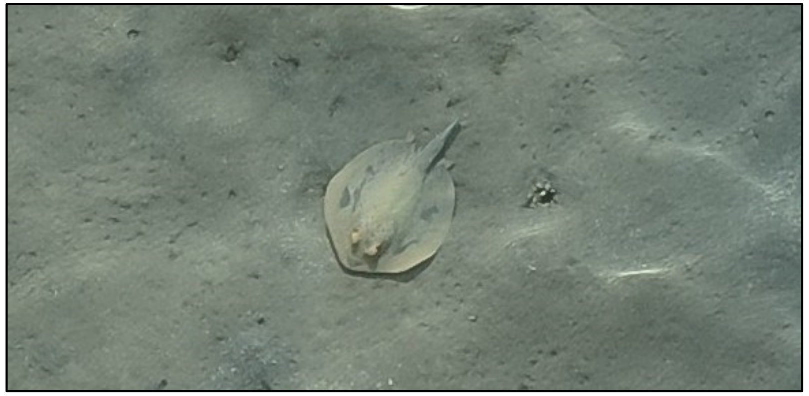

| | |
|---|---|
| Species: | *Taeniura lymma* |
| Date: | 2024-05-23 (a) |
| Disc width: | 27 cm |
| Disc length: | 27 cm |
| Total length: | 40 cm |
| Notes: | |

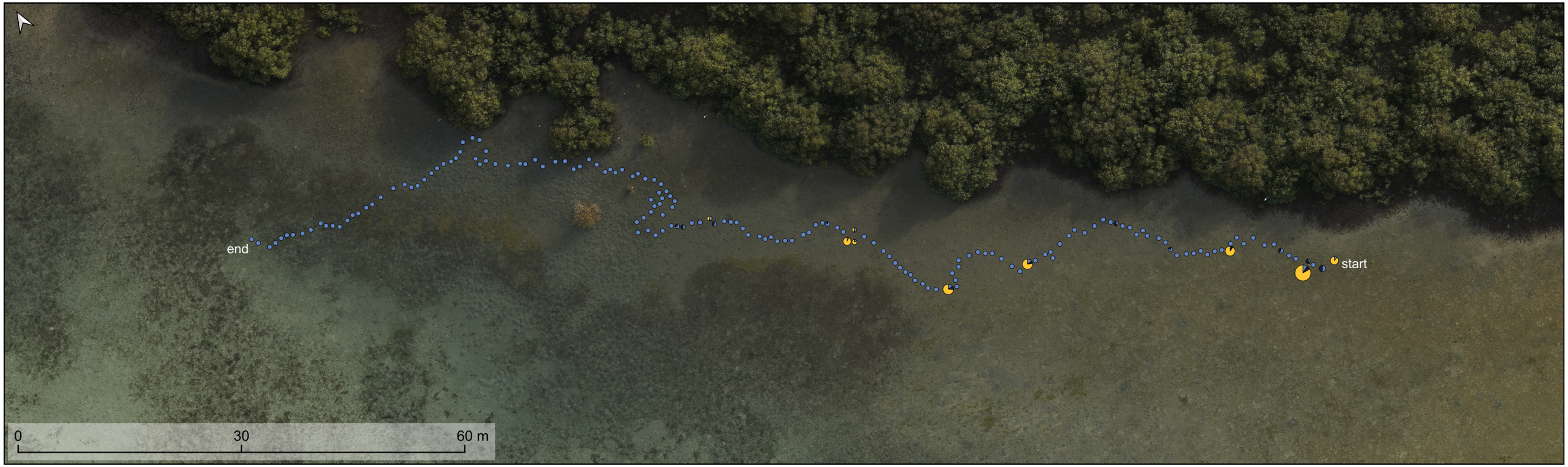


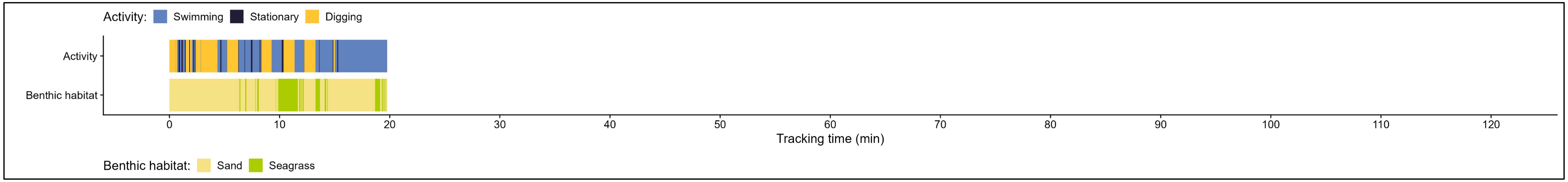

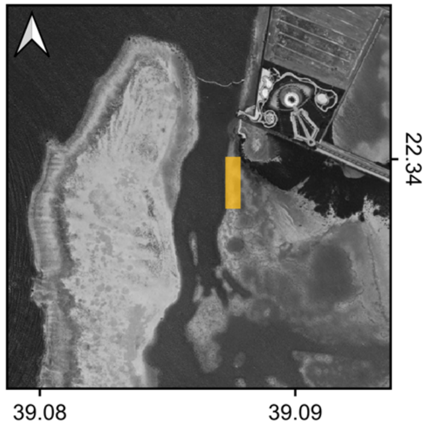


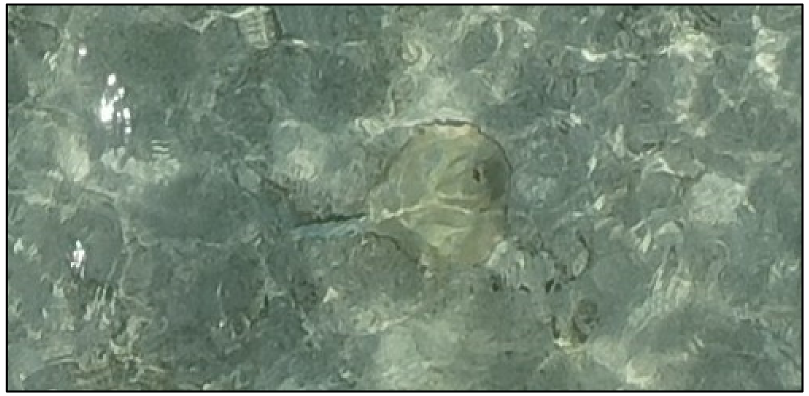

| | |
|---|---|
| Species: | *Taeniura lymma* |
| Date: | 2024-05-23 (b) |
| Disc width: | 29 cm |
| Disc length: | 29 cm |
| Total length: | NA not measurable, due to too much wave distortion. |
| Notes: | |

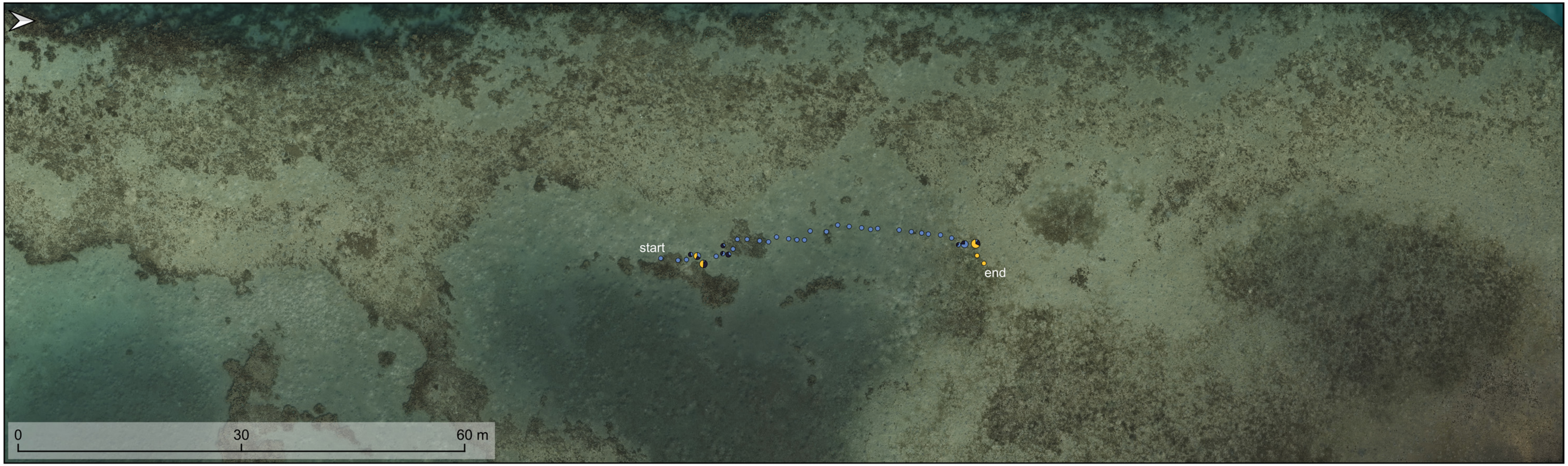


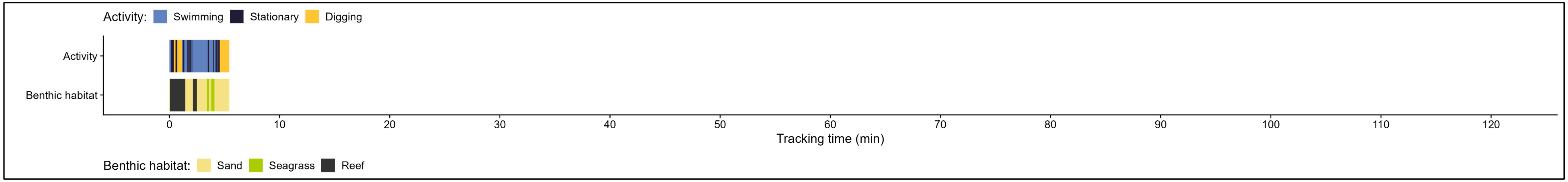

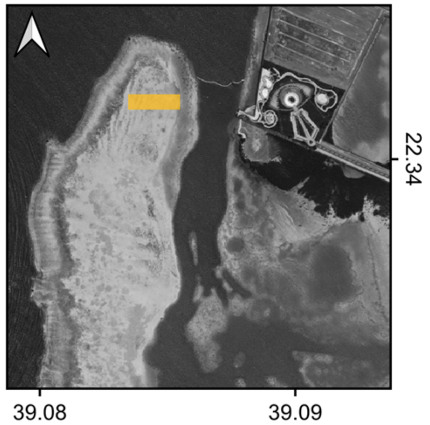

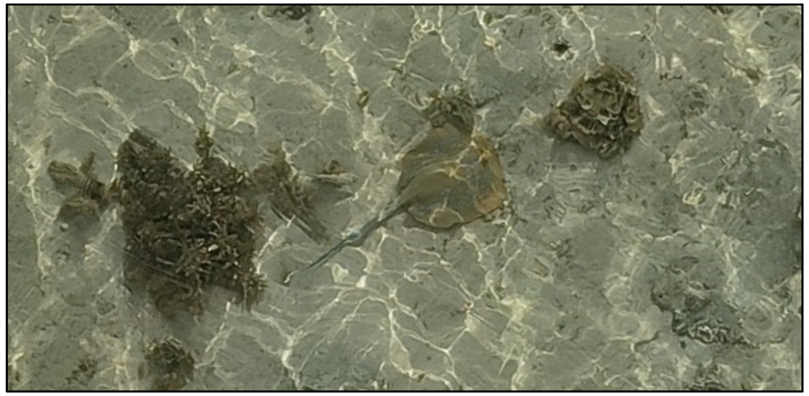

Species: *Taeniura lymma*
Date: 2024-06-03 (a)
Disc width: 24 cm
Disc length: 23 cm
Total length: 60 cm
Notes:

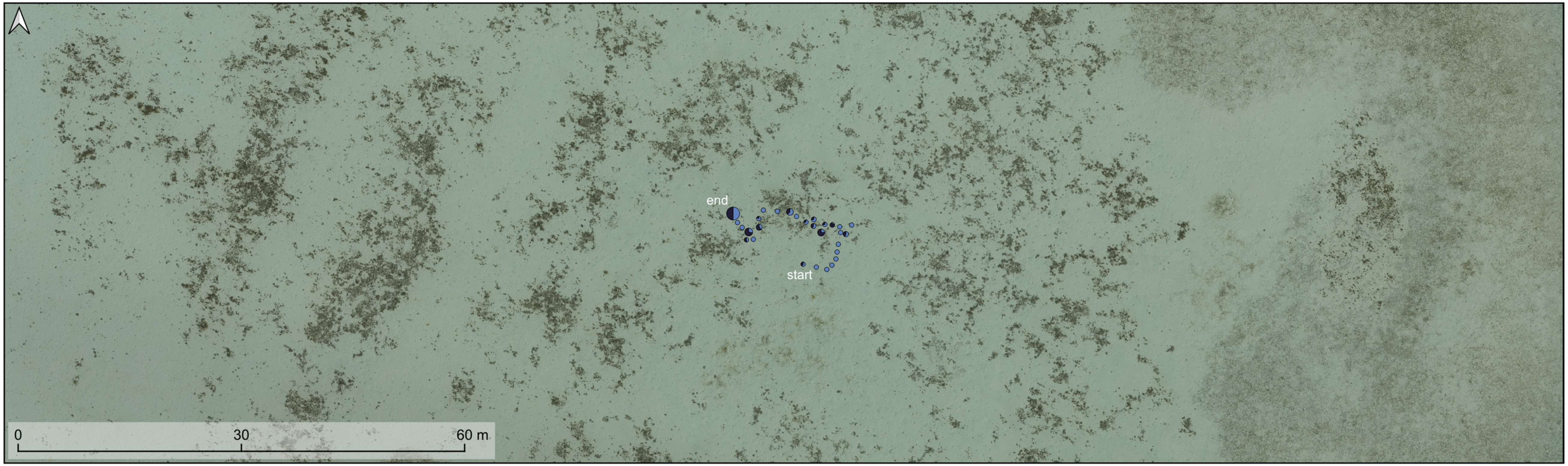

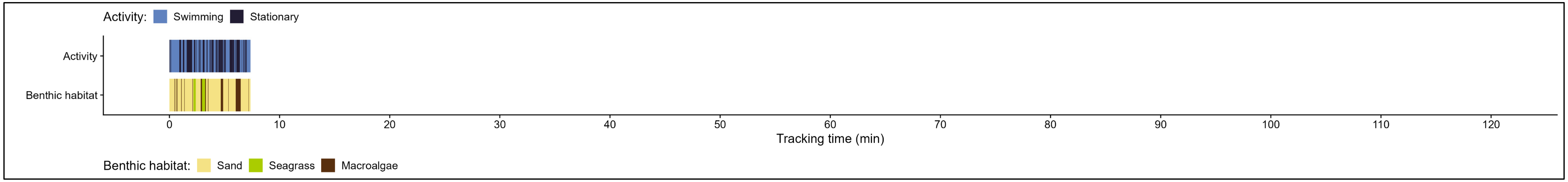

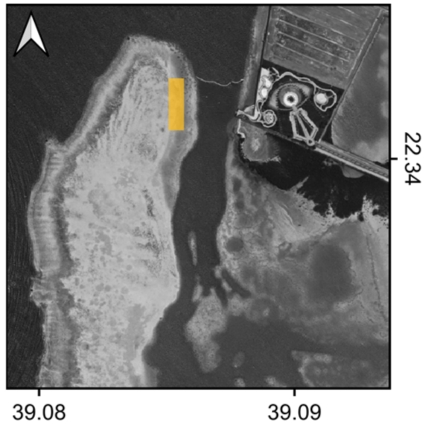


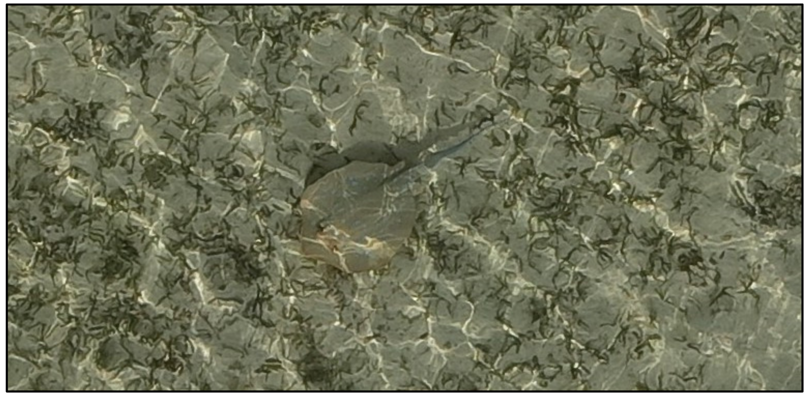

| | |
|---|---|
| Species: | *Taeniura lymma* |
| Date: | 2024-06-03 (b) |
| Disc width: | 26 cm |
| Disc length: | 30 cm |
| Total length: | 62 cm |
| Notes: | |

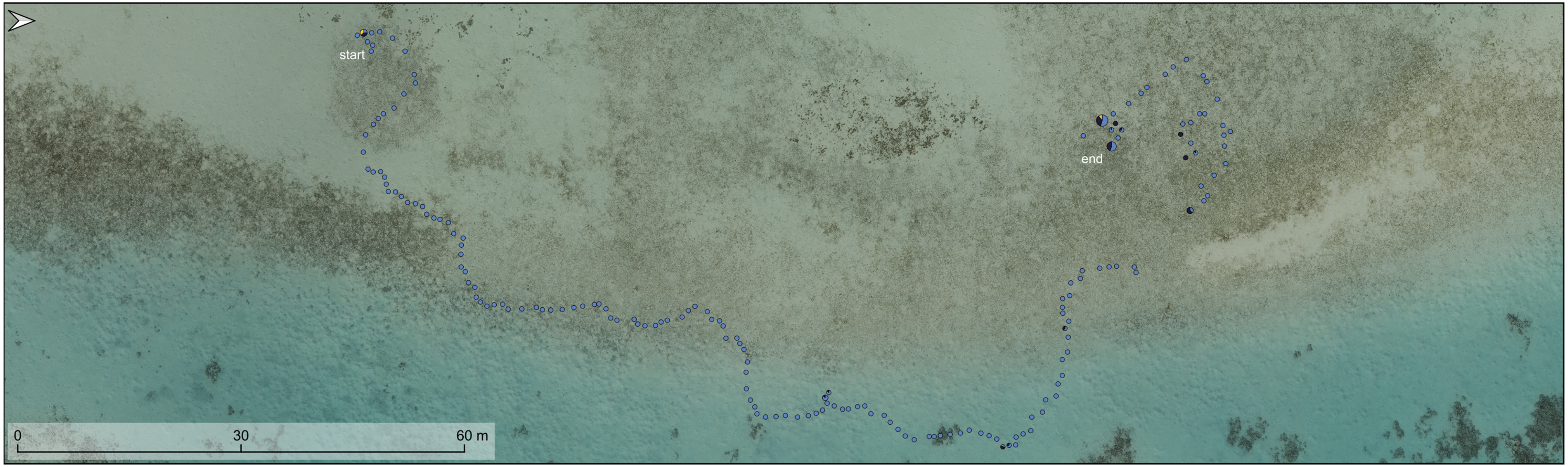


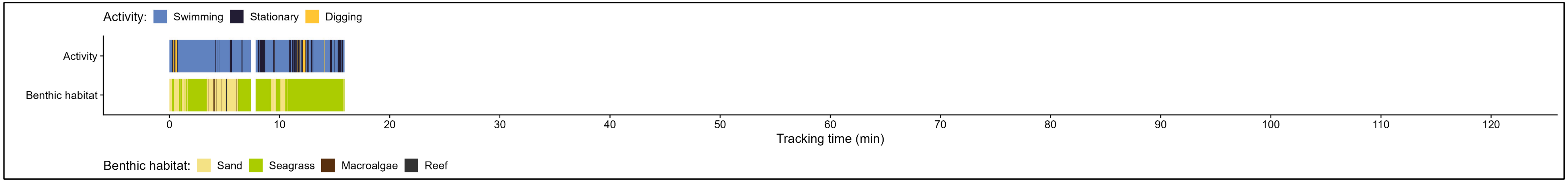

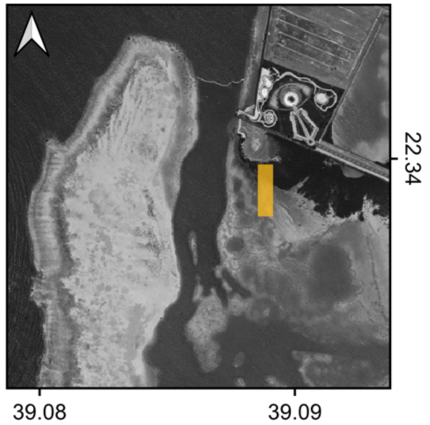


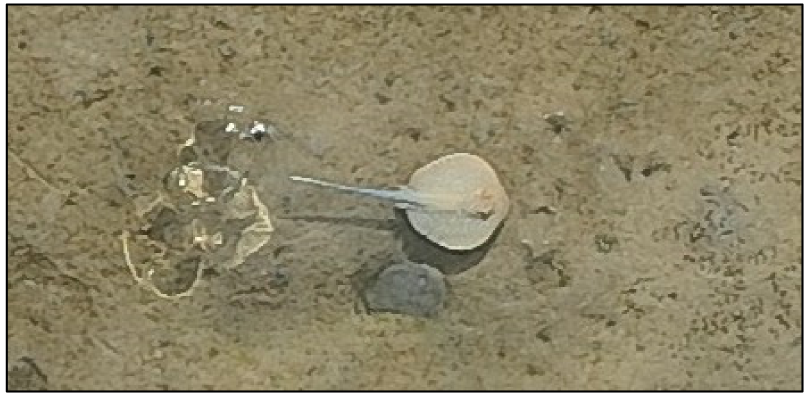

Species: *Taeniura lymma*

Date: 2024-06-03 (c)

Disc width: 16 cm

Disc length: 16 cm

Total length: 37 cm

Notes:

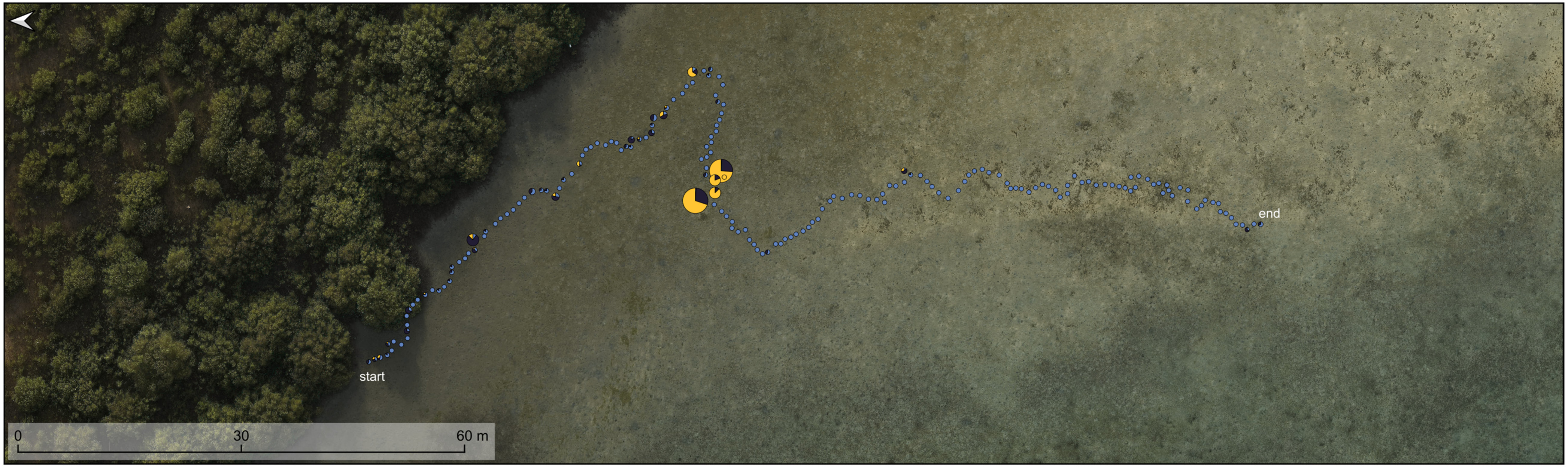


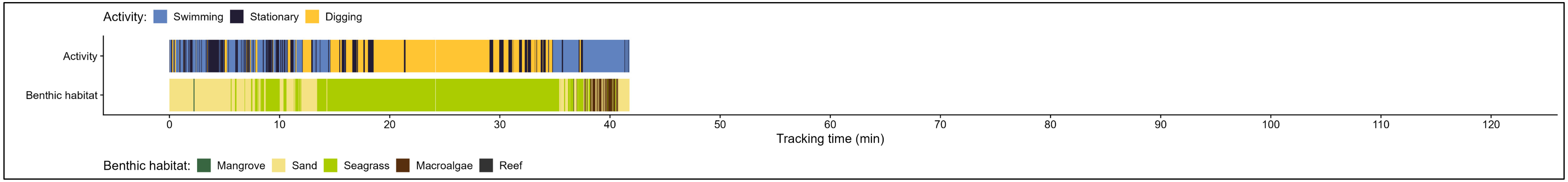

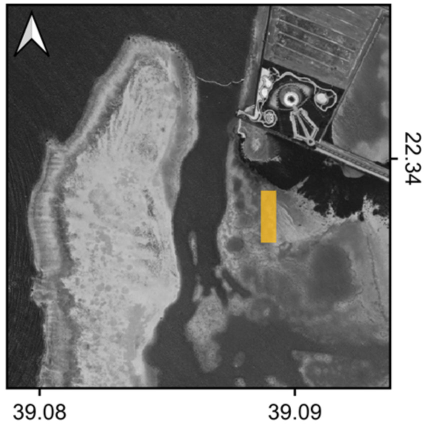


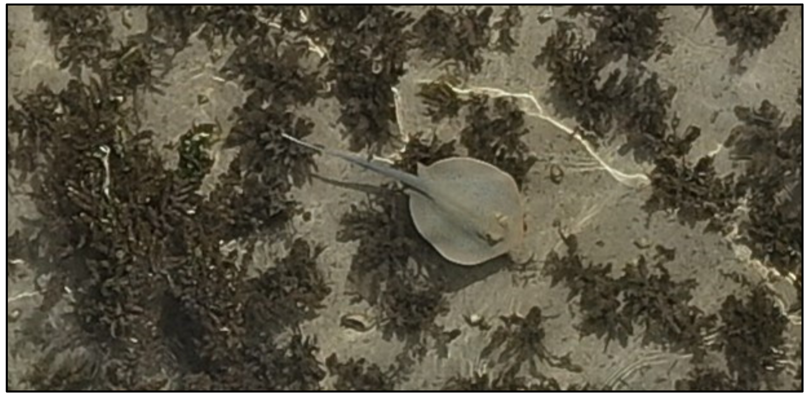

Species: *Taeniura lymma*

Date: 2024-06-03 (d)

Disc width: 19 cm

Disc length: 21 cm

Total length: 49 cm

Notes:

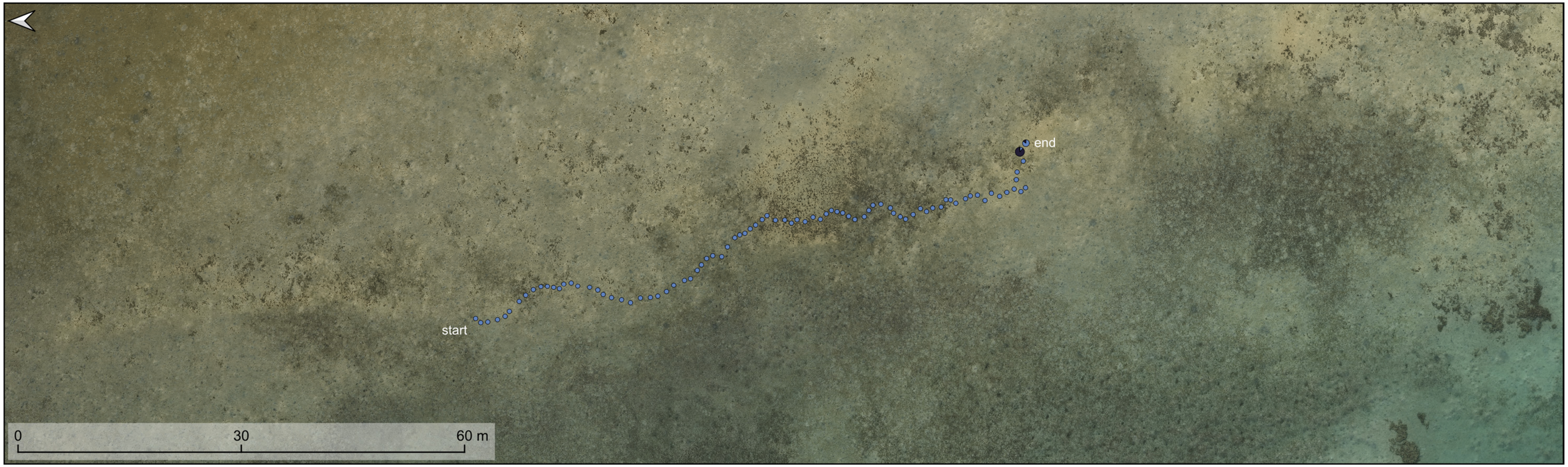


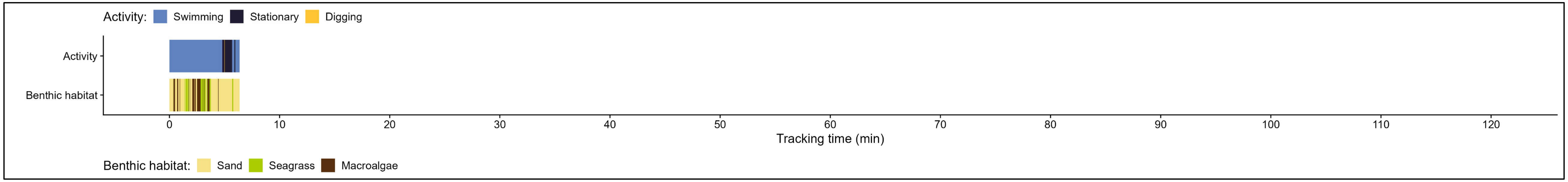

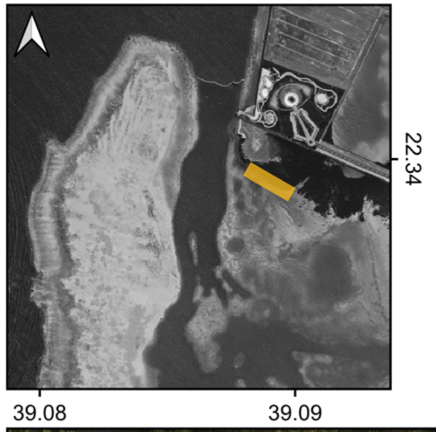


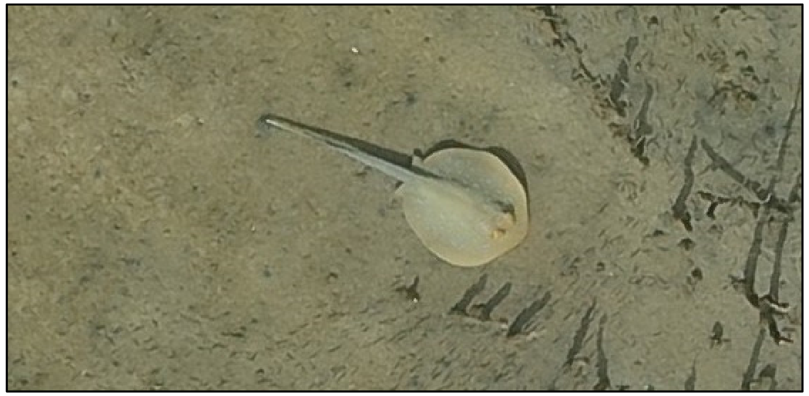

Species: *Taeniura lymma*

Date: 2024-06-06

Disc width: 24 cm

Disc length: 27 cm

Total length: 61 cm

Notes:

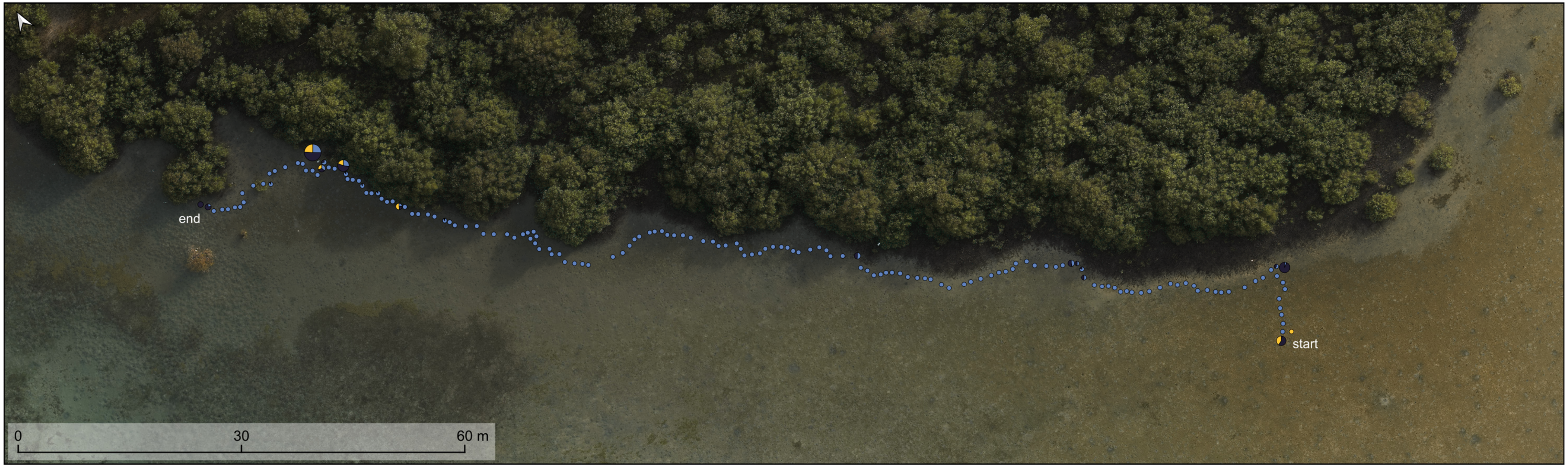


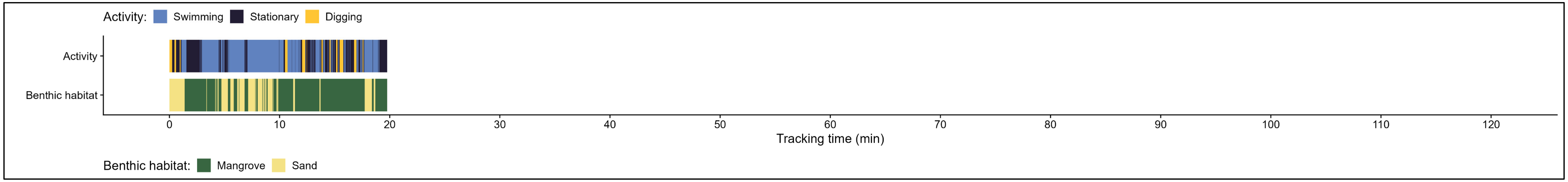

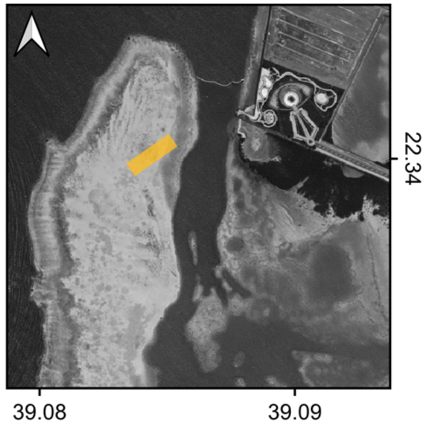


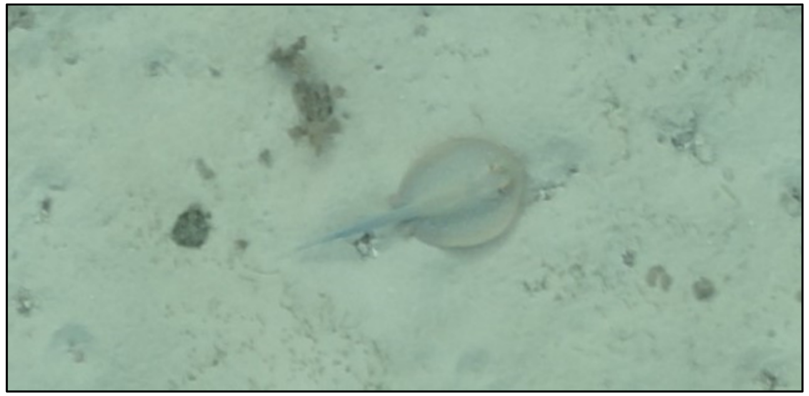

Species: *Taeniura lymma*

Date: 2024-06-10 (a)

Disc width: 27 cm

Disc length: 30 cm

Total length: 65 cm

Notes:

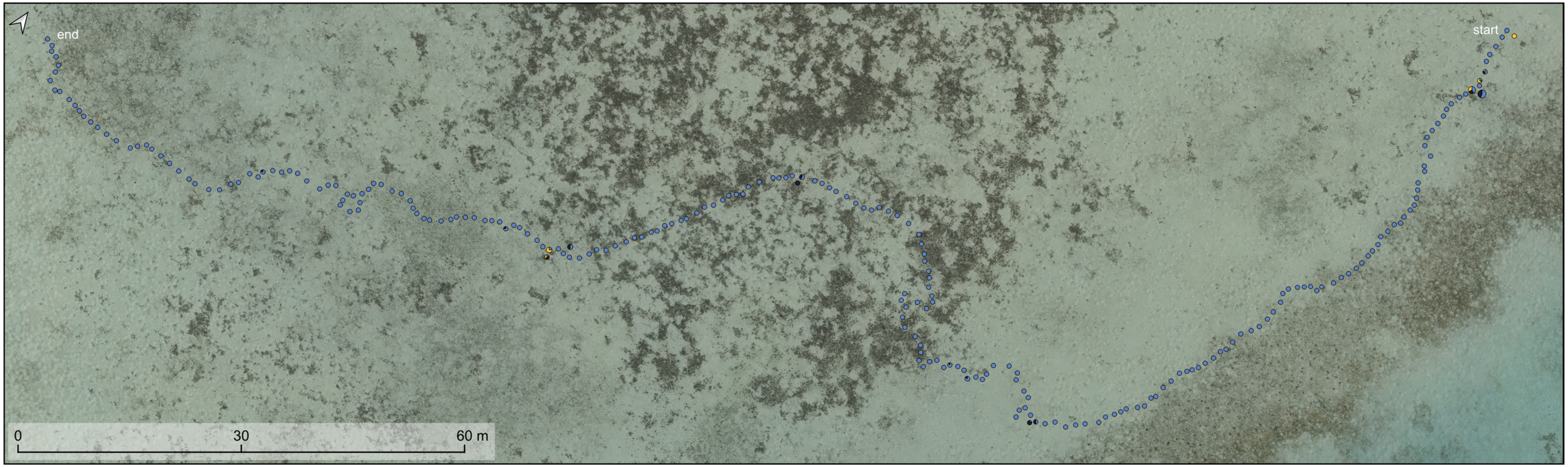


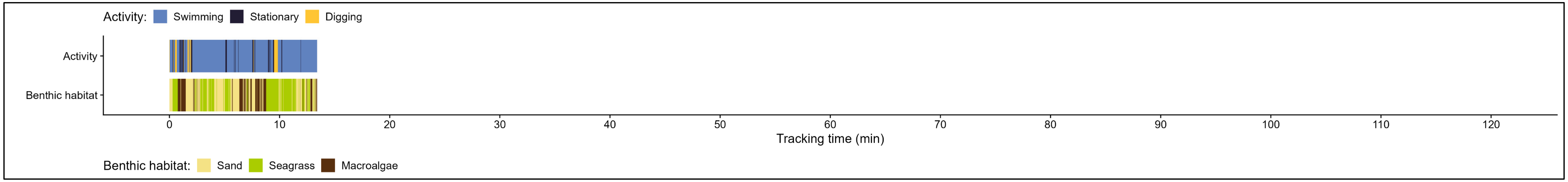

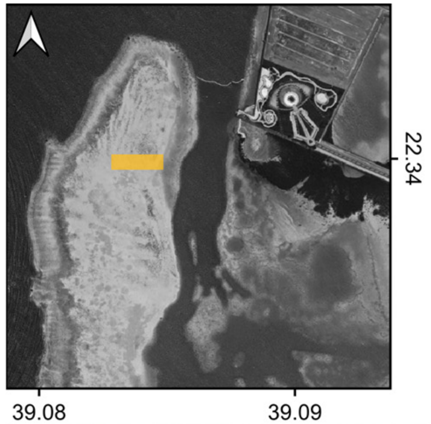


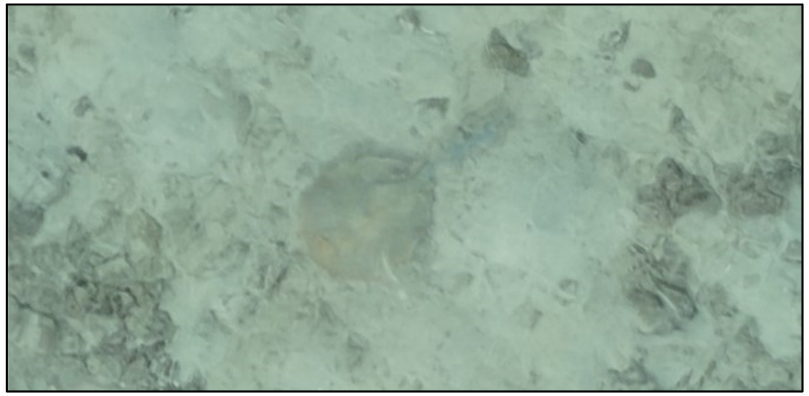

| | |
|---|---|
| Species: | *Taeniura lymma* |
| Date: | 2024-06-10 (b) |
| Disc width: | 32 cm |
| Disc length: | 33 cm |
| Total length: | NA not measurable, due to too much wave distortion. |
| Notes: | |

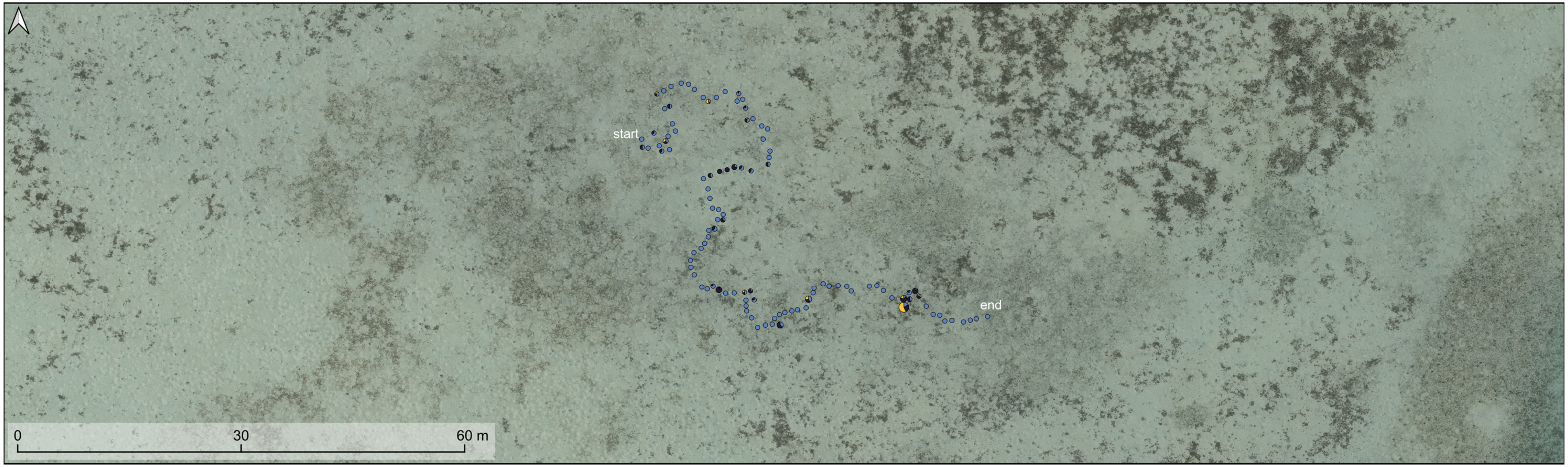


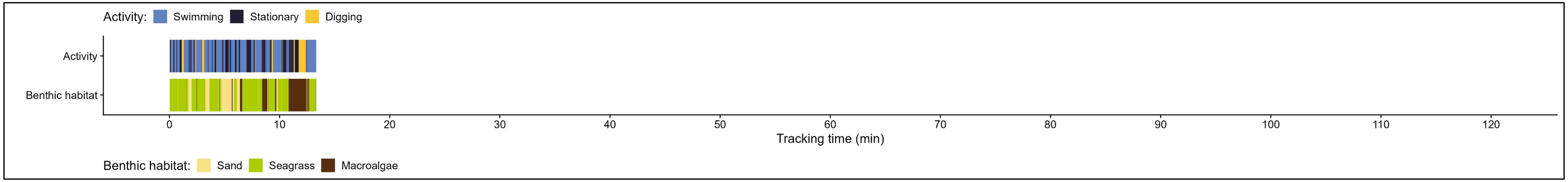

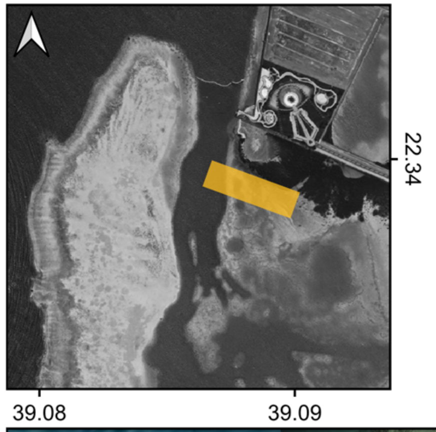


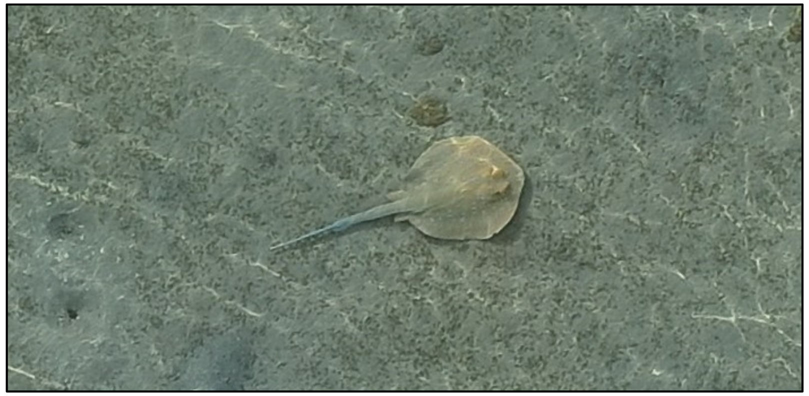

| | |
|---|---|
| Species: | *Taeniura lymma* |
| Date: | 2024-06-11 |
| Disc width: | 27 cm |
| Disc length: | 30 cm |
| Total length: | 68 cm |
| Notes: | |

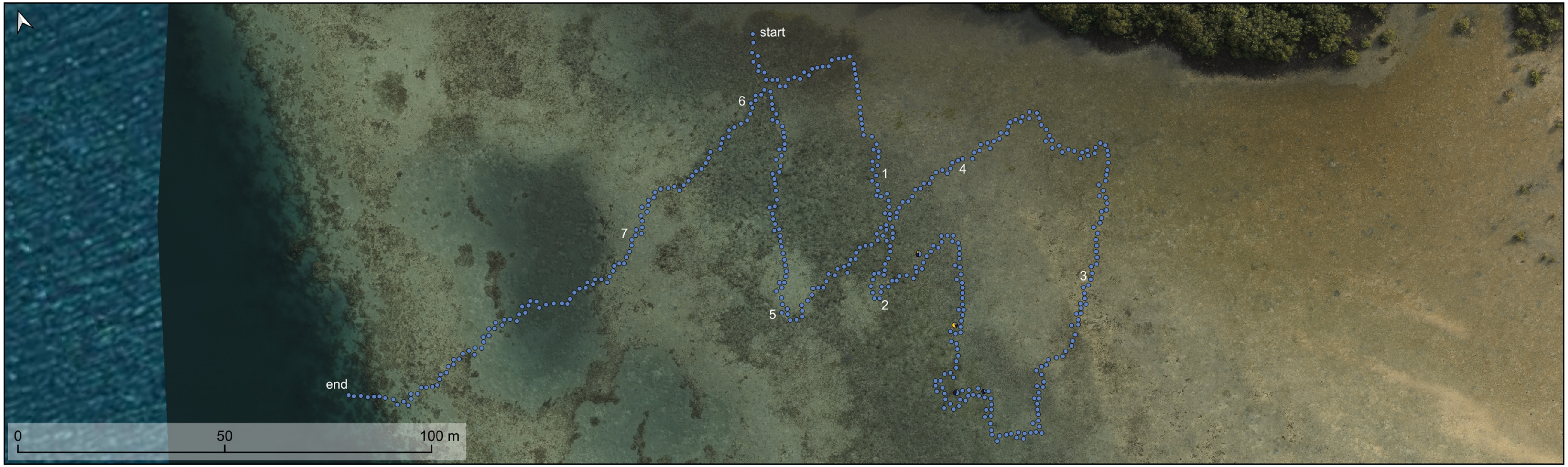


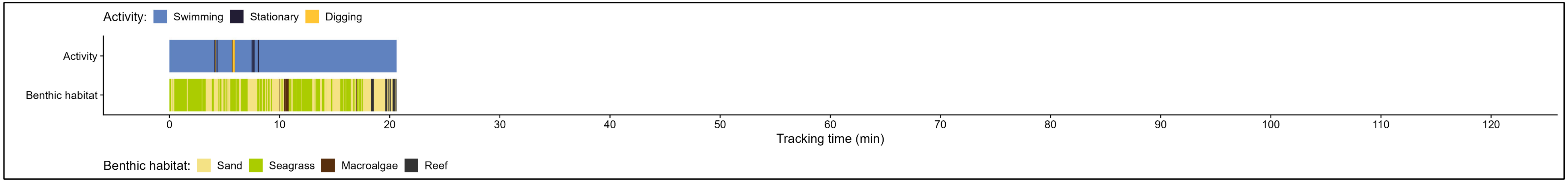

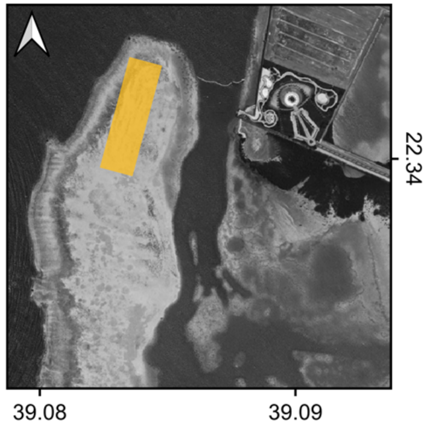


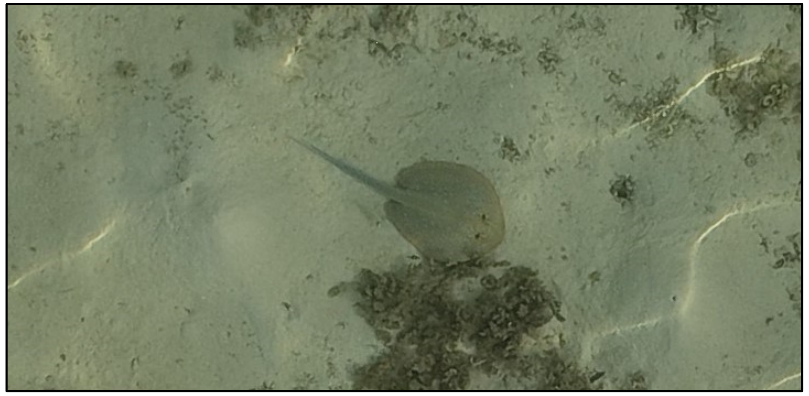

| | |
|---|---|
| Species: | *Taeniura lymma* |
| Date: | 2024-06-19 |
| Disc width: | 28 cm |
| Disc length: | 31 cm |
| Total length: | 67 cm |
| Notes: | |

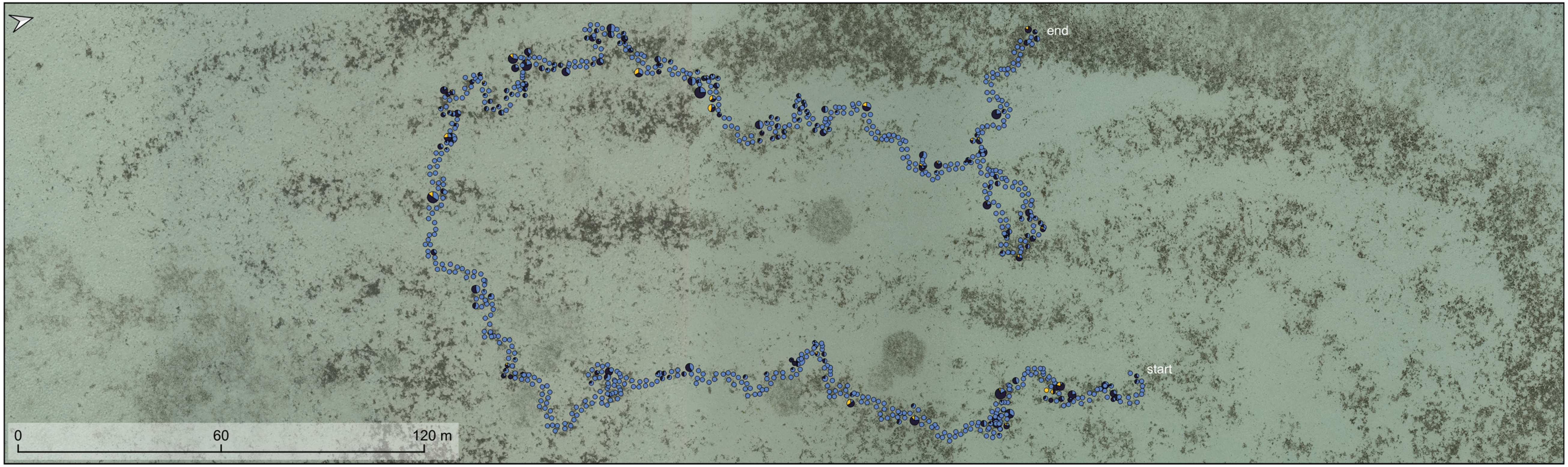


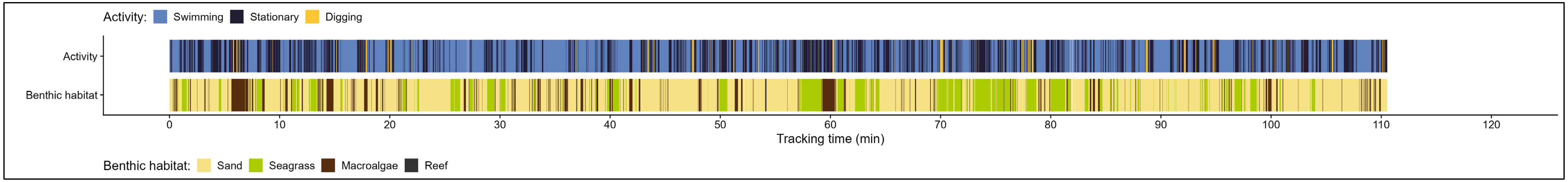

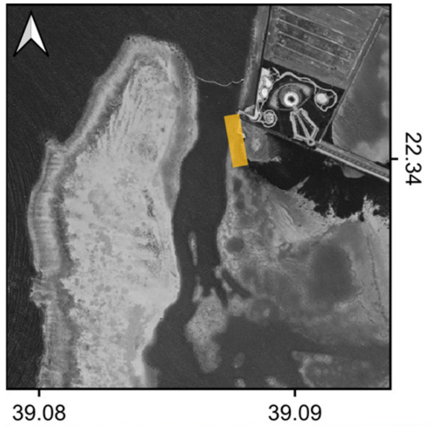


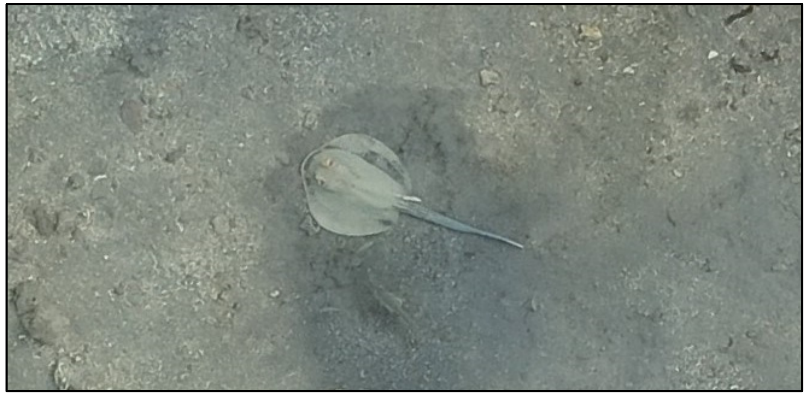

| | |
|---|---|
| Species: | *Taeniura lymma* |
| Date: | 2024-06-24 |
| Disc width: | 25 cm |
| Disc length: | 24 cm |
| Total length: | 57 cm |
| Notes: | |

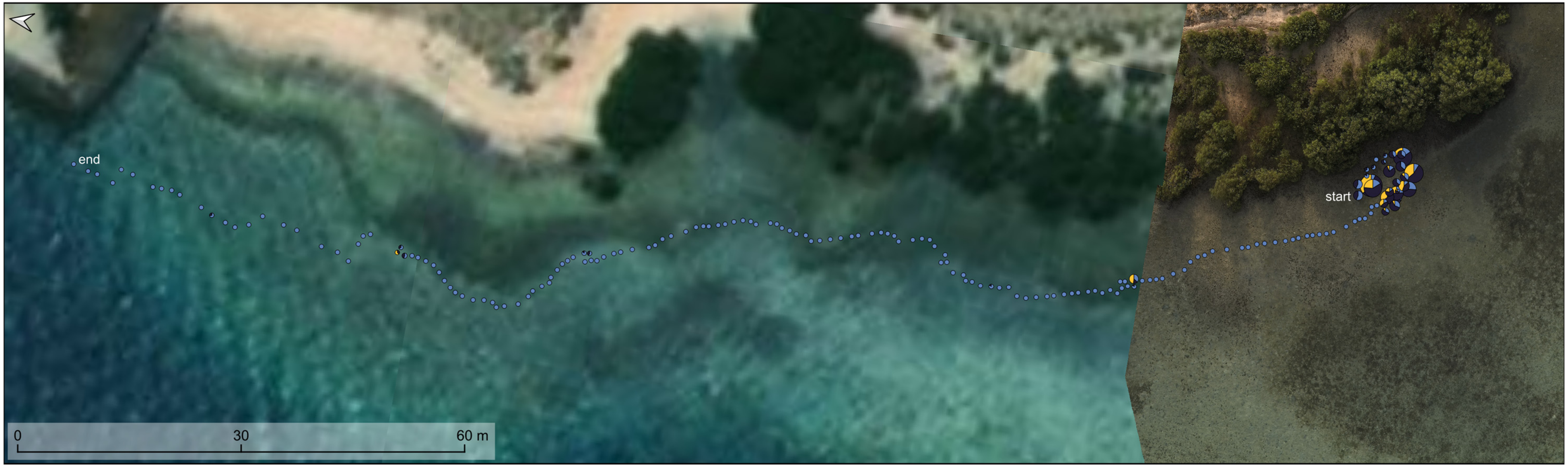


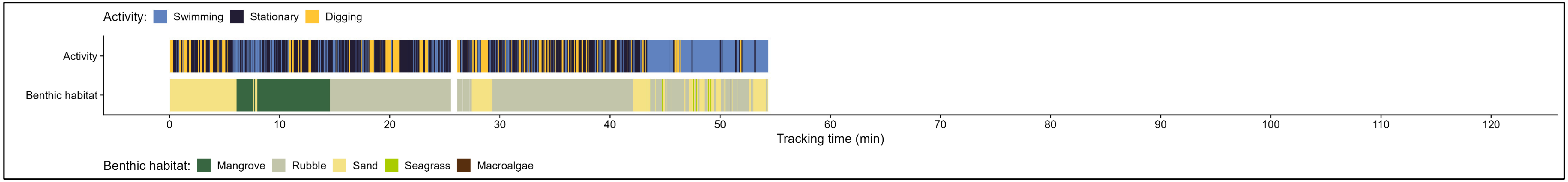

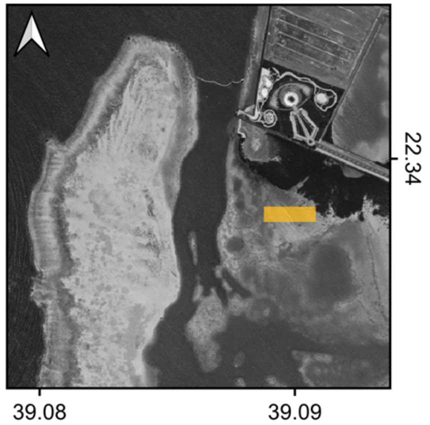


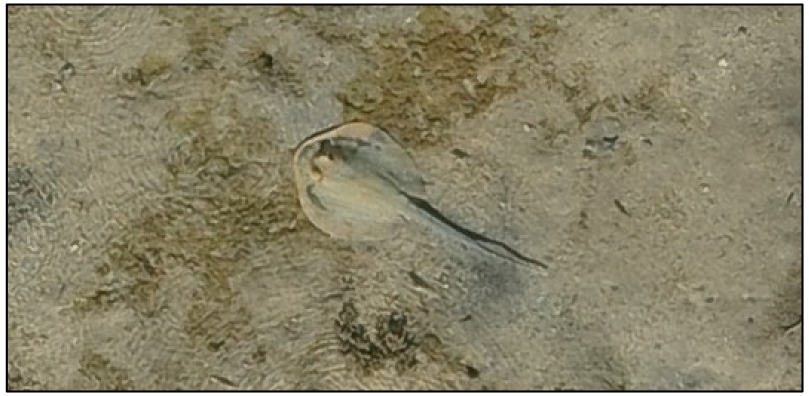

| | |
|---|---|
| Species: | *Taeniura lymma* |
| Date: | 2024-07-03 |
| Disc width: | 25 cm |
| Disc length: | 27 cm |
| Total length: | 61 cm |
| Notes: | |

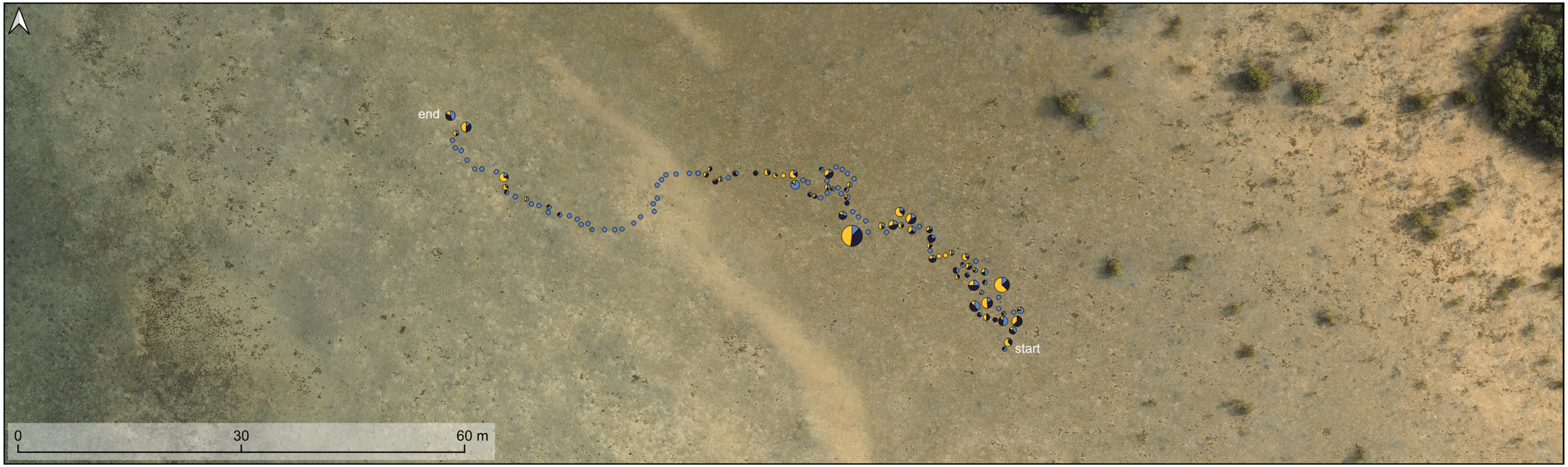


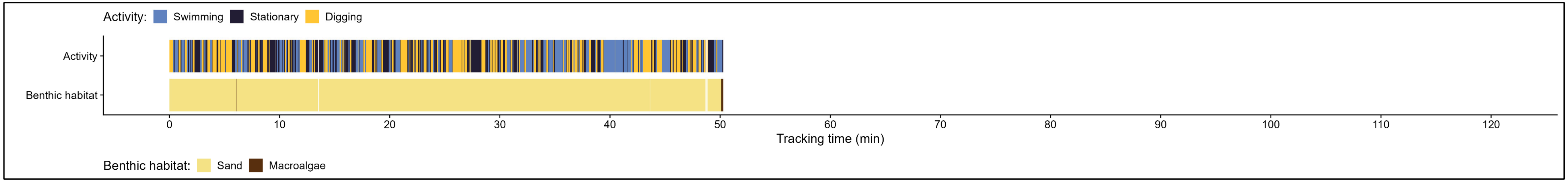

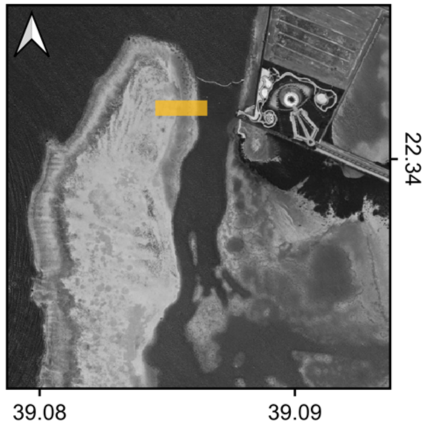


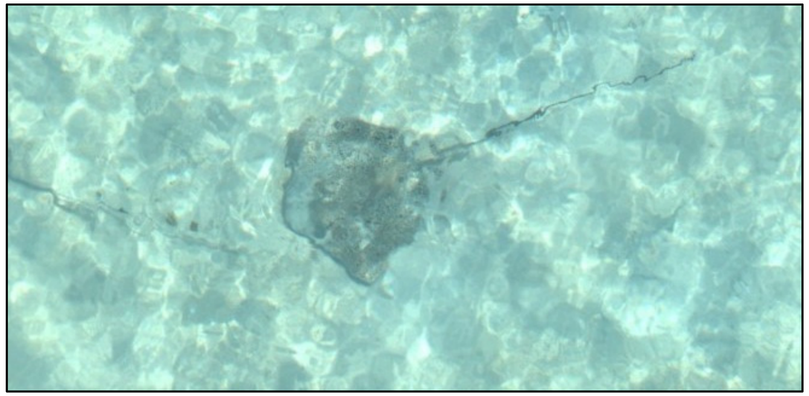

| | |
|---|---|
| Species: | *Himantura uarnak* |
| Date: | 2024-03-27 |
| Disc width: | 73 cm |
| Disc length: | 61 cm |
| Total length: | 184 cm |
| Notes: | |

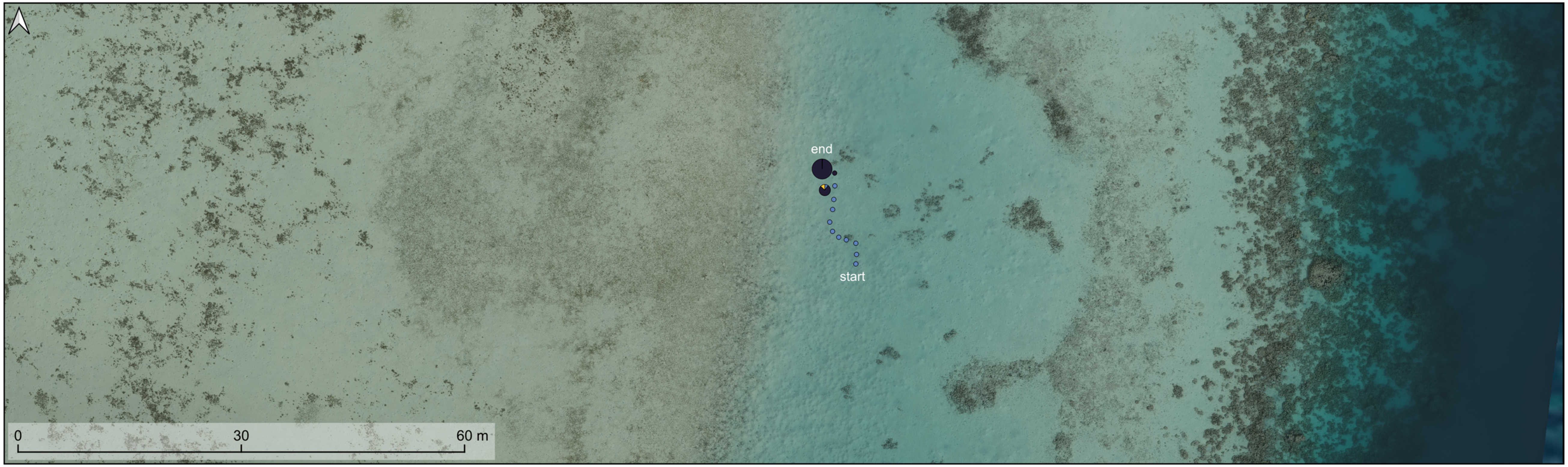


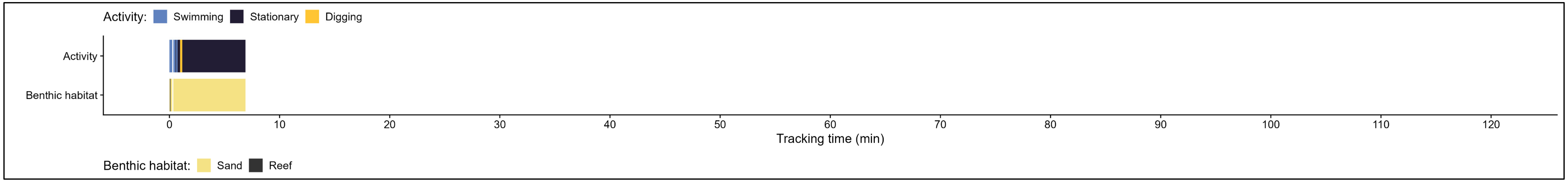

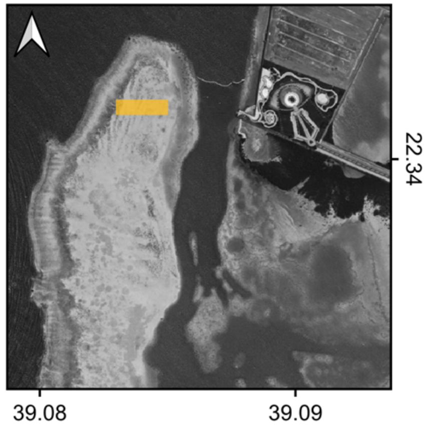


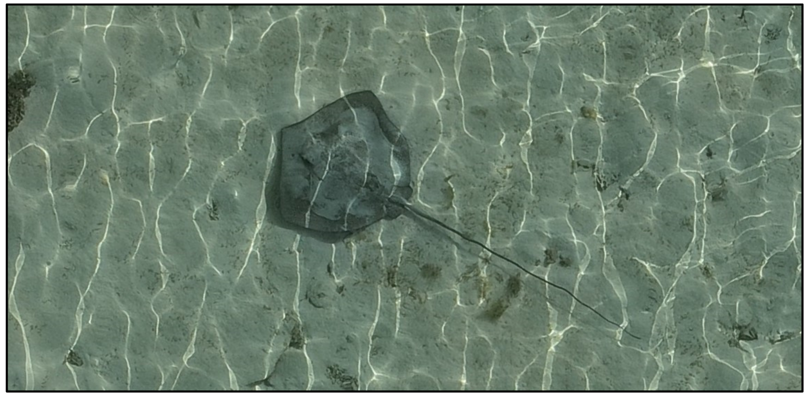

| | |
|---|---|
| Species: | *Himantura uarnak* |
| Date: | 2024-04-28 |
| Disc width: | 86 cm |
| Disc length: | 73 cm |
| Total length: | 232 cm |
| Notes: | |

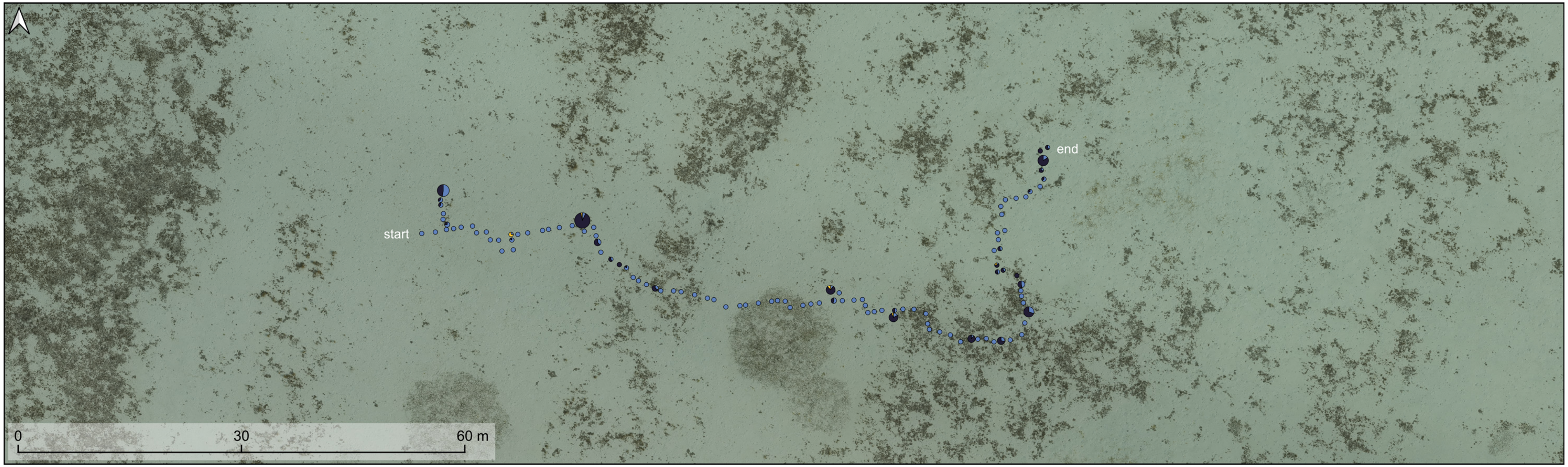


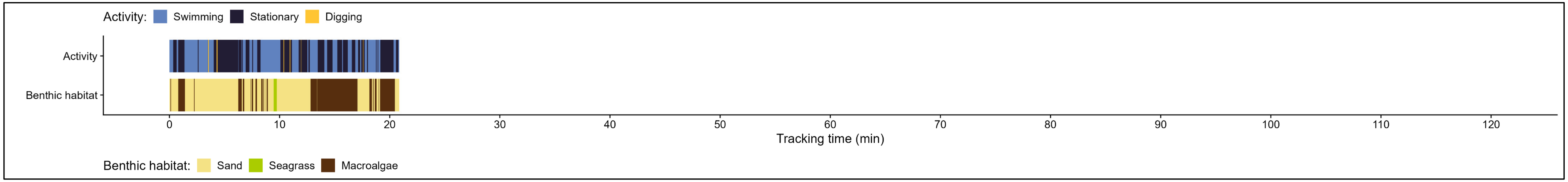

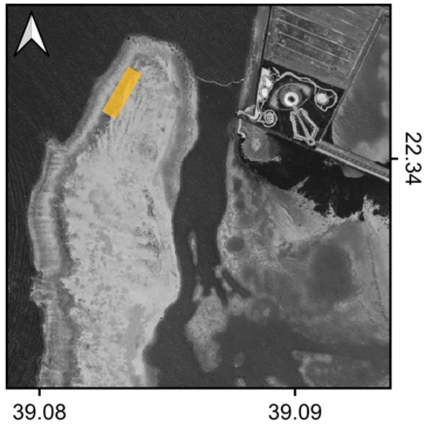


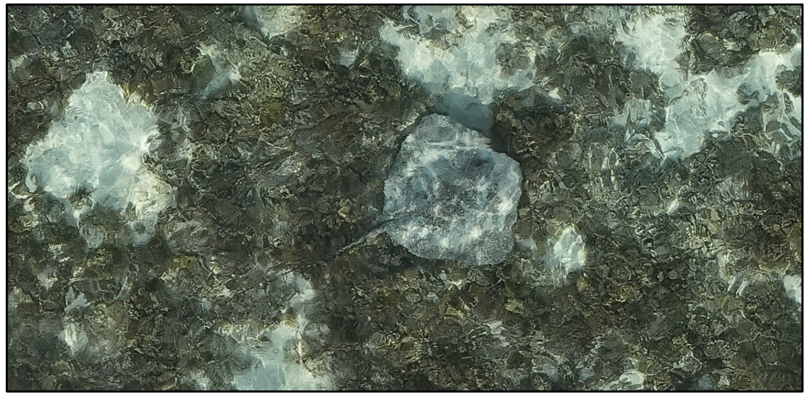

| | |
|---|---|
| Species: | *Himantura uarnak* |
| Date: | 2024-05-01 |
| Disc width: | 96 cm |
| Disc length: | 81 cm |
| Total length: | 193 cm |
| Notes: | |

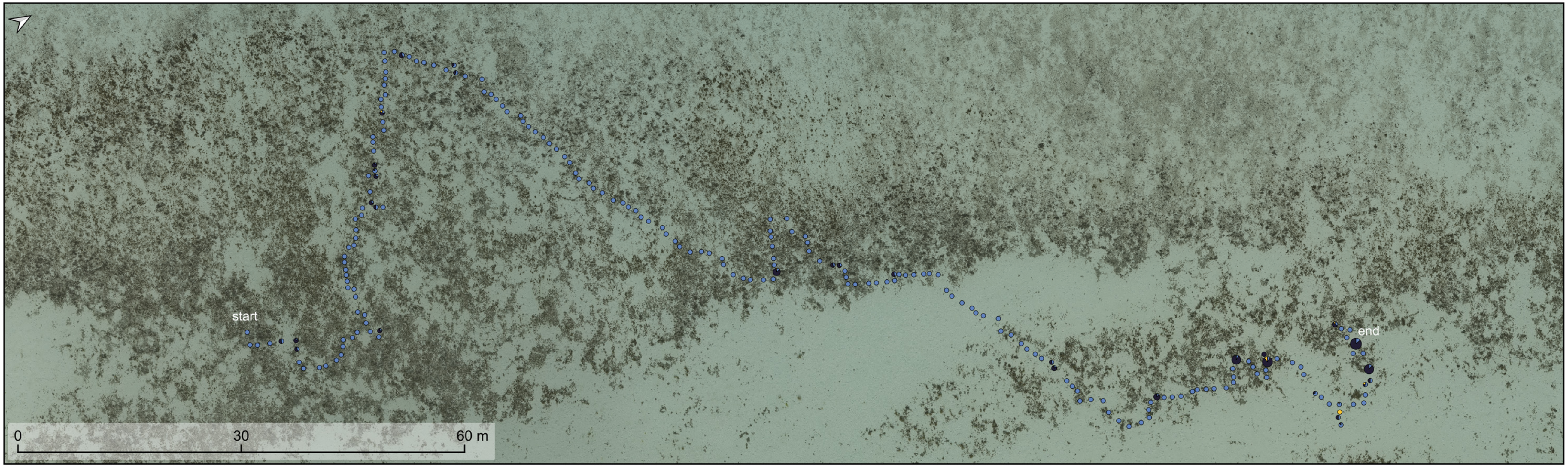


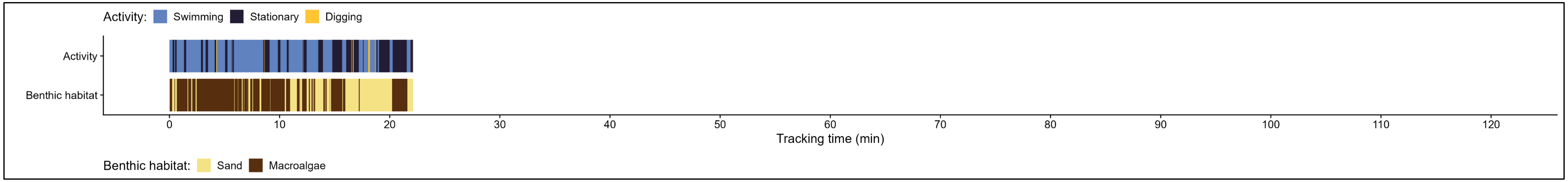

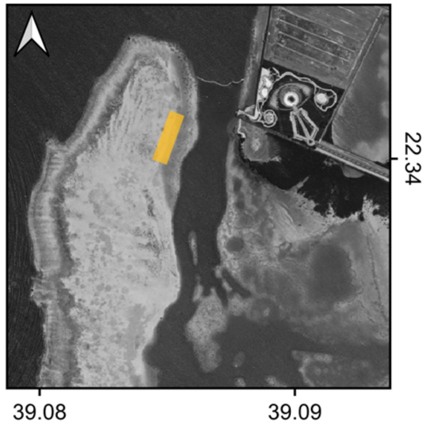


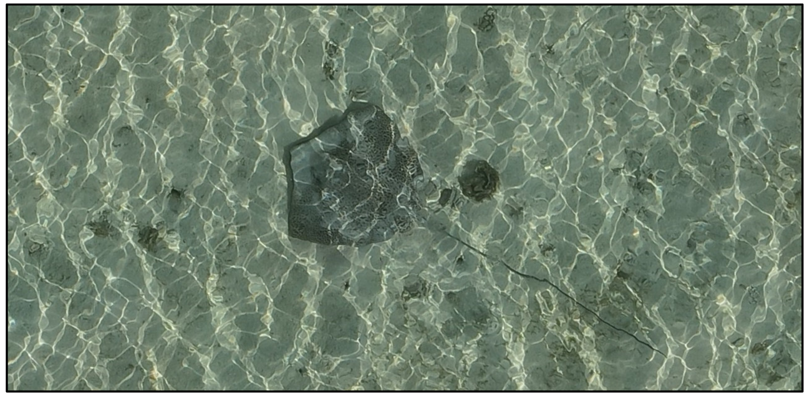

| | |
|---|---|
| Species: | *Himantura uarnak* |
| Date: | 2024-05-09 (a) |
| Disc width: | 106 cm |
| Disc length: | 92 cm |
| Total length: | 292 cm |
| Notes: | |

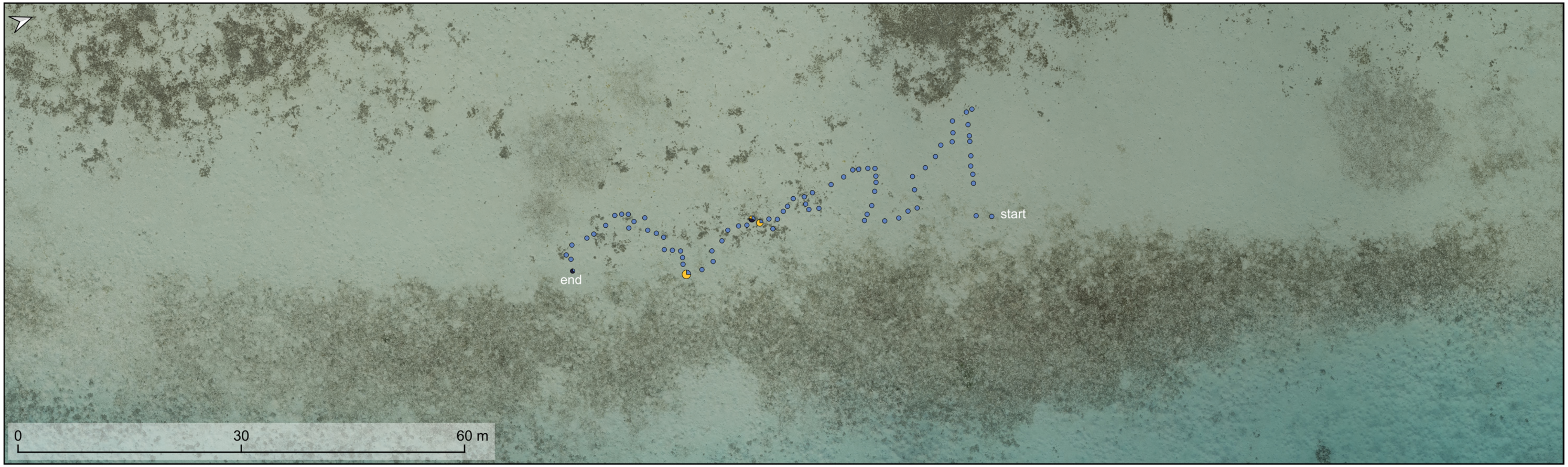


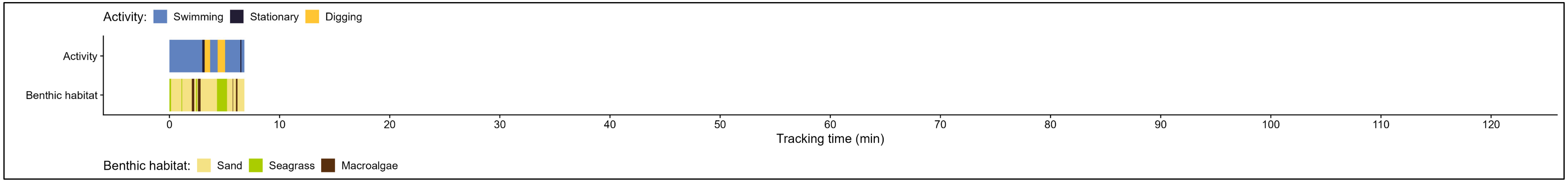

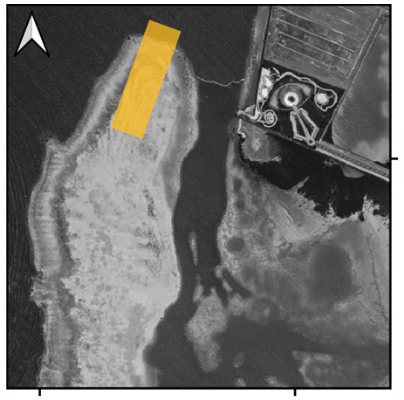


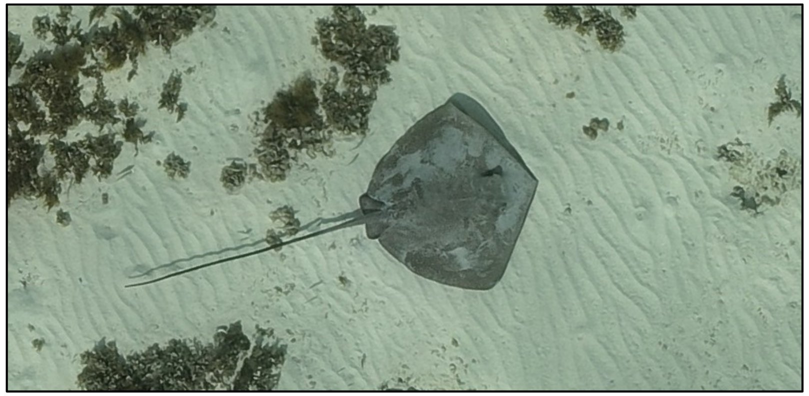

| | |
|---|---|
| Species: | *Himantura uarnak* |
| Date: | 2024-05-09 (b) |
| Disc width: | 92 cm |
| Disc length: | 75 cm |
| Total length: | 203 cm |
| Notes: | Numbers show travel direction. |

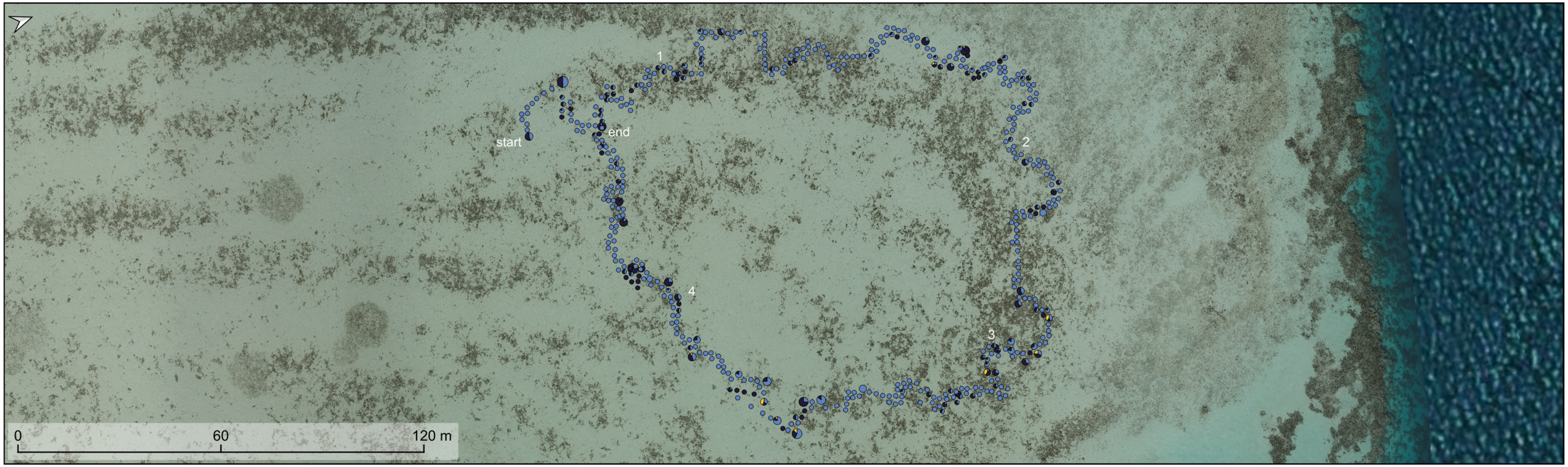


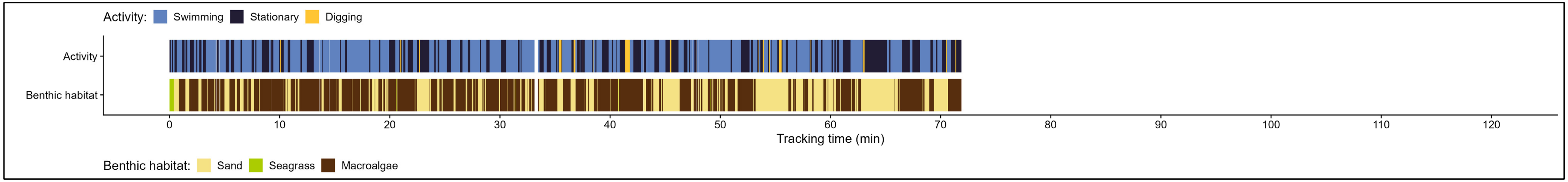

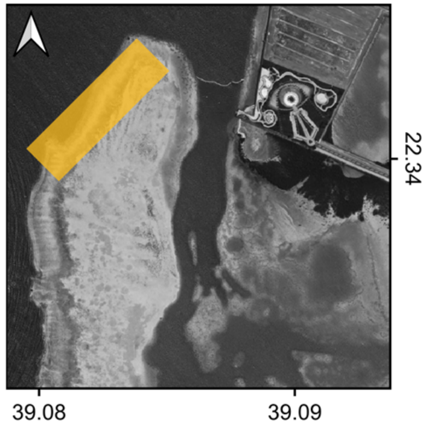


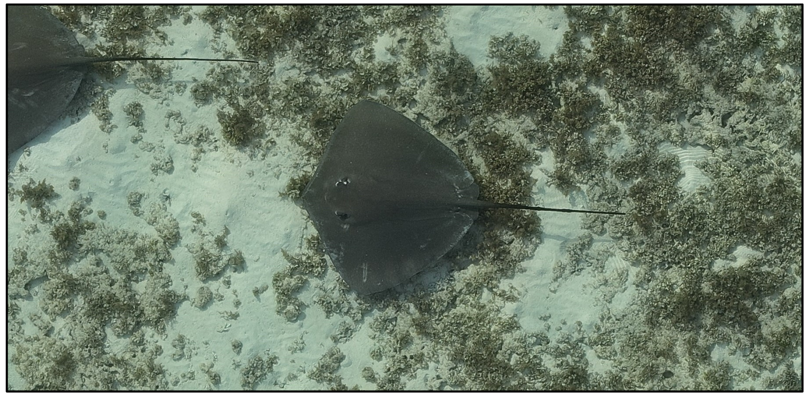

Species: *cf. Pateobatis fai*

Date: 2024-05-10

Disc width: 122 cm

Disc length: 107 cm

Total length: 213 cm

Notes: Followed by a second individual and two trevally (Video E3). Likely also seen as duo along the transects on 05-22 (with trevally) and 06-03 (white dots). On 06-11 a similar looking individual was spotted, again with a trevally alongside.

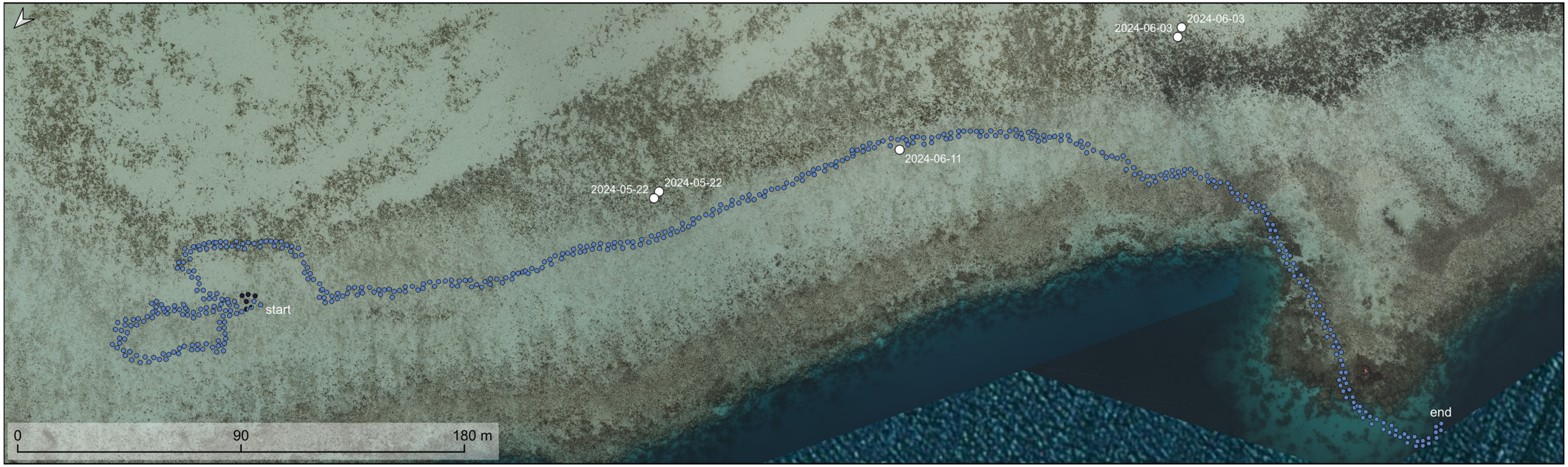


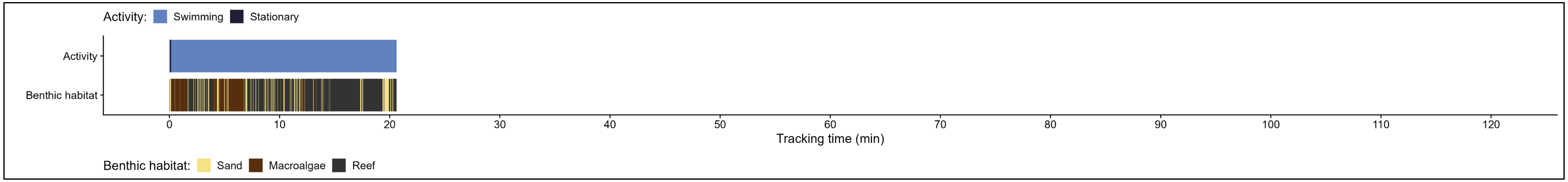

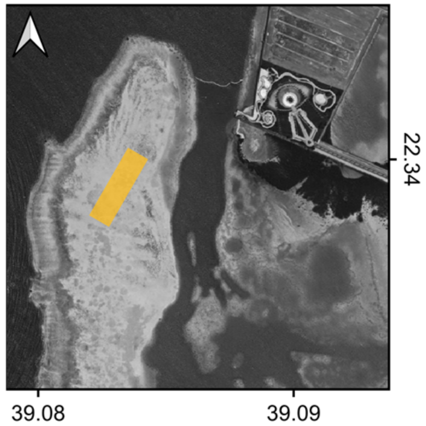


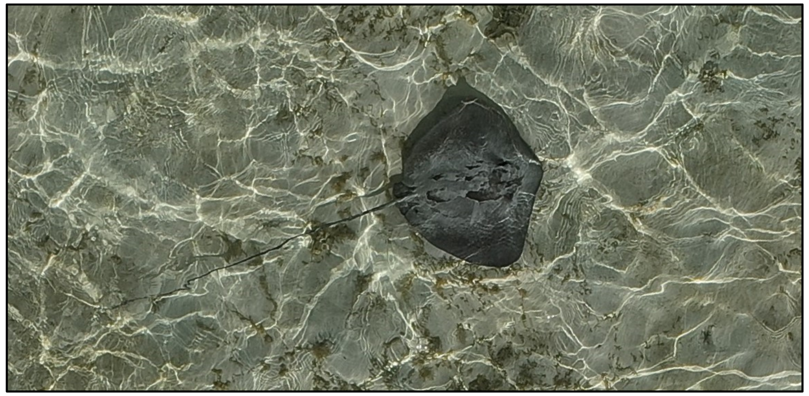

Species: cf. *Pateobatis fai*

Date: 2024-05-17

Disc width: 75 cm

Disc length: 59 cm

Total length: 203 cm

Notes:

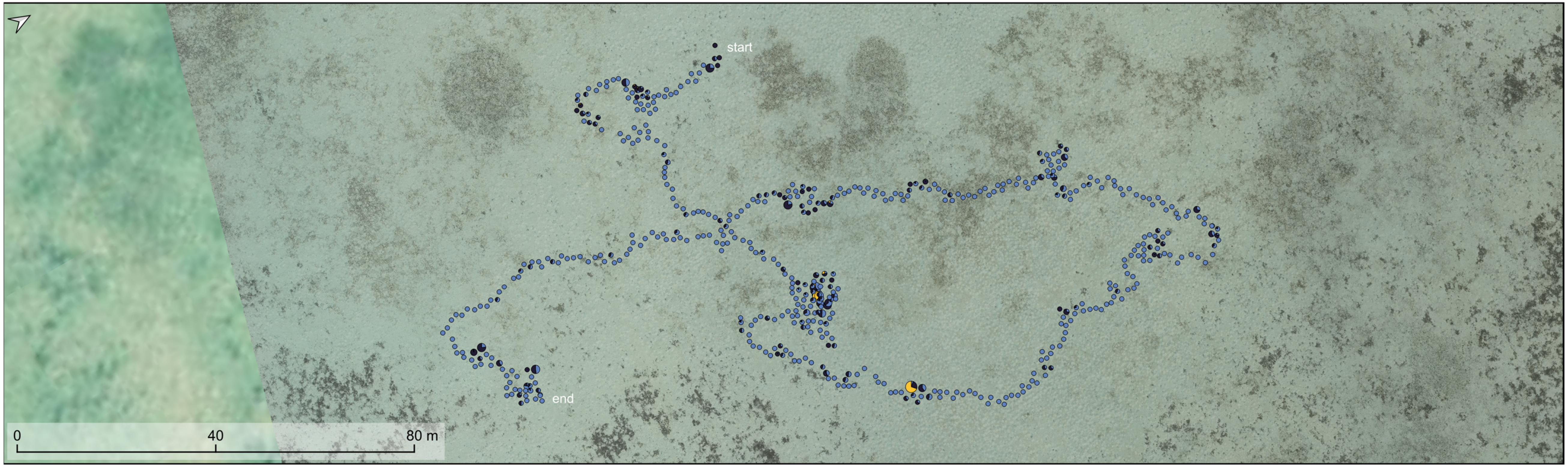


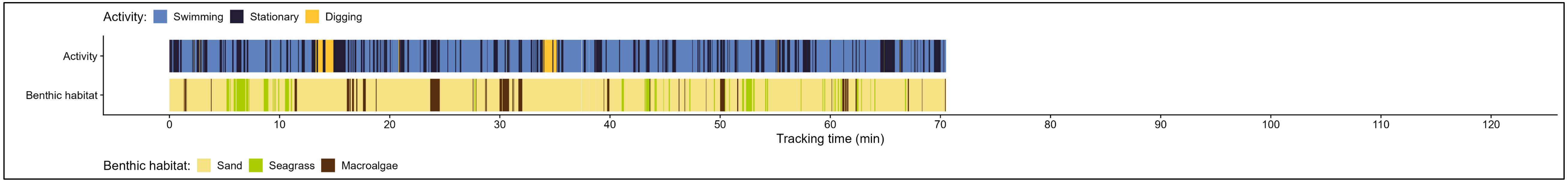

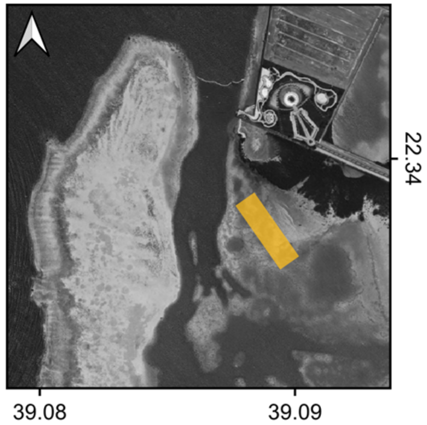


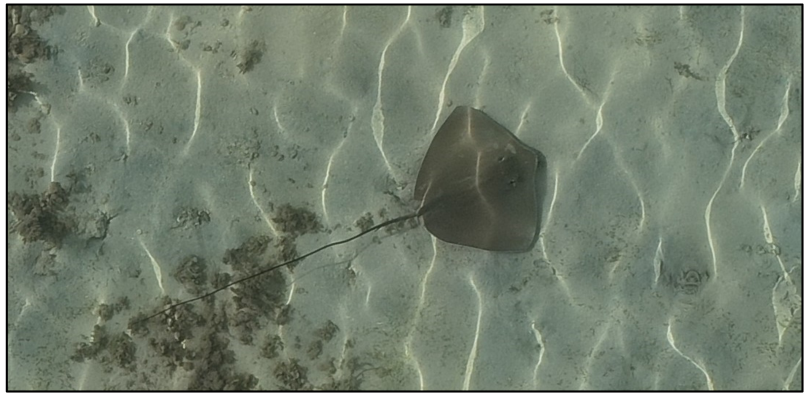

| | |
|---|---|
| Species: | cf. *Pateobatis fai* |
| Date: | 2024-05-20 |
| Disc width: | 69 cm |
| Disc length: | 54 cm |
| Total length: | 191 cm |
| Notes: | For this individual stationary reflects resting behaviour. |

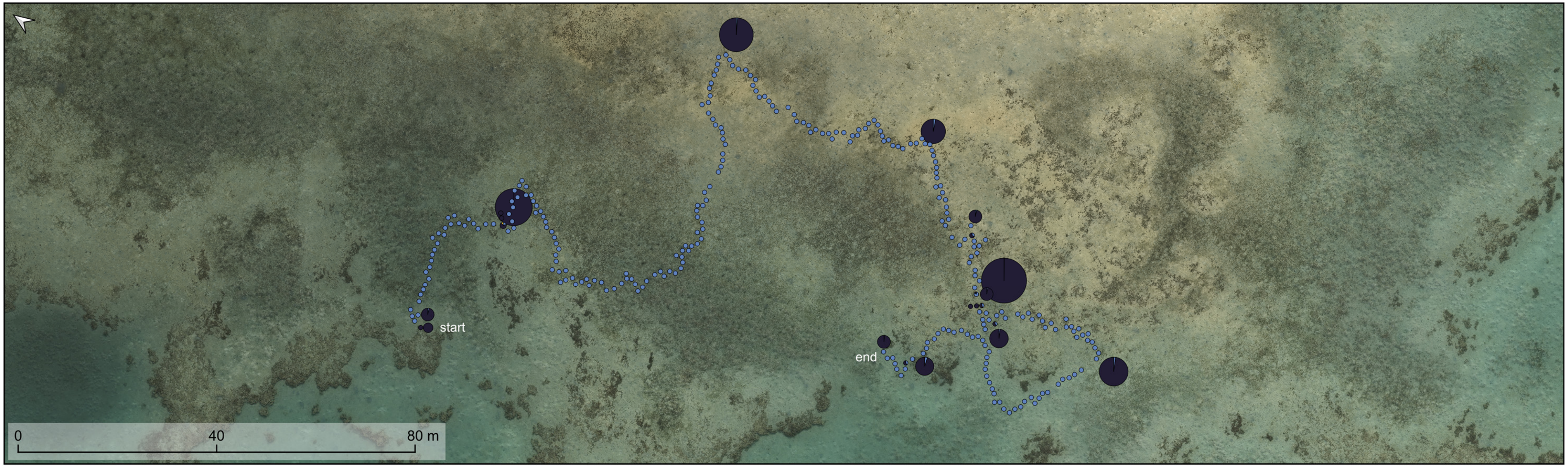


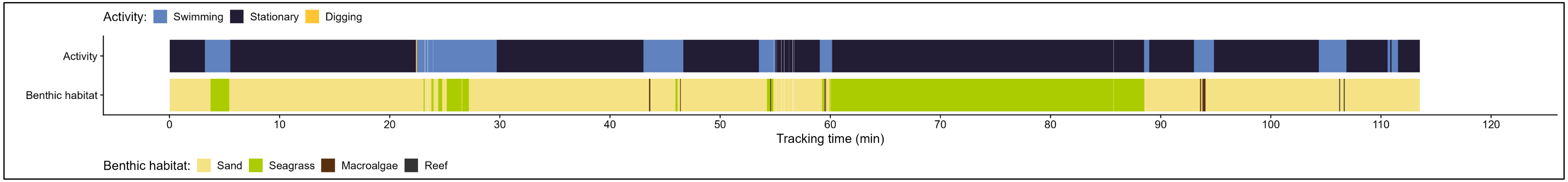

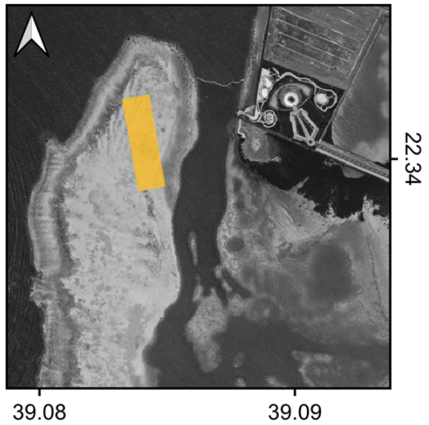


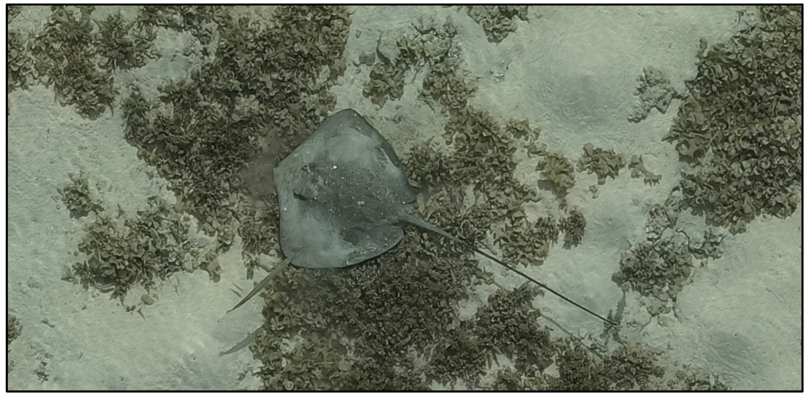

| | |
|---|---|
| Species: | *Himantura uarnak* |
| Date: | 2024-05-22 |
| Disc width: | 80 cm |
| Disc length: | 68 cm |
| Total length: | 183 cm |
| Notes: | Track was picked up again, ID confirmed through unique pattern of white spots. Numbers show travel direction. White dots show re-observations of the same individual in the transect surveys. This individual was also tracked on June 11th. |

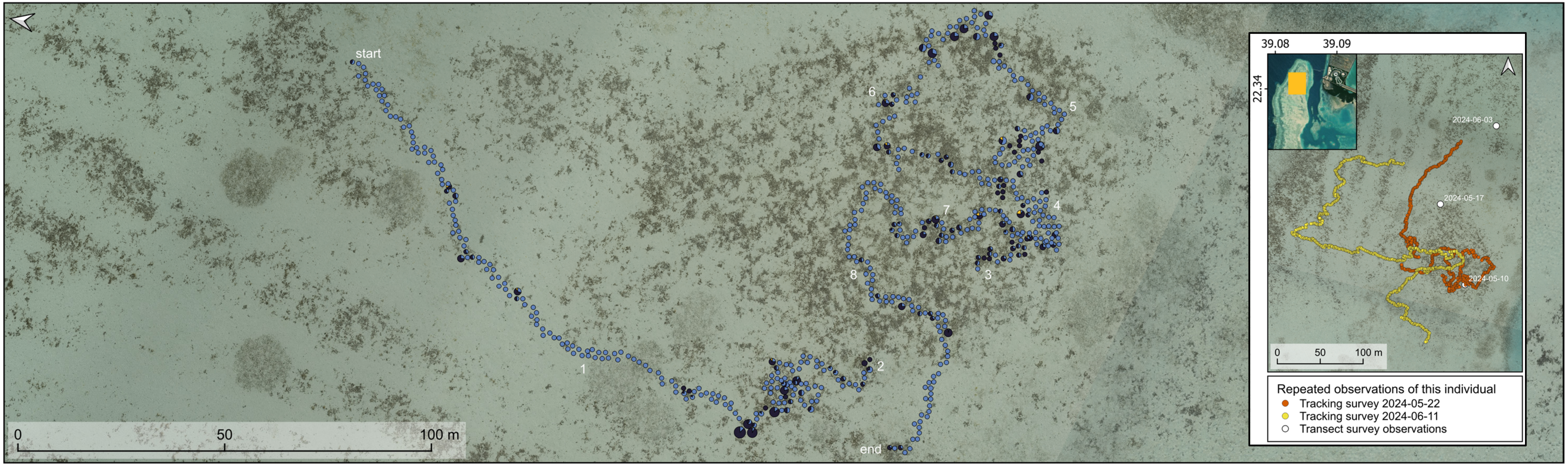


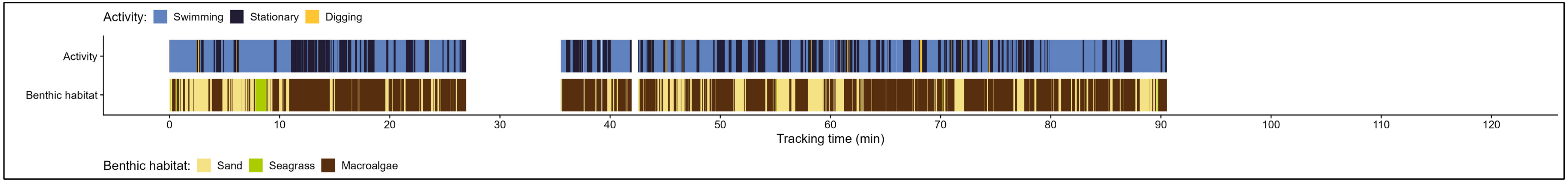

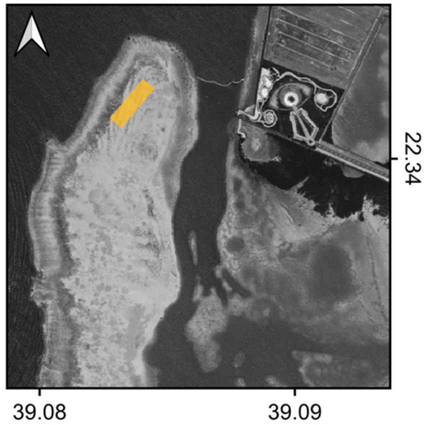


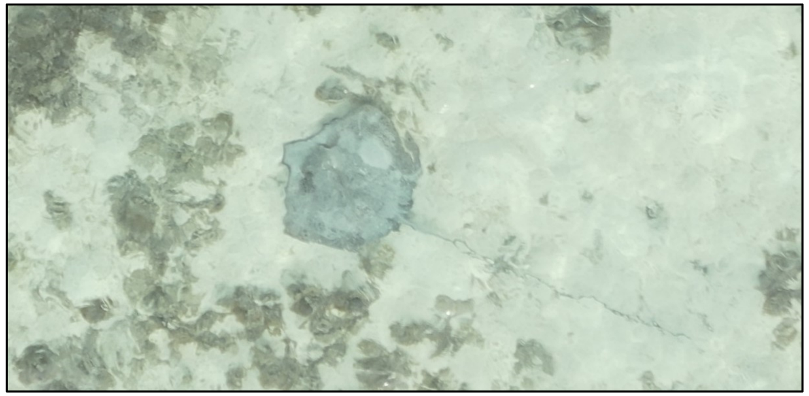

| | |
|---|---|
| Species: | Whipray |
| Date: | 2024-06-10 |
| Disc width: | 73 cm |
| Disc length: | 59 cm |
| Total length: | 209 cm |
| Notes: | Likely *Himantura uarnak* as it seems to have spots. However, this is difficult to see due to waves. |

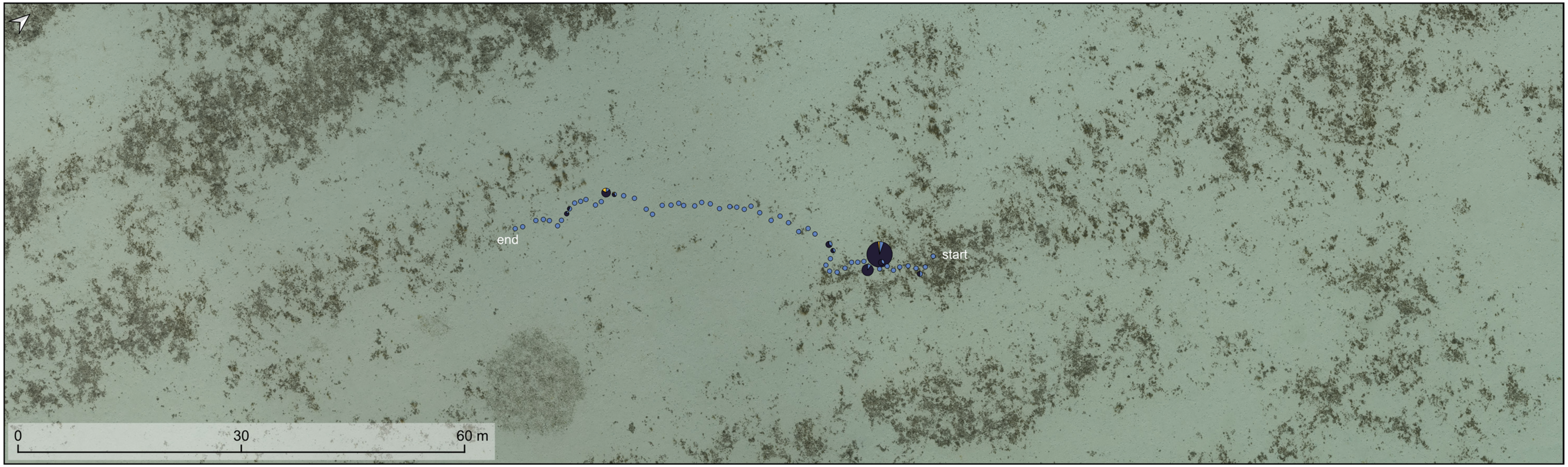


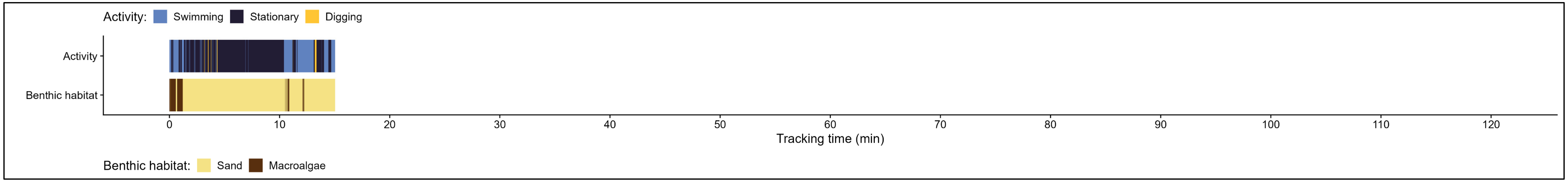

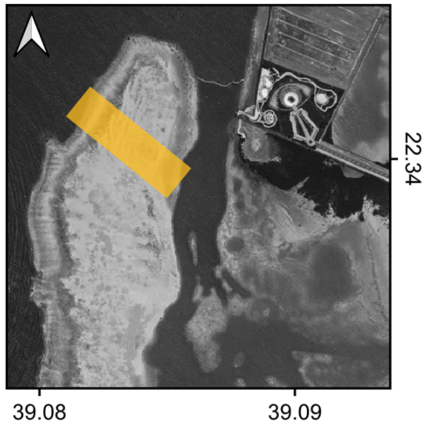


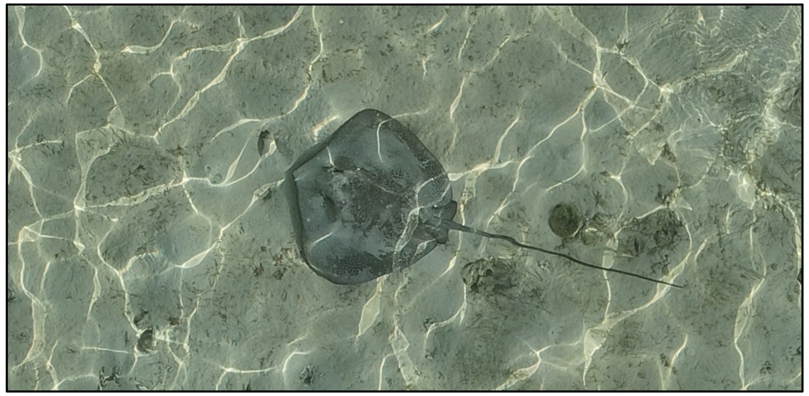

| | |
|---|---|
| Species: | *Himantura uarnak* |
| Date: | 2024-06-11 |
| Disc width: | 80 cm |
| Disc length: | 70 cm |
| Total length: | 185 cm |
| Notes: | The same individual has been tracked on May 22nd and was observed in the transect surveys on May 10th, May 17th, and June 3rd. |

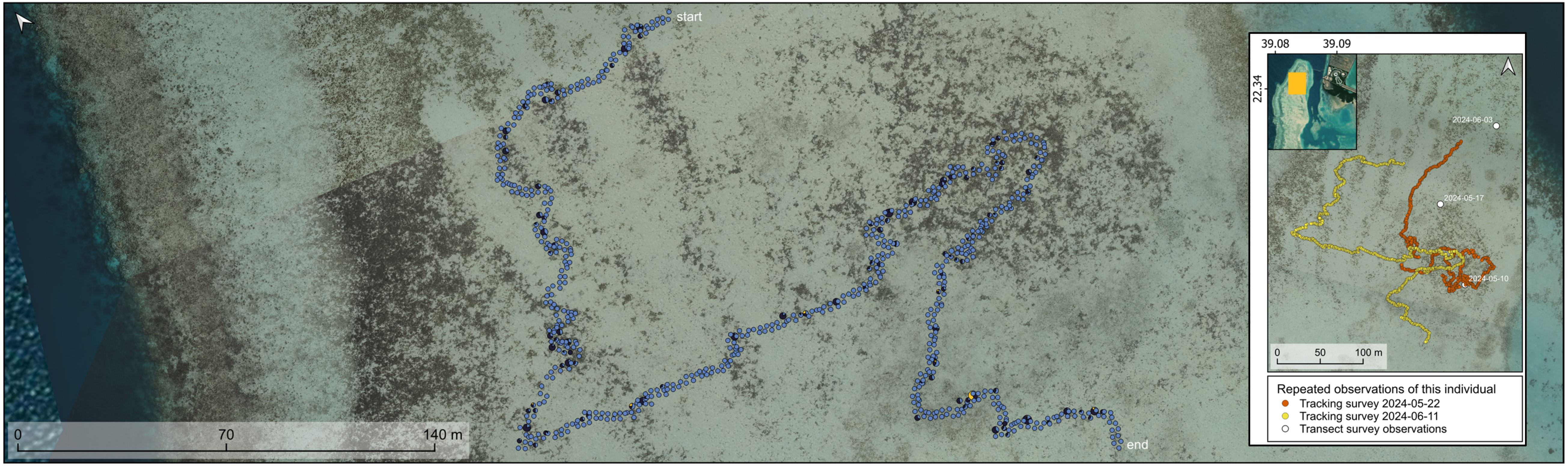


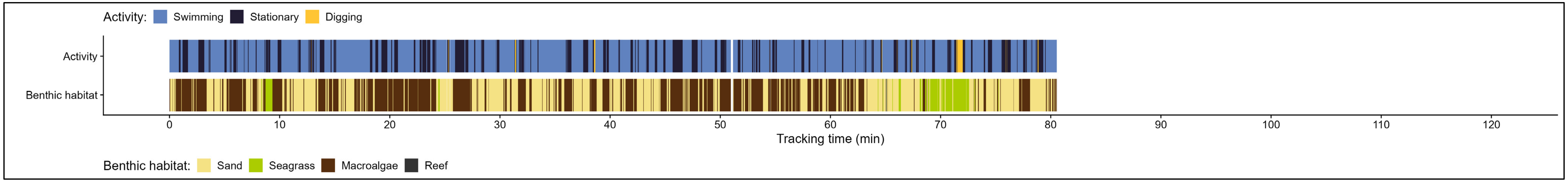

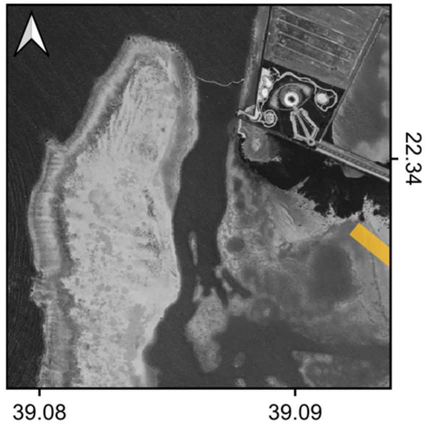


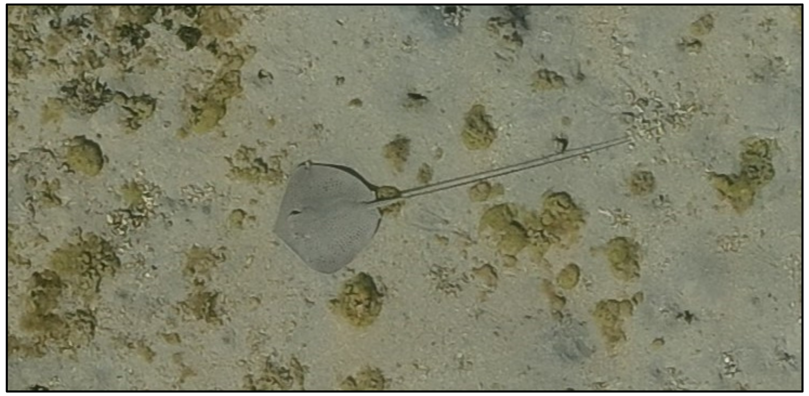

Species: *Himantura uarnak*

Date: 2024-06-17

Disc width: 32 cm

Disc length: 29 cm

Total length: 67 cm

Notes:

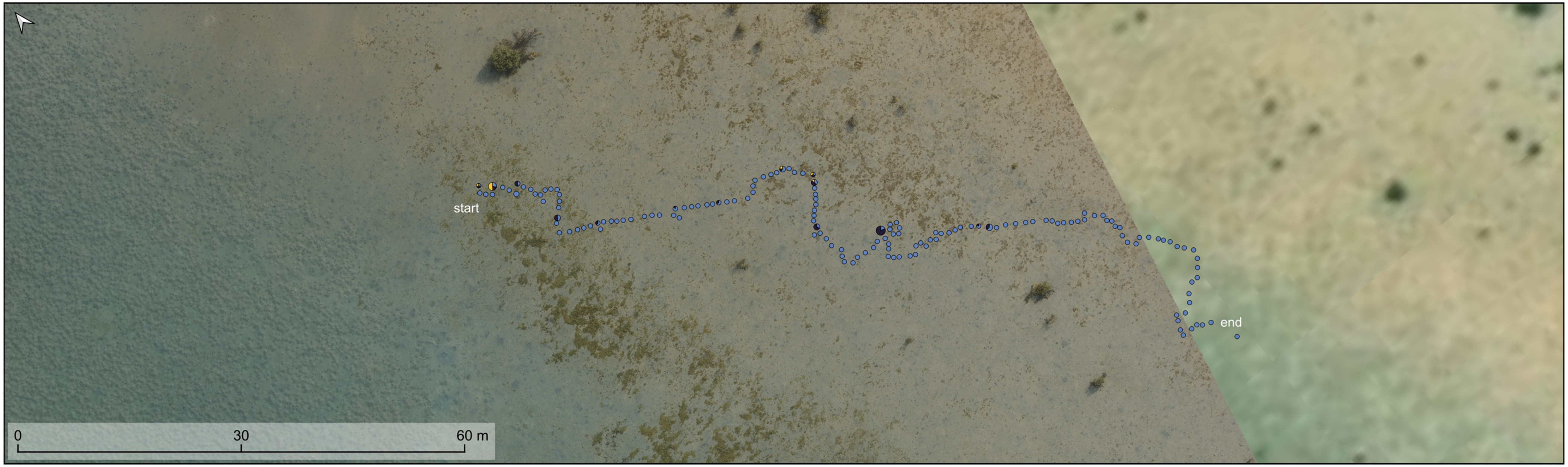


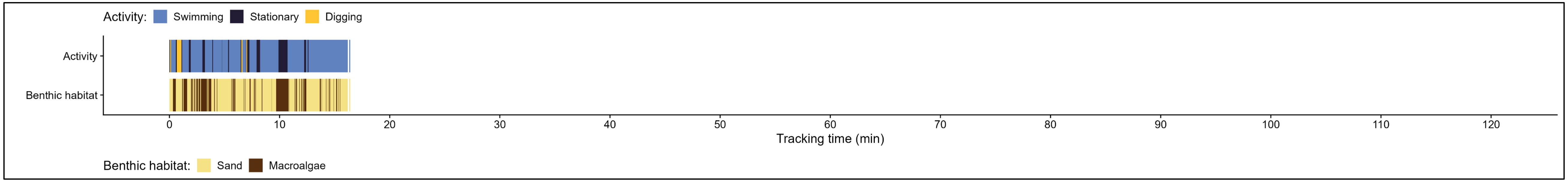

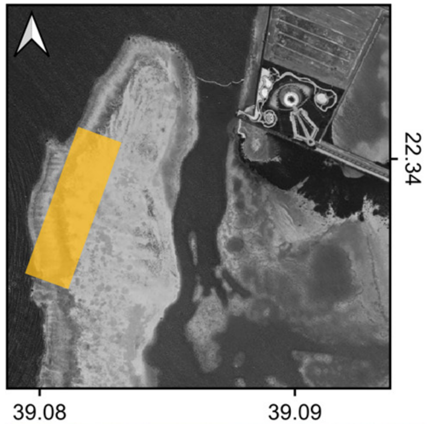


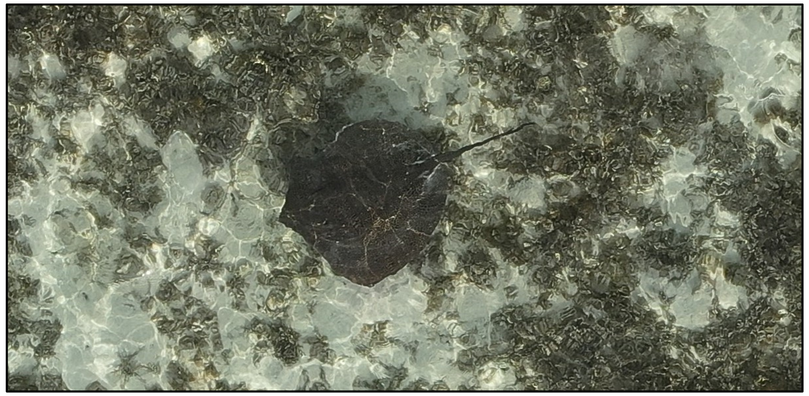

| | |
|---|---|
| Species: | *Himantura uarnak* |
| Date: | 2024-06-23 |
| Disc width: | 80 cm (if not wounded, estimated based on symmetry) |
| Disc length: | 74 cm |
| Total length: | 125 cm |
| Notes: | Right side of the pectoral disc is partly missing, potentially from a recent predation event. |

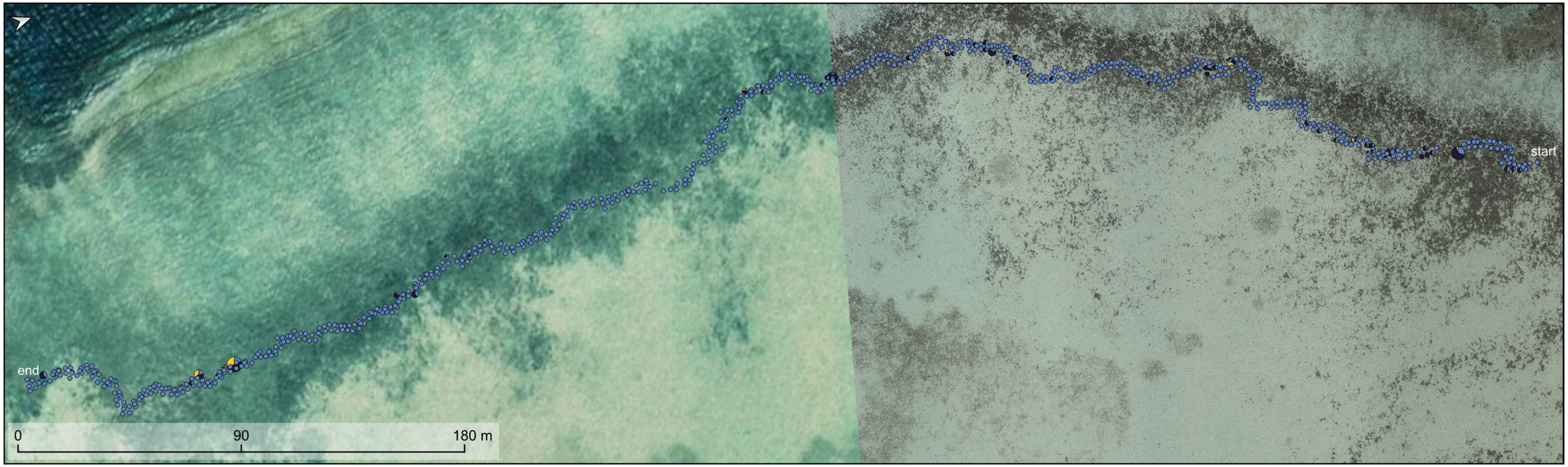


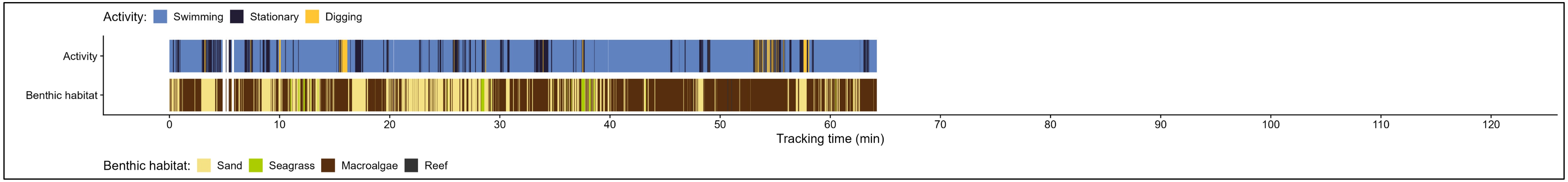

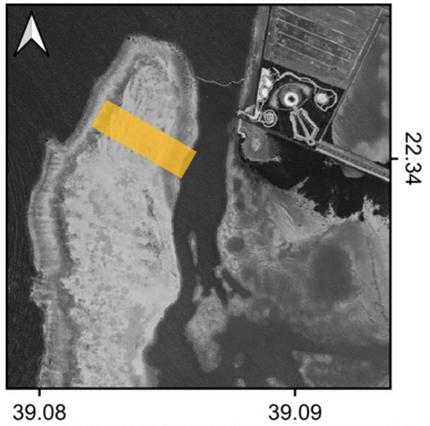


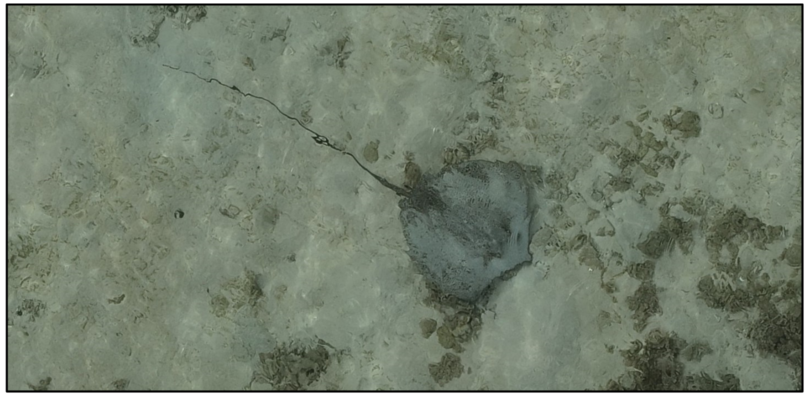

Species: *Himantura uarnak*

Date: 2024-06-24

Disc width: 82 cm

Disc length: 70 cm

Total length: 223 cm

Notes:

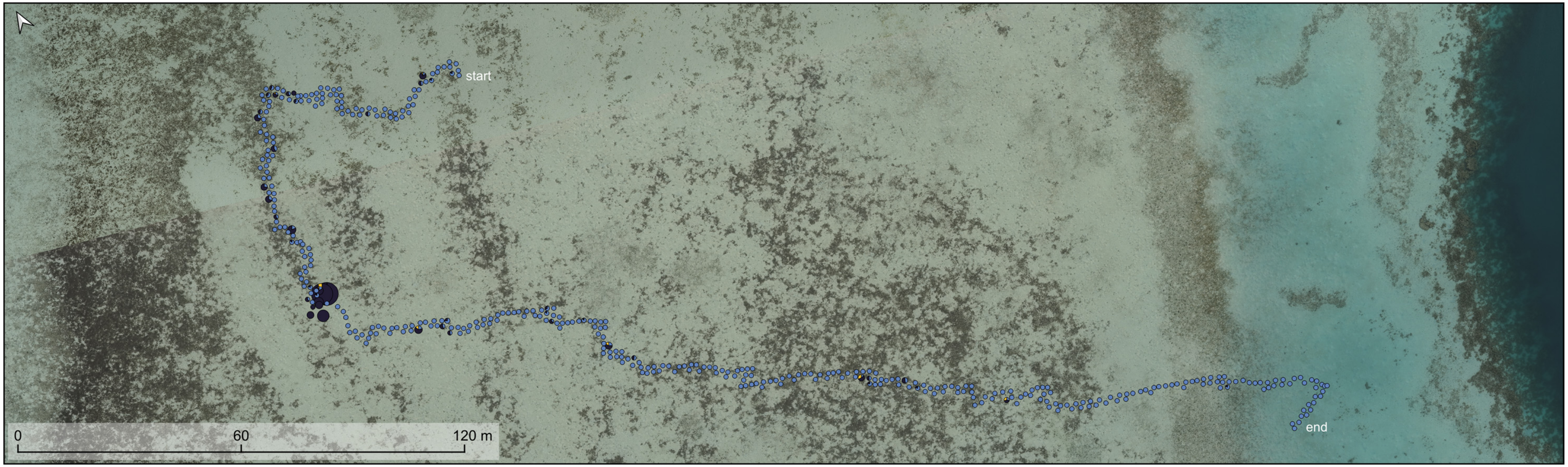


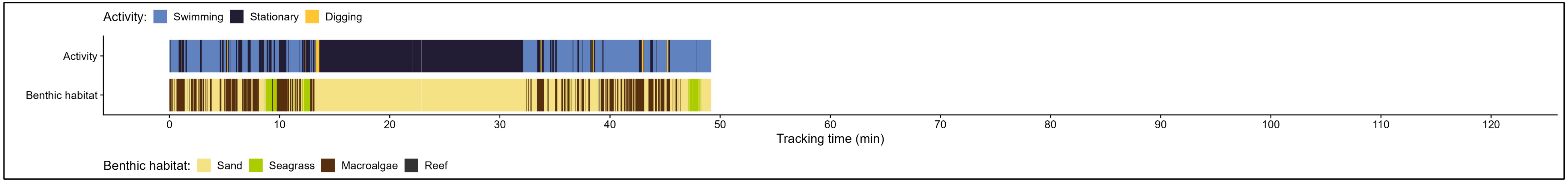

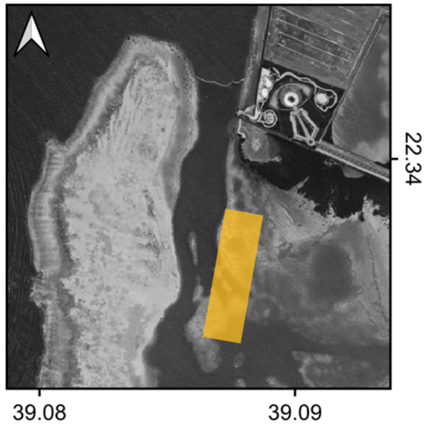


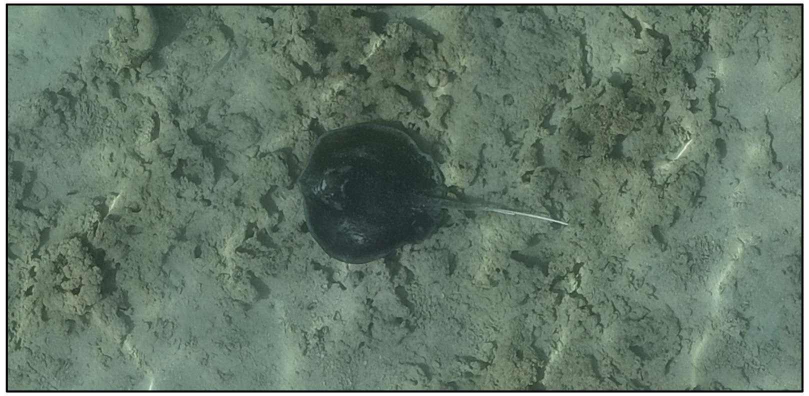

Species: *Urogymnus granulatus*

Date: 2024-05-02

Disc width: 80 cm

Disc length: 78 cm

Total length: 155 cm

Notes:

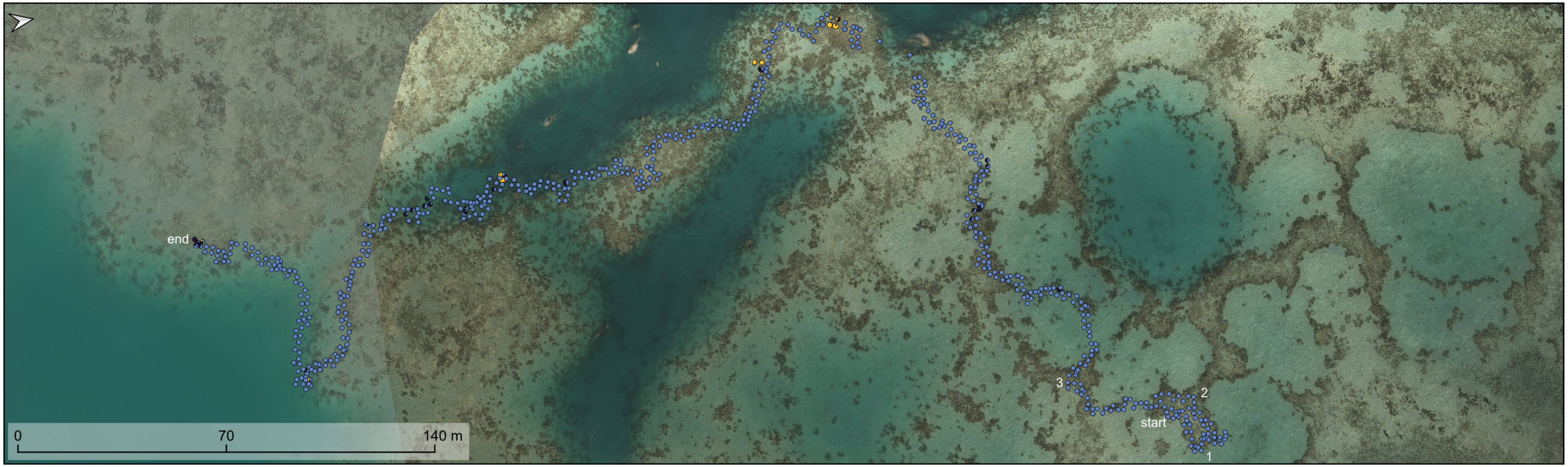


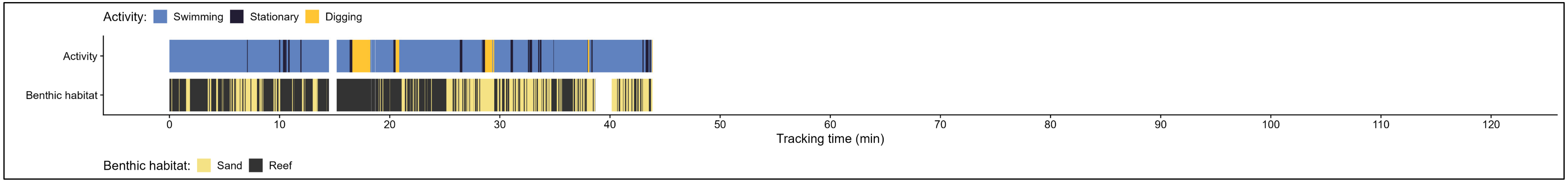

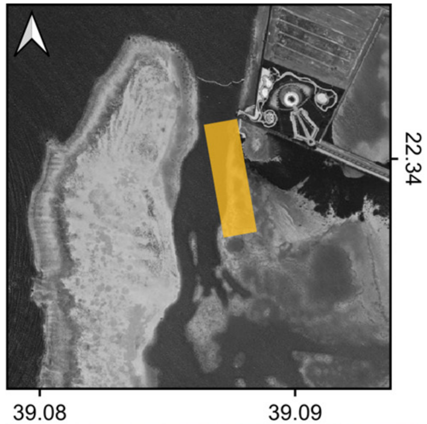


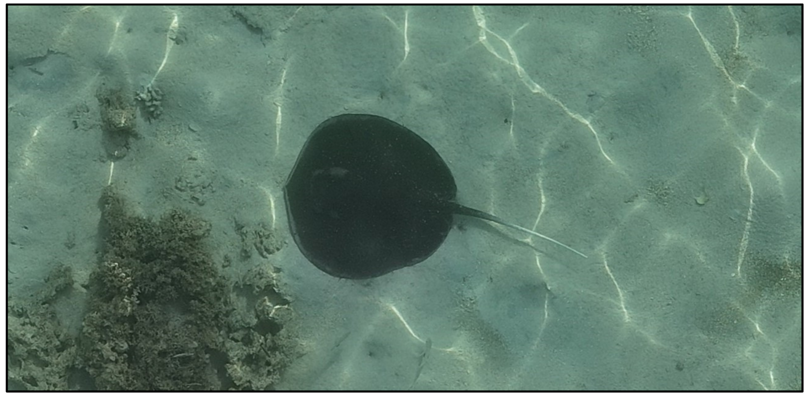

| | |
|---|---|
| Species: | *Urogymnus granulatus* |
| Date: | 2024-06-06 |
| Disc width: | 81 cm |
| Disc length: | 79 cm |
| Total length: | 152 cm |
| Notes: | |

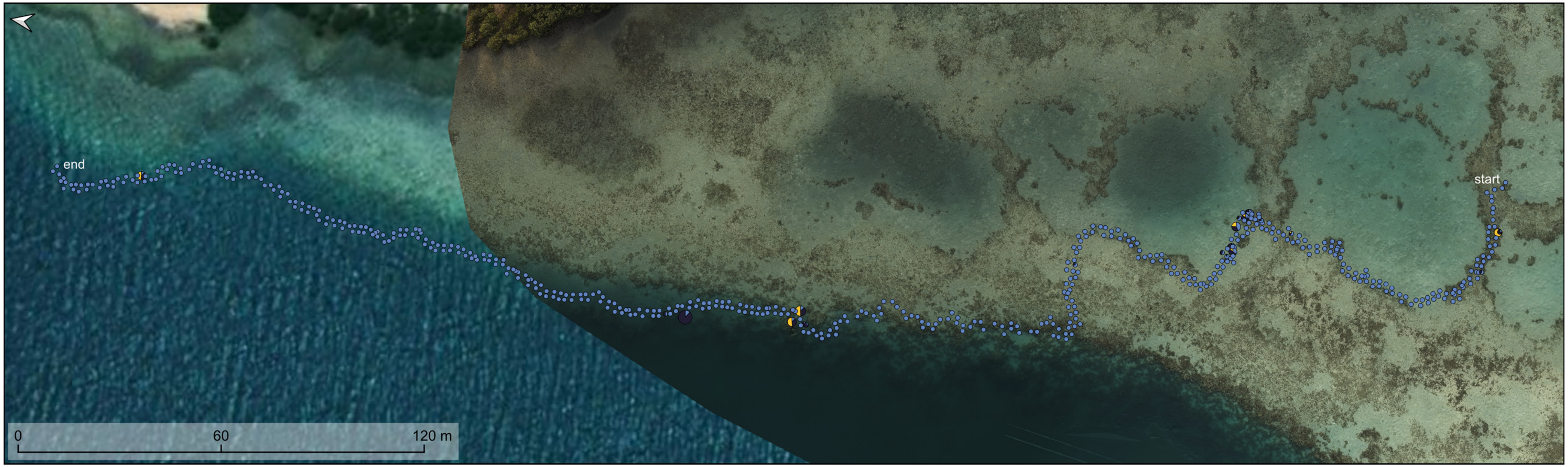


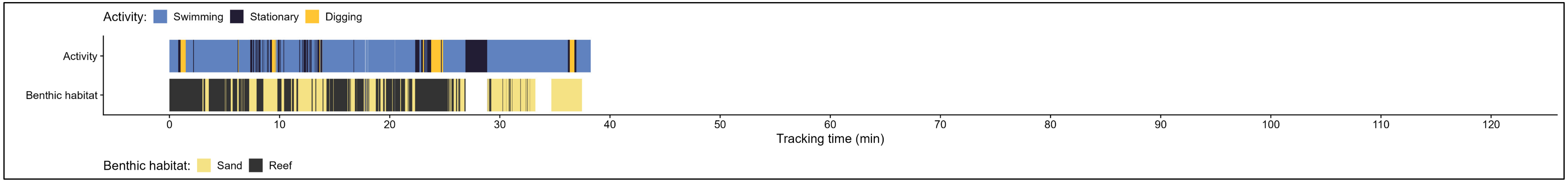

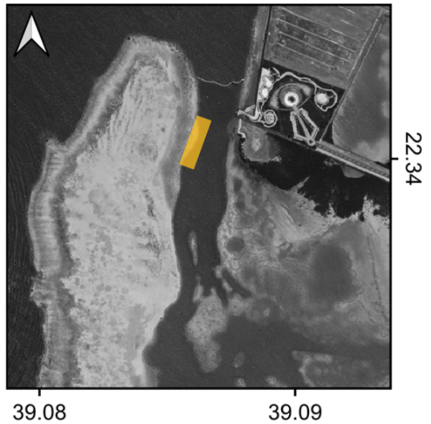


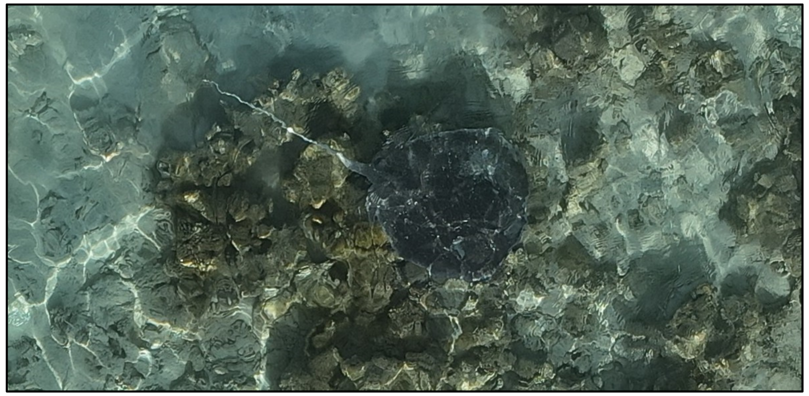

| | |
|---|---|
| Species: | *Urogymnus granulatus* |
| Date: | 2024-07-02 |
| Disc width: | 71 cm |
| Disc length: | 74 cm |
| Total length: | 160 cm |
| Notes: | |

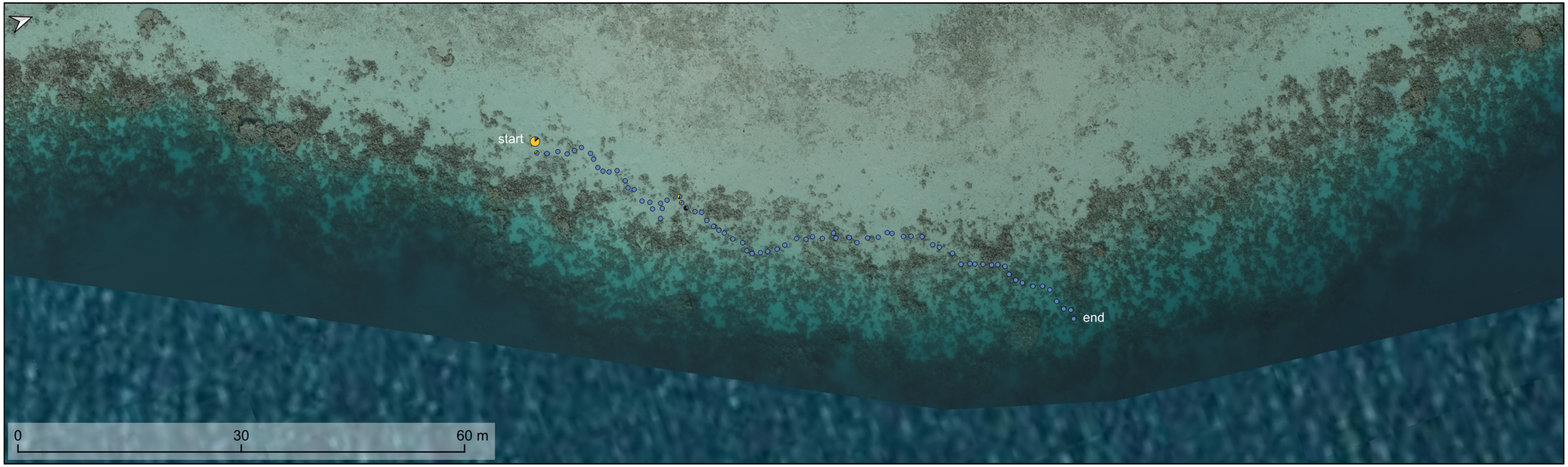


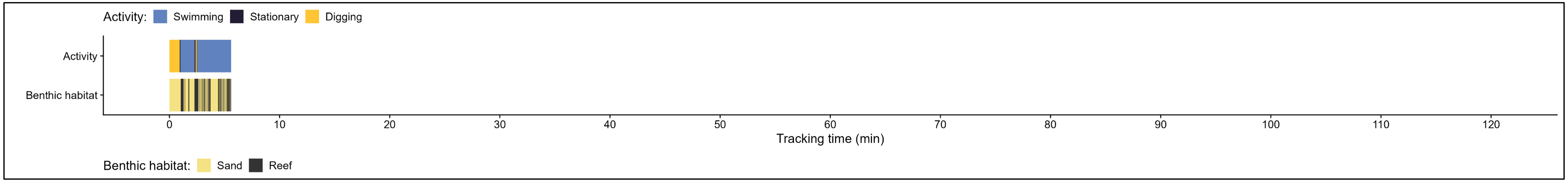

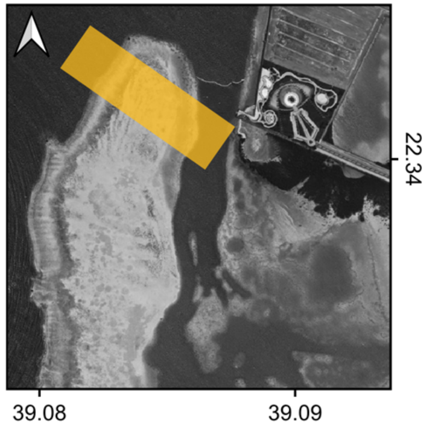


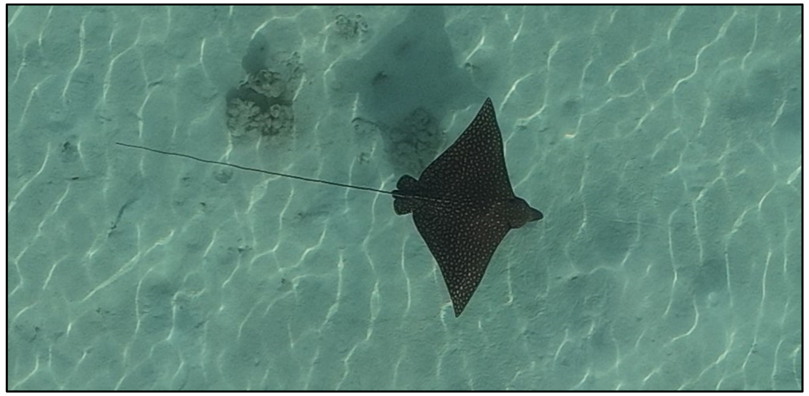

| | |
|---|---|
| Species: | *Aetobatus ocellatus* |
| Date: | 2024-05-09 |
| Disc width: | 86 cm |
| Disc length: | 55 cm |
| Total length: | 166 cm |
| Notes: | |

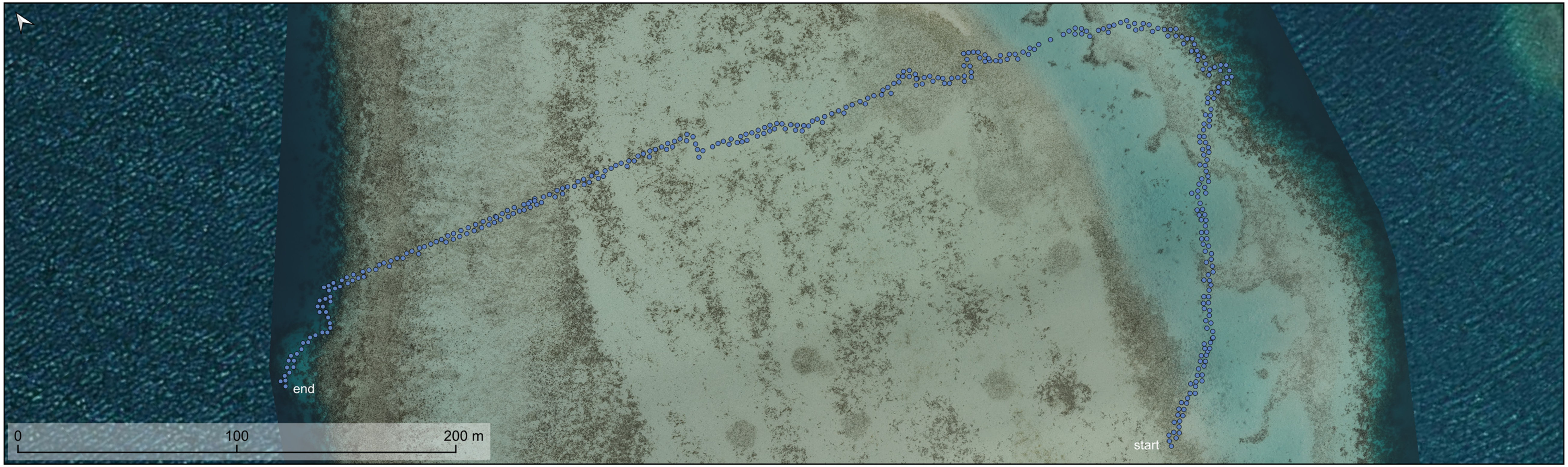


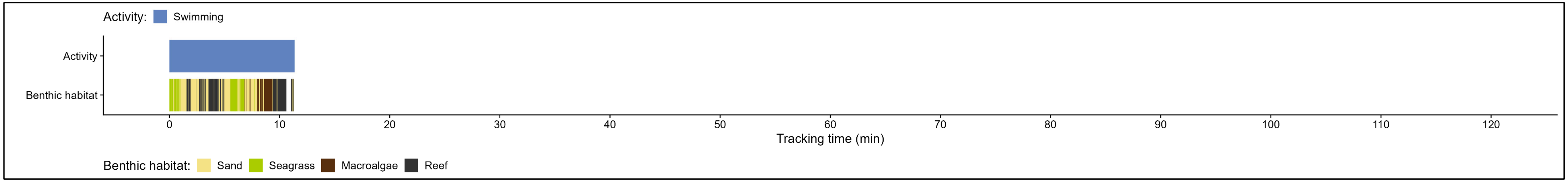

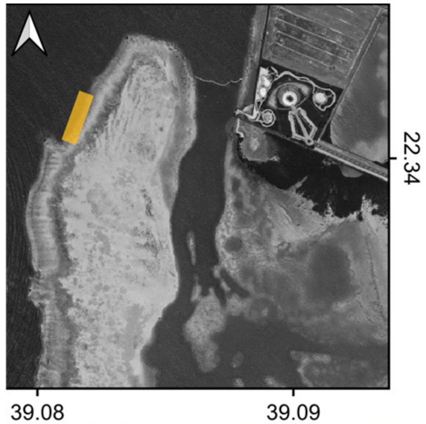


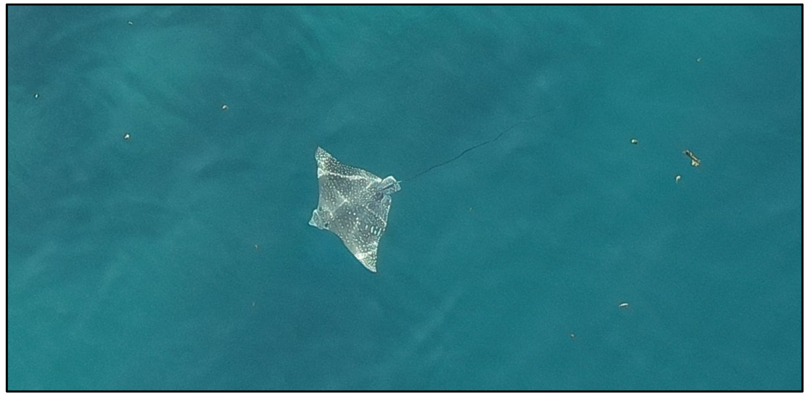

| | |
|---|---|
| Species: | *Aetobatus ocellatus* |
| Date: | 2024-06-17 |
| Disc width: | 132 cm |
| Disc length: | 80 cm |
| Total length: | 262 cm |
| Notes: | Quickly lost in deep water, retained as separate track to maximise training data for the AI model. |

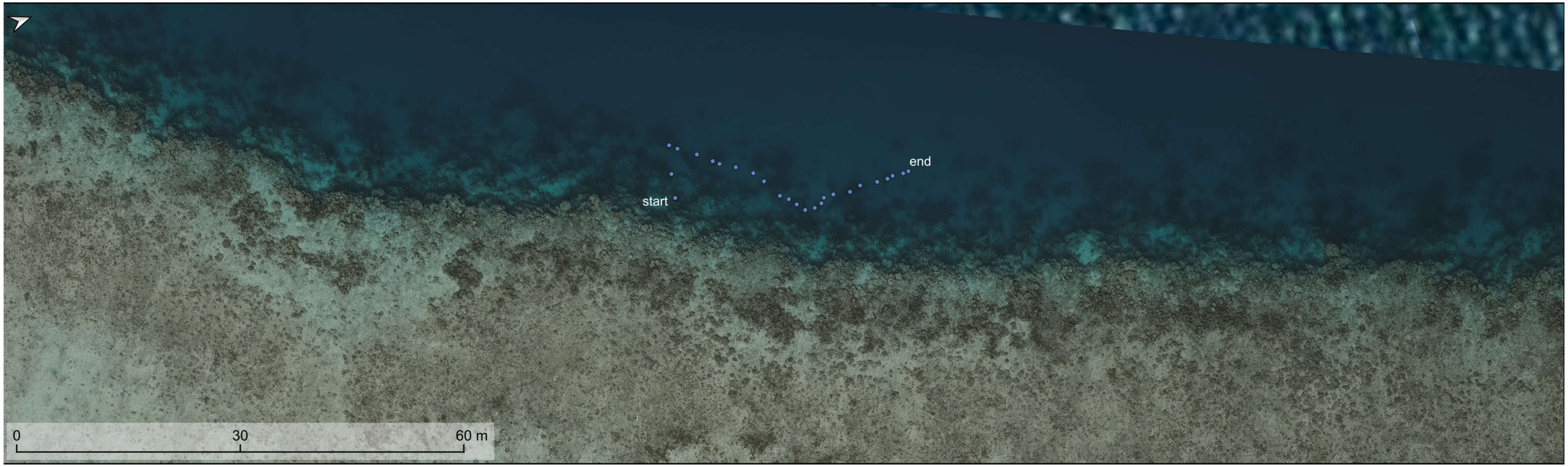


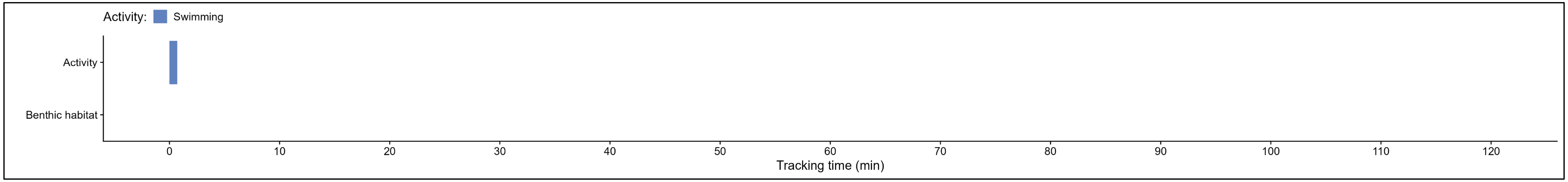

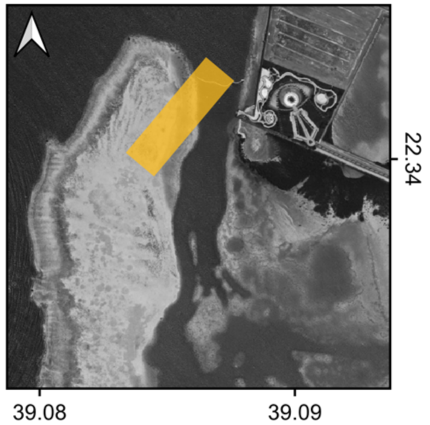


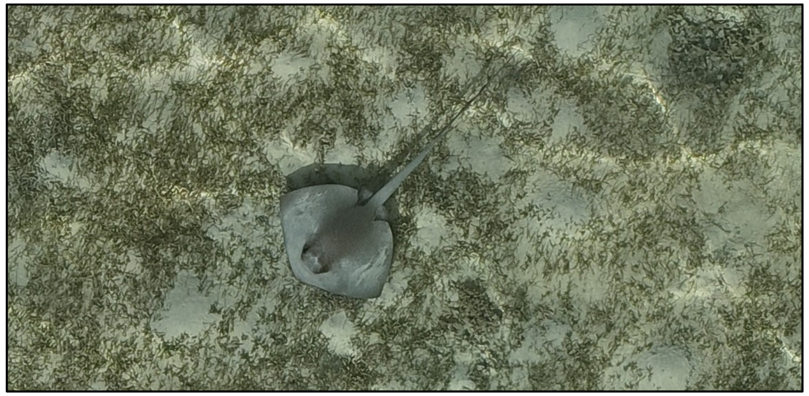

| | |
|---|---|
| Species: | *Pastinachus sephen* |
| Date: | 2024-05-24 |
| Disc width: | 61 cm |
| Disc length: | 43 cm |
| Total length: | 139 cm |
| Notes: | |

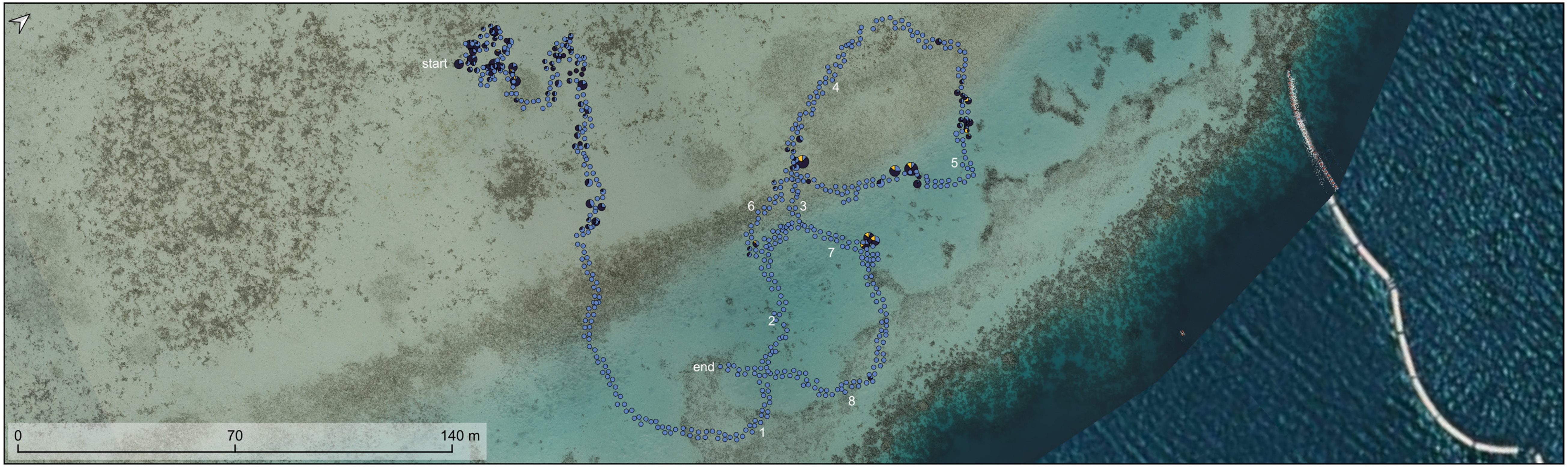


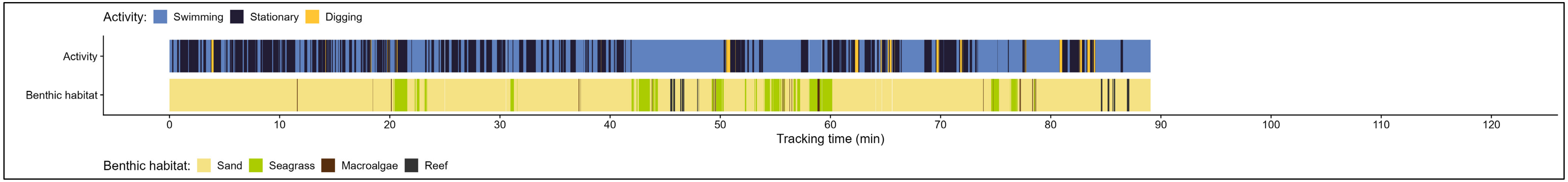

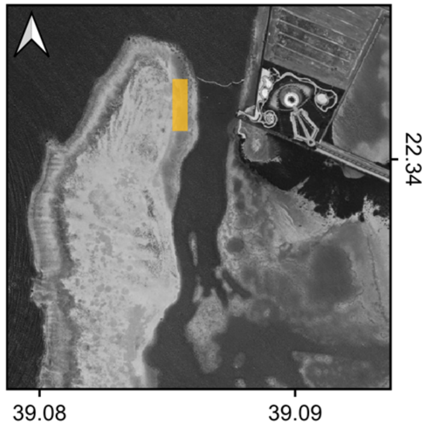


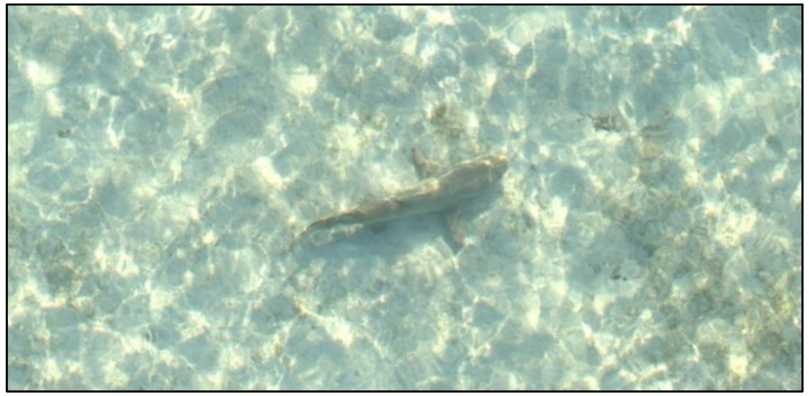

Species: *Negaprion acutidens*

Date: 2024-02-21

Disc width:

Disc length:

Total length: 57 cm

Notes:

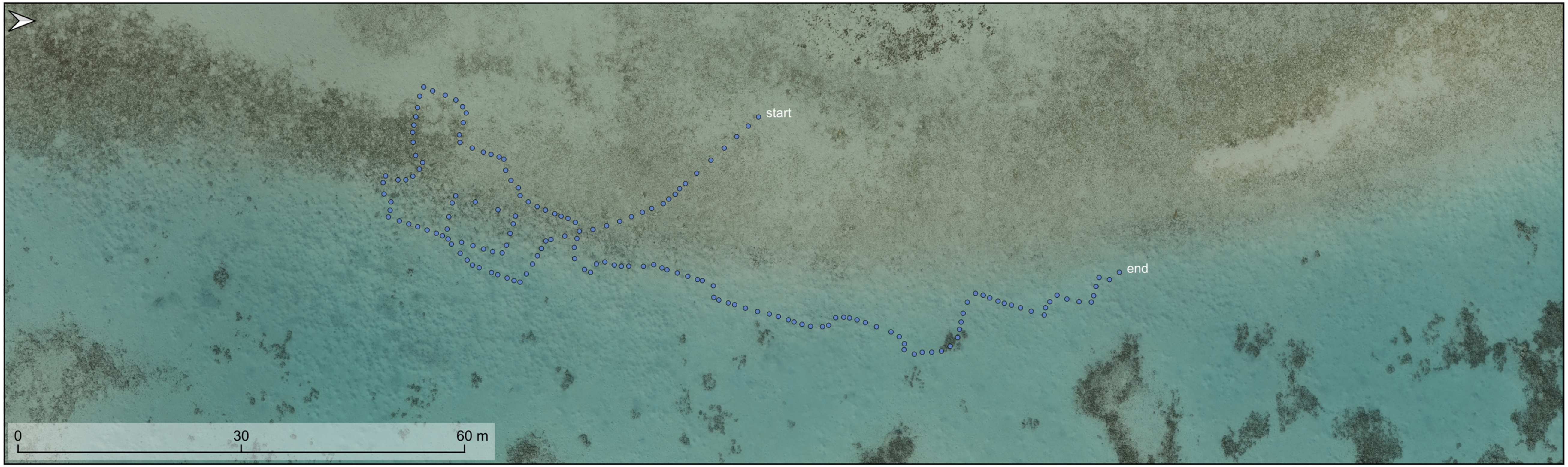


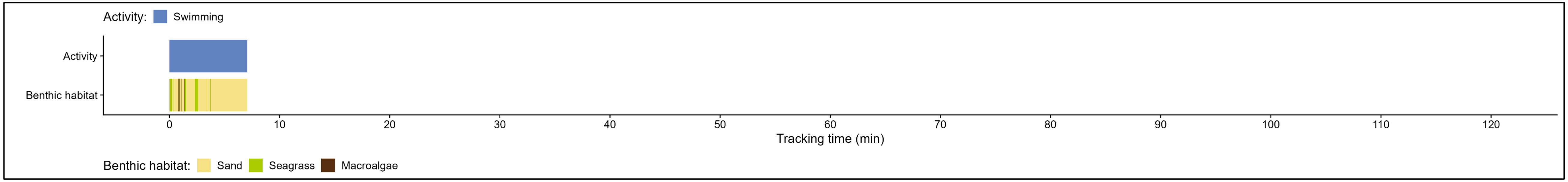
